# ProteoForge: An Imputation-Aware Framework for Differential Proteoform Discovery in Bottom-Up Proteomics

**DOI:** 10.64898/2025.12.12.694008

**Authors:** Enes K. Ergin, Agustina Conrrero, Kirsty M. Ferguson, Philipp F. Lange

## Abstract

The human genome contains approximately 20,000 protein-coding genes. However, millions of diverse protein variants, called proteoforms, exist. Despite originating from the same gene, proteoforms often have distinct biological roles. In bottom-up proteomics, the aggregation of peptide measurements into protein-level quantities often obscures this information. Existing methods for proteoform deconvolution are limited by their handling of missing data, which can introduce significant bias. To address this we developed ProteoForge, which builds on an imputation-aware statistical model to identify and group co-varying peptides into quantitatively differential proteoforms (dPFs). Benchmarking against existing deconvolution methods demonstrated that ProteoForge provides high accuracy and stability in datasets with high rates of missing values, complex experimental designs, or varying signal strengths. Application of ProteoForge to proteomics data from lung cancer cells under hypoxia revealed extensive proteoform-level regulation hidden by standard protein-level analysis.

## 1. INTRODUCTION

The human genome contains approximately 20,000 protein-coding genes. However, contrary to a simplified view of the central dogma, millions of diverse protein forms exist^1,2^. Made through mechanisms such as alternative splicing and post-translational modifications (PTMs)^3,4^, proteoforms originating from the same gene are not functionally redundant and can exhibit different or even antagonistic roles ^5–7^. For instance, distinct proteoforms of the oncogene KRAS have opposing effects on MAPK signalling. This underscores the limitations of gene or RNA-level analysis alone in explaining biological phenomena and the need for analysis methods that encompass protein diversity^8,9^.

Bottom-up proteomics is a standard method for large-scale protein analysis^10^ where proteins are digested into smaller peptides before measurement. This process introduces the “protein inference problem”: a single peptide sequence can map to multiple proteins, isoforms or proteoforms, making it difficult to trace back to its specific origin^11–13^. To manage this ambiguity, peptide measurements are often aggregated into “protein groups” following the Occam’s-razor principle^14^, which simplifies data analysis but obscures the quantitative behaviour of individual proteoforms^15–17^. Top-down proteomics analyzes intact proteoforms, however its application remains limited by technical constraints^18^. As an alternative, several computational methods have therefore been developed to infer proteoform-level changes from bottom-up data^19–23^. Established methods such as PeCorA (Peptide Correlation Analysis) detect quantitative changes in single modified peptides, while COPF (Correlation-based functional ProteoForm assessment) identifies co-regulated peptide groups that define a proteoform^21,22^. More recently, advanced frameworks have emerged to refine this resolution: AlphaQuant utilizes a hierarchical tree-based structure to propagate quantitative confidence from fragment ions to genes, while MSstatsWeightedSummary employs convex optimization to disentangle protein abundances from shared peptides^20,23^. However, a critical barrier for these methods is their handling of missing data, which often requires removal of incomplete measurements or the use of imputation strategies that can bias results.

Here, we present ProteoForge, a computational framework using an imputation-aware statistical model to identify quantitatively differential proteoforms (dPFs) from bottom-up proteomics data. We evaluate ProteoForge against COPF and PeCorA, which represent the current gold standards for correlation-based proteoform discovery and single-peptide outlier detection, respectively. While newer tools like AlphaQuant and MSstatsWeightedSummary offer powerful quantification capabilities, they represent concurrent developments with distinct operational focuses, such as resolving known shared peptides or utilizing specific counting statistics for missing values, whereas COPF and PeCorA share ProteoForge’s primary objective of identifying de novo peptide discordance in standard processed datasets. In this study, we apply ProteoForge to both simulated and publicly available proteomics datasets to demonstrate its performance and its ability to uncover subtle yet critical biological insights that are often obscured by conventional protein-level quantification.

## 2. MATERIALS AND METHODS

### 2.1 ProteoForge framework

The core workflow of ProteoForge involves four modules: (1) data processing and normalization, (2) identification of significantly discordant peptides, (3) clustering of quantitatively similar peptides, and (4) construction of differential proteoforms (dPFs) (Figure 1A).

**Figure 1.**
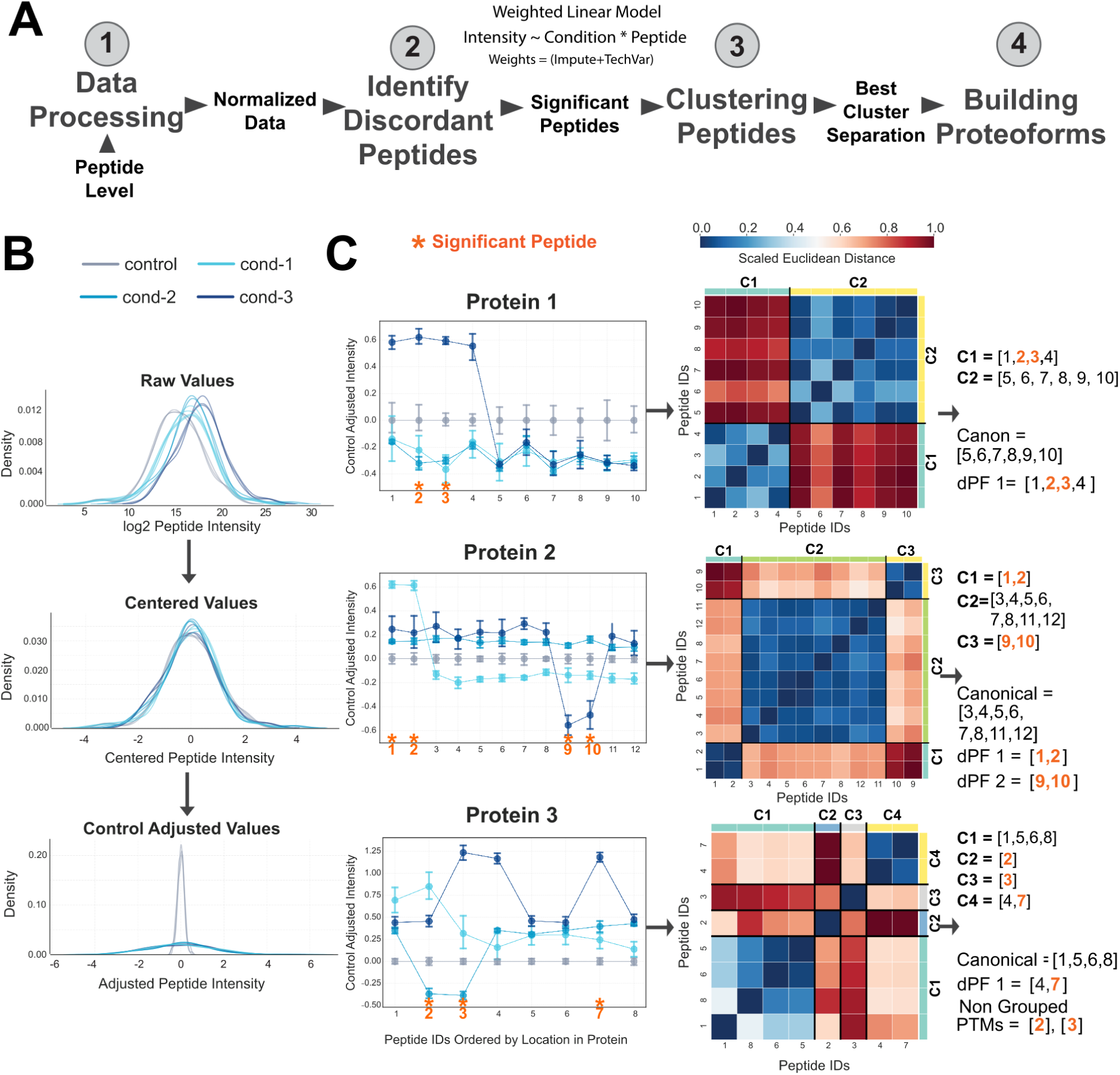
Overview of ProteoForge framework. **(A)** Schematic representation of the four primary stages of the ProteoForge analytical workflow: (1) Data Processing, (2) Identifying Discordant Peptides, (3) Clustering Peptides, and (4) Building Proteoforms. **(B)** Density plots demonstrating the impact of different normalization stages on the peptide intensity distributions for four conditions (control, cond-1, cond-2, cond-3). **(C)** Examples for three proteins (Protein 1, Protein 2, and Protein 3) illustrating the identification of Significantly Discordant Peptides. Plots show the Control Adjusted Intensity (Y-axis) for individual peptides (X-axis, ordered by location) across the conditions. Error bars represent the 95% confidence interval (CI) around the mean of replicates. The weighted linear model used to identify significant peptide-condition interaction; orange stars indicate significantly discordant peptides (p < 0.001). Examples of Peptide Clustering (three heatmaps) and Proteoform Building (right text). Heatmaps show the scaled Euclidean distance between all peptide pairs (color bar, 0.0 to 1.0) for each protein. The resulting Canonical peptide groups, final differentially expressed Proteoforms (dPFs) and non-grouped (Single peptide PTMs) are listed to the right.

#### 2.1.1 Module 1: Data processing and normalization

This module selects fully quantified peptides from proteins with at least four peptides. Prior imputation is required if there are missing values. The data is normalized to highlight changes relative to the control, standard, or normal samples. Raw non-log intensities are transformed into a logarithmic scale. Each sample undergoes z-score normalization: *z_i_ = (y_i_ - μ) / σ*, where *y_i_* is the log-transformed value, *μ* is the mean, and *σ* is the standard deviation, to ensure a mean of zero and a standard deviation of one^21^. To establish a baseline for each peptide relative to its control average, the average z-scored intensity of each peptide across control samples *(C_p_)* is calculated and subtracted from the corresponding peptide’s z-scored intensity in each sample: *A_p,i_ = z_p,i_ - C_p_*. This facilitates more precise comparisons between experimental conditions and highlights potential changes in peptide abundance (Figure 1B).

#### 2.1.2 Module 2: Identification of significantly discordant peptides

A linear model with weights is used to identify peptides that are significantly discordant within proteins across conditions. The process iteratively compares each peptide with its sibling peptides, fitting a linear model that includes an interaction term between these groups. Each model can be represented as *Intensity ∼ Condition * Peptide*, where *Intensity* represents the standardized peptide intensity, *Condition* denotes the experimental condition, and *Peptide* refers to the individual peptide being analyzed. There is a variety of linear models to choose from, and based on the model, the custom weights calculated from various known artifacts, such as imputation or technical variation, can be passed as a weight matrix or a robust linear model, which calculates internal robust weights based on the data (Supplementary Note 1). The main model we have chosen to highlight ProteoForge’s application and benchmarks is RLM; more details on each model can be found in Supplementary Note 1.

The statistical significance of all interaction terms was determined using a Wald test, with the specific test statistic and its underlying distribution tailored to the linear model used. We adjusted the resulting p-values to control the False Discovery Rate (FDR) using the Benjamini-Hochberg (BH) procedure in a two-step approach, which first groups and adjusts p-values on a per-protein basis, followed by a global adjustment. This effectively mitigates the statistical burden from the large number of simultaneous tests, balancing statistical power with rigorous error control.

#### 2.1.3 Module 3: Clustering of quantitatively similar peptides

Peptides are first grouped by calculating a distance matrix based on the Euclidean distance between their median adjusted intensity profiles across all samples. This distance matrix is used for hierarchical clustering with the Ward linkage method. While the module provides several functions, the default (*hybrid_outlier_cut)*, automatically determines the optimal cluster number.

#### 2.1.4 Module 4: Construction of differential proteoforms (dPFs)

To call a set of peptides a differential proteoform (dPF), the algorithm requires the significantly discordant peptides to belong to a cluster containing at least two peptides (Figure 1C). For proteins without such clusters, all peptides are assigned to the non-significant representation or canonical (dPF_0). When a cluster with significant peptides is identified, its peptides form a new differential proteoform (dPF_1), while the remaining peptides stay in dPF_0. Due to insufficient evidence, significantly discordant peptides in singleton clusters are excluded from both groups, denoted as dPF_-1. These cases, potentially representing single peptide PTMs, are reported separately for further analysis. Finally, a peptide-to-dPF mapping table is generated. This table uses the naming convention “proteinacc_dPF_N” to represent each proteoform, where N is the cluster, and dPF_ denotes the non-significant representation, excluding significantly discordant peptides.

### 2.2 Benchmarking performance of ProteoForge against PeCorA and COPF

ProteoForge’s performance was benchmarked against COPF and PeCorA using both a publicly available mass spectrometry dataset with in-silico modifications and a series of controlled simulations. In these data, the intensity values of varying numbers of peptides per protein were perturbed to enable assessment of proteoform prediction.

#### 2.2.1 SWATH-MS interlab dataset

A benchmark dataset was recreated from the COPF manuscript^22^, which originated from a SWATH-MS inter-laboratory study of HEK293 cell lysates across 21 replicate runs performed on days 1, 3, and 5^24^. After filtering, a ground truth was established by introducing two types of in-silico changes. First, to simulate differential expression, peptide intensities for samples from days 3 and 5 were multiplied by random factors ranging from 1 to 6. Second, to create artificial proteoforms, a subset of peptides from 1,000 randomly selected proteins had their day 5 intensities modified by a random factor between 0.01 and 0.9. The number of modified peptides per protein was varied to be one, two, a random fraction between 2% and 50%, or 50% (rounded up per protein (e.g., perturbing three out of five peptides). The first three datasets (one, two or random) directly mirrored the COPF benchmark.

#### 2.2.2 Simulated scenarios

To assess performance under various conditions, a series of controlled simulations was performed. A base dataset was first established by generating data for 500 proteins across 10 replicates in 3 conditions, with each protein containing between 5 and 50 peptides. The simulation process began by sampling mean protein abundances from a normal distribution (mean = 20, standard deviation = 2) and protein-level coefficients of variation (CVs) from a log-normal distribution. The number of peptides per protein was determined using a beta-binomial distribution (α=0.5, β=3). Peptide-level abundances were subsequently generated from these protein values, incorporating random noise and a small fraction of outliers (0.01% of peptides) to simulate experimental data. This base dataset then served as the foundation for four distinct simulation scenarios.

Simulation 1 was conducted to evaluate the impact of data imputation in the context of different peptide perturbation levelsl. Perturbation was modeled by varying the number of altered peptides per protein across four categories: two, random (2-50%), 50%, and >50%. For each perturbation level, two data versions were generated: a complete version and an imputed version. Generation of the imputed version involved the introduction and subsequent correction of missing values across 200 proteins. Specifically, 100 proteins featured random missing values in up to 35% of replicates, which were imputed using the k-Nearest Neighbors (kNN) method. The remaining 100 proteins contained complete missing values, replicates entirely absent within one of the three experimental conditions. This complete missingness was imputed with downshifted low values, a method commonly employed for data considered missing not at random^25^. Simulation 2 systematically tests the effect of data missingness, by creating 25 distinct datasets combining different levels of protein and peptide missingness (ranging from 0% to 80%). Simulation 3 investigates the magnitude of perturbation by applying log_2_ fold changes drawn from six different intervals, ranging from (0.1, 0.25) to (1.25, 1.50). Finally, Simulation 4 examines multi-condition complexity by generating datasets with two to six conditions, testing the effects of peptide perturbation overlap and directionality. Simulation 2 uses the ‘random’ peptide perturbation level as its base, while Simulations 3 and 4 use imputed data and random peptide perturbation, followed by modification of other parameters to establish the experiments.

#### 2.2.3 Performance metrics

Performance was quantified using Receiver Operating Characteristic (ROC) curves, the Area Under the Curve (AUC), and the Matthews Correlation Coefficient (MCC). These metrics were calculated from a confusion matrix (True/False Positives/Negatives) by comparing method outputs against the *in-silico* ground truth across a range of significance thresholds. The standard formulas for these metrics were applied as described in Chicco & Jurman (2020, 2023)^26,27^. The MCC, ranging from −1 to +1, was selected for its balanced, class-imbalance-robust performance: +1 for perfect prediction, 0 for random prediction, and −1 for inverse prediction.

### 2.3 Analysis of lung cancer cell hypoxia dataset

Quantitative proteomics data were sourced from Tomin et al.^28^, who used bottom-up proteomics to investigate the response of lung cancer cells to hypoxia (PRIDE repository, PXD062503). Our analysis used the precursor-level quantitative data (report.parquet) generated by the DIANN software as input.

#### 2.3.1 Data preprocessing

Modified peptide-level data were first calculated by summing precursor intensities for each peptide. The initial dataset consisted of 16 samples (four biological replicates per group, across two time points (48 hr vs 72 hr) and two conditions (Hypoxia vs Normoxia)). Data were filtered: i) to include only peptides quantified in at least four biological replicates in one or more experimental groups, and ii) to retain only proteins identified by four or more peptides. This resulted in a final matrix of 95,369 peptides across 7,161 proteins.

Missing values were handled using a two-step imputation strategy. First, peptides with values completely missing across all replicates of a condition were imputed using values drawn from a normal distribution derived from the 0.1 percentile of observed intensities, shifted down by a magnitude of two. Second, all remaining sparse missing values were imputed using a k-Nearest Neighbours (kNN) approach. Finally, the complete data matrix was normalized using the ProteoForge control-based protocol, with the ‘Normoxia (21% O_2_) - 72 Hr’ samples serving as the reference control group.

#### 2.3.2 ProteoForge application

ProteoForge was applied to the preprocessed data to make three comparisons: (1) Hypoxia vs. Normoxia at 48 hours, (2) Hypoxia vs. Normoxia at 72 hours, and (3) a time-course analysis comparing Hypoxia 48hr, Hypoxia 72hr, and Normoxia 72hr samples.

For all analyses, the robust linear model (RLM) was used. A two-step multiple testing correction was applied, using a Bonferroni correction at the protein level followed by a global Benjamini-Hochberg FDR correction (fdr_bh)^29^. Peptides with an adjusted p-value below 0.001 were classified as significantly discordant. Peptide clusters were generated using the hybrid_outlier_cut method, and these clusters were used to construct differential proteoforms (dPFs) as previously described.

To interpret the biological relevance of significant findings, peptides within dPFs were annotated using a custom utility (annotate.py). This module integrates feature annotations from UniProt (May 2024)^30^, protease-substrate information from MEROPS^31^, and post-translational modification data from iPTMnet^32^. All significant peptides were also manually cross-referenced with the TopFind database^33,34^ to identify potential neo-N and C-termini.

#### 2.3.3 Downstream analysis

To examine the impact of ProteoForge on global protein-level analyses, four different protein-level quantification strategies were compared: (1) Original, the protein-level output from DIANN, subset to match the proteins analyzed by ProteoForge; (2) mean(pep), a summarization method based on averaging top peptide intensities per protein; (3) mean(all), a summarization method based on averaging all identified peptide intensities per protein; and (4) dPFs, based on averaging peptide intensities within each identified dPF group.

These four datasets were analyzed using QuEStVar^35,36^ to identify statistically different or equivalent proteins, using an equivalence boundary of [-0.25, 0.25] LFC (log2 fold change), a difference boundary of [-0.5, 0.5] LFC, and an FDR of 0.05. The resulting upregulated, downregulated, and equivalent proteins from each strategy were then subjected to pathway analysis using g:Profiler^37^, with all identified proteins as the background and an FDR threshold of 0.001.

### 2.4 Technical details

ProteoForge was built and tested using Python (version 3.13.2). Numpy (v.1.24.3)^38^ and Pandas (v.1.5.2)^39^ facilitated data manipulation, analysis. The Statsmodels (v.0.14.1)^40^ package was used to build and test the linear models, whereas scikit-learn (v.1.5.2)^41^ was used for performance metrics and kNN imputation. The SciPy (v.1.10.1)^42^ served as the backbone in the clustering analyses. All visualizations were generated using Matplotlib (version 3.7.2)^43^ and Seaborn (version 0.11.2)^44^. Benchmarking against COPF and PeCorA utilized custom R (v4.4.2) scripts interfacing with CCProfiler (v0.99.1)^22,45^ for COPF, and PeCorA (v0.0.90)^21^. The complete analysis codes, associated datasets, computational notebooks, and their rendered outputs are permanently archived on GitHub and can be accessed via https://github.com/LangeLab/ProteoForge_Analysis/.

## 3. RESULTS

### 3.1 ProteoForge: a four-module framework to identify differential proteoforms

Identifying distinct proteoforms from bottom-up proteomics data is challenging as peptides are often shared between proteoforms and information is lost during protein inference. Existing proteoform deconvolution methods are frequently limited by their handling of missing data. We therefore developed ProteoForge, a framework that leverages peptide-level quantitative data to identify and assemble theoretical peptide groups, termed quantitatively differential proteoforms (dPFs). This framework distinguishes peptides that behave differently across conditions and defines dPFs as a set of at least two peptides from a protein with similar behaviour patterns. Importantly, ProteoForge is an imputation-aware statistical model, minimizing bias from missing values by incorporating information from auto-calculated or user directed weights during linear modelling.

ProteoForge has four key modules. First, it normalizes peptide intensity data (Figure 1A.1). Next, a robust or weighted linear model identifies peptides with significantly discordant quantitative patterns (Figure 1A.2). The third module clusters peptides with similar quantitative profiles (Figure 1A.3). Finally, dPFs are built by combining statistical and clustering data: a cluster with at least one significant discordant peptide and a minimum of two peptides is assigned as a dPF. Significant singletons are flagged and all other peptides not in a dPF are assigned to form a canonical proteoform (dPF_0) (Figure 1A.4).

### 3.2 ProteoForge demonstrates more robust identification and grouping of peptides than PeCorA and COPF

We set out to assess ProteoForge’s performance in two key areas, compared to two alternative proteoform deconvolution methods, PeCorA and COPF: i) quantitative assessment of significantly discordant peptide identification (referred to here as ‘peptide identification’) and, ii) grouping of peptides with similar variance and intensity profiles within parent proteins to infer potential proteoforms (referred to here as ‘peptide grouping’). For peptide grouping, only COPF was used for comparison as PeCorA does not provide this functionality. To model both complete data and data with imputed missing values, we used real data with known perturbations as well as simulated datasets.

First, we used the COPF benchmarking approach of Bludau et al. (2021)^22^ and a published HEK293 dataset^24^, to generate four distinct in-silico datasets of increasing perturbation complexity (referred to as “the SWATH-MS Interlab benchmark”). For 1,000 randomly selected proteins, either one, two, a random number (2%-50%), or half (50%) of peptides were perturbed per protein.

For peptide identification, at one-peptide, two-peptide, and random conditions ProteoForge outperformed all methods, however when the half of peptides perturbed ProteoForge drastically reduced in prediction performance to 0.06 MCC (Figure 2A). Half peptide condition is also COPF, performing the best up to 0.14 MCC. The trends observed in Figure 2A were supported by Receiver Operating Characteristic (ROC) curve analysis, where ProteoForge maintained a high area under the curve (AUC 0.79–0.94) across all scenarios except 50% perturbation condition (Figure S1A). ProteoForge’s MCC performance curve formed a broad plateau across a wide range of p-values (Figure S1B); this demonstrates that, while peak performance is achieved at stringent thresholds (-log10(p)> 4), a lower stringency threshold, such as -log10(p) = 3, still captures a substantial fraction of the maximum predictive accuracy (Figure S1B). ProteoForge therefore provides a robust strategy for peptide identification that balances high precision with broad discovery, avoiding true proteoform loss due to overly conservative filtering.

**Figure 2.**
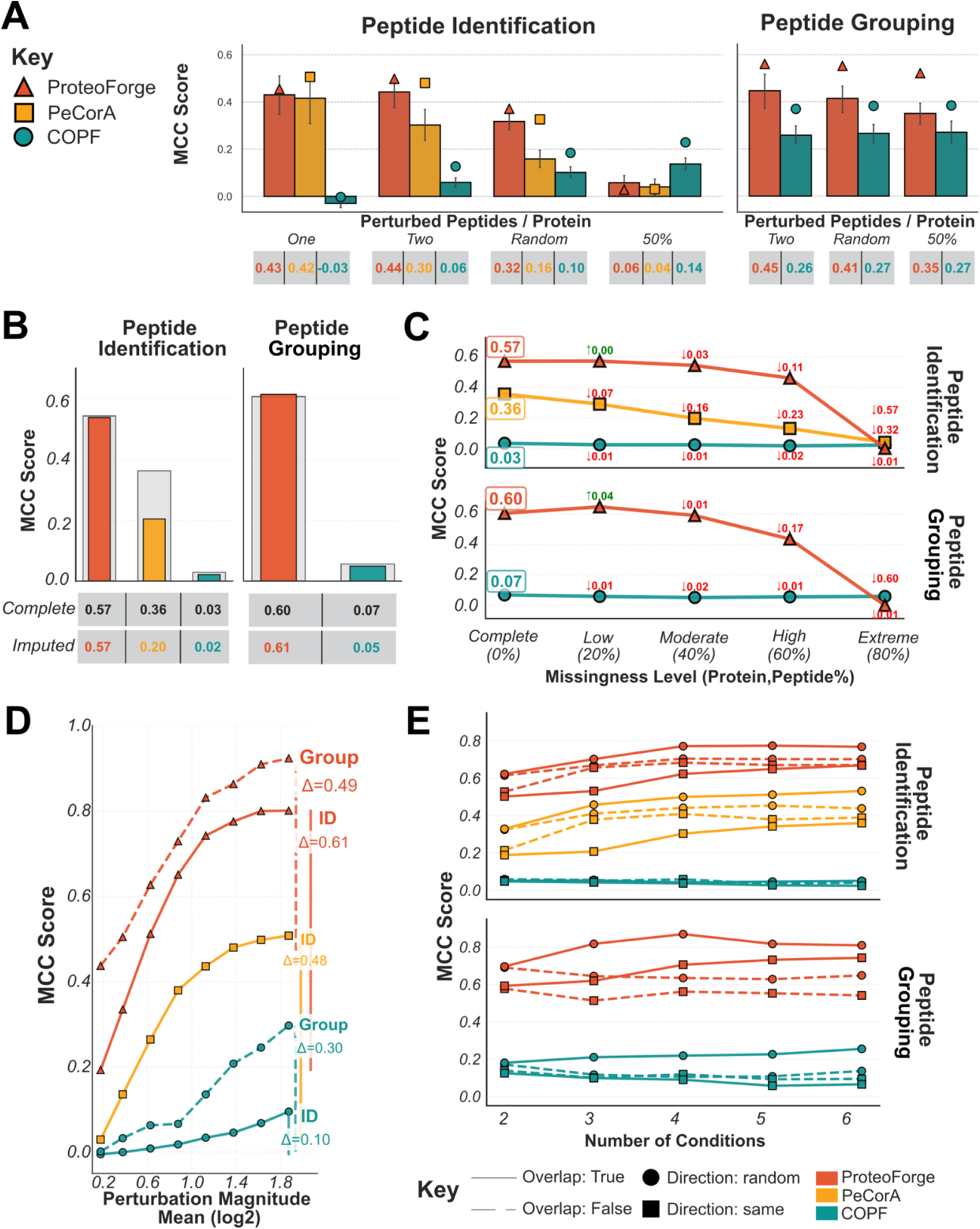
ProteoForge demonstrates more robust identification and grouping of peptides across diverse benchmark conditions than PeCorA and COPF. The performance of ProteoForge, PeCorA, and COPF were evaluated using peptide identification (detecting discordant peptides) and peptide grouping (clustering co-varying peptides). Unless otherwise specified, the method is indicated by colour and shape (ProteoForge: triangle, red, PeCorA: square, yellow, COPF: circle, teal). Performance is reported as the Matthews Correlation Coefficient (MCC). PeCorA was excluded from peptide grouping analyses. **(A)** Performance on a real-world benchmark SWATH-MS Interlab dataset. Bar heights represent the mean MCC score. Overlaid points (triangles, squares, circles) indicate the MCC score achieved at a Benjamini-Hochberg adjusted p-value of 0.001. Error bars: 95% confidence interval calculated from 1000 bootstrap samples. **(B)** Simulation 1: MCC comparing complete data (grey) and imputed data with random number of perturbed peptides (coloured). **(C)** Simulation 2: MCC scores on imputed simulated data across a range of increasing missingness levels, from 0% (complete) to 80%. **(D)** Simulation 3: Method performance as a function of the mean log2-fold change (Perturbation Magnitude) for perturbed peptides in a simulation. **(E)** Simulation 4: Performance on simulated data with an increasing number of experimental conditions. Method is encoded by colour, the data overlap is indicated by linestyle (solid: True, dashed: False), and the direction of peptide change is represented by shape (square: same direction, circle: random direction).

Next, we evaluated the performance of ProteoForge and COPF in peptide grouping. The same dataset was used with three perturbation scenarios (2, 2-50% or 50% peptides perturbed) (Figure 2A). Across all conditions, ProteoForge maintained a stable and high mean MCC of approximately 0.45, whereas COPF’s scores were consistently around 0.27. With increased perturbations up to half ProteForge’s performance reduced to 0.35. This performance difference was also reflected in a consistently higher AUC for ProteoForge (0.85) compared to COPF (∼0.77) (Figure S2A). The similar broad plateau observed for peptide identification, was also prevalent for peptide grouping, but this time reaching its optimal MCC score within a clear range of stringent adjusted p-values (-log10(p)≈ 4–6). COPF’s peak performance occurred at a less stringent threshold (-log10(p)≈ 1) (Figure S2B). This suggests that ProteoForge provides more statistically robust peptide grouping than COPF, and the performance reduction from peptide identification at 50% peptide perturbation condition is mitigated by better cluster accuracy.

To thoroughly evaluate ProteoForge’s performance against COPF and PeCorA and understand how various parameters affect its results, including data quality, experimental design, and pattern changes, we next created four simulated data scenarios (Figure 2B-E). 500 proteins were used with 250 proteins perturbed to investigate the impact of: i) data imputation in the context of different peptide perturbation levels (Simulation 1), ii) the level of data missingness (Simulation 2), iii) the magnitude of peptide perturbations (Simulation 3) and, iv) multi-condition complexity including peptide perturbation overlap and directionality (Simulation 4). First, we assessed the effect of data imputation. For data with random number (2-50%) perturbed peptides per protein, ProteoForge’s peptide identification performance was largely unaffected by imputed data; after 35% of the data was removed and imputed, ProteoForge’s MCC score decreased by only 0.005 (from 0.574 to 0.569). In contrast, PeCorA’s MCC score decreased by 0.158 (Figure 2B). This stability was consistent across different peptide perturbation scenarios, where ProteoForge maintained high AUC values while PeCorA’s scores decreased with data imputation (Figure S3A). The reduction of MCC across different perturbation scenarios and not just random was also present but it was negligible and minimal, and for more challenging half and more than half cases the MCC improved slightly for ProteoForge while others notably decreased (Figure S3B).

For peptide grouping, ProteoForge achieved higher MCC scores than COPF (Figure 2B). In fact, ProteoForge’s performance marginally increased on the imputed dataset, as shown by an improvement in the mean MCC score from 0.60 to 0.61 (Figure 2B). This increase is attributed to the imputation of completely missing values within a condition; these are replaced with low values, creating a strong signal that the algorithm correctly identifies as a significant perturbation. ProteoForge also performed well even in scenarios where individual peptide identification was challenging; in the ‘>50% Peptides’ perturbed condition, it maintained a high AUC (∼0.91), demonstrating its ability to cluster peptides effectively (Figure S4A). This is due to at least one or two peptides that are statistically discordant clustering with other peptides that would have been missed by ProteoForge peptide identification. At an adjusted p-value of 0.001, ProteoForge also consistently achieved moderate MCC scores (∼0.55-61) while COPF’s were near zero (Figure S4B).

For the second simulation, we assessed the impact of increasing data missingness. For peptide identification, ProteoForge’s performance was resilient, with its MCC score declining modestly from 0.57 to 0.46 as the percentage of missing values increased from 0% to 60%, while going from 60% to 80% missingness, a very sharp decline dropped MCC around 0. (Figure 2C). In contrast, PeCorA’s accuracy was highly dependent on data completeness and degraded sharply as the level of missingness increased (Figure 2C). Heatmaps of performance at a p-value of 0.001 confirmed that ProteoForge maintains high accuracy across combined protein- and peptide-level missingness, except the 80% missingness where it drops dramatically but more so for peptide-level. In contrast, PeCorA’s performance is primarily impacted by protein-level missingness (Figure S5A). For peptide grouping, ProteoForge’s MCC scores also decreased with higher missingness but were consistently higher than those of COPF, except for 80% which highest levels of missingness effectively set MCC at around 0 (Figure 2C and Figure S5B). These extremely sharp declines at very extreme missigness levels are due to RLM’s internal auto-calculation of weights to consider imputed values as the real ones, due to them being the dominant values, which creates an issue. However with custom imputation weights and WLS or GLM models even at these extreme levels prediction accuracy for both identification and grouping is higher than other methods (Supplementary Note 1).

Next, we evaluated the impact of peptide perturbation magnitude on method performance, using imputed data with random (2-50%) peptide perturbations (Figure 2D). As expected, the performance of all methods improved with a stronger biological signal; however, ProteoForge achieved higher MCC scores than the other methods across the entire tested range (Figure 2D). For peptide identification, ProteoForge’s MCC increased to 0.91 at the highest perturbation magnitude, while PeCorA reached 0.52. A similar trend was observed for peptide grouping, where ProteoForge’s MCC reached 0.91 while COPF’s remained below 0.29 (Figure 2D, Figure S6).

Finally, the impact of increasing experimental complexity was assessed by varying the number of conditions from two to six, again using imputed data with ‘random’ peptide perturbations. ProteoForge’s performance remained high and stable for both peptide identification and grouping as the number of conditions increased (Figure 2E). This stability was maintained across various complex scenarios, including those with random directional changes and overlapping perturbations between conditions (Figure S7).

### 3.3 ProteoForge identifies alternative hypoxia-induced dPFs in lung cancer cells

Having evaluated ProteoForge’s performance under a variety of simulated scenarios, we next tested if our framework could bring new biological insights to a published proteomics dataset. To assess the influence of hypoxia on the glyoxalase pathway, Tomin et al.^28^ performed bottom-up proteomics on H358 lung cancer cells exposed to normal (21% O_2_) or hypoxic (1% O_2_) conditions for 48 or 72 hours. Following data preparation and processing, the dataset contained 95,369 peptides across 7,161 proteins. ProteoForge ran for both 48 and 72 hour conditions separately to be able to determine and clearly display hypoxia vs normoxia proteoforms at both conditions. At 48h comparison, ProteoForge found ∼55% (3,920) of proteins had not significantly discordant peptides, ∼30% (2,185) had a single significantly discordant peptide, and the remaining ∼15% (1,056) had multiple significantly discordant peptides (Figure 3A). At 72hr comparison, ProteoForge found ∼29% (2,109) of proteins had not significantly discordant peptides, ∼35% (2,477) had a single significantly discordant peptide, and the remaining ∼36% (2,575) had multiple significantly discordant peptides (Figure 3A).

**Figure 3.**
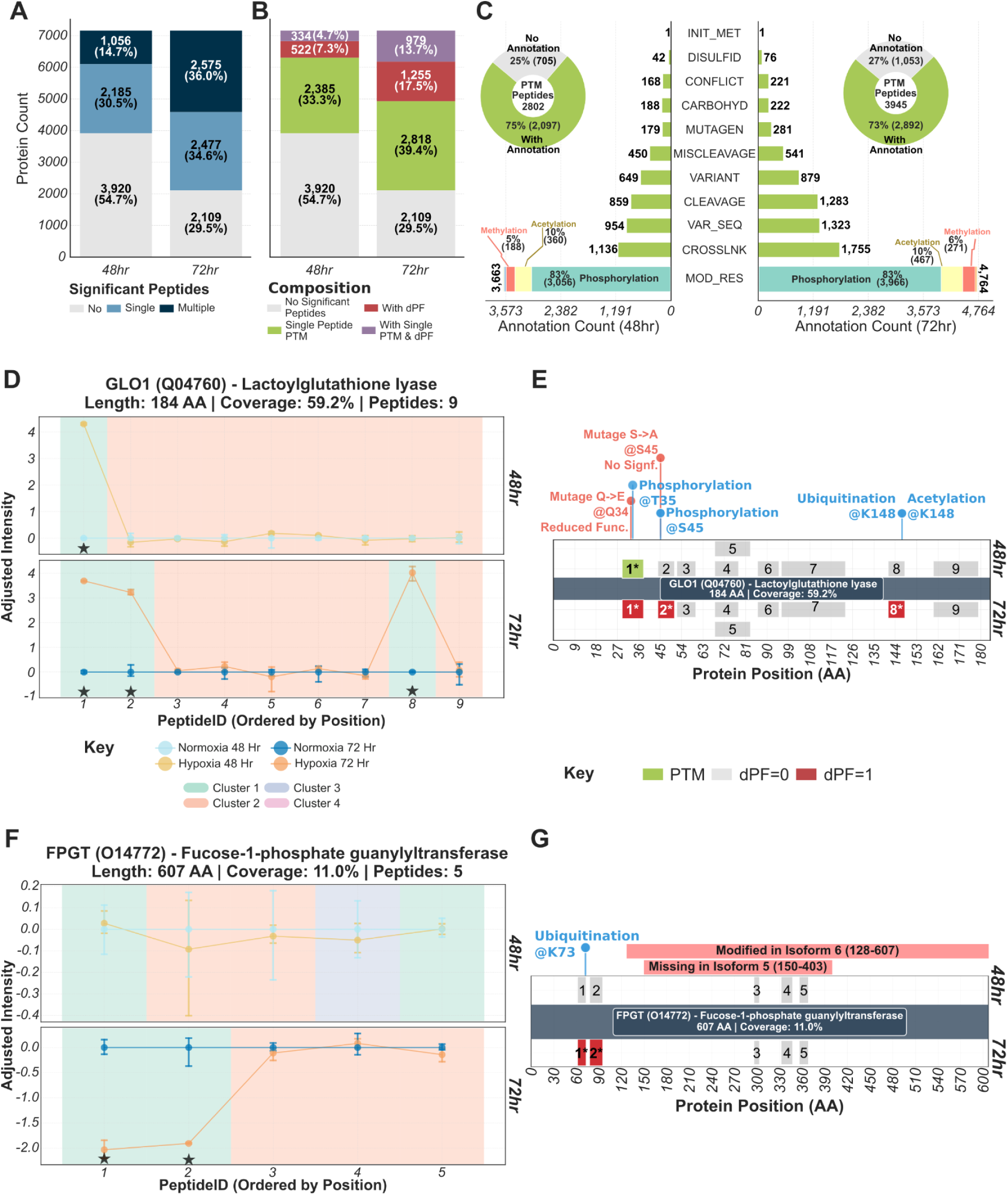
ProteoForge deconvolutes proteoform heterogeneity to identify specific PTMs and potential proteoform-like events. ProteoForge was applied to a lung cancer cell hypoxia dataset to identify and characterize peptide-level regulation. **(A)** Stacked bar charts show the distribution of protein count and the corresponding total peptide count (48hr and 72hr) based on the number of significantly discordant peptides identified (adj. p < 0.001): no single significant peptide, multiple significant peptides or single significant peptide. **(B)** Stacked bar charts further classify proteins into functional categories: no significant peptides, single peptide PTM, with dPF or with single PTM and dPF. **(C)** Bar plots detail the UniProt annotations for the significant peptides from PTM and dPF proteins (48hr and 72hr). An inset pie chart shows the proportion of these peptides that match a known biological feature, with phosphorylation, ubiquitination, and acetylation being predominant modifications. (**D, F)** Quantitative profiles for peptides from Glyoxalase I (GLO1) and Fucose-1-phosphate guanylyltransferase (FPGT) are shown across four experimental conditions. Error bars: 95% confidence interval. **(F, H)** Schematics for GLO1 and FPGT mapping the locations of the quantified peptides and known protein features.

Peptides were then clustered, based on the distance values using adjusted intensity, to build proteoforms. At 48hr, ∼7% (522) of proteins showed to have at least one additional dPF, while ∼33% (2,385) of proteins contained significantly discordant peptides in singleton clusters predicted to be PTMs (named ‘single peptide PTMs’). Furthermore, ∼5% (334) of proteins were found to have both a single peptide PTM and at least one additional dPF (Figure 3B). At 72hr, ∼17% (1,255) of proteins showed to have at least one additional dPF, while ∼39% (2,818) of proteins contained at least one ‘single peptide PTMs’. Furthermore, ∼14% (979) of proteins were found to have both a single peptide PTM and at least one additional dPF (Figure 3B).

To identify the molecular basis for discordant peptide behaviour, we first examined single peptide PTMs across two timepoints, where 48hr had 2,802 peptides identified as potential PTMs across 2,719 proteins, and at 72hr 3,945 peptides across 3,797 proteins. It is important to note that a single protein could have multiple single peptide PTMs belonging to distinct clusters. For these peptides, we cross-referenced their genomic locations with annotations in a custom database. This revealed that the majority of peptides across both timepoints (75% for 48hr and 73% for 72hr) matched a known biological feature, while the remaining had no corresponding annotation (Figure 3C). The most frequent feature type was a modified residue (MOD_RES), with phosphorylation being the most prevalent (∼83% of the modified residues) followed by acetylation (∼10%) and methylation (∼5-6%) across both timepoints summaries (Figure 3C).

The enzyme Glyoxalase I (GLO1; Q04760) exemplifies the granularity provided by this approach. While the original study observed increased levels of the GLO1 product (S-lactoylglutathione) under hypoxia, they reported no correlation between hypoxia and GLO1 protein abundance. ProteoForge analysis resolved this discrepancy. At 48 hours, we identified a single discordant peptide (Peptide 1) that remained abundant under hypoxia despite low levels in normoxia (Figure 3D). This peptide harbors known phosphorylation (T35) and mutagenesis (Q34) sites, suggesting a condition-specific PTM (Figure 3E). By 72 hours, this response changed into a multi-peptide dPF (Peptides 1, 2, and 8) characterized by elevated abundance in hypoxia (Figure 3D). Peptides 1 and 2 share similar annotations, containing mutagenized sites (Q34 and S45) and phosphorylation sites (T35 and S45) (Figure 3E). Although Peptide 8 carries distinct modifications (ubiquitination/acetylation at K148) compared to Peptides 1 and 2, ProteoForge grouped them into a single dPF based on their synchronized upregulation under hypoxic stress (Figure 3E). This demonstrates that GLO1 undergoes complex, multi-site regulation that standard protein-level averaging obscures.

Fucose-1-phosphate guanylyltransferase (FPGT; Q14772) is an enzyme involved in the salvage pathway of L-fucose metabolism. The analysis at 48 hours showed all no discordant peptides, indicating a stable protein status at this initial time point. Conversely, the 72-hour comparison yielded a two peptide dPF, signifying peptide heterogeneity (Figure 3F). The dPF was defined by two discordant peptides (Peptides 1 and 2) which span amino acid positions 60–95, that showed a reduction in hypoxia relative to normoxia, while the other peptides maintained similar intensities. Notable annotations for these two peptides only include a ubiquitination even at K73 residue, however what makes these two peptides likely behave distinctly is the existence of isoforms that span the rest of the protein. Isoform 5 for the FPGT protein, is noted to be missing between 150-403 amino-acids, while Isoform 6 does generate a modified version of the protein from 128-607 amino-acids, a product of an alternative splicing event (Figure 3G). This finding highlights ProteoForge’s capacity to distinguish isoform-driven variance from PTM-driven variance without prior knowledge.

Collectively, these analyses demonstrate ProteoForge’s ability to deconvolute complex quantitative peptide signals into proteoform patterns and successfully distinguish variations linked to known annotations from other unannotated, condition-dependent peptide behaviours.

### 3.4 Proteoform deconvolution improves quantitative reproducibility and enhances statistical power for differential analysis

We next sought to examine the impact of proteoform deconvolution on a typical quantitative global protein analysis. Four different protein-level quantification strategies were applied to Tomin et al’s dataset^28^: DIANN’s protein group quantification (‘Original’), the mean of the top three most intense peptides (‘mean(top)’), the mean of all peptides (‘mean(all)’), and the mean of peptides within each ProteoForge-defined dPF (‘dPFs’).

First, we assessed quantitative reproducibility by calculating the median coefficient of variation (CV) across replicates (Figure 4A), where a lower CV indicates higher reproducibility. The ‘Original’ and ‘mean(top)’ methods showed the highest variability, with median CVs ranging from 7.7% to 9.8%. In contrast, the ‘mean(all)’ method consistently yielded the most reproducible results, with the lowest CVs across all conditions (ranging from 6.0% to 7.7%), likely as averaging all peptides has a variance-smoothing effect. While not surpassing ‘mean(all)’, the ‘dPFs’ method showed strong reproducibility, with median CVs ranging from 7.0% to 8.1%, and a robust foundation for subsequent quantitative differential analyses.

**Figure 4.**
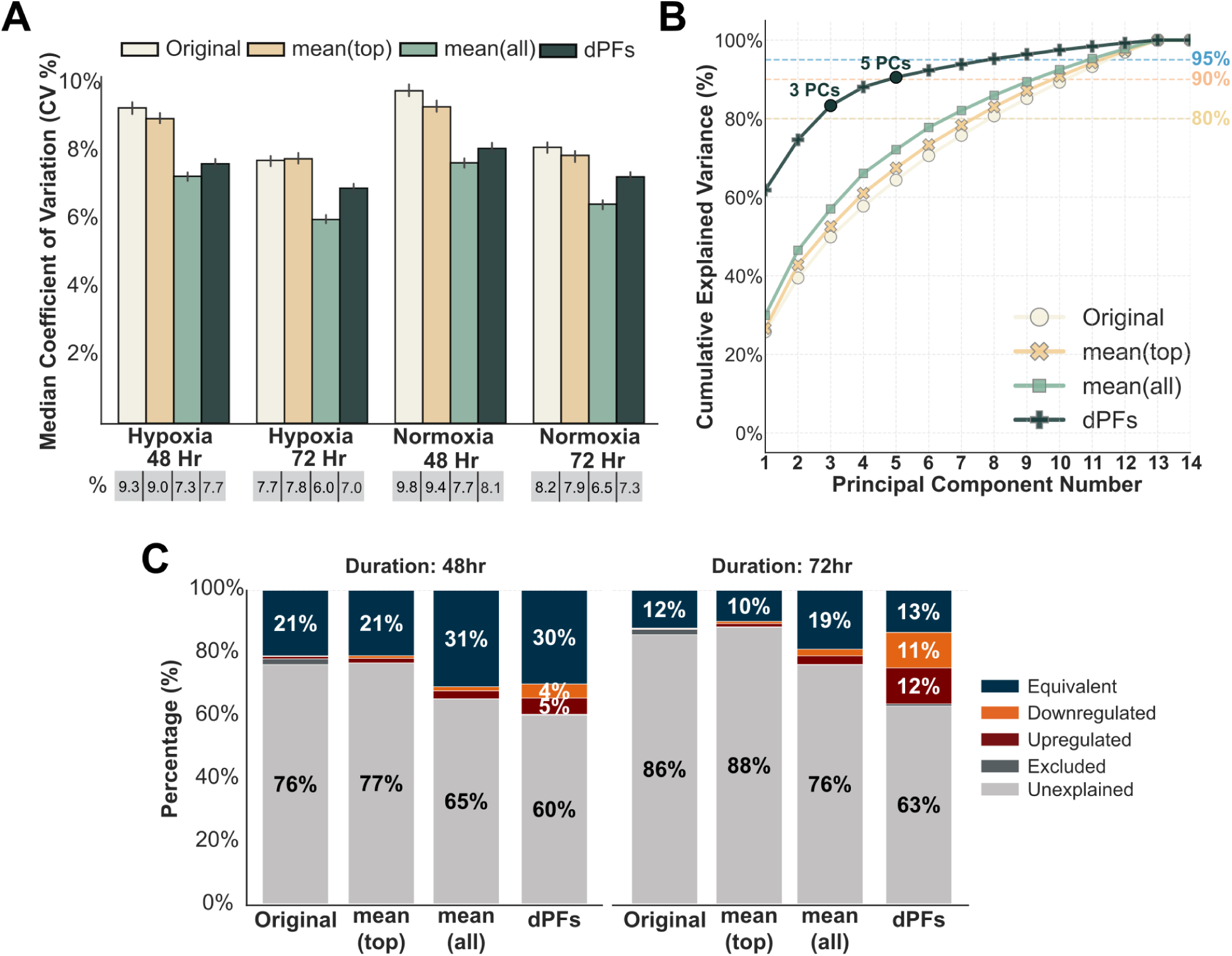
Proteoform deconvolution improves quantitative reproducibility and enhances statistical power for differential analysis. The performance of four protein-level quantification strategies were compared using the lung cancer hypoxia dataset^28^: Original (DIANN output), mean(top) (mean of top 3 peptides), mean(all) (mean of all peptides), and dPFs (mean of peptides within each ProteoForge-defined proteoform). **(A)** A bar chart compares the median coefficient of variation (CV) across biological replicates for each method and condition. Error bars: 95% confidence interval. **(B)** A line graph shows the cumulative variance explained by principal components for each quantification strategy. **(C)** Stacked bar charts show the percentage of proteins classified into different regulatory states (Equivalent, Downregulated, Upregulated, etc.) following differential analysis at 48 and 72 hours. The dPFs method identifies a substantially larger fraction of regulated proteins compared to the other three methods, demonstrating greater statistical power.

Next, we examined the data variance structure with a principal component analysis (PCA) (Figure 4B). The ‘dPFs’ method captures the most variance with the fewest components, explaining more than 80% of the cumulative variance with just three principal components. This indicates that deconvolving proteins into refined proteoform units more effectively distinguishes experimental groups, further visualized by the improved separation of groups in the PCA score plot (Figure S8).

This improved separation translates into greater statistical power for differential analyses (Figure 4C). At 72-hours, the ‘dPFs’ method classified a combined 23% of proteins as either upregulated (12%) or downregulated (11%), a substantial increase compared to the ‘mean(all)’ (4%), ‘mean(top)’ (2%), and ‘Original’ (2%) methods. By providing a clearer biological signal for a larger portion of the proteome, the ‘dPFs’ method also reduces proteins with an ambiguous “Unexplained” status, maximizing the biological insights derived from the dataset.

Finally, to understand the functional consequences of proteoform-level changes, we performed a global pathway enrichment analysis on quantitative data from the ‘Original’ or ‘dPFs’ method. While both methods identified a similar total number of enriched terms per statistical status query, their distribution was strikingly different (Figure 5A). The ‘Original’ method predominantly classified pathways as “Equivalent,” particularly at the 72hr timepoint, where almost no “Elevated” terms (those enriched in the upregulated set) were found. In contrast, ‘dPFs’ uncovered a substantial number of both “Elevated” and “Reduced” terms (those enriched in the downregulated set) at both 48hr and 72hr. This trend was consistent across all GO categories. For instance, at 72 hours the ‘dPFs’ method identified 87 “Elevated” and 181 “Reduced” GO Biological Process (GO:BP) terms, whereas the ‘Original’ method found no GO:BP terms to be “Reduced” or “Elevated” (Figure 5B).

**Figure 5.**
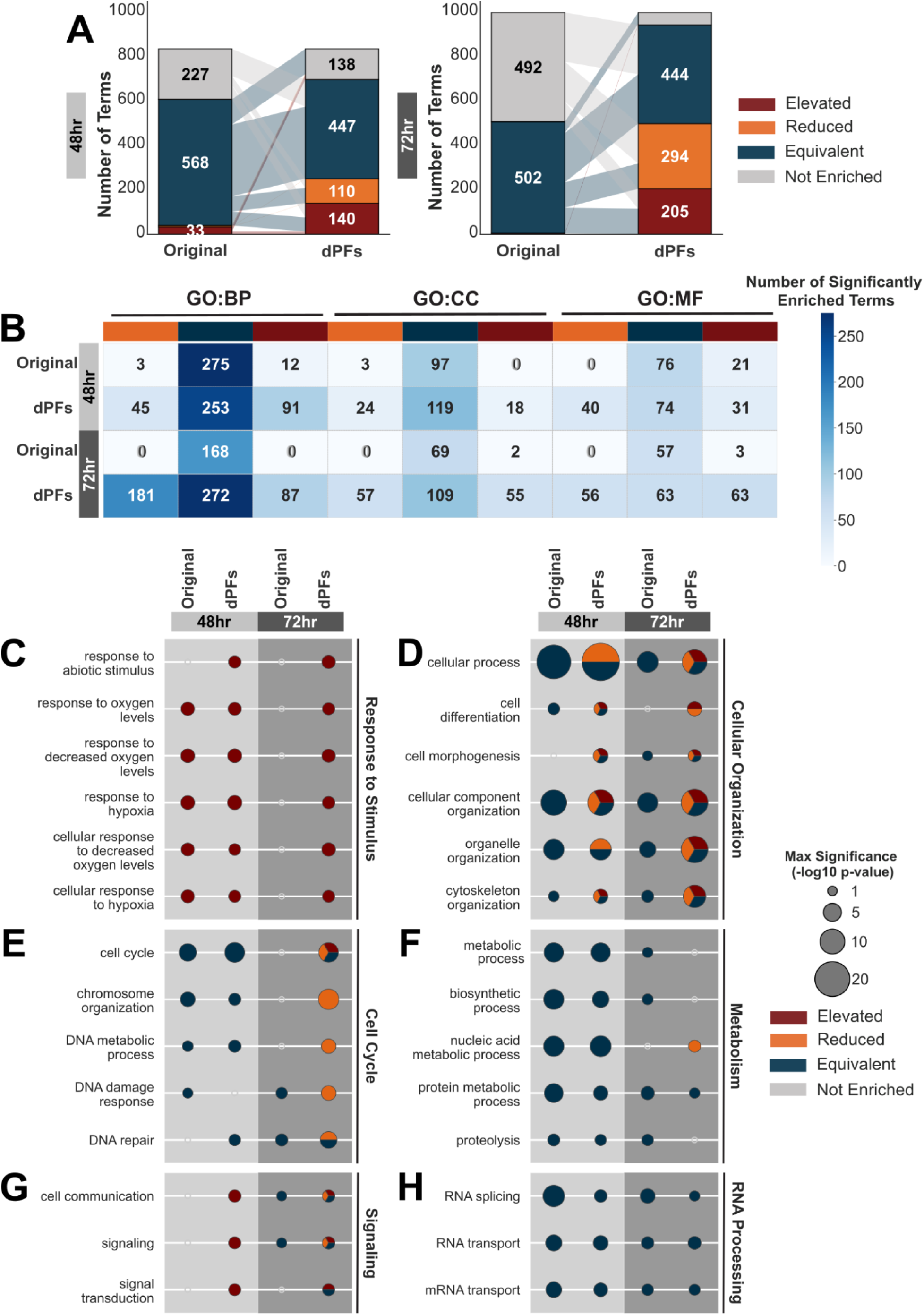
dPF-based analysis reveals dynamic pathway regulation masked by conventional protein quantification. Pathway enrichment analysis was performed on regulated protein sets derived from the Original (DIANN) and dPFs (ProteoForge) quantification methods at 48 and 72 hours of hypoxia. **(A)** An alluvial plot shows the number and enrichment status (Elevated, Reduced, Equivalent) of enriched Gene Ontology (GO) terms. **(B)** A heatmap detailing the number of enriched terms by GO category (BP: Biological Process, CC: Cellular Component, MF: Molecular Function), confirming the trend observed in (A). **(C-H)** Dot plots show the enrichment status and significance for selected GO term groups. Dot colour indicates the regulatory status (red: Elevated, blue: Reduced, orange: Equivalent), and dot size corresponds to the statistical significance (-log10 p-value). The dPFs method correctly identifies the elevation of canonical hypoxia pathways in **(C)** “Response to Stimulus” and reveals complex patterns in **(D)** “Cellular Organization,” **(E)** “Cell Cycle,” **(F)** “Metabolism,” **(G)** “Signaling,” and **(H)** “RNA Processing”.

Further analysis of the enrichment status and significance of several GO terms found distinct differences between the ‘dPFs’ and ‘Original’ methods. For example, in the “Response to Stimulus” category, ’dPFs’ identified a distinct, time-dependent elevation of canonical hypoxia pathways, specifically “cellular response to hypoxia” and “response to decreased oxygen levels” at 72 hours (Figure 5C). The ’Original’ method did not detect this response. A similarly striking contrast appeared in “Cellular Organization” and “Cell Cycle” (Figure 5D, E). While the ’Original’ method showed broad equivalence, ’dPFs’ revealed a marked reduction in cell cyled, alongside multiple dPFs showing elevation, reduction, and equivalence in Cellular Organization at 72 hours (Figure 5D, E). A notable divergence was also observed in “Metabolism”: while the 48 hours timepoints both methods suggested equivalence for metabolic processes, at 72 hours ’dPFs’ lost any enrichment for these terms, suggesting the protein-level signal may not reflect proteoform behaviour (Figure 5F). Another interesting category showed the “Signalling” terms to be generally elevated in “dPFs”, while not enriched in or equivalent in Original, indicating, while the enrichment significance is lower for these terms, the deconvolution is expanding the insights (Figure 5G). Finally, provided are some terms within the RNA processing category where both methods at both timepoints agree to be equivalent, which shows the mainly housekeeping processes didn’t change meaningfully under hypoxic conditions and didn’t result in distinct proteoform profiles (Figure 5H). Together, these results highlight that proteoform-level resolution enhances detection of pathway changes and uncovers distinct functional signatures that are averaged out in standard protein-level assessments.

## 4. DISCUSSION

In this study, we present ProteoForge, a framework that detects and groups peptides with distinct quantitative patterns into differential proteoforms (dPFs). The primary goal of ProteoForge is to move beyond conventional protein-level quantification and provide a more granular view of peptide-level dynamics, enabling the identification of candidate proteoforms for further biological validation. Importantly, ProteoForge uses an imputation-aware robust statistical model to navigate the problem of imputed missing data, a limitation for existing proteoform deconvolution methods.

Comprehensive benchmarking on both real-world and simulated datasets demonstrated that ProteoForge provides stable performance across a wide range of analytical challenges. Its accuracy was largely unaffected by high rates of missing and imputed values, a condition which substantially degrades the performance of other methods, such as PeCorA. Notably, our analysis identified a critical stability threshold: ProteoForge maintains consistent identification and grouping performance (MCC 0.55-0.65) up to 60% missingness. Beyond this point, while all methods eventually succumb to data sparsity, ProteoForge’s conservative handling of outliers prevents the rapid performance collapse observed in standard models. Furthermore, ProteoForge achieved higher accuracy across all tested signal strengths and maintained its performance as experimental complexity increased through more conditions or random directional changes (Figure 2).

The robust performance of ProteoForge can be attributed to its underlying statistical framework, specifically the selection of a Robust Linear Model (RLM) with Huber M-estimation. Our benchmarking in Supplementary Note 1 revealed that RLM offers the optimal balance between performance and usability. By effectively treating the artificial variance introduced by imputation as ’outliers’, RLM automatically down-weights their influence without requiring the user to manually specify complex weight matrices, which are prone to misconfiguration. This approach enables the incorporation of measurement variance and accounts for the uncertainty of imputed values without the fragility of standard weighting schemes. This also explains the observed performance increase after imputing completely missing conditions; the introduction of low values creates a strong, unambiguous signal of perturbation that the algorithm correctly identifies.

The deconvolution of peptide signals into dPFs has broader implications for proteomics, particularly in addressing the protein inference problem, the challenge of accurately assigning peptides to their originating proteins^13,46^. The presence of a dPF can help resolve ambiguity for shared peptides; if a peptide common to multiple proteins exhibits a discordant pattern, it provides evidence that it originates from a specific, regulated proteoform. This concept complements other advanced methods that utilize weighted networks or peptide-level evidence to enhance confidence in protein identification and quantification^20,23^. For instance, the recently introduced AlphaQuant framework leverages a hierarchical tree structure to propagate error models from the fragment ion level up to the gene, offering high sensitivity by utilizing counting statistics rather than imputation for missing values^20^. Similarly, MSstatsWeightedSummary addresses the specific challenge of quantifying proteins with known shared peptides by estimating contribution weights via convex optimization, though it currently ignores missing feature intensities^23^. By retaining and interpreting these signals, ProteoForge provides an additional layer of evidence that can refine protein-level conclusions.

Using a published proteomics dataset in lung cancer cells^28^, we demonstrated ProteoForge’s ability to successfully distinguish hypoxia-induced variations linked to known annotations, such as the condition-specific PTM in GLO1 and the potential isoform driven differences at FPGT at 72-hour only. This provides a clear rationale to prioritize proteoform candidates for functional validation. This ability provides a clear rationale to prioritize specific proteoform candidates for functional validation. Furthermore, by retaining peptide-level variance, the analysis revealed a directional reorganization of cell cycle and signaling machinery (Figure 5E, H) that was obscured by the assumption of protein-level equivalence in standard, protein-level analyses. The resulting divergence in pathway enrichment conclusions between ProteoForge (‘dPF’) and the standard analysis (‘Original’) for the ‘Metabolism’ and ‘Signaling’ categories (Figure 5F, H) highlights the importance of retaining and interpreting proteoform-level data for accurate biological conclusions.

An alternative approach for handling discordant peptides, suggested by the PeCorA framework, is to remove them to improve the accuracy of parent protein quantification^21^. While this strategy is valid for researchers focused solely on protein-level abundance, ProteoForge operates on the principle that these peptides are not errors but sources of novel biological information. Interpreting this information still has limitations and deconvolution of similar abundance patterns remains a significant challenge for the field^47,48^. Furthermore, technical artifacts from sample preparation, such as missed cleavages, can confound dPF detection. The development of centralized, comprehensive proteoform databases would greatly aid high-throughput annotation of identified dPFs and reduce the need for manual verification^3,30,49^.

Future work will focus on refining the peptide grouping logic to incorporate context-aware adaptive weighting. While RLM currently serves as the robust default, our data indicates that weighting schemes incorporating biological priors, such as limit of detection or local data density, could further enhance sensitivity. This would allow more flexible models like WLS or GLM to surpass current robust baselines in complex edge cases where data-driven weights can be reliably calculated.

## Supporting information

Analysis notebooks as html render

## Data Availability

The data and scripts related to this study are available at https://doi.org/10.5281/zenodo.17795845 and https://github.com/LangeLab/ProteoForge_Analysis/ under an CC BY-NC 4.0 license.

## Supporting information

- **Supplementary Figures:** include Figures S1 and S2, illustrating the performance of discordant peptide identification and grouping on the SWATH-MS benchmark. Figures S3 and S4 demonstrate the framework’s resilience to data imputation. Figures S5 to S7 display the performance stability across varying levels of missingness, perturbation magnitudes, and experimental complexities. Lastly, Figure S8 highlights the improved separation of experimental groups in PCA using dPF-based quantification. ***(PDF)***
- **Supplementary Note 1:** presents the evaluation and justification for selecting linear estimators for imputation-aware peptide analysis. ***(PDF)***
- **Supplementary Notebooks:** contains the collection of analysis notebooks rendered in HTML. Notebooks S1 and S2 cover the SWATH-MS benchmarking. Notebooks S4 through S6 detail the generation of simulated datasets and subsequent performance benchmarking, while Notebook S7 compares the available linear models. Notebooks S9 to S13 document the data preparation, application, and exploration of ProteoForge on the lung cancer hypoxia dataset. Notebook S14 compares protein-level downstream analysis strategies. Finally, Notebooks S3, S8, and S15 contain the code for generating the main figures. ***(ZIP)***

## Acknowledgments

We thank all Lange lab members for their valuable feedback upon testing the framework in various projects and experimental settings.

## Abbreviations

AUC: Area Under the Curve
BH: Benjamini-Hochberg
COPF: Correlation-based functional Proteoform
CV: Coefficient of Variation
dPFs: Quantitatively Differential Proteoforms
FDR: False Discovery Rate
kNN: k-Nearest Neighbours
LFC: Log Fold Change
MCC: Matthews Correlation Coefficient
PeCorA: Peptide Correlation Analysis
PTM: Post-Translational Modifications
QuEStVar: Quantitative Exploration of Stability and Variability
ROC: Receiver Operating Characteristic
WLS: Weighted Least Squares

## References

(1) Smith, L. M.; Agar, J. N.; Chamot-Rooke, J.; Danis, P. O.; Ge, Y.; Loo, J. A.; Paša-Tolić, L.; Tsybin, Y. O.; Kelleher, N. L.; Proteomics, C. for T.-D. The Human Proteoform Project: Defining the Human Proteome. Sci. Adv. 2021, 7 (46), eabk0734. 10.1126/sciadv.abk0734.

(2) Smith, L. M.; Kelleher, N. L.; Consortium for Top Down Proteomics. Proteoform: A Single Term Describing Protein Complexity. Nat. Methods 2013, 10 (3), 186–187. 10.1038/nmeth.2369.

(3) Aebersold, R.; Agar, J. N.; Amster, I. J.; Baker, M. S.; Bertozzi, C. R.; Boja, E. S.; Costello, C. E.; Cravatt, B. F.; Fenselau, C.; Garcia, B. A.; Ge, Y.; Gunawardena, J.; Hendrickson, R. C.; Hergenrother, P. J.; Huber, C. G.; Ivanov, A. R.; Jensen, O. N.; Jewett, M. C.; Kelleher, N. L.; Kiessling, L. L.; Krogan, N. J.; Larsen, M. R.; Loo, J. A.; Ogorzalek Loo, R. R.; Lundberg, E.; MacCoss, M. J.; Mallick, P.; Mootha, V. K.; Mrksich, M.; Muir, T. W.; Patrie, S. M.; Pesavento, J. J.; Pitteri, S. J.; Rodriguez, H.; Saghatelian, A.; Sandoval, W.; Schlüter, H.; Sechi, S.; Slavoff, S. A.; Smith, L. M.; Snyder, M. P.; Thomas, P. M.; Uhlén, M.; Van Eyk, J. E.; Vidal, M.; Walt, D. R.; White, F. M.; Williams, E. R.; Wohlschlager, T.; Wysocki, V. H.; Yates, N. A.; Young, N. L.; Zhang, B. How Many Human Proteoforms Are There? Nat. Chem. Biol. 2018, 14 (3), 206–214. 10.1038/nchembio.2576.

(4) Carbonara, K.; Andonovski, M.; Coorssen, J. R. Proteomes Are of Proteoforms: Embracing the Complexity. Proteomes 2021, 9 (3), 38. 10.3390/proteomes9030038.

(5) Faria, M.; Félix, D.; Domingues, R.; Bugalho, M. J.; Matos, P.; Silva, A. L. Antagonistic Effects of RAC1 and Tumor-Related RAC1b on NIS Expression in Thyroid. J. Mol. Endocrinol. 2019, 63 (4), 309–320. 10.1530/JME-19-0195.

(6) Degan, S. E.; Gelman, I. H. Emerging Roles for AKT Isoform Preference in Cancer Progression Pathways. Mol. Cancer Res. 2021, 19 (8), 1251–1257. 10.1158/1541-7786.MCR-20-1066.

(7) McCool, E. N.; Xu, T.; Chen, W.; Beller, N. C.; Nolan, S. M.; Hummon, A. B.; Liu, X.; Sun, L. Deep Top-down Proteomics Revealed Significant Proteoform-Level Differences between Metastatic and Nonmetastatic Colorectal Cancer Cells. Sci. Adv. 2022, 8 (51), eabq6348. 10.1126/sciadv.abq6348.

(8) Adams, L. M.; DeHart, C. J.; Drown, B. S.; Anderson, L. C.; Bocik, W.; Boja, E. S.; Hiltke, T. M.; Hendrickson, C. L.; Rodriguez, H.; Caldwell, M.; Vafabakhsh, R.; Kelleher, N. L. Mapping the KRAS Proteoform Landscape in Colorectal Cancer Identifies Truncated KRAS4B That Decreases MAPK Signaling. J. Biol. Chem. 2022, 299 (1), 102768. 10.1016/j.jbc.2022.102768.

(9) Chetta, M.; Basile, A.; Tarsitano, M.; Rivieccio, M.; Oro, M.; Capitanio, N.; Bukvic, N.; Priolo, M.; Rosati, A. The Target Therapy Hyperbole: “KRAS (p.G12C)”—the Simplification of a Complex Biological Problem. Cancers 2024, 16 (13), 2389. 10.3390/cancers16132389.

(10) Jiang, Y.; Rex, D. A. B.; Schuster, D.; Neely, B. A.; Rosano, G. L.; Volkmar, N.; Momenzadeh, A.; Peters-Clarke, T. M.; Egbert, S. B.; Kreimer, S.; Doud, E. H.; Crook, O. M.; Yadav, A. K.; Vanuopadath, M.; Hegeman, A. D.; Mayta, M. L.; Duboff, A. G.; Riley, N. M.; Moritz, R. L.; Meyer, J. G. Comprehensive Overview of Bottom-up Proteomics Using Mass Spectrometry. ACS Meas. Sci. Au 2024, 4 (4), 338–417. 10.1021/acsmeasuresciau.3c00068.

(11) Ivanov, M. V.; Solovyeva, E. M.; Bubis, J. A.; Gorshkov, M. V. Improving the Protein Inference from Bottom-up Proteomic Data Using Identifications from MS1 Spectra. J. Am. Soc. Mass Spectrom. 2021, 32 (5), 1258–1262. 10.1021/jasms.1c00061.

(12) Nesvizhskii, A. I.; Aebersold, R. Interpretation of Shotgun Proteomic Data: The Protein Inference Problem. Mol. Cell. Proteomics 2005, 4 (10), 1419–1440. 10.1074/mcp.R500012-MCP200.

(13) He, Z.; Huang, T.; Liu, X.; Zhu, P.; Teng, B.; Deng, S. Protein Inference: A Protein Quantification Perspective. Comput. Biol. Chem. 2016, 63, 21–29. 10.1016/j.compbiolchem.2016.02.006.

(14) Nesvizhskii, A. I.; Keller, A.; Kolker, E.; Aebersold, R. A Statistical Model for Identifying Proteins by Tandem Mass Spectrometry. Anal. Chem. 2003, 75 (17), 4646–4658. 10.1021/ac0341261.

(15) Plubell, D. L.; Käll, L.; Webb-Robertson, B.-J.; Bramer, L. M.; Ives, A.; Kelleher, N. L.; Smith, L. M.; Montine, T. J.; Wu, C. C.; MacCoss, M. J. Putting Humpty Dumpty Back Together Again: What Does Protein Quantification Mean in Bottom-up Proteomics? J. Proteome Res. 2022, 21 (4), 891–898. 10.1021/acs.jproteome.1c00894.

(16) Saltzman, A. B.; Leng, M.; Bhatt, B.; Singh, P.; Chan, D. W.; Dobrolecki, L.; Chandrasekaran, H.; Choi, J. M.; Jain, A.; Jung, S. Y.; Lewis, M. T.; Ellis, M. J.; Malovannaya, A. gpGrouper: A Peptide Grouping Algorithm for Gene-Centric Inference and Quantitation of Bottom-up Proteomics Data. Mol. Cell. Proteomics 2018, 17 (11), 2270–2283. 10.1074/mcp.TIR118.000850.

(17) Schork, K.; Turewicz, M.; Uszkoreit, J.; Rahnenführer, J.; Eisenacher, M. Characterization of Peptide-Protein Relationships in Protein Ambiguity Groups via Bipartite Graphs. PLOS One 2022, 17 (10), e0276401. 10.1371/journal.pone.0276401.

(18) Brown, K. A.; Melby, J. A.; Roberts, D. S.; Ge, Y. Top-down Proteomics: Challenges, Innovations, and Applications in Basic and Clinical Research. Expert Rev. Proteomics 2020, 17 (10), 719–733. 10.1080/14789450.2020.1855982.

(19) Dou, Y.; Liu, Y.; Yi, X.; Olsen, L. K.; Zhu, H.; Gao, Q.; Zhou, H.; Zhang, B. SEPepQuant Enhances the Detection of Possible Isoform Regulations in Shotgun Proteomics. Nat. Commun. 2023, 14 (1), 5809. 10.1038/s41467-023-41558-2.

(20) Ammar, C.; Thielert, M.; Weiss, C. A. M.; Rodriguez, E. H.; Strauss, M. T.; Rosenberger, F. A.; Zeng, W.-F.; Mann, M. Tree-Based Quantification Infers Proteoform Regulation in Bottom-up Proteomics Data. Biorxiv 2025. 10.1101/2025.03.06.641844.

(21) Dermit, M.; Peters-Clarke, T. M.; Shishkova, E.; Meyer, J. G. Peptide Correlation Analysis (Pecora) Reveals Differential Proteoform Regulation. J. Proteome Res. 2021, 20 (4), 1972–1980. 10.1021/acs.jproteome.0c00602.

(22) Bludau, I.; Frank, M.; Dörig, C.; Cai, Y.; Heusel, M.; Rosenberger, G.; Picotti, P.; Collins, B. C.; Röst, H.; Aebersold, R. Systematic Detection of Functional Proteoform Groups from Bottom-up Proteomic Datasets. Nat. Commun. 2021, 12 (1), 3810. 10.1038/s41467-021-24030-x.

(23) Staniak, M.; Huang, T.; Figueroa-Navedo, A. M.; Kohler, D.; Choi, M.; Hinkle, T.; Kleinheinz, T.; Blake, R.; Rose, C. M.; Xu, Y.; Jean Beltran, P. M.; Xue, L.; Bogdan, M.; Vitek, O. Relative Quantification of Proteins and Post-Translational Modifications in Proteomic Experiments with Shared Peptides: A Weight-Based Approach. Bioinformatics 2025, 41 (3), btaf046. 10.1093/bioinformatics/btaf046.

(24) Collins, B. C.; Hunter, C. L.; Liu, Y.; Schilling, B.; Rosenberger, G.; Bader, S. L.; Chan, D. W.; Gibson, B. W.; Gingras, A.-C.; Held, J. M.; Hirayama-Kurogi, M.; Hou, G.; Krisp, C.; Larsen, B.; Lin, L.; Liu, S.; Molloy, M. P.; Moritz, R. L.; Ohtsuki, S.; Schlapbach, R.; Selevsek, N.; Thomas, S. N.; Tzeng, S.-C.; Zhang, H.; Aebersold, R. Multi-Laboratory Assessment of Reproducibility, Qualitative and Quantitative Performance of SWATH-Mass Spectrometry. Nat. Commun. 2017, 8 (1), 291. 10.1038/s41467-017-00249-5.

(25) Tyanova, S.; Cox, J. Perseus: A Bioinformatics Platform for Integrative Analysis of Proteomics Data in Cancer Research. Methods Mol. Biol. 2018, 1711, 133–148. 10.1007/978-1-4939-7493-1_7.

(26) Chicco, D.; Jurman, G. The Advantages of the Matthews Correlation Coefficient (MCC) over F1 Score and Accuracy in Binary Classification Evaluation. BMC Genomics 2020, 21 (1), 6. 10.1186/s12864-019-6413-7.

(27) Chicco, D.; Jurman, G. The Matthews Correlation Coefficient (MCC) Should Replace the ROC AUC as the Standard Metric for Assessing Binary Classification. Biodata Min. 2023, 16 (1), 4. 10.1186/s13040-023-00322-4.

(28) Tomin, T.; Honeder, S. E.; Liesinger, L.; Gremel, D.; Retzl, B.; Lindenmann, J.; Brcic, L.; Schittmayer, M.; Birner-Gruenberger, R. Increased Antioxidative Defense and Reduced Advanced Glycation End-Product Formation by Metabolic Adaptation in Non-Small-Cell-Lung-Cancer Patients. Nat. Commun. 2025, 16 (1), 5157. 10.1038/s41467-025-60326-y.

(29) Benjamini, Y.; Hochberg, Y. Controlling the False Discovery Rate: A Practical and Powerful Approach to Multiple Testing. J. R. Stat. Soc. Ser. B Methodol. 1995, 57 (1), 289–300. 10.1111/j.2517-6161.1995.tb02031.x.

(30) The UniProt Consortium. UniProt: The Universal Protein Knowledgebase in 2025. Nucleic Acids Res. 2025, 53 (D1), D609–D617. 10.1093/nar/gkae1010.

(31) Rawlings, N. D.; Barrett, A. J.; Thomas, P. D.; Huang, X.; Bateman, A.; Finn, R. D. The MEROPS Database of Proteolytic Enzymes, Their Substrates and Inhibitors in 2017 and a Comparison with Peptidases in the PANTHER Database. Nucleic Acids Res. 2018, 46 (D1), D624–D632. 10.1093/nar/gkx1134.

(32) Huang, H.; Arighi, C. N.; Ross, K. E.; Ren, J.; Li, G.; Chen, S.-C.; Wang, Q.; Cowart, J.; Vijay-Shanker, K.; Wu, C. H. iPTMnet: An Integrated Resource for Protein Post-Translational Modification Network Discovery. Nucleic Acids Res. 2018, 46 (D1), D542–D550. 10.1093/nar/gkx1104.

(33) Lange, P. F.; Overall, C. M. TopFIND, a Knowledgebase Linking Protein Termini with Function. Nat. Methods 2011, 8 (9), 703–704. 10.1038/nmeth.1669.

(34) Fortelny, N.; Yang, S.; Pavlidis, P.; Lange, P. F.; Overall, C. M. Proteome TopFIND 3.0 with TopFINDer and PathFINDer: Database and Analysis Tools for the Association of Protein Termini to Pre- and Post-Translational Events. Nucleic Acids Res. 2015, 43 (Database issue), D290–7. 10.1093/nar/gku1012.

(35) Ergin, E. K.; Myung, J. J. K.; Lange, P. F. Snapshot of the Quantitative Protein Stability Analysis in Cancer Cell Lines Using QuEStVar. Zenodo 2024. 10.5281/zenodo.10694635.

(36) Ergin, E. K.; Myung, J. J. K.; Lange, P. F. Statistical Testing for Protein Equivalence Identifies Core Functional Modules Conserved across 360 Cancer Cell Lines and Presents a General Approach to Investigating Biological Systems. J. Proteome Res. 2024, 23 (6), 2169–2185. 10.1021/acs.jproteome.4c00131.

(37) Kolberg, L.; Raudvere, U.; Kuzmin, I.; Vilo, J.; Peterson, H. Gprofiler2 -- an R Package for Gene List Functional Enrichment Analysis and Namespace Conversion Toolset g:Profiler. F1000Research 2020, 9, 709. 10.12688/f1000research.24956.2.

(38) Harris, C. R.; Millman, K. J.; van der Walt, S. J.; Gommers, R.; Virtanen, P.; Cournapeau, D.; Wieser, E.; Taylor, J.; Berg, S.; Smith, N. J.; Kern, R.; Picus, M.; Hoyer, S.; van Kerkwijk, M. H.; Brett, M.; Haldane, A.; Del Río, J. F.; Wiebe, M.; Peterson, P.; Gérard-Marchant, P.; Sheppard, K.; Reddy, T.; Weckesser, W.; Abbasi, H.; Gohlke, C.; Oliphant, T. E. Array Programming with NumPy. Nature 2020, 585 (7825), 357–362. 10.1038/s41586-020-2649-2.

(39) McKinney, W. Data Structures for Statistical Computing in Python. In Proceedings of the 9th Python in Science Conference; Proceedings of the python in science conference; SciPy: Austin, Texas, 2010; pp 56–61. 10.25080/Majora-92bf1922-00a.

(40) Seabold, S.; Perktold, J. Statsmodels: Econometric and Statistical Modeling with Python. In Proceedings of the 9th Python in Science Conference; Proceedings of the python in science conference; SciPy: Austin, Texas, 2010; pp 92–96. 10.25080/Majora-92bf1922-011.

(41) Pedregosa, F.; Varoquaux, G.; Gramfort, A.; Michel, V.; Thirion, B.; Grisel, O.; Blondel, M.; Prettenhofer, P.; Weiss, R.; Dubourg, V.; Vanderplas, J.; Passos, A.; Cournapeau, D.; Brucher, M.; Perrot, M.; Duchesnay, É. Scikit-Learn: Machine Learning in Python. J. Mach. Learn. Res. 2011.

(42) Virtanen, P.; Gommers, R.; Oliphant, T. E.; Haberland, M.; Reddy, T.; Cournapeau, D.; Burovski, E.; Peterson, P.; Weckesser, W.; Bright, J.; van der Walt, S. J.; Brett, M.; Wilson, J.; Millman, K. J.; Mayorov, N.; Nelson, A. R. J.; Jones, E.; Kern, R.; Larson, E.; Carey, C. J.; Polat, İ.; Feng, Y.; Moore, E. W.; VanderPlas, J.; Laxalde, D.; Perktold, J.; Cimrman, R.; Henriksen, I.; Quintero, E. A.; Harris, C. R.; Archibald, A. M.; Ribeiro, A. H.; Pedregosa, F.; van Mulbregt, P.; SciPy 1.0 Contributors. SciPy 1.0: Fundamental Algorithms for Scientific Computing in Python. Nat. Methods 2020, 17 (3), 261–272. 10.1038/s41592-019-0686-2.

(43) Hunter, J. D. Matplotlib: A 2D Graphics Environment. Comput. Sci. Eng. 2007, 9 (3), 90–95. 10.1109/MCSE.2007.55.

(44) Waskom, M. Seaborn: Statistical Data Visualization. J. Open Source Softw. 2021, 6 (60), 3021. 10.21105/joss.03021.

(45) Heusel, M.; Bludau, I.; Rosenberger, G.; Hafen, R.; Frank, M.; Banaei-Esfahani, A.; van Drogen, A.; Collins, B. C.; Gstaiger, M.; Aebersold, R. Complex-Centric Proteome Profiling by SEC-SWATH-MS. Mol. Syst. Biol. 2019, 15 (1), e8438. 10.15252/msb.20188438.

(46) Audain, E.; Uszkoreit, J.; Sachsenberg, T.; Pfeuffer, J.; Liang, X.; Hermjakob, H.; Sanchez, A.; Eisenacher, M.; Reinert, K.; Tabb, D. L.; Kohlbacher, O.; Perez-Riverol, Y. In-Depth Analysis of Protein Inference Algorithms Using Multiple Search Engines and Well-Defined Metrics. J. Proteomics 2017, 150, 170–182. 10.1016/j.jprot.2016.08.002.

(47) Bamberger, C.; Martínez-Bartolomé, S.; Montgomery, M.; Pankow, S.; Hulleman, J. D.; Kelly, J. W.; Yates, J. R. Deducing the Presence of Proteins and Proteoforms in Quantitative Proteomics. Nat. Commun. 2018, 9 (1), 2320. 10.1038/s41467-018-04411-5.

(48) Po, A.; Eyers, C. E. Top-down Proteomics and the Challenges of True Proteoform Characterization. J. Proteome Res. 2023, 22 (12), 3663–3675. 10.1021/acs.jproteome.3c00416.

(49) Zahn-Zabal, M.; Michel, P.-A.; Gateau, A.; Nikitin, F.; Schaeffer, M.; Audot, E.; Gaudet, P.; Duek, P. D.; Teixeira, D.; Rech de Laval, V.; Samarasinghe, K.; Bairoch, A.; Lane, L. The neXtProt Knowledgebase in 2020: Data, Tools and Usability Improvements. Nucleic Acids Res. 2020, 48 (D1), D328–D334. 10.1093/nar/gkz995.

