## Supplementary figures for "ProteoForge: An Imputation-Aware Framework for Differential Proteoform Discovery in Bottom-Up Proteomics"

### Table of Contents

**Supplemental Figures**


- **Figure S1:** ProteoForge demonstrates superior performance for discordant peptide identification in the SWATH-MS InterLab benchmark.
- **Figure S2:** ProteoForge accurately groups co-varying peptides in the SWATH-MS Interlab benchmark dataset.
- **Figure S3:** ProteoForge maintains high performance on imputed data in a discordant peptide identification benchmark.
- **Figure S4:** ProteoForge's peptide grouping performance is resilient to data imputation.
- **Figure S5:** ProteoForge's performance remains robust even at extreme levels of missingness in protein and peptide data.
- **Figure S6:** Higher perturbations lead to better performance across methods and benchmarks.
- **Figure S7:** Performance sensitivity to experimental design complexity, perturbation direction, and feature overlap.
- **Figure S8:** Differential proteoform (dPF) based protein quantification improves the separation of experimental groups in PCA.


**Supplemental Notes**


- **Note 1:** Selection of linear estimators for imputation-aware peptide analysis within ProteoForge


**Supplemental Analysis Notebooks**


- **Notebook S1:** Discordant peptide identification benchmark using SWATH-MS Interlab Data
- **Notebook S2:** Peptide grouping benchmark using SWATH-MS Interlab Data
- **Notebook S3:** Demo figure assembly for Figure 1
- **Notebook S4:** Generation of simulated dataset for benchmarking
- **Notebook S5:** Peptide identification benchmark on simulated datasets
- **Notebook S6:** Peptide grouping benchmark on simulated datasets
- **Notebook S7:** Comparison of linear models offered by ProteoForge
- **Notebook S8:** Highly summarized figure assembly for Figure 2
- **Notebook S9:** Data preparation for Hypoxia Experiment of H358 cell lines
- **Notebook S10:** Applying ProteoForge to Hypoxia vs Normoxia (48hr timepoint)
- **Notebook S11:** Applying ProteoForge to Hypoxia vs Normoxia (72hr timepoint)
- **Notebook S12:** Exploring ProteoForge results from 48hr timepoint
- **Notebook S13:** Exploring ProteoForge results from 72hr timepoint
- **Notebook S14:** Protein-level downstream analysis comparison
- **Notebook S15:** Figure assembly for 48hr and 72hr timepoint summaries

### Abbreviations

| | |
|---|---|
| **AUC** | Area Under the Curve |
| **COPF** | **Co**rrelation-based functional **P**roteo**F**orm assessment |
| **dPF** | Quantitatively Differential Proteoforms |
| **FPR** | False Positive Rate |
| **MCC** | Matthews Correlation Coefficient |
| **PCA** | Principal Component Analysis |
| **PeCorA** | **Pe**ptide **Cor**relation **A**nalysis |
| **ProteoForge** | **Proteo**form **Forge**r |
| **ROC** | Receiver Operating Characteristic |
| **TPR** | True Positive Rate |

### Supplemental Figures

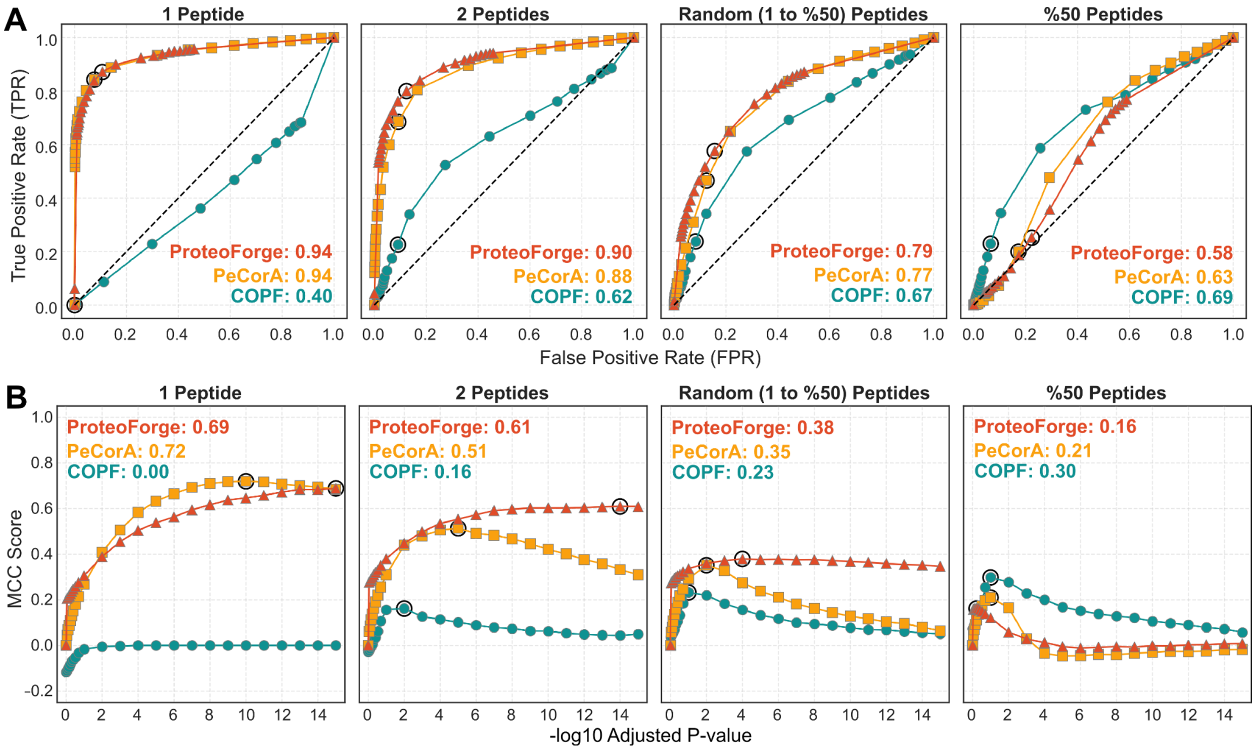


**Figure S1.** ProteoForge shows superior performance for discordant peptide identification in the SWATH-MS Interlab benchmark.

The performance of ProteoForge (red triangles), PeCorA (yellow squares), and COPF (teal circles) for identifying discordant peptides was evaluated in the SWATH-MS Interlab benchmark dataset. The analysis covered four perturbation scenarios: (i) 1 peptide, (ii) 2 peptides, (iii) 1-50% peptides (random), or (iv) 50% of peptides perturbed per protein. **(A)** ROC curves plot the TPR against the FPR for each scenario, with the AUC listed for each method. The dashed diagonal line indicates a random classifier (AUC = 0.5). **(B)** The MCC score is plotted against the significance threshold ($-\log_{10}$ Adjusted P-value), with the maximum MCC score for each method listed. In all panels, the black circle on each curve indicates the performance at the threshold corresponding to the maximum MCC score.

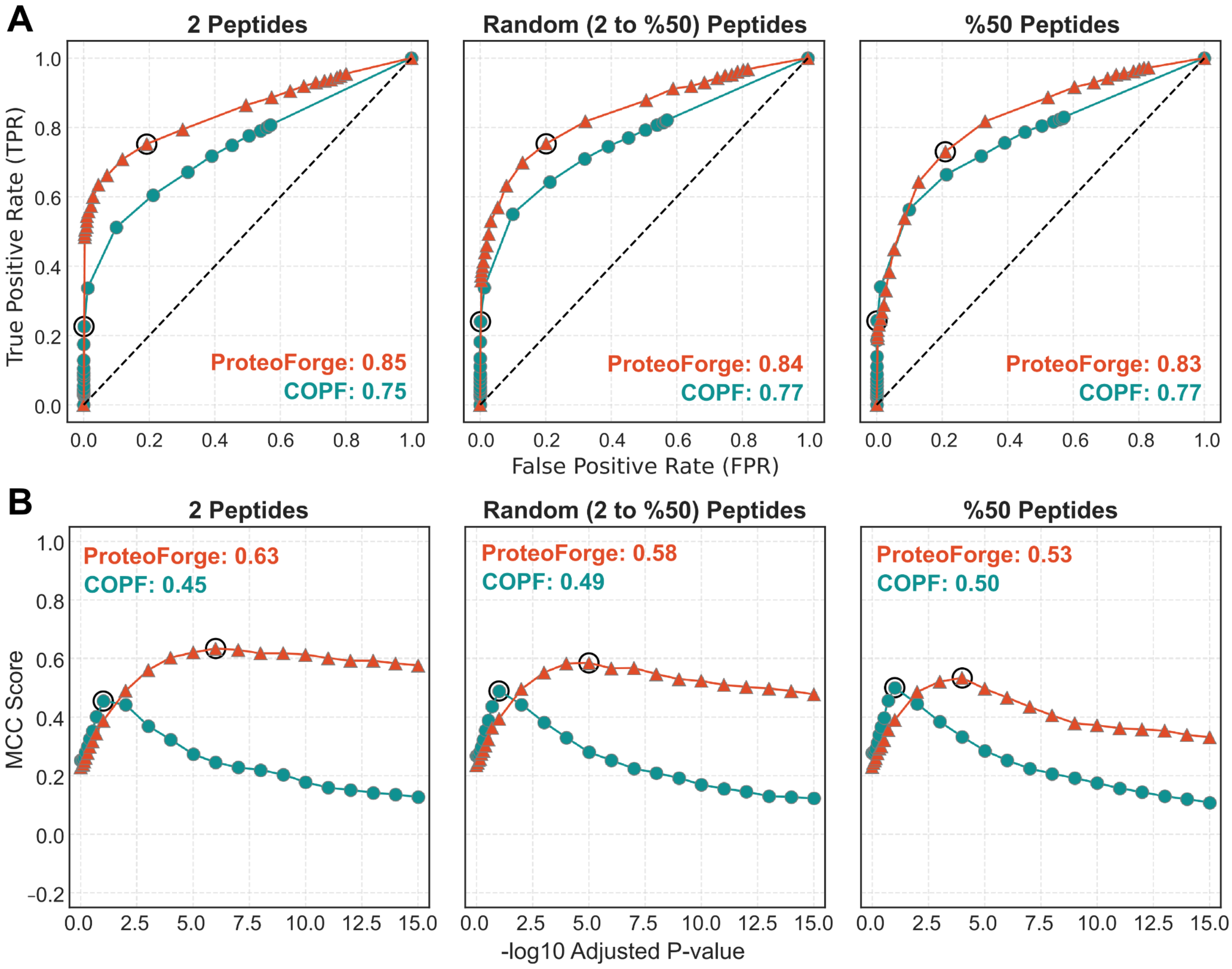


**Figure S2.** ProteoForge accurately groups co-varying peptides in the SWATH-MS Interlab benchmark dataset.

The ability of ProteoForge and COPF to group co-varying peptides and infer potential proteoforms was assessed using the SWATH-MS Interlab benchmark dataset across three perturbation scenarios: (i) 2 peptides, (ii) a random number of peptides (2-50%), or (iii) 50% of peptides perturbed per target protein. PeCorA was excluded due to its lack of peptide grouping functionality. **(A)** ROC curves compare the TPR and FPR for peptide grouping, with AUC values reported for ProteoForge (red triangles) and COPF (teal circles). The dashed diagonal line represents the performance of a random classifier. **(B)** The MCC score is plotted against the significance cutoff (-$\log_{10}$ Adjusted P-value), with the maximum MCC for each method reported. Black circles highlight performance at the threshold yielding the maximum MCC score.

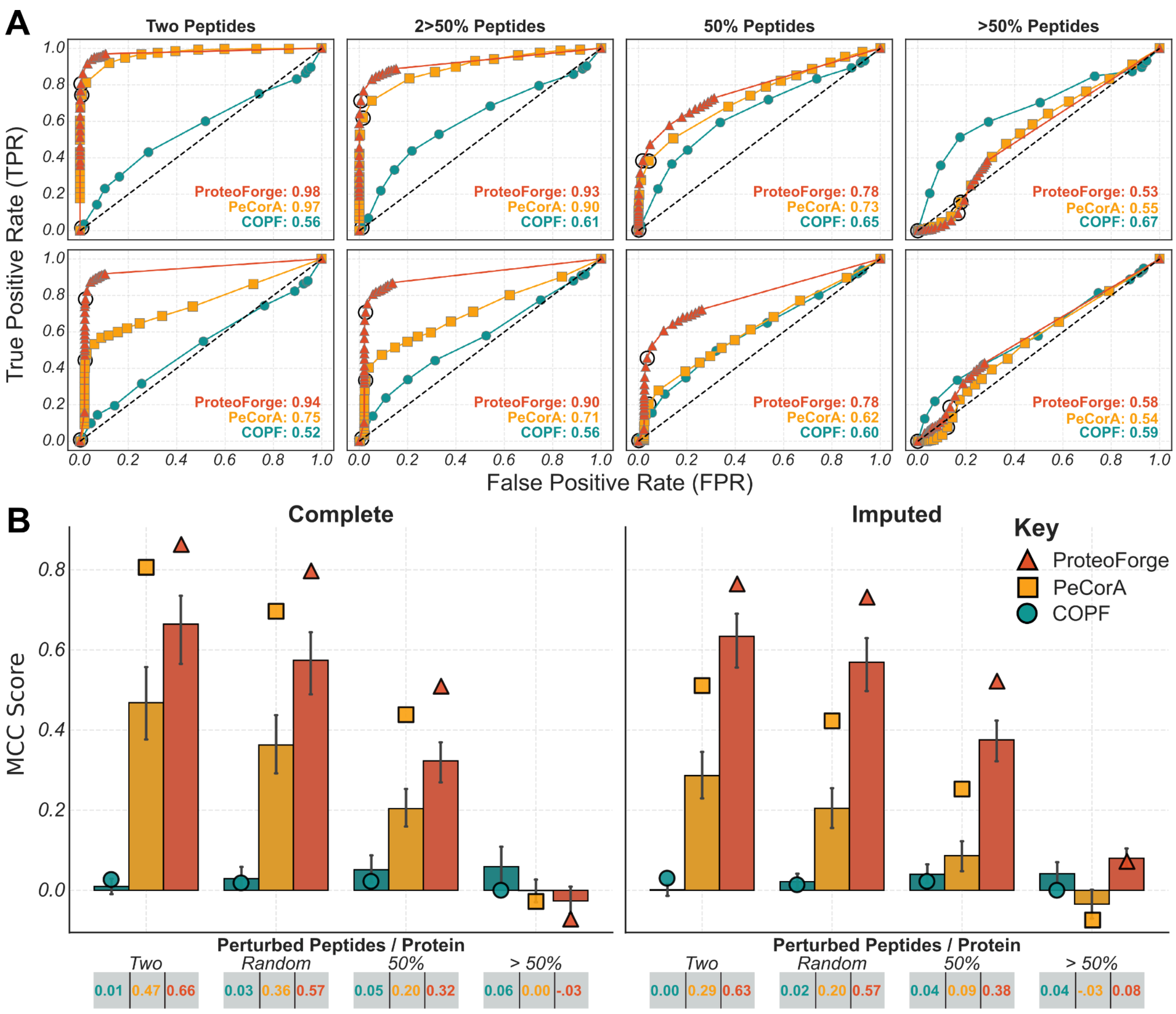


**Figure S3.** ProteoForge maintains high performance on imputed data in a discordant peptide identification benchmark.

To assess the impact of data imputation on **discordant peptide identification**, three methods were benchmarked on a fully observed 'Complete' dataset and a version where missing values were programmatically introduced and then imputed ('Imputed'). This comparison was performed across four scenarios with varying percentages of peptides perturbed per protein: (i) Two Peptides, (ii) 2 to 50%, (iii) exactly 50%, and (iv) more than 50%. **(A)** ROC curves plot the TPR versus the FPR for the 'Complete' (top row) and 'Imputed' (bottom row) datasets, with the AUC reported for ProteoForge (red triangles), PeCorA (yellow squares), and COPF (teal circles). **(B)** Bar plots show the mean MCC scores across all significance thresholds for both datasets, with error bars representing the 95% confidence interval. Circular points indicate the MCC score at an adjusted p-value of 0.001, and horizontal dashed lines provide a qualitative performance reference.

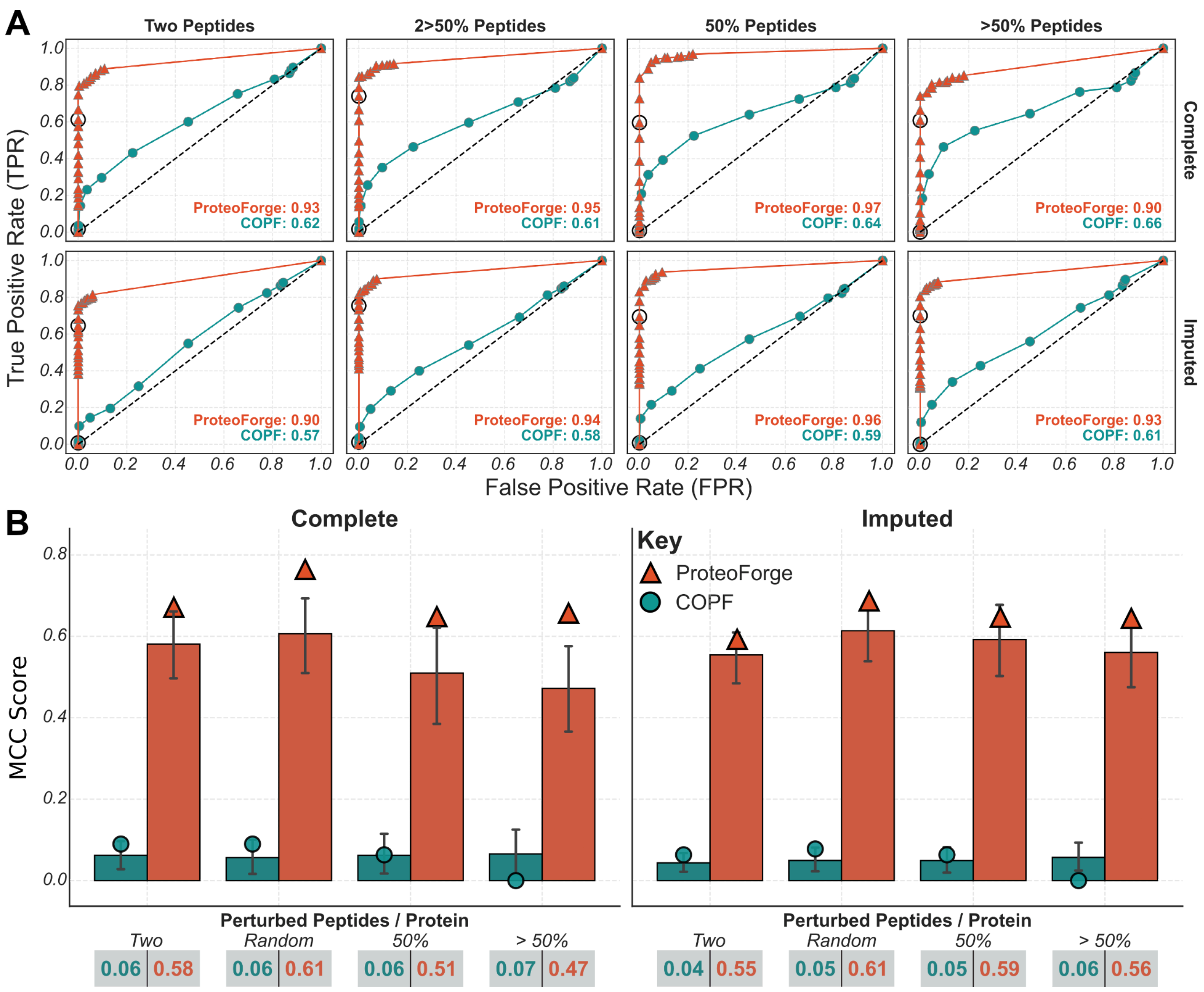


**Figure S4.** ProteoForge's peptide grouping performance is resilient to data imputation.

To assess the impact of data imputation on the **peptide grouping task**, the performance of ProteoForge and COPF was benchmarked on a fully observed 'Complete' dataset and an 'Imputed' version where missing values were programmatically introduced and then filled. This comparison was performed across four perturbation scenarios: (i) Two Peptides, (ii) 2 to 50% of peptides, (iii) exactly 50% of peptides, and (iv) more than 50% of peptides perturbed per protein. PeCorA was omitted as it lacks peptide grouping functionality. **(A)** ROC curves plot the TPR versus the FPR for the 'Complete' (top row) and 'Imputed' (bottom row) datasets, with the AUC reported for ProteoForge (red triangles) and COPF (teal circles). **(B)** Bar plots show the mean MCC scores across all significance thresholds, with error bars representing the 95% confidence interval. Circular points indicate the MCC score at an adjusted p-value of 0.001, and horizontal dashed lines serve as a qualitative performance reference.

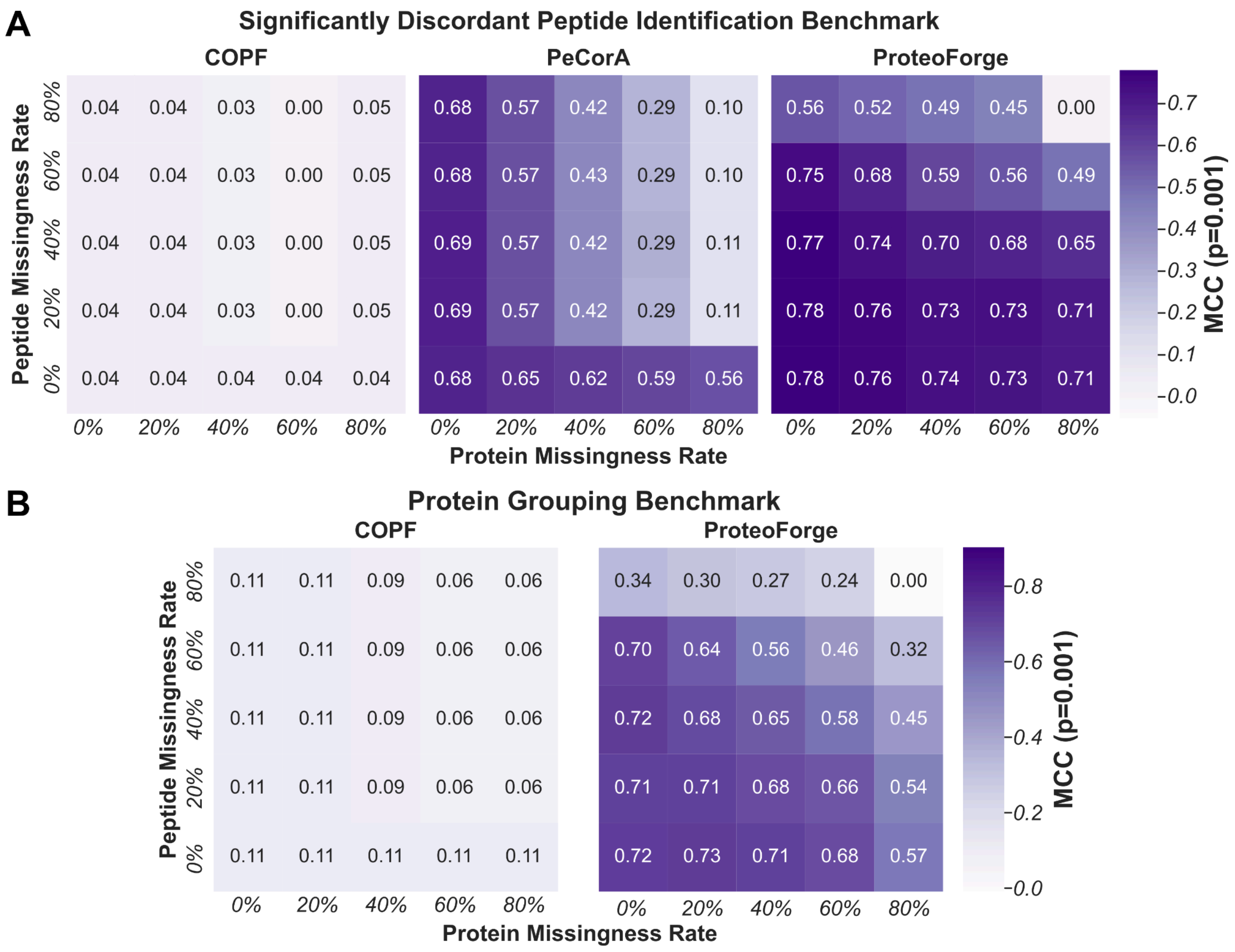


**Figure S5.** ProteoForge's performance remains robust even at extreme levels of protein and peptide data missingness.

To evaluate robustness to data missingness, a simulation was performed using the "Random (2-50%) Peptides" perturbation scenario as a base, with missingness introduced at both the protein (0-80%) and peptide (0-80%) levels. **(A)** Heatmaps show the MCC at a fixed adjusted p-value of 0.001 for the **Significantly Discordant Peptide Identification** benchmark for COPF, PeCorA, and ProteoForge. The colour scale indicates the MCC score, with darker purple representing better performance. **(B)** Heatmaps display the MCC score (adjusted p-value = 0.001) for the **Protein Grouping** benchmark for COPF and ProteoForge, where darker shades indicate higher MCC scores.

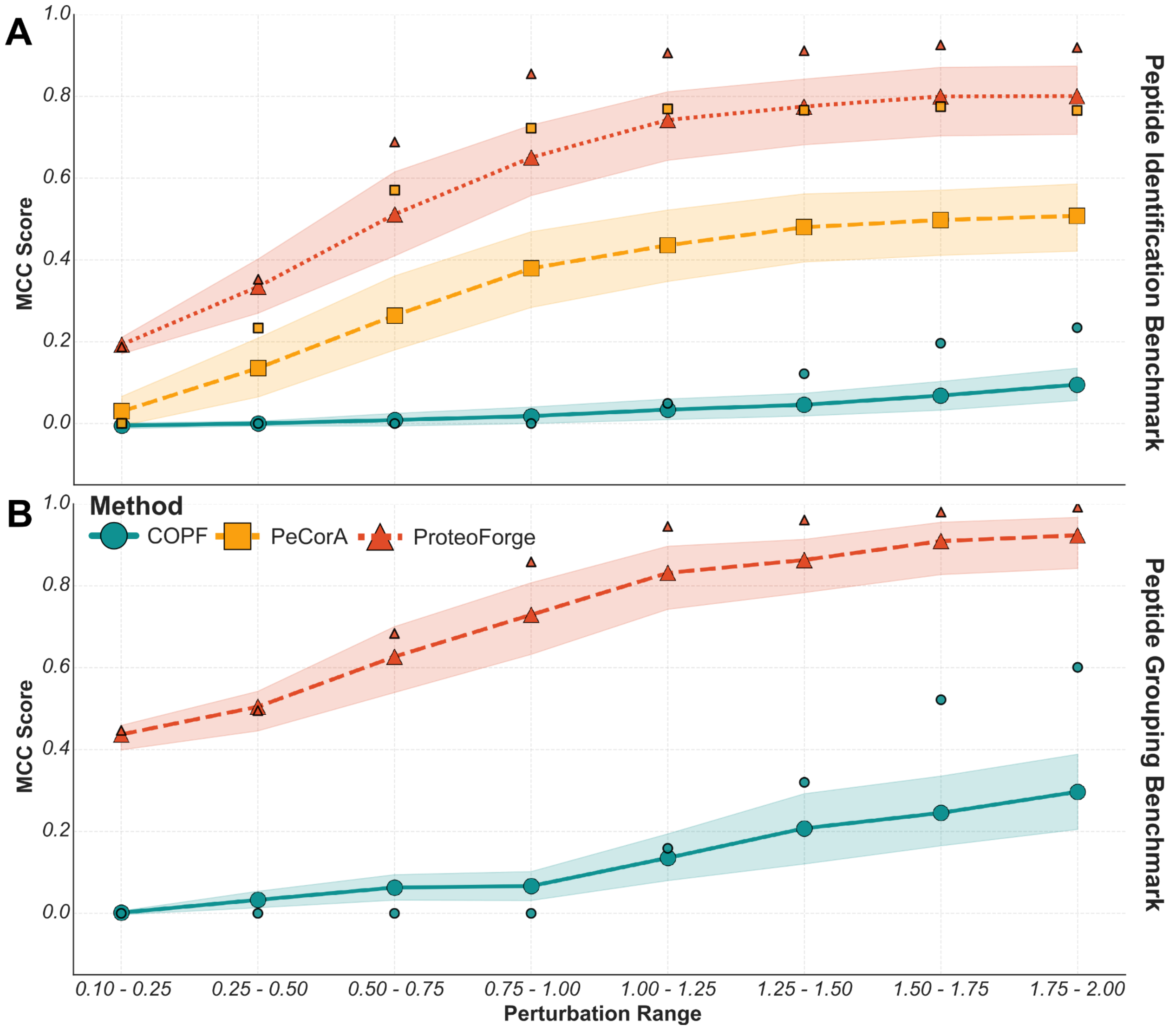


**Figure S6.** Higher perturbations lead to better performance across methods and benchmarks.

Using the imputed 'Random (2-50%) Peptides' dataset, this analysis evaluates method performance as a function of perturbation magnitude, where the x-axis represents the $\log_2$ fold change range from which perturbations were randomly selected. Lines represent the mean MCC scores, with shaded error bands indicating the 95% confidence interval; points with black edges denote the MCC score at an adjusted p-value of 0.001. Horizontal dashed lines provide a qualitative performance reference. **(A)** Performance on the **Discordant Peptide Identification** benchmark is shown for ProteoForge (red triangles), PeCorA (yellow squares), and COPF (teal circles). **(B)** Performance on the **Peptide Grouping** benchmark is shown for ProteoForge and COPF; PeCorA is omitted as it does not support this functionality.

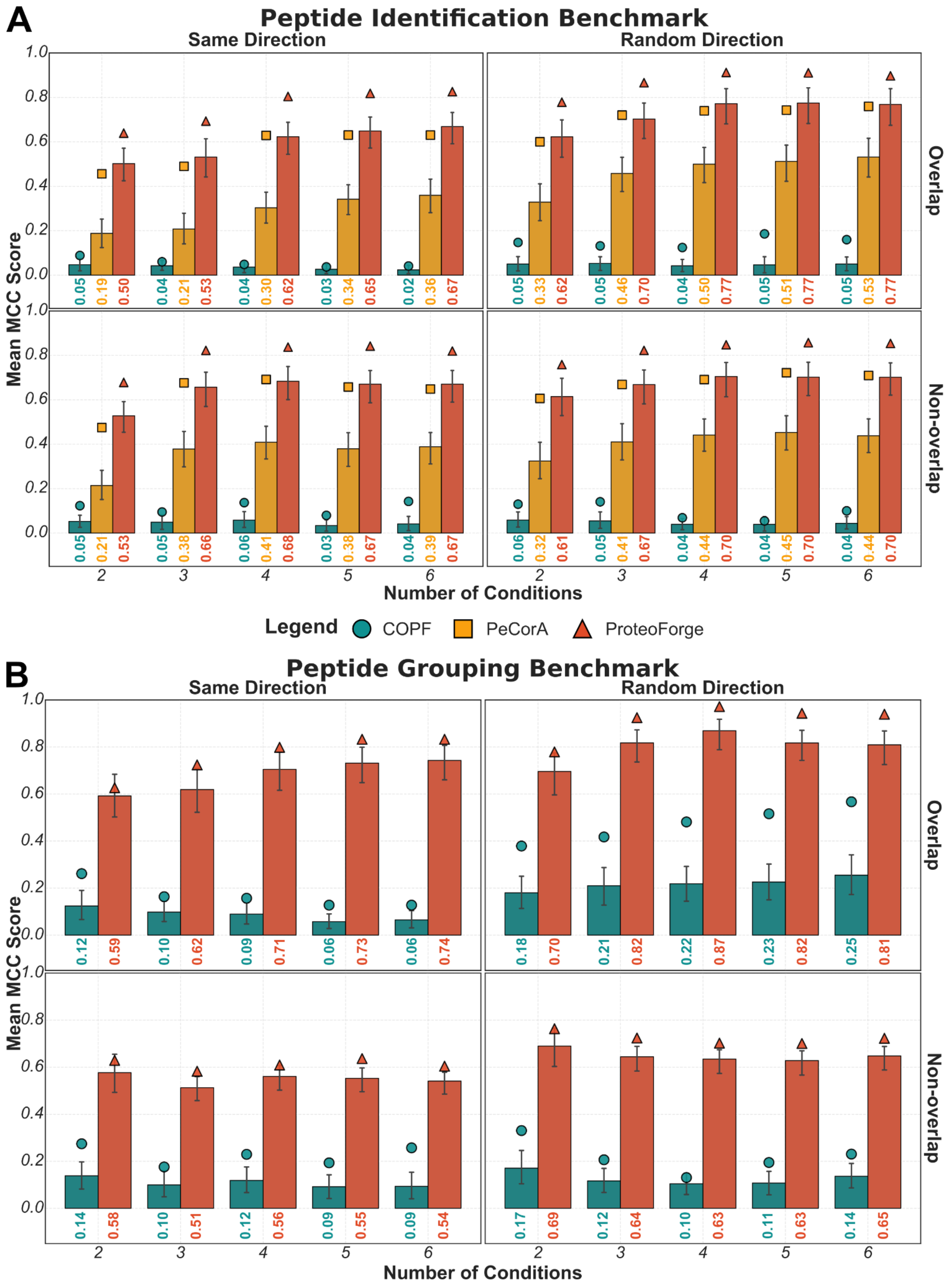


**Figure S7.** Performance sensitivity to experimental design complexity, perturbation direction, and feature overlap.

This simulation evaluates method performance on an imputed dataset as the number of experimental conditions increases from 2 to 6 (x-axis). The analysis is stratified in a 2x2 grid comparing perturbation direction (all 'Same Direction' vs. 'Random Direction') and perturbation overlap ('Overlap' vs. 'Non-overlap' between conditions). Bar height represents the mean MCC across all significance thresholds, with error bars indicating the 95% confidence interval; overlaid points show the MCC at an adjusted p-value of 0.001. **(A)** Results for the **Discordant Peptide Identification** benchmark are shown for COPF (teal), PeCorA (yellow), and ProteoForge (red). **(B)** Results for the **Peptide Grouping** benchmark are shown for COPF and ProteoForge.

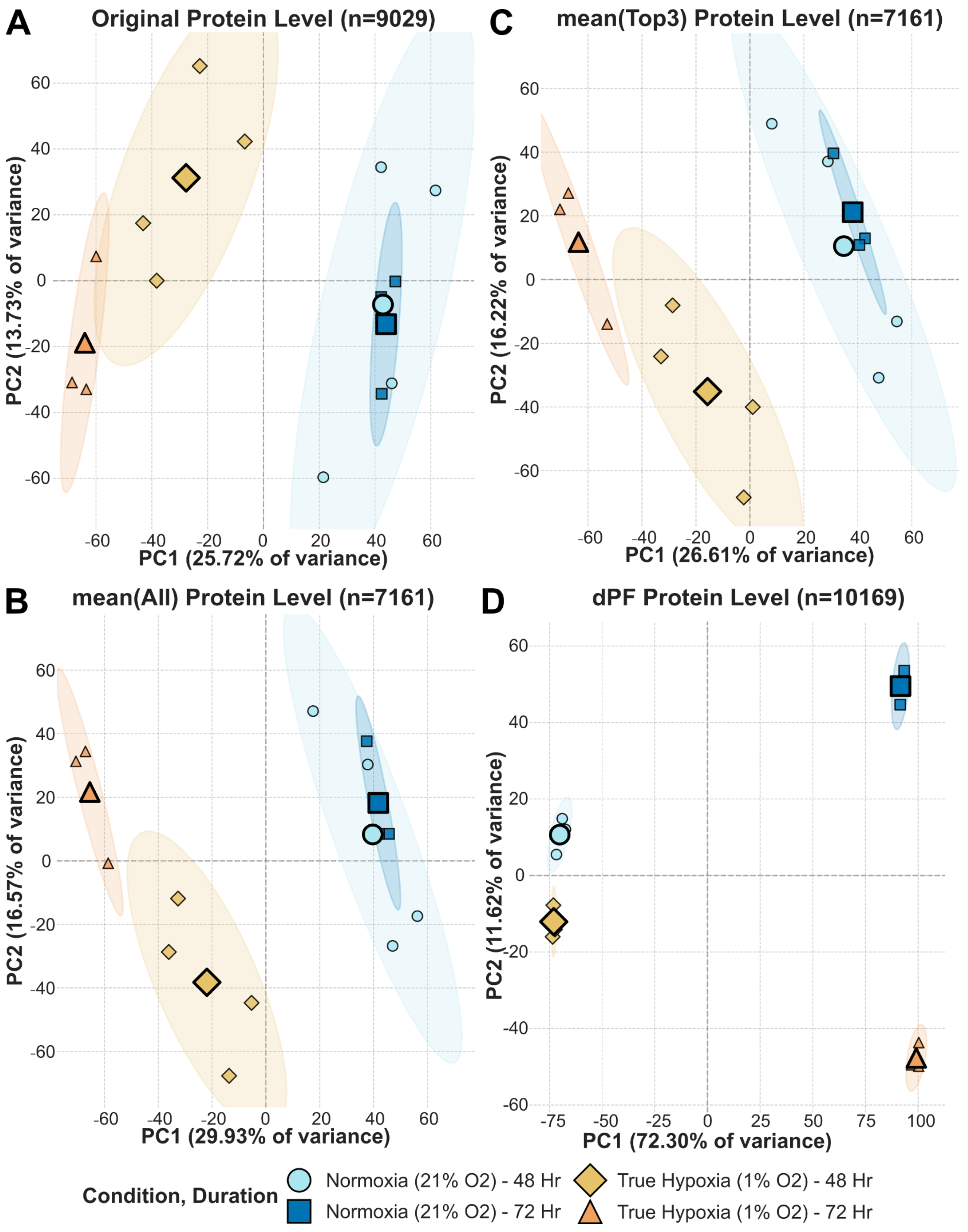


**Figure S8.** Differential proteoform (dPF) based protein quantification improves the separation of experimental groups in PCA.

PCA was used to visualize the clustering of biological replicates based on protein abundance from four experimental conditions: Normoxia (21% $O_2$) - 48 Hr (light-blue circles), Normoxia (21% $O_2$) - 72 Hr (dark blue squares), True Hypoxia (1% $O_2$) - 48 Hr (yellow diamonds), and True Hypoxia (1% $O_2$) - 72 Hr (orange triangles). Each point represents a replicate, ellipses indicate the 95% confidence interval, and larger black-edged markers represent group centroids. The percentage of variance explained by the principal components is indicated on the axes. Each panel shows a PCA based on a different protein quantification strategy, with the number of features (n) noted: **(A)** protein quantities as reported by DIANN (n=9029); **(B)** protein quantities from the mean of all peptides (n=7161); **(C)** protein quantities from the mean of the top 3 peptides (n=7161); and **(D)** protein quantities from dPF groups, which separate signals from distinct proteoforms to reduce variance (n=10169).
