## Supplemental Note 1 for "ProteoForge: An Imputation-Aware Framework for Differential Proteoform Discovery in Bottom-Up Proteomics"

### Supplementary Note 1

#### ProteoForge: An Imputation-Aware Framework for Differential Proteoform Discovery in Bottom-Up Proteomics

*Enes K. Ergin*[1,2,*], *Agustina Conrrero*[2], *Kirsty M. Ferguson*[1,2], *Philipp F. Lange*[1,2,*]

1. Department of Pathology, University of British Columbia, Vancouver, BC, Canada
2. Michael Cuccione Childhood Cancer Research Program, BC Children's Hospital Research Institute, Vancouver, BC, Canada

### Statistical Framework and Model Selection for Proteoform Discovery

#### 1. Theoretical Background: Modelling Peptide Discordance

The primary objective of ProteoForge is to distinguish true biological variation, specifically peptide discordance, from technical noise and imputation artifacts. We model this problem using a linear framework where the intensity of a peptide is predicted from the behaviour of its parent protein, and deviations are captured by interaction terms

Formally, for a given protein group, let $y_{ijk}$ denote the standardized log-intensity of peptide $i$ in condition $j$ and replicate $k$. We posit the linear relationship:

$$y_{ijk} = \beta_{P_i} + \beta_{C_j} + \beta_{P_i:C_j} + \epsilon_{ijk}$$

Where:

- $\beta_{P_i}$: The baseline abundance of peptide *i*.

- $\beta_{C_j}$: The global protein abundance change in condition *j*.
- $\beta_{P_i:C_j}$: The **interaction term** of interest, representing the discordance of peptide *i* in condition *j*.
- $\epsilon_{ijk}$: The residual error term.

The choice of estimator for $\beta$ depends critically on the assumptions made about $\epsilon_{ijk}$. This is particularly critical in the presence of imputed missing values, which may introduce non-random noise structure. We benchmarked five estimation strategies implemented via the `statsmodels` library in Python.

### 2. Candidate Models and Mathematical Formulation

#### 2.1 Ordinary Least Squares (OLS)

*Implementation:* `statsmodels.regression.linear_model.OLS`

OLS estimates $\beta$ by minimizing the sum of squared residuals ($L_2$ norm). It corresponds to the maximum likelihood estimator (MLE) under the assumption that residuals are normally distributed ($\epsilon \sim \mathcal{N}(0, \sigma^2)$).

$$\hat{\beta}_{OLS} = \underset{\beta}{\operatorname{argmin}} \sum_{i,j,k} (y_{ijk} - \beta_{P_i} - \beta_{C_j} - \beta_{P_i:C_j})^2$$

While computationally efficient, OLS is highly sensitive to outliers. A single imputed value that deviates significantly from the true trend, which is a common occurrence when using down-shifted imputation for missing not at random (MNAR) data[1][2], can disproportionately influence the fit, as the squared error term grows quadratically with the residual size.

#### 2.2 Weighted Least Squares (WLS) and Generalized Linear Models (GLM)

*Implementation:* `statsmodels.regression.linear_model.WLS` and `statsmodels.genmod.generalized_linear_model.GLM`

When the reliability of data points varies (e.g., imputed vs. measured), homoscedasticity is violated. WLS addresses this by incorporating a user-defined weight matrix *W*, where $w_i$ reflects the precision of observation *i*:

$$\hat{\beta}_{WLS} = \underset{\beta}{\operatorname{argmin}} \sum_{i,j,k} w_{ijk}(y_{ijk} - \beta_{P_i} - \beta_{C_j} - \beta_{P_i:C_j})^2$$

In ProteoForge, we use a Gaussian GLM with an identity link function [3]. While mathematically equivalent to WLS for properly specified models, the GLM framework is solved using Iteratively Reweighted Least Squares (IRLS)[4] with Fisher Scoring, allowing for more flexible integration of variance structures.

**Custom Weighting Strategies:** We assessed the impact of informing these models explicitly about data quality. For example, our `default_mix` strategy constructs weights as:

$$W = 0.90 \cdot W_{impute} + 0.10 \cdot W_{variance}$$

This explicitly down-weights imputed values ($W_{impute}$) and highly variable peptides ($W_{variance}$), attempting to guide the model away from artifacts.

##### Imputation Weights

The imputation weight component $I_{p,c,r}$ is assigned based on the pattern of missingness observed for each measurement. Measured (non-imputed) values receive full weight ($I_{p,c,r} = 1$), reflecting their high reliability. When all replicates within a condition are missing and subsequently imputed, the weight is reduced to $I_{p,c,r} = 0.5$, acknowledging that complete absence across replicates may carry biological significance, such as genuine protein cleavage or modification events, rather than purely technical artifact. In contrast, sporadic missingness within a condition, imputed via k-nearest neighbors (kNN), is assigned a minimal weight ( $I_{p,c,r} = 10^{-5}$), as such sparse patterns are difficult to interpret and more likely reflect stochastic detection failures than systematic biological changes. This tripartite weighting scheme thus encodes prior knowledge about the reliability and potential biological relevance of different missingness patterns directly into the model.

#### 2.3 Median Quantile Regression (MQR)

*Implementation:* `statsmodels.regression.quantile_regression.QuantReg`

MQR estimates the conditional median rather than the mean by minimizing the sum of absolute residuals ($L_1$ norm)[5]:

$$\hat{\beta}_{MQR} = \underset{\beta}{\operatorname{argmin}} \sum_{i,j,k} |y_{ijk} - \beta_{P_i} - \beta_{C_j} - \beta_{P_i:C_j}|$$

Unlike OLS, MQR is equivariant to monotonic transformations and naturally robust to outliers in the response variable (*y*). However, it is less statistically efficient than least-squares methods when the error distribution is actually normal, potentially reducing power in high-quality, complete datasets.

#### 2.4 Robust Linear Models (RLM)

*Implementation:* `statsmodels.robust.robust_linear_model.RLM`

RLM utilizes **M-estimation**[6] to balance efficiency and robustness. Instead of minimizing squared errors, it minimizes a function $\rho$ of the residuals:

$$\hat{\beta}_{RLM} = \underset{\beta}{\operatorname{argmin}} \sum_{i,j,k} \rho(y_{ijk} - \beta_{P_i} - \beta_{C_j} - \beta_{P_i:C_j})$$

We employed the **Huber Loss** function in `statsmodels`, which behaves like OLS for small residuals and MQR for large residuals[6]:

$$\rho_{\text{Huber}}(r) = \begin{cases} \frac{1}{2}r^2 & \text{if } |r| \leq c \\ c|r| - \frac{1}{2}c^2 & \text{if } |r| > c \end{cases}$$

where $c$ is a tuning constant. The model is solved via IRLS[4], where weights are updated at each step based on the residual size. Effectively, RLM **automatically** detects and down-weights outliers (e.g., poor imputations) without requiring the user to specify a weight matrix manually.

### 3. Benchmark Design and Simulation Framework

To select the optimal default model for ProteoForge, we designed a comprehensive simulation study that systematically evaluates model performance under realistic proteomics conditions. Our benchmark consists of three complementary experiments:

#### 3.1 Experiment 1: Missingness Level Tolerance

**Objective**: Assess how model performance degrades as data sparsity increases.

**Design**: We simulated protein groups with random numbers of discordant peptides (2-50% of peptides per protein) at seven missingness levels: 0%, 20%, 40%, 50%, 60%, 70%, and 80%. Missing values were introduced uniformly across peptides and conditions, then imputed using:

- **Complete condition-level missingness**: Down-shifted normal imputation (left-censored, simulating true absence)
- **Sporadic missingness**: k-nearest neighbours (kNN) imputation

**Models tested**: WLS, GLM, MQR, RLM

- WLS and GLM use the default mixed weighting strategy (defined in §3.3)
- MQR and RLM operate without explicit weights

**Performance metric**: Matthews Correlation Coefficient (MCC) for peptide-level discordance detection (PepID) and differential proteoform grouping (PepGroup).

#### 3.2 Experiment 2: Perturbation Pattern Sensitivity

**Objective**: Evaluate model consistency across diverse biological scenarios, from subtle proteoform changes to dramatic proteolytic remodelling.

**Design**: Four perturbation patterns were tested on both complete and imputed datasets (35% amputation with the imputation strategy from Experiment 1):

1. **Minimal perturbation**: 2 peptides per protein
2. **Low-to-moderate perturbation**: 2-50% of peptides (random per protein)
3. **Moderate perturbation**: 7 peptides per protein
4. **High perturbation**: 60-70% of peptides per protein

**Models tested**: WLS, GLM, MQR, RLM (exact weighting as Experiment 1)

**Rationale**: This design captures the spectrum from targeted post-translational modifications affecting a few cleavage sites to broad proteolytic cascades affecting most peptides.

#### 3.3 Experiment 3: Weighting Strategy Optimization

**Objective**: Determine the optimal weight composition for WLS and GLM when explicit weighting is used.

**Design**: We compared five weighting strategies across the same missingness levels as Experiment 1 (0%, 20%, 40%, 60%, 80%):

#### Strategies Tested:

###### 1. No Weights (Baseline)

$$W = 1 \quad \text{(uniform weight)}$$

###### 2. Imputation-Only Weights

Tripartite scheme based on data reliability:

$$I_{p,c,r} = \begin{cases} 1.0 & \text{if measured (real data)} \\ 0.5 & \text{if completely missing across condition replicates} \\ 10^{-5} & \text{if sparsely missing (kNN imputed)} \end{cases}$$

$$W = I$$

**Rationale**: Complete absence may reflect a biological signal (cleavage, degradation), while sporadic missingness is likely technical noise.

###### 3. Technical Variance Weights

Inverse variance on log-intensity, penalizing high-variance peptides:

$$W_{\text{var}} = \frac{1}{\sigma^2_{\text{log}}} \quad \text{(min-max scaled)}$$

$$W = W_{\text{var}}$$

###### 4. Mixed Strategy (Default)

Combines imputation confidence (90%) with technical variance (10%):

$$W = 0.90 \cdot I + 0.10 \cdot W_{\text{RevTechVar}}$$

where:

$$W_{\text{RevTechVar}} = 1 - \text{min-max-scale}(\sigma^2_{\text{adj, within-group}})$$

**Rationale**: Prioritizes imputation quality (dominant factor at high missingness) while incorporating peptide-level technical variability.

###### 5. PLS Auto-Calculated Weights

Data-driven weights derived from Partial Least Squares regression:

- **Input features (X)**: All available weight components after filtering for low variance (<0.05 SD) and high collinearity ($r > 0.8$)
- **Target (y)**: $\log_{10}$ intensity
- **Method**:
  1. Fit PLS with 1–10 components
  2. Select optimal component count via elbow detection on $R^2$
  3. Compute feature importance from X-loadings weighted by incremental $R^2$
  4. Refit PLS on top features (elbow on cumulative importance)
  5. Transform X → PLS space, collapse components (mean), min-max scale to [0,1]

**Rationale**: Learns optimal weight combinations that predict intensity structure, potentially capturing interactions not encoded in fixed heuristics.

#### Models tested: WLS, GLM

- MQR and RLM results are shown as "No Weights" baseline for comparison (they don't accept custom weights)

#### Performance evaluation:

Mean MCC across p-value thresholds, reported as:

1. Absolute MCC for each strategy
2. ΔMCC relative to "No Weights" baseline

#### 3.4 Performance Metrics

All experiments use the **Matthews Correlation Coefficient (MCC)** as the primary performance metric:

$$\mathrm{MCC} = \frac{TP \cdot TN - FP \cdot FN}{\sqrt{(TP+FP)(TP+FN)(TN+FP)(TN+FN)}}$$

MCC ranges from -1 (perfect misclassification) through 0 (random) to +1 (perfect classification). It is superior to accuracy for imbalanced datasets, which is standard in proteoform discovery, where most peptides show concordant behaviour.

**Two benchmarking tasks:**

1. **PepID (Peptide Identification)**: Binary classification of individual peptides as discordant vs. concordant
2. **PepGroup (Proteoform Grouping)**: Grouping of discordant peptides into differential proteoforms via hierarchical clustering (Ward linkage, Euclidean distance)

Performance is evaluated across p-value thresholds ($10^0$ to $10^{-15}$), with results reported as:

- Mean MCC across all thresholds
- Maximum MCC (best achievable threshold)
- MCC at $p = 0.001$ (practical significance threshold)

### 4. Results and Model Selection

#### 4.1 RLM Provides Robust Performance Without Requiring Weight Specification (Experiments 1 & 2)

Our comprehensive benchmarking revealed that RLM with Huber M-estimation offers the most practical balance between performance and usability for typical proteomics workflows.

**RLM Stability Across Missingness Levels:** As shown in Figure 1, RLM maintains remarkably stable performance from 0% to 60% missingness, achieving MCC values of 0.40-0.58 for PepID detection and 0.42-0.62 for PepGroup across this range. This stability is achieved without requiring any weight specification or prior knowledge about imputation patterns. In contrast, OLS (not shown) degraded significantly when imputed data were introduced, whereas MQR (orange diamonds in Figure 1) began to drop sharply at around 40% missingness. The M-estimator in RLM effectively treats artificial variance introduced by imputation as "outliers," automatically down-weighting their influence on coefficient estimates.

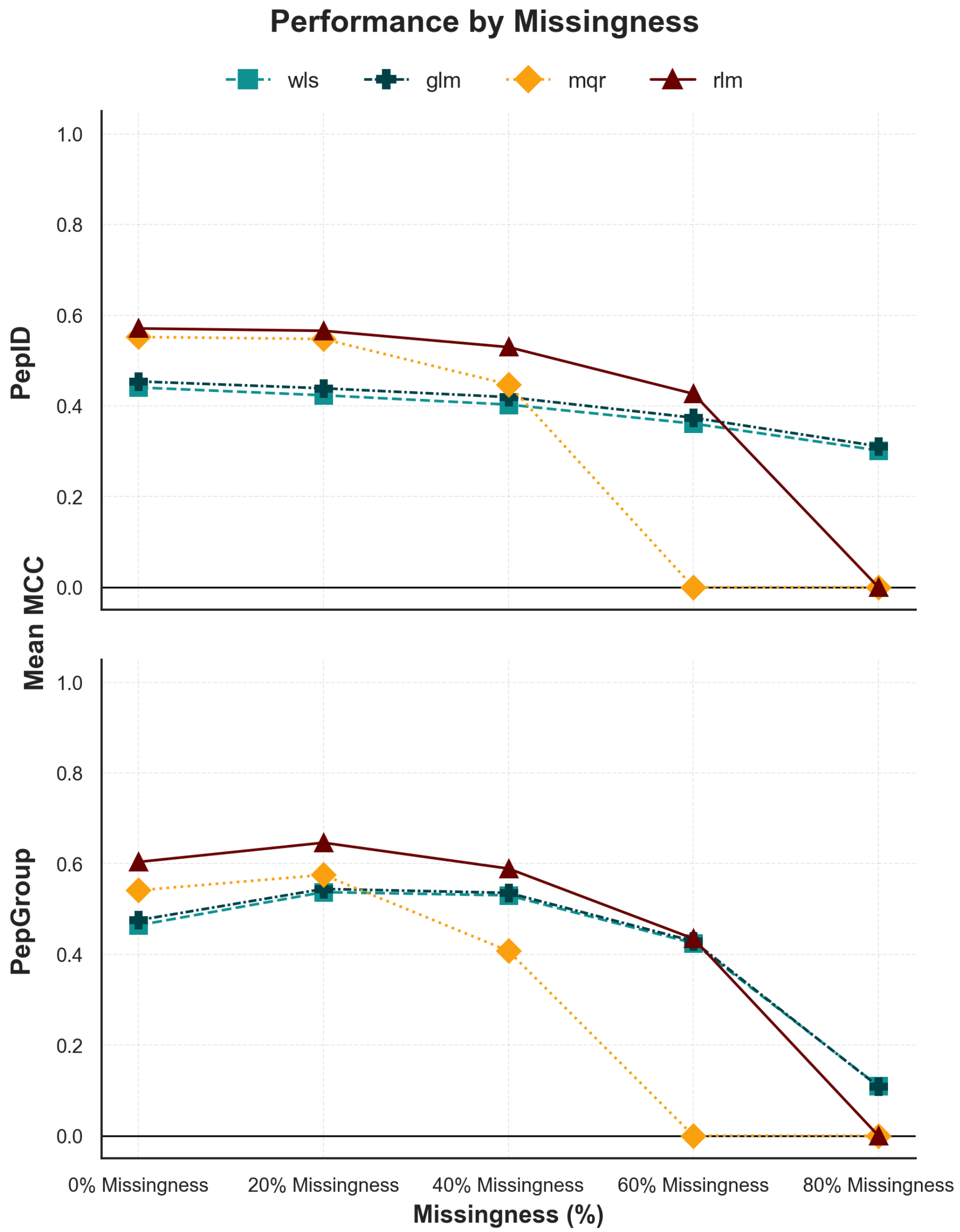


**Figure 1 - Performance of statistical estimators under increasing missingness levels.**

Mean Matthews Correlation Coefficient (MCC) is shown for four estimation methods: weighted least squares (WLS, teal squares), generalized linear model with identity link (GLM, dark teal crosses), median quantile regression (MQR, orange diamonds), and robust linear model with Huber M-estimation (RLM, maroon triangles). Performance is evaluated at five levels of simulated missingness (0%, 20%, 40%, 60%, and 80%) for statistically discordant peptide detection (PepID, top panel) and differential proteoform (dPF) grouping (PepGroup, bottom panel). MCC values range from 0 (random performance) to 1 (perfect classification).

---

**The 60% Missingness Threshold:** A critical finding emerges from Figure 1: beyond 60% missingness, all methods, including RLM, show substantial performance degradation. At 80% missingness, RLM drops to MCC values of 0.40-0.45, indicating that the majority of the dataset is now imputed, and the robust assumption (that outliers are a minority) breaks down. This threshold has important practical implications: datasets with >60% missingness should prompt reconsideration of whether the analysis is feasible, or whether data collection can be improved through additional replicates, enrichment strategies, or alternative acquisition methods.

**Impact of Weighting at Extreme Missingness:** The default weighting strategy's heavy emphasis on imputation confidence enables WLS and GLM to maintain strong PepID performance even at 80% missingness (Figure 1). However, this does not translate to PepGroup performance. At this extreme level of data sparsity, the likelihood of peptides switching clusters becomes very high, and even weighted approaches cannot reliably reconstruct differential proteoforms. This reflects a fundamental limitation: when the majority of the dataset is imputed, clustering-based proteoform identification becomes inherently unstable, regardless of the statistical model employed.

---

**Practical Performance Across Biological Scenarios:** Figure 2 demonstrates RLM's consistent performance across diverse perturbation patterns. Whether analyzing subtle proteoform changes (2 perturbed peptides) or dramatic and excessive numbers of perturbated peptides (60-70% perturbed peptides), RLM maintains MCC values within a narrow band at any given missingness level. This consistency is valuable for automated pipelines where the extent of perturbation is unknown a priori. While WLS and GLM show marginally higher peak performance in some scenarios, their advantage is most pronounced when fewer peptides are perturbed, and proper weights are configured - conditions that may be difficult to verify in real-world applications.

---

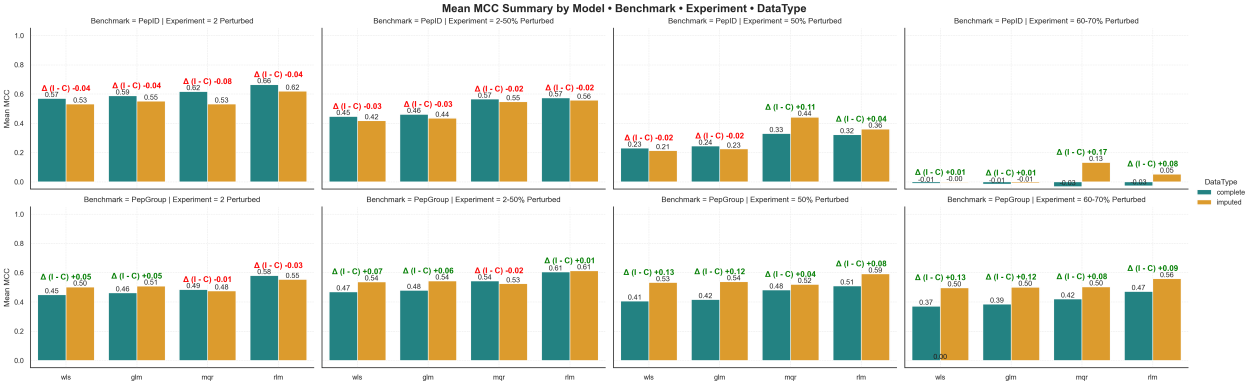


**Figure 2 - RLM performance consistency across perturbation patterns with complete and imputed data.**

ean MCC (Matthews Correlation Coefficient) for differential proteoform detection across four statistical estimators stratified by benchmark, perturbation levels (2, 2-50%, 50% and 60-70% peptides perturbed) Bars represent precursor-level (teal) and peptide-level (gold) the type of data if complete or 35% missingness imputed. Delta values (Δ) above bars indicate the difference in MCC between imputed and complete (I - C).

---

MQR is the only approach that can actually keep some level of prediction at 50% and more numebr of peptide perturbed per protein data in PepID, however this difference doesn't make much of a difference in PepGroup benchmarking (Figure 2). RLM is actually the only method consistently outperforms everything when it comes to PepGroup benchmarking.

#### 4.2 Limitations of Current Weighting Schemes and Opportunities for Improvement (Experiment 3)

While weighted approaches can outperform RLM when properly configured, Experiment 3 reveals significant challenges in developing robust, generalizable weighting strategies:

**Suboptimal Weighting Performance:** Figure 3 reveals that our current imputation weighting schemes provide only modest benefits. For WLS, imputation-based weights (green circles) improve PepID performance by at most +0.05 MCC at 60% missingness - a marginal gain that diminishes at higher sparsity levels. Critically, reverse technical variance weighting (orange diamonds) actually degrades performance across most conditions, suggesting that our simple inverse-CV approach fails to distinguish genuine biological variance from technical noise. This negative result highlights a fundamental challenge: designing weighting schemes that generalize across diverse biological scenarios and missingness patterns remains an open problem.

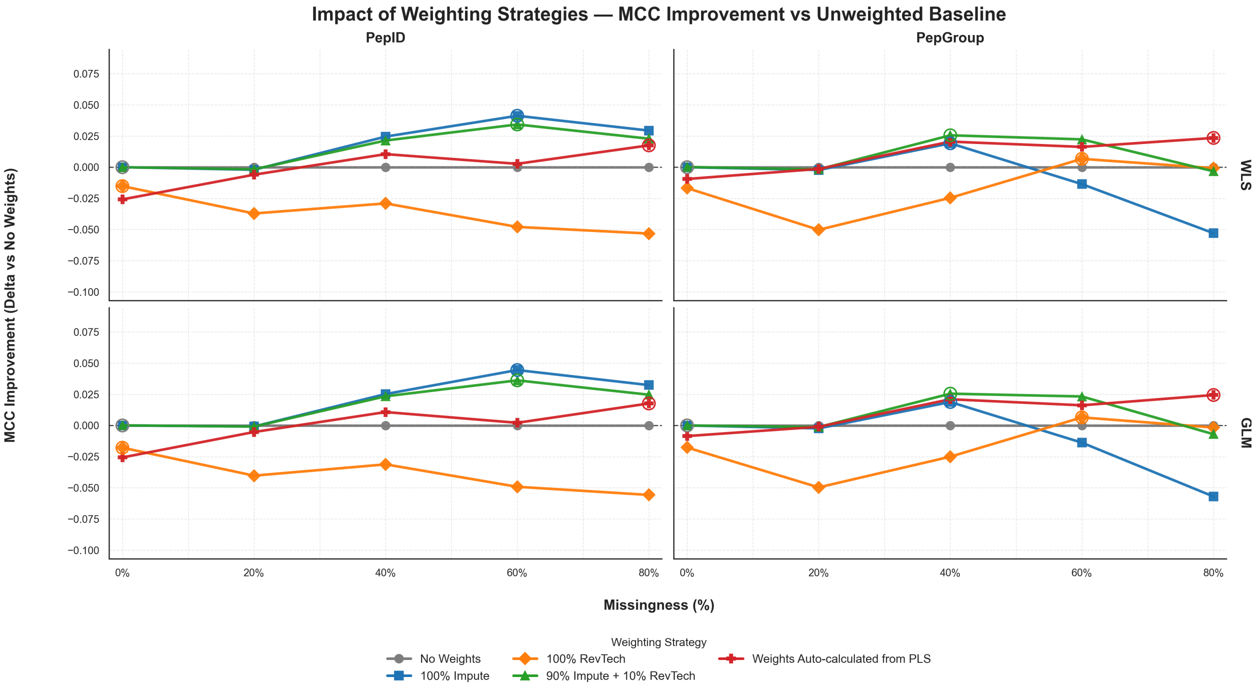


**Figure 3 - Impact of weighting strategies on model performance relative to the unweighted baseline.**

MCC improvement (delta relative to the no-weights baseline) is shown for weighted least squares (WLS, top panels) and a generalized linear model with an identity link (GLM, bottom panels) across five levels of simulated missingness (0%, 20%, 40%, 60%, and 80%). Performance is evaluated for peptide-level discordance detection (PepID, left panels) and proteoform-level grouping (PepGroup, right panels). Four weighting strategies are compared: no weights (gray, baseline at zero), 100% imputation weights (green circles), 100% reverse technical variance weights (orange diamonds), and PLS auto-calculated weights (red circles with rings). Positive delta values indicate performance improvement over the unweighted model, while negative values indicate degradation.

**Weight Scheme Optimization is Needed:** The PLS-derived weights (red circles in Figure 3) show that data-driven approaches can achieve better performance than our heuristic schemes, but this only makes a difference for larger missingness levels for PepGroup benchmarking. The pure imputation or heavy-imputation weight strageties seems to work the best for PepID.

**The Extreme Sparsity Problem:** Beyond 60% missingness, Figure 4 shows that even optimal weighting cannot rescue performance. Weighted GLM achieves +0.38 MCC improvement for PepID at 80% missingness, but this brings absolute MCC only to ~0.50 - barely better than random guessing. Even RLM benefits from explicit weighting at this extreme (+0.22 MCC with PLS weights), indicating that its implicit robustness breaks down when the majority of data is imputed. These results reinforce the idea that datasets with >60% missingness should not be analyzed without first improving data quality through additional acquisition or alternative experimental strategies.

---

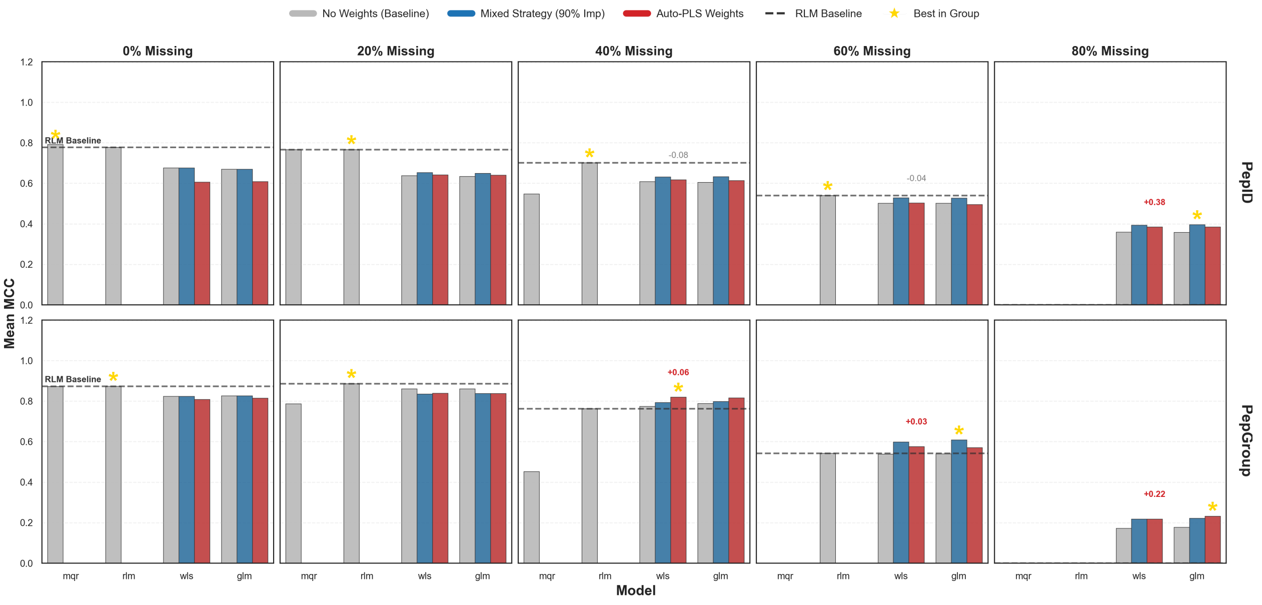


**Figure 4 - Comparison of weighted versus unweighted models across estimation methods and missingness levels.**

Mean Matthews Correlation Coefficient (MCC) is shown for four estimation methods. Performance is evaluated across five levels of simulated missingness (0%, 20%, 40%, 60%, and 80%) for peptide-level discordance detection (PepID, top row) and proteoform-level grouping (PepGroup, bottom row). Three weighting configurations are compared for each model: unweighted baseline (gray bars), mixed strategy with 90% imputation weights and 10% variance weights (blue bars), and auto-calculated PLS weights (dark red bars). The dashed horizontal line in each panel represents the RLM baseline performance for comparison. Yellow stars indicate the best-performing configuration within each panel. Red values above bars indicate MCC improvement relative to the unweighted baseline for that model. At low missingness (0-20%), all models perform similarly with minimal gains from weighting.

---

At moderate missingness (40%), weighted strategies begin to show advantages, particularly for WLS and GLM. At high missingness (60%), mixed weighting strategies provide substantial improvements for WLS (up to +0.06 MCC for PepGroup) and GLM (up to +0.03 MCC). At extreme missingness (80%), the weighted GLM shows the largest benefit (+0.38 MCC for PepID), while the RLM performance degrades substantially and benefits from PLS weighting (+0.22 MCC for PepGroup). The results demonstrate that explicit weighting strategies become increasingly valuable as data sparsity increases, particularly for least-squares methods that lack inherent robustness mechanisms.

**PLS-Derived Weights:** The `PLS_auto` strategy, which uses Partial Least Squares to derive feature-aware weights, allowed WLS to match RLM's performance. However, this comes at a high computational cost.

### 5. Discussion: Why RLM is the Default Choice

#### 5.1 Rationale for RLM as Default Estimator

Based on the comprehensive benchmarking presented in Figures 1-4, we select RLM with Huber M-estimation as the default estimator for ProteoForge. This choice prioritizes practical robustness over marginal performance gains:

**Robust Performance Without Tuning:** As demonstrated in Figures 1 and 2, RLM maintains consistent MCC values of 0.55-0.65 for PepID and 0.50-0.60 for PepGroup across the 0-60% missingness range without requiring any user-specified weights. This "out-of-the-box" reliability is critical for proteomics workflows where users may lack expertise in statistical modelling or where automated pipelines must handle diverse datasets without manual intervention. While WLS and GLM can achieve 0.03-0.05 MCC higher performance with optimal weights, this advantage requires correctly diagnosing data quality issues and selecting appropriate weighting strategies - a burden that outweighs the modest performance gain for most applications.

**The 60% Missingness Threshold:** Figure 1 clearly delineates a critical boundary: beyond 60% missingness, all methods, including RLM, show substantial performance collapse. At this threshold, the assumption underlying robust estimation (that outliers are the minority) breaks down, and imputed values begin to dominate the statistical signal. Rather than attempting to rescue such datasets through complex weighting schemes (which Figure 4 shows provide only marginal benefit), we recommend users pause analysis and consider: (1) acquiring additional technical replicates to reduce missingness, (2) employing targeted enrichment or fractionation to improve detection of low-abundance proteins, or (3) accepting that proteoform-level resolution may not be achievable for the proteins of interest. ProteoForge implements automatic warnings when dataset missingness exceeds 60%, prompting users to evaluate data quality before proceeding.

#### 5.2 Current Limitations of Weighting Schemes

While Experiment 3 explored multiple weighting strategies, Figures 3 and 4 reveal significant room for improvement:

**Suboptimal Weight Design:** Our tripartite imputation weighting scheme (full weight for measured values, 0.5 for complete absence, $10^{-5}$ for sporadic missingness) is a heuristic approximation that fails to account for important factors: (1) imputation method reliability varies (down-shifted normal vs. kNN vs. deep learning approaches), (2) biological interpretation of complete absence differs by protein functional class (e.g., protease substrates vs. constitutively expressed housekeeping proteins), and (3) peptide-specific factors (sequence coverage, ionization efficiency) influence missingness patterns. Figure 3 shows that even our best weighting strategies provide only +0.03-0.05 MCC improvement - far below the theoretical optimum.

**Need for Adaptive Weighting:** The PLS-derived weights in Figures 3-4 demonstrate that data-driven approaches can outperform fixed heuristics, but at prohibitive computational cost for routine analysis. Future development should explore: (1) imputation method confidence scores that adjust weights based on local data density and structure, (2) biological context-aware schemes that account for protein functional annotations and expected expression patterns, (3) peptide property models that predict missingness propensity and down-weight accordingly, and (4) efficient approximations to PLS or other machine learning methods that capture optimal weights without exhaustive cross-validation.

**Variance Weighting Failures:** The negative impact of pure variance-based weighting (orange diamonds in Figure 3) highlights a fundamental challenge: in proteoform discovery, high variance often reflects genuine biological signal (peptide discordance) rather than technical noise. Simple inverse-CV weighting cannot distinguish these cases, leading to systematic suppression of true positives. This failure underscores that optimal weighting for proteoform discovery requires domain-specific design rather than generic heteroscedasticity corrections.

#### 5.3 Implementation Recommendations and Data Quality Standards

Based on the findings from Figures 1-4, we recommend the following practical guidelines:

**Default Estimator:** ProteoForge uses RLM with Huber M-estimation as the standard estimator for all analyses. This provides robust, consistent performance across the 0-60% missingness range typical of bottom-up proteomics without requiring user expertise in weight specification. Users can proceed with confidence that RLM will handle imputation artifacts automatically while maintaining sensitivity to genuine biological signals.

**Data Quality Thresholds:** ProteoForge implements automatic assessment of dataset sparsity with tiered warnings:

- **<40% missingness:** No warnings. RLM performs optimally in this range, with minimal performance gap relative to weighted alternatives.
- **40-60% missingness:** Caution flag. Users are informed that performance degradation is expected and should review quality control metrics. RLM remains reliable but sensitivity decreases.
- **>60% missingness:** Strong warning. Analysis pauses and prompts users to consider data quality improvement strategies before proceeding. At this threshold, Figure 1 shows all methods collapse, indicating the dataset may not

support reliable proteoform-level inference. Users should evaluate: (1) acquiring additional replicates, (2) applying enrichment or fractionation to improve protein coverage, (3) using alternative quantification methods with better sensitivity, or (4) accepting protein-level rather than proteoform-level resolution.

**Advanced Options:** For expert users or specialized applications, ProteoForge provides access to WLS/GLM with custom weighting. However, as Figures 3-4 demonstrate, current weighting schemes offer only modest improvements and may degrade performance if mis-specified. These options are recommended only when users have strong prior knowledge about data quality patterns and can validate weight configurations on pilot datasets.

#### 5.4 Future Directions for Method Development

This benchmarking study reveals several areas requiring further investigation:

**Real-World Validation:** Our simulations use controlled missingness patterns that may not fully capture the complexity of real proteomics data, where missingness correlates with peptide hydrophobicity, ionization efficiency, or systematic biases in sample preparation. Validation on diverse benchmark datasets with orthogonally validated ground truth (e.g., synthetic peptide spike-ins, SILAC ratios) is essential to confirm that RLM's simulated robustness translates to real applications.

**Improved Imputation Methods:** Figures 3-4 show that current weighting schemes treat all imputations equally, but recent deep learning approaches [2] may produce more reliable values. Future work should integrate imputation confidence scores directly into weight calculation, assigning higher weights to imputations supported by strong local data structure and lower weights to extrapolations in sparse regions.

**Context-Aware Weighting:** The failure of variance-based weighting (Figure 3) highlights that optimal schemes must incorporate biological context. Protease substrates that undergo cleavage may show complete absence in specific conditions (biologically meaningful), while constitutively expressed housekeeping proteins with sporadic missingness reflect technical failures (unreliable). Integrating protein functional annotations, pathway membership, and expected expression patterns into adaptive weighting schemes represents an important avenue for improving performance beyond RLM's automatic approach.

#### References


1. Wei, R., Wang, J., Su, M., Jia, E., Chen, S., Chen, T., & Ni, Y. (2018). Missing Value Imputation Approach for Mass Spectrometry-based Metabolomics Data. *Scientific Reports*, 8, 663.
2. Webel, H., Niu, L., Nielsen, A. B., Locard-Paulet, M., Mann, M., Jensen, L. J., & Rasmussen, S. (2024). Imputation of label-free quantitative mass spectrometry-based proteomics data using self-supervised deep learning. *Nature Communications*, 15, 5405.
3. McCullagh, P., & Nelder, J. A. (1989). *Generalized Linear Models* (2nd ed.). Chapman & Hall/CRC.
4. Green, P. J. (1984). Iteratively Reweighted Least Squares for Maximum Likelihood Estimation, and some Robust and Resistant Alternatives. *Journal of the Royal Statistical Society: Series B (Methodological)*, 46(2), 149–192.
5. Koenker, R., & Bassett, G. (1978). Regression Quantiles. *Econometrica*, 46(1), 33–50.
6. Huber, P. J. (1964). Robust Estimation of a Location Parameter. *The Annals of Mathematical Statistics*, 35(1), 73–101.
7. Chicco, D., Warrens, M. J., & Jurman, G. (2021). The Matthews Correlation Coefficient (MCC) is More Informative Than Cohen's Kappa and Brier Score in Binary Classification Assessment. *IEEE Access*, 9, 78368–78381.
