## Supplementary material for "ProteoForge: An Imputation-Aware Framework for Differential Proteoform Discovery in Bottom-Up Proteomics": Analysis notebooks as html render: Notebook S1.html

### Notebook S1

###### Enes Kemal Ergin

#### 2025-12-08 23:00:16

### Supplementary Notebook 1: Discordant peptide identification benchmark using SWATH-MS Interlab Data¶

- **License:** Creative Commons Attribution-NonCommercial 4.0 International License
- **Version:** 0.2
- **Edit Log:**
  - 2025-11-28: Initial version of the notebook
  - 2025-12-08: Revise the whole notebook, ensuring clarity and correctness

---

**Requirements:**

The following preprocessing and method runs must be completed (in order) before executing this notebook:

1. `01-DataProcessing.R` — Prepares SWATH‑MS Interlab data and writes processed datasets to `data/prepared/` as `.feather` files
2. `02-runCOPF.R` — Runs COPF; saves results to `data/results/` as `COPF_{dataID}_result.feather`
3. `03-runPeCorA.R` — Runs PeCorA; saves results to `data/results/` as `PeCorA_{dataID}_result.feather`
4. `04-runProteoForge.py` — Runs ProteoForge; saves results to `data/results/` as `ProteoForge_{dataID}_result.feather`

**Expected files:** `data/results/COPF_*`, `data/results/PeCorA_*`, `data/results/ProteoForge_*` (feather format)

> **Note:** Feather (`.feather`) is a fast, cross-language binary table format readable from both R and Python—ideal for exchanging processed data, but not human-readable.

---

**Data Information:**

This analysis uses four perturbation datasets created by `01-DataProcessing.R`. Each dataset corresponds to a perturbation scenario with a short identifier (used in filenames) and a display name (used in figures):

| Identifier | Display Name | Description |
| --- | --- | --- |
| `1pep` | 1 Peptide | Perturb exactly 1 peptide per selected protein |
| `2pep` | 2 Peptides | Perturb exactly 2 peptides per selected protein |
| `050pep` | %50 Peptides | Perturb ~50% of peptides per selected protein |
| `random` | Random (1 to %50) Peptides | Perturb 1–50% of peptides per protein (randomly) |

> `050pep` is tricky since proteins with 5 peptides will default to perturbing 3 peptides (60%), while those with 4 peptides will perturb 2 (50%). The actual percentage varies by protein, but the original COPF publication used the ceiling to round up, which we also follow here.

**How perturbations are generated:**

1. `01-DataProcessing.R` reads the SWATH‑MS interlab table, filters proteins/peptides, aggregates charge states, median-normalizes intensities, and selects proteins with ≥4 peptides and sufficient replicates
2. A set of proteins to perturb is sampled, a per-protein reduction factor (`red_fac`) is drawn uniformly, and `generatePerturbedProfiles()` marks peptides as `perturbed_peptide` and reduces their intensities in `day5`
3. Prepared feather files are written to `./data/prepared/` and used by method scripts (`02`–`04`) to produce result files

---

**Purpose:**

This notebook compares three methods (**COPF**, **PeCorA**, **ProteoForge**) across four simulated perturbation scenarios and generates publication-quality performance figures (ROC, PR, F1, MCC). Note that COPF does not inherently support discordant peptide detection—it outputs peptide groups, which are back-calculated for this benchmark.

---

#### Setup¶

This section imports required libraries, configures display settings, and defines paths for data and figures.

> **Note:** The HTML rendering of this notebook hides code cells by default. Click the "Code" buttons on the right to expand them.

##### Libraries¶

```
import os
import sys
import warnings
warnings.filterwarnings("ignore")

import numpy as np # Numerical computing
import pandas as pd # Data manipulatio

import seaborn as sns # R-like high-level plots
import matplotlib.pyplot as plt # Python's base plotting 

sys.path.append('../')

# Utility imports for this analysis
from src import utils, plots

# Initialize the timer
startTime = utils.getTime()
```

##### Display Settings¶

The cell below configures pandas, matplotlib, and seaborn display options for improved readability of tables and figures, including color palettes and figure export settings.

```
## Figure Settings

# Define default colors and styles for plots
def_colors = [
    "#139593", "#fca311", "#e54f2a",
    "#c3c3c3", "#555555",
    "#690000", "#5f4a00", "#004549"
]

# Set seaborn style
sns.set_theme(
    style="white",
    context="paper",
    palette=def_colors,
    font_scale=1,
    rc={
        "figure.figsize": (6, 4),
        "font.family": "sans-serif",
        "font.sans-serif": ["Arial", "Ubuntu Mono"],
    }
)

# Figure Saving Settings
figure_formats = ["pdf"]
save_to_folder = True
transparent_bg = True
figure_dpi = 300

## Configure dataframe displaying
pd.set_option('display.float_format', lambda x: '%.4f' % x)
pd.set_option('display.max_rows', 50)
pd.set_option('display.max_columns', 50)
pd.set_option('display.width', 500)  # Set a wider display width

## Printing Settings
verbose = True

## Show the color-palettes
plots.color_palette( def_colors, save=False )
```

##### Data and Result Paths¶

Data and figures are organized in separate folders:

- `data_path` — Input directory containing method results (COPF, PeCorA, ProteoForge)
- `output_path` — Output directory for aggregated performance data
- `figure_path` — Directory for generated plots and figures

```
notebook_name = "01-IdentificationBenchmark"
data_path = "./data/results/"
output_path = f"./data/results/"
figure_path = f"./figures/{notebook_name}/"

# Create the output folder if it does not exist
if not os.path.exists(output_path):
    os.makedirs(output_path)

# Create figure folder structure, if needed
if save_to_folder:
    for i in figure_formats:
        cur_folder = figure_path + i + "/"
        if not os.path.exists(cur_folder):
            os.makedirs(cur_folder)
```

##### Analysis Parameters¶

Key parameters for the evaluation framework:

- `seed = 42` — Random seed for reproducibility
- `pthr = 1e-3` — P-value threshold highlighted on ROC and PR curves (conventional cutoff)
- `thresholds` — Range of p-value thresholds from `1` to `1e-15` for calculating performance metrics

```
# Global variables
seed = 42 # Seed for reproducibility  
pthr = 10**-3  # p-value threshold for significance
thresholds = list(utils.generate_thresholds(10.0, -15, 1, 0, 1, 0.1)) # Thresholds for the analysis

method_palette = {
    "COPF": "#139593",
    "PeCorA": "#fca311",
    "ProteoForge": "#e54f2a",
}
method_styles = {
    "COPF": "--",
    "PeCorA": "-.",
    "ProteoForge": ":",
}
method_markers = {
    "COPF": "o",
    "PeCorA": "s",
    "ProteoForge": "^",
}

data_order = [
    "1 Peptide", 
    "2 Peptides", 
    "Random (1 to %50) Peptides", 
    "%50 Peptides"
]
```

#### Performance Analysis¶

##### Data Aggregation and Preprocessing¶

The first step aggregates results from all methods and scenarios. The loop below:

1. Reads each method's `.feather` result files for all four perturbation scenarios
2. Calculates performance metrics (TP, FP, TN, FN) across p-value thresholds
3. Combines everything into a single DataFrame (`results_data`)

This unified dataset serves as the foundation for all subsequent performance visualizations.

```
results_data = pd.DataFrame()
# Read and combine the outputs:
for dataID in ["1pep", "2pep", "050pep", "random"]:
    for curMethod in ["COPF", "PeCorA", "ProteoForge"]:
        # Determine the p-value column
        if curMethod == "COPF": 
            pval_col = "proteoform_score_pval"
        else:  
            pval_col = "adj_pval"
        # Read the data
        data = pd.read_feather(f"{data_path}{curMethod}_{dataID}_result.feather")
        if curMethod == "ProteoForge":
            data = data[['protein_id', 'peptide_id', 'perturbed_peptide', pval_col]].drop_duplicates()
            
        # Create the metrics data across the thresholds
        metric_data = utils.create_metric_data(
            data,
            thresholds,
            label_col="perturbed_peptide",
            pvalue_col=pval_col,
        )
        metric_data["method"] = curMethod
        metric_data["perturbation"] = dataID
    
        results_data = pd.concat([results_data, metric_data], axis=0, ignore_index=True)


print(f"Data shape: {results_data.shape}")
results_data["perturbation"] = results_data["perturbation"].replace({
    "1pep": "1 Peptide",
    "2pep": "2 Peptides",
    "050pep": "%50 Peptides",
    "random": "Random (1 to %50) Peptides",
})
# Save the processed data
output_file = f"{output_path}peptide_identification_performance_data.feather"
results_data.to_feather(output_file)
results_data.to_csv(output_file.replace('.feather', '.csv'), index=False)

results_data.tail()
```

```
Data shape: (300, 14)
```

|  | TP | FP | TN | FN | TPR | FPR | FDR | MCC | Precision | Recall | F1 | threshold | method | perturbation |
| --- | --- | --- | --- | --- | --- | --- | --- | --- | --- | --- | --- | --- | --- | --- |
| 295 | 3101 | 8497 | 10120 | 529 | 0.8543 | 0.4564 | 0.7326 | 0.2943 | 0.2674 | 0.8543 | 0.4073 | 0.6000 | ProteoForge | Random (1 to %50) Peptides |
| 296 | 3125 | 8823 | 9794 | 505 | 0.8609 | 0.4739 | 0.7384 | 0.2868 | 0.2616 | 0.8609 | 0.4012 | 0.7000 | ProteoForge | Random (1 to %50) Peptides |
| 297 | 3142 | 9096 | 9521 | 488 | 0.8656 | 0.4886 | 0.7433 | 0.2800 | 0.2567 | 0.8656 | 0.3960 | 0.8000 | ProteoForge | Random (1 to %50) Peptides |
| 298 | 3162 | 9345 | 9272 | 468 | 0.8711 | 0.5020 | 0.7472 | 0.2749 | 0.2528 | 0.8711 | 0.3919 | 0.9000 | ProteoForge | Random (1 to %50) Peptides |
| 299 | 3630 | 18617 | 0 | 0 | 1.0000 | 1.0000 | 0.8368 | 0.0000 | 0.1632 | 1.0000 | 0.2806 | 1.0000 | ProteoForge | Random (1 to %50) Peptides |

The table above shows the compiled data structure. Each row captures performance for a specific method and perturbation scenario at a given p-value threshold.

##### ROC Analysis¶

The **Receiver Operating Characteristic (ROC) curve** evaluates how well each method distinguishes between perturbed and unperturbed peptides by plotting True Positive Rate (TPR/sensitivity) against False Positive Rate (FPR/1-specificity).

**Interpretation:**

- A perfect classifier hugs the top-left corner
- **Area Under the Curve (AUC):** `1.0` = perfect, `0.5` = random chance
- Black circles indicate performance at `p = 1e-3`

```
# ROC Curves for different perturbations and methods
fig, axes = plt.subplots(1, 4, figsize=(16, 4), sharey=True, gridspec_kw={"wspace": 0.05})

for i, pert in enumerate(data_order):
    cur_data = results_data[results_data["perturbation"] == pert]
    # Ensure the data is complete for ROC curves
    cur_data = cur_data.groupby("method").apply(
        lambda x: utils.complete_curve_data(x, 'ROC', 'FPR', 'TPR')
    ).reset_index(drop=True)
    
    # Calculate AUC per method from TPR and FPR
    auc_data = cur_data.groupby("method").apply(
        lambda x: np.trapezoid(
            x.sort_values("FPR")["TPR"], x.sort_values("FPR")["FPR"]
        ), 
        include_groups=False
    )
    # Plot the ROC curve
    sns.lineplot(
        data=cur_data,
        x="FPR",
        y="TPR",
        hue="method",
        style="method",
        palette=method_palette,
        ax=axes[i],
        # Show the points
        markers=method_markers,
        dashes=False,
        markersize=7.5,
        markeredgewidth=0.5,
        markeredgecolor="gray", 
        legend=False,
        rasterized=True,
        estimator=None,
    )

    # Add AUC values as legend like text
    for j, method in enumerate(auc_data.index):
        auc = auc_data[method]
        # Add color to match the palette
        color = method_palette[method]
        axes[i].text(
            0.95,
            0.05 + j * 0.075,
            f"{method}: {auc:.2f}",
            color=color,
            transform=axes[i].transAxes,
            ha="right",
            va="bottom",
            fontsize=12,
            fontweight="bold",
        )
        # Add a large no-facecolor marker to the pthr value for each method
        pthr_data = cur_data[(cur_data["method"] == method) & (cur_data["threshold"] == pthr)]
        axes[i].scatter(
            pthr_data["FPR"],
            pthr_data["TPR"],
            color=color,
            s=125,
            edgecolor="black",
            linewidth=1.25,
            marker="o",
            facecolors="none",
        )

    axes[i].plot([0, 1], [0, 1], color="black", linestyle="--")
    axes[i].set_title(f"{pert}", fontsize=14, fontweight="bold")
    axes[i].set_xlabel("FPR", fontsize=12)
    axes[i].set_ylabel("TPR", fontsize=12)
    axes[i].set_xlim(-0.05, 1.05)
    axes[i].set_ylim(-0.05, 1.05)
    axes[i].grid("both", linestyle="--", linewidth=0.75, alpha=0.5, color="lightgrey")

# Add a text indicating the circle is the p-value threshold under fig.title and above the first subplot
fig.text(
    0.9,
    .97,
    f"Black circles represent the TPR and FPR values at p-value threshold of {pthr}",
    ha="right",
    va="center",
    fontsize=12,
    fontstyle="italic",
    color="gray",
)

# Finalize the plot
plt.tight_layout()
plots.finalize_plot(
    fig, 
    show=True,
    save=save_to_folder,
    filename=f"ROC_curves_perturbations_methods", 
    filepath=figure_path,
    formats=figure_formats, 
    transparent=transparent_bg,
    dpi=figure_dpi,
)
```

**ROC Analysis Results:**

The ROC curves reveal consistent performance patterns across scenarios:

- **Overall Performance:** **ProteoForge** and **PeCorA** occupy the top tier, with **ProteoForge** generally providing the best discrimination across thresholds except the 50% peptide perturbation scenario, it is the worst performer.
- **Behavior at `p = 0.001`:** The black circles show a practical trade-off—**ProteoForge** achieves higher sensitivity (TPR) at the cost of higher false positive rate (FPR) compared to **PeCorA**
- **COPF Characteristics:** **COPF** performs poorly with single-peptide perturbations since its clustering approach requires multiple discordant peptides. Performance improves as more peptides are perturbed (max=0.69 AUC at 50% perturbation).

**Practical Recommendations:**

| Scenario | Recommended Method | Rationale |
| --- | --- | --- |
| Maximize sensitivity | **ProteoForge** | Higher TPR across thresholds |
| Control false positives | **PeCorA** | More specific at fixed cutoffs |
| Multiple perturbed peptides | **COPF** | Clustering benefits from multiple signals |

##### Precision-Recall (PR) Analysis¶

ROC curves can be overly optimistic with class imbalance (far more unperturbed than perturbed peptides). **Precision-Recall (PR) curves** provide a more focused evaluation.

**Interpretation:**

- Goal: stay in the **top-right corner** (high precision + high recall)
- More informative for understanding the trustworthiness of positive identifications

```
## Precision-Recall Curves for different perturbations and methods
fig, axes = plt.subplots(1, 4, figsize=(16, 4), sharey=True, gridspec_kw={"wspace": 0.05})

for i, pert in enumerate(data_order):
    cur_data = results_data[results_data["perturbation"] == pert].copy()
    # # Add a small value to avoid division by zero
    cur_data["Precision"] = cur_data["TP"] / (cur_data["TP"] + cur_data["FP"] + 1e-6)
    cur_data["Recall"] = cur_data["TP"] / (cur_data["TP"] + cur_data["FN"] + 1e-6)
    
    cur_data = cur_data.sort_values("Recall", ascending=False)
    cur_data = cur_data[~((cur_data["Recall"] == 0) & (cur_data["Precision"] == 0))]
    
    # Ensure the data is complete for PR curves
    cur_data = cur_data.groupby("method").apply(
        lambda x: utils.complete_curve_data(x, 'PR', 'Recall', 'Precision')
    ).reset_index(drop=True)
    
    # Calculate AUC per method from Precision and Recall
    # Here we use the trapezoidal rule to calculate the area under the curve
    f1_data = cur_data.groupby("method").apply(
        lambda x: np.trapezoid(
            x.sort_values("Recall")["Precision"], x.sort_values("Recall")["Recall"]
        ),
        include_groups=False
    )

    # Plot the ROC curve
    sns.lineplot(
        data=cur_data,
        x="Recall",
        y="Precision",
        hue="method",
        style="method",
        palette=method_palette,
        ax=axes[i],
        # Show the points
        markers=method_markers,
        dashes=False,
        markersize=7.5,
        markeredgewidth=0.5,
        markeredgecolor="gray", 
        legend=False,
        rasterized=True,
        estimator=None,
    )

    # Add AUC values as legend like text
    for j, method in enumerate(f1_data.index):
        auc = f1_data[method]
        # Add color to match the palette
        color = method_palette[method]
        axes[i].text(
            0.05,
            0.05 + j * 0.075,
            f"{method}: {auc:.2f}",
            color=color,
            transform=axes[i].transAxes,
            ha="left",
            va="bottom",
            fontsize=12,
            fontweight="bold",
        )
        # Add a large no-facecolor marker to the pthr value for each method
        pthr_data = cur_data[(cur_data["method"] == method) & (cur_data["threshold"] == pthr)]
        axes[i].scatter(
            pthr_data["Recall"],
            pthr_data["Precision"],
            color=color,
            s=125,
            edgecolor="black",
            linewidth=1.25,
            marker="o",
            facecolors="none",
        )

    axes[i].set_title(f"{pert}", fontsize=14, fontweight="bold")
    axes[i].set_xlabel("Recall", fontsize=12)
    axes[i].set_ylabel("Precision", fontsize=12)
    axes[i].set_xlim(-0.05, 1.05)
    axes[i].set_ylim(-0.05, 1.05)
    axes[i].grid("both", linestyle="--", linewidth=0.75, alpha=0.5, color="lightgrey")

# Add a text indicating the circle is the p-value threshold under fig.title and above the first subplot
fig.text(
    0.9,
    .97,
    f"Black circles represent the Precision and Recall values at p-value threshold of {pthr}",
    ha="right",
    va="center",
    fontsize=12,
    fontstyle="italic",
    color="gray",
)

# Finalize the plot
plt.tight_layout()
plots.finalize_plot(
    fig, 
    show=True,
    save=save_to_folder,
    filename=f"PR_curves_perturbations_methods", 
    filepath=figure_path,
    formats=figure_formats, 
    transparent=transparent_bg,
    dpi=figure_dpi,
)
```

**PR Analysis Results:**

The PR curves reveal performance differences not obvious from ROC analysis:

**Simple Scenarios (1 or 2 Peptides):**

- **PeCorA** and **ProteoForge** are the best choices
- **PeCorA** shows a slight edge in single-peptide cases (AUPRC = `0.76`)
- **COPF** is ineffective (AUPRC = `0.03`)—its clustering algorithm has no pattern to detect

**Random Scenario:**

- **ProteoForge** stands out (AUPRC = `0.48`)—maintains high precision across a wide recall range
- Outperforms **PeCorA** (`0.40`) and **COPF** (`0.29`) in this realistic, mixed-signal environment

**Extreme Perturbation (%50 Peptides):**

- Results invert: **COPF** (AUPRC = `0.42`) is the top performer
- Widespread perturbation creates the large clusters **COPF** is designed to find
- **PeCorA** and **ProteoForge** performance degrades significantly, this is due to 50% sometimes is more than half of the peptides perturbed per protein, making it hard to distinguish discordance,

**Summary:**

| Method | Strength | Best Use Case |
| --- | --- | --- |
| **ProteoForge** | Most versatile and robust | General-purpose, complex scenarios |
| **PeCorA** | Highly specific detection | Singular, isolated changes |
| **COPF** | Specialized clustering | Large fraction of discordant peptides |

##### F1 Score Across P-Value Thresholds¶

###### Optimal Thresholds at Maximum F1¶

For practical use, a single optimal p-value threshold is often needed. The table below shows the threshold that maximizes the **F1 score** (harmonic mean of precision and recall) for each method and scenario.

```
# Identify the best threshold for each method and perturbation
best_thresholds = results_data.groupby(["method", "perturbation"]).apply(
    lambda x: x.loc[x["F1"].idxmax(), ["threshold", "F1"]]
).reset_index()
best_thresholds.columns = ["method", "perturbation", "threshold", "F1"]
best_thresholds
```

|  | method | perturbation | threshold | F1 |
| --- | --- | --- | --- | --- |
| 0 | COPF | %50 Peptides | 0.1000 | 0.4876 |
| 1 | COPF | 1 Peptide | 0.8000 | 0.0679 |
| 2 | COPF | 2 Peptides | 0.0100 | 0.2506 |
| 3 | COPF | Random (1 to %50) Peptides | 0.1000 | 0.3820 |
| 4 | PeCorA | %50 Peptides | 0.1000 | 0.4455 |
| 5 | PeCorA | 1 Peptide | 0.0000 | 0.7231 |
| 6 | PeCorA | 2 Peptides | 0.0000 | 0.5524 |
| 7 | PeCorA | Random (1 to %50) Peptides | 0.0100 | 0.4690 |
| 8 | ProteoForge | %50 Peptides | 0.7000 | 0.4215 |
| 9 | ProteoForge | 1 Peptide | 0.0000 | 0.6980 |
| 10 | ProteoForge | 2 Peptides | 0.0000 | 0.6374 |
| 11 | ProteoForge | Random (1 to %50) Peptides | 0.0001 | 0.4859 |

The table provides concrete p-values to maximize F1 for any given perturbation.

###### Visualizing F1 Across Thresholds¶

The plots below show F1 score sensitivity to threshold choice. The x-axis displays `-log10(p-value)`, so smaller (more significant) p-values appear on the right. Black circles mark optimal F1 values from the table above.

```
# F1 Score for different perturbations and methods at the best threshold
fig, axes = plt.subplots(1, 4, figsize=(16, 4), sharey=True, gridspec_kw={"wspace": 0.05})
# Run the loop for each perturbation
for i, pert in enumerate(data_order):
    cur_data = results_data[results_data["perturbation"] == pert]
    cur_data["-log10(threshold)"] = -np.log10(cur_data["threshold"])
    sns.lineplot(
        data=cur_data,
        x="-log10(threshold)",
        y="F1",
        hue="method",
        style="method",
        palette=method_palette,
        ax=axes[i],
        markers=method_markers,
        dashes=False,
        markersize=7.5,
        markeredgewidth=0.5,
        markeredgecolor="gray",
        legend=False,
        rasterized=True,
        estimator=None,
    )

    # Add The highest F1 values as legend like text 
    for j, method in enumerate(best_thresholds[best_thresholds["perturbation"] == pert]["method"]):
        f1 = best_thresholds[(best_thresholds["perturbation"] == pert) & (best_thresholds["method"] == method)]["F1"].values[0]
        
        # Add a large no-facecolor marker to the p-value that gives the highest F1 score for each method
        pthr_data = cur_data[
            (cur_data["method"] == method) & 
            (cur_data["threshold"] == best_thresholds[
                (best_thresholds["perturbation"] == pert) & 
                (best_thresholds["method"] == method)
            ]["threshold"].values[0])
        ]
        axes[i].scatter(
            -np.log10(pthr_data["threshold"]),
            pthr_data["F1"],
            color=color,
            s=125,
            edgecolor="black",
            linewidth=1.25,
            marker="o",
            facecolors="none",
        )
        # Add color to match the palette
        color = method_palette[method]
        axes[i].text(
            0.05,
            0.75 + j * 0.075,
            f"{method}: {f1:.2f}",
            color=color,
            transform=axes[i].transAxes,
            ha="left",
            va="bottom",
            fontsize=12,
            fontweight="bold",
        )

    axes[i].set_title(f"{pert}", fontsize=14, fontweight="bold")
    axes[i].set_xlabel("-log 10 P-value Threshold", fontsize=12)
    axes[i].set_ylabel("F1 Score", fontsize=12)
    axes[i].set_xlim(-0.5, 15.5)
    axes[i].set_ylim(-0.05, 1.05)
    axes[i].grid("both", linestyle="--", linewidth=0.75, alpha=0.5, color="lightgrey")

fig.text(
    0.9,
    .97,
    f"Black circles represent the F1 values at the best threshold for each method",
    ha="right",
    va="center",
    fontsize=12,
    fontstyle="italic",
    color="gray",
)
# Finalize the plot
plt.tight_layout()
plots.finalize_plot(
    fig, 
    show=True,
    save=save_to_folder,
    filename=f"F1_scores_perturbations_methods", 
    filepath=figure_path,
    formats=figure_formats, 
    transparent=transparent_bg,
    dpi=figure_dpi,
)
```

**F1 Analysis Results:**

These plots highlight a critical difference between methods: **robustness to threshold choice**.

**Peak Performance:**

| Scenario | Best Method | F1 Score |
| --- | --- | --- |
| 1 Peptide | **PeCorA** | 0.72 |
| 2 Peptides | **ProteoForge** | 0.64 |
| Random | **ProteoForge** | 0.49 |
| %50 Peptides | **COPF** | 0.49 |

**Key Insight — Curve Shape Matters:**

- **ProteoForge** (red triangles): Broad, flat plateau—maintains optimal F1 across a wide range of thresholds. *Practical advantage: stability and reliability without precise tuning*
- **PeCorA** (orange squares): Sharp peak—performance is highly sensitive to threshold choice. Slightly too-stringent thresholds cause F1 to collapse
- **COPF**: Requires widespread perturbation to be effective

**Practical Implication:** Prefer methods with wide plateaus (robust) for consistent cross-dataset performance; use sharply peaked methods only when the optimal threshold can be reliably determined.

##### MCC Score Across P-Value Thresholds¶

###### Optimal Thresholds at Maximum MCC¶

The **Matthews Correlation Coefficient (MCC)** provides a robust single-number summary accounting for all four confusion-matrix values:

| MCC Value | Interpretation |
| --- | --- |
| `+1` | Perfect prediction |
| `0` | No better than random |
| `-1` | Total disagreement |

The table below lists thresholds that maximize MCC—use these as alternative operating points when balanced performance is the priority.

```
best_thresholds = results_data.groupby(["method", "perturbation"]).apply(
    lambda x: x.loc[x["MCC"].idxmax(), ["threshold", "MCC"]]
).reset_index()
best_thresholds.columns = ["method", "perturbation", "threshold", "MCC"]
best_thresholds
```

|  | method | perturbation | threshold | MCC |
| --- | --- | --- | --- | --- |
| 0 | COPF | %50 Peptides | 0.1000 | 0.2981 |
| 1 | COPF | 1 Peptide | 0.0000 | 0.0000 |
| 2 | COPF | 2 Peptides | 0.0100 | 0.1620 |
| 3 | COPF | Random (1 to %50) Peptides | 0.1000 | 0.2322 |
| 4 | PeCorA | %50 Peptides | 0.1000 | 0.2100 |
| 5 | PeCorA | 1 Peptide | 0.0000 | 0.7193 |
| 6 | PeCorA | 2 Peptides | 0.0000 | 0.5133 |
| 7 | PeCorA | Random (1 to %50) Peptides | 0.0100 | 0.3515 |
| 8 | ProteoForge | %50 Peptides | 0.6000 | 0.1620 |
| 9 | ProteoForge | 1 Peptide | 0.0000 | 0.6885 |
| 10 | ProteoForge | 2 Peptides | 0.0000 | 0.6093 |
| 11 | ProteoForge | Random (1 to %50) Peptides | 0.0001 | 0.3794 |

This table provides MCC-optimized operating points, which often differ from F1-optimized thresholds depending on research goals.

###### Visualizing MCC Across Thresholds¶

The plots below show MCC score across the full range of p-value thresholds. Black circles mark maximum MCC values for each method.

```
# MCC Score for different perturbations and methods at the best threshold
fig, axes = plt.subplots(1, 4, figsize=(16, 4), sharey=True, gridspec_kw={"wspace": 0.05})
# Run the loop for each perturbation
for i, pert in enumerate(data_order):
    cur_data = results_data[results_data["perturbation"] == pert].copy()
    cur_data["-log10(threshold)"] = -np.log10(cur_data["threshold"])
    sns.lineplot(
        data=cur_data,
        x="-log10(threshold)",
        y="MCC",
        hue="method",
        style="method",
        palette=method_palette,
        ax=axes[i],
        markers=method_markers,
        dashes=False,
        markersize=7.5,
        markeredgewidth=0.5,
        markeredgecolor="gray",
        legend=False,
        rasterized=True,
        estimator=None,
    )

    # Add The highest MCC values as legend like text 
    for j, method in enumerate(best_thresholds[best_thresholds["perturbation"] == pert]["method"]):
        mcc = best_thresholds[(best_thresholds["perturbation"] == pert) & (best_thresholds["method"] == method)]["MCC"].values[0]
        
        # Add a large no-facecolor marker to the p-value that gives the highest MCC score for each method
        pthr_data = cur_data[
            (cur_data["method"] == method) & 
            (cur_data["threshold"] == best_thresholds[
                (best_thresholds["perturbation"] == pert) & 
                (best_thresholds["method"] == method)
            ]["threshold"].values[0])
        ]
        axes[i].scatter(
            -np.log10(pthr_data["threshold"]),
            pthr_data["MCC"],
            color=color,
            s=125,
            edgecolor="black",
            linewidth=1.25,
            marker="o",
            facecolors="none",
        )
        # Add color to match the palette
        color = method_palette[method]
        axes[i].text(
            0.05,
            0.75 + j * 0.075,
            f"{method}: {mcc:.2f}",
            color=color,
            transform=axes[i].transAxes,
            ha="left",
            va="bottom",
            fontsize=12,
            fontweight="bold",
        )

    axes[i].set_title(f"{pert}", fontsize=14, fontweight="bold")
    axes[i].set_xlabel("-log 10 P-value Threshold", fontsize=12)
    axes[i].set_ylabel("MCC Score", fontsize=12)
    axes[i].set_xlim(-0.5, 15.5)
    axes[i].set_ylim(-0.25, 1.05)
    axes[i].grid("both", linestyle="--", linewidth=0.75, alpha=0.5, color="lightgrey")
    
fig.text(
    0.9,
    .97,
    f"Black circles represent the MCC values at the best threshold for each method",
    ha="right",
    va="center",
    fontsize=12,
    fontstyle="italic",
    color="gray",
)
# Finalize the plot
plt.tight_layout()
plots.finalize_plot(
    fig, 
    show=True,
    save=save_to_folder,
    filename=f"MCC_scores_perturbations_methods", 
    filepath=figure_path,
    formats=figure_formats, 
    transparent=transparent_bg,
    dpi=figure_dpi,
)
```

**MCC Analysis Results:**

The MCC analysis confirms performance trends observed with other metrics.

**Peak Performance:**

| Scenario | Best Method | MCC |
| --- | --- | --- |
| 1 Peptide | **PeCorA** | 0.72 |
| 2 Peptides | **ProteoForge** | 0.61 |
| Random | **ProteoForge** | 0.38 |
| %50 Peptides | **COPF** | 0.30 |

**Robustness Analysis:**

- **ProteoForge:** Wide, stable plateau—performance stays high across a large range of p-value cutoffs → *robust and easy to use*
- **PeCorA:** Sharp peak—excellent at a narrow range but sensitive to threshold choice.
- **COPF:** Requires 50% perturbation to increase around 0.3; otherwise MCC is low

---

#### Conclusions¶

Across all four evaluation metrics (**ROC**, **Precision-Recall**, **F1**, **MCC**), a consistent picture emerges:

| Method | Characteristics | Recommended Use |
| --- | --- | --- |
| **ProteoForge** | Most robust, high-performing, versatile | General choice—stable across scenarios and thresholds |
| **PeCorA** | Excellent specialist | Isolated discordance with careful threshold tuning |
| **COPF** | Specialized clustering | Large fraction of discordant peptides expected |

> Note: **50% Peptides Scenario:** This scenario is particularly challenging since it can involve perturbing more than half the peptides per protein, making it difficult to distinguish discordance effectively for PeCorA and ProteoForge.

---

#### Execution Time¶

```
print("Notebook Execution Time:", utils.prettyTimer(utils.getTime() - startTime))
```

```
Notebook Execution Time: 00h:00m:07s
```
