## Supplementary material for "ProteoForge: An Imputation-Aware Framework for Differential Proteoform Discovery in Bottom-Up Proteomics": Analysis notebooks as html render: Notebook S2.html

### Notebook S2

###### Enes Kemal Ergin

#### 2025-12-08 23:00:14

### Supplementary Notebook 2: Peptide grouping benchmark using SWATH-MS Interlab Data¶

- **License:** Creative Commons Attribution-NonCommercial 4.0 International License
- **Version:** 0.2
- **Edit Log:**
  - 2025-11-28: Initial version of the notebook
  - 2025-12-08: Revise the whole notebook, ensuring clarity and correctness

---

**Requirements:**

The following preprocessing and method runs must be completed (in order) before executing this notebook:

1. `01-DataProcessing.R` — Prepares SWATH‑MS Interlab data and writes processed datasets to `data/prepared/` as `.feather` files
2. `02-runCOPF.R` — Runs COPF; saves results to `data/results/` as `COPF_{dataID}_result.feather`
3. `04-runProteoForge.py` — Runs ProteoForge; saves results to `data/results/` as `ProteoForge_{dataID}_result.feather`

**Expected files:** `data/results/COPF_*`, `data/results/ProteoForge_*` (feather format)

> **Note:** Feather (`.feather`) is a fast, cross-language binary table format readable from both R and Python—ideal for exchanging processed data, but not human-readable.

---

**Data Information:**

This analysis uses three perturbation datasets created by `01-DataProcessing.R`. Each dataset corresponds to a perturbation scenario with a short identifier (used in filenames) and a display name (used in figures):

| Identifier | Display Name | Description |
| --- | --- | --- |
| `2pep` | 2 Peptides | Perturb exactly 2 peptides per selected protein |
| `050pep` | %50 Peptides | Perturb ~50% of peptides per selected protein |
| `random` | Random (2 to %50) Peptides | Perturb 2–50% of peptides per protein (randomly) |

> **Note:** The `1pep` scenario was excluded since a single perturbed peptide should not group with unperturbed ones, making it trivial.
>
> `050pep` is tricky since proteins with 5 peptides will default to perturbing 3 peptides (60%), while those with 4 peptides will perturb 2 (50%). The actual percentage varies by protein, but the original COPF publication used the ceiling to round up, which we also follow here.

**How perturbations are generated:**

1. `01-DataProcessing.R` reads the SWATH‑MS interlab table, filters proteins/peptides, aggregates charge states, median-normalizes intensities, and selects proteins with ≥4 peptides and sufficient replicates
2. A set of proteins to perturb is sampled, a per-protein reduction factor (`red_fac`) is drawn uniformly, and `generatePerturbedProfiles()` marks peptides as `perturbed_peptide` and reduces their intensities in `day5`
3. Prepared feather files are written to `./data/prepared/` and used by method scripts (`02`–`04`) to produce result files

---

**Purpose:**

This notebook compares two methods (**COPF**, **ProteoForge**) across three simulated perturbation scenarios and generates publication-quality performance figures (ROC, PR, F1, MCC). Note that PeCorA is not included because it does not output peptide groups—only flags individual discordant peptides—making direct comparison difficult.

---

#### Setup¶

This section imports required libraries, configures display settings, and defines paths for data and figures.

> **Note:** The HTML rendering of this notebook hides code cells by default. Click the "Code" buttons on the right to expand them.

##### Libraries¶

```
import os
import sys
import warnings

import numpy as np # Numerical computing
import pandas as pd # Data manipulation

import seaborn as sns # R-like high-level plots
import matplotlib.pyplot as plt # Python's base plotting 

sys.path.append('../')
from src import utils, plots

warnings.filterwarnings('ignore')
# Initialize the timer
startTime = utils.getTime()
```

##### Display Settings¶

The cell below configures pandas, matplotlib, and seaborn display options for improved readability of tables and figures, including color palettes and figure export settings.

```
## Figure Settings

# Define default colors and styles for plots
def_colors = [
    "#139593", "#fca311", "#e54f2a",
    "#c3c3c3", "#555555",
    "#690000", "#5f4a00", "#004549"
]

# Set seaborn style
sns.set_theme(
    style="white",
    context="paper",
    palette=def_colors,
    font_scale=1,
    rc={
        "figure.figsize": (6, 4),
        "font.family": "sans-serif",
        "font.sans-serif": ["Arial", "Ubuntu Mono"],
    }
)

# Figure Saving Settings
figure_formats = ["pdf"]
save_to_folder = True
transparent_bg = True
figure_dpi = 300

## Configure dataframe displaying
pd.set_option('display.float_format', lambda x: '%.4f' % x)
pd.set_option('display.max_rows', 50)
pd.set_option('display.max_columns', 50)
pd.set_option('display.width', 500)  # Set a wider display width

## Printing Settings
verbose = True

## Show the color-palettes
plots.color_palette( def_colors, save=False )
```

##### Data and Result Paths¶

Data and figures are organized in separate folders:

- `data_path` — Input directory containing method results (COPF, ProteoForge)
- `output_path` — Output directory for aggregated performance data
- `figure_path` — Directory for generated plots and figures

```
notebook_name = "02-GroupingBenchmark"
data_path = "./data/results/"
output_path = f"./data/results/"
figure_path = f"./figures/{notebook_name}/"

# Create the output folder if it does not exist
if not os.path.exists(output_path):
    os.makedirs(output_path)

# Create figure folder structure, if needed
if save_to_folder:
    for i in figure_formats:
        cur_folder = figure_path + i + "/"
        if not os.path.exists(cur_folder):
            os.makedirs(cur_folder)
```

##### Analysis Parameters¶

Key parameters for the evaluation framework:

- `seed = 42` — Random seed for reproducibility
- `pthr = 1e-3` — P-value threshold highlighted on ROC and PR curves (conventional cutoff)
- `thresholds` — Range of p-value thresholds from `1` to `1e-15` for calculating performance metrics
- `update_proteoform_grouping_in_COPF` — Custom function to build proteoform groups from COPF results per threshold (COPF returns proteoform groups for a single threshold only)

```
# Global variables
seed = 42 # Seed for reproducibility  
pthr = 10**-3  # p-value threshold for significance
thresholds = list(utils.generate_thresholds(10.0, -15, 1, 0, 1, 0.1)) # Thresholds for the analysis
data_order = ["2 Peptides", "Random (2 to %50) Peptides", "%50 Peptides"]

method_palette = {
    "COPF": "#139593",
    "ProteoForge": "#e54f2a",
}
method_styles = {
    "COPF": "--",
    "ProteoForge": ":",
}
method_markers = {
    "COPF": "o",
    "ProteoForge": "^",
}

def update_proteoform_grouping_in_COPF(
        data: pd.DataFrame,
        score_thr: float = 0.5,      # Specific to COPF if None, it will not be used
        pval_thr: float = 0.05,
        protein_col: str = "protein_id",
        cluster_col: str = "cluster",
        pval_col: str = "proteoform_score_pval_adj",
        score_col: str = "proteoform_score",
        sep: str = "_"
    ):
    # Create initial proteoform_id based on score and pval thresholds
    if score_thr is not None:
        condition = (data[score_col] >= score_thr) & (data[pval_col] <= pval_thr)
    else:
        condition = data[pval_col] <= pval_thr
    
    data['proteoform_id'] = data[protein_col]
    data.loc[condition, 'proteoform_id'] = data.loc[condition].apply(
        lambda row: f"{row[protein_col]}{sep}{int(row[cluster_col])}", axis=1
    )
    # If proteoform_id is exactly the same as protein_id, add "_0" to the cluster 0
    data.loc[data[protein_col] == data['proteoform_id'], 'proteoform_id'] = data[protein_col] + f"{sep}0"
    
    # Special case for cluster 100
    data.loc[data[cluster_col] == 100, 'proteoform_id'] = data[protein_col] + f"{sep}0"
    
    # Count unique proteoform_id per protein_id
    n_proteoforms = data.groupby(protein_col)['proteoform_id'].transform('nunique')
    
    # Adjust n_proteoforms if "_0" exists
    has_zero_cluster = data['proteoform_id'].str.endswith(f'{sep}0')
    n_proteoforms -= has_zero_cluster.groupby(data[protein_col]).transform('sum')
    
    # Final adjustment of proteoform_id
    data.loc[n_proteoforms.isin([0, 1]), 'proteoform_id'] = data[protein_col]
    
    return data
```

#### Performance Analysis¶

##### Data Aggregation and Preprocessing¶

The first step aggregates results from all methods and scenarios. The loop below:

1. Reads each method's `.feather` result files for all three perturbation scenarios
2. Calculates performance metrics (TP, FP, TN, FN) across p-value thresholds
3. Combines everything into a single DataFrame (`results_data`)

This unified dataset serves as the foundation for all subsequent performance visualizations.

```
st = utils.getTime()

results_data = pd.DataFrame()

## Comment out below for unneccessary re-run
# Variables
pval_col = "adj_pval"
peptide_col = "peptide_id"
protein_col = "protein_id"
# pepID_col = "peptideID" # Numerical ID for peptides
perturbed_protein_col = "perturbed_protein"
perturbed_peptide_col = "perturbed_peptide"
proteoform_group_col =  "PTM_id"
cluster_col = "ClusterID"
## Proteoform Grouping Benchmark of COPF and ProteoForge
# Read and combine the outputs:
for dataID in ["2pep", "050pep", "random"]:
    for curMethod in ["COPF", "ProteoForge"]:
        # Read the data
        data = pd.read_feather(f"{data_path}{curMethod}_{dataID}_result.feather")
            
        # Loop through the thresholds
        metric_data = []
        for thr in thresholds:
            # Determine the p-value column
            if curMethod == "COPF": 
                result = update_proteoform_grouping_in_COPF(
                    data, 
                    score_thr = None,
                    pval_thr = thr,
                    protein_col = protein_col,
                    cluster_col = 'cluster',
                    pval_col= 'proteoform_score_pval',
                    score_col = 'proteoform_score',
                )
                result['n_proteoforms'] = result.groupby(protein_col)['proteoform_id'].transform( lambda x: x.nunique() )
                result['WithProteoform'] = result['n_proteoforms'] > 1

                result = result[[protein_col, perturbed_protein_col, 'WithProteoform', ]].drop_duplicates().reset_index(drop=True)
            elif curMethod == "ProteoForge":
                result = data[[
                    protein_col, peptide_col,
                    perturbed_protein_col, perturbed_peptide_col, 
                    pval_col, cluster_col
                ]].drop_duplicates().reset_index(drop=True).copy()

                result[proteoform_group_col] = -1
                
                result.loc[
                    result[pval_col] < thr,  proteoform_group_col
                ] = result.loc[ result[pval_col] < thr,  cluster_col ]
                result['isPTM'] = result[proteoform_group_col] > 0

                result['isGroup'] = result.groupby([protein_col, proteoform_group_col])[peptide_col].transform('count') > 1
                result['dPF'] = result['isGroup'] & result['isPTM']
                # If per protein at least one True in dPF, then the protein has proteoform
                result['WithProteoform'] = result.groupby(protein_col)['dPF'].transform('any')
                # Drop the duplicates
                result = result[[protein_col, perturbed_protein_col, 'WithProteoform', ]].drop_duplicates().reset_index(drop=True)

            true_labels = result[perturbed_protein_col]
            pred_labels = result['WithProteoform']

            metrics = utils.calculate_metrics(
                true_labels=true_labels, pred_labels=pred_labels, 
                verbose=False, return_metrics=True
            )
            metrics['threshold'] = thr
            metric_data.append(pd.DataFrame([metrics]))
        metric_data = pd.concat(metric_data, axis=0, ignore_index=True)
        metric_data["method"] = curMethod
        metric_data["perturbation"] = dataID

        results_data = pd.concat([results_data, metric_data], axis=0, ignore_index=True)

print(f"Data shape: {results_data.shape}")
results_data["perturbation"] = results_data["perturbation"].replace({
    # "1pep": "1 Peptide",
    "2pep": "2 Peptides",
    "050pep": "%50 Peptides",
    "random": "Random (2 to %50) Peptides",
})

# Save the processed data
output_file = f"{output_path}peptide_grouping_performance_data.feather"
results_data.to_feather(output_file)
results_data.to_csv(output_file.replace('.feather', '.csv'), index=False)

results_data.tail()
```

```
Data shape: (150, 14)
```

|  | TP | FP | TN | FN | TPR | FPR | FDR | MCC | Precision | Recall | F1 | threshold | method | perturbation |
| --- | --- | --- | --- | --- | --- | --- | --- | --- | --- | --- | --- | --- | --- | --- |
| 145 | 950 | 909 | 308 | 50 | 0.9500 | 0.7469 | 0.4890 | 0.2746 | 0.5110 | 0.9500 | 0.6646 | 0.6000 | ProteoForge | Random (2 to %50) Peptides |
| 146 | 952 | 935 | 282 | 48 | 0.9520 | 0.7683 | 0.4955 | 0.2568 | 0.5045 | 0.9520 | 0.6595 | 0.7000 | ProteoForge | Random (2 to %50) Peptides |
| 147 | 960 | 956 | 261 | 40 | 0.9600 | 0.7855 | 0.4990 | 0.2534 | 0.5010 | 0.9600 | 0.6584 | 0.8000 | ProteoForge | Random (2 to %50) Peptides |
| 148 | 965 | 978 | 239 | 35 | 0.9650 | 0.8036 | 0.5033 | 0.2440 | 0.4967 | 0.9650 | 0.6558 | 0.9000 | ProteoForge | Random (2 to %50) Peptides |
| 149 | 968 | 994 | 223 | 32 | 0.9680 | 0.8168 | 0.5066 | 0.2359 | 0.4934 | 0.9680 | 0.6536 | 1.0000 | ProteoForge | Random (2 to %50) Peptides |

The table above shows the compiled data structure. Each row captures performance for a specific method and perturbation scenario at a given p-value threshold.

##### ROC Analysis¶

The **Receiver Operating Characteristic (ROC) curve** evaluates how well each method distinguishes between perturbed and unperturbed peptides by plotting True Positive Rate (TPR/sensitivity) against False Positive Rate (FPR/1-specificity).

**Interpretation:**

- A perfect classifier hugs the top-left corner
- **Area Under the Curve (AUC):** `1.0` = perfect, `0.5` = random chance
- Black circles indicate performance at `p = 1e-3`

```
# ROC Curves for different perturbations and methods
fig, axes = plt.subplots(1, 3, figsize=(12, 4), sharey=True, gridspec_kw={"wspace": 0.1})

for i, pert in enumerate(data_order):
    cur_data = results_data[results_data["perturbation"] == pert].copy()
    # Ensure the data is complete for ROC curves
    cur_data = cur_data.groupby("method").apply(
        lambda x: utils.complete_curve_data(x, 'ROC', 'FPR', 'TPR')
    ).reset_index(drop=True)
    
    # Calculate AUC per method from TPR and FPR
    auc_data = cur_data.groupby("method").apply(
        lambda x: np.trapezoid(
            x.sort_values("FPR")["TPR"], x.sort_values("FPR")["FPR"]
        ), 
        include_groups=False
    )
    # Plot the ROC curve
    sns.lineplot(
        data=cur_data,
        x="FPR",
        y="TPR",
        hue="method",
        style="method",
        palette=method_palette,
        ax=axes[i],
        # Show the points
        markers=method_markers,
        dashes=False,
        markersize=7.5,
        markeredgewidth=0.5,
        markeredgecolor="gray", 
        legend=False,
        rasterized=True, 
        estimator=None,
    )
    # Add AUC values as legend like text
    for j, method in enumerate(auc_data.index):
        auc = auc_data[method]
        # Add color to match the palette
        color = method_palette[method]
        axes[i].text(
            0.95,
            0.05 + j * 0.075,
            f"{method}: {auc:.2f}",
            color=color,
            transform=axes[i].transAxes,
            ha="right",
            va="bottom",
            fontsize=12,
            fontweight="bold",
        )
        # Add a large no-facecolor marker to the pthr value for each method
        pthr_data = cur_data[(cur_data["method"] == method) & (cur_data["threshold"] == pthr)]
        axes[i].scatter(
            pthr_data["FPR"],
            pthr_data["TPR"],
            color=color,
            s=125,
            edgecolor="black",
            linewidth=1.25,
            marker="o",
            facecolors="none",
        )

    axes[i].plot([0, 1], [0, 1], color="black", linestyle="--")
    axes[i].set_title(f"{pert}", fontsize=14, fontweight="bold")
    axes[i].set_xlabel("FPR", fontsize=12)
    axes[i].set_ylabel("TPR", fontsize=12)
    axes[i].set_xlim(-0.05, 1.05)
    axes[i].set_ylim(-0.05, 1.05)
    axes[i].grid("both", linestyle="--", linewidth=0.75, alpha=0.5, color="lightgrey")

# Add a text indicating the circle is the p-value threshold under fig.title and above the first subplot
fig.text(
    0.9,
    .97,
    f"Black circles represent the TPR and FPR values at p-value threshold of {pthr}",
    ha="right",
    va="center",
    fontsize=12,
    fontstyle="italic",
    color="gray",
)

# Finalize the plot
plt.tight_layout()
plots.finalize_plot(
    fig, 
    show=True,
    save=save_to_folder,
    filename=f"ROC_curves_perturbations_methods", 
    filepath=figure_path,
    formats=figure_formats, 
    transparent=transparent_bg,
    dpi=figure_dpi,
)
```

**ROC Analysis Results:**

The ROC curves reveal consistent performance patterns across scenarios:

- **Overall Performance:** **ProteoForge** consistently outperforms **COPF** across all perturbation scenarios, achieving higher AUC values (~0.85-83 vs ~0.77-75)
- **Behavior at `p = 0.001`:** **ProteoForge** reaches ~0.78 TPR but with higher FPR, while **COPF** shows lower TPR (~0.2) with lower FPR
- **Interpretation:** **ProteoForge** is more sensitive (finds more true positives) but may produce more false positives at this threshold. **COPF** is more conservative but misses many perturbed proteins

##### Precision-Recall (PR) Analysis¶

ROC curves can be overly optimistic with class imbalance (far more unperturbed than perturbed peptides). **Precision-Recall (PR) curves** provide a more focused evaluation.

**Interpretation:**

- Goal: stay in the **top-right corner** (high precision + high recall)
- More informative for understanding the trustworthiness of positive identifications

```
## Precision-Recall Curves for different perturbations and methods
fig, axes = plt.subplots(1, 3, figsize=(12, 4), sharey=True, gridspec_kw={"wspace": 0.1})

for i, pert in enumerate(data_order):
    cur_data = results_data[results_data["perturbation"] == pert].copy()
    # Add a small value to avoid division by zero
    cur_data["Precision"] = cur_data["TP"] / (cur_data["TP"] + cur_data["FP"] + 1e-6)
    cur_data["Recall"] = cur_data["TP"] / (cur_data["TP"] + cur_data["FN"] + 1e-6)
    
    cur_data = cur_data.sort_values("Recall", ascending=False)
    cur_data = cur_data[~((cur_data["Recall"] == 0) & (cur_data["Precision"] == 0))]
    
    # Ensure the data is complete for PR curves
    cur_data = cur_data.groupby("method").apply(
        lambda x: utils.complete_curve_data(x, 'PR', 'Recall', 'Precision')
    ).reset_index(drop=True)
    
    # Calculate AUC per method from Precision and Recall
    # Here we use the trapezoidal rule to calculate the area under the curve
    f1_data = cur_data.groupby("method").apply(
        lambda x: np.trapezoid(
            x.sort_values("Recall")["Precision"], x.sort_values("Recall")["Recall"]
        ),
        include_groups=False
    )
    
    # Plot the ROC curve
    sns.lineplot(
        data=cur_data,
        x="Recall",
        y="Precision",
        hue="method",
        style="method",
        palette=method_palette,
        ax=axes[i],
        # Show the points
        markers=method_markers,
        dashes=False,
        markersize=7.5,
        markeredgewidth=0.5,
        markeredgecolor="gray", 
        legend=False,
        rasterized=True,
        estimator=None,
    )

    # Add AUC values as legend like text
    for j, method in enumerate(f1_data.index):
        auc = f1_data[method]
        # Add color to match the palette
        color = method_palette[method]
        axes[i].text(
            0.05,
            0.05 + j * 0.075,
            f"{method}: {auc:.2f}",
            color=color,
            transform=axes[i].transAxes,
            ha="left",
            va="bottom",
            fontsize=12,
            fontweight="bold",
        )
        # Add a large no-facecolor marker to the pthr value for each method
        pthr_data = cur_data[(cur_data["method"] == method) & (cur_data["threshold"] == pthr)]
        axes[i].scatter(
            pthr_data["Recall"],
            pthr_data["Precision"],
            color=color,
            s=125,
            edgecolor="black",
            linewidth=1.25,
            marker="o",
            facecolors="none",
        )

    axes[i].set_title(f"{pert}", fontsize=14, fontweight="bold")
    axes[i].set_xlabel("Recall", fontsize=12)
    axes[i].set_ylabel("Precision", fontsize=12)
    axes[i].set_xlim(-0.05, 1.05)
    axes[i].set_ylim(-0.05, 1.05)
    axes[i].grid("both", linestyle="--", linewidth=0.75, alpha=0.5, color="lightgrey")

# Add a text indicating the circle is the p-value threshold under fig.title and above the first subplot
fig.text(
    0.9,
    .97,
    f"Black circles represent the Precision and Precision values at p-value threshold of {pthr}",
    ha="right",
    va="center",
    fontsize=12,
    fontstyle="italic",
    color="gray",
)

# Finalize the plot
plt.tight_layout()
plots.finalize_plot(
    fig, 
    show=True,
    save=save_to_folder,
    filename=f"PR_curves_perturbations_methods", 
    filepath=figure_path,
    formats=figure_formats, 
    transparent=transparent_bg,
    dpi=figure_dpi,
)
```

**PR Analysis Results:**

The PR curves reveal performance differences:

- **ProteoForge:** Achieves 0.87-81 PR-AUC across all scenarios
- **COPF:** Ranges from 0.73 (2pep) to 0.76 (%50)—worst in 2pep, best in %50
- **At `p = 1e-3`:** ProteoForge has ~0.78 precision and ~0.75 recall, while COPF has ~1.0 precision and ~0.2 recall
- **Key observation:** COPF shows sharp precision decline around 0.2 recall in 2pep → many false positives when trying to find more true positives. ProteoForge maintains higher precision even as recall increases

##### F1 Score Across P-Value Thresholds¶

###### Optimal Thresholds at Maximum F1¶

For practical use, a single optimal p-value threshold is often needed. The table below shows the threshold that maximizes the **F1 score** (harmonic mean of precision and recall) for each method and scenario.

```
# Identify the best threshold for each method and perturbation
best_thresholds = results_data.groupby(["method", "perturbation"]).apply(
    lambda x: x.loc[x["F1"].idxmax(), ["threshold", "F1"]]
).reset_index()
best_thresholds.columns = ["method", "perturbation", "threshold", "F1"]
best_thresholds
```

|  | method | perturbation | threshold | F1 |
| --- | --- | --- | --- | --- |
| 0 | COPF | %50 Peptides | 0.2000 | 0.6909 |
| 1 | COPF | 2 Peptides | 0.4000 | 0.6542 |
| 2 | COPF | Random (2 to %50) Peptides | 0.3000 | 0.6765 |
| 3 | ProteoForge | %50 Peptides | 0.0100 | 0.7360 |
| 4 | ProteoForge | 2 Peptides | 0.0001 | 0.7646 |
| 5 | ProteoForge | Random (2 to %50) Peptides | 0.0010 | 0.7538 |

The table provides concrete p-values to maximize F1 for any given situation. For example, using **ProteoForge** on the "Random" dataset, a threshold of `1e-3` achieves the best precision-recall balance.

###### Visualizing F1 Across Thresholds¶

The plots below show F1 score sensitivity to threshold choice. The x-axis displays `-log10(p-value)`, so smaller (more significant) p-values appear on the right. Black circles mark optimal F1 values from the table above.

```
# F1 Score for different perturbations and methods at the best threshold
fig, axes = plt.subplots(1, 3, figsize=(12, 4), sharey=True, gridspec_kw={"wspace": 0.1})

for i, pert in enumerate(data_order):
    cur_data = results_data[results_data["perturbation"] == pert].copy()
    cur_data["-log10(threshold)"] = -np.log10(cur_data["threshold"])
    sns.lineplot(
        data=cur_data,
        x="-log10(threshold)",
        y="F1",
        hue="method",
        style="method",
        palette=method_palette,
        ax=axes[i],
        markers=method_markers,
        dashes=False,
        markersize=7.5,
        markeredgewidth=0.5,
        markeredgecolor="gray",
        legend=False,
        rasterized=True,
        estimator=None,
    )

    # Add The highest F1 values as legend like text 
    for j, method in enumerate(best_thresholds[best_thresholds["perturbation"] == pert]["method"]):
        f1 = best_thresholds[(best_thresholds["perturbation"] == pert) & (best_thresholds["method"] == method)]["F1"].values[0]
        
        # Add a large no-facecolor marker to the p-value that gives the highest F1 score for each method
        pthr_data = cur_data[
            (cur_data["method"] == method) & 
            (cur_data["threshold"] == best_thresholds[
                (best_thresholds["perturbation"] == pert) & 
                (best_thresholds["method"] == method)
            ]["threshold"].values[0])
        ]
        axes[i].scatter(
            -np.log10(pthr_data["threshold"]),
            pthr_data["F1"],
            color=color,
            s=125,
            edgecolor="black",
            linewidth=1.25,
            marker="o",
            facecolors="none",
        )
        # Add color to match the palette
        color = method_palette[method]
        axes[i].text(
            0.05,
            0.8 + j * 0.1,
            f"{method}: {f1:.2f}",
            color=color,
            transform=axes[i].transAxes,
            ha="left",
            va="bottom",
            fontsize=12,
            fontweight="bold",
        )

    axes[i].set_title(f"{pert}", fontsize=14, fontweight="bold")
    axes[i].set_xlabel("-log 10 P-value Threshold", fontsize=12)
    axes[i].set_ylabel("F1 Score", fontsize=12)
    axes[i].set_xlim(-0.5, 15.5)
    axes[i].set_ylim(-0.05, 1.05)
    axes[i].grid("both", linestyle="--", linewidth=0.75, alpha=0.5, color="lightgrey")

fig.text(
    0.9,
    .97,
    f"Black circles represent the F1 values at the best threshold for each method",
    ha="right",
    va="center",
    fontsize=12,
    fontstyle="italic",
    color="gray",
)
# Finalize the plot
plt.tight_layout()
plots.finalize_plot(
    fig, 
    show=True,
    save=save_to_folder,
    filename=f"F1_scores_perturbations_methods", 
    filepath=figure_path,
    formats=figure_formats, 
    transparent=transparent_bg,
    dpi=figure_dpi,
)
```

**F1 Analysis Results:**

These plots highlight method robustness to threshold choice:

- **ProteoForge** consistently achieves higher F1-scores across all perturbation scenarios (peak ~0.74–0.76) compared to **COPF** (peak ~0.65–0.69)
- Optimal thresholds are generally more stringent for ProteoForge (~1e-3), indicating good precision-recall balance even at stricter cutoffs
- **Practical implication:** ProteoForge demonstrates robustness to threshold selection, making it a safer choice when the optimal cutoff is uncertain

##### MCC Score Across P-Value Thresholds¶

###### Optimal Thresholds at Maximum MCC¶

The **Matthews Correlation Coefficient (MCC)** provides a robust single-number summary accounting for all four confusion-matrix values:

- `+1` = Perfect prediction
- `0` = No better than random
- `-1` = Total disagreement

The table below lists thresholds that maximize MCC—use these as alternative operating points when balanced performance is the priority.

```
best_thresholds = results_data.groupby(["method", "perturbation"]).apply(
    lambda x: x.loc[x["MCC"].idxmax(), ["threshold", "MCC"]]
).reset_index()
best_thresholds.columns = ["method", "perturbation", "threshold", "MCC"]
best_thresholds
```

|  | method | perturbation | threshold | MCC |
| --- | --- | --- | --- | --- |
| 0 | COPF | %50 Peptides | 0.1000 | 0.5005 |
| 1 | COPF | 2 Peptides | 0.1000 | 0.4548 |
| 2 | COPF | Random (2 to %50) Peptides | 0.1000 | 0.4891 |
| 3 | ProteoForge | %50 Peptides | 0.0001 | 0.5336 |
| 4 | ProteoForge | 2 Peptides | 0.0000 | 0.6342 |
| 5 | ProteoForge | Random (2 to %50) Peptides | 0.0000 | 0.5850 |

This table provides MCC-optimized operating points, which often differ from F1-optimized thresholds depending on research goals.

###### Visualizing MCC Across Thresholds¶

The plots below show MCC score across the full range of p-value thresholds. Black circles mark maximum MCC values for each method.

```
# MCC Score for different perturbations and methods at the best threshold
fig, axes = plt.subplots(1, 3, figsize=(12, 4), sharey=True, gridspec_kw={"wspace": 0.1, "hspace": 0.1})

for i, pert in enumerate(data_order):
    cur_data = results_data[results_data["perturbation"] == pert].copy()
    cur_data["-log10(threshold)"] = -np.log10(cur_data["threshold"])
    sns.lineplot(
        data=cur_data,
        x="-log10(threshold)",
        y="MCC",
        hue="method",
        style="method",
        palette=method_palette,
        ax=axes[i],
        markers=method_markers,
        dashes=False,
        markersize=7.5,
        markeredgewidth=0.5,
        markeredgecolor="gray",
        legend=False,
        rasterized=True,
        estimator=None,
    )

    # Add The highest MCC values as legend like text 
    for j, method in enumerate(best_thresholds[best_thresholds["perturbation"] == pert]["method"]):
        mcc = best_thresholds[(best_thresholds["perturbation"] == pert) & (best_thresholds["method"] == method)]["MCC"].values[0]
        
        # Add a large no-facecolor marker to the p-value that gives the highest MCC score for each method
        pthr_data = cur_data[
            (cur_data["method"] == method) & 
            (cur_data["threshold"] == best_thresholds[
                (best_thresholds["perturbation"] == pert) & 
                (best_thresholds["method"] == method)
            ]["threshold"].values[0])
        ]
        axes[i].scatter(
            -np.log10(pthr_data["threshold"]),
            pthr_data["MCC"],
            color=color,
            s=125,
            edgecolor="black",
            linewidth=1.25,
            marker="o",
            facecolors="none",
        )
        # Add color to match the palette
        color = method_palette[method]
        axes[i].text(
            0.05,
            0.75 + j * 0.075,
            f"{method}: {mcc:.2f}",
            color=color,
            transform=axes[i].transAxes,
            ha="left",
            va="bottom",
            fontsize=12,
            fontweight="bold",
        )

    axes[i].set_title(f"{pert}", fontsize=14, fontweight="bold")
    axes[i].set_xlabel("-log 10 P-value Threshold", fontsize=12)
    axes[i].set_ylabel("MCC Score", fontsize=12)
    axes[i].set_xlim(-0.5, 15.5)
    axes[i].set_ylim(-0.25, 1.05)
    axes[i].grid("both", linestyle="--", linewidth=0.75, alpha=0.5, color="lightgrey")
    
fig.text(
    0.9,
    .97,
    f"Black circles represent the MCC values at the best threshold for each method",
    ha="right",
    va="center",
    fontsize=12,
    fontstyle="italic",
    color="gray",
)
# Finalize the plot
plt.tight_layout()
plots.finalize_plot(
    fig, 
    show=True,
    save=save_to_folder,
    filename=f"MCC_scores_perturbations_methods", 
    filepath=figure_path,
    formats=figure_formats, 
    transparent=transparent_bg,
    dpi=figure_dpi,
)
```

**MCC Analysis Results:**

The MCC analysis confirms performance trends observed with other metrics:

- **ProteoForge** consistently achieves higher MCC scores (peak ~0.63–0.53) compared to **COPF** (peak ~0.45–0.50)
- **ProteoForge** also from 0.1 to even lower p-value thresholds maintains better MCC than COPF
- This indicates a substantial performance gap between the two methods

---

#### Conclusions¶

Across all four evaluation metrics (**ROC**, **Precision-Recall**, **F1**, **MCC**), a consistent picture emerges:

| Method | Characteristics | Recommended Use |
| --- | --- | --- |
| **ProteoForge** | Most robust, high-performing, versatile | General choice—stable across scenarios and thresholds |
| **COPF** | Specialized clustering | Large fraction of discordant peptides expected |

> Note: **50% Peptides Scenario:** Even though this scenario had very negatively effected ProteoForge in identification benchmarks, when clusters are formed the performance improves significantly. This shows how well ProteoForges multiple-steps to identify potential proteoform groups work even in difficult scenarios.

---

#### Execution Time¶

```
print("Notebook Execution Time:", utils.prettyTimer(utils.getTime() - startTime))
```

```
Notebook Execution Time: 00h:00m:25s
```
