## Supplementary material for "ProteoForge: An Imputation-Aware Framework for Differential Proteoform Discovery in Bottom-Up Proteomics": Analysis notebooks as html render: Notebook S3.html

### Notebook S3

###### Enes Kemal Ergin

#### 2025-12-08 23:00:30

### Supplementary Notebook 3: Demo figure assembly for Figure 1¶

- **License:** Creative Commons Attribution-NonCommercial 4.0 International License
- **Version:** 0.2
- **Edit Log:**
  - 2025-11-28: Initial version of the notebook
  - 2025-12-08: Revise the whole notebook, ensuring clarity and correctness

---

**Requirements:**

No requirements beyond what can already be reached and done within the notebook.

---

**Data Information:**

The data used in this notebook is simulated data generated specifically for demonstration purposes. It does not represent real experimental data.

---

**Purpose:**

This notebook demonstrates the assembly of Figure 1 using randomly generated data to mimic a real proteomics data. The focus is to generate small plots that can be combined into a larger figure, showcasing the overview of the ProteoForge analysis workflow.

---

#### Setup¶

This section imports required libraries, configures display settings, and defines paths for data and figures.

> **Note:** The HTML rendering of this notebook hides code cells by default. Click the "Code" buttons on the right to expand them.

##### Libraries¶

```
import os
import sys
import warnings

import numpy as np # Numerical computing
import pandas as pd # Data manipulatio

import seaborn as sns # R-like high-level plots
import matplotlib.pyplot as plt # Python's base plotting 

sys.path.append('../')
from src import utils, plots

warnings.filterwarnings('ignore')

# Initialize the timer
startTime = utils.getTime()
```

##### Display Settings¶

The cell below configures pandas, matplotlib, and seaborn display options for improved readability of tables and figures, including color palettes and figure export settings.

```
# Create a dictionary for a greyscale color palette with 9 colors
def_colors = [
    "#139593", "#fca311", "#e54f2a",
    "#c3c3c3", "#555555",
    "#690000", "#5f4a00", "#004549"
]
demo_palette = {
    "control": "#8d99ae" ,
    "cond-1": "#48cae4" ,
    "cond-2": "#0096c7",
    "cond-3": "#023e8a",
}

# Set seaborn style
sns.set_theme(
    style="white",
    context="paper",
    palette=def_colors,
    font_scale=1,
    rc={
        "figure.figsize": (6, 4),
        "font.family": "sans-serif",
        "font.sans-serif": ["Arial", "Ubuntu Mono"],
    }
)

# Figure Saving Settings
figure_formats = ["pdf"]
save_to_folder = True
transparent_bg = True
figure_dpi = 300

## Configure dataframe displaying
pd.set_option('display.float_format', lambda x: '%.4f' % x)
pd.set_option('display.max_rows', 25)
pd.set_option('display.max_columns', 25)

plots.color_palette(
    def_colors, 
    save=False
)

plots.color_palette(
    demo_palette,
    name="Demo",
    save=False
)
```


##### Data and Result Paths¶

Data and figures are organized in separate folders:

- `output_path` — Output directory to store the demo data used in this notebook (`./data/prepared/`)
- `figure_path` — Directory for generated plots and figures (`./figures/`)

```
notebook_name = "demo"
output_path = f"./data/prepared/"
figure_path = f"./figures/{notebook_name}/"

# Create the output folder if it does not exist
if not os.path.exists(output_path):
    os.makedirs(output_path)

# Create figure folder structure, if needed
if save_to_folder:
    for i in figure_formats:
        cur_folder = figure_path + i + "/"
        if not os.path.exists(cur_folder):
            os.makedirs(cur_folder)
```

#### Demo Data¶

##### Data Generation¶

A simple data is generated for the sake of demonstration from 250 proteins, across 4 conditions with 5 replicates each. Each condition shifts by 1 log2 from control to condition 4.

```
# Parameters
n_proteins = 250
n_replicates = 5
conditions = ["control", "cond-1", "cond-2", "cond-3"]
shifts = {"control": 0, "cond-1": 1, "cond-2": 2, "cond-3": 3}  # Increasing differences
cond_filename_dict = {
    "control": [f"control_{i+1}" for i in range(n_replicates)],
    "cond-1": [f"cond-1_{i+1}" for i in range(n_replicates)],
    "cond-2": [f"cond-2_{i+1}" for i in range(n_replicates)],
    "cond-3": [f"cond-3_{i+1}" for i in range(n_replicates)],
}
cond_palette = {}
for condition, samples in cond_filename_dict.items():
    for sample in samples:
        cond_palette[sample] = demo_palette[condition]


# Generate protein IDs
protein_ids = [f"Protein_{i+1}" for i in range(n_proteins)]

# Function to generate base intensity values with more outliers closer to IQR
def generate_base_intensity():
    if np.random.rand() < 0.25:  # 30% chance to be an outlier
        return np.random.normal(loc=15, scale=4)  # Outliers closer to IQR
    else:
        return np.random.normal(loc=15, scale=2)  # Main distribution

# Generate data
data = []
for condition in conditions:
    for protein in protein_ids:
        base_intensity_log2 = generate_base_intensity() + shifts[condition]  # Apply shift
        for replicate in range(1, n_replicates + 1):
            replicate_intensity_log2 = base_intensity_log2 + np.random.normal(loc=0, scale=0.5)  # Smaller noise for replicates
            intensity = 2 ** replicate_intensity_log2  # Convert log2 intensity to linear scale
            data.append([protein, condition, replicate, intensity])

# Create DataFrame
df = pd.DataFrame(data, columns=["Protein", "Condition", "Replicate", "Intensity"])
df["Sample"] = df["Condition"] + "_" + df["Replicate"].astype(str)
df["log2_Intensity"] = np.log2(df["Intensity"])

# Display the first few rows of the DataFrame
print("Generated DataFrame (first 5 rows):")
print(df.head())
```

```
Generated DataFrame (first 5 rows):
     Protein Condition  Replicate  Intensity     Sample  log2_Intensity
0  Protein_1   control          1 27000.6421  control_1         14.7207
1  Protein_1   control          2 24273.2402  control_2         14.5671
2  Protein_1   control          3 18063.8069  control_3         14.1408
3  Protein_1   control          4 32948.5489  control_4         15.0079
4  Protein_1   control          5 50222.6445  control_5         15.6161
```

##### Boxplot of the Generated Data¶

The boxplot below displays the distribution of log2 peptide intensities across all samples. Each box represents a sample, colored by condition. Notice how the median intensity increases progressively from control through cond-3, reflecting the systematic shifts introduced during data generation.

```
# Initialize the figure
fig, ax = plt.subplots(
    figsize=(8, 4)
)

# Create the boxplot
sns.boxplot(
    x="Sample",
    y="log2_Intensity",
    hue="Condition",
    data=df,
    ax=ax,
    dodge=False,
    palette=demo_palette,
    # showfliers=False
    fliersize=0.5,
    linewidth=0.5,
)
# Set the x-axis labels
ax.set_xlabel("Samples", fontsize=14)
ax.set_ylabel("log2 - Peptide Intensity", fontsize=14)
# ax.set_xticklabels(ax.get_xticklabels(), rotation=90, horizontalalignment="right")
ax.set_xticklabels([])
ax.grid("both", linestyle="--", linewidth=0.75, alpha=0.5, color="lightgrey")
# place legend to upper right with 2 columns
ax.legend(
    ncol=4, frameon=False, title="Condition", 
    title_fontsize=12, fontsize=10, 
    # Center the legend on top
    bbox_to_anchor=(0.5, 1.1), loc="center"
)
sns.despine(left=True, bottom=True)
plt.tight_layout()

# Save the figure
plots.finalize_plot(
    plt.gcf(),
    filename="Demo_PeptideBoxplot",
    show=True,
    save=save_to_folder,
    filepath=figure_path,
    formats=figure_formats, 
    transparent=transparent_bg,
    dpi=figure_dpi,
)
```

##### ProteoForge's Processing Step:¶

###### 1. Initial State (Log2 Transformation)¶

The data is first log2 transformed to stabilize variance and make the data more normally distributed, which is a common preprocessing step in proteomics data analysis.

```
log2_data = df.pivot_table(
    index="Protein",
    columns="Sample",
    values="log2_Intensity",
    aggfunc="mean"
)

print("Generated log2 transformed data (first 5 rows):")
print(log2_data.head())
print()


# Plot the log2 raw values with a density plot
# Initialize the figure and axis
fig, ax = plt.subplots(
    figsize=(6, 3)
)
sns.kdeplot(
    data = log2_data,
    ax=ax,
    # fill=True,
    palette=cond_palette,
    linewidth=1.5,
    alpha=0.5,
    legend=False,
)

ax.set_xlabel("log2(Peptide Intensity)", fontsize=14)
ax.set_ylabel("Density", fontsize=14)
ax.set_title("Log2 Raw Values", fontsize=16, fontweight="bold")
ax.grid("both", linestyle="--", linewidth=0.75, alpha=0.5, color="lightgrey")
# Assemble the legend from status colors dictionary tumor and normal two
handles = []
labels = []
for k, v in demo_palette.items():
    handles.append(plt.Line2D([0], [0], color=v, lw=2))
    labels.append(k)

ax.legend(
    handles,
    labels,
    title="Condition",
    title_fontsize=12,
    fontsize=10,
    frameon=False,
    loc="upper right",
)
sns.despine(left=True, bottom=True)
plt.tight_layout()

# Save the figure
plots.finalize_plot(
    plt.gcf(),
    filename="Demo_Log2DensityPlot",
    show=True,
    save=save_to_folder,
    filepath=figure_path,
    formats=figure_formats, 
    transparent=transparent_bg,
    dpi=figure_dpi,
)
```

```
Generated log2 transformed data (first 5 rows):
Sample       cond-1_1  cond-1_2  cond-1_3  cond-1_4  cond-1_5  cond-2_1  \
Protein                                                                   
Protein_1     16.2662   16.4575   16.4714   15.3654   16.8313   18.0592   
Protein_10    15.8992   16.1557   16.4512   16.5807   14.8331    8.0537   
Protein_100   19.4083   19.3189   19.5896   18.5018   18.9829   18.4722   
Protein_101   15.1038   14.6603   16.1692   14.7098   14.8481   18.4904   
Protein_102   11.3785   11.2580   10.3157   10.7738   10.2850   17.7710   

Sample       cond-2_2  cond-2_3  cond-2_4  cond-2_5  cond-3_1  cond-3_2  \
Protein                                                                   
Protein_1     18.3328   18.2812   18.5381   18.1538   16.8453   16.9935   
Protein_10     8.1197    7.1558    7.5572    7.2730   15.4596   15.8069   
Protein_100   17.3684   18.6259   16.9120   16.8202   16.3610   17.2472   
Protein_101   18.4551   17.5342   17.5006   18.8558   19.4874   18.7560   
Protein_102   18.5215   18.3965   18.0426   17.8879   14.0855   13.0718   

Sample       cond-3_3  cond-3_4  cond-3_5  control_1  control_2  control_3  \
Protein                                                                      
Protein_1     16.9099   15.6672   16.4889    14.7207    14.5671    14.1408   
Protein_10    14.3366   15.7251   15.1585    16.0345    16.9961    16.1488   
Protein_100   17.1650   17.1074   16.9162    15.9323    16.0972    15.1698   
Protein_101   19.0051   19.4204   18.5448    18.4981    19.3284    18.7445   
Protein_102   14.3244   13.3429   12.9039    12.7125    13.8791    13.9723   

Sample       control_4  control_5  
Protein                            
Protein_1      15.0079    15.6161  
Protein_10     16.5650    17.0498  
Protein_100    15.3078    16.7295  
Protein_101    18.1039    18.2287  
Protein_102    13.7302    13.1367
```

###### 2. Centering¶

This standardizes the log2 intensities per sample (column) by computing z-scores: for each sample, subtract its mean and divide by its standard deviation. The result has mean ≈ 0 and unit variance per sample, which removes sample-specific location/scale differences (while preserving relative differences between proteins) and makes distributions comparable across samples for plotting and clustering.

```
centered_data = (log2_data - log2_data.mean()) / log2_data.std()
print("Generated centered data (first 5 rows):")
print(centered_data.head())
print()

# Plot the log2 centered values with a density plot
# Initialize the figure and axis
fig, ax = plt.subplots(
    figsize=(6, 3)
)

sns.kdeplot(
    data=centered_data,
    ax=ax,
    # fill=True,
    palette=cond_palette,
    linewidth=1.5,
    alpha=0.5,
    legend=False,
)

ax.set_xlabel("log2(Peptide Intensity)", fontsize=14)
ax.set_ylabel("Density", fontsize=14)
ax.set_title("Centered Values", fontsize=16, fontweight="bold")
ax.grid("both", linestyle="--", linewidth=0.75, alpha=0.5, color="lightgrey")
# Assemble the legend from status colors dictionary tumor and normal two
handles = []
labels = []
for k, v in demo_palette.items():
    handles.append(plt.Line2D([0], [0], color=v, lw=2))
    labels.append(k)

ax.legend(
    handles,
    labels,
    title="Condition",
    title_fontsize=12,
    fontsize=10,
    frameon=False,
    loc="upper right",
)
sns.despine(left=True, bottom=True)
plt.tight_layout()

# Save the figure
plots.finalize_plot(
    plt.gcf(),
    filename="Demo_CenteredDensityPlot",
    show=True,
    save=save_to_folder,
    filepath=figure_path,
    formats=figure_formats, 
    transparent=transparent_bg,
    dpi=figure_dpi,
)
```

```
Generated centered data (first 5 rows):
Sample       cond-1_1  cond-1_2  cond-1_3  cond-1_4  cond-1_5  cond-2_1  \
Protein                                                                   
Protein_1      0.0453    0.1013    0.1188   -0.3264    0.2584    0.4669   
Protein_10    -0.0973   -0.0167    0.1111    0.1401   -0.5149   -3.2792   
Protein_100    1.2660    1.2197    1.3008    0.8776    1.0910    0.6215   
Protein_101   -0.4064   -0.6012    0.0042   -0.5781   -0.5090    0.6283   
Protein_102   -1.8537   -1.9310   -2.2147   -2.0892   -2.2748    0.3590   

Sample       cond-2_2  cond-2_3  cond-2_4  cond-2_5  cond-3_1  cond-3_2  \
Protein                                                                   
Protein_1      0.5600    0.5563    0.6653    0.4961   -0.4616   -0.4161   
Protein_10    -3.2670   -3.6819   -3.4735   -3.6399   -0.9875   -0.8605   
Protein_100    0.1986    0.6876    0.0524   -0.0108   -0.6454   -0.3211   
Protein_101    0.6058    0.2717    0.2743    0.7630    0.5411    0.2439   
Protein_102    0.6307    0.6002    0.4786    0.3951   -1.5090   -1.8847   

Sample       cond-3_3  cond-3_4  cond-3_5  control_1  control_2  control_3  \
Protein                                                                      
Protein_1     -0.4694   -0.9044   -0.5983    -0.0881    -0.1388    -0.3234   
Protein_10    -1.4374   -0.8825   -1.0960     0.3626     0.6829     0.3685   
Protein_100   -0.3734   -0.3590   -0.4385     0.3275     0.3788     0.0311   
Protein_101    0.3189    0.5170    0.1708     1.2077     1.4718     1.2628   
Protein_102   -1.4420   -1.7846   -1.9394    -0.7770    -0.3715    -0.3814   

Sample       control_4  control_5  
Protein                            
Protein_1       0.0158     0.2020  
Protein_10      0.5488     0.6999  
Protein_100     0.1184     0.5887  
Protein_101     1.0756     1.1093  
Protein_102    -0.4216    -0.6590
```

###### 3. Control Adjusted Intensities¶

To highlight changes relative to the control condition, the mean log2 intensity of the control samples is subtracted from each protein's intensity across all conditions. This centers the data around the control, making it easier to visualize deviations in protein expression levels in response to different conditions.

```
# Adjusted data
# Calculate the mean of each peptide across the day1 samples
cntrPepMean = centered_data[cond_filename_dict["control"]].mean(axis=1)
# Substract cntrPepMean from each sample row-wise in centered_data
adjusted_dat = centered_data.subtract(cntrPepMean, axis=0)

print("Generated control adjusted data (first 5 rows):")
print(adjusted_dat.head())
print()

# Plot the log2 adjusted values with a density plot
# Initialize the figure and axis
fig, ax = plt.subplots(
    figsize=(6, 3)
)

sns.kdeplot(
    data=adjusted_dat,
    ax=ax,
    # fill=True,
    palette=cond_palette,
    linewidth=1.5,
    alpha=0.5,
    legend=False,
)

ax.set_xlabel("log2(Peptide Intensity)", fontsize=14)
ax.set_ylabel("Density", fontsize=14)
ax.set_title("Control Adjusted Values", fontsize=16, fontweight="bold")
ax.grid("both", linestyle="--", linewidth=0.75, alpha=0.5, color="lightgrey")
# Assemble the legend from status colors dictionary tumor and normal two
handles = []
labels = []
for k, v in demo_palette.items():
    handles.append(plt.Line2D([0], [0], color=v, lw=2))
    labels.append(k)

ax.legend(
    handles,
    labels,
    title="Condition",
    title_fontsize=12,
    fontsize=10,
    frameon=False,
    loc="upper right",
)
sns.despine(left=True, bottom=True)
plt.tight_layout()

# Save the figure
plots.finalize_plot(
    plt.gcf(),
    filename="Demo_AdjustedDensityPlot",
    show=True,
    save=save_to_folder,
    filepath=figure_path,
    formats=figure_formats, 
    transparent=transparent_bg,
    dpi=figure_dpi,
)
```

```
Generated control adjusted data (first 5 rows):
Sample       cond-1_1  cond-1_2  cond-1_3  cond-1_4  cond-1_5  cond-2_1  \
Protein                                                                   
Protein_1      0.1118    0.1678    0.1853   -0.2599    0.3249    0.5334   
Protein_10    -0.6298   -0.5492   -0.4214   -0.3924   -1.0474   -3.8117   
Protein_100    0.9771    0.9308    1.0119    0.5887    0.8021    0.3326   
Protein_101   -1.6318   -1.8266   -1.2212   -1.8036   -1.7345   -0.5971   
Protein_102   -1.3316   -1.4089   -1.6926   -1.5671   -1.7527    0.8811   

Sample       cond-2_2  cond-2_3  cond-2_4  cond-2_5  cond-3_1  cond-3_2  \
Protein                                                                   
Protein_1      0.6265    0.6228    0.7318    0.5626   -0.3951   -0.3496   
Protein_10    -3.7995   -4.2145   -4.0061   -4.1724   -1.5201   -1.3930   
Protein_100   -0.0903    0.3987   -0.2365   -0.2997   -0.9343   -0.6100   
Protein_101   -0.6196   -0.9538   -0.9512   -0.4625   -0.6843   -0.9815   
Protein_102    1.1528    1.1223    1.0007    0.9172   -0.9869   -1.3626   

Sample       cond-3_3  cond-3_4  cond-3_5  control_1  control_2  control_3  \
Protein                                                                      
Protein_1     -0.4029   -0.8379   -0.5318    -0.0216    -0.0723    -0.2569   
Protein_10    -1.9700   -1.4150   -1.6285    -0.1699     0.1503    -0.1640   
Protein_100   -0.6623   -0.6479   -0.7274     0.0386     0.0899    -0.2578   
Protein_101   -0.9066   -0.7084   -1.0547    -0.0178     0.2464     0.0373   
Protein_102   -0.9199   -1.2625   -1.4173    -0.2548     0.1506     0.1407   

Sample       control_4  control_5  
Protein                            
Protein_1       0.0823     0.2685  
Protein_10      0.0163     0.1674  
Protein_100    -0.1705     0.2998  
Protein_101    -0.1498    -0.1161  
Protein_102     0.1005    -0.1369
```

Now that the data has been processed, we can move on to examining how ProteoForge identifies peptide-level outliers and clusters peptides that behave differently across conditions.

---

##### ProteoForge's Main Analysis Steps¶

This section presents three example proteins with varying peptide behaviors and complexities to illustrate how ProteoForge performs its analysis. For each example, we provide both a peptide-level intensity plot and a clustered heatmap to visualize which peptides are grouped together based on their behavioral patterns.

###### Example 1: Simple Single-Condition Perturbation¶

This is the simplest case where all conditions show slightly decreased intensities relative to control as the baseline. However, peptides 1-4 in cond-3 exhibit a large increase (+0.75) as a perturbation, causing these peptides to behave distinctly compared to the remaining peptides of the same protein.

```
nPeps = 10
nReps = 5

# Generate for control mean is 0 and std is 0.25
control_data = np.random.normal(loc=0, scale=0.15, size=(nPeps, nReps))
control_data
# Move the mean to 1 and std to 0.5
cond1_data = np.random.normal(loc=-0.25, scale=0.15, size=(nPeps, nReps))
cond1_data
# Move the mean to 2 and std to 0.75
cond2_data = np.random.normal(loc=-0.27, scale=0.05, size=(nPeps, nReps))
cond2_data
# Move the mean to 3 and std to 1
cond3_data = np.random.normal(loc=-0.23, scale=0.1, size=(nPeps, nReps))
cond3_data

# put all the data together in a DataFrame
data = np.concatenate([control_data, cond1_data, cond2_data, cond3_data], axis=1)
data = pd.DataFrame(
    data,
    index=[f"Pep{i+1}" for i in range(nPeps)],
    columns=[
        "control_1", "control_2", "control_3", "control_4", "control_5",
        "cond-1_1", "cond-1_2", "cond-1_3", "cond-1_4", "cond-1_5",
        "cond-2_1", "cond-2_2", "cond-2_3", "cond-2_4", "cond-2_5",
        "cond-3_1", "cond-3_2", "cond-3_3", "cond-3_4", "cond-3_5"
    ]
)
# Adjust the data by control mean
data = data - data.loc[:, "control_1":"control_5"].mean(axis=1).values[:, None]

data = data.reset_index().melt(
    id_vars="index",
    var_name="Sample",
    value_name="Intensity"
)
data["Condition"] = data["Sample"].str.split("_", expand=True)[0]
# Introduce peptide moves
# data.loc[
#     ((data["Condition"] == "cond-1") & (data["index"].isin(["Pep1", "Pep2"]))), "Intensity"] += .5
# data.loc[
#     ((data["Condition"] == "cond-2") & (data["index"].isin(["Pep2", "Pep3"]))), "Intensity"] -= .75
data.loc[
    ((data["Condition"] == "cond-3") & (data["index"].isin(["Pep1", "Pep2", "Pep3", "Pep4"]))), "Intensity"] += .75

# Initialize the figure
fig, ax = plt.subplots(figsize=(6, 4))
sns.lineplot(
    ax=ax,
    data=data,
    x="index",
    y="Intensity",
    hue="Condition",
    palette=demo_palette,
    rasterized=True,
    # Add styling
    alpha=0.75,

    ## Markers
    marker="o",
    markersize=10,
    markeredgewidth=0,
    # markeredgecolor="black",
    ## Line style
    linewidth=1,
    linestyle="-",
    dashes=False,
    # errorbar
    err_style="bars",
    err_kws={"capsize": 5, "elinewidth": 2.5, "capthick": 2.5, "zorder": 0, "linewidth": 1.5},
)
ax.set_xlabel(f"Ordered Peptides (PeptideID)", fontsize=12)
ax.set_ylabel("Control Adjusted Intensity", fontsize=12)
ax.set_title(
    "",
    fontsize=16,
    fontweight="bold",
)
ax.grid("both", linestyle="--", linewidth=0.5, alpha=0.5, color="lightgrey")
plt.tight_layout()
# Save the figure
plots.finalize_plot(
    plt.gcf(),
    filename="Demo_Example1_PeptideConditionLineplot",
    show=True,
    save=save_to_folder,
    filepath=figure_path,
    formats=figure_formats, 
    transparent=transparent_bg,
    dpi=figure_dpi,
)
```

The line plot with control-adjusted intensities reveals that peptides 1-4 show a significant increase in cond-3 compared to the control, while peptides 5-10 remain relatively stable across all conditions. This indicates that peptides 1-4 are behaving differently from the rest of the peptides in this protein. Depending on the p-value threshold, either all peptides 1-4 would be marked as outliers or some might not be flagged by the linear model. However, with ProteoForge's clustering approach, these peptides would be grouped together based on their similar behavioral patterns.

---

Now moving on to the clustered heatmap, we highlight the clusters identified by hierarchical clustering based on Euclidean distance:

```
from scipy.spatial.distance import pdist, squareform

# Create a mapping from integer to unique peptide index (e.g., 1: 'Pep1', 2: 'Pep2', ...)
unique_indices = data['index'].unique()
int_to_index = {i+1: idx for i, idx in enumerate(unique_indices)}
index_to_int = {v: k for k, v in int_to_index.items()}

data['PeptideID'] = data['index'].map(index_to_int)
data = data.sort_values(by=['PeptideID', 'Condition', 'Sample']).reset_index(drop=True)
df = data.groupby(["PeptideID", "Condition"])["Intensity"].mean().unstack(level="Condition")

distance_vector = pdist(df.values, metric='euclidean')

# squareform converts it into a full, symmetric distance matrix for easier interpretation
dist_corr = pd.DataFrame(squareform(distance_vector), index=df.index, columns=df.index)

dist_labels = np.array([1,1,1,1,2,2,2,2,2,2])

# Rescale the distance matrix to a 0-1 range for better comparability with correlation matrices
dist_min = dist_corr.values.min()
dist_max = dist_corr.values.max()
if dist_max > dist_min:
    scaled_distance_matrix = (dist_corr - dist_min) / (dist_max - dist_min)
else:
    scaled_distance_matrix = dist_corr.copy()

ax = plots.clustering_check_with_single_heatmap(
    corr_matrix=scaled_distance_matrix,
    labels=dist_labels,
    protein_id='Protein 1',
    vmin=0, vmax=1
)
# Set the axis title
ax.set_title("Distance Matrix with Clustering Check", fontsize=14)
# Cbar label
cbar = ax.collections[0].colorbar
cbar.set_label('Scaled Euclidean Distance', labelpad=15, fontsize=12)

# Increase label size
ax.set_xlabel("Peptide ID", fontsize=14)
ax.set_ylabel("Peptide ID", fontsize=14)

# Increase tick label size
ax.tick_params(axis='x', labelsize=12)
ax.tick_params(axis='y', labelsize=12)

plt.tight_layout()
plots.finalize_plot(
    plt.gcf(),
    filename="Demo_Example1_PeptideConditionCorrelation",
    show=True,
    save=save_to_folder,
    filepath=figure_path,
    formats=figure_formats, 
    transparent=transparent_bg,
    dpi=figure_dpi,
)
```

The heatmap shows two distinct clusters: Cluster 1 (C1) contains peptides 1-4, which show a significant increase in cond-3, while Cluster 2 (C2) contains peptides 5-10, which remain stable across all conditions. This clustering confirms that peptides 1-4 are behaving differently from the rest of the peptides in this protein, successfully identifying a potential proteoform group.

---

###### Example 2: Multi-Condition Perturbations¶

The second example increases in complexity by introducing perturbations in two different conditions: peptides 1-2 show an increase (+0.75) in cond-1, while peptides 9-10 show a decrease (-0.75) in cond-3. This creates two distinct proteoform-like profiles affecting different regions of the protein, resulting in at least 3 clusters.

```
nPeps = 12
nReps = 5

# Generate for control mean is 0 and std is 0.25
control_data = np.random.normal(loc=0, scale=0.05, size=(nPeps, nReps))
control_data
# Move the mean to 1 and std to 0.5
cond1_data = np.random.normal(loc=-0.15, scale=0.05, size=(nPeps, nReps))
cond1_data
# Move the mean to 2 and std to 0.75
cond2_data = np.random.normal(loc=0.15, scale=0.05, size=(nPeps, nReps))
cond2_data
# Move the mean to 3 and std to 1
cond3_data = np.random.normal(loc=0.25, scale=0.14, size=(nPeps, nReps))
cond3_data

# put all the data together in a DataFrame
data = np.concatenate([control_data, cond1_data, cond2_data, cond3_data], axis=1)
data = pd.DataFrame(
    data,
    index=[f"Pep{i+1}" for i in range(nPeps)],
    columns=[
        "control_1", "control_2", "control_3", "control_4", "control_5",
        "cond-1_1", "cond-1_2", "cond-1_3", "cond-1_4", "cond-1_5",
        "cond-2_1", "cond-2_2", "cond-2_3", "cond-2_4", "cond-2_5",
        "cond-3_1", "cond-3_2", "cond-3_3", "cond-3_4", "cond-3_5"
    ]
)
# Adjust the data by control mean
data = data - data.loc[:, "control_1":"control_5"].mean(axis=1).values[:, None]

data = data.reset_index().melt(
    id_vars="index",
    var_name="Sample",
    value_name="Intensity"
)
data["Condition"] = data["Sample"].str.split("_", expand=True)[0]
# Introduce peptide moves
data.loc[
    ((data["Condition"] == "cond-1") & (data["index"].isin(["Pep1", "Pep2"]))), "Intensity"] += .75
# data.loc[
#     ((data["Condition"] == "cond-2") & (data["index"].isin(["Pep2", "Pep3"]))), "Intensity"] -= .75
data.loc[
    ((data["Condition"] == "cond-3") & (data["index"].isin(["Pep9", "Pep10"]))), "Intensity"] -= .75

# Initialize the figure
fig, ax = plt.subplots(figsize=(6, 4))
sns.lineplot(
    ax=ax,
    data=data,
    x="index",
    y="Intensity",
    hue="Condition",
    palette=demo_palette,
    rasterized=True,
    # Add styling
    alpha=0.75,

    ## Markers
    marker="o",
    markersize=10,
    markeredgewidth=0,
    # markeredgecolor="black",
    ## Line style
    linewidth=1,
    linestyle="-",
    dashes=False,
    # errorbar
    err_style="bars",
    err_kws={"capsize": 5, "elinewidth": 2.5, "capthick": 2.5, "zorder": 0, "linewidth": 1.5},
)
ax.set_xlabel(f"Ordered Peptides (PeptideID)", fontsize=12)
ax.set_ylabel("Control Adjusted Intensity", fontsize=12)
ax.set_title(
    "",
    fontsize=16,
    fontweight="bold",
)
ax.grid("both", linestyle="--", linewidth=0.5, alpha=0.5, color="lightgrey")
plt.tight_layout()
# Save the figure
plots.finalize_plot(
    plt.gcf(),
    filename="Demo_Example2_PeptideConditionLineplot",
    show=True,
    save=save_to_folder,
    filepath=figure_path,
    formats=figure_formats, 
    transparent=transparent_bg,
    dpi=figure_dpi,
)
```

The line plot demonstrates the increased complexity with two distinct perturbation events occurring in different conditions. We observe that:

- **Pep1 and Pep2** show a significant increase (+0.75) in cond-1, deviating from the baseline trend
- **Pep9 and Pep10** show a notable decrease (-0.75) in cond-3, moving in the opposite direction
- **Pep3-8 and Pep11-12** maintain the expected baseline behavior across all conditions

This creates an interesting scenario where outlier peptides exist at both ends of the protein (early and late peptides) and respond differently to distinct conditions. Such patterns could indicate condition-specific post-translational modifications or alternative proteoforms that are activated under different experimental treatments.

---

Now moving on to the clustered heatmap, we visualize how these distinct perturbations result in three separate clusters:

```
# Create a mapping from integer to unique peptide index (e.g., 1: 'Pep1', 2: 'Pep2', ...)
unique_indices = data['index'].unique()
int_to_index = {i+1: idx for i, idx in enumerate(unique_indices)}
index_to_int = {v: k for k, v in int_to_index.items()}

data['PeptideID'] = data['index'].map(index_to_int)
data = data.sort_values(by=['PeptideID', 'Condition', 'Sample']).reset_index(drop=True)
df = data.groupby(["PeptideID", "Condition"])["Intensity"].mean().unstack(level="Condition")

distance_vector = pdist(df.values, metric='euclidean')

# squareform converts it into a full, symmetric distance matrix for easier interpretation
dist_corr = pd.DataFrame(squareform(distance_vector), index=df.index, columns=df.index)

dist_labels = np.array([1,1,2,2,2,2,2,2,3,3,2,2])

# Rescale the distance matrix to a 0-1 range for better comparability with correlation matrices
dist_min = dist_corr.values.min()
dist_max = dist_corr.values.max()
if dist_max > dist_min:
    scaled_distance_matrix = (dist_corr - dist_min) / (dist_max - dist_min)
else:
    scaled_distance_matrix = dist_corr.copy()

ax = plots.clustering_check_with_single_heatmap(
    corr_matrix=scaled_distance_matrix,
    labels=dist_labels,
    protein_id='Protein 1',
    vmin=0, vmax=1
)
# Set the axis title
ax.set_title("Distance Matrix with Clustering Check", fontsize=14)
# Cbar label
cbar = ax.collections[0].colorbar
cbar.set_label('Scaled Euclidean Distance', labelpad=15, fontsize=12)

# Increase label size
ax.set_xlabel("Peptide ID", fontsize=14)
ax.set_ylabel("Peptide ID", fontsize=14)

# Increase tick label size
ax.tick_params(axis='x', labelsize=12)
ax.tick_params(axis='y', labelsize=12)

plt.tight_layout()
plots.finalize_plot(
    plt.gcf(),
    filename="Demo_Example2_PeptideConditionCorrelation",
    show=True,
    save=save_to_folder,
    filepath=figure_path,
    formats=figure_formats, 
    transparent=transparent_bg,
    dpi=figure_dpi,
)
```

The heatmap reveals three distinct clusters as expected: Cluster 1 (C1) contains peptides 1-2, which respond specifically to cond-1; Cluster 2 (C2) encompasses the baseline peptides (3-8 and 11-12) that maintain consistent behavior; and Cluster 3 (C3) contains peptides 9-10, which respond specifically to cond-3. This demonstrates ProteoForge's ability to identify multiple independent proteoform-like behaviors within a single protein.

---

###### Example 3: Complex Overlapping Perturbations¶

This is the most complex example, where multiple outlier peptides exist with different behaviors and some even overlap across conditions with variations. This example is provided to introduce the concept of real-world complexity where not all proteins clearly show 2-3 distinct and easy to identify proteoform profiles.

In this example, we simulate 8 peptides across 4 conditions with 5 replicates each. The baseline behavior shows a gradual increase from control through conditions 1-3. However, we introduce multiple perturbations:

- **Peptides 1-2** show an increase (+0.5) specifically in cond-1
- **Peptides 2-3** show a decrease (-0.75) in cond-2, with Pep2 affected in both cond-1 and cond-2
- **Peptides 3, 4, and 7** show an increase (+0.75) in cond-3, with Pep3 showing complex behavior across multiple conditions

This overlapping pattern creates a challenging clustering scenario where peptides may belong to multiple behavioral groups depending on the condition, resulting in at least 4 distinct clusters that reflect the intricate proteoform landscape often observed in real biological data.

```
nPeps = 8
nReps = 5

# Generate for control mean is 0 and std is 0.25
control_data = np.random.normal(loc=0, scale=0.05, size=(nPeps, nReps))
control_data
# Move the mean to 1 and std to 0.5
cond1_data = np.random.normal(loc=0.25, scale=0.15, size=(nPeps, nReps))
cond1_data
# Move the mean to 2 and std to 0.75
cond2_data = np.random.normal(loc=0.35, scale=0.05, size=(nPeps, nReps))
cond2_data
# Move the mean to 3 and std to 1
cond3_data = np.random.normal(loc=0.45, scale=0.1, size=(nPeps, nReps))
cond3_data

# put all the data together in a DataFrame
data = np.concatenate([control_data, cond1_data, cond2_data, cond3_data], axis=1)
data = pd.DataFrame(
    data,
    index=[f"Pep{i+1}" for i in range(nPeps)],
    columns=[
        "control_1", "control_2", "control_3", "control_4", "control_5",
        "cond-1_1", "cond-1_2", "cond-1_3", "cond-1_4", "cond-1_5",
        "cond-2_1", "cond-2_2", "cond-2_3", "cond-2_4", "cond-2_5",
        "cond-3_1", "cond-3_2", "cond-3_3", "cond-3_4", "cond-3_5"
    ]
)
# Adjust the data by control mean
data = data - data.loc[:, "control_1":"control_5"].mean(axis=1).values[:, None]

data = data.reset_index().melt(
    id_vars="index",
    var_name="Sample",
    value_name="Intensity"
)
data["Condition"] = data["Sample"].str.split("_", expand=True)[0]
# Introduce peptide moves
data.loc[
    ((data["Condition"] == "cond-1") & (data["index"].isin(["Pep1", "Pep2"]))), "Intensity"] += .5
data.loc[
    ((data["Condition"] == "cond-2") & (data["index"].isin(["Pep2", "Pep3"]))), "Intensity"] -= .75
data.loc[
    ((data["Condition"] == "cond-3") & (data["index"].isin(["Pep3", "Pep4", 'Pep7']))), "Intensity"] += .75

# Initialize the figure
fig, ax = plt.subplots(figsize=(6, 4))
sns.lineplot(
    ax=ax,
    data=data,
    x="index",
    y="Intensity",
    hue="Condition",
    palette=demo_palette,
    rasterized=True,
    # Add styling
    alpha=0.75,

    ## Markers
    marker="o",
    markersize=10,
    markeredgewidth=0,
    # markeredgecolor="black",
    ## Line style
    linewidth=1,
    linestyle="-",
    dashes=False,
    # errorbar
    err_style="bars",
    err_kws={"capsize": 5, "elinewidth": 2.5, "capthick": 2.5, "zorder": 0, "linewidth": 1.5},
)
ax.set_xlabel(f"Ordered Peptides (PeptideID)", fontsize=12)
ax.set_ylabel("Control Adjusted Intensity", fontsize=12)
ax.set_title(
    "",
    fontsize=16,
    fontweight="bold",
)
ax.grid("both", linestyle="--", linewidth=0.5, alpha=0.5, color="lightgrey")
plt.tight_layout()
# Save the figure
plots.finalize_plot(
    plt.gcf(),
    filename="Demo_Example3_PeptideConditionLineplot",
    show=True,
    save=save_to_folder,
    filepath=figure_path,
    formats=figure_formats, 
    transparent=transparent_bg,
    dpi=figure_dpi,
)
```

The line plot reveals the complex interplay of peptide behaviors across conditions. We observe that:

- **Pep1** shows elevated intensity in cond-1 but follows the general trend in other conditions
- **Pep2** displays a unique pattern with increased intensity in cond-1 and decreased intensity in cond-2
- **Pep3** exhibits the most complex behavior: decreased in cond-2 but sharply increased in cond-3
- **Pep4 and Pep7** show specific increases only in cond-3
- **Pep5, Pep6, and Pep8** maintain the baseline protein behavior across all conditions

This heterogeneous response pattern makes it challenging to assign peptides to simple binary groups (outlier vs. non-outlier). The overlapping behaviors suggest that traditional statistical approaches may miss these nuanced patterns, whereas ProteoForge's clustering approach can capture the multi-dimensional relationships between peptides.

---

Now moving on to the clustered heatmap, we visualize how these complex peptide behaviors translate into distance-based clustering:

```
# Create a mapping from integer to unique peptide index (e.g., 1: 'Pep1', 2: 'Pep2', ...)
unique_indices = data['index'].unique()
int_to_index = {i+1: idx for i, idx in enumerate(unique_indices)}
index_to_int = {v: k for k, v in int_to_index.items()}

data['PeptideID'] = data['index'].map(index_to_int)
data = data.sort_values(by=['PeptideID', 'Condition', 'Sample']).reset_index(drop=True)
df = data.groupby(["PeptideID", "Condition"])["Intensity"].mean().unstack(level="Condition")

distance_vector = pdist(df.values, metric='euclidean')

# squareform converts it into a full, symmetric distance matrix for easier interpretation
dist_corr = pd.DataFrame(squareform(distance_vector), index=df.index, columns=df.index)

dist_labels = np.array([1,2,3,4,1,1,4,1])

# Rescale the distance matrix to a 0-1 range for better comparability with correlation matrices
dist_min = dist_corr.values.min()
dist_max = dist_corr.values.max()
if dist_max > dist_min:
    scaled_distance_matrix = (dist_corr - dist_min) / (dist_max - dist_min)
else:
    scaled_distance_matrix = dist_corr.copy()

ax = plots.clustering_check_with_single_heatmap(
    corr_matrix=scaled_distance_matrix,
    labels=dist_labels,
    protein_id='Protein 1',
    vmin=0, vmax=1
)
# Set the axis title
ax.set_title("Distance Matrix with Clustering Check", fontsize=14)
# Cbar label
cbar = ax.collections[0].colorbar
cbar.set_label('Scaled Euclidean Distance', labelpad=15, fontsize=12)


# Increase label size
ax.set_xlabel("Peptide ID", fontsize=14)
ax.set_ylabel("Peptide ID", fontsize=14)

# Increase tick label size
ax.tick_params(axis='x', labelsize=12)
ax.tick_params(axis='y', labelsize=12)

plt.tight_layout()
plots.finalize_plot(
    plt.gcf(),
    filename="Demo_Example3_PeptideConditionCorrelation",
    show=True,
    save=save_to_folder,
    filepath=figure_path,
    formats=figure_formats, 
    transparent=transparent_bg,
    dpi=figure_dpi,
)
```

The heatmap confirms the presence of multiple distinct peptide behaviors resulting in four clusters:

- **Cluster 1 (C1)**: Peptides 1, 5, 6, and 8 — follow the baseline protein behavior
- **Cluster 2 (C2)**: Peptide 2 — unique pattern with effects in both cond-1 and cond-2
- **Cluster 3 (C3)**: Peptide 3 — complex behavior affected in cond-2 and cond-3
- **Cluster 4 (C4)**: Peptides 4 and 7 — specific increases in cond-3

This clustering effectively captures the intricate relationships among peptides, highlighting ProteoForge's capability to discern complex proteoform profiles in proteomics data. Based on the clustering combined with the linear model's outlier detection, peptides 2, 3, and 7 would be flagged as outliers (depending on the p-value threshold), with peptides 2 and 3 potentially indicating PTM/modified peptides, while peptides 4 and 7 display proteoform-like behavior that can be grouped together.

---

#### Conclusion¶

This notebook demonstrates the assembly of Figure 1 using simulated proteomics data. It showcases the data processing steps of ProteoForge and illustrates its ability to identify peptide-level outliers and cluster peptides based on their behavior across conditions. The three examples provided highlight progressively complex scenarios of peptide behavior:

1. **Example 1** — Simple case with a single perturbation in one condition (2 clusters)
2. **Example 2** — Intermediate complexity with independent perturbations in different conditions (3 clusters)
3. **Example 3** — Real-world complexity with overlapping, condition-dependent behaviors (4 clusters)

These demonstrations showcase the versatility and effectiveness of the ProteoForge analysis workflow in capturing both straightforward and intricate proteoform profiles.

> **Important:** This notebook is for demonstration purposes only. The figures generated here were used to build the overview figure of ProteoForge and do not represent real experimental data. The data is randomly generated each time you re-run the notebook, so exact values and plots will vary. In real scenarios, seed values would be used to ensure reproducibility.

```
print("Notebook Execution Time:", utils.prettyTimer(utils.getTime() - startTime))
```

```
Notebook Execution Time: 00h:00m:04s
```
