## Supplementary material for "ProteoForge: An Imputation-Aware Framework for Differential Proteoform Discovery in Bottom-Up Proteomics": Analysis notebooks as html render: Notebook S4.html

### Notebook S4

###### Enes Kemal Ergin

#### 2025-12-08 23:00:31

### Supplementary Notebook 4: Generating Simulation Datasets for Benchmarking¶

- **License:** Creative Commons Attribution-NonCommercial 4.0 International License
- **Version:** 0.2
- **Edit Log:**
  - 2025-11-28: Initial version of the notebook
  - 2025-12-08: Revise the whole notebook, ensuring clarity and correctness

---

**Requirements:**

This notebook doesn't need any data requirements as it generates synthetic datasets from scratch.

**Data Information:**  
This notebook generates synthetic proteomics datasets from scratch—no input files required. All simulated data is saved to `./data/Sim{1-4}/` folders as Feather files.

**Purpose:**  
Generate controlled synthetic proteomics datasets to benchmark proteoform detection methods (COPF, PeCorA, ProteoForge) under systematically varied conditions. Four simulation scenarios are created:

| Simulation | Focus | Variations |
| --- | --- | --- |
| **Sim1** | Imputation effects | 4 perturbation patterns × 2 data types (complete/imputed) |
| **Sim2** | Missingness tolerance | 5 protein × 5 peptide missingness levels = 25 combinations |
| **Sim3** | Detection sensitivity | 8 perturbation magnitude ranges |
| **Sim4** | Experimental complexity | 5 conditions × 2 overlap × 2 direction = 20 combinations |

---

#### Setup¶

This section imports libraries, configures display settings, and defines paths for outputs.

> **Note:** The HTML rendering of this notebook hides code cells by default. Click the "Code" buttons to expand them.

##### Libraries¶

```
import os
import sys
import warnings
warnings.filterwarnings("ignore")

import numpy as np # Numerical computing
import pandas as pd # Data manipulatio

import seaborn as sns # R-like high-level plots
import matplotlib.pyplot as plt # Python's base plotting 


sys.path.append('../')
# Utility imports for this analysis
from src import utils, plots

from Simulation import sims # Simulation functions 

# Initialize the timer
startTime = utils.getTime()
```

##### Display Settings¶

The cell below configures pandas, matplotlib, and seaborn display options for improved readability of tables and figures, including color palettes and figure export settings.

```
## Figure Settings

# Define default colors and styles for plots
def_colors = [
    "#139593", "#fca311", "#e54f2a",
    "#c3c3c3", "#555555",
    "#690000", "#5f4a00", "#004549"
]

# Set seaborn style
sns.set_theme(
    style="white",
    context="paper",
    palette=def_colors,
    font_scale=1,
    rc={
        "figure.figsize": (6, 4),
        "font.family": "sans-serif",
        "font.sans-serif": ["Arial", "Ubuntu Mono"],
    }
)

# Figure Saving Settings
figure_formats = ["pdf"]
save_to_folder = True
transparent_bg = True
figure_dpi = 300

## Configure dataframe displaying
pd.set_option('display.float_format', lambda x: '%.4f' % x)
pd.set_option('display.max_rows', 50)
pd.set_option('display.max_columns', 50)
pd.set_option('display.width', 500)  # Set a wider display width

## Printing Settings
verbose = True

## Reproducibility Settings
seed = 42

## Show the color-palettes
plots.color_palette( def_colors, save=False )
```

##### Paths and Global Variables¶

```
# Establish paths and 
notebook_name = "01-SimulatedDatasets"
input_path = f"./data/"
mainFig_path = f"./figures/"

def setup_simulation_paths( simID ):
    """
    Create output and figure directories for a simulation, if they do not exist. 
    (Uses global variable for save_to_folder and figure_formats)

    Args:
        simID (str): Simulation ID.
        
        (Global Variables)
            input_path (str): Base path for input data.
            mainFig_path (str): Base path for figures.
            save_to_folder (bool): Whether to save figures to folders.
            figure_formats (list): List of formats to save figures in.
    
    Returns:
        tuple: (output_path, figure_path)
    """
    output_path = f"{input_path}{simID}/"
    if not os.path.exists(output_path):
        os.makedirs(output_path)

    figure_path = f"{mainFig_path}{simID}/"
    if save_to_folder:
        for fmt in figure_formats:
            cur_folder = os.path.join(figure_path, fmt)
            if not os.path.exists(cur_folder):
                os.makedirs(cur_folder)
    return output_path, figure_path
```

---

#### Simulation 1: Complete vs. Imputed Data Comparison¶

**Objective:** Assess whether imputing missing values affects method performance compared to complete (fully quantified) datasets.

**Experimental Design:**

- **Base dataset**: 500 proteins × 3 conditions × 10 replicates, 5–50 peptides per protein
- **Perturbation**: 250 proteins perturbed in condition 2 with magnitude 0.5–1.5 log2
- **Four perturbation patterns tested**:
  1. **twoPep** — Exactly 2 peptides perturbed per protein
  2. **halfPep** — 50% of peptides perturbed per protein
  3. **halfPlusPep** — 70% of peptides perturbed per protein
  4. **randomPep** — Random 10–50% of peptides perturbed per protein
- **Data versions**: Each pattern generates both complete and imputed versions
- **Imputation**: Downshifted imputation with 35% missingness rate

**Output Files:** `./data/Sim1/2_{pattern}_{complete|imputed}_InputData.feather`

##### Base Dataset Generation¶

The code below generates the foundational protein/peptide data structure used across all Sim1 perturbation scenarios.

```
stTime = utils.getTime()

# --- Simulation Parameters ---
simID = "Sim1"
seed = 42  # Seed for reproducibility
n_condition = 3  # Number of conditions (1 is control, others are cond-N-1)
n_proteins = 500  # Number of proteins in the dataset
n_replicates = 10  # Number of replicates per condition
n_peptides = (5, 50)  # (min, max) for peptides per protein
condition_shifts = [0, 1, 2]  # Shifts for the conditions
nPro_to_perturb = 250  # Number of proteins to perturb
perturb_conds = ['cond2']  # Conditions to perturb
perturb_overlap = False  # Allow overlap between perturbed peptides
pertMag_range = (.5, 1.5)  # Range of values to perturb the peptides with (uniform)

# Generate the paths for it.
output_path, figure_path = setup_simulation_paths( simID )

print(f"\n{'='*50}")
print(f"{'Simulation ID:':<18} {simID}")
print(f"{'='*50}")
print("• Simulation Data:")
print(f"    - Generates a synthetic proteomics dataset with {n_proteins} proteins.")
print(f"    - Each protein has {n_peptides[0]}–{n_peptides[1]} peptides, across {n_condition} conditions.")
print(f"    - The first condition is the control; the rest are experimental.")
print("• Simulation Goal:")
print("    - Compare the effects of perturbations on the data, both in complete and imputed forms.")
print("• Parameters:")
print(f"    - RNG seed: {seed}")
print(f"    - {n_condition} conditions (control + {n_condition-1} treatments), log2 shifts: {condition_shifts}")
print(f"    - {n_proteins} proteins × {n_replicates} replicates, {n_peptides[0]}–{n_peptides[1]} peptides/protein")
print(f"    - Perturbing {nPro_to_perturb}/{n_proteins} proteins (overlap={perturb_overlap})")
print(f"    - Perturbation in conditions: {perturb_conds}, magnitude range: {pertMag_range}")
print(f"\n{'='*50}")
print("• Step 1: Generate Protein Mean Values (with outliers)")
print(f"    - Using seed: {seed}")
mean_values = sims.normal_distribution_with_outliers(
    mu=20,
    sd=2,
    size=n_proteins,
    is_log2=False,
    outlier_fraction=0.10,
    outlier_sd_multiplier=1.0,
    seed=seed
)
print(f"    - Generated {len(mean_values)} protein mean values (log2 scale)")
print(f"    - Stats: mean={np.mean(mean_values):.2f}, std={np.std(mean_values):.2f}, min={np.min(mean_values):.2f}, max={np.max(mean_values):.2f}")
print(f"    - Outliers: fraction=10.0%, sd-multiplier=1.0")
mean_values = 2 ** mean_values

print(f"\n{'-'*50}")
print("• Step 2: Generate Protein Coefficient of Variation (CV) Values")
cv_values = sims.lognormal_distribution(
    mu=10,
    med=8,
    size=n_proteins,
    seed=seed
)
print(f"    - Log-normal distribution: mean=10, median=8")
print(f"    - Generated {len(cv_values)} CV values")
print(f"    - Stats: mean={np.mean(cv_values):.2f}, std={np.std(cv_values):.2f}, min={np.min(cv_values):.2f}, max={np.max(cv_values):.2f}")

print(f"\n{'-'*50}")
print("• Step 3: Generate Replicates for Control (Biological Reference)")
control_data = sims.generate_replicates(
    mean_values,
    cv_values,
    meanScale='raw',
    cvType='percent',
    nReps=n_replicates,
    randomizeCV=True,
    as_dataframe=True,
    seed=seed
)
print(f"    - Generated {control_data.shape[1]} control replicates for {control_data.shape[0]} proteins")
print(f"    - This subset will serve as the control for all treatments.")

print(f"\n{'-'*50}")
print("• Step 4: Generate Number of Peptides per Protein")
pepN_cnts = sims.generate_peptide_counts(
    n_proteins=n_proteins,
    min_peptides=n_peptides[0],
    max_peptides=n_peptides[1],
    alpha=0.5,
    beta=3,
)
print(f"    - Beta-binomial: min={n_peptides[0]}, max={n_peptides[1]}, alpha=0.5, beta=3")
print(f"    - Generated {len(pepN_cnts)} peptide counts")

print(f"\n{'-'*50}")
print("• Step 5: Generate Peptide-Level Data from Protein-Level Data")
pep_data = sims.generate_peptide_level(
    control_data,
    pepN_cnts,
    is_log2=False,
    repStd=(0.1, 0.25),
    outlier_fraction=0.0001,
    outlier_multiplier=0.01,
    add_noise=True,
    noise_sd=0.10,
    seed=seed
)
print(f"    - Peptide-level data: {pep_data.shape[0]} peptides × {pep_data.shape[1]} samples")
print(f"    - Noise: sd=0.10, Outliers: fraction=0.0001, multiplier=0.01")

print(f"\n{'-'*50}")
print("• Step 6: Generate Condition Mappers")
condition_sample_map, condition_shifts = sims.generate_condition_mappers(
    n_condition=n_condition-1,
    n_replicates=n_replicates,
    condition_shifts=condition_shifts[1:],
    control_name='control',
    condition_suffix='cond',
)
print(f"    - Condition sample map keys: {list(condition_sample_map.keys())}")
print(f"    - Condition shifts: {condition_shifts}")

print(f"\n{'-'*50}")
print("• Step 7: Generate Complete Peptide-Level Data for All Conditions")
generated_data = sims.generate_complete_data(
    pep_data,
    condition_shifts=condition_shifts,
    condition_sample_map=condition_sample_map,
    shift_scale=0.10,
    is_log2=False,
    add_noise=False,
    noise_sd=0.10,
    seed=seed
)
print(f"    - Complete dataset: {generated_data.shape[0]} peptides × {generated_data.shape[1]} samples")
print(f"    - Noise: sd=0.10, shift_scale=0.10, no additional noise added")
print(f"{'='*50}\n")

# generated_data.to_feather(f"{output_path}1_GeneratedData.feather")
```

```
==================================================
Simulation ID:     Sim1
==================================================
• Simulation Data:
    - Generates a synthetic proteomics dataset with 500 proteins.
    - Each protein has 5–50 peptides, across 3 conditions.
    - The first condition is the control; the rest are experimental.
• Simulation Goal:
    - Compare the effects of perturbations on the data, both in complete and imputed forms.
• Parameters:
    - RNG seed: 42
    - 3 conditions (control + 2 treatments), log2 shifts: [0, 1, 2]
    - 500 proteins × 10 replicates, 5–50 peptides/protein
    - Perturbing 250/500 proteins (overlap=False)
    - Perturbation in conditions: ['cond2'], magnitude range: (0.5, 1.5)

==================================================
• Step 1: Generate Protein Mean Values (with outliers)
    - Using seed: 42
    - Generated 500 protein mean values (log2 scale)
    - Stats: mean=20.01, std=1.96, min=13.52, max=27.71
    - Outliers: fraction=10.0%, sd-multiplier=1.0

--------------------------------------------------
• Step 2: Generate Protein Coefficient of Variation (CV) Values
    - Log-normal distribution: mean=10, median=8
    - Generated 500 CV values
    - Stats: mean=10.07, std=8.36, min=0.92, max=104.93

--------------------------------------------------
• Step 3: Generate Replicates for Control (Biological Reference)
    - Generated 10 control replicates for 500 proteins
    - This subset will serve as the control for all treatments.

--------------------------------------------------
• Step 4: Generate Number of Peptides per Protein
    - Beta-binomial: min=5, max=50, alpha=0.5, beta=3
    - Generated 500 peptide counts

--------------------------------------------------
• Step 5: Generate Peptide-Level Data from Protein-Level Data
    - Peptide-level data: 5583 peptides × 12 samples
    - Noise: sd=0.10, Outliers: fraction=0.0001, multiplier=0.01

--------------------------------------------------
• Step 6: Generate Condition Mappers
    - Condition sample map keys: ['control', 'cond1', 'cond2']
    - Condition shifts: {'control': 0, 'cond1': 1, 'cond2': 2}

--------------------------------------------------
• Step 7: Generate Complete Peptide-Level Data for All Conditions
    - Complete dataset: 5583 peptides × 30 samples
    - Noise: sd=0.10, shift_scale=0.10, no additional noise added
==================================================
```

##### Perturbation Pattern 1: Two Peptides (twoPep)¶

Perturbs exactly **2 peptides** per protein in the same direction. This scenario tests detection of minimal proteoform changes—the smallest signal that might represent a post-translational modification affecting a localized region.

```
print("Two Peptides Perturbation Setup")
perturb_name = "twoPep"  # Perturbation scenario name
nPep_to_perturb = 2  # Number of peptides to perturb; if > 1 then it is a fraction
perturb_dir_setup = "same"  # Direction of perturbation: [random, same]
np.random.seed(seed)  # Set the seed for reproducibility

print(f"\n{'='*50}")
print("• Step 1: Perturbation Setup")
simulated_data = generated_data.copy()
print(f"    - Perturbing {nPro_to_perturb}/{n_proteins} proteins with {nPep_to_perturb} peptides each")
print(f"    - Conditions: {perturb_conds}, Overlap: {perturb_overlap}, Magnitude range: {pertMag_range}")

unique_proteins = simulated_data.reset_index()["Protein"].unique()
proteins_to_perturb = np.random.choice(unique_proteins, nPro_to_perturb, replace=False)
print(f"    - Proteins selected for perturbation: {proteins_to_perturb[:5]} ... (total: {len(proteins_to_perturb)})")


# Subset the data for perturbation
tmp = np.log2(simulated_data).copy()
perturb_data = tmp.loc[proteins_to_perturb]
unchanged_data = tmp.drop(proteins_to_perturb)

perturb_values = sims.generate_perturb_values(
    nPro_to_perturb=nPro_to_perturb,
    nCond=nPep_to_perturb,
    pertMag_range=pertMag_range,
    direction="same",
)

# Find the condition for each protein
perturb_conditions = np.random.choice(perturb_conds, nPro_to_perturb)

# Go through the proteins and perturb single peptide in each
dictCnt = 0
perturbation_map = {}
print("    - Applying perturbations to selected proteins and peptides...")
for i, protein in enumerate(proteins_to_perturb):
    # Get the current protein data
    cur_data = perturb_data.loc[protein]
    # Get the perturbation value
    perturb_value = perturb_values[i]
    # Select n number of non-consecutive peptides to perturb
    pep_pos = np.random.choice(cur_data.index, nPep_to_perturb, replace=False)
    cond = perturb_conditions[i]
    # Get the perturbation condition
    perturb_samples = condition_sample_map[cond]
    for j in range(nPep_to_perturb):
        # Get the peptides to perturb for the current condition
        perturb_peptide = pep_pos[j]
        # Get the perturbation peptide intensity
        perturb_intensity = cur_data.loc[perturb_peptide, perturb_samples]
        # Update the perturbation peptide intensity
        perturb_intensity += perturb_value[j]
        perturb_data.loc[(protein, perturb_peptide), perturb_samples] = perturb_intensity
        perturbation_map[dictCnt] = {
            "Protein": protein,
            "Peptide": perturb_peptide,
            "pertCondition": cond,
            "pertShift": perturb_value[j],
        }
        dictCnt += 1

print(f"    - Total perturbations applied: {dictCnt}")

perturbed_data = pd.concat([perturb_data, unchanged_data], axis=0)
perturbed_data = (np.power(2, perturbed_data))

print(f"    - Perturbed data shape: {perturbed_data.shape}")

# --- Build Test Data Object ---
print("    - Building test data object with perturbation map...")
test_data = sims.build_test_data(
    data=perturbed_data,
    condition_sample_map=condition_sample_map,
    perturbation_map=perturbation_map,
    proteins_to_perturb=proteins_to_perturb,
    missing_data=None,
)
test_data.to_feather(f"{output_path}2_{perturb_name}_complete_InputData.feather")
complete_data = test_data.copy()

print(f"\n{'='*50}")
print("• Step 2: Imputation Setup")

# --- Simulate Missing Data (Amputation) ---
print("    - Simulating missing data (amputation)...")
missing_data = sims.amputation(
    data=perturbed_data,
    unique_proteins=unique_proteins,
    proteins_to_perturb=proteins_to_perturb,
    condition_shifts=condition_shifts,
    condition_sample_map=condition_sample_map,
    n_amputate_1=100,
    n_amputate_2=100,
    n_amputate_3=100,
    missing_rate=0.35,
    seed=seed
)
print(f"    - Introduced missing data with rate=35.0%")

# --- Impute Missing Data ---
print("    - Imputing missing data using downshifted imputation...")
imputed_data = sims.downshifted_imputation(
    data=missing_data,
    condition_sample_map=condition_sample_map,
    is_log2=False,
    shiftMag=2,
    lowPct=0.15,
    minValue=8,
    impute_all=False,
    seed=seed
)
print(f"    - Imputed missing data with downshift=2.0 and lowPct=0.15")

# --- Build Final Test Data Object ---
print("    - Building final test data object with imputed data...")
test_data = sims.build_test_data(
    data=imputed_data,
    condition_sample_map=condition_sample_map,
    perturbation_map=perturbation_map,
    proteins_to_perturb=proteins_to_perturb,
    missing_data=missing_data,
)
test_data.to_feather(f"{output_path}2_{perturb_name}_imputed_InputData.feather")
imputed_data = test_data.copy()
print(f"{'='*50}\n")
```

```
Two Peptides Perturbation Setup

==================================================
• Step 1: Perturbation Setup
    - Perturbing 250/500 proteins with 2 peptides each
    - Conditions: ['cond2'], Overlap: False, Magnitude range: (0.5, 1.5)
    - Proteins selected for perturbation: ['pro_362' 'pro_74' 'pro_375' 'pro_156' 'pro_105'] ... (total: 250)
    - Applying perturbations to selected proteins and peptides...
    - Total perturbations applied: 500
    - Perturbed data shape: (5583, 30)
    - Building test data object with perturbation map...

==================================================
• Step 2: Imputation Setup
    - Simulating missing data (amputation)...
    - Introduced missing data with rate=35.0%
    - Imputing missing data using downshifted imputation...
    - Imputed missing data with downshift=2.0 and lowPct=0.15
    - Building final test data object with imputed data...
==================================================
```

##### Perturbation Pattern 2: Half Peptides (halfPep)¶

Perturbs **50%** of peptides per protein. This simulates a scenario where half the protein is affected—potentially representing domain-specific changes or partial degradation.

```
print("Half of the Peptides (50%) Perturbation Setup")
perturb_name = "halfPep"  # Perturbation scenario name
nPep_to_perturb = 0.50  # Number of peptides to perturb; if > 1 then it is a fraction
perturb_dir_setup = "same"  # Direction of perturbation: [random, same]
np.random.seed(seed)  # Set the seed for reproducibility

print(f"\n{'='*50}")
print("• Step 1: Perturbation Setup")
simulated_data = generated_data.copy()
print(f"    - Perturbing {nPro_to_perturb}/{n_proteins} proteins with {nPep_to_perturb*100:.0f}% peptides each")
print(f"    - Conditions: {perturb_conds}, Overlap: {perturb_overlap}, Magnitude range: {pertMag_range}")
unique_proteins = simulated_data.reset_index()["Protein"].unique()
proteins_to_perturb = np.random.choice(unique_proteins, nPro_to_perturb, replace=False)
print(f"    - Proteins selected for perturbation: {proteins_to_perturb[:5]} ... (total: {len(proteins_to_perturb)})")

# Subset the data for perturbation
tmp = np.log2(simulated_data).copy()
perturb_data = tmp.loc[proteins_to_perturb]
unchanged_data = tmp.drop(proteins_to_perturb)

# Go through the proteins and perturb single peptide in each
dictCnt = 0
perturbation_map = {}
print("    - Applying perturbations to selected proteins and peptides...")
for i, protein in enumerate(proteins_to_perturb):
    cur_data = perturb_data.loc[protein]
    n = int( np.floor(nPep_to_perturb * len(cur_data) ))
    if n < 2: n = 2
    # print(f"Protein: {protein} - Peptides: {n}, Total Peptides: {len(cur_data)}")
    pepNums = np.random.choice(cur_data.index, n, replace=False)
    # Identify the conditions to perturb
    conds = np.random.choice(perturb_conds, len(pepNums), replace=True)
    # Identify the magnitude of perturbation
    pertMag = np.random.uniform(pertMag_range[0], pertMag_range[1], len(pepNums))
    #
    if perturb_dir_setup == "random":
        pertDir = np.random.choice([-1, 1], len(pepNums), replace=True)
    elif perturb_dir_setup == "same":
        # Pick random direction
        pertDir = np.random.choice([-1, 1], 1, replace=True)
        pertDir = np.repeat(pertDir, len(pepNums))
    
    for j in range(len(pepNums)):
        perturb_samples = condition_sample_map[conds[j]]
        # Get the peptides to perturb for the current condition
        perturb_peptide = pepNums[j]
        # Get the perturbation peptide intensity
        perturb_intensity = cur_data.loc[perturb_peptide, perturb_samples]
        # Update the perturbation peptide intensity
        perturb_intensity += pertMag[j] * pertDir[j]
        perturb_data.loc[(protein, perturb_peptide), perturb_samples] = perturb_intensity
        perturbation_map[dictCnt] = {
            "Protein": protein,
            "Peptide": perturb_peptide,
            "pertCondition": conds[j],
            "pertShift": pertMag[j] * pertDir[j],
        }
        dictCnt += 1

print(f"    - Total perturbations applied: {dictCnt}")

perturbed_data = pd.concat([perturb_data, unchanged_data], axis=0)
perturbed_data = (np.power(2, perturbed_data))
print(f"    - Perturbed data shape: {perturbed_data.shape}")

# --- Build Test Data Object ---
print("    - Building test data object with perturbation map...")
test_data = sims.build_test_data(
    data=perturbed_data,
    condition_sample_map=condition_sample_map,
    perturbation_map=perturbation_map,
    proteins_to_perturb=proteins_to_perturb,
    missing_data=None,
)
test_data.to_feather(f"{output_path}2_{perturb_name}_complete_InputData.feather")
complete_data = test_data.copy()

print(f"\n{'='*50}")
print("• Step 2: Imputation Setup")

# --- Simulate Missing Data (Amputation) ---
print("    - Simulating missing data (amputation)...")
missing_data = sims.amputation(
    data=perturbed_data,
    unique_proteins=unique_proteins,
    proteins_to_perturb=proteins_to_perturb,
    condition_shifts=condition_shifts,
    condition_sample_map=condition_sample_map,
    n_amputate_1=100,
    n_amputate_2=100,
    n_amputate_3=100,
    missing_rate=0.35,
    seed=seed
)
print(f"    - Introduced missing data with rate=35.0%")

# --- Impute Missing Data ---
print("    - Imputing missing data using downshifted imputation...")
imputed_data = sims.downshifted_imputation(
    data=missing_data,
    condition_sample_map=condition_sample_map,
    is_log2=False,
    shiftMag=2,
    lowPct=0.15,
    minValue=8,
    impute_all=False,
    seed=seed
)
print(f"    - Imputed missing data with downshift=2.0 and lowPct=0.15")

# --- Build Final Test Data Object ---
print("    - Building final test data object with imputed data...")
test_data = sims.build_test_data(
    data=imputed_data,
    condition_sample_map=condition_sample_map,
    perturbation_map=perturbation_map,
    proteins_to_perturb=proteins_to_perturb,
    missing_data=missing_data,
)
test_data.to_feather(f"{output_path}2_{perturb_name}_imputed_InputData.feather")
imputed_data = test_data.copy()
print(f"{'='*50}\n")
```

```
Half of the Peptides (50%) Perturbation Setup

==================================================
• Step 1: Perturbation Setup
    - Perturbing 250/500 proteins with 50% peptides each
    - Conditions: ['cond2'], Overlap: False, Magnitude range: (0.5, 1.5)
    - Proteins selected for perturbation: ['pro_362' 'pro_74' 'pro_375' 'pro_156' 'pro_105'] ... (total: 250)
    - Applying perturbations to selected proteins and peptides...
    - Total perturbations applied: 1343
    - Perturbed data shape: (5583, 30)
    - Building test data object with perturbation map...

==================================================
• Step 2: Imputation Setup
    - Simulating missing data (amputation)...
    - Introduced missing data with rate=35.0%
    - Imputing missing data using downshifted imputation...
    - Imputed missing data with downshift=2.0 and lowPct=0.15
    - Building final test data object with imputed data...
==================================================
```

##### Perturbation Pattern 3: Majority Peptides (halfPlusPep)¶

Perturbs **70%** of peptides per protein. Tests near-complete protein-level changes where only a minority of peptides remain unaffected—approaching traditional differential expression scenarios.

```
print("More than Half of the Peptides (70%) Perturbation Setup")
perturb_name = "halfPlusPep"  # Perturbation scenario name
nPep_to_perturb = 0.70  # Number of peptides to perturb; if > 1 then it is a fraction
perturb_dir_setup = "same"  # Direction of perturbation: [random, same]
np.random.seed(seed)  # Set the seed for reproducibility
print(f"\n{'='*50}")
print("• Step 1: Perturbation Setup")
simulated_data = generated_data.copy()
print(f"    - Perturbing {nPro_to_perturb}/{n_proteins} proteins with {nPep_to_perturb*100:.0f}% peptides each")
print(f"    - Conditions: {perturb_conds}, Overlap: {perturb_overlap}, Magnitude range: {pertMag_range}")
unique_proteins = simulated_data.reset_index()["Protein"].unique()
proteins_to_perturb = np.random.choice(unique_proteins, nPro_to_perturb, replace=False)
print(f"    - Proteins selected for perturbation: {proteins_to_perturb[:5]} ... (total: {len(proteins_to_perturb)})")   

# Subset the data for perturbation
tmp = np.log2(simulated_data).copy()
perturb_data = tmp.loc[proteins_to_perturb]
unchanged_data = tmp.drop(proteins_to_perturb)

# Go through the proteins and perturb single peptide in each
dictCnt = 0
perturbation_map = {}
print("    - Applying perturbations to selected proteins and peptides...")
for i, protein in enumerate(proteins_to_perturb):
    cur_data = perturb_data.loc[protein]
    n = int( np.ceil(nPep_to_perturb * len(cur_data) ))
    if n < 2: n = 2
    # print(f"Protein: {protein} - Peptides: {n}, Total Peptides: {len(cur_data)}")
    pepNums = np.random.choice(cur_data.index, n, replace=False)
    # Identify the conditions to perturb
    conds = np.random.choice(perturb_conds, len(pepNums), replace=True)
    # Identify the magnitude of perturbation
    pertMag = np.random.uniform(pertMag_range[0], pertMag_range[1], len(pepNums))
    #
    if perturb_dir_setup == "random":
        pertDir = np.random.choice([-1, 1], len(pepNums), replace=True)
    elif perturb_dir_setup == "same":
        # Pick random direction
        pertDir = np.random.choice([-1, 1], 1, replace=True)
        pertDir = np.repeat(pertDir, len(pepNums))
    
    for j in range(len(pepNums)):
        perturb_samples = condition_sample_map[conds[j]]
        # Get the peptides to perturb for the current condition
        perturb_peptide = pepNums[j]
        # Get the perturbation peptide intensity
        perturb_intensity = cur_data.loc[perturb_peptide, perturb_samples]
        # Update the perturbation peptide intensity
        perturb_intensity += pertMag[j] * pertDir[j]
        perturb_data.loc[(protein, perturb_peptide), perturb_samples] = perturb_intensity
        perturbation_map[dictCnt] = {
            "Protein": protein,
            "Peptide": perturb_peptide,
            "pertCondition": conds[j],
            "pertShift": pertMag[j] * pertDir[j],
        }
        dictCnt += 1

print(f"    - Total perturbations applied: {dictCnt}")

# --- Combine Perturbed and Unchanged Data ---
perturbed_data = pd.concat([perturb_data, unchanged_data], axis=0)
perturbed_data = (np.power(2, perturbed_data))
print(f"    - Perturbed data shape: {perturbed_data.shape}")

# --- Build Test Data Object ---
print("    - Building test data object with perturbation map...")
test_data = sims.build_test_data(
    data=perturbed_data,
    condition_sample_map=condition_sample_map,
    perturbation_map=perturbation_map,
    proteins_to_perturb=proteins_to_perturb,
    missing_data=None,
)
test_data.to_feather(f"{output_path}2_{perturb_name}_complete_InputData.feather")
complete_data = test_data.copy()

print(f"\n{'='*50}")
print("• Step 2: Imputation Setup")

# --- Simulate Missing Data (Amputation) ---
print("    - Simulating missing data (amputation)...")
missing_data = sims.amputation(
    data=perturbed_data,
    unique_proteins=unique_proteins,
    proteins_to_perturb=proteins_to_perturb,
    condition_shifts=condition_shifts,
    condition_sample_map=condition_sample_map,
    n_amputate_1=100,
    n_amputate_2=100,
    n_amputate_3=100,
    missing_rate=0.35,
    seed=seed
)
print(f"    - Introduced missing data with rate=35.0%")

# --- Impute Missing Data ---
print("    - Imputing missing data using downshifted imputation...")
imputed_data = sims.downshifted_imputation(
    data=missing_data,
    condition_sample_map=condition_sample_map,
    is_log2=False,
    shiftMag=2,
    lowPct=0.15,
    minValue=8,
    impute_all=False,
    seed=seed
)
print(f"    - Imputed missing data with downshift=2.0 and lowPct=0.15")

# --- Build Final Test Data Object ---
print("    - Building final test data object with imputed data...")
test_data = sims.build_test_data(
    data=imputed_data,
    condition_sample_map=condition_sample_map,
    perturbation_map=perturbation_map,
    proteins_to_perturb=proteins_to_perturb,
    missing_data=missing_data,
)
test_data.to_feather(f"{output_path}2_{perturb_name}_imputed_InputData.feather")
imputed_data = test_data.copy()
print(f"{'='*50}\n")
```

```
More than Half of the Peptides (70%) Perturbation Setup

==================================================
• Step 1: Perturbation Setup
    - Perturbing 250/500 proteins with 70% peptides each
    - Conditions: ['cond2'], Overlap: False, Magnitude range: (0.5, 1.5)
    - Proteins selected for perturbation: ['pro_362' 'pro_74' 'pro_375' 'pro_156' 'pro_105'] ... (total: 250)
    - Applying perturbations to selected proteins and peptides...
    - Total perturbations applied: 2104
    - Perturbed data shape: (5583, 30)
    - Building test data object with perturbation map...

==================================================
• Step 2: Imputation Setup
    - Simulating missing data (amputation)...
    - Introduced missing data with rate=35.0%
    - Imputing missing data using downshifted imputation...
    - Imputed missing data with downshift=2.0 and lowPct=0.15
    - Building final test data object with imputed data...
==================================================
```

##### Perturbation Pattern 4: Random Peptides (randomPep)¶

Perturbs a **random 10–50%** of peptides per protein (minimum 2). This is the most realistic scenario, simulating the variable nature of biological perturbations where the extent of change varies across proteins.

```
print("Random Number of Peptides (2 to 50%) Perturbation Setup")
perturb_name = "randomPep"  # Perturbation scenario name
nPep_to_perturb = -1  # Number of peptides to perturb; if > 1 then it is a fraction
perturb_dir_setup = "same"  # Direction of perturbation: [random, same]
np.random.seed(seed)  # Set the seed for reproducibility

print(f"\n{'='*50}")
print("• Step 1: Perturbation Setup")
simulated_data = generated_data.copy()
print(f"    - Perturbing {nPro_to_perturb}/{n_proteins} proteins with {nPep_to_perturb} peptides each")
print(f"    - Conditions: {perturb_conds}, Overlap: {perturb_overlap}, Magnitude range: {pertMag_range}")

# Get the unique proteins
unique_proteins = simulated_data.reset_index()["Protein"].unique()
# Select random proteins to perturb
proteins_to_perturb = np.random.choice(unique_proteins, nPro_to_perturb, replace=False)
print(f"    - Proteins selected for perturbation: {proteins_to_perturb[:5]} ... (total: {len(proteins_to_perturb)})")
# Pick a fraction between 0.1 and 0.5 of the peptides to perturb
if nPep_to_perturb == -1:
    nPep_array = np.random.uniform(0.1, 0.5, len(unique_proteins))
elif nPep_to_perturb < 1 and nPep_to_perturb > 0:
    nPep_array = np.repeat(nPep_to_perturb, len(unique_proteins))
elif nPep_to_perturb >= 1:
    nPep_array = np.repeat(nPep_to_perturb, len(unique_proteins))

# --- Subset Data for Perturbation ---
tmp = np.log2(simulated_data).copy()
perturb_data = tmp.loc[proteins_to_perturb]
unchanged_data = tmp.drop(proteins_to_perturb)

# --- Apply Perturbations to Selected Proteins and Peptides ---
dictCnt = 0
perturbation_map = {}
print("    - Applying perturbations to selected proteins and peptides...")
for i, protein in enumerate(proteins_to_perturb):
    cur_data = perturb_data.loc[protein]
    if nPep_to_perturb < 1:
        n = int(nPep_array[i] * len(cur_data))
        # Ensure minimum of 1 peptide is perturbed
        if n < 2: n = 2
    elif nPep_to_perturb >= 1:
        n = int(nPep_to_perturb)
    # print(f"Protein: {protein} - Peptides: {n}, Total Peptides: {len(cur_data)}")
    pepNums = np.random.choice(cur_data.index, n, replace=False)
    # Identify the conditions to perturb
    conds = np.random.choice(perturb_conds, len(pepNums), replace=True)
    # Identify the magnitude of perturbation
    pertMag = np.random.uniform(pertMag_range[0], pertMag_range[1], len(pepNums))
    #
    if perturb_dir_setup == "random":
        pertDir = np.random.choice([-1, 1], len(pepNums), replace=True)
    elif perturb_dir_setup == "same":
        # Pick random direction
        pertDir = np.random.choice([-1, 1], 1, replace=True)
        pertDir = np.repeat(pertDir, len(pepNums))
    
    for j in range(len(pepNums)):
        perturb_samples = condition_sample_map[conds[j]]
        # Get the peptides to perturb for the current condition
        perturb_peptide = pepNums[j]
        # Get the perturbation peptide intensity
        perturb_intensity = cur_data.loc[perturb_peptide, perturb_samples]
        # Update the perturbation peptide intensity
        perturb_intensity += pertMag[j] * pertDir[j]
        perturb_data.loc[(protein, perturb_peptide), perturb_samples] = perturb_intensity
        perturbation_map[dictCnt] = {
            "Protein": protein,
            "Peptide": perturb_peptide,
            "pertCondition": conds[j],
            "pertShift": pertMag[j] * pertDir[j],
        }
        dictCnt += 1
print(f"    - Total perturbations applied: {dictCnt}")

perturbed_data = pd.concat([perturb_data, unchanged_data], axis=0)
perturbed_data = (np.power(2, perturbed_data))
print(f"    - Perturbed data shape: {perturbed_data.shape}")

# --- Build Test Data Object ---
print("    - Building test data object with perturbation map...")
test_data = sims.build_test_data(
    data=perturbed_data,
    condition_sample_map=condition_sample_map,
    perturbation_map=perturbation_map,
    proteins_to_perturb=proteins_to_perturb,
    missing_data=None,
)
test_data.to_feather(f"{output_path}2_{perturb_name}_complete_InputData.feather")
complete_data = test_data.copy()

print(f"\n{'='*50}")
print("• Step 2: Imputation Setup")

# --- Simulate Missing Data (Amputation) ---
print("    - Simulating missing data (amputation)...")
missing_data = sims.amputation(
    data=perturbed_data,
    unique_proteins=unique_proteins,
    proteins_to_perturb=proteins_to_perturb,
    condition_shifts=condition_shifts,
    condition_sample_map=condition_sample_map,
    n_amputate_1=100,
    n_amputate_2=100,
    n_amputate_3=100,
    missing_rate=0.35,
    seed=seed
)
print(f"    - Introduced missing data with rate=35.0%")

# --- Impute Missing Data ---
print("    - Imputing missing data using downshifted imputation...")
imputed_data = sims.downshifted_imputation(
    data=missing_data,
    condition_sample_map=condition_sample_map,
    is_log2=False,
    shiftMag = 2,
    lowPct = 0.15,
    minValue = 8,
    impute_all=False,
    seed=seed
)
print(f"    - Imputed missing data with downshift=2.5 and lowPct=0.10")

# --- Build Final Test Data Object ---
print("    - Building final test data object with imputed data...")
test_data = sims.build_test_data(
    data=imputed_data,
    condition_sample_map=condition_sample_map,
    perturbation_map=perturbation_map,
    proteins_to_perturb=proteins_to_perturb,
    missing_data=missing_data,
)
test_data.to_feather(f"{output_path}2_{perturb_name}_imputed_InputData.feather")
imputed_data = test_data.copy()
print(f"{'='*50}\n")
```

```
Random Number of Peptides (2 to 50%) Perturbation Setup

==================================================
• Step 1: Perturbation Setup
    - Perturbing 250/500 proteins with -1 peptides each
    - Conditions: ['cond2'], Overlap: False, Magnitude range: (0.5, 1.5)
    - Proteins selected for perturbation: ['pro_362' 'pro_74' 'pro_375' 'pro_156' 'pro_105'] ... (total: 250)
    - Applying perturbations to selected proteins and peptides...
    - Total perturbations applied: 835
    - Perturbed data shape: (5583, 30)
    - Building test data object with perturbation map...

==================================================
• Step 2: Imputation Setup
    - Simulating missing data (amputation)...
    - Introduced missing data with rate=35.0%
    - Imputing missing data using downshifted imputation...
    - Imputed missing data with downshift=2.5 and lowPct=0.10
    - Building final test data object with imputed data...
==================================================
```

```
nVersions = 4 * 2  # 4 perturbation types, each with complete and imputed data
elapsed = utils.prettyTimer(utils.getTime() - stTime)
print(f"\n{'-'*50}")
print(f"• Completed generation of {nVersions} versions for simulation: {simID}")
print(f"    - Elapsed time: {elapsed}")
print(f"{'-'*50}\n")
```

```
--------------------------------------------------
• Completed generation of 8 versions for simulation: Sim1
    - Elapsed time: 00h:00m:09s
--------------------------------------------------
```

---

#### Simulation 2: Missingness Level Impact¶

**Objective:** Quantify how increasing proportions of missing data affect detection performance after imputation.

**Experimental Design:**

- **Base dataset**: Same structure as Sim1 (500 proteins, 3 conditions, 10 replicates)
- **Perturbation**: Random 10–50% pattern (same as randomPep from Sim1)
- **Missingness levels**: Factorial design with 5 × 5 = 25 combinations
  - Protein-level: 0%, 20%, 40%, 60%, 80% of proteins have missing values
  - Peptide-level: 0%, 20%, 40%, 60%, 80% of peptides within affected proteins are missing
- **Imputation**: Downshifted imputation applied uniformly

**Key Question:** At what missingness threshold does performance critically degrade?

**Output Files:** `./data/Sim2/2_Pro{rate}_Pep{rate}_imputed_InputData.feather`

##### Base Dataset and Perturbation Setup¶

```
stTime = utils.getTime()

# --- Simulation Parameters ---
simID = "Sim2"
seed = 42  # Seed for reproducibility
n_condition = 3  # Number of conditions (1 is control, others are cond-N-1)
n_proteins = 500  # Number of proteins in the dataset
n_replicates = 10  # Number of replicates per condition
n_peptides = (5, 50)  # (min, max) for peptides per protein
condition_shifts = [0, 1, 2]  # Shifts for the conditions
nPro_to_perturb = 250  # Number of proteins to perturb
perturb_conds = ['cond2']  # Conditions to perturb
perturb_overlap = False  # Allow overlap between perturbed peptides
pertMag_range = (.5, 1.5)  # Range of values to perturb the peptides with (uniform)

# Generate the paths for it.
output_path, figure_path = setup_simulation_paths( simID )

print(f"\n{'='*50}")
print(f"{'Simulation ID:':<18} {simID}")
print(f"{'='*50}")
print("• Simulation Data:")
print(f"    - Generates a synthetic proteomics dataset with {n_proteins} proteins.")
print(f"    - Each protein has {n_peptides[0]}–{n_peptides[1]} peptides, across {n_condition} conditions.")
print(f"    - The first condition is the control; the rest are experimental.")
print(f"    - Uses the random (2 to 50%) peptide perturbation strategy as the base and adds different levels of missingness.")
print("• Simulation Goal:")
print("    - Compare the effects of of missingness rates in protein and peptide levels with imputation.")
print("• Parameters:")
print(f"    - RNG seed: {seed}")
print(f"    - {n_condition} conditions (control + {n_condition-1} treatments), log2 shifts: {condition_shifts}")
print(f"    - {n_proteins} proteins × {n_replicates} replicates, {n_peptides[0]}–{n_peptides[1]} peptides/protein")
print(f"    - Perturbing {nPro_to_perturb}/{n_proteins} proteins (overlap={perturb_overlap})")
print(f"    - Perturbation in conditions: {perturb_conds}, magnitude range: {pertMag_range}")
print(f"{'='*50}\n")
print("• Step 1: Generate Protein Mean Values (with outliers)")
print(f"    - Using seed: {seed}")
mean_values = sims.normal_distribution_with_outliers(
    mu=20,
    sd=2,
    size=n_proteins,
    is_log2=False,
    outlier_fraction=0.10,
    outlier_sd_multiplier=1.0,
    seed=seed
)
print(f"    - Generated {len(mean_values)} protein mean values (log2 scale)")
print(f"    - Stats: mean={np.mean(mean_values):.2f}, std={np.std(mean_values):.2f}, min={np.min(mean_values):.2f}, max={np.max(mean_values):.2f}")
print(f"    - Outliers: fraction=10.0%, sd-multiplier=1.0")
mean_values = 2 ** mean_values

print(f"\n{'-'*50}")
print("• Step 2: Generate Protein Coefficient of Variation (CV) Values")
cv_values = sims.lognormal_distribution(
    mu=10,
    med=8,
    size=n_proteins,
    seed=seed
)
print(f"    - Log-normal distribution: mean=10, median=8")
print(f"    - Generated {len(cv_values)} CV values")
print(f"    - Stats: mean={np.mean(cv_values):.2f}, std={np.std(cv_values):.2f}, min={np.min(cv_values):.2f}, max={np.max(cv_values):.2f}")

print(f"\n{'-'*50}")
print("• Step 3: Generate Replicates for Control (Biological Reference)")
control_data = sims.generate_replicates(
    mean_values,
    cv_values,
    meanScale='raw',
    cvType='percent',
    nReps=n_replicates,
    randomizeCV=True,
    as_dataframe=True,
    seed=seed
)
print(f"    - Generated {control_data.shape[1]} control replicates for {control_data.shape[0]} proteins")
print(f"    - This subset will serve as the control for all treatments.")

print(f"\n{'-'*50}")
print("• Step 4: Generate Number of Peptides per Protein")
pepN_cnts = sims.generate_peptide_counts(
    n_proteins=n_proteins,
    min_peptides=n_peptides[0],
    max_peptides=n_peptides[1],
    alpha=0.5,
    beta=3,
)
print(f"    - Beta-binomial: min={n_peptides[0]}, max={n_peptides[1]}, alpha=0.5, beta=3")
print(f"    - Generated {len(pepN_cnts)} peptide counts")

print(f"\n{'-'*50}")
print("• Step 5: Generate Peptide-Level Data from Protein-Level Data")
pep_data = sims.generate_peptide_level(
    control_data,
    pepN_cnts,
    is_log2=False,
    repStd=(0.1, 0.25),
    outlier_fraction=0.0001,
    outlier_multiplier=0.01,
    add_noise=True,
    noise_sd=0.10,
    seed=seed
)
print(f"    - Peptide-level data: {pep_data.shape[0]} peptides × {pep_data.shape[1]} samples")
print(f"    - Noise: sd=0.10, Outliers: fraction=0.0001, multiplier=0.01")

print(f"\n{'-'*50}")
print("• Step 6: Generate Condition Mappers")
condition_sample_map, condition_shifts = sims.generate_condition_mappers(
    n_condition=n_condition-1,
    n_replicates=n_replicates,
    condition_shifts=condition_shifts[1:],
    control_name='control',
    condition_suffix='cond',
)
print(f"    - Condition sample map keys: {list(condition_sample_map.keys())}")
print(f"    - Condition shifts: {condition_shifts}")

print(f"\n{'-'*50}")
print("• Step 7: Generate Complete Peptide-Level Data for All Conditions")
generated_data = sims.generate_complete_data(
    pep_data,
    condition_shifts=condition_shifts,
    condition_sample_map=condition_sample_map,
    shift_scale=0.10,
    is_log2=False,
    add_noise=False,
    noise_sd=0.10,
    seed=seed
)
print(f"    - Complete dataset: {generated_data.shape[0]} peptides × {generated_data.shape[1]} samples")
print(f"    - Noise: sd=0.10, shift_scale=0.10, no additional noise added")

perturb_name = "randomPep"  # Perturbation scenario name
nPep_to_perturb = -1  # Number of peptides to perturb; if > 1 then it is a fraction
perturb_dir_setup = "same"  # Direction of perturbation: [random, same]
np.random.seed(seed)  # Set the seed for reproducibility

print(f"\n{'-'*50}")
print("• Step 8: Setup Random Number of Peptides to Perturb per Protein")
simulated_data = generated_data.copy()
print(f"    - Perturbing {nPro_to_perturb}/{n_proteins} proteins with {nPep_to_perturb} peptides each")
print(f"    - Conditions: {perturb_conds}, Overlap: {perturb_overlap}, Magnitude range: {pertMag_range}")
unique_proteins = simulated_data.reset_index()["Protein"].unique()
proteins_to_perturb = np.random.choice(unique_proteins, nPro_to_perturb, replace=False)
print(f"    - Proteins selected for perturbation: {proteins_to_perturb[:5]} ... (total: {len(proteins_to_perturb)})")

# --- Determine Number of Peptides to Perturb per Protein ---
if nPep_to_perturb == -1:
    nPep_array = np.random.uniform(0.1, 0.5, len(proteins_to_perturb))
elif 0 < nPep_to_perturb < 1:
    nPep_array = np.repeat(nPep_to_perturb, len(proteins_to_perturb))
else:
    nPep_array = np.repeat(int(nPep_to_perturb), len(proteins_to_perturb))
print(f"    - nPep_array example: {nPep_array[:5]}... (total: {len(nPep_array)})")

# --- Subset Data for Perturbation ---
tmp = np.log2(simulated_data).copy()
perturb_data = tmp.loc[proteins_to_perturb]
unchanged_data = tmp.drop(proteins_to_perturb)
# --- Apply Perturbations to Selected Proteins and Peptides ---
dictCnt = 0
perturbation_map = {}
print("    - Applying perturbations to selected proteins and peptides...")
for i, protein in enumerate(proteins_to_perturb):   
    cur_data = perturb_data.loc[protein]
    if nPep_to_perturb < 1:
        n = int(nPep_array[i] * len(cur_data))
        if n < 2: n = 2  # Ensure minimum of 2 peptides are perturbed
    elif nPep_to_perturb >= 1:
        n = int(nPep_to_perturb)
    pepNums = np.random.choice(cur_data.index, n, replace=False)
    conds = np.random.choice(perturb_conds, len(pepNums), replace=True)
    pertMag = np.random.uniform(pertMag_range[0], pertMag_range[1], len(pepNums))
    if perturb_dir_setup == "random":
        pertDir = np.random.choice([-1, 1], len(pepNums), replace=True)
    elif perturb_dir_setup == "same":
        pertDir = np.random.choice([-1, 1], 1, replace=True)
        pertDir = np.repeat(pertDir, len(pepNums))
    for j in range(len(pepNums)):
        perturb_samples = condition_sample_map[conds[j]]
        perturb_peptide = pepNums[j]
        perturb_intensity = cur_data.loc[perturb_peptide, perturb_samples]
        perturb_intensity += pertMag[j] * pertDir[j]
        perturb_data.loc[(protein, perturb_peptide), perturb_samples] = perturb_intensity
        perturbation_map[dictCnt] = {
            "Protein": protein,
            "Peptide": perturb_peptide,
            "pertCondition": conds[j],
            "pertShift": pertMag[j] * pertDir[j],
        }
        dictCnt += 1
print(f"    - Total perturbations applied: {dictCnt}")
# --- Combine Perturbed and Unchanged Data ---
perturbed_data = pd.concat([perturb_data, unchanged_data], axis=0)
perturbed_data = (np.power(2, perturbed_data))
print(f"    - Perturbed data shape: {perturbed_data.shape}")
# --- Build Test Data Object ---
print("    - Building test data object with perturbation map...")
test_data = sims.build_test_data(
    data=perturbed_data,
    condition_sample_map=condition_sample_map,
    perturbation_map=perturbation_map,
    proteins_to_perturb=proteins_to_perturb,
    missing_data=None,
)
complete_data = test_data.copy()
```

```
==================================================
Simulation ID:     Sim2
==================================================
• Simulation Data:
    - Generates a synthetic proteomics dataset with 500 proteins.
    - Each protein has 5–50 peptides, across 3 conditions.
    - The first condition is the control; the rest are experimental.
    - Uses the random (2 to 50%) peptide perturbation strategy as the base and adds different levels of missingness.
• Simulation Goal:
    - Compare the effects of of missingness rates in protein and peptide levels with imputation.
• Parameters:
    - RNG seed: 42
    - 3 conditions (control + 2 treatments), log2 shifts: [0, 1, 2]
    - 500 proteins × 10 replicates, 5–50 peptides/protein
    - Perturbing 250/500 proteins (overlap=False)
    - Perturbation in conditions: ['cond2'], magnitude range: (0.5, 1.5)
==================================================

• Step 1: Generate Protein Mean Values (with outliers)
    - Using seed: 42
    - Generated 500 protein mean values (log2 scale)
    - Stats: mean=20.01, std=1.96, min=13.52, max=27.71
    - Outliers: fraction=10.0%, sd-multiplier=1.0

--------------------------------------------------
• Step 2: Generate Protein Coefficient of Variation (CV) Values
    - Log-normal distribution: mean=10, median=8
    - Generated 500 CV values
    - Stats: mean=10.07, std=8.36, min=0.92, max=104.93

--------------------------------------------------
• Step 3: Generate Replicates for Control (Biological Reference)
    - Generated 10 control replicates for 500 proteins
    - This subset will serve as the control for all treatments.

--------------------------------------------------
• Step 4: Generate Number of Peptides per Protein
    - Beta-binomial: min=5, max=50, alpha=0.5, beta=3
    - Generated 500 peptide counts

--------------------------------------------------
• Step 5: Generate Peptide-Level Data from Protein-Level Data
    - Peptide-level data: 5583 peptides × 12 samples
    - Noise: sd=0.10, Outliers: fraction=0.0001, multiplier=0.01

--------------------------------------------------
• Step 6: Generate Condition Mappers
    - Condition sample map keys: ['control', 'cond1', 'cond2']
    - Condition shifts: {'control': 0, 'cond1': 1, 'cond2': 2}

--------------------------------------------------
• Step 7: Generate Complete Peptide-Level Data for All Conditions
    - Complete dataset: 5583 peptides × 30 samples
    - Noise: sd=0.10, shift_scale=0.10, no additional noise added

--------------------------------------------------
• Step 8: Setup Random Number of Peptides to Perturb per Protein
    - Perturbing 250/500 proteins with -1 peptides each
    - Conditions: ['cond2'], Overlap: False, Magnitude range: (0.5, 1.5)
    - Proteins selected for perturbation: ['pro_362' 'pro_74' 'pro_375' 'pro_156' 'pro_105'] ... (total: 250)
    - nPep_array example: [0.47337452 0.30041595 0.31575098 0.37358551 0.34634047]... (total: 250)
    - Applying perturbations to selected proteins and peptides...
    - Total perturbations applied: 835
    - Perturbed data shape: (5583, 30)
    - Building test data object with perturbation map...
```

```
print(f"\n{'-'*50}")
print("• Step 9: Setup and Run for Different Levels of Missingness")
missRatePeps = [0, 0.2, 0.4, 0.6, 0.8]
missRatePros = [0, 0.2, 0.4, 0.6, 0.8]
nVersions = len(missRatePeps) * len(missRatePros)
print(f"    - Running for {nVersions} combinations of protein and peptide missingness:")
for i in missRatePros:
    # Calculate the numbers based on the protein missingness
    # n_amputate_1 and n_amputate_2 should add to non-perturbed proteins
    n_amputate_2 = int(i * len(proteins_to_perturb))
    n_amputate_1 = len(proteins_to_perturb) - n_amputate_2
    non_perturbed_proteins = np.setdiff1d(unique_proteins, proteins_to_perturb)
    n_amputate_3 = int(i * len(non_perturbed_proteins))
    for j in missRatePeps:
        print(f"     - Version = Protein Missingness: {i}, Peptide Missingness: {j}, saving as: 2_Pro{i}_Pep{j}_imputed_InputData_feather")
        # Find out the number of 
        missing_data = sims.amputation(
            data = perturbed_data,
            unique_proteins = unique_proteins,
            proteins_to_perturb = proteins_to_perturb,
            condition_shifts = condition_shifts,
            condition_sample_map = condition_sample_map,
            n_amputate_1 = n_amputate_1,
            n_amputate_2 = n_amputate_2,
            n_amputate_3 = n_amputate_3,
            missing_rate = j, # Peptide missingness rate
            seed = seed
        )

        imputed_data = sims.downshifted_imputation(
            data = missing_data,
            condition_sample_map = condition_sample_map,
            is_log2 = False,
            shiftMag = 2,
            lowPct = 0.15,
            minValue = 8,
            impute_all = False,
            seed = seed
        )
        test_data = sims.build_test_data(
            data = imputed_data,
            condition_sample_map = condition_sample_map,
            perturbation_map = perturbation_map,
            proteins_to_perturb = proteins_to_perturb,
            missing_data = missing_data,
        )

        test_data.to_feather(f"{output_path}2_Pro{i}_Pep{j}_imputed_InputData.feather")

elapsed = utils.prettyTimer(utils.getTime() - stTime)
print(f"\n{'-'*50}")
print(f"• Completed generation of {nVersions} versions for simulation: {simID}")
print(f"    - Elapsed time: {elapsed}")
print(f"{'-'*50}\n")
```

```
--------------------------------------------------
• Step 9: Setup and Run for Different Levels of Missingness
    - Running for 25 combinations of protein and peptide missingness:
     - Version = Protein Missingness: 0, Peptide Missingness: 0, saving as: 2_Pro0_Pep0_imputed_InputData_feather
     - Version = Protein Missingness: 0, Peptide Missingness: 0.2, saving as: 2_Pro0_Pep0.2_imputed_InputData_feather
     - Version = Protein Missingness: 0, Peptide Missingness: 0.4, saving as: 2_Pro0_Pep0.4_imputed_InputData_feather
     - Version = Protein Missingness: 0, Peptide Missingness: 0.6, saving as: 2_Pro0_Pep0.6_imputed_InputData_feather
     - Version = Protein Missingness: 0, Peptide Missingness: 0.8, saving as: 2_Pro0_Pep0.8_imputed_InputData_feather
     - Version = Protein Missingness: 0.2, Peptide Missingness: 0, saving as: 2_Pro0.2_Pep0_imputed_InputData_feather
     - Version = Protein Missingness: 0.2, Peptide Missingness: 0.2, saving as: 2_Pro0.2_Pep0.2_imputed_InputData_feather
     - Version = Protein Missingness: 0.2, Peptide Missingness: 0.4, saving as: 2_Pro0.2_Pep0.4_imputed_InputData_feather
     - Version = Protein Missingness: 0.2, Peptide Missingness: 0.6, saving as: 2_Pro0.2_Pep0.6_imputed_InputData_feather
     - Version = Protein Missingness: 0.2, Peptide Missingness: 0.8, saving as: 2_Pro0.2_Pep0.8_imputed_InputData_feather
     - Version = Protein Missingness: 0.4, Peptide Missingness: 0, saving as: 2_Pro0.4_Pep0_imputed_InputData_feather
     - Version = Protein Missingness: 0.4, Peptide Missingness: 0.2, saving as: 2_Pro0.4_Pep0.2_imputed_InputData_feather
     - Version = Protein Missingness: 0.4, Peptide Missingness: 0.4, saving as: 2_Pro0.4_Pep0.4_imputed_InputData_feather
     - Version = Protein Missingness: 0.4, Peptide Missingness: 0.6, saving as: 2_Pro0.4_Pep0.6_imputed_InputData_feather
     - Version = Protein Missingness: 0.4, Peptide Missingness: 0.8, saving as: 2_Pro0.4_Pep0.8_imputed_InputData_feather
     - Version = Protein Missingness: 0.6, Peptide Missingness: 0, saving as: 2_Pro0.6_Pep0_imputed_InputData_feather
     - Version = Protein Missingness: 0.6, Peptide Missingness: 0.2, saving as: 2_Pro0.6_Pep0.2_imputed_InputData_feather
     - Version = Protein Missingness: 0.6, Peptide Missingness: 0.4, saving as: 2_Pro0.6_Pep0.4_imputed_InputData_feather
     - Version = Protein Missingness: 0.6, Peptide Missingness: 0.6, saving as: 2_Pro0.6_Pep0.6_imputed_InputData_feather
     - Version = Protein Missingness: 0.6, Peptide Missingness: 0.8, saving as: 2_Pro0.6_Pep0.8_imputed_InputData_feather
     - Version = Protein Missingness: 0.8, Peptide Missingness: 0, saving as: 2_Pro0.8_Pep0_imputed_InputData_feather
     - Version = Protein Missingness: 0.8, Peptide Missingness: 0.2, saving as: 2_Pro0.8_Pep0.2_imputed_InputData_feather
     - Version = Protein Missingness: 0.8, Peptide Missingness: 0.4, saving as: 2_Pro0.8_Pep0.4_imputed_InputData_feather
     - Version = Protein Missingness: 0.8, Peptide Missingness: 0.6, saving as: 2_Pro0.8_Pep0.6_imputed_InputData_feather
     - Version = Protein Missingness: 0.8, Peptide Missingness: 0.8, saving as: 2_Pro0.8_Pep0.8_imputed_InputData_feather

--------------------------------------------------
• Completed generation of 25 versions for simulation: Sim2
    - Elapsed time: 00h:00m:19s
--------------------------------------------------
```

---

#### Simulation 3: Perturbation Magnitude Sensitivity¶

**Objective:** Determine the minimum perturbation magnitude required for reliable detection across methods.

**Experimental Design:**

- **Base dataset**: Same structure as Sim1/Sim2
- **Perturbation**: Random 10–50% pattern with **random direction** (up or down)
- **Magnitude ranges tested** (log2 fold-change):
  | Range | Interpretation |
  |-------|----------------|
  | 0.10–0.25 | Subtle (e.g., phosphorylation stoichiometry) |
  | 0.25–0.50 | Small |
  | 0.50–0.75 | Moderate |
  | 0.75–1.00 | Medium |
  | 1.00–1.25 | Substantial |
  | 1.25–1.50 | Large |
  | 1.50–1.75 | Strong |
  | 1.75–2.00 | Very strong (e.g., isoform switching) |
- **Data**: Imputed versions only (25% baseline missingness)

**Key Question:** What is the practical detection threshold for each method?

**Output Files:** `./data/Sim3/2_{low}_{high}_InputData.feather`

##### Base Dataset Generation¶

```
stTime = utils.getTime()

# --- Simulation Parameters ---
simID = "Sim3"
seed = 42  # Seed for reproducibility
n_condition = 3  # Number of conditions (1 is control, others are cond-N-1)
n_proteins = 500  # Number of proteins in the dataset
n_replicates = 10  # Number of replicates per condition
n_peptides = (5, 50)  # (min, max) for peptides per protein
condition_shifts = [0, 1, 2]  # Shifts for the conditions
nPro_to_perturb = 250  # Number of proteins to perturb
perturb_conds = ['cond2']  # Conditions to perturb
perturb_overlap = False  # Allow overlap between perturbed peptides
# Magnitude of perturbations to simulate
perturbationRanges = [
    (0.1, 0.25),
    (0.25, 0.50),
    (0.50, 0.75),
    (0.75, 1.0),
    (1.0, 1.25),
    (1.25, 1.50),
    (1.50, 1.75),
    (1.75, 2.0),
]

# Generate the paths for it.
output_path, figure_path = setup_simulation_paths( simID )

print(f"\n{'='*50}")
print(f"{'Simulation ID:':<18} {simID}")
print(f"{'='*50}")
print("• Simulation Data:")
print(f"    - Generates a synthetic proteomics dataset with {n_proteins} proteins.")
print(f"    - Each protein has {n_peptides[0]}–{n_peptides[1]} peptides, across {n_condition} conditions.")
print(f"    - The first condition is the control; the rest are experimental.")
print(f"    - Uses the random (2 to 50%) peptide perturbation strategy as the base and adds different levels of missingness.")
print("• Simulation Goal:")
print("    - To evaluate the effect of different perturbation magnitude ranges on downstream analyses.")
print("• Parameters:")
print(f"    - RNG seed: {seed}")
print(f"    - {n_condition} conditions (control + {n_condition-1} treatments), log2 shifts: {condition_shifts}")
print(f"    - {n_proteins} proteins × {n_replicates} replicates, {n_peptides[0]}–{n_peptides[1]} peptides/protein")
print(f"    - Perturbing {nPro_to_perturb}/{n_proteins} proteins (overlap={perturb_overlap})")
print(f"    - Perturbation in conditions: {perturb_conds}")
print(f"    - Perturbation magnitude ranges tested: \n       {perturbationRanges}")
print(f"{'='*50}\n")
print("• Step 1: Generate Protein Mean Values (with outliers)")
print(f"    - Using seed: {seed}")
print(f"{'='*50}\n")
print("• Step 1: Generate Protein Mean Values (with outliers)")
print(f"    - Using seed: {seed}")
mean_values = sims.normal_distribution_with_outliers(
    mu=20,
    sd=2,
    size=n_proteins,
    is_log2=False,
    outlier_fraction=0.10,
    outlier_sd_multiplier=1.0,
    seed=seed
)
print(f"    - Generated {len(mean_values)} protein mean values (log2 scale)")
print(f"    - Stats: mean={np.mean(mean_values):.2f}, std={np.std(mean_values):.2f}, min={np.min(mean_values):.2f}, max={np.max(mean_values):.2f}")
print(f"    - Outliers: fraction=10.0%, sd-multiplier=1.0")
mean_values = 2 ** mean_values

print(f"\n{'-'*50}")
print("• Step 2: Generate Protein Coefficient of Variation (CV) Values")
cv_values = sims.lognormal_distribution(
    mu=10,
    med=8,
    size=n_proteins,
    seed=seed
)
print(f"    - Log-normal distribution: mean=10, median=8")
print(f"    - Generated {len(cv_values)} CV values")
print(f"    - Stats: mean={np.mean(cv_values):.2f}, std={np.std(cv_values):.2f}, min={np.min(cv_values):.2f}, max={np.max(cv_values):.2f}")

print(f"\n{'-'*50}")
print("• Step 3: Generate Replicates for Control (Biological Reference)")
control_data = sims.generate_replicates(
    mean_values,
    cv_values,
    meanScale='raw',
    cvType='percent',
    nReps=n_replicates,
    randomizeCV=True,
    as_dataframe=True,
    seed=seed
)
print(f"    - Generated {control_data.shape[1]} control replicates for {control_data.shape[0]} proteins")
print(f"    - This subset will serve as the control for all treatments.")

print(f"\n{'-'*50}")
print("• Step 4: Generate Number of Peptides per Protein")
pepN_cnts = sims.generate_peptide_counts(
    n_proteins=n_proteins,
    min_peptides=n_peptides[0],
    max_peptides=n_peptides[1],
    alpha=0.5,
    beta=3,
)
print(f"    - Beta-binomial: min={n_peptides[0]}, max={n_peptides[1]}, alpha=0.5, beta=3")
print(f"    - Generated {len(pepN_cnts)} peptide counts")

print(f"\n{'-'*50}")
print("• Step 5: Generate Peptide-Level Data from Protein-Level Data")
pep_data = sims.generate_peptide_level(
    control_data,
    pepN_cnts,
    is_log2=False,
    repStd=(0.1, 0.25),
    outlier_fraction=0.0001,
    outlier_multiplier=0.01,
    add_noise=True,
    noise_sd=0.10,
    seed=seed
)
print(f"    - Peptide-level data: {pep_data.shape[0]} peptides × {pep_data.shape[1]} samples")
print(f"    - Noise: sd=0.10, Outliers: fraction=0.0001, multiplier=0.01")

print(f"\n{'-'*50}")
print("• Step 6: Generate Condition Mappers")
condition_sample_map, condition_shifts = sims.generate_condition_mappers(
    n_condition=n_condition-1,
    n_replicates=n_replicates,
    condition_shifts=condition_shifts[1:],
    control_name='control',
    condition_suffix='cond',
)
print(f"    - Condition sample map keys: {list(condition_sample_map.keys())}")
print(f"    - Condition shifts: {condition_shifts}")

print(f"\n{'-'*50}")
print("• Step 7: Generate Complete Peptide-Level Data for All Conditions")
generated_data = sims.generate_complete_data(
    pep_data,
    condition_shifts=condition_shifts,
    condition_sample_map=condition_sample_map,
    shift_scale=0.10,
    is_log2=False,
    add_noise=False,
    noise_sd=0.10,
    seed=seed
)
print(f"    - Complete dataset: {generated_data.shape[0]} peptides × {generated_data.shape[1]} samples")
print(f"    - Noise: sd=0.10, shift_scale=0.10, no additional noise added")
```

```
==================================================
Simulation ID:     Sim3
==================================================
• Simulation Data:
    - Generates a synthetic proteomics dataset with 500 proteins.
    - Each protein has 5–50 peptides, across 3 conditions.
    - The first condition is the control; the rest are experimental.
    - Uses the random (2 to 50%) peptide perturbation strategy as the base and adds different levels of missingness.
• Simulation Goal:
    - To evaluate the effect of different perturbation magnitude ranges on downstream analyses.
• Parameters:
    - RNG seed: 42
    - 3 conditions (control + 2 treatments), log2 shifts: [0, 1, 2]
    - 500 proteins × 10 replicates, 5–50 peptides/protein
    - Perturbing 250/500 proteins (overlap=False)
    - Perturbation in conditions: ['cond2']
    - Perturbation magnitude ranges tested: 
       [(0.1, 0.25), (0.25, 0.5), (0.5, 0.75), (0.75, 1.0), (1.0, 1.25), (1.25, 1.5), (1.5, 1.75), (1.75, 2.0)]
==================================================

• Step 1: Generate Protein Mean Values (with outliers)
    - Using seed: 42
==================================================

• Step 1: Generate Protein Mean Values (with outliers)
    - Using seed: 42
    - Generated 500 protein mean values (log2 scale)
    - Stats: mean=20.01, std=1.96, min=13.52, max=27.71
    - Outliers: fraction=10.0%, sd-multiplier=1.0

--------------------------------------------------
• Step 2: Generate Protein Coefficient of Variation (CV) Values
    - Log-normal distribution: mean=10, median=8
    - Generated 500 CV values
    - Stats: mean=10.07, std=8.36, min=0.92, max=104.93

--------------------------------------------------
• Step 3: Generate Replicates for Control (Biological Reference)
    - Generated 10 control replicates for 500 proteins
    - This subset will serve as the control for all treatments.

--------------------------------------------------
• Step 4: Generate Number of Peptides per Protein
    - Beta-binomial: min=5, max=50, alpha=0.5, beta=3
    - Generated 500 peptide counts

--------------------------------------------------
• Step 5: Generate Peptide-Level Data from Protein-Level Data
    - Peptide-level data: 5583 peptides × 12 samples
    - Noise: sd=0.10, Outliers: fraction=0.0001, multiplier=0.01

--------------------------------------------------
• Step 6: Generate Condition Mappers
    - Condition sample map keys: ['control', 'cond1', 'cond2']
    - Condition shifts: {'control': 0, 'cond1': 1, 'cond2': 2}

--------------------------------------------------
• Step 7: Generate Complete Peptide-Level Data for All Conditions
    - Complete dataset: 5583 peptides × 30 samples
    - Noise: sd=0.10, shift_scale=0.10, no additional noise added
```

```
nVersions = len(perturbationRanges) * 2  # Each with complete and imputed data
simulated_data = generated_data.copy()
nPep_to_perturb = -1            # Number of peptides to perturb if > 1 then it is a fraction
perturb_dir_setup = "random"    # Randomly perturb the direction of the perturbation [random, same]
np.random.seed(seed)            # Set the seed

print(f"\n{'-'*50}")
print("• Step 9: Setup and Run for Different Levels of Missingness")

print(f"    - Running for {nVersions} combinations of perturbation magnitude ranges:")
for pertMag_range in perturbationRanges:
    print(f"     - Version = Perturbation Magnitude Range: {pertMag_range}, saving as: 2_{pertMag_range[0]}_{pertMag_range[1]}_InputData.feather") 
    # Select random proteins to perturb
    proteins_to_perturb = np.random.choice(unique_proteins, nPro_to_perturb, replace=False)
    # Pick a fraction between 0.1 and 0.5 of the peptides to perturb
    if nPep_to_perturb == -1:
        nPep_array = np.random.uniform(0.1, 0.5, len(unique_proteins))
    elif nPep_to_perturb < 1 and nPep_to_perturb > 0:
        nPep_array = np.repeat(nPep_to_perturb, len(unique_proteins))
    elif nPep_to_perturb >= 1:
        nPep_array = np.repeat(nPep_to_perturb, len(unique_proteins))

    # Subset the data for perturbation
    tmp = np.log2(simulated_data).copy()
    perturb_data = tmp.loc[proteins_to_perturb]
    unchanged_data = tmp.drop(proteins_to_perturb)

    # Go through the proteins and perturb single peptide in each
    dictCnt = 0
    perturbation_map = {}

    for i, protein in enumerate(proteins_to_perturb):
        cur_data = perturb_data.loc[protein]
        if nPep_to_perturb < 1:
            n = int(nPep_array[i] * len(cur_data))
            # Ensure minimum of 1 peptide is perturbed
            if n < 2: n = 2
        elif nPep_to_perturb >= 1:
            n = int(nPep_to_perturb)
        # print(f"Protein: {protein} - Peptides: {n}, Total Peptides: {len(cur_data)}")
        pepNums = np.random.choice(cur_data.index, n, replace=False)
        # Identify the conditions to perturb
        conds = np.random.choice(perturb_conds, len(pepNums), replace=True)
        # Identify the magnitude of perturbation
        pertMag = np.random.uniform(pertMag_range[0], pertMag_range[1], len(pepNums))
        #
        if perturb_dir_setup == "random":
            pertDir = np.random.choice([-1, 1], len(pepNums), replace=True)
        elif perturb_dir_setup == "same":
            # Pick random direction
            pertDir = np.random.choice([-1, 1], 1, replace=True)
            pertDir = np.repeat(pertDir, len(pepNums))
        
        for j in range(len(pepNums)):
            perturb_samples = condition_sample_map[conds[j]]
            # Get the peptides to perturb for the current condition
            perturb_peptide = pepNums[j]
            # Get the perturbation peptide intensity
            perturb_intensity = cur_data.loc[perturb_peptide, perturb_samples]
            # Update the perturbation peptide intensity
            perturb_intensity += pertMag[j] * pertDir[j]
            perturb_data.loc[(protein, perturb_peptide), perturb_samples] = perturb_intensity
            perturbation_map[dictCnt] = {
                "Protein": protein,
                "Peptide": perturb_peptide,
                "pertCondition": conds[j],
                "pertShift": pertMag[j] * pertDir[j],
            }
            dictCnt += 1

    perturbed_data = pd.concat([perturb_data, unchanged_data], axis=0)
    perturbed_data = (np.power(2, perturbed_data))

    missing_data = sims.amputation(
        data = perturbed_data,
        unique_proteins = unique_proteins,
        proteins_to_perturb = proteins_to_perturb,
        condition_shifts = condition_shifts,
        condition_sample_map = condition_sample_map,
        n_amputate_1 = 50,
        n_amputate_2 = 100,
        n_amputate_3 = 50,
        missing_rate = 0.25,
        seed = seed
    )

    imputed_data = sims.downshifted_imputation(
        data = missing_data,
        condition_sample_map = condition_sample_map,
        is_log2 = False,
        shiftMag = 2,
        lowPct = 0.15,
        minValue = 8,
        impute_all = False,
        seed = seed
    )
    test_data = sims.build_test_data(
        data = imputed_data,
        condition_sample_map = condition_sample_map,
        perturbation_map = perturbation_map,
        proteins_to_perturb = proteins_to_perturb,
        missing_data = missing_data,
    )
    test_data.to_feather(
        f"{output_path}2_{pertMag_range[0]}_{pertMag_range[1]}_InputData.feather"
    )

elapsed = utils.prettyTimer(utils.getTime() - stTime)
print(f"\n{'-'*50}")
print(f"• Completed generation of {nVersions} versions for simulation: {simID}")
print(f"    - Elapsed time: {elapsed}")
print(f"{'-'*50}\n")
```

```
--------------------------------------------------
• Step 9: Setup and Run for Different Levels of Missingness
    - Running for 16 combinations of perturbation magnitude ranges:
     - Version = Perturbation Magnitude Range: (0.1, 0.25), saving as: 2_0.1_0.25_InputData.feather
     - Version = Perturbation Magnitude Range: (0.25, 0.5), saving as: 2_0.25_0.5_InputData.feather
     - Version = Perturbation Magnitude Range: (0.5, 0.75), saving as: 2_0.5_0.75_InputData.feather
     - Version = Perturbation Magnitude Range: (0.75, 1.0), saving as: 2_0.75_1.0_InputData.feather
     - Version = Perturbation Magnitude Range: (1.0, 1.25), saving as: 2_1.0_1.25_InputData.feather
     - Version = Perturbation Magnitude Range: (1.25, 1.5), saving as: 2_1.25_1.5_InputData.feather
     - Version = Perturbation Magnitude Range: (1.5, 1.75), saving as: 2_1.5_1.75_InputData.feather
     - Version = Perturbation Magnitude Range: (1.75, 2.0), saving as: 2_1.75_2.0_InputData.feather

--------------------------------------------------
• Completed generation of 16 versions for simulation: Sim3
    - Elapsed time: 00h:00m:12s
--------------------------------------------------
```

---

#### Simulation 4: Complex Multi-Variable Experimental Designs¶

**Objective:** Evaluate method robustness across experimental designs with multiple interacting factors.

**Experimental Design:**

- **Base dataset**: Same protein/peptide structure, but variable number of conditions
- **Factors varied**:

| Factor | Levels | Description |
| --- | --- | --- |
| **Conditions** | 2, 3, 4, 5, 6 | Number of experimental conditions (first is control) |
| **Overlap** | True, False | Same peptides perturbed across conditions vs. different peptides |
| **Direction** | same, random | All perturbations in same direction vs. random up/down |

- **Total combinations**: 5 × 2 × 2 = **20 datasets**
- **Condition strategy**:
  - 2 conditions → perturb last condition only
  - 3+ conditions → perturb last two conditions
- **Perturbation**: Random 10–50% pattern, magnitude 0.5–1.5 log2
- **Data**: Imputed versions (25% baseline missingness)

**Key Questions:**

- Does adding more conditions improve detection?
- Are overlapping perturbation patterns easier or harder to detect?
- Does perturbation direction consistency affect accuracy?

**Output Files:** `./data/Sim4/2_{N}Cond_{Overlap|NonOverlap}_{same|random}Dir_InputData.feather`

##### Systematic Dataset Generation¶

```
stTime = utils.getTime()

## Global Variables to be Used for the Simulation
simID = "Sim4"                  # Simulation ID
n_proteins = 500                # Number of proteins in the dataset
n_replicates = 10               # Number of replicates per condition
n_peptides = (5, 50)            # (min, max) for peptides per protein
nPro_to_perturb = 250           # Number of proteins to perturb
pertMag_range = (.5, 1.5)       # Range of values to perturb the peptides with (uniform)
nPep_to_perturb = -1            # Randomly perturb between 2 and 50% of the peptides

# Systematic parameter combinations
overlap_types = [True, False]    # Overlap vs Non-overlap
direction_types = ["random", "same"]  # Direction of perturbations

# Number of conditions to generate (with shifts)
conditions = {
    2: [0, .5],
    3: [0, .5, 1],
    4: [0, .5, 1, 1.5],
    5: [0, .5, 1, 1.5, 2],
    6: [0, .5, 1, 1.5, 2, 2.5]
}

# Calculate total versions
total_combinations = len(conditions) * len(overlap_types) * len(direction_types)

# Seed for reproducibility
seed = 42                       

# Generate the paths for it.
output_path, figure_path = setup_simulation_paths( simID )

print(f"\n{'='*50}")
print(f"{'Simulation ID:':<18} {simID}")
print(f"{'='*50}")
print("• Simulation Data:")
print(f"    - Generates a synthetic proteomics dataset with {n_proteins} proteins.")
print(f"    - Each protein has {n_peptides[0]}–{n_peptides[1]} peptides, across multiple conditions.")
print(f"    - The first condition is the control; the rest are experimental.")
print(f"    - Uses systematic combinations of perturbation parameters.")
print("• Simulation Goal:")
print("    - To evaluate the effect of different perturbation strategies on downstream analyses.")
print("• Parameters:")
print(f"    - RNG seed: {seed}")
print(f"    - {n_proteins} proteins × {n_replicates} replicates, {n_peptides[0]}–{n_peptides[1]} peptides/protein")
print(f"    - Perturbing {nPro_to_perturb}/{n_proteins} proteins")
print(f"    - Number of conditions: {list(conditions.keys())}")
print(f"    - Overlap types: {overlap_types}")
print(f"    - Direction types: {direction_types}")
print(f"    - Condition strategy: 2 conditions uses last only; 3+ conditions use last two")
print(f"    - Perturbation magnitude range: {pertMag_range}")
print(f"    - Total combinations: {total_combinations}")
print(f"{'='*50}\n")

print(f"• Step 1: Generate systematic combinations of datasets")
version_count = 0

for n_condition, condition_shifts in conditions.items():
    print(f"\n--- Processing {n_condition} conditions with shifts: {condition_shifts} ---")
    
    # Set seed for each condition set to ensure reproducibility
    np.random.seed(seed)
    
    # Create Protein Mean Values
    mean_values = sims.normal_distribution_with_outliers(
        mu=20, sd=2, size=n_proteins, 
        is_log2=False, outlier_fraction=0.10, 
        outlier_sd_multiplier=1.0, seed=seed
    )
    mean_values = 2**mean_values

    # Create Protein CV Values
    cv_values = sims.lognormal_distribution(
        mu=10, med=8, size=n_proteins, seed=seed
    )

    # Generate replicates to have a biological sample as reference
    control_data = sims.generate_replicates(
        mean_values, cv_values, meanScale="raw", cvType="percent", 
        nReps=n_replicates, randomizeCV=True, as_dataframe=True, seed=seed
    )

    # Generate the number of peptides per protein
    pepN_cnts = sims.generate_peptide_counts(
        n_proteins=n_proteins, min_peptides=n_peptides[0],
        max_peptides=n_peptides[1], alpha=.5, beta=3,
    )

    # Generate the peptide level data using the protein level data 
    pep_data = sims.generate_peptide_level(
        control_data, pepN_cnts, is_log2=False, repStd=(0.1, 0.25),
        outlier_fraction=0.0001, outlier_multiplier=0.01,
        add_noise=True, noise_sd=0.10, seed=seed
    )

    # Generate Condition Mappers
    condition_sample_map, condition_shifts_adj = sims.generate_condition_mappers(
        n_condition=n_condition-1, n_replicates=n_replicates,
        condition_shifts=condition_shifts[1:], control_name='control',
        condition_suffix='cond',
    )

    # Generate the complete peptide level data with all conditions
    complete_data = sims.generate_complete_data(
        pep_data, condition_shifts=condition_shifts_adj,
        condition_sample_map=condition_sample_map, shift_scale=0.10,
        is_log2=False, add_noise=False, noise_sd=0.10, seed=seed
    )
    
    # Get available conditions for perturbation
    available_conditions = list(condition_sample_map.keys())
    unique_proteins = complete_data.reset_index()["Protein"].unique()
    
    # Determine conditions to perturb based on number of conditions
    if n_condition == 2:
        # For 2 conditions: use the last (and only) treatment condition
        perturb_conds = [available_conditions[-1]]
    else:
        # For 3+ conditions: use the last two treatment conditions
        perturb_conds = available_conditions[-2:]
    
    # Iterate through all parameter combinations
    for overlap_type in overlap_types:
        for direction_type in direction_types:
            version_count += 1
            
            # Create descriptive filename
            overlap_str = "Overlap" if overlap_type else "NonOverlap"
            filename = f"2_{n_condition}Cond_{overlap_str}_{direction_type}Dir_InputData.feather"
            
            print(f"    [{version_count:2d}/{total_combinations}] Generating: {filename}")
            print(f"        - Available conditions: {available_conditions}")
            print(f"        - Conditions to perturb: {perturb_conds}")
            print(f"        - Overlap: {overlap_type}, Direction: {direction_type}")

            
            # Reset random seed for consistent protein selection
            np.random.seed(seed + version_count)
            
            simulated_data = complete_data.copy()
            proteins_to_perturb = np.random.choice(unique_proteins, nPro_to_perturb, replace=False)

            # Determine number of peptides to perturb per protein
            if nPep_to_perturb == -1:
                nPep_array = np.random.uniform(0.1, 0.5, len(proteins_to_perturb))
            elif 0 < nPep_to_perturb < 1:
                nPep_array = np.repeat(nPep_to_perturb, len(proteins_to_perturb))
            else:
                nPep_array = np.repeat(int(nPep_to_perturb), len(proteins_to_perturb))

            # Subset the data for perturbation
            tmp = np.log2(simulated_data).copy()
            perturb_data = tmp.loc[proteins_to_perturb]
            unchanged_data = tmp.drop(proteins_to_perturb)

            # Apply perturbations
            dictCnt = 0
            perturbation_map = {}
            
            for i, protein in enumerate(proteins_to_perturb):
                cur_data = perturb_data.loc[protein]
                
                # Determine number of peptides to perturb
                if nPep_to_perturb < 1:
                    n = int(nPep_array[i] * len(cur_data))
                    if n < 2: n = 2  # Minimum 2 peptides
                else:
                    n = int(nPep_to_perturb)
                
                pepNums = np.random.choice(cur_data.index, n, replace=False)
                
                # Configure perturbation based on overlap and direction
                if overlap_type:
                    # Overlap: same peptides perturbed across multiple conditions
                    pertMag = np.random.uniform(pertMag_range[0], pertMag_range[1], 
                                              (len(pepNums), len(perturb_conds)))
                    if direction_type == "random":
                        pertDir = np.random.choice([-1, 1], 
                                                 (len(pepNums), len(perturb_conds)), 
                                                 replace=True)
                    else:  # "same"
                        pertDir = np.random.choice([-1, 1], 1, replace=True)
                        pertDir = np.repeat(pertDir, len(pepNums) * len(perturb_conds)).reshape(
                            (len(pepNums), len(perturb_conds)))
                    
                    perturb_shift = pertMag * pertDir
                    
                    # Apply perturbations to all conditions
                    for j, cond in enumerate(perturb_conds):
                        perturb_samples = condition_sample_map[cond]
                        perturb_intensity = cur_data.loc[pepNums, perturb_samples]
                        perturb_intensity += perturb_shift[:, j][:, np.newaxis]
                        perturb_data.loc[(protein, pepNums), perturb_samples] = perturb_intensity.values
                    
                    # Update perturbation map
                    for p, peptide in enumerate(pepNums):
                        perturbation_map[dictCnt] = {
                            "Protein": protein,
                            "Peptide": peptide,
                            "pertCondition": perturb_conds,
                            "pertShift": perturb_shift[p, :].tolist(),
                            "overlapType": overlap_str,
                            "directionType": direction_type
                        }
                        dictCnt += 1
                        
                else:
                    # Non-overlap: different peptides for different conditions
                    pertMag = np.random.uniform(pertMag_range[0], pertMag_range[1], len(pepNums))
                    
                    if direction_type == "random":
                        pertDir = np.random.choice([-1, 1], len(pepNums), replace=True)
                    else:  # "same"
                        pertDir = np.random.choice([-1, 1], 1, replace=True)
                        pertDir = np.repeat(pertDir, len(pepNums))
                    
                    # Assign peptides to conditions (distribute evenly)
                    peptide_conditions = np.random.choice(perturb_conds, len(pepNums), replace=True)
                    
                    for p, (peptide, cond) in enumerate(zip(pepNums, peptide_conditions)):
                        perturb_samples = condition_sample_map[cond]
                        perturb_intensity = cur_data.loc[peptide, perturb_samples]
                        shift_value = pertMag[p] * pertDir[p]
                        perturb_intensity += shift_value
                        perturb_data.loc[(protein, peptide), perturb_samples] = perturb_intensity
                        
                        perturbation_map[dictCnt] = {
                            "Protein": protein,
                            "Peptide": peptide,
                            "pertCondition": [cond],
                            "pertShift": [shift_value],
                            "overlapType": overlap_str,
                            "directionType": direction_type
                        }
                        dictCnt += 1

            # Combine perturbed and unchanged data
            perturbed_data = pd.concat([perturb_data, unchanged_data], axis=0)
            perturbed_data = np.power(2, perturbed_data)

            # Apply missing data simulation
            missing_data = sims.amputation(
                data=perturbed_data, unique_proteins=unique_proteins,
                proteins_to_perturb=proteins_to_perturb,
                condition_shifts=condition_shifts_adj,
                condition_sample_map=condition_sample_map,
                n_amputate_1=50, n_amputate_2=100, n_amputate_3=50,
                missing_rate=0.25, seed=seed + version_count
            )

            # Impute missing data
            imputed_data = sims.downshifted_imputation(
                data=missing_data, condition_sample_map=condition_sample_map,
                is_log2=False, shiftMag=2, lowPct=0.15, minValue=8,
                impute_all=False, seed=seed + version_count
            )

            # Build final test data
            test_data = sims.build_test_data(
                data=imputed_data, condition_sample_map=condition_sample_map,
                perturbation_map=perturbation_map,
                proteins_to_perturb=proteins_to_perturb,
                missing_data=missing_data,
            )

            # Save the data
            test_data.to_feather(f"{output_path}{filename}")

print(f"\n{'-'*50}")
print(f"• Completed generation of {version_count} versions for simulation: {simID}")
elapsed = utils.prettyTimer(utils.getTime() - stTime)
print(f"    - Elapsed time: {elapsed}")
print(f"{'-'*50}\n")
```

```
==================================================
Simulation ID:     Sim4
==================================================
• Simulation Data:
    - Generates a synthetic proteomics dataset with 500 proteins.
    - Each protein has 5–50 peptides, across multiple conditions.
    - The first condition is the control; the rest are experimental.
    - Uses systematic combinations of perturbation parameters.
• Simulation Goal:
    - To evaluate the effect of different perturbation strategies on downstream analyses.
• Parameters:
    - RNG seed: 42
    - 500 proteins × 10 replicates, 5–50 peptides/protein
    - Perturbing 250/500 proteins
    - Number of conditions: [2, 3, 4, 5, 6]
    - Overlap types: [True, False]
    - Direction types: ['random', 'same']
    - Condition strategy: 2 conditions uses last only; 3+ conditions use last two
    - Perturbation magnitude range: (0.5, 1.5)
    - Total combinations: 20
==================================================

• Step 1: Generate systematic combinations of datasets

--- Processing 2 conditions with shifts: [0, 0.5] ---
    [ 1/20] Generating: 2_2Cond_Overlap_randomDir_InputData.feather
        - Available conditions: ['control', 'cond1']
        - Conditions to perturb: ['cond1']
        - Overlap: True, Direction: random
    [ 2/20] Generating: 2_2Cond_Overlap_sameDir_InputData.feather
        - Available conditions: ['control', 'cond1']
        - Conditions to perturb: ['cond1']
        - Overlap: True, Direction: same
    [ 3/20] Generating: 2_2Cond_NonOverlap_randomDir_InputData.feather
        - Available conditions: ['control', 'cond1']
        - Conditions to perturb: ['cond1']
        - Overlap: False, Direction: random
    [ 4/20] Generating: 2_2Cond_NonOverlap_sameDir_InputData.feather
        - Available conditions: ['control', 'cond1']
        - Conditions to perturb: ['cond1']
        - Overlap: False, Direction: same

--- Processing 3 conditions with shifts: [0, 0.5, 1] ---
    [ 5/20] Generating: 2_3Cond_Overlap_randomDir_InputData.feather
        - Available conditions: ['control', 'cond1', 'cond2']
        - Conditions to perturb: ['cond1', 'cond2']
        - Overlap: True, Direction: random
    [ 6/20] Generating: 2_3Cond_Overlap_sameDir_InputData.feather
        - Available conditions: ['control', 'cond1', 'cond2']
        - Conditions to perturb: ['cond1', 'cond2']
        - Overlap: True, Direction: same
    [ 7/20] Generating: 2_3Cond_NonOverlap_randomDir_InputData.feather
        - Available conditions: ['control', 'cond1', 'cond2']
        - Conditions to perturb: ['cond1', 'cond2']
        - Overlap: False, Direction: random
    [ 8/20] Generating: 2_3Cond_NonOverlap_sameDir_InputData.feather
        - Available conditions: ['control', 'cond1', 'cond2']
        - Conditions to perturb: ['cond1', 'cond2']
        - Overlap: False, Direction: same

--- Processing 4 conditions with shifts: [0, 0.5, 1, 1.5] ---
    [ 9/20] Generating: 2_4Cond_Overlap_randomDir_InputData.feather
        - Available conditions: ['control', 'cond1', 'cond2', 'cond3']
        - Conditions to perturb: ['cond2', 'cond3']
        - Overlap: True, Direction: random
    [10/20] Generating: 2_4Cond_Overlap_sameDir_InputData.feather
        - Available conditions: ['control', 'cond1', 'cond2', 'cond3']
        - Conditions to perturb: ['cond2', 'cond3']
        - Overlap: True, Direction: same
    [11/20] Generating: 2_4Cond_NonOverlap_randomDir_InputData.feather
        - Available conditions: ['control', 'cond1', 'cond2', 'cond3']
        - Conditions to perturb: ['cond2', 'cond3']
        - Overlap: False, Direction: random
    [12/20] Generating: 2_4Cond_NonOverlap_sameDir_InputData.feather
        - Available conditions: ['control', 'cond1', 'cond2', 'cond3']
        - Conditions to perturb: ['cond2', 'cond3']
        - Overlap: False, Direction: same

--- Processing 5 conditions with shifts: [0, 0.5, 1, 1.5, 2] ---
    [13/20] Generating: 2_5Cond_Overlap_randomDir_InputData.feather
        - Available conditions: ['control', 'cond1', 'cond2', 'cond3', 'cond4']
        - Conditions to perturb: ['cond3', 'cond4']
        - Overlap: True, Direction: random
    [14/20] Generating: 2_5Cond_Overlap_sameDir_InputData.feather
        - Available conditions: ['control', 'cond1', 'cond2', 'cond3', 'cond4']
        - Conditions to perturb: ['cond3', 'cond4']
        - Overlap: True, Direction: same
    [15/20] Generating: 2_5Cond_NonOverlap_randomDir_InputData.feather
        - Available conditions: ['control', 'cond1', 'cond2', 'cond3', 'cond4']
        - Conditions to perturb: ['cond3', 'cond4']
        - Overlap: False, Direction: random
    [16/20] Generating: 2_5Cond_NonOverlap_sameDir_InputData.feather
        - Available conditions: ['control', 'cond1', 'cond2', 'cond3', 'cond4']
        - Conditions to perturb: ['cond3', 'cond4']
        - Overlap: False, Direction: same

--- Processing 6 conditions with shifts: [0, 0.5, 1, 1.5, 2, 2.5] ---
    [17/20] Generating: 2_6Cond_Overlap_randomDir_InputData.feather
        - Available conditions: ['control', 'cond1', 'cond2', 'cond3', 'cond4', 'cond5']
        - Conditions to perturb: ['cond4', 'cond5']
        - Overlap: True, Direction: random
    [18/20] Generating: 2_6Cond_Overlap_sameDir_InputData.feather
        - Available conditions: ['control', 'cond1', 'cond2', 'cond3', 'cond4', 'cond5']
        - Conditions to perturb: ['cond4', 'cond5']
        - Overlap: True, Direction: same
    [19/20] Generating: 2_6Cond_NonOverlap_randomDir_InputData.feather
        - Available conditions: ['control', 'cond1', 'cond2', 'cond3', 'cond4', 'cond5']
        - Conditions to perturb: ['cond4', 'cond5']
        - Overlap: False, Direction: random
    [20/20] Generating: 2_6Cond_NonOverlap_sameDir_InputData.feather
        - Available conditions: ['control', 'cond1', 'cond2', 'cond3', 'cond4', 'cond5']
        - Conditions to perturb: ['cond4', 'cond5']
        - Overlap: False, Direction: same

--------------------------------------------------
• Completed generation of 20 versions for simulation: Sim4
    - Elapsed time: 00h:00m:36s
--------------------------------------------------
```

---

#### Summary: Generated Datasets¶

This notebook generated **61 synthetic datasets** across four simulation scenarios:

| Simulation | Focus | # Datasets | Output Location |
| --- | --- | --- | --- |
| **Sim1** | Imputation effects | 8 | `./data/Sim1/` |
| **Sim2** | Missingness tolerance | 25 | `./data/Sim2/` |
| **Sim3** | Detection sensitivity | 8 | `./data/Sim3/` |
| **Sim4** | Experimental complexity | 20 | `./data/Sim4/` |

##### Common Dataset Properties¶

- **Proteins**: 500 (250 perturbed, 250 unchanged)
- **Peptides per protein**: 5–50 (beta-binomial distribution)
- **Replicates**: 10 per condition
- **Ground truth**: Each dataset includes `isPerturbed` and `isOutlier` columns for benchmarking

##### Next Steps¶

The generated datasets are used by:

1. `02-runCOPF.R` — Run COPF method
2. `03-runPeCorA.R` — Run PeCorA method
3. `04-runProteoForge.py` — Run ProteoForge method

```
print("Notebook Execution Time:", utils.prettyTimer(utils.getTime() - startTime))
```

```
Notebook Execution Time: 00h:03m:04s
```
