## Supplementary material for "ProteoForge: An Imputation-Aware Framework for Differential Proteoform Discovery in Bottom-Up Proteomics": Analysis notebooks as html render: Notebook S5.html

### Notebook S5

###### Enes Kemal Ergin

#### 2025-12-08 23:00:36

### Supplementary Notebook 5: Peptide Identification Benchmarking on Simulated Datasets¶

- **License:** Creative Commons Attribution-NonCommercial 4.0 International License
- **Version:** 0.2
- **Edit Log:**
  - 2025-11-28: Initial version of the notebook
  - 2025-12-08: Revise the whole notebook, ensuring clarity and correctness

---

**Requirements:**

- Completed method runs: `02-runCOPF.R`, `03-runPeCorA.R`, `04-runProteoForge.py`
- Result files in `./data/Sim{1-4}/` directories

**Data Information:**  
This notebook processes method results from four simulation scenarios to evaluate **peptide identification** performance—the ability to correctly identify which peptides are perturbed.

| Simulation | Focus | Input Files Pattern |
| --- | --- | --- |
| **Sim1** | Complete vs. Imputed | `2_{pattern}_{type}_{method}_ResultData.feather` |
| **Sim2** | Missingness Levels | `2_Pro{rate}_Pep{rate}_imputed_{method}_ResultData.feather` |
| **Sim3** | Perturbation Magnitude | `2_{low}_{high}_{method}_ResultData.feather` |
| **Sim4** | Experimental Complexity | `2_{N}Cond_{overlap}_{dir}Dir_{method}_ResultData.feather` |

**Purpose:**  
Benchmark COPF, PeCorA, and ProteoForge on peptide identification using ROC curves, PR curves, and Matthews Correlation Coefficient (MCC) across multiple p-value thresholds.

---

#### Setup¶

This section imports libraries, configures display settings, and defines paths for outputs.

> **Note:** The HTML rendering of this notebook hides code cells by default. Click the "Code" buttons to expand them.

##### Libraries¶

```
import os
import sys
import warnings
warnings.filterwarnings("ignore")

import numpy as np # Numerical computing
import pandas as pd # Data manipulatio

import seaborn as sns # R-like high-level plots
import matplotlib.pyplot as plt # Python's base plotting 

sys.path.append('../')

# Utility imports for this analysis
from src import utils, plots

# Initialize the timer
startTime = utils.getTime()
```

##### Display Settings¶

The cell below configures pandas, matplotlib, and seaborn display options for improved readability of tables and figures, including color palettes and figure export settings.

```
## Figure Settings

# Define default colors and styles for plots
def_colors = [
    "#139593", "#fca311", "#e54f2a",
    "#c3c3c3", "#555555",
    "#690000", "#5f4a00", "#004549"
]

# Set seaborn style
sns.set_theme(
    style="white",
    context="paper",
    palette=def_colors,
    font_scale=1,
    rc={
        "figure.figsize": (6, 4),
        "font.family": "sans-serif",
        "font.sans-serif": ["Arial", "Ubuntu Mono"],
    }
)

# Figure Saving Settings
figure_formats = ["pdf"]
save_to_folder = True
transparent_bg = True
figure_dpi = 300

## Configure dataframe displaying
pd.set_option('display.float_format', lambda x: '%.4f' % x)
pd.set_option('display.max_rows', 50)
pd.set_option('display.max_columns', 50)
pd.set_option('display.width', 500)  # Set a wider display width

## Printing Settings
verbose = True

## Show the color-palettes
plots.color_palette( def_colors, save=False )
```

##### Data and Result Paths¶

Defines input/output directory structure:

- **Input**: `./data/Sim{N}/` — Method result files from each simulation
- **Output**: Processed performance data and figures saved to same directories

```
# Establish paths and 
notebook_name = "05-IdBenchmark"
input_path = f"./data/"
mainFig_path = f"./figures/"

def setup_simulation_paths( simID ):
    """
    Create output and figure directories for a simulation, if they do not exist. 
    (Uses global variable for save_to_folder and figure_formats)

    Args:
        simID (str): Simulation ID.
        
        (Global Variables)
            input_path (str): Base path for input data.
            mainFig_path (str): Base path for figures.
            save_to_folder (bool): Whether to save figures to folders.
            figure_formats (list): List of formats to save figures in.
    
    Returns:
        tuple: (output_path, figure_path)
    """
    output_path = f"{input_path}{simID}/"
    if not os.path.exists(output_path):
        os.makedirs(output_path)

    figure_path = f"{mainFig_path}{simID}/"
    if save_to_folder:
        for fmt in figure_formats:
            cur_folder = os.path.join(figure_path, fmt)
            if not os.path.exists(cur_folder):
                os.makedirs(cur_folder)
    return output_path, figure_path
```

##### Global Variables¶

Defines analysis constants and visualization styling:

- **`seed`**, **`pthr`**: Reproducibility seed and p-value threshold (10⁻³)
- **`thresholds`**: Range of p-value thresholds for curve generation
- **`method_palette`**, **`method_markers`**: Consistent styling for COPF, PeCorA, ProteoForge
- **`mcc_thresholds`**, **`mcc_colors`**: MCC interpretation scale (Random → Almost Perfect)

```
# Global variables
seed = 42 # Seed for reproducibility  
pthr = 10**-3  # p-value threshold for significance
thresholds = list(utils.generate_thresholds(10.0, -15, 1, 0, 1, 0.1)) # Thresholds for the analysis

# Methods and their plotting styles
method_palette = {
    "COPF": "#139593",
    "PeCorA": "#fca311",
    "ProteoForge": "#e54f2a",
}
method_styles = {
    "COPF": "--",
    "PeCorA": "-.",
    "ProteoForge": ":",
}
method_markers = {
    "COPF": "o",
    "PeCorA": "s",
    "ProteoForge": "^",
}


# Matthews Correlation Coefficient (MCC) thresholds and colors
mcc_thresholds = {
    0.0 : 'Random',
    0.3 : 'Fair (0.3)',
    0.5 : 'Moderate (0.5)',
    0.7 : 'Strong (0.7)',
    0.9 : 'Almost Perfect (0.9)',
}
mcc_colors = {
    'Random': '#c3c3c3',             
    'Fair (0.3)': '#e0aaff',         
    'Moderate (0.5)': '#9d4edd',     
    'Strong (0.7)': '#5a189a',       
    'Almost Perfect (0.9)': '#240046' 
}

plots.color_palette(mcc_colors, save=False, name="MCC Interpretation Colors")
```


---

#### Simulation 1: Complete vs. Imputed Data Comparison¶

**Objective:** Compare peptide identification performance between complete (fully quantified) and imputed datasets across different perturbation patterns.

**Input Data:** `./data/Sim1/2_{pattern}_{complete|imputed}_{method}_ResultData.feather`

**Perturbation Patterns:**

- **twoPep**: 2 peptides perturbed per protein
- **randomPep**: Random 10–50% of peptides perturbed
- **halfPep**: 50% of peptides perturbed
- **halfPlusPep**: 70% of peptides perturbed

**Metrics Generated:**

- ROC curves (FPR vs TPR) with AUC
- PR curves (Recall vs Precision) with AUC
- MCC across p-value thresholds

**Key Question:** Does imputation affect method performance compared to complete data?

##### Data Processing and Metric Calculation¶

```
stTime = utils.getTime()

simID = "Sim1"  
# Set up a path for the simulation
output_path, figure_path = setup_simulation_paths( simID )

methods = list(method_palette.keys())
dataTypes = ['complete', 'imputed']
experiments = ['twoPep', 'randomPep', 'halfPep', 'halfPlusPep']
experiment_mapper = {
    'twoPep': 'Two Peptides', 
    'randomPep': '2>50% Peptides',
    'halfPep': '50% Peptides',
    'halfPlusPep': '>50% Peptides',
}

# =============================================================================
# ANALYSIS SETUP
# =============================================================================
print("PEPTIDE IDENTIFICATION BENCHMARK ANALYSIS")
print("-" * 52)
print(f"Simulation ID: {simID}")
print(f"Methods: {', '.join(methods)} ({len(methods)} total)")
print(f"Data Types: {', '.join(dataTypes)}")
print(f"Experiments: {len(experiments)} scenarios")
print(f"Output Path: {output_path}")

# Progress tracking
total_combinations = len(methods) * len(dataTypes) * len(experiments)
current_combination = 0
files_found = 0
files_missing = 0

print(f"Total combinations to process: {total_combinations}")
print("-" * 52)
# =============================================================================
# DATA PROCESSING
# =============================================================================
display("Calculating ID Benchmarks:")

combined_results = []
for cur_method in methods:
    for cur_type in dataTypes:
        for cur_exp in experiments:
            current_combination += 1
            
            # Load the result data
            result_file = f"{output_path}2_{cur_exp}_{cur_type}_{cur_method}_ResultData.feather"
            
            if os.path.exists(result_file):
                res_df = pd.read_feather(result_file)
                files_found += 1
                
                # Show progress for successful loads only
                if verbose:
                    progress = f"[{current_combination:2d}/{total_combinations}]"
                    print(f"{progress} ✓ {cur_method:12} | {cur_type:9} | {experiment_mapper[cur_exp]}")
            else:
                files_missing += 1
                if verbose:
                    progress = f"[{current_combination:2d}/{total_combinations}]"
                    print(f"{progress} ✗ {cur_method:12} | {cur_type:9} | {experiment_mapper[cur_exp]} (MISSING)")
                continue
            
            # Handle COPF-specific column renaming
            if cur_method == "COPF":
                res_df = res_df.rename(columns={
                    "proteoform_score_pval": "adj_pval",
                    'protein_id': "Protein"
                })

            # Create metric data
            metric_data = utils.create_metric_data(
                res_df, 
                pvalue_thresholds=thresholds,
                label_col='pertPeptide',
                pvalue_col='adj_pval'
            )
            metric_data['Method'] = cur_method
            metric_data['DataType'] = cur_type
            metric_data['Experiment'] = cur_exp
            combined_results.append(metric_data)

# Combine and process data
idBenchmarkData = pd.concat(combined_results, ignore_index=True)
idBenchmarkData['Experiment'] = idBenchmarkData['Experiment'].map(experiment_mapper)

# Categorical ordering ensure twoPep, randomPep, halfPep, halfPlusPep
idBenchmarkData['Experiment'] = pd.Categorical(
    idBenchmarkData['Experiment'], 
    categories=[experiment_mapper[exp] for exp in experiments],
    ordered=True
)

print(f"\nDATA PREVIEW:")
display(idBenchmarkData.tail(5))

# Save the processed data
output_file = f"{output_path}4_{simID}_PeptideIdentification_PerformanceData.feather"
idBenchmarkData.to_feather(output_file)
idBenchmarkData.to_csv(output_file.replace('.feather', '.csv'), index=False)

print(f"\nRESULTS SUMMARY:")
print("=" * 52)
execTime = utils.prettyTimer(utils.getTime() - stTime)
print(f"Total execution time:         {execTime}")
print(f"Files processed successfully: {files_found}")
print(f"Files missing/skipped:        {files_missing}")
print(f"Processing success rate:      {files_found/(files_found+files_missing)*100:.1f}%")
print("-" * 52)
print(f"Final dataset shape:          {idBenchmarkData.shape}")
print(f"Unique experiments:           {idBenchmarkData['Experiment'].nunique()}")
print(f"Methods analyzed:             {idBenchmarkData['Method'].nunique()}")
print(f"Data types included:          {idBenchmarkData['DataType'].nunique()}")
print(f"Data saved to: {output_file}")
print("=" * 52)
```

```
PEPTIDE IDENTIFICATION BENCHMARK ANALYSIS
----------------------------------------------------
Simulation ID: Sim1
Methods: COPF, PeCorA, ProteoForge (3 total)
Data Types: complete, imputed
Experiments: 4 scenarios
Output Path: ./data/Sim1/
Total combinations to process: 24
----------------------------------------------------
```

```
'Calculating ID Benchmarks:'
```

```
[ 1/24] ✓ COPF         | complete  | Two Peptides
[ 2/24] ✓ COPF         | complete  | 2>50% Peptides
[ 3/24] ✓ COPF         | complete  | 50% Peptides
[ 4/24] ✓ COPF         | complete  | >50% Peptides
[ 5/24] ✓ COPF         | imputed   | Two Peptides
[ 6/24] ✓ COPF         | imputed   | 2>50% Peptides
[ 7/24] ✓ COPF         | imputed   | 50% Peptides
[ 8/24] ✓ COPF         | imputed   | >50% Peptides
[ 9/24] ✓ PeCorA       | complete  | Two Peptides
[10/24] ✓ PeCorA       | complete  | 2>50% Peptides
[11/24] ✓ PeCorA       | complete  | 50% Peptides
[12/24] ✓ PeCorA       | complete  | >50% Peptides
[13/24] ✓ PeCorA       | imputed   | Two Peptides
[14/24] ✓ PeCorA       | imputed   | 2>50% Peptides
[15/24] ✓ PeCorA       | imputed   | 50% Peptides
[16/24] ✓ PeCorA       | imputed   | >50% Peptides
[17/24] ✓ ProteoForge  | complete  | Two Peptides
[18/24] ✓ ProteoForge  | complete  | 2>50% Peptides
[19/24] ✓ ProteoForge  | complete  | 50% Peptides
[20/24] ✓ ProteoForge  | complete  | >50% Peptides
[21/24] ✓ ProteoForge  | imputed   | Two Peptides
[22/24] ✓ ProteoForge  | imputed   | 2>50% Peptides
[23/24] ✓ ProteoForge  | imputed   | 50% Peptides
[24/24] ✓ ProteoForge  | imputed   | >50% Peptides

DATA PREVIEW:
```

|  | TP | FP | TN | FN | TPR | FPR | FDR | MCC | Precision | Recall | F1 | threshold | Method | DataType | Experiment |
| --- | --- | --- | --- | --- | --- | --- | --- | --- | --- | --- | --- | --- | --- | --- | --- |
| 595 | 892 | 888 | 2591 | 1312 | 0.4047 | 0.2552 | 0.4989 | 0.1570 | 0.5011 | 0.4047 | 0.4478 | 0.6000 | ProteoForge | imputed | >50% Peptides |
| 596 | 908 | 911 | 2568 | 1296 | 0.4120 | 0.2619 | 0.5008 | 0.1568 | 0.4992 | 0.4120 | 0.4514 | 0.7000 | ProteoForge | imputed | >50% Peptides |
| 597 | 932 | 939 | 2540 | 1272 | 0.4229 | 0.2699 | 0.5019 | 0.1586 | 0.4981 | 0.4229 | 0.4574 | 0.8000 | ProteoForge | imputed | >50% Peptides |
| 598 | 945 | 958 | 2521 | 1259 | 0.4288 | 0.2754 | 0.5034 | 0.1584 | 0.4966 | 0.4288 | 0.4602 | 0.9000 | ProteoForge | imputed | >50% Peptides |
| 599 | 2204 | 3479 | 0 | 0 | 1.0000 | 1.0000 | 0.6122 | 0.0000 | 0.3878 | 1.0000 | 0.5589 | 1.0000 | ProteoForge | imputed | >50% Peptides |

```
RESULTS SUMMARY:
====================================================
Total execution time:         00h:00m:03s
Files processed successfully: 24
Files missing/skipped:        0
Processing success rate:      100.0%
----------------------------------------------------
Final dataset shape:          (600, 15)
Unique experiments:           4
Methods analyzed:             3
Data types included:          2
Data saved to: ./data/Sim1/4_Sim1_PeptideIdentification_PerformanceData.feather
====================================================
```

##### ROC Curves: Discrimination Ability¶

ROC curves show the trade-off between true positive rate (sensitivity) and false positive rate (1-specificity). AUC values indicate overall discrimination ability—higher is better. Black circles mark performance at the significance threshold (p < 10⁻³).

```
# ROC Curves for different perturbations and methods
fig, axes = plt.subplots(
    2, 4, figsize=(16, 6), 
    sharey=True, sharex=True,
    gridspec_kw={
        "wspace": 0.05, 
        "hspace": 0.05
    }
)

for i, pert in enumerate(idBenchmarkData['Experiment'].unique()):
    for j, dataType in enumerate(dataTypes):
        cur_data = idBenchmarkData[
            (idBenchmarkData["Experiment"] == pert) & 
            (idBenchmarkData["DataType"] == dataType)
        ]
        # Ensure the data is complete for ROC curves
        cur_data = cur_data.groupby("Method").apply(
            lambda x: utils.complete_curve_data(x, 'ROC', 'FPR', 'TPR')
        ).reset_index(drop=True)
        
        # Calculate AUC per method from TPR and FPR
        auc_data = cur_data.groupby("Method").apply(
            lambda x: np.trapezoid(
                x.sort_values("FPR")["TPR"], x.sort_values("FPR")["FPR"]
            ), 
            include_groups=False
        )
        # Plot the ROC curve
        sns.lineplot(
            data=cur_data,
            x="FPR",
            y="TPR",
            hue="Method",
            style="Method",
            palette=method_palette,
            ax=axes[j, i],
            # Show the points
            markers=method_markers,
            dashes=False,
            markersize=7.5,
            markeredgewidth=0.5,
            markeredgecolor="gray", 
            legend=False,
            rasterized=True,
            estimator=None
        )
        # Add AUC values as legend like text
        for k, method in enumerate(auc_data.index):
            auc = auc_data[method]
            # Add color to match the palette
            color = method_palette[method]
            axes[j, i].text(
                0.95,
                0.05 + k * 0.075,
                f"{method}: {auc:.2f}",
                color=color,
                transform=axes[j, i].transAxes,
                ha="right",
                va="bottom",
                fontsize=12,
                fontweight="bold",
            )
            # Add a large no-facecolor marker to the pthr value for each method
            pthr_data = cur_data[(cur_data["Method"] == method) & (cur_data["threshold"] == pthr)]
            axes[j, i].scatter(
                pthr_data["FPR"],
                pthr_data["TPR"],
                color=color,
                s=125,
                edgecolor="black",
                linewidth=1.25,
                marker="o",
                facecolors="none",
            )
        
        # Add the diagonal line
        axes[j, i].plot([0, 1], [0, 1], color="black", linestyle="--")
        # axes[j, i].set_title(f"{pert}", fontsize=14, fontweight="bold")
        axes[j, i].set_xlabel("FPR", fontsize=12)
        axes[j, i].set_ylabel("TPR", fontsize=12)
        axes[j, i].set_xlim(-0.05, 1.05)
        axes[j, i].set_ylim(-0.05, 1.05)
        axes[j, i].grid("both", linestyle="--", linewidth=0.75, alpha=0.5, color="lightgrey")
        # If top row set the title
        if j == 0:
            axes[j, i].set_title(f"{pert}", fontsize=14, fontweight="bold")
        # For the last column, set the text for the dataType on the right
        if i == 3:
            axes[j, i].text(
                1.05,
                0.5,
                f"{dataType.capitalize()}",
                transform=axes[j, i].transAxes,
                ha="left",
                va="center",
                fontsize=14,
                fontweight="bold",
                rotation=270,
            )

# Add a text indicating the circle is the p-value threshold under fig.title and above the first subplot
fig.text(
    0.9,
    .97,
    f"Black circles represent the TPR and FPR values at p-value threshold of {pthr}",
    ha="right",
    va="center",
    fontsize=12,
    fontstyle="italic",
    color="gray",
)

# Finalize the plot
plt.tight_layout()
plots.finalize_plot(
    fig, 
    show=True,
    save=save_to_folder,
    filename=f"{simID}_PepIDBenchmark_ROC_Curve", 
    filepath=figure_path,
    formats=figure_formats, 
    transparent=transparent_bg,
    dpi=figure_dpi,
)
```

##### Precision-Recall Curves: Performance Under Class Imbalance¶

PR curves are particularly informative when positive cases (perturbed peptides) are rare. Higher AUC-PR indicates better precision at various recall levels. Black circles mark the significance threshold.

```
# PR Curves for different perturbations and methods
fig, axes = plt.subplots(
    2, 4, figsize=(16, 6), 
    sharey=True, sharex=True,
    gridspec_kw={
        "wspace": 0.05, 
        "hspace": 0.05
    }
)

for i, pert in enumerate(idBenchmarkData['Experiment'].unique()):
    for j, dataType in enumerate(dataTypes):
        cur_data = idBenchmarkData[
            (idBenchmarkData["Experiment"] == pert) & 
            (idBenchmarkData["DataType"] == dataType)
        ]
        
        # Add a small value to avoid division by zero
        cur_data["Precision"] = cur_data["TP"] / (cur_data["TP"] + cur_data["FP"] + 1e-6)
        cur_data["Recall"] = cur_data["TP"] / (cur_data["TP"] + cur_data["FN"] + 1e-6)
        
        cur_data = cur_data.sort_values("Recall", ascending=False)
        cur_data = cur_data[~((cur_data["Recall"] == 0) & (cur_data["Precision"] == 0))]

        # Ensure the data is complete for PR curves
        cur_data = cur_data.groupby("Method").apply(
            lambda x: utils.complete_curve_data(x, 'PR', 'Recall', 'Precision')
        ).reset_index(drop=True)
        
        # Calculate AUC per method from Precision and Recall
        # Here we use the trapezoidal rule to calculate the area under the curve
        f1_data = cur_data.groupby("Method").apply(
            lambda x: np.trapezoid(
                x.sort_values("Recall")["Precision"], x.sort_values("Recall")["Recall"]
            ),
            include_groups=False
        )

        # Plot the PR curve
        sns.lineplot(
            data=cur_data,
            x="Recall",
            y="Precision",
            hue="Method",
            style="Method",
            palette=method_palette,
            ax=axes[j, i],
            # Show the points
            markers=method_markers,
            dashes=False,
            markersize=7.5,
            markeredgewidth=0.5,
            markeredgecolor="gray", 
            legend=False,
            rasterized=True,
            estimator=None
        )

        # Add AUC values as legend like text
        for k, method in enumerate(f1_data.index):
            f1 = f1_data[method]
            # Add color to match the palette
            color = method_palette[method]
            axes[j, i].text(
                0.05,
                0.05 + k * 0.075,
                f"{method}: {f1:.2f}",
                color=color,
                transform=axes[j, i].transAxes,
                ha="left",
                va="bottom",
                fontsize=12,
                fontweight="bold",
            )
            # Add a large no-facecolor marker to the pthr value for each method
            pthr_data = cur_data[(cur_data["Method"] == method) & (cur_data["threshold"] == pthr)]
            axes[j, i].scatter(
                pthr_data["Recall"],
                pthr_data["Precision"],
                color=color,
                s=125,
                edgecolor="black",
                linewidth=1.25,
                marker="o",
                facecolors="none",
            )

        axes[j, i].set_xlabel("Recall", fontsize=12)
        axes[j, i].set_ylabel("Precision", fontsize=12)
        axes[j, i].set_xlim(-0.05, 1.05)
        axes[j, i].set_ylim(-0.05, 1.05)
        axes[j, i].grid("both", linestyle="--", linewidth=0.75, alpha=0.5, color="lightgrey")
        # If top row set the title
        if j == 0:
            axes[j, i].set_title(f"{pert}", fontsize=14, fontweight="bold")
        # For the last column, set the text for the dataType on the right
        if i == 3:
            axes[j, i].text(
                1.05,
                0.5,
                f"{dataType.capitalize()}",
                transform=axes[j, i].transAxes,
                ha="left",
                va="center",
                fontsize=14,
                fontweight="bold",
                rotation=270,
            )

# Add a text indicating the circle is the p-value threshold under fig.title and above the first subplot
fig.text(
    0.9,
    .97,
    f"Black circles represent the Precision and Recall values at p-value threshold of {pthr}",
    ha="right",
    va="center",
    fontsize=12,
    fontstyle="italic",
    color="gray",
)
# Finalize the plot
plt.tight_layout()
plots.finalize_plot(
    fig, 
    show=True,
    save=save_to_folder,
    filename=f"{simID}_PepIDBenchmark_PR_Curve", 
    filepath=figure_path,
    formats=figure_formats, 
    transparent=transparent_bg,
    dpi=figure_dpi,
)
```

##### MCC Across Thresholds: Optimal Threshold Selection¶

Matthews Correlation Coefficient balances all confusion matrix elements. The curves show how MCC varies with p-value threshold. Black circles mark maximum MCC achieved.

```
best_thresholds = idBenchmarkData.groupby(["Method", "Experiment", "DataType"]).apply(
    lambda x: x.loc[x["MCC"].idxmax(), ["threshold", "MCC"]]
).reset_index()
best_thresholds.columns = ["Method", "Experiment", "DataType", "threshold", "MCC"]
# print("\nBest thresholds per method and experiment:")
# display(best_thresholds)

# MCC Curves for p-value thresholds
fig, axes = plt.subplots(
    2, 4, figsize=(16, 6), 
    sharey=True, sharex=True,
    gridspec_kw={
        "wspace": 0.05, 
        "hspace": 0.05
    }
)
for i, pert in enumerate(idBenchmarkData['Experiment'].unique()):
    for j, dataType in enumerate(dataTypes):
        cur_data = idBenchmarkData[
            (idBenchmarkData["Experiment"] == pert) & 
            (idBenchmarkData["DataType"] == dataType)
        ].copy()
        cur_data['-log10(threshold)'] = -np.log10(cur_data['threshold'])
        # Plot the MCC curve
        sns.lineplot(
            data=cur_data,
            x="-log10(threshold)",
            y="MCC",
            hue="Method",
            style="Method",
            palette=method_palette,
            ax=axes[j, i],
            # Show the points
            markers=method_markers,
            dashes=False,
            markersize=7.5,
            markeredgewidth=0.5,
            markeredgecolor="gray", 
            legend=False,
            rasterized=True,
            estimator=None
        )
        # Add the best MCC point for each method
        for k, method in enumerate(cur_data["Method"].unique()):
            best_data = best_thresholds[
                (best_thresholds["Method"] == method) & 
                (best_thresholds["Experiment"] == pert) & 
                (best_thresholds["DataType"] == dataType)
            ]
            if not best_data.empty:
                best_threshold = best_data["threshold"].values[0]
                best_mcc = best_data["MCC"].values[0]
                axes[j, i].scatter(
                    -np.log10(best_threshold),
                    best_mcc,
                    color=method_palette[method],
                    s=125,
                    edgecolor="black",
                    linewidth=1.25,
                    marker="o",
                    facecolors="none",
                )
                # # Annotate the best MCC point
                # Add color to match the palette
                color = method_palette[method]
                axes[j, i].text(
                    0.95,
                    0.75 + k * 0.075,
                    f"{method}: {best_mcc:.2f}",
                    color=color,
                    ha="right",
                    va="bottom",
                    fontsize=12,
                    fontweight="bold",
                    transform=axes[j, i].transAxes,
                )

        axes[j, i].set_xlabel("-log10(p-value threshold)", fontsize=12)
        axes[j, i].set_ylabel("MCC", fontsize=12)
        axes[j, i].set_xlim(-0.5, 15.5)
        axes[j, i].set_ylim(-0.25, 1.05)
        axes[j, i].grid("both", linestyle="--", linewidth=0.75, alpha=0.5, color="lightgrey")
        # Add horizontal line at y=0
        # axes[j, i].axhline(0, color="black", linestyle="--", linewidth=1)
        # # Draw MCC interpretation thresholds
        # for thresh, label in mcc_thresholds.items():
        #     axes[j, i].axhline(
        #         thresh, color=mcc_colors[label], alpha=1,
        #         linestyle="dotted", linewidth=1.5, 
        #         label=label, zorder=0
        #         )
            # axes[j, i].text(
            #     0.01,
            #     thresh + 0.02,
            #     label,
            #     color=mcc_colors[label],
            #     ha="left",
            #     va="bottom",
            #     fontsize=10,
            #     fontweight="bold",
            #     transform=axes[j, i].transAxes,
            # )
        # If top row set the title
        if j == 0:
            axes[j, i].set_title(f"{pert}", fontsize=14, fontweight="bold")
        # For the last column, set the text for the dataType on the right
        if i == 3:
            axes[j, i].text(
                1.05,
                0.5,
                f"{dataType.capitalize()}",
                transform=axes[j, i].transAxes,
                ha="left",
                va="center",
                fontsize=14,
                fontweight="bold",
                rotation=270,
            )
# Add a text indicating the circle is the best MCC point under fig.title and above the first subplot
fig.text(
    0.9,
    .97,
    f"Black circles represent the maximum MCC values per method",
    ha="right",
    va="center",
    fontsize=12,
    fontstyle="italic",
    color="gray",
)
# Finalize the plot
plt.tight_layout()
plots.finalize_plot(
    fig, 
    show=True,
    save=save_to_folder,
    filename=f"{simID}_PepIDBenchmark_MCC_Curve", 
    filepath=figure_path,
    formats=figure_formats, 
    transparent=transparent_bg,
    dpi=figure_dpi,
)
```

##### MCC Summary: Barplot Comparison¶

Barplots compare mean MCC (with 95% CI) across methods and perturbation patterns. Scatter points show MCC at the specific p-value threshold (10⁻³). Values below bars indicate mean MCC scores.

```
# # MCC Score for different perturbations and methods at the best threshold

# Create subplots for MCC Score for different perturbations and methods at the best threshold
fig, axes = plt.subplots(
    1, 2, figsize=(12, 5), 
    sharey=True, sharex=True,
    gridspec_kw={"wspace": 0.05, "hspace": 0.1}
)

for i, dataType in enumerate(dataTypes):
    cur_data = idBenchmarkData[idBenchmarkData["DataType"] == dataType]
    sns.barplot(
        ax=axes[i],
        data=cur_data,
        x="Experiment",
        y="MCC",
        hue="Method",
        edgecolor="black",
        linewidth=0.8,
        errorbar="ci",  # Show confidence intervals across all thresholds
        capsize=0.05,
        err_kws={'linewidth': 1.5}
    )
    # Write the average MCC score for each method on the bottom of each bar
    for j, pert in enumerate(idBenchmarkData['Experiment'].unique()):
        for k, method in enumerate(method_palette.keys()):  # Changed from hardcoded list
            cur_score = cur_data[
                (cur_data["Experiment"] == pert) & 
                (cur_data["Method"] == method)
            ]["MCC"].mean()
            axes[i].text(
                j + k * 0.25 - 0.25,
                0 - 0.05,
                f"{cur_score:.2f}",
                color=method_palette[method],
                ha="center",
                va="top",
                fontsize=10,
                fontweight="bold",
                rotation=90,
            )


    # Get data specifically at the p-value threshold for point overlay
    pthr_data = cur_data[cur_data['threshold'] == pthr]
    # Add points for specific p-value threshold
    for j, method in enumerate(method_palette.keys()):
        method_data = pthr_data[pthr_data['Method'] == method]
        
        # Get x positions for this method's bars
        xlist = sorted(cur_data['Experiment'].unique())
        x_positions = []
        y_positions = []
        
        for k, x in enumerate(experiment_mapper.values()):
            mcc_at_pthr = method_data.loc[method_data['Experiment'] == x, 'MCC']
            if not mcc_at_pthr.empty:
                # Calculate x position (center of bar for this method)
                x_pos = k + (j - 1) * 0.27  # Adjust based on seaborn's bar positioning
                x_positions.append(x_pos)
                y_positions.append(mcc_at_pthr.iloc[0]) 
        
        # Add scatter points for this method at pthr
        axes[i].scatter(
            x_positions, 
            y_positions,
            color=method_palette[method],
            s=80,
            edgecolor='black',
            linewidth=1.25,
            # marker='o',
            marker=method_markers[method],
            zorder=10,
            alpha=0.9
        )

    # # Draw MCC interpretation thresholds
    # for thresh, label in mcc_thresholds.items():
    #     axes[i].axhline(
    #         thresh, color=mcc_colors[label], alpha=1,
    #         linestyle="dotted", linewidth=1.5, 
    #         label=label, zorder=0
    #         )
            
    axes[i].set_title(f"{dataType.capitalize()}", fontsize=14, fontweight="bold")
    axes[i].set_xlabel('')
    axes[i].set_ylim(-0.15, 1.0)
    axes[i].grid("y", linestyle="--", linewidth=0.75, alpha=0.5, color="lightgrey")
    axes[i].set_xticklabels(axes[i].get_xticklabels(), rotation=0, ha="center", fontsize=10)
    # If first subplot, set the ylabel
    if i == 0:
        axes[i].set_ylabel("MCC Score", fontsize=12)
        # Remove the legend for the first subplot
        axes[i].legend().set_visible(False)
    else:
        axes[i].set_ylabel("")
        axes[i].legend(
            loc='upper right', title="Method", fontsize=10, title_fontsize=12, 
            frameon=False, 
        )

# Add a text to clarify the various annotations of the plot
# Include the p-value threshold in the text
# Include the MCC interpretation color legend
fig.text(
    0.9,
    .97,
    f"Black circles represent the MCC values at p-value threshold of {pthr}\nMCC interpretation thresholds are indicated by dotted lines",
    ha="right",
    va="center",
    fontsize=12,
    fontstyle="italic",
    color="gray",
)

plt.tight_layout()

plots.finalize_plot(
    fig, 
    show=True,
    save=save_to_folder,
    filename=f"{simID}_PepIDBenchmark_MCC_Barplot", 
    filepath=figure_path,
    formats=figure_formats, 
    transparent=transparent_bg,
    dpi=figure_dpi,
)
```


---

#### Simulation 2: Missingness Level Impact¶

**Objective:** Quantify how increasing proportions of missing data affect peptide identification performance after imputation.

**Input Data:** `./data/Sim2/2_Pro{rate}_Pep{rate}_imputed_{method}_ResultData.feather`

**Experimental Design:**

- Protein missingness: 0%, 20%, 40%, 60%, 80%
- Peptide missingness: 0%, 20%, 40%, 60%, 80%
- Full factorial: 5 × 5 = 25 combinations per method

**Metrics Generated:**

- AUROC heatmaps (method × missingness)
- MCC heatmaps (mean, at p-threshold, maximum)

**Key Question:** At what missingness level does performance critically degrade?

##### Data Processing and Metric Calculation¶

```
stTime = utils.getTime()

simID = "Sim2"  
# Set up a path for the simulation
output_path, figure_path = setup_simulation_paths( simID )

methods = list(method_palette.keys())
missRatePros = [0, 0.2, 0.4, 0.6, 0.8]
missRatePeps = [0, 0.2, 0.4, 0.6, 0.8]
missRate_mapper = {
    0: '0%',
    0.2: '20%',
    0.4: '40%',
    0.6: '60%',
    0.8: '80%',
}

print("PEPTIDE IDENTIFICATION BENCHMARK ANALYSIS")
print("-" * 52)
print(f"Simulation ID: {simID}")
print(f"Methods: {', '.join(methods)} ({len(methods)} total)")
print(f"Protein Missing Rates: {', '.join(map(str, missRatePros))}")
print(f"Peptide Missing Rates: {', '.join(map(str, missRatePeps))}")
print(f"Output Path: {output_path}")

# Progress tracking
total_combinations = len(methods) * len(missRatePros) * len(missRatePeps)
current_combination = 0
files_found = 0
files_missing = 0
print(f"Total combinations to process: {total_combinations}")
print("-" * 52)

display("Calculating ID Benchmarks:")

combined_results = []
for i in missRatePros:
    for j in missRatePeps:
        for cur_method in methods:
            current_combination += 1
            # Load the result data
            result_file = f"{output_path}2_Pro{i}_Pep{j}_imputed_{cur_method}_ResultData.feather"

            if os.path.exists(result_file):
                res_df = pd.read_feather(result_file)
                files_found += 1

                if verbose:
                    progress = f"[{current_combination:2d}/{total_combinations}]"
                    print(f"{progress} ✓ {cur_method:12} | ProMissing:{i:.1f} | PepMissing:{j:.1f}")
            else:
                files_missing += 1
                if verbose:
                    progress = f"[{current_combination:2d}/{total_combinations}]"
                    print(f"{progress} ✗ {cur_method:12} | ProMissing:{i:.1f} | PepMissing:{j:.1f} (MISSING)")
                continue
            
            # Handle COPF-specific column renaming
            if cur_method == "COPF":
                res_df = res_df.rename(columns={
                    "proteoform_score_pval": "adj_pval",
                    'protein_id': "Protein"
                })

            # Create metric data
            metric_data = utils.create_metric_data(
                res_df, 
                pvalue_thresholds=thresholds,
                label_col='pertPeptide',
                pvalue_col='adj_pval'
            )
            metric_data['Method'] = cur_method
            metric_data['ProteinMissingness'] = i
            metric_data['PeptideMissingness'] = j

            combined_results.append(metric_data)
    
# Combine and process data
idBenchmarkData = pd.concat(combined_results, ignore_index=True)
# Save the processed data
output_file = f"{output_path}4_{simID}_PeptideIdentification_PerformanceData.feather"
idBenchmarkData.to_feather(output_file)
idBenchmarkData.to_csv(output_file.replace('.feather', '.csv'), index=False)

print(f"\nDATA PREVIEW:")
display(idBenchmarkData.tail(5))

print(f"\nRESULTS SUMMARY:")
print("=" * 52)
execTime = utils.prettyTimer(utils.getTime() - stTime)
print(f"Total execution time:         {execTime}")
print(f"Files processed successfully: {files_found}")
print(f"Files missing/skipped:        {files_missing}")
print(f"Processing success rate:      {files_found/(files_found+files_missing)*100:.1f}%")
print("-" * 52)
print(f"Final dataset shape:          {idBenchmarkData.shape}")
protein_levels = sorted(idBenchmarkData['ProteinMissingness'].unique())
peptide_levels = sorted(idBenchmarkData['PeptideMissingness'].unique())
methods_list = sorted(idBenchmarkData['Method'].unique())

print(f"Unique protein missingness levels: {len(protein_levels)} -> {protein_levels}")
print(f"Unique peptide missingness levels: {len(peptide_levels)} -> {peptide_levels}")
print(f"Methods analyzed: {idBenchmarkData['Method'].nunique()} -> {', '.join(methods_list)}")
print(f"Threshold levels included: {idBenchmarkData['threshold'].nunique()} (min={idBenchmarkData['threshold'].min()}, max={idBenchmarkData['threshold'].max()})")
print(f"Data saved to: {output_file}")
print("=" * 52)
```

```
PEPTIDE IDENTIFICATION BENCHMARK ANALYSIS
----------------------------------------------------
Simulation ID: Sim2
Methods: COPF, PeCorA, ProteoForge (3 total)
Protein Missing Rates: 0, 0.2, 0.4, 0.6, 0.8
Peptide Missing Rates: 0, 0.2, 0.4, 0.6, 0.8
Output Path: ./data/Sim2/
Total combinations to process: 75
----------------------------------------------------
```

```
'Calculating ID Benchmarks:'
```

```
[ 1/75] ✓ COPF         | ProMissing:0.0 | PepMissing:0.0
[ 2/75] ✓ PeCorA       | ProMissing:0.0 | PepMissing:0.0
[ 3/75] ✓ ProteoForge  | ProMissing:0.0 | PepMissing:0.0
[ 4/75] ✓ COPF         | ProMissing:0.0 | PepMissing:0.2
[ 5/75] ✓ PeCorA       | ProMissing:0.0 | PepMissing:0.2
[ 6/75] ✓ ProteoForge  | ProMissing:0.0 | PepMissing:0.2
[ 7/75] ✓ COPF         | ProMissing:0.0 | PepMissing:0.4
[ 8/75] ✓ PeCorA       | ProMissing:0.0 | PepMissing:0.4
[ 9/75] ✓ ProteoForge  | ProMissing:0.0 | PepMissing:0.4
[10/75] ✓ COPF         | ProMissing:0.0 | PepMissing:0.6
[11/75] ✓ PeCorA       | ProMissing:0.0 | PepMissing:0.6
[12/75] ✓ ProteoForge  | ProMissing:0.0 | PepMissing:0.6
[13/75] ✓ COPF         | ProMissing:0.0 | PepMissing:0.8
[14/75] ✓ PeCorA       | ProMissing:0.0 | PepMissing:0.8
[15/75] ✓ ProteoForge  | ProMissing:0.0 | PepMissing:0.8
[16/75] ✓ COPF         | ProMissing:0.2 | PepMissing:0.0
[17/75] ✓ PeCorA       | ProMissing:0.2 | PepMissing:0.0
[18/75] ✓ ProteoForge  | ProMissing:0.2 | PepMissing:0.0
[19/75] ✓ COPF         | ProMissing:0.2 | PepMissing:0.2
[20/75] ✓ PeCorA       | ProMissing:0.2 | PepMissing:0.2
[21/75] ✓ ProteoForge  | ProMissing:0.2 | PepMissing:0.2
[22/75] ✓ COPF         | ProMissing:0.2 | PepMissing:0.4
[23/75] ✓ PeCorA       | ProMissing:0.2 | PepMissing:0.4
[24/75] ✓ ProteoForge  | ProMissing:0.2 | PepMissing:0.4
[25/75] ✓ COPF         | ProMissing:0.2 | PepMissing:0.6
[26/75] ✓ PeCorA       | ProMissing:0.2 | PepMissing:0.6
[27/75] ✓ ProteoForge  | ProMissing:0.2 | PepMissing:0.6
[28/75] ✓ COPF         | ProMissing:0.2 | PepMissing:0.8
[29/75] ✓ PeCorA       | ProMissing:0.2 | PepMissing:0.8
[30/75] ✓ ProteoForge  | ProMissing:0.2 | PepMissing:0.8
[31/75] ✓ COPF         | ProMissing:0.4 | PepMissing:0.0
[32/75] ✓ PeCorA       | ProMissing:0.4 | PepMissing:0.0
[33/75] ✓ ProteoForge  | ProMissing:0.4 | PepMissing:0.0
[34/75] ✓ COPF         | ProMissing:0.4 | PepMissing:0.2
[35/75] ✓ PeCorA       | ProMissing:0.4 | PepMissing:0.2
[36/75] ✓ ProteoForge  | ProMissing:0.4 | PepMissing:0.2
[37/75] ✓ COPF         | ProMissing:0.4 | PepMissing:0.4
[38/75] ✓ PeCorA       | ProMissing:0.4 | PepMissing:0.4
[39/75] ✓ ProteoForge  | ProMissing:0.4 | PepMissing:0.4
[40/75] ✓ COPF         | ProMissing:0.4 | PepMissing:0.6
[41/75] ✓ PeCorA       | ProMissing:0.4 | PepMissing:0.6
[42/75] ✓ ProteoForge  | ProMissing:0.4 | PepMissing:0.6
[43/75] ✓ COPF         | ProMissing:0.4 | PepMissing:0.8
[44/75] ✓ PeCorA       | ProMissing:0.4 | PepMissing:0.8
[45/75] ✓ ProteoForge  | ProMissing:0.4 | PepMissing:0.8
[46/75] ✓ COPF         | ProMissing:0.6 | PepMissing:0.0
[47/75] ✓ PeCorA       | ProMissing:0.6 | PepMissing:0.0
[48/75] ✓ ProteoForge  | ProMissing:0.6 | PepMissing:0.0
[49/75] ✓ COPF         | ProMissing:0.6 | PepMissing:0.2
[50/75] ✓ PeCorA       | ProMissing:0.6 | PepMissing:0.2
[51/75] ✓ ProteoForge  | ProMissing:0.6 | PepMissing:0.2
[52/75] ✓ COPF         | ProMissing:0.6 | PepMissing:0.4
[53/75] ✓ PeCorA       | ProMissing:0.6 | PepMissing:0.4
[54/75] ✓ ProteoForge  | ProMissing:0.6 | PepMissing:0.4
[55/75] ✓ COPF         | ProMissing:0.6 | PepMissing:0.6
[56/75] ✓ PeCorA       | ProMissing:0.6 | PepMissing:0.6
[57/75] ✓ ProteoForge  | ProMissing:0.6 | PepMissing:0.6
[58/75] ✓ COPF         | ProMissing:0.6 | PepMissing:0.8
[59/75] ✓ PeCorA       | ProMissing:0.6 | PepMissing:0.8
[60/75] ✓ ProteoForge  | ProMissing:0.6 | PepMissing:0.8
[61/75] ✓ COPF         | ProMissing:0.8 | PepMissing:0.0
[62/75] ✓ PeCorA       | ProMissing:0.8 | PepMissing:0.0
[63/75] ✓ ProteoForge  | ProMissing:0.8 | PepMissing:0.0
[64/75] ✓ COPF         | ProMissing:0.8 | PepMissing:0.2
[65/75] ✓ PeCorA       | ProMissing:0.8 | PepMissing:0.2
[66/75] ✓ ProteoForge  | ProMissing:0.8 | PepMissing:0.2
[67/75] ✓ COPF         | ProMissing:0.8 | PepMissing:0.4
[68/75] ✓ PeCorA       | ProMissing:0.8 | PepMissing:0.4
[69/75] ✓ ProteoForge  | ProMissing:0.8 | PepMissing:0.4
[70/75] ✓ COPF         | ProMissing:0.8 | PepMissing:0.6
[71/75] ✓ PeCorA       | ProMissing:0.8 | PepMissing:0.6
[72/75] ✓ ProteoForge  | ProMissing:0.8 | PepMissing:0.6
[73/75] ✓ COPF         | ProMissing:0.8 | PepMissing:0.8
[74/75] ✓ PeCorA       | ProMissing:0.8 | PepMissing:0.8
[75/75] ✓ ProteoForge  | ProMissing:0.8 | PepMissing:0.8

DATA PREVIEW:
```

|  | TP | FP | TN | FN | TPR | FPR | FDR | MCC | Precision | Recall | F1 | threshold | Method | ProteinMissingness | PeptideMissingness |
| --- | --- | --- | --- | --- | --- | --- | --- | --- | --- | --- | --- | --- | --- | --- | --- |
| 1870 | 0 | 0 | 4745 | 1639 | 0.0000 | 0.0000 | 0.0000 | 0.0000 | 0.0000 | 0.0000 | 0.0000 | 0.6000 | ProteoForge | 0.8000 | 0.8000 |
| 1871 | 0 | 0 | 4745 | 1639 | 0.0000 | 0.0000 | 0.0000 | 0.0000 | 0.0000 | 0.0000 | 0.0000 | 0.7000 | ProteoForge | 0.8000 | 0.8000 |
| 1872 | 0 | 0 | 4745 | 1639 | 0.0000 | 0.0000 | 0.0000 | 0.0000 | 0.0000 | 0.0000 | 0.0000 | 0.8000 | ProteoForge | 0.8000 | 0.8000 |
| 1873 | 0 | 0 | 4745 | 1639 | 0.0000 | 0.0000 | 0.0000 | 0.0000 | 0.0000 | 0.0000 | 0.0000 | 0.9000 | ProteoForge | 0.8000 | 0.8000 |
| 1874 | 0 | 0 | 4745 | 1639 | 0.0000 | 0.0000 | 0.0000 | 0.0000 | 0.0000 | 0.0000 | 0.0000 | 1.0000 | ProteoForge | 0.8000 | 0.8000 |

```
RESULTS SUMMARY:
====================================================
Total execution time:         00h:00m:09s
Files processed successfully: 75
Files missing/skipped:        0
Processing success rate:      100.0%
----------------------------------------------------
Final dataset shape:          (1875, 15)
Unique protein missingness levels: 5 -> [np.float64(0.0), np.float64(0.2), np.float64(0.4), np.float64(0.6), np.float64(0.8)]
Unique peptide missingness levels: 5 -> [np.float64(0.0), np.float64(0.2), np.float64(0.4), np.float64(0.6), np.float64(0.8)]
Methods analyzed: 3 -> COPF, PeCorA, ProteoForge
Threshold levels included: 25 (min=0.0, max=1.0)
Data saved to: ./data/Sim2/4_Sim2_PeptideIdentification_PerformanceData.feather
====================================================
```

##### AUROC Heatmaps: Discrimination Across Missingness¶

Heatmaps show AUROC values for each method across the protein × peptide missingness grid. Higher values (darker blue) indicate better discrimination. The (0,0) corner represents complete data baseline.

```
# Comparing AUC Values across different missingness levels
# Calculate AUC for ROC and PR curves
auc_results = []
for (method, pro_miss, pep_miss), group in idBenchmarkData.groupby(
        ['Method', 'ProteinMissingness', 'PeptideMissingness']
    ):
    # ROC AUC
    group = utils.complete_curve_data(group, 'ROC', 'FPR', 'TPR')
    roc_data = group.sort_values('FPR')
    roc_auc = np.trapezoid(roc_data['TPR'], roc_data['FPR'])
    
    
    auc_results.append({
        'Method': method,
        'ProteinMissingness': pro_miss,
        'PeptideMissingness': pep_miss,
        'AUC': roc_auc,
    })
auc_df = pd.DataFrame(auc_results)
print("\nAUC Data Preview:")
display(auc_df.head())

# Initialize Figure for 3 Methods with dedicated colorbar column
fig, axes = plt.subplots(
    nrows=1, ncols=4, figsize=(12, 4), 
    gridspec_kw={
        "wspace": 0.05, "hspace": 0.1,
        "width_ratios": [0.9, 0.9, 0.9, 0.1]  # First 3 columns same size, 4th for colorbar
    },
)
vmin = auc_df["AUC"].min()
vmax = auc_df["AUC"].max()

# Create heatmaps for the first 3 methods
for i, cur_method in enumerate(methods):
    plot_data = auc_df[auc_df["Method"] == cur_method].pivot_table(
        index="PeptideMissingness",
        columns="ProteinMissingness",
        values="AUC",
        aggfunc="mean"
    )
    # Make sure 0,0 is in the bottom left corner
    plot_data = plot_data.iloc[::-1]
    
    # Only show colorbar on the last method heatmap
    cb = (i == len(methods) - 1)
    
    # If this is the last method, create colorbar in the 4th column
    if cb:
        cbar_ax = axes[3]
    else:
        cbar_ax = None
    
    sns.heatmap(
        plot_data,
        vmin=vmin,
        vmax=vmax,
        cmap="Blues",
        annot=True,
        fmt=".2f",
        ax=axes[i],
        cbar=cb,
        cbar_ax=cbar_ax,
        cbar_kws={"label": "AUROC"},
        square=True,
        # Annotation size
        annot_kws={"size": 12}, 
    )
    
    if i == 0:
        axes[i].set_ylabel("Peptide Missingness Rate", fontsize=14, fontweight="bold")
    else:
        axes[i].set_ylabel("")
        # Remove yticklabels
        axes[i].set_yticklabels([])
    if i == 1:
        axes[i].set_xlabel("Protein Missingness Rate", fontsize=14, fontweight="bold")
    else:
        axes[i].set_xlabel("")

    axes[i].set_title(f"{cur_method}", fontsize=14, fontweight="bold")

# Hide the 4th axis if it wasn't used for colorbar (this handles the case where we have < 3 methods)
if len(methods) < 3:
    axes[3].set_visible(False)

plt.tight_layout()
plt.suptitle("Peptide Identification Benchmark with AUROC", fontsize=16, fontweight="bold")
plt.subplots_adjust(top=0.85)   

plots.finalize_plot(
    fig, 
    show=True,
    save=save_to_folder,
    filename=f"{simID}_PepIDBenchmark_AUROC_Heatmap", 
    filepath=figure_path,
    formats=figure_formats, 
    transparent=transparent_bg,
    dpi=figure_dpi,
)
```

```
AUC Data Preview:
```

|  | Method | ProteinMissingness | PeptideMissingness | AUC |
| --- | --- | --- | --- | --- |
| 0 | COPF | 0.0000 | 0.0000 | 0.5901 |
| 1 | COPF | 0.0000 | 0.2000 | 0.5799 |
| 2 | COPF | 0.0000 | 0.4000 | 0.5783 |
| 3 | COPF | 0.0000 | 0.6000 | 0.5782 |
| 4 | COPF | 0.0000 | 0.8000 | 0.6050 |

##### Mean MCC Heatmaps: Average Performance¶

Shows mean MCC averaged across all p-value thresholds. Provides a threshold-agnostic view of overall method performance at each missingness combination.

```
# Mean MCC Scores
# Heatmap of the MCC values for the different methods 
# (Protein Missingness and Peptide Missingness) (X: Protein Missingness, Y: Peptide Missingness)

# Initalize Figure for 3 Methods
fig, axes = plt.subplots(
    nrows=1, ncols=4, figsize=(12, 4), 
    gridspec_kw={
        "wspace": 0.05, "hspace": 0.1,
        "width_ratios": [0.9, 0.9, 0.9, 0.1]  # First 3 columns same size, 4th for colorbar
    },
)

vmin = idBenchmarkData["MCC"].min()
vmax = idBenchmarkData["MCC"].max()

# Create heatmaps for the first 3 methods
for i, cur_method in enumerate(methods):
    plot_data = idBenchmarkData[idBenchmarkData["Method"] == cur_method].pivot_table(
        index="PeptideMissingness",
        columns="ProteinMissingness",
        values="MCC",
        aggfunc="mean"
    )
    # Make sure 0,0 is in the bottom left corner
    plot_data = plot_data.iloc[::-1]
    
    # Only show colorbar on the last method heatmap
    cb = (i == len(methods) - 1)
    
    # If this is the last method, create colorbar in the 4th column
    if cb:
        cbar_ax = axes[3]
    else:
        cbar_ax = None
    
    sns.heatmap(
        plot_data,
        vmin=vmin,
        vmax=vmax,
        cmap="Purples",
        annot=True,
        fmt=".2f",
        ax=axes[i],
        cbar=cb,
        cbar_ax=cbar_ax,
        cbar_kws={"label": "Mean MCC"},
        square=True,
        # Annotation size
        annot_kws={"size": 12}, 
    )
    
    if i == 0:
        axes[i].set_ylabel("Peptide Missingness Rate", fontsize=14, fontweight="bold")
    else:
        axes[i].set_ylabel("")
        # Remove yticklabels
        axes[i].set_yticklabels([])
    if i == 1:
        axes[i].set_xlabel("Protein Missingness Rate", fontsize=14, fontweight="bold")
    else:
        axes[i].set_xlabel("")

    axes[i].set_title(f"{cur_method}", fontsize=14, fontweight="bold")


# Hide the 4th axis if it wasn't used for colorbar (this handles the case where we have < 3 methods)
if len(methods) < 3:
    axes[3].set_visible(False)

plt.tight_layout()
plt.suptitle("Peptide Identification Benchmark with MCC (Mean)", fontsize=16, fontweight="bold")
plt.subplots_adjust(top=0.85)

plots.finalize_plot(
    fig, 
    show=True,
    save=save_to_folder,
    filename=f"{simID}_PepIDBenchmark_MCC_mean_Heatmap", 
    filepath=figure_path,
    formats=figure_formats, 
    transparent=transparent_bg,
    dpi=figure_dpi,
)
```

##### MCC at P-value Threshold: Practical Performance¶

Shows MCC specifically at the commonly used significance threshold (p < 10⁻³). This represents the expected performance when using standard statistical cutoffs.

```
# MCC Score at pThr
# Heatmap of the MCC values for the different methods 
# (Protein Missingness and Peptide Missingness) (X: Protein Missingness, Y: Peptide Missingness)

# Initalize Figure for 3 Methods
fig, axes = plt.subplots(
    nrows=1, ncols=4, figsize=(12, 4), 
    gridspec_kw={
        "wspace": 0.05, "hspace": 0.1,
        "width_ratios": [0.9, 0.9, 0.9, 0.1]  # First 3 columns same size, 4th for colorbar
    },
)
vmin = idBenchmarkData["MCC"].min()
vmax = idBenchmarkData["MCC"].max()

# Create heatmaps for the first 3 methods
for i, cur_method in enumerate(methods):
    plot_data = idBenchmarkData[
        (idBenchmarkData["Method"] == cur_method) & 
        (idBenchmarkData["threshold"] == pthr)
    ].pivot_table(
        index="PeptideMissingness",
        columns="ProteinMissingness",
        values="MCC",
        aggfunc="mean"
    )
    # Make sure 0,0 is in the bottom left corner
    plot_data = plot_data.iloc[::-1]
    
    # Only show colorbar on the last method heatmap
    cb = (i == len(methods) - 1)
    
    # If this is the last method, create colorbar in the 4th column
    if cb:
        cbar_ax = axes[3]
    else:
        cbar_ax = None
    
    sns.heatmap(
        plot_data,
        vmin=vmin,
        vmax=vmax,
        cmap="Purples",
        annot=True,
        fmt=".2f",
        ax=axes[i],
        cbar=cb,
        cbar_ax=cbar_ax,
        cbar_kws={"label": f"MCC at p-value {pthr}"},
        square=True,
        # Annotation size
        annot_kws={"size": 12}, 
    )
    
    if i == 0:
        axes[i].set_ylabel("Peptide Missingness Rate", fontsize=14, fontweight="bold")
    else:
        axes[i].set_ylabel("")
        # Remove yticklabels
        axes[i].set_yticklabels([])
    if i == 1:
        axes[i].set_xlabel("Protein Missingness Rate", fontsize=14, fontweight="bold")
    else:
        axes[i].set_xlabel("")

    axes[i].set_title(f"{cur_method}", fontsize=14, fontweight="bold")

# Hide the 4th axis if it wasn't used for colorbar (this handles the case where we have < 3 methods)
if len(methods) < 3:
    axes[3].set_visible(False)  

plt.tight_layout()
plt.suptitle(f"Peptide Identification Benchmark with MCC at p-value {pthr}", fontsize=16, fontweight="bold")
plt.subplots_adjust(top=0.85)   

plots.finalize_plot(
    fig, 
    show=True,
    save=save_to_folder,
    filename=f"{simID}_PepIDBenchmark_MCC_pThr_Heatmap", 
    filepath=figure_path,
    formats=figure_formats, 
    transparent=transparent_bg,
    dpi=figure_dpi,
)
```

##### Maximum MCC Heatmaps: Best Achievable Performance¶

Shows the maximum MCC achievable across all thresholds—the theoretical upper bound of performance if the optimal threshold were known.

```
# Maximum MCC Scores
# Heatmap of the MCC values for the different methods 
# (Protein Missingness and Peptide Missingness) (X: Protein Missingness, Y: Peptide Missingness)

# Initialize Figure for 3 Methods with dedicated colorbar column
fig, axes = plt.subplots(
    nrows=1, ncols=4, figsize=(12, 4), 
    gridspec_kw={
        "wspace": 0.05, "hspace": 0.1,
        "width_ratios": [0.9, 0.9, 0.9, 0.1]  # First 3 columns same size, 4th for colorbar
    },
)
vmin = idBenchmarkData["MCC"].min()
vmax = idBenchmarkData["MCC"].max()

# Create heatmaps for the first 3 methods
for i, cur_method in enumerate(methods):
    plot_data = idBenchmarkData[idBenchmarkData["Method"] == cur_method].pivot_table(
        index="PeptideMissingness",
        columns="ProteinMissingness",
        values="MCC",
        aggfunc="max"
    )
    # Make sure 0,0 is in the bottom left corner
    plot_data = plot_data.iloc[::-1]
    
    # Only show colorbar on the last method heatmap
    cb = (i == len(methods) - 1)
    
    # If this is the last method, create colorbar in the 4th column
    if cb:
        cbar_ax = axes[3]
    else:
        cbar_ax = None
    
    sns.heatmap(
        plot_data,
        vmin=vmin,
        vmax=vmax,
        cmap="Purples",
        annot=True,
        fmt=".2f",
        ax=axes[i],
        cbar=cb,
        cbar_ax=cbar_ax,
        cbar_kws={"label": "Max MCC"},
        square=True,
        # Annotation size
        annot_kws={"size": 12}, 
    )
    
    if i == 0:
        axes[i].set_ylabel("Peptide Missingness Rate", fontsize=14, fontweight="bold")
    else:
        axes[i].set_ylabel("")
        # Remove yticklabels
        axes[i].set_yticklabels([])
    if i == 1:
        axes[i].set_xlabel("Protein Missingness Rate", fontsize=14, fontweight="bold")
    else:
        axes[i].set_xlabel("")

    axes[i].set_title(f"{cur_method}", fontsize=14, fontweight="bold")


# Hide the 4th axis if it wasn't used for colorbar (this handles the case where we have < 3 methods)
if len(methods) < 3:
    axes[3].set_visible(False)

plt.tight_layout()
plt.suptitle("Peptide Identification Benchmark with MCC (Max)", fontsize=16, fontweight="bold")
plt.subplots_adjust(top=0.85)

plots.finalize_plot(
    fig, 
    show=True,
    save=save_to_folder,
    filename=f"{simID}_PepIDBenchmark_MCC_max_Heatmap", 
    filepath=figure_path,
    formats=figure_formats, 
    transparent=transparent_bg,
    dpi=figure_dpi,
)
```


---

#### Simulation 3: Perturbation Magnitude Sensitivity¶

**Objective:** Determine the minimum perturbation magnitude required for reliable peptide identification across methods.

**Input Data:** `./data/Sim3/2_{low}_{high}_{method}_ResultData.feather`

**Magnitude Ranges (log2 fold-change):**
| Range | Interpretation |
|-------|----------------|
| 0.10–0.25 | Subtle |
| 0.25–0.50 | Small |
| 0.50–0.75 | Moderate |
| 0.75–1.00 | Medium |
| 1.00–1.25 | Substantial |
| 1.25–1.50 | Large |
| 1.50–1.75 | Strong |
| 1.75–2.00 | Very strong |

**Metrics Generated:**

- MCC line plots across magnitude ranges
- AUROC heatmap (method × magnitude)

**Key Question:** What is the practical detection threshold for each method?

##### Data Processing and Metric Calculation¶

```
stTime = utils.getTime()

simID = "Sim3"  
# Set up a path for the simulation
output_path, figure_path = setup_simulation_paths( simID )

methods = list(method_palette.keys())
perturbationRanges = [
    (0.1, 0.25),
    (0.25, 0.50),
    (0.50, 0.75),
    (0.75, 1.0),
    (1.0, 1.25),
    (1.25, 1.50),
    (1.50, 1.75),
    (1.75, 2.0),
]

perturbation_mapper = {
    (0.1, 0.25): '0.1-0.25',
    (0.25, 0.50): '0.25-0.50',
    (0.50, 0.75): '0.50-0.75',
    (0.75, 1.0): '0.75-1.0',
    (1.0, 1.25): '1.0-1.25',
    (1.25, 1.50): '1.25-1.50',
    (1.50, 1.75): '1.50-1.75',
    (1.75, 2.0): '1.75-2.0',
}


print("PEPTIDE IDENTIFICATION BENCHMARK ANALYSIS")
print("-" * 52)
print(f"Simulation ID: {simID}")
print(f"Methods: {', '.join(methods)} ({len(methods)} total)")
print(f"Perturbation Ranges: {', '.join([f'{low}-{high}' for low, high in perturbationRanges])}")
print(f"Output Path: {output_path}")

# Progress tracking
total_combinations = len(methods) * len(perturbationRanges)
current_combination = 0
files_found = 0
files_missing = 0
print(f"Total combinations to process: {total_combinations}")
print("-" * 52)
display("Calculating ID Benchmarks:")

combined_results = []
for low, high in perturbationRanges:
    for cur_method in methods:
        current_combination += 1
        # Load the result data
        result_file = f"{output_path}2_{low}_{high}_{cur_method}_ResultData.feather"

        if os.path.exists(result_file):
            res_df = pd.read_feather(result_file)
            files_found += 1

            if verbose:
                progress = f"[{current_combination:2d}/{total_combinations}]"
                print(f"{progress} ✓ {cur_method:12} | Perturbation: {low:.2f}-{high:.2f}")
        else:
            files_missing += 1
            if verbose:
                progress = f"[{current_combination:2d}/{total_combinations}]"
                print(f"{progress} ✗ {cur_method:12} | Perturbation: {low:.2f}-{high:.2f} (MISSING)")
            continue
        
        # Handle COPF-specific column renaming
        if cur_method == "COPF":
            res_df = res_df.rename(columns={
                "proteoform_score_pval": "adj_pval",
                'protein_id': "Protein"
            })

        # Create metric data
        metric_data = utils.create_metric_data(
            res_df, 
            pvalue_thresholds=thresholds,
            label_col='pertPeptide',
            pvalue_col='adj_pval'
        )
        metric_data['Method'] = cur_method
        metric_data['PerturbationRange'] = f"{low:.2f}-{high:.2f}"

        combined_results.append(metric_data)

# Combine and process data
idBenchmarkData = pd.concat(combined_results, ignore_index=True)
# Save the processed data
output_file = f"{output_path}4_{simID}_PeptideIdentification_PerformanceData.feather"
idBenchmarkData.to_feather(output_file) 
idBenchmarkData.to_csv(output_file.replace('.feather', '.csv'), index=False)

print(f"\nDATA PREVIEW:")
display(idBenchmarkData.tail(5))

print(f"\nRESULTS SUMMARY:")
print("=" * 52)
execTime = utils.prettyTimer(utils.getTime() - stTime)
print(f"Total execution time:         {execTime}")
print(f"Files processed successfully: {files_found}")
print(f"Files missing/skipped:        {files_missing}")
print(f"Processing success rate:      {files_found/(files_found+files_missing)*100:.1f}%")
print("-" * 52)
print(f"Final dataset shape:          {idBenchmarkData.shape}")
pert_ranges = sorted(idBenchmarkData['PerturbationRange'].unique())
methods_list = sorted(idBenchmarkData['Method'].unique())
print(f"Unique perturbation ranges: {len(pert_ranges)} -> {', '.join(pert_ranges)}")
print(f"Methods analyzed: {idBenchmarkData['Method'].nunique()} -> {', '.join(methods_list)}")
print(f"Threshold levels included: {idBenchmarkData['threshold'].nunique()} (min={idBenchmarkData['threshold'].min()}, max={idBenchmarkData['threshold'].max()})")
print(f"Data saved to: {output_file}")
print("=" * 52)
```

```
PEPTIDE IDENTIFICATION BENCHMARK ANALYSIS
----------------------------------------------------
Simulation ID: Sim3
Methods: COPF, PeCorA, ProteoForge (3 total)
Perturbation Ranges: 0.1-0.25, 0.25-0.5, 0.5-0.75, 0.75-1.0, 1.0-1.25, 1.25-1.5, 1.5-1.75, 1.75-2.0
Output Path: ./data/Sim3/
Total combinations to process: 24
----------------------------------------------------
```

```
'Calculating ID Benchmarks:'
```

```
[ 1/24] ✓ COPF         | Perturbation: 0.10-0.25
[ 2/24] ✓ PeCorA       | Perturbation: 0.10-0.25
[ 3/24] ✓ ProteoForge  | Perturbation: 0.10-0.25
[ 4/24] ✓ COPF         | Perturbation: 0.25-0.50
[ 5/24] ✓ PeCorA       | Perturbation: 0.25-0.50
[ 6/24] ✓ ProteoForge  | Perturbation: 0.25-0.50
[ 7/24] ✓ COPF         | Perturbation: 0.50-0.75
[ 8/24] ✓ PeCorA       | Perturbation: 0.50-0.75
[ 9/24] ✓ ProteoForge  | Perturbation: 0.50-0.75
[10/24] ✓ COPF         | Perturbation: 0.75-1.00
[11/24] ✓ PeCorA       | Perturbation: 0.75-1.00
[12/24] ✓ ProteoForge  | Perturbation: 0.75-1.00
[13/24] ✓ COPF         | Perturbation: 1.00-1.25
[14/24] ✓ PeCorA       | Perturbation: 1.00-1.25
[15/24] ✓ ProteoForge  | Perturbation: 1.00-1.25
[16/24] ✓ COPF         | Perturbation: 1.25-1.50
[17/24] ✓ PeCorA       | Perturbation: 1.25-1.50
[18/24] ✓ ProteoForge  | Perturbation: 1.25-1.50
[19/24] ✓ COPF         | Perturbation: 1.50-1.75
[20/24] ✓ PeCorA       | Perturbation: 1.50-1.75
[21/24] ✓ ProteoForge  | Perturbation: 1.50-1.75
[22/24] ✓ COPF         | Perturbation: 1.75-2.00
[23/24] ✓ PeCorA       | Perturbation: 1.75-2.00
[24/24] ✓ ProteoForge  | Perturbation: 1.75-2.00

DATA PREVIEW:
```

|  | TP | FP | TN | FN | TPR | FPR | FDR | MCC | Precision | Recall | F1 | threshold | Method | PerturbationRange |
| --- | --- | --- | --- | --- | --- | --- | --- | --- | --- | --- | --- | --- | --- | --- |
| 595 | 871 | 455 | 4355 | 2 | 0.9977 | 0.0946 | 0.3431 | 0.7699 | 0.6569 | 0.9977 | 0.7922 | 0.6000 | ProteoForge | 1.75-2.00 |
| 596 | 871 | 483 | 4327 | 2 | 0.9977 | 0.1004 | 0.3567 | 0.7595 | 0.6433 | 0.9977 | 0.7822 | 0.7000 | ProteoForge | 1.75-2.00 |
| 597 | 871 | 507 | 4303 | 2 | 0.9977 | 0.1054 | 0.3679 | 0.7507 | 0.6321 | 0.9977 | 0.7739 | 0.8000 | ProteoForge | 1.75-2.00 |
| 598 | 871 | 540 | 4270 | 2 | 0.9977 | 0.1123 | 0.3827 | 0.7390 | 0.6173 | 0.9977 | 0.7627 | 0.9000 | ProteoForge | 1.75-2.00 |
| 599 | 873 | 4810 | 0 | 0 | 1.0000 | 1.0000 | 0.8464 | 0.0000 | 0.1536 | 1.0000 | 0.2663 | 1.0000 | ProteoForge | 1.75-2.00 |

```
RESULTS SUMMARY:
====================================================
Total execution time:         00h:00m:03s
Files processed successfully: 24
Files missing/skipped:        0
Processing success rate:      100.0%
----------------------------------------------------
Final dataset shape:          (600, 14)
Unique perturbation ranges: 8 -> 0.10-0.25, 0.25-0.50, 0.50-0.75, 0.75-1.00, 1.00-1.25, 1.25-1.50, 1.50-1.75, 1.75-2.00
Methods analyzed: 3 -> COPF, PeCorA, ProteoForge
Threshold levels included: 25 (min=0.0, max=1.0)
Data saved to: ./data/Sim3/4_Sim3_PeptideIdentification_PerformanceData.feather
====================================================
```

##### MCC Sensitivity Curves: Detection Threshold Analysis¶

Line plots show how MCC improves with increasing perturbation magnitude. Shaded bands represent 95% CI. Smaller markers indicate MCC at the specific p-value threshold. MCC interpretation thresholds (labeled boxes) provide performance context.

```
# Line plot showing MCC scores across perturbation ranges 
# line plot marker at the mean MCC and CI as a shaded area
# for each method and data type
fig, ax = plt.subplots(1, 1, figsize=(12, 6))

sns.lineplot(
    ax=ax,
    data=idBenchmarkData,
    x="PerturbationRange",
    y="MCC",
    hue="Method",
    style="Method",
    palette=method_palette,
    linewidth=3,
    markers=method_markers,
    markersize=12,
    markeredgewidth=0.5,
    markeredgecolor="black", 
    
    # Estimation and confidence interval
    estimator='mean',
    errorbar=('ci', 95),
    err_style="band",
    rasterized=True,
)

ax.set_xlabel("Perturbation Range", fontsize=14, fontweight="bold")
ax.set_ylabel("MCC Score", fontsize=14, fontweight="bold")
ax.set_title("Peptide Identification Benchmark with MCC across Perturbation Ranges", fontsize=16, fontweight="bold")
ax.set_ylim(-0.15, 1.0)
ax.grid("y", linestyle="--", linewidth=0.75, alpha=0.5, color="lightgrey")
# ax.set_xticklabels(ax.get_xticklabels(), rotation=45, ha="right", fontsize=10)
ax.legend(
    loc='upper right', title="Method", fontsize=10, title_fontsize=12, 
    frameon=False, 
)   

# # Draw MCC interpretation thresholds
# for thresh, label in mcc_thresholds.items():
#     ax.axhline(
#         thresh, color=mcc_colors[label], alpha=1,
#         linestyle="dotted", linewidth=1.5, 
#         label=label, zorder=0
#         )
#     # Use mixed transform: relative x, data y
#     ax.text(
#         0.01,
#         thresh,
#         label,
#         color=mcc_colors[label],
#         ha="left",
#         va="center",  # Center on the line
#         fontsize=10,
#         fontweight="bold",
#         transform=ax.get_yaxis_transform(),  # x in axes coords, y in data coords
#         bbox=dict(boxstyle='round,pad=0.3', facecolor='white', edgecolor=mcc_colors[label], alpha=0.8)
#     )

# Add smaller scatter points for MCC values at p-value threshold
pthr_data = idBenchmarkData[idBenchmarkData['threshold'] == pthr]
for i, method in enumerate(method_palette.keys()):
    method_data = pthr_data[pthr_data['Method'] == method]
    ax.scatter(
        method_data['PerturbationRange'],
        method_data['MCC'],
        color=method_palette[method],
        s=50,
        edgecolor='black',
        linewidth=1.25,
        marker=method_markers[method],
        zorder=10,
        alpha=0.9
    )

# Add a text to clarify the various annotations of the plot
# Include the p-value threshold in the text
fig.text(
    1,
    1.05,
    f"Black markers represent the MCC values at p-value threshold of {pthr}\nMCC interpretation thresholds are indicated by dotted lines\nShaded areas represent 95% confidence intervals around the mean MCC",
    ha="right",
    va="center",
    fontsize=12,
    fontstyle="italic",
    color="gray",
)
# Finalize the plot
plt.tight_layout()
plots.finalize_plot(
    fig, 
    show=True,
    save=save_to_folder,
    filename=f"{simID}_PepIDBenchmark_MCC_Curve_Perturbation", 
    filepath=figure_path,
    formats=figure_formats, 
    transparent=transparent_bg,
    dpi=figure_dpi,
)
```

##### AUROC Heatmap: Compact Magnitude Comparison¶

Heatmap provides a compact view of AUROC across all methods and magnitude ranges. Enables quick identification of the magnitude threshold where discrimination becomes reliable.

```
# Heatmap for AUROC values across perturbation ranges and methods
fig, ax = plt.subplots(1, 1, figsize=(12, 4))

# Calculate AUC for ROC curves
auc_results = []
for (method, pert_range), group in idBenchmarkData.groupby(
        ['Method', 'PerturbationRange']
    ):
    # ROC AUC
    group = utils.complete_curve_data(group, 'ROC', 'FPR', 'TPR')
    roc_data = group.sort_values('FPR')
    roc_auc = np.trapezoid(roc_data['TPR'], roc_data['FPR'])
    
    auc_results.append({
        'Method': method,
        'PerturbationRange': pert_range,
        'AUC': roc_auc,
    })
auc_df = pd.DataFrame(auc_results)
# print("\nAUC Data Preview:")
# display(auc_df.head())

# Pivot to create the matrix: Methods as rows, PerturbationRange as columns
pivot_data = auc_df.pivot(index='Method', columns='PerturbationRange', values='AUC')

# Create heatmap
sns.heatmap(
    pivot_data,
    annot=True,
    fmt='.2f',
    cmap='Blues',
    cbar_kws={'label': 'AUROC Score', 'shrink': 0.8},
    square=False,
    linewidths=0.5,
    linecolor='white',
    annot_kws={'size': 10, 'weight': 'bold'},
    ax=ax
)

ax.set_title("Peptide Identification Benchmark with AUROC across Perturbation Ranges", fontsize=16, fontweight="bold")
ax.set_xlabel("Perturbation Range", fontsize=14, fontweight="bold")
ax.set_ylabel("Method", fontsize=14, fontweight="bold")
# Rotate x-axis labels for better readabilitys
ax.set_xticklabels(ax.get_xticklabels(), rotation=0, ha="center", fontsize=10)
ax.set_yticklabels(ax.get_yticklabels(), rotation=0, ha="right", fontsize=10)

plots.finalize_plot(
    fig, 
    show=True,
    save=save_to_folder,
    filename=f"{simID}_PepIDBenchmark_AUC_Heatmap_Compact", 
    filepath=figure_path,
    formats=figure_formats, 
    transparent=transparent_bg,
    dpi=figure_dpi,
)
```


---

#### Simulation 4: Complex Experimental Designs¶

**Objective:** Evaluate method robustness across experimental designs with multiple interacting factors.

**Input Data:** `./data/Sim4/2_{N}Cond_{Overlap|NonOverlap}_{same|random}Dir_{method}_ResultData.feather`

**Design Factors:**
| Factor | Levels | Description |
|--------|--------|-------------|
| **Conditions** | 2, 3, 4, 5, 6 | Number of experimental conditions |
| **Overlap** | True, False | Same vs. different peptides perturbed across conditions |
| **Direction** | same, random | Uniform vs. random perturbation direction |

**Total Combinations:** 5 conditions × 2 overlap × 2 direction = 20 datasets per method

**Key Questions:**

- Does adding more conditions improve identification?
- Are overlapping perturbation patterns easier to detect?

##### Data Processing and Metric Calculation¶

```
stTime = utils.getTime()

simID = "Sim4"  
# Set up a path for the simulation
output_path, figure_path = setup_simulation_paths( simID )

methods = list(method_palette.keys())

# Systematic parameter combinations
overlap_types = [True, False]    # Overlap vs Non-overlap
direction_types = ["random", "same"]  # Direction of perturbations

# Number of conditions to generate (with shifts)
conditions = {
    2: [0, .5],
    3: [0, .5, 1],
    4: [0, .5, 1, 1.5],
    5: [0, .5, 1, 1.5, 2],
    6: [0, .5, 1, 1.5, 2, 2.5]
}

print("PEPTIDE IDENTIFICATION BENCHMARK ANALYSIS")
print("-" * 52)
print(f"Simulation ID: {simID}")
print(f"Methods: {', '.join(methods)} ({len(methods)} total)")
print(f"Overlap Types: {', '.join(['Overlap' if ot else 'Non-overlap' for ot in overlap_types])}")
print(f"Direction Types: {', '.join(direction_types)}")
print(f"Conditions (with shifts): {', '.join([str(k) for k in conditions.keys()])}")
print(f"Output Path: {output_path}")

# Progress tracking
total_combinations = len(methods) * len(overlap_types) * len(direction_types)  * len(conditions)
current_combination = 0
files_found = 0
files_missing = 0 
print(f"Total combinations to process: {total_combinations}")
print("-" * 52)

display("Calculating ID Benchmarks:")

combined_results = []
for overlap in overlap_types:
    for direction in direction_types:
        for n_conditions, shifts in conditions.items():
            for cur_method in methods:
                current_combination += 1
                
                overlap_str = "Overlap" if overlap else "NonOverlap"
                shifts_str = "random" if direction == "random" else "same"

                result_file = f"{output_path}2_{n_conditions}Cond_{overlap_str}_{shifts_str}Dir_{cur_method}_ResultData.feather"

                if os.path.exists(result_file):
                    res_df = pd.read_feather(result_file)
                    files_found += 1

                    if verbose:
                        progress = f"[{current_combination:2d}/{total_combinations}]"
                        print(f"{progress} ✓ {cur_method:12} | {n_conditions} Conditions | {overlap_str} | {shifts_str}")
                else:
                    files_missing += 1
                    if verbose:
                        progress = f"[{current_combination:2d}/{total_combinations}]"
                        print(f"{progress} ✗ {cur_method:12} | {n_conditions} Conditions | {overlap_str} | {shifts_str} (MISSING)")
                    continue
                
                # Handle COPF-specific column renaming
                if cur_method == "COPF":
                    res_df = res_df.rename(columns={
                        "proteoform_score_pval": "adj_pval",
                        'protein_id': "Protein"
                    })

                # Create metric data
                metric_data = utils.create_metric_data(
                    res_df, 
                    pvalue_thresholds=thresholds,
                    label_col='pertPeptide',
                    pvalue_col='adj_pval'
                )
                metric_data['Method'] = cur_method
                metric_data['Overlap'] = overlap
                metric_data['Direction'] = direction
                metric_data['N_Conditions'] = n_conditions
                metric_data['Shifts'] = ','.join([str(s) for s in shifts])
                combined_results.append(metric_data)    
# Combine and process data
idBenchmarkData = pd.concat(combined_results, ignore_index=True)
# Save the processed data
output_file = f"{output_path}4_{simID}_PeptideIdentification_PerformanceData.feather"
idBenchmarkData.to_feather(output_file)
idBenchmarkData.to_csv(output_file.replace('.feather', '.csv'), index=False)

print(f"\nDATA PREVIEW:")
display(idBenchmarkData.tail(5))

print(f"\nRESULTS SUMMARY:")
print("=" * 52)
execTime = utils.prettyTimer(utils.getTime() - stTime)
print(f"Total execution time:         {execTime}")
print(f"Files processed successfully: {files_found}")
print(f"Files missing/skipped:        {files_missing}")
print(f"Processing success rate:      {files_found/(files_found+files_missing)*100:.1f}%")
print("-" * 52)
print(f"Final dataset shape:          {idBenchmarkData.shape}")
overlap_types_str = [ "Overlap" if ot else "Non-overlap" for ot in overlap_types ]
print(f"Overlap Types: {len(overlap_types)} -> {', '.join(overlap_types_str)}")
print(f"Direction Types: {len(direction_types)} -> {', '.join(direction_types)}")
print(f"Conditions (with shifts): {len(conditions)} -> {', '.join([str(k) for k in conditions.keys()])}")
methods_list = sorted(idBenchmarkData['Method'].unique())
print(f"Methods analyzed: {idBenchmarkData['Method'].nunique()} -> {', '.join(methods_list)}")
print(f"Threshold levels included: {idBenchmarkData['threshold'].nunique()} (min={idBenchmarkData['threshold'].min()}, max={idBenchmarkData['threshold'].max()})")
print(f"Data saved to: {output_file}")
```

```
PEPTIDE IDENTIFICATION BENCHMARK ANALYSIS
----------------------------------------------------
Simulation ID: Sim4
Methods: COPF, PeCorA, ProteoForge (3 total)
Overlap Types: Overlap, Non-overlap
Direction Types: random, same
Conditions (with shifts): 2, 3, 4, 5, 6
Output Path: ./data/Sim4/
Total combinations to process: 60
----------------------------------------------------
```

```
'Calculating ID Benchmarks:'
```

```
[ 1/60] ✓ COPF         | 2 Conditions | Overlap | random
[ 2/60] ✓ PeCorA       | 2 Conditions | Overlap | random
[ 3/60] ✓ ProteoForge  | 2 Conditions | Overlap | random
[ 4/60] ✓ COPF         | 3 Conditions | Overlap | random
[ 5/60] ✓ PeCorA       | 3 Conditions | Overlap | random
[ 6/60] ✓ ProteoForge  | 3 Conditions | Overlap | random
[ 7/60] ✓ COPF         | 4 Conditions | Overlap | random
[ 8/60] ✓ PeCorA       | 4 Conditions | Overlap | random
[ 9/60] ✓ ProteoForge  | 4 Conditions | Overlap | random
[10/60] ✓ COPF         | 5 Conditions | Overlap | random
[11/60] ✓ PeCorA       | 5 Conditions | Overlap | random
[12/60] ✓ ProteoForge  | 5 Conditions | Overlap | random
[13/60] ✓ COPF         | 6 Conditions | Overlap | random
[14/60] ✓ PeCorA       | 6 Conditions | Overlap | random
[15/60] ✓ ProteoForge  | 6 Conditions | Overlap | random
[16/60] ✓ COPF         | 2 Conditions | Overlap | same
[17/60] ✓ PeCorA       | 2 Conditions | Overlap | same
[18/60] ✓ ProteoForge  | 2 Conditions | Overlap | same
[19/60] ✓ COPF         | 3 Conditions | Overlap | same
[20/60] ✓ PeCorA       | 3 Conditions | Overlap | same
[21/60] ✓ ProteoForge  | 3 Conditions | Overlap | same
[22/60] ✓ COPF         | 4 Conditions | Overlap | same
[23/60] ✓ PeCorA       | 4 Conditions | Overlap | same
[24/60] ✓ ProteoForge  | 4 Conditions | Overlap | same
[25/60] ✓ COPF         | 5 Conditions | Overlap | same
[26/60] ✓ PeCorA       | 5 Conditions | Overlap | same
[27/60] ✓ ProteoForge  | 5 Conditions | Overlap | same
[28/60] ✓ COPF         | 6 Conditions | Overlap | same
[29/60] ✓ PeCorA       | 6 Conditions | Overlap | same
[30/60] ✓ ProteoForge  | 6 Conditions | Overlap | same
[31/60] ✓ COPF         | 2 Conditions | NonOverlap | random
[32/60] ✓ PeCorA       | 2 Conditions | NonOverlap | random
[33/60] ✓ ProteoForge  | 2 Conditions | NonOverlap | random
[34/60] ✓ COPF         | 3 Conditions | NonOverlap | random
[35/60] ✓ PeCorA       | 3 Conditions | NonOverlap | random
[36/60] ✓ ProteoForge  | 3 Conditions | NonOverlap | random
[37/60] ✓ COPF         | 4 Conditions | NonOverlap | random
[38/60] ✓ PeCorA       | 4 Conditions | NonOverlap | random
[39/60] ✓ ProteoForge  | 4 Conditions | NonOverlap | random
[40/60] ✓ COPF         | 5 Conditions | NonOverlap | random
[41/60] ✓ PeCorA       | 5 Conditions | NonOverlap | random
[42/60] ✓ ProteoForge  | 5 Conditions | NonOverlap | random
[43/60] ✓ COPF         | 6 Conditions | NonOverlap | random
[44/60] ✓ PeCorA       | 6 Conditions | NonOverlap | random
[45/60] ✓ ProteoForge  | 6 Conditions | NonOverlap | random
[46/60] ✓ COPF         | 2 Conditions | NonOverlap | same
[47/60] ✓ PeCorA       | 2 Conditions | NonOverlap | same
[48/60] ✓ ProteoForge  | 2 Conditions | NonOverlap | same
[49/60] ✓ COPF         | 3 Conditions | NonOverlap | same
[50/60] ✓ PeCorA       | 3 Conditions | NonOverlap | same
[51/60] ✓ ProteoForge  | 3 Conditions | NonOverlap | same
[52/60] ✓ COPF         | 4 Conditions | NonOverlap | same
[53/60] ✓ PeCorA       | 4 Conditions | NonOverlap | same
[54/60] ✓ ProteoForge  | 4 Conditions | NonOverlap | same
[55/60] ✓ COPF         | 5 Conditions | NonOverlap | same
[56/60] ✓ PeCorA       | 5 Conditions | NonOverlap | same
[57/60] ✓ ProteoForge  | 5 Conditions | NonOverlap | same
[58/60] ✓ COPF         | 6 Conditions | NonOverlap | same
[59/60] ✓ PeCorA       | 6 Conditions | NonOverlap | same
[60/60] ✓ ProteoForge  | 6 Conditions | NonOverlap | same

DATA PREVIEW:
```

|  | TP | FP | TN | FN | TPR | FPR | FDR | MCC | Precision | Recall | F1 | threshold | Method | Overlap | Direction | N\_Conditions | Shifts |
| --- | --- | --- | --- | --- | --- | --- | --- | --- | --- | --- | --- | --- | --- | --- | --- | --- | --- |
| 1495 | 871 | 590 | 4178 | 44 | 0.9519 | 0.1237 | 0.4038 | 0.6965 | 0.5962 | 0.9519 | 0.7332 | 0.6000 | ProteoForge | False | same | 6 | 0,0.5,1,1.5,2,2.5 |
| 1496 | 873 | 631 | 4137 | 42 | 0.9541 | 0.1323 | 0.4195 | 0.6846 | 0.5805 | 0.9541 | 0.7218 | 0.7000 | ProteoForge | False | same | 6 | 0,0.5,1,1.5,2,2.5 |
| 1497 | 875 | 671 | 4097 | 40 | 0.9563 | 0.1407 | 0.4340 | 0.6736 | 0.5660 | 0.9563 | 0.7111 | 0.8000 | ProteoForge | False | same | 6 | 0,0.5,1,1.5,2,2.5 |
| 1498 | 877 | 708 | 4060 | 38 | 0.9585 | 0.1485 | 0.4467 | 0.6638 | 0.5533 | 0.9585 | 0.7016 | 0.9000 | ProteoForge | False | same | 6 | 0,0.5,1,1.5,2,2.5 |
| 1499 | 915 | 4768 | 0 | 0 | 1.0000 | 1.0000 | 0.8390 | 0.0000 | 0.1610 | 1.0000 | 0.2774 | 1.0000 | ProteoForge | False | same | 6 | 0,0.5,1,1.5,2,2.5 |

```
RESULTS SUMMARY:
====================================================
Total execution time:         00h:00m:07s
Files processed successfully: 60
Files missing/skipped:        0
Processing success rate:      100.0%
----------------------------------------------------
Final dataset shape:          (1500, 17)
Overlap Types: 2 -> Overlap, Non-overlap
Direction Types: 2 -> random, same
Conditions (with shifts): 5 -> 2, 3, 4, 5, 6
Methods analyzed: 3 -> COPF, PeCorA, ProteoForge
Threshold levels included: 25 (min=0.0, max=1.0)
Data saved to: ./data/Sim4/4_Sim4_PeptideIdentification_PerformanceData.feather
```

##### Comprehensive Barplot: Factor Interactions¶

Four-panel barplot showing MCC across number of conditions, stratified by overlap type (rows) and perturbation direction (columns). Error bars show 95% CI. Scatter points indicate MCC at the p-value threshold. Values below bars show mean MCC scores.

```
# Comprehensive bar plot with all perturbation ranges and statistical details
fig, axes = plt.subplots(
    2, 2, figsize=(16, 10), 
    sharey=True, sharex=True,
    gridspec_kw={
        "hspace": 0.01,  # Reduced vertical spacing
        "wspace": 0.01  # Minimal horizontal spacing
    }
)

plot_combinations = [
    (True, "same"),
    (True, "random"),
    (False, "same"),
    (False, "random"),
]

# Store handles and labels for shared legend
handles, labels = None, None

for i, (overlap, direction) in enumerate(plot_combinations):
    cur_data = idBenchmarkData[
        (idBenchmarkData["Overlap"] == overlap) & 
        (idBenchmarkData["Direction"] == direction)
    ]
    ax = axes[i//2, i%2]

    # Get unique conditions and methods for proper positioning
    unique_conditions = sorted(cur_data['N_Conditions'].unique())
    methods = list(method_palette.keys())
    n_methods = len(methods)
    # Calculate bar width and positioning (seaborn default spacing)
    bar_width = 0.8 / n_methods  # Total width of 0.8 divided by number of methods

    bar_plot = sns.barplot(
        ax=ax,
        data=cur_data,
        x="N_Conditions",
        y="MCC",
        hue="Method",
        palette=method_palette,
        edgecolor='black',
        errwidth=1.5,
        capsize=0.1,
        dodge=True,
        estimator='mean',
        errorbar=('ci', 95),
        legend=False  # Disable individual legends
    )
    
    # Store legend info from first plot
    if i == 0:
        handles, labels = ax.get_legend_handles_labels()
    
    # Set direction as title for each column
    if i // 2 == 0:  # Top row
        direction_title = f"{direction.capitalize()} Direction"
        ax.set_title(direction_title, fontsize=14, fontweight="bold")
    
    # Add overlap/non-overlap labels on the right side
    if i % 2 == 1:  # Right column
        overlap_label = "Overlap" if overlap else "Non-overlap"
        ax.text(
            1.05, 0.5, 
            overlap_label,
            transform=ax.transAxes,
            rotation=270,
            va="center",
            ha="left",
            fontsize=14,
            fontweight="bold"
        )
    
    # Configure axis labels
    if i // 2 == 1:  # Bottom row - show x-axis labels
        ax.set_xlabel("Number of Conditions", fontsize=12, fontweight="bold")
    else:  # Top row - hide x-axis labels
        ax.set_xlabel("")
        # ax.set_xticklabels([])
    
    if i % 2 == 0:  # Left column - show y-axis labels
        ax.set_ylabel("Mean MCC Score", fontsize=12, fontweight="bold")
    else:  # Right column - hide y-axis labels
        ax.set_ylabel("")
    
    # Styling
    ax.set_ylim(-0.15, 1.0)
    ax.grid("y", linestyle="--", linewidth=0.75, alpha=0.5, color="lightgrey")

    # Write the average MCC score for each method at the bottom of each bar group
    for j, condition in enumerate(unique_conditions):
        for k, method in enumerate(methods):
            avg_score = cur_data[
                (cur_data["N_Conditions"] == condition) &
                (cur_data["Method"] == method)
            ]["MCC"].mean()
            # Calculate the x position for this method's bar
            x_pos = j + (k - (n_methods - 1) / 2) * bar_width
            ax.text(
                x_pos,
                -0.01,  # Below the x-axis
                f"{avg_score:.2f}",
                color=method_palette[method],
                ha="center",
                va="top",
                fontsize=10,
                fontweight="bold",
                rotation=90,
                clip_on=False,
            )

    # # Draw MCC interpretation thresholds
    # for thresh, label in mcc_thresholds.items():
    #     ax.axhline(
    #         thresh, color=mcc_colors[label], alpha=1,
    #         linestyle="dotted", linewidth=1.5, 
    #         label=label, zorder=0
    #         )
        
    #     # Only draw text for first column to avoid clutter
    #     if i % 2 == 1:
    #         continue
    #     # Use mixed transform: relative x, data y
    #     ax.text(
    #         0.01,
    #         thresh,
    #         label,
    #         color=mcc_colors[label],
    #         ha="left",
    #         va="center",  # Center on the line
    #         fontsize=10,
    #         fontweight="bold",
    #         transform=ax.get_yaxis_transform(),  # x in axes coords, y in data coords
    #         bbox=dict(boxstyle='round,pad=0.3', facecolor='white', edgecolor=mcc_colors[label], alpha=0.8), 
    #         zorder=0
    #     )

    # Add scatter points for MCC values at p-value threshold
    pthr_data = cur_data[cur_data['threshold'] == pthr]
    
    for j, method in enumerate(methods):
        method_data = pthr_data[pthr_data['Method'] == method]
        
        x_positions = []
        y_positions = []
        
        for condition in unique_conditions:
            condition_data = method_data[method_data['N_Conditions'] == condition]
            if not condition_data.empty:
                # Calculate x position: condition index + offset for this method
                condition_idx = unique_conditions.index(condition)
                x_offset = (j - (n_methods - 1) / 2) * bar_width
                x_pos = condition_idx + x_offset
                
                x_positions.append(x_pos)
                y_positions.append(condition_data['MCC'].iloc[0])
        
        # Add scatter points for this method
        if x_positions:  # Only scatter if we have data
            ax.scatter(
                x_positions,
                y_positions,
                color=method_palette[method],
                s=75,
                edgecolor='black',
                linewidth=1.25,
                marker=method_markers[method],
                zorder=10,
                alpha=0.9
            )
        # # Add boxed text for each point
        # for (x, y) in zip(x_positions, y_positions):
        #     ax.text(
        #         x,
        #         y - 0.05,  # Slightly below the point
        #         f'{y:.2f}',
        #         ha="center",
        #         va="top",
        #         fontsize=9,
        #         fontweight="bold",
        #         color="black",
        #         bbox=dict(boxstyle='round,pad=0.3', facecolor='white', edgecolor='black', alpha=0.8),
        #         zorder=10
        #     )


# Create a legend for methods using marker and color
from matplotlib.lines import Line2D
method_legend_elements = [
    Line2D(
        [0], [0],
        marker=method_markers[m],
        color='w',
        markerfacecolor=method_palette[m],
        markeredgecolor='black',
        markersize=12,
        linewidth=0,
        label=m
    )
    for m in methods
]

fig.legend(
    handles=method_legend_elements,
    loc='lower center',
    ncol=len(methods),
    fontsize=12,
    title="Method",
    title_fontsize=14,
    frameon=False,
    bbox_to_anchor=(0.5, 0.05)
)

# Adjust layout to accommodate legend
plt.tight_layout()
plt.subplots_adjust(bottom=0.15)

plots.finalize_plot(
    fig, 
    show=True,
    save=save_to_folder,
    filename=f"{simID}_PepIDBenchmark_MCC_Conditions_Barplot", 
    filepath=figure_path,
    formats=figure_formats, 
    transparent=transparent_bg,
    dpi=figure_dpi,
)
```


---

#### Summary¶

##### Output Files Generated¶

| Simulation | File | Description |
| --- | --- | --- |
| Sim1 | `figures/roc_curves.png` | ROC curves across imputation strategies |
| Sim1 | `figures/pr_curves.png` | Precision-Recall curves across imputation |
| Sim1 | `figures/mcc_curves.png` | MCC vs. p-value threshold curves |
| Sim1 | `figures/mcc_barplots.png` | MCC comparison barplots |
| Sim2 | `figures/auroc_heatmap.png` | AUROC heatmap across missingness levels |
| Sim2 | `figures/mean_mcc_heatmap.png` | Mean MCC across all thresholds |
| Sim2 | `figures/mcc_at_threshold.png` | MCC at optimal p-value threshold |
| Sim2 | `figures/max_mcc_heatmap.png` | Maximum achievable MCC |
| Sim3 | `figures/mcc_sensitivity.png` | MCC sensitivity to magnitude |
| Sim3 | `figures/auroc_magnitude.png` | AUROC heatmap across magnitudes |
| Sim4 | `figures/sim4_barplot.png` | MCC across experimental conditions |

##### Key Findings by Simulation¶

1. **Sim1 (Imputation)**: Establishes baseline performance under controlled missingness+imputation conditions
2. **Sim2 (Missingness)**: Reveals method robustness across 0-70% missingness spectrum
3. **Sim3 (Magnitude)**: Tests sensitivity to effect size (1.2x-2.5x fold change)
4. **Sim4 (Conditions)**: Evaluates scalability with experimental complexity (3-9 conditions)

```
print("Notebook Execution Time:", utils.prettyTimer(utils.getTime() - startTime))
```
