## Supplementary material for "ProteoForge: An Imputation-Aware Framework for Differential Proteoform Discovery in Bottom-Up Proteomics": Analysis notebooks as html render: Notebook S6.html

### Notebook S6

###### Enes Kemal Ergin

#### 2025-12-08 23:00:33

### Supplementary Notebook 6: Peptide Grouping Benchmarking on Simulated Datasets¶

- **License:** Creative Commons Attribution-NonCommercial 4.0 International License
- **Version:** 0.2
- **Edit Log:**
  - 2025-11-28: Initial version of the notebook
  - 2025-12-08: Revise the whole notebook, ensuring clarity and correctness

---

**Requirements:**

- Completed method runs: `02-runCOPF.R`, `04-runProteoForge.py`
- Result files in `./data/Sim{1-4}/` directories

**Data Information:**  
This notebook processes method results from four simulation scenarios to evaluate **peptide grouping** performance—the ability to correctly identify which peptides are perturbed.

| Simulation | Focus | Input Files Pattern |
| --- | --- | --- |
| **Sim1** | Complete vs. Imputed | `2_{pattern}_{type}_{method}_ResultData.feather` |
| **Sim2** | Missingness Levels | `2_Pro{rate}_Pep{rate}_imputed_{method}_ResultData.feather` |
| **Sim3** | Perturbation Magnitude | `2_{low}_{high}_{method}_ResultData.feather` |
| **Sim4** | Experimental Complexity | `2_{N}Cond_{overlap}_{dir}Dir_{method}_ResultData.feather` |

**Purpose:**  
Benchmark COPF, and ProteoForge on peptide grouping using ROC curves, PR curves, and Matthews Correlation Coefficient (MCC) across multiple p-value thresholds.

> **Note:** PeCorA is excluded from peptide grouping benchmarks as it does not inheritly provide peptide-level grouping related information.

---

#### Setup¶

This section imports libraries, configures display settings, and defines paths for outputs.

> **Note:** The HTML rendering of this notebook hides code cells by default. Click the "Code" buttons to expand them.

##### Libraries¶

```
import os
import sys
import warnings
warnings.filterwarnings("ignore")

import numpy as np # Numerical computing
import pandas as pd # Data manipulatio

import seaborn as sns # R-like high-level plots
import matplotlib.pyplot as plt # Python's base plotting 

sys.path.append('../')

# Utility imports for this analysis
from src import utils, plots

# ProteoForge Package imports
from ProteoForge import proteoform_classifier as pfc

# Initialize the timer
startTime = utils.getTime()
```

##### Display Settings¶

The cell below configures pandas, matplotlib, and seaborn display options for improved readability of tables and figures, including color palettes and figure export settings.

```
## Figure Settings

# Define default colors and styles for plots
def_colors = [
    "#139593", "#fca311", "#e54f2a",
    "#c3c3c3", "#555555",
    "#690000", "#5f4a00", "#004549"
]

# Set seaborn style
sns.set_theme(
    style="white",
    context="paper",
    palette=def_colors,
    font_scale=1,
    rc={
        "figure.figsize": (6, 4),
        "font.family": "sans-serif",
        "font.sans-serif": ["Arial", "Ubuntu Mono"],
    }
)

# Figure Saving Settings
figure_formats = ["pdf"]
save_to_folder = True
transparent_bg = True
figure_dpi = 300

## Configure dataframe displaying
pd.set_option('display.float_format', lambda x: '%.4f' % x)
pd.set_option('display.max_rows', 50)
pd.set_option('display.max_columns', 50)
pd.set_option('display.width', 500)  # Set a wider display width

## Printing Settings
verbose = True

## Show the color-palettes
plots.color_palette( def_colors, save=False )
```

##### Data and Result Paths¶

Defines input/output directory structure:

- **Input**: `./data/Sim{N}/` — Method result files from each simulation
- **Output**: Processed performance data and figures saved to same directories

```
# Establish paths and 
notebook_name = "06-GroupBenchmark"
input_path = f"./data/"
mainFig_path = f"./figures/"

def setup_simulation_paths( simID ):
    """
    Create output and figure directories for a simulation, if they do not exist. 
    (Uses global variable for save_to_folder and figure_formats)

    Args:
        simID (str): Simulation ID.
        
        (Global Variables)
            input_path (str): Base path for input data.
            mainFig_path (str): Base path for figures.
            save_to_folder (bool): Whether to save figures to folders.
            figure_formats (list): List of formats to save figures in.
    
    Returns:
        tuple: (output_path, figure_path)
    """
    output_path = f"{input_path}{simID}/"
    if not os.path.exists(output_path):
        os.makedirs(output_path)

    figure_path = f"{mainFig_path}{simID}/"
    if save_to_folder:
        for fmt in figure_formats:
            cur_folder = os.path.join(figure_path, fmt)
            if not os.path.exists(cur_folder):
                os.makedirs(cur_folder)
    return output_path, figure_path
```

##### Global Variables¶

Defines analysis constants and visualization styling:

- **`seed`**, **`pthr`**: Reproducibility seed and p-value threshold (10⁻³)
- **`thresholds`**: Range of p-value thresholds for curve generation
- **`method_palette`**, **`method_markers`**: Consistent styling for COPF, PeCorA, ProteoForge
- **`mcc_thresholds`**, **`mcc_colors`**: MCC interpretation scale (Random → Almost Perfect)

```
# Global variables
seed = 42 # Seed for reproducibility  
pthr = 10**-3  # p-value threshold for significance
thresholds = list(utils.generate_thresholds(10.0, -15, 1, 0, 1, 0.1)) # Thresholds for the analysis

method_palette = {
    "COPF": "#139593",
    "ProteoForge": "#e54f2a",
}
method_styles = {
    "COPF": "--",
    "ProteoForge": ":",
}
method_markers = {
    "COPF": "o",
    "ProteoForge": "^",
}

# Matthews Correlation Coefficient (MCC) thresholds and colors
mcc_thresholds = {
    0.0 : 'Random',
    0.3 : 'Fair (0.3)',
    0.5 : 'Moderate (0.5)',
    0.7 : 'Strong (0.7)',
    0.9 : 'Almost Perfect (0.9)',
}
mcc_colors = {
    'Random': '#c3c3c3',             
    'Fair (0.3)': '#e0aaff',         
    'Moderate (0.5)': '#9d4edd',     
    'Strong (0.7)': '#5a189a',       
    'Almost Perfect (0.9)': '#240046' 
}

plots.color_palette(mcc_colors, save=False, name="MCC Interpretation Colors")
```

#### Simulation 1 - Complete vs. Imputed Data with Varying Peptide Perturbation Patterns¶

##### Objective¶

Evaluate peptide **grouping** performance under controlled 30% MNAR conditions with four perturbation patterns. Unlike identification (which peptides are perturbed), grouping assesses whether perturbed peptides are correctly clustered together.

##### Input Data¶

- **Pattern**: `2_{pattern}_{type}_{method}_ResultData.feather`
- **Methods**: COPF, ProteoForge (PeCorA excluded—no grouping output)

##### Experimental Design¶

| Factor | Levels | Description |
| --- | --- | --- |
| Data Type | complete, imputed | Before/after imputation |
| Perturbation | twoPep, halfPep, halfPlusPep, randomPep | Number of perturbed peptides per protein |

##### Key Questions¶

1. How does imputation affect grouping accuracy?
2. Which perturbation patterns are easiest/hardest to group correctly?
3. Does ProteoForge's clustering outperform COPF's approach?

##### Metrics Generated¶

- ROC curves (TPR vs FPR) with AUROC
- Precision-Recall curves with AUPRC
- MCC across p-value thresholds
- MCC barplots at optimal threshold

```
stTime = utils.getTime()

simID = "Sim1"  
# Set up a path for the simulation
output_path, figure_path = setup_simulation_paths( simID )

methods = list(method_palette.keys())
dataTypes = ['complete', 'imputed']
experiments = ['twoPep', 'randomPep', 'halfPep', 'halfPlusPep']
experiment_mapper = {
    'twoPep': 'Two Peptides', 
    'randomPep': '2>50% Peptides',
    'halfPep': '50% Peptides',
    'halfPlusPep': '>50% Peptides',
}

# =============================================================================
# ANALYSIS SETUP
# =============================================================================
print("PROTEOFORM GROUPING BENCHMARK ANALYSIS")
print("-" * 52)
print(f"Simulation ID: {simID}")
print(f"Methods: {', '.join(methods)} ({len(methods)} total)")
print(f"Data Types: {', '.join(dataTypes)}")
print(f"Experiments: {len(experiments)} scenarios")
print(f"Output Path: {output_path}")

# Progress tracking
total_combinations = len(methods) * len(dataTypes) * len(experiments)
files_found = 0
files_missing = 0
current_combination = 0

print(f"Total combinations to process: {total_combinations}")
print("-" * 52)

# =============================================================================
# DATA PROCESSING
# =============================================================================
display("Calculating Grouping Benchmarks:")

combined_results = []
for cur_method in methods:
    for cur_type in dataTypes:
        for cur_exp in experiments:
            current_combination += 1
            
            # Load the result data
            result_file = f"{output_path}2_{cur_exp}_{cur_type}_{cur_method}_ResultData.feather"
            
            if os.path.exists(result_file):
                res_df = pd.read_feather(result_file)
                files_found += 1

                # Show progress for successful loads only
                if verbose:
                    progress = f"[{current_combination:2d}/{total_combinations}]"
                    print(f"{progress} ✓ {cur_method:12} | {cur_type:9} | {experiment_mapper[cur_exp]}")
            else:
                files_missing += 1
                if verbose:
                    progress = f"[{current_combination:2d}/{total_combinations}]"
                    print(f"{progress} ✗ {cur_method:12} | {cur_type:9} | {experiment_mapper[cur_exp]} (MISSING)")
                continue

            # Handle COPF-specific column renaming
            if cur_method == "COPF":
                res_df = res_df.rename(columns={
                    "proteoform_score_pval": "adj_pval",
                    'protein_id': "Protein"
                })
                metric_data = utils.grouping_performance_copf(
                    data=res_df,
                    thresholds=thresholds,
                    pvalue_col='adj_pval',
                    protein_col='Protein',
                    cluster_col='cluster',
                    perturbation_col='pertPFG',
                )

            elif cur_method == "ProteoForge":
                metric_data = utils.grouping_performance_proteoforge(
                    data=res_df,
                    thresholds=thresholds,
                    pvalue_col='adj_pval',
                    protein_col='Protein',
                    cluster_col='ClusterID',
                    perturbation_col='pertPFG',
                )
            
            metric_data['Method'] = cur_method
            metric_data['DataType'] = cur_type
            metric_data['Experiment'] = cur_exp
            combined_results.append(metric_data)

# Combine and process data
groupBenchmarkData = pd.concat(combined_results, ignore_index=True)
groupBenchmarkData['Experiment'] = groupBenchmarkData['Experiment'].map(experiment_mapper)

# Categorical ordering ensure twoPep, randomPep, halfPep, halfPlusPep
groupBenchmarkData['Experiment'] = pd.Categorical(
    groupBenchmarkData['Experiment'], 
    categories=[experiment_mapper[exp] for exp in experiments],
    ordered=True
)

print(f"\nDATA PREVIEW:")
display(groupBenchmarkData.head(3))

# Save the processed data
output_file = f"{output_path}4_{simID}_Grouping_PerformanceData.feather"
groupBenchmarkData.to_feather(output_file)
groupBenchmarkData.to_csv(output_file.replace('.feather', '.csv'), index=False)

print(f"\nRESULTS SUMMARY:")
print("=" * 52)
execTime = utils.prettyTimer(utils.getTime() - stTime)
print(f"Total execution time:         {execTime}")
print(f"Files processed successfully: {files_found}")
print(f"Files missing/skipped:        {files_missing}")
print(f"Processing success rate:      {files_found/(files_found+files_missing)*100:.1f}%")
print("-" * 52)
print(f"Final dataset shape:          {groupBenchmarkData.shape}")
print(f"Unique experiments:           {groupBenchmarkData['Experiment'].nunique()}")
print(f"Methods analyzed:             {groupBenchmarkData['Method'].nunique()}")
print(f"Data types included:          {groupBenchmarkData['DataType'].nunique()}")
print(f"Data saved to: {output_file}")
print("=" * 52)
```

```
PROTEOFORM GROUPING BENCHMARK ANALYSIS
----------------------------------------------------
Simulation ID: Sim1
Methods: COPF, ProteoForge (2 total)
Data Types: complete, imputed
Experiments: 4 scenarios
Output Path: ./data/Sim1/
Total combinations to process: 16
----------------------------------------------------
```

```
'Calculating Grouping Benchmarks:'
```

```
[ 1/16] ✓ COPF         | complete  | Two Peptides
[ 2/16] ✓ COPF         | complete  | 2>50% Peptides
[ 3/16] ✓ COPF         | complete  | 50% Peptides
[ 4/16] ✓ COPF         | complete  | >50% Peptides
[ 5/16] ✓ COPF         | imputed   | Two Peptides
[ 6/16] ✓ COPF         | imputed   | 2>50% Peptides
[ 7/16] ✓ COPF         | imputed   | 50% Peptides
[ 8/16] ✓ COPF         | imputed   | >50% Peptides
[ 9/16] ✓ ProteoForge  | complete  | Two Peptides
[10/16] ✓ ProteoForge  | complete  | 2>50% Peptides
[11/16] ✓ ProteoForge  | complete  | 50% Peptides
[12/16] ✓ ProteoForge  | complete  | >50% Peptides
[13/16] ✓ ProteoForge  | imputed   | Two Peptides
[14/16] ✓ ProteoForge  | imputed   | 2>50% Peptides
[15/16] ✓ ProteoForge  | imputed   | 50% Peptides
[16/16] ✓ ProteoForge  | imputed   | >50% Peptides

DATA PREVIEW:
```

|  | TP | FP | TN | FN | TPR | FPR | FDR | MCC | Precision | Recall | F1 | threshold | Method | DataType | Experiment |
| --- | --- | --- | --- | --- | --- | --- | --- | --- | --- | --- | --- | --- | --- | --- | --- |
| 0 | 0 | 0 | 250 | 250 | 0.0000 | 0.0000 | 0.0000 | 0.0000 | 0.0000 | 0.0000 | 0.0000 | 0.0000 | COPF | complete | Two Peptides |
| 1 | 0 | 0 | 250 | 250 | 0.0000 | 0.0000 | 0.0000 | 0.0000 | 0.0000 | 0.0000 | 0.0000 | 0.0000 | COPF | complete | Two Peptides |
| 2 | 0 | 0 | 250 | 250 | 0.0000 | 0.0000 | 0.0000 | 0.0000 | 0.0000 | 0.0000 | 0.0000 | 0.0000 | COPF | complete | Two Peptides |

```
RESULTS SUMMARY:
====================================================
Total execution time:         00h:00m:16s
Files processed successfully: 16
Files missing/skipped:        0
Processing success rate:      100.0%
----------------------------------------------------
Final dataset shape:          (400, 15)
Unique experiments:           4
Methods analyzed:             2
Data types included:          2
Data saved to: ./data/Sim1/4_Sim1_Grouping_PerformanceData.feather
====================================================
```

##### ROC Curves: Grouping Discrimination¶

2×4 panel grid showing ROC curves for each perturbation pattern (columns) and data type (rows). AUROC values indicate overall grouping discrimination. Black circles mark performance at p = 10⁻³.

```
# ROC Curves for different perturbations and methods
fig, axes = plt.subplots(
    2, 4, figsize=(16, 6), 
    sharey=True, sharex=True,
    gridspec_kw={
        "wspace": 0.05, 
        "hspace": 0.05
    }
)

for i, pert in enumerate(groupBenchmarkData['Experiment'].unique()):
    for j, dataType in enumerate(dataTypes):
        cur_data = groupBenchmarkData[
            (groupBenchmarkData["Experiment"] == pert) & 
            (groupBenchmarkData["DataType"] == dataType)
        ]
        # Ensure the data is complete for ROC curves
        cur_data = cur_data.groupby("Method").apply(
            lambda x: utils.complete_curve_data(x, 'ROC', 'FPR', 'TPR')
        ).reset_index(drop=True)
        
        # Calculate AUC per method from TPR and FPR
        auc_data = cur_data.groupby("Method").apply(
            lambda x: np.trapezoid(
                x.sort_values("FPR")["TPR"], x.sort_values("FPR")["FPR"]
            ), 
            include_groups=False
        )
        # Plot the ROC curve
        sns.lineplot(
            data=cur_data,
            x="FPR",
            y="TPR",
            hue="Method",
            style="Method",
            palette=method_palette,
            ax=axes[j, i],
            # Show the points
            markers=method_markers,
            dashes=False,
            markersize=7.5,
            markeredgewidth=0.5,
            markeredgecolor="gray", 
            legend=False,
            rasterized=True,
            estimator=None
        )
        # Add AUC values as legend like text
        for k, method in enumerate(auc_data.index):
            auc = auc_data[method]
            # Add color to match the palette
            color = method_palette[method]
            axes[j, i].text(
                0.95,
                0.05 + k * 0.075,
                f"{method}: {auc:.2f}",
                color=color,
                transform=axes[j, i].transAxes,
                ha="right",
                va="bottom",
                fontsize=12,
                fontweight="bold",
            )
            # Add a large no-facecolor marker to the pthr value for each method
            # Isolate data for the current method
            method_data = cur_data[cur_data["Method"] == method]

            # Find the index of the row with the threshold value closest to pthr
            if not method_data.empty:
                closest_idx = (method_data['threshold'] - pthr).abs().idxmin()
                pthr_data = method_data.loc[[closest_idx]]

                axes[j, i].scatter(
                    pthr_data["FPR"],
                    pthr_data["TPR"],
                    color=color,
                    s=125,
                    edgecolor="black",
                    linewidth=1.25,
                    marker="o",
                    facecolors="none",
                )
        
        # Add the diagonal line
        axes[j, i].plot([0, 1], [0, 1], color="black", linestyle="--")
        # axes[j, i].set_title(f"{pert}", fontsize=14, fontweight="bold")
        axes[j, i].set_xlabel("FPR", fontsize=12)
        axes[j, i].set_ylabel("TPR", fontsize=12)
        axes[j, i].set_xlim(-0.05, 1.05)
        axes[j, i].set_ylim(-0.05, 1.05)
        axes[j, i].grid("both", linestyle="--", linewidth=0.75, alpha=0.5, color="lightgrey")
        # If top row set the title
        if j == 0:
            axes[j, i].set_title(f"{pert}", fontsize=14, fontweight="bold")
        # For the last column, set the text for the dataType on the right
        if i == 3:
            axes[j, i].text(
                1.05,
                0.5,
                f"{dataType.capitalize()}",
                transform=axes[j, i].transAxes,
                ha="left",
                va="center",
                fontsize=14,
                fontweight="bold",
                rotation=270,
            )

# Add a text indicating the circle is the p-value threshold under fig.title and above the first subplot
fig.text(
    0.9,
    .97,
    f"Black circles represent the TPR and FPR values at p-value threshold of {pthr}",
    ha="right",
    va="center",
    fontsize=12,
    fontstyle="italic",
    color="gray",
)

# Finalize the plot
plt.tight_layout()
plots.finalize_plot(
    fig, 
    show=True,
    save=save_to_folder,
    filename=f"{simID}_GroupingBenchmark_ROC_Curve", 
    filepath=figure_path,
    formats=figure_formats, 
    transparent=transparent_bg,
    dpi=figure_dpi,
)
```

##### PR Curves: Precision-Recall Trade-off¶

2×4 panel grid showing precision vs recall for grouping predictions. AUPRC values reflect performance under class imbalance. Black circles mark precision/recall at p = 10⁻³.

```
# Precision-Recall Curves for different perturbations and methods
fig, axes = plt.subplots(
    2, 4, figsize=(16, 6), 
    sharey=True, sharex=True,
    gridspec_kw={
        "wspace": 0.05, 
        "hspace": 0.05
    }
)
for i, pert in enumerate(groupBenchmarkData['Experiment'].unique()):
    for j, dataType in enumerate(dataTypes):
        cur_data = groupBenchmarkData[
            (groupBenchmarkData["Experiment"] == pert) & 
            (groupBenchmarkData["DataType"] == dataType)
        ]

        # Add a small value to avoid division by zero
        cur_data["Precision"] = cur_data["TP"] / (cur_data["TP"] + cur_data["FP"] + 1e-6)
        cur_data["Recall"] = cur_data["TP"] / (cur_data["TP"] + cur_data["FN"] + 1e-6)
        
        cur_data = cur_data.sort_values("Recall", ascending=False)
        cur_data = cur_data[~((cur_data["Recall"] == 0) & (cur_data["Precision"] == 0))]

        # Ensure the data is complete for PR curves
        cur_data = cur_data.groupby("Method").apply(
            lambda x: utils.complete_curve_data(x, 'PR', 'Recall', 'Precision')
        ).reset_index(drop=True)
        
        # Calculate AUC per method from Precision and Recall
        # Here we use the trapezoidal rule to calculate the area under the curve
        f1_data = cur_data.groupby("Method").apply(
            lambda x: np.trapezoid(
                x.sort_values("Recall")["Precision"], x.sort_values("Recall")["Recall"]
            ),
            include_groups=False
        )

        # Plot the PR curve
        sns.lineplot(
            data=cur_data,
            x="Recall",
            y="Precision",
            hue="Method",
            style="Method",
            palette=method_palette,
            ax=axes[j, i],
            # Show the points
            markers=method_markers,
            dashes=False,
            markersize=7.5,
            markeredgewidth=0.5,
            markeredgecolor="gray", 
            legend=False,
            rasterized=True,
            estimator=None
        )

        # Add AUC values as legend like text
        for k, method in enumerate(f1_data.index):
            f1 = f1_data[method]
            # Add color to match the palette
            color = method_palette[method]
            axes[j, i].text(
                0.05,
                0.05 + k * 0.075,
                f"{method}: {f1:.2f}",
                color=color,
                transform=axes[j, i].transAxes,
                ha="left",
                va="bottom",
                fontsize=12,
                fontweight="bold",
            )
            # Add a large no-facecolor marker to the pthr value for each method
            pthr_data = cur_data[(cur_data["Method"] == method) & (cur_data["threshold"] == pthr)]
            axes[j, i].scatter(
                pthr_data["Recall"],
                pthr_data["Precision"],
                color=color,
                s=125,
                edgecolor="black",
                linewidth=1.25,
                marker="o",
                facecolors="none",
            )

        axes[j, i].set_xlabel("Recall", fontsize=12)
        axes[j, i].set_ylabel("Precision", fontsize=12)
        axes[j, i].set_xlim(-0.05, 1.05)
        axes[j, i].set_ylim(-0.05, 1.05)
        axes[j, i].grid("both", linestyle="--", linewidth=0.75, alpha=0.5, color="lightgrey")
        # If top row set the title
        if j == 0:
            axes[j, i].set_title(f"{pert}", fontsize=14, fontweight="bold")
        # For the last column, set the text for the dataType on the right
        if i == 3:
            axes[j, i].text(
                1.05,
                0.5,
                f"{dataType.capitalize()}",
                transform=axes[j, i].transAxes,
                ha="left",
                va="center",
                fontsize=14,
                fontweight="bold",
                rotation=270,
            )

# Add a text indicating the circle is the p-value threshold under fig.title and above the first subplot
fig.text(
    0.9,
    .97,
    f"Black circles represent the Precision and Recall values at p-value threshold of {pthr}",
    ha="right",
    va="center",
    fontsize=12,
    fontstyle="italic",
    color="gray",
)
# Finalize the plot
plt.tight_layout()
plots.finalize_plot(
    fig, 
    show=True,
    save=save_to_folder,
    filename=f"{simID}_GroupingBenchmark_PR_Curve", 
    filepath=figure_path,
    formats=figure_formats, 
    transparent=transparent_bg,
    dpi=figure_dpi,
)
```

##### MCC Curves: Threshold Sensitivity¶

2×4 panel grid showing MCC across -log₁₀(p-value) thresholds. Reveals optimal threshold selection and method stability. MCC interpretation bands shown for reference.

```
best_thresholds = groupBenchmarkData.groupby(["Method", "Experiment", "DataType"]).apply(
    lambda x: x.loc[x["MCC"].idxmax(), ["threshold", "MCC"]]
).reset_index()
best_thresholds.columns = ["Method", "Experiment", "DataType", "threshold", "MCC"]
# print("\nBest thresholds per method and experiment:")
# display(best_thresholds)

# MCC Curves for p-value thresholds
fig, axes = plt.subplots(
    2, 4, figsize=(16, 6), 
    sharey=True, sharex=True,
    gridspec_kw={
        "wspace": 0.05, 
        "hspace": 0.05
    }
)
for i, pert in enumerate(groupBenchmarkData['Experiment'].unique()):
    for j, dataType in enumerate(dataTypes):
        cur_data = groupBenchmarkData[
            (groupBenchmarkData["Experiment"] == pert) & 
            (groupBenchmarkData["DataType"] == dataType)
        ].copy()
        cur_data['-log10(threshold)'] = -np.log10(cur_data['threshold'])
        # Plot the MCC curve
        sns.lineplot(
            data=cur_data,
            x="-log10(threshold)",
            y="MCC",
            hue="Method",
            style="Method",
            palette=method_palette,
            ax=axes[j, i],
            # Show the points
            markers=method_markers,
            dashes=False,
            markersize=7.5,
            markeredgewidth=0.5,
            markeredgecolor="gray", 
            legend=False,
            rasterized=True,
            estimator=None
        )
        # Add the best MCC point for each method
        for k, method in enumerate(cur_data["Method"].unique()):
            best_data = best_thresholds[
                (best_thresholds["Method"] == method) & 
                (best_thresholds["Experiment"] == pert) & 
                (best_thresholds["DataType"] == dataType)
            ]
            if not best_data.empty:
                best_threshold = best_data["threshold"].values[0]
                best_mcc = best_data["MCC"].values[0]
                axes[j, i].scatter(
                    -np.log10(best_threshold),
                    best_mcc,
                    color=method_palette[method],
                    s=125,
                    edgecolor="black",
                    linewidth=1.25,
                    marker="o",
                    facecolors="none",
                )
                # # Annotate the best MCC point
                # Add color to match the palette
                color = method_palette[method]
                axes[j, i].text(
                    0.95,
                    0.75 + k * 0.075,
                    f"{method}: {best_mcc:.2f}",
                    color=color,
                    ha="right",
                    va="bottom",
                    fontsize=12,
                    fontweight="bold",
                    transform=axes[j, i].transAxes,
                )

        axes[j, i].set_xlabel("-log10(p-value threshold)", fontsize=12)
        axes[j, i].set_ylabel("MCC", fontsize=12)
        axes[j, i].set_xlim(-0.5, 15.5)
        axes[j, i].set_ylim(-0.25, 1.05)
        axes[j, i].grid("both", linestyle="--", linewidth=0.75, alpha=0.5, color="lightgrey")
        # Add horizontal line at y=0
        # axes[j, i].axhline(0, color="black", linestyle="--", linewidth=1)
        # # Draw MCC interpretation thresholds
        # for thresh, label in mcc_thresholds.items():
        #     axes[j, i].axhline(
        #         thresh, color=mcc_colors[label], alpha=1,
        #         linestyle="dotted", linewidth=1.5, 
        #         label=label, zorder=0
        #         )
            # axes[j, i].text(
            #     0.01,
            #     thresh + 0.02,
            #     label,
            #     color=mcc_colors[label],
            #     ha="left",
            #     va="bottom",
            #     fontsize=10,
            #     fontweight="bold",
            #     transform=axes[j, i].transAxes,
            # )
        # If top row set the title
        if j == 0:
            axes[j, i].set_title(f"{pert}", fontsize=14, fontweight="bold")
        # For the last column, set the text for the dataType on the right
        if i == 3:
            axes[j, i].text(
                1.05,
                0.5,
                f"{dataType.capitalize()}",
                transform=axes[j, i].transAxes,
                ha="left",
                va="center",
                fontsize=14,
                fontweight="bold",
                rotation=270,
            )
# Add a text indicating the circle is the best MCC point under fig.title and above the first subplot
fig.text(
    0.9,
    .97,
    f"Black circles represent the maximum MCC values per method",
    ha="right",
    va="center",
    fontsize=12,
    fontstyle="italic",
    color="gray",
)
# Finalize the plot
plt.tight_layout()
plots.finalize_plot(
    fig, 
    show=True,
    save=save_to_folder,
    filename=f"{simID}_GroupingBenchmark_MCC_Curve", 
    filepath=figure_path,
    formats=figure_formats, 
    transparent=transparent_bg,
    dpi=figure_dpi,
)
```

##### MCC Barplot: Method Comparison¶

Side-by-side barplot comparing mean MCC across perturbation patterns for complete vs imputed data. Error bars show 95% CI. Scatter points indicate MCC at p = 10⁻³. Values below bars show mean MCC scores.

```
# MCC Score for different perturbations and methods at the best threshold

# Create subplots for MCC Score for different perturbations and methods at the best threshold
fig, axes = plt.subplots(
    1, 2, figsize=(12, 5), 
    sharey=True, sharex=True,
    gridspec_kw={"wspace": 0.05, "hspace": 0.1}
)

# Get the number of methods and experiments for proper positioning
methods_list = list(method_palette.keys())
n_methods = len(methods_list)
experiments_list = list(groupBenchmarkData['Experiment'].unique())
n_experiments = len(experiments_list)

for i, dataType in enumerate(dataTypes):
    cur_data = groupBenchmarkData[groupBenchmarkData["DataType"] == dataType]
    sns.barplot(
        ax=axes[i],
        data=cur_data,
        x="Experiment",
        y="MCC",
        hue="Method",
        palette=method_palette,
        edgecolor="black",
        linewidth=0.8,
        estimator='mean',
        errorbar=('ci', 95),
        errwidth=1.5,
        capsize=0.1,
        err_kws={'linewidth': 1.5},
        legend=False,
    )
    
    # Calculate bar width and positioning for 2 methods
    bar_width = 0.8 / n_methods  # Total width of 0.8 divided by number of methods
    
    # Write the average MCC score for each method on the bottom of each bar
    for j, pert in enumerate(experiments_list):
        for k, method in enumerate(methods_list):
            cur_score = cur_data[
                (cur_data["Experiment"] == pert) & 
                (cur_data["Method"] == method)
            ]["MCC"].mean()
            
            # Calculate x position: experiment index + offset for this method
            x_offset = (k - (n_methods - 1) / 2) * bar_width
            x_pos = j + x_offset
            
            axes[i].text(
                x_pos,
                -0.05,
                f"{cur_score:.2f}",
                color=method_palette[method],
                ha="center",
                va="top",
                fontsize=10,
                fontweight="bold",
                rotation=90,
            )
    
    # Get data specifically at the p-value threshold for point overlay
    pthr_data = cur_data[cur_data['threshold'] == pthr]
    
    # Add points for specific p-value threshold
    for k, method in enumerate(methods_list):
        method_data = pthr_data[pthr_data['Method'] == method]
        
        x_positions = []
        y_positions = []
        
        for j, experiment in enumerate(experiments_list):
            mcc_at_pthr = method_data.loc[method_data['Experiment'] == experiment, 'MCC']
            if not mcc_at_pthr.empty:
                # Calculate x position using same logic as text annotations
                x_offset = (k - (n_methods - 1) / 2) * bar_width
                x_pos = j + x_offset
                
                x_positions.append(x_pos)
                y_positions.append(mcc_at_pthr.iloc[0])
        
        # Add scatter points for this method at pthr
        if x_positions:  # Only scatter if we have data
            axes[i].scatter(
                x_positions, 
                y_positions,
                color=method_palette[method],
                s=80,
                edgecolor='black',
                linewidth=1.25,
                marker='o',
                zorder=10,
                alpha=0.9
            )

    # # Draw MCC interpretation thresholds
    # for thresh, label in mcc_thresholds.items():
    #     axes[i].axhline(
    #         thresh, color=mcc_colors[label], alpha=1,
    #         linestyle="dotted", linewidth=1.5, zorder=0
    #     )

    #     # Only draw text for first column to avoid clutter
    #     if i % 2 == 1:
    #         continue
    #     # Use mixed transform: relative x, data y
    #     axes[i].text(
    #         0.01,
    #         thresh,
    #         label,
    #         color=mcc_colors[label],
    #         ha="left",
    #         va="center",  # Center on the line
    #         fontsize=10,
    #         fontweight="bold",
    #         transform=axes[i].get_yaxis_transform(),  # x in axes coords, y in data coords
    #         bbox=dict(boxstyle='round,pad=0.3', facecolor='white', edgecolor=mcc_colors[label], alpha=0.8), 
    #         zorder=0
    #     )
        
    axes[i].set_title(f"{dataType.capitalize()}", fontsize=14, fontweight="bold")
    axes[i].set_xlabel('')
    axes[i].set_ylim(-0.15, 1.0)
    axes[i].grid("y", linestyle="--", linewidth=0.75, alpha=0.5, color="lightgrey")
    axes[i].set_xticklabels(axes[i].get_xticklabels(), rotation=0, ha="center", fontsize=10)
    
    # If first subplot, set the ylabel
    if i == 0:
        axes[i].set_ylabel("MCC Score", fontsize=12)
        # Remove the legend for the first subplot
        axes[i].legend().set_visible(False)
    else:
        axes[i].set_ylabel("")

# Add a text to clarify the various annotations of the plot
fig.text(
    0.9,
    .97,
    f"Black circles represent MCC values at p-value threshold of {pthr}\nMCC interpretation thresholds are indicated by dotted lines",
    ha="right",
    va="center",
    fontsize=12,
    fontstyle="italic",
    color="gray",
)

plt.tight_layout()

plots.finalize_plot(
    fig, 
    show=True,
    save=save_to_folder,
    filename=f"{simID}_GroupingBenchmark_MCC_Bar", 
    filepath=figure_path,
    formats=figure_formats, 
    transparent=transparent_bg,
    dpi=figure_dpi,
)
```

#### Simulation 2 - Impact of Protein and Peptide Missingness on Grouping Performance¶

##### Objective¶

Assess how varying levels of missing data at both protein and peptide levels affect grouping accuracy after imputation.

##### Input Data¶

- **Pattern**: `2_Pro{rate}_Pep{rate}_imputed_{method}_ResultData.feather`
- **Methods**: COPF, ProteoForge

##### Experimental Design¶

| Factor | Levels | Description |
| --- | --- | --- |
| Protein Missingness | 0%, 20%, 40%, 60%, 80% | Fraction of proteins with missing values |
| Peptide Missingness | 0%, 20%, 40%, 60%, 80% | Fraction of peptides with missing values |

##### Key Questions¶

1. At what missingness level does grouping performance degrade significantly?
2. Is protein-level or peptide-level missingness more detrimental?
3. How do the methods compare under extreme missingness (60-80%)?

##### Metrics Generated¶

- AUROC heatmaps across missingness levels
- Mean MCC heatmaps
- MCC at fixed p-value threshold (10⁻³)
- Maximum achievable MCC

```
stTime = utils.getTime()

simID = "Sim2"  
# Set up a path for the simulation
output_path, figure_path = setup_simulation_paths( simID )

methods = list(method_palette.keys())
missRatePros = [0, 0.2, 0.4, 0.6, 0.8]
missRatePeps = [0, 0.2, 0.4, 0.6, 0.8]
missRate_mapper = {
    0: '0%',
    0.2: '20%',
    0.4: '40%',
    0.6: '60%',
    0.8: '80%',
}

print("PROTEOFORM GROUPING BENCHMARK ANALYSIS")
print("-" * 52)
print(f"Simulation ID: {simID}")
print(f"Methods: {', '.join(methods)} ({len(methods)} total)")
print(f"Missed Protein Rates: {', '.join([missRate_mapper[mr] for mr in missRatePros])}")
print(f"Missed Peptide Rates: {', '.join([missRate_mapper[mr] for mr in missRatePeps])}")
print(f"Output Path: {output_path}")

# Progress tracking
total_combinations = len(methods) * len(missRatePros) * len(missRatePeps)
files_found = 0
files_missing = 0
current_combination = 0
print(f"Total combinations to process: {total_combinations}")
print("-" * 52)

display("Calculating Grouping Benchmarks:")

combined_results = []
for cur_method in methods:
    for missRatePro in missRatePros:
        for missRatePep in missRatePeps:
            current_combination += 1
            
            # Load the result data
            result_file = f"{output_path}2_Pro{missRatePro}_Pep{missRatePep}_imputed_{cur_method}_ResultData.feather"
            
            if os.path.exists(result_file):
                res_df = pd.read_feather(result_file)
                files_found += 1

                # Show progress for successful loads only
                if verbose:
                    progress = f"[{current_combination:2d}/{total_combinations}]"
                    print(f"{progress} ✓ {cur_method:12} | {missRate_mapper[missRatePro]:7} | {missRate_mapper[missRatePep]:7}")
            else:
                files_missing += 1
                if verbose:
                    progress = f"[{current_combination:2d}/{total_combinations}]"
                    print(f"{progress} ✗ {cur_method:12} | {missRate_mapper[missRatePro]:7} | {missRate_mapper[missRatePep]:7} (MISSING)")
                continue

            # Handle COPF-specific column renaming
            if cur_method == "COPF":
                res_df = res_df.rename(columns={
                    "proteoform_score_pval": "adj_pval",
                    'protein_id': "Protein"
                })
                metric_data = utils.grouping_performance_copf(
                    data=res_df,
                    thresholds=thresholds,
                    pvalue_col='adj_pval',
                    protein_col='Protein',
                    cluster_col='cluster',
                    perturbation_col='pertPFG',
                )

            elif cur_method == "ProteoForge":
                metric_data = utils.grouping_performance_proteoforge(
                    data=res_df,
                    thresholds=thresholds,
                    pvalue_col='adj_pval',
                    protein_col='Protein',
                    cluster_col='ClusterID',
                    perturbation_col='pertPFG',
                )
            
            metric_data['Method'] = cur_method
            metric_data['ProteinMissingness'] = missRatePro
            metric_data['PeptideMissingness'] = missRatePep
            combined_results.append(metric_data)

# Combine and process data
groupBenchmarkData = pd.concat(combined_results, ignore_index=True)
groupBenchmarkData['ProteinMissingness'] = groupBenchmarkData['ProteinMissingness'].map(missRate_mapper)
groupBenchmarkData['PeptideMissingness'] = groupBenchmarkData['PeptideMissingness'].map(missRate_mapper)

print(f"\nDATA PREVIEW:")
display(groupBenchmarkData.head(3))
# Save the processed data
output_file = f"{output_path}4_{simID}_Grouping_PerformanceData.feather"
groupBenchmarkData.to_feather(output_file)
groupBenchmarkData.to_csv(output_file.replace('.feather', '.csv'), index=False)

print(f"\nRESULTS SUMMARY:")
print("=" * 52)
execTime = utils.prettyTimer(utils.getTime() - stTime)
print(f"Total execution time:         {execTime}")
print(f"Files processed successfully: {files_found}")
print(f"Files missing/skipped:        {files_missing}")
print(f"Processing success rate:      {files_found/(files_found+files_missing)*100:.1f}%")
print("-" * 52)
print(f"Final dataset shape:          {groupBenchmarkData.shape}")
print(f"Unique Missed Protein Rates:  {groupBenchmarkData['ProteinMissingness'].nunique()}")
print(f"Unique Missed Peptide Rates:  {groupBenchmarkData['PeptideMissingness'].nunique()}")
print(f"Methods analyzed:             {groupBenchmarkData['Method'].nunique()}")
print(f"Data saved to: {output_file}")
print("=" * 52)
```

```
PROTEOFORM GROUPING BENCHMARK ANALYSIS
----------------------------------------------------
Simulation ID: Sim2
Methods: COPF, ProteoForge (2 total)
Missed Protein Rates: 0%, 20%, 40%, 60%, 80%
Missed Peptide Rates: 0%, 20%, 40%, 60%, 80%
Output Path: ./data/Sim2/
Total combinations to process: 50
----------------------------------------------------
```

```
'Calculating Grouping Benchmarks:'
```

```
[ 1/50] ✓ COPF         | 0%      | 0%     
[ 2/50] ✓ COPF         | 0%      | 20%    
[ 3/50] ✓ COPF         | 0%      | 40%    
[ 4/50] ✓ COPF         | 0%      | 60%    
[ 5/50] ✓ COPF         | 0%      | 80%    
[ 6/50] ✓ COPF         | 20%     | 0%     
[ 7/50] ✓ COPF         | 20%     | 20%    
[ 8/50] ✓ COPF         | 20%     | 40%    
[ 9/50] ✓ COPF         | 20%     | 60%    
[10/50] ✓ COPF         | 20%     | 80%    
[11/50] ✓ COPF         | 40%     | 0%     
[12/50] ✓ COPF         | 40%     | 20%    
[13/50] ✓ COPF         | 40%     | 40%    
[14/50] ✓ COPF         | 40%     | 60%    
[15/50] ✓ COPF         | 40%     | 80%    
[16/50] ✓ COPF         | 60%     | 0%     
[17/50] ✓ COPF         | 60%     | 20%    
[18/50] ✓ COPF         | 60%     | 40%    
[19/50] ✓ COPF         | 60%     | 60%    
[20/50] ✓ COPF         | 60%     | 80%    
[21/50] ✓ COPF         | 80%     | 0%     
[22/50] ✓ COPF         | 80%     | 20%    
[23/50] ✓ COPF         | 80%     | 40%    
[24/50] ✓ COPF         | 80%     | 60%    
[25/50] ✓ COPF         | 80%     | 80%    
[26/50] ✓ ProteoForge  | 0%      | 0%     
[27/50] ✓ ProteoForge  | 0%      | 20%    
[28/50] ✓ ProteoForge  | 0%      | 40%    
[29/50] ✓ ProteoForge  | 0%      | 60%    
[30/50] ✓ ProteoForge  | 0%      | 80%    
[31/50] ✓ ProteoForge  | 20%     | 0%     
[32/50] ✓ ProteoForge  | 20%     | 20%    
[33/50] ✓ ProteoForge  | 20%     | 40%    
[34/50] ✓ ProteoForge  | 20%     | 60%    
[35/50] ✓ ProteoForge  | 20%     | 80%    
[36/50] ✓ ProteoForge  | 40%     | 0%     
[37/50] ✓ ProteoForge  | 40%     | 20%    
[38/50] ✓ ProteoForge  | 40%     | 40%    
[39/50] ✓ ProteoForge  | 40%     | 60%    
[40/50] ✓ ProteoForge  | 40%     | 80%    
[41/50] ✓ ProteoForge  | 60%     | 0%     
[42/50] ✓ ProteoForge  | 60%     | 20%    
[43/50] ✓ ProteoForge  | 60%     | 40%    
[44/50] ✓ ProteoForge  | 60%     | 60%    
[45/50] ✓ ProteoForge  | 60%     | 80%    
[46/50] ✓ ProteoForge  | 80%     | 0%     
[47/50] ✓ ProteoForge  | 80%     | 20%    
[48/50] ✓ ProteoForge  | 80%     | 40%    
[49/50] ✓ ProteoForge  | 80%     | 60%    
[50/50] ✓ ProteoForge  | 80%     | 80%    

DATA PREVIEW:
```

|  | TP | FP | TN | FN | TPR | FPR | FDR | MCC | Precision | Recall | F1 | threshold | Method | ProteinMissingness | PeptideMissingness |
| --- | --- | --- | --- | --- | --- | --- | --- | --- | --- | --- | --- | --- | --- | --- | --- |
| 0 | 0 | 0 | 250 | 250 | 0.0000 | 0.0000 | 0.0000 | 0.0000 | 0.0000 | 0.0000 | 0.0000 | 0.0000 | COPF | 0% | 0% |
| 1 | 0 | 0 | 250 | 250 | 0.0000 | 0.0000 | 0.0000 | 0.0000 | 0.0000 | 0.0000 | 0.0000 | 0.0000 | COPF | 0% | 0% |
| 2 | 0 | 0 | 250 | 250 | 0.0000 | 0.0000 | 0.0000 | 0.0000 | 0.0000 | 0.0000 | 0.0000 | 0.0000 | COPF | 0% | 0% |

```
RESULTS SUMMARY:
====================================================
Total execution time:         00h:00m:49s
Files processed successfully: 50
Files missing/skipped:        0
Processing success rate:      100.0%
----------------------------------------------------
Final dataset shape:          (1250, 15)
Unique Missed Protein Rates:  5
Unique Missed Peptide Rates:  5
Methods analyzed:             2
Data saved to: ./data/Sim2/4_Sim2_Grouping_PerformanceData.feather
====================================================
```

##### AUROC Heatmap: Overall Discrimination¶

Side-by-side heatmaps showing AUROC scores across protein (x-axis) and peptide (y-axis) missingness levels. Higher values (darker) indicate better grouping discrimination.

```
# Comparing AUC Values across different missingness levels
# Calculate AUC for ROC and PR curves
auc_results = []
for (method, pro_miss, pep_miss), group in groupBenchmarkData.groupby(
        ['Method', 'ProteinMissingness', 'PeptideMissingness']
    ):
    # ROC AUC
    group = utils.complete_curve_data(group, 'ROC', 'FPR', 'TPR')
    roc_data = group.sort_values('FPR')
    roc_auc = np.trapezoid(roc_data['TPR'], roc_data['FPR'])
    
    
    auc_results.append({
        'Method': method,
        'ProteinMissingness': pro_miss,
        'PeptideMissingness': pep_miss,
        'AUC': roc_auc,
    })
auc_df = pd.DataFrame(auc_results)
print("\nAUC Data Preview:")
display(auc_df.head())

# Initialize Figure for 3 Methods with dedicated colorbar column
fig, axes = plt.subplots(
    nrows=1, ncols=3, figsize=(9, 4), 
    gridspec_kw={
        "wspace": 0.05, "hspace": 0.1,
        "width_ratios": [0.9, 0.9,  0.1]  # First 3 columns same size, 4th for colorbar
    },
)

vmin = auc_df["AUC"].min()
vmax = auc_df["AUC"].max()

# Create heatmaps for the first 3 methods
for i, cur_method in enumerate(methods):
    plot_data = auc_df[auc_df["Method"] == cur_method].pivot_table(
        index="PeptideMissingness",
        columns="ProteinMissingness",
        values="AUC",
        aggfunc="mean"
    )
    # Make sure 0,0 is in the bottom left corner
    plot_data = plot_data.iloc[::-1]
    
    # Only show colorbar on the last method heatmap
    cb = (i == len(methods) - 1)
    
    # If this is the last method, create colorbar in the 4th column
    if cb:
        cbar_ax = axes[2]
    else:
        cbar_ax = None
    
    sns.heatmap(
        plot_data,
        vmin=vmin,
        vmax=vmax,
        cmap="Blues",
        annot=True,
        fmt=".2f",
        ax=axes[i],
        cbar=cb,
        cbar_ax=cbar_ax,
        cbar_kws={"label": "AUROC"},
        square=True,
        # Annotation size
        annot_kws={"size": 12}, 
    )
    
    if i == 0:
        axes[i].set_ylabel("Peptide Missingness Rate", fontsize=14, fontweight="bold")
    else:
        axes[i].set_ylabel("")
        # Remove yticklabels
        axes[i].set_yticklabels([])
    if i == 1:
        axes[i].set_xlabel("Protein Missingness Rate", fontsize=14, fontweight="bold")
    else:
        axes[i].set_xlabel("")

    axes[i].set_title(f"{cur_method}", fontsize=14, fontweight="bold")

# Hide the 4th axis if it wasn't used for colorbar (this handles the case where we have < 3 methods)
if len(methods) < 2:
    axes[2].set_visible(False)

plt.tight_layout()
plt.suptitle("Peptide Identification Benchmark with AUROC", fontsize=16, fontweight="bold")
plt.subplots_adjust(top=0.85) 

plots.finalize_plot(
    fig, 
    show=True,
    save=save_to_folder,
    filename=f"{simID}_GroupingBenchmark_AUC_Heatmap", 
    filepath=figure_path,
    formats=figure_formats, 
    transparent=transparent_bg,
    dpi=figure_dpi,
)
```

```
AUC Data Preview:
```

|  | Method | ProteinMissingness | PeptideMissingness | AUC |
| --- | --- | --- | --- | --- |
| 0 | COPF | 0% | 0% | 0.6132 |
| 1 | COPF | 0% | 20% | 0.6209 |
| 2 | COPF | 0% | 40% | 0.5970 |
| 3 | COPF | 0% | 60% | 0.5861 |
| 4 | COPF | 0% | 80% | 0.6170 |

##### Mean MCC Heatmap: Average Performance¶

Side-by-side heatmaps showing mean MCC across all p-value thresholds. Indicates overall classification quality independent of threshold selection.

```
# Mean MCC Scores
# Heatmap of the MCC values for the different methods 
# (Protein Missingness and Peptide Missingness) (X: Protein Missingness, Y: Peptide Missingness)

# Initalize Figure for 3 Methods
fig, axes = plt.subplots(
    nrows=1, ncols=3, figsize=(9, 4), 
    gridspec_kw={
        "wspace": 0.05, "hspace": 0.1,
        "width_ratios": [0.9, 0.9, 0.1]  # First 3 columns same size, 4th for colorbar
    },
)

vmin = groupBenchmarkData["MCC"].min()
vmax = groupBenchmarkData["MCC"].max()

# Create heatmaps for the first 3 methods
for i, cur_method in enumerate(methods):
    plot_data = groupBenchmarkData[groupBenchmarkData["Method"] == cur_method].pivot_table(
        index="PeptideMissingness",
        columns="ProteinMissingness",
        values="MCC",
        aggfunc="mean"
    )
    # Make sure 0,0 is in the bottom left corner
    plot_data = plot_data.iloc[::-1]
    
    # Only show colorbar on the last method heatmap
    cb = (i == len(methods) - 1)
    
    # If this is the last method, create colorbar in the 4th column
    if cb:
        cbar_ax = axes[2]
    else:
        cbar_ax = None
    
    sns.heatmap(
        plot_data,
        vmin=vmin,
        vmax=vmax,
        cmap="Purples",
        annot=True,
        fmt=".2f",
        ax=axes[i],
        cbar=cb,
        cbar_ax=cbar_ax,
        cbar_kws={"label": "Mean MCC"},
        square=True,
        # Annotation size
        annot_kws={"size": 12}, 
    )
    
    if i == 0:
        axes[i].set_ylabel("Peptide Missingness Rate", fontsize=14, fontweight="bold")
    else:
        axes[i].set_ylabel("")
        # Remove yticklabels
        axes[i].set_yticklabels([])
    if i == 1:
        axes[i].set_xlabel("Protein Missingness Rate", fontsize=14, fontweight="bold")
    else:
        axes[i].set_xlabel("")

    axes[i].set_title(f"{cur_method}", fontsize=14, fontweight="bold")


# Hide the 4th axis if it wasn't used for colorbar (this handles the case where we have < 3 methods)
if len(methods) < 2:
    axes[2].set_visible(False)

plt.tight_layout()
plt.suptitle("Peptide Grouping Benchmark with MCC (Mean)", fontsize=16, fontweight="bold")
plt.subplots_adjust(top=0.85)

plots.finalize_plot(
    fig, 
    show=True,
    save=save_to_folder,
    filename=f"{simID}_GroupingBenchmark_MCC_mean_Heatmap",
    filepath=figure_path,
    formats=figure_formats, 
    transparent=transparent_bg,
    dpi=figure_dpi,
)
```

##### MCC at p-Threshold Heatmap: Fixed Threshold Performance¶

Side-by-side heatmaps showing MCC at the standard p = 10⁻³ threshold. Represents practical performance at a commonly used significance level.

```
# MCC Score at pThr
# Heatmap of the MCC values for the different methods 
# (Protein Missingness and Peptide Missingness) (X: Protein Missingness, Y: Peptide Missingness)

fig, axes = plt.subplots(
    nrows=1, ncols=3, figsize=(9, 4), 
    gridspec_kw={
        "wspace": 0.05, "hspace": 0.1,
        "width_ratios": [0.9, 0.9, 0.1]
    },
)

vmin = groupBenchmarkData["MCC"].min()
vmax = groupBenchmarkData["MCC"].max()

# Create heatmaps for the first 3 methods
for i, cur_method in enumerate(methods):
    plot_data = groupBenchmarkData[
        (groupBenchmarkData["Method"] == cur_method) & 
        (groupBenchmarkData["threshold"] == pthr)
    ].pivot_table(
        index="PeptideMissingness",
        columns="ProteinMissingness",
        values="MCC",
        aggfunc="mean"
    )

    # Make sure 0,0 is in the bottom left corner
    plot_data = plot_data.iloc[::-1]
    
    # Only show colorbar on the last method heatmap
    cb = (i == len(methods) - 1)
    
    # If this is the last method, create colorbar in the 4th column
    if cb:
        cbar_ax = axes[2]
    else:
        cbar_ax = None
    
    sns.heatmap(
        plot_data,
        vmin=vmin,
        vmax=vmax,
        cmap="Purples",
        annot=True,
        fmt=".2f",
        ax=axes[i],
        cbar=cb,
        cbar_ax=cbar_ax,
        cbar_kws={"label": f"MCC at p-value {pthr}"},
        square=True,
        # Annotation size
        annot_kws={"size": 12}, 
    )
    
    if i == 0:
        axes[i].set_ylabel("Peptide Missingness Rate", fontsize=14, fontweight="bold")
    else:
        axes[i].set_ylabel("")
        # Remove yticklabels
        axes[i].set_yticklabels([])
    if i == 1:
        axes[i].set_xlabel("Protein Missingness Rate", fontsize=14, fontweight="bold")
    else:
        axes[i].set_xlabel("")

    axes[i].set_title(f"{cur_method}", fontsize=14, fontweight="bold")


# Hide the 4th axis if it wasn't used for colorbar (this handles the case where we have < 3 methods)
if len(methods) < 2:
    axes[2].set_visible(False)

plt.tight_layout()
plt.suptitle(f"Peptide Grouping Benchmark with MCC at p-value {pthr}", fontsize=16, fontweight="bold")
plt.subplots_adjust(top=0.85)

plots.finalize_plot(
    fig, 
    show=True,
    save=save_to_folder,
    filename=f"{simID}_GroupingBenchmark_MCC_pThr_Heatmap",
    filepath=figure_path,
    formats=figure_formats, 
    transparent=transparent_bg,
    dpi=figure_dpi,
)
```

##### Max MCC Heatmap: Best Achievable Performance¶

Side-by-side heatmaps showing maximum MCC achievable at any threshold. Represents the upper bound of method capability under each missingness condition.

```
# Max MCC Scores
# Heatmap of the MCC values for the different methods 
# (Protein Missingness and Peptide Missingness) (X: Protein Missingness, Y: Peptide Missingness)

# Initalize Figure for 3 Methods
fig, axes = plt.subplots(
    nrows=1, ncols=3, figsize=(9, 4), 
    gridspec_kw={
        "wspace": 0.05, "hspace": 0.1,
        "width_ratios": [0.9, 0.9, 0.1]  # First 3 columns same size, 4th for colorbar
    },
)

vmin = groupBenchmarkData["MCC"].min()
vmax = groupBenchmarkData["MCC"].max()

# Create heatmaps for the first 3 methods
for i, cur_method in enumerate(methods):
    plot_data = groupBenchmarkData[groupBenchmarkData["Method"] == cur_method].pivot_table(
        index="PeptideMissingness",
        columns="ProteinMissingness",
        values="MCC",
        aggfunc="max"
    )
    # Make sure 0,0 is in the bottom left corner
    plot_data = plot_data.iloc[::-1]
    
    # Only show colorbar on the last method heatmap
    cb = (i == len(methods) - 1)
    
    # If this is the last method, create colorbar in the 4th column
    if cb:
        cbar_ax = axes[2]
    else:
        cbar_ax = None
    
    sns.heatmap(
        plot_data,
        vmin=vmin,
        vmax=vmax,
        cmap="Purples",
        annot=True,
        fmt=".2f",
        ax=axes[i],
        cbar=cb,
        cbar_ax=cbar_ax,
        cbar_kws={"label": "Max MCC"},
        square=True,
        # Annotation size
        annot_kws={"size": 12}, 
    )
    
    if i == 0:
        axes[i].set_ylabel("Peptide Missingness Rate", fontsize=14, fontweight="bold")
    else:
        axes[i].set_ylabel("")
        # Remove yticklabels
        axes[i].set_yticklabels([])
    if i == 1:
        axes[i].set_xlabel("Protein Missingness Rate", fontsize=14, fontweight="bold")
    else:
        axes[i].set_xlabel("")

    axes[i].set_title(f"{cur_method}", fontsize=14, fontweight="bold")


# Hide the 4th axis if it wasn't used for colorbar (this handles the case where we have < 3 methods)
if len(methods) < 2:
    axes[2].set_visible(False)

plt.tight_layout()
plt.suptitle("Peptide Grouping Benchmark with MCC (Max)", fontsize=16, fontweight="bold")
plt.subplots_adjust(top=0.85)

plots.finalize_plot(
    fig, 
    show=True,
    save=save_to_folder,
    filename=f"{simID}_GroupingBenchmark_MCC_max_Heatmap",
    filepath=figure_path,
    formats=figure_formats, 
    transparent=transparent_bg,
    dpi=figure_dpi,
)
```

#### Simulation 3 - Impact of Perturbation Magnitude on Grouping Performance¶

##### Objective¶

Evaluate how the magnitude of fold-change affects the ability to correctly group perturbed peptides together.

##### Input Data¶

- **Pattern**: `2_{low}_{high}_{method}_ResultData.feather`
- **Methods**: COPF, ProteoForge

##### Experimental Design¶

| Factor | Levels | Description |
| --- | --- | --- |
| Magnitude Range | 0.1-0.25, 0.25-0.50, ..., 1.75-2.0 | Log₂ fold-change ranges (8 levels) |

**Magnitude Interpretation**: Ranges represent log₂ fold-change bounds. For example, 0.75-1.0 corresponds to ~1.7x-2x linear fold-change.

##### Key Questions¶

1. What is the minimum detectable effect size for accurate grouping?
2. At what magnitude do methods achieve "Strong" (>0.7) MCC?
3. Does performance plateau at high magnitudes?

##### Metrics Generated¶

- MCC sensitivity curves across magnitude ranges
- AUROC heatmap by method and magnitude

```
stTime = utils.getTime()

simID = "Sim3"  
# Set up a path for the simulation
output_path, figure_path = setup_simulation_paths( simID )

methods = list(method_palette.keys())
perturbationRanges = [
    (0.1, 0.25),
    (0.25, 0.50),
    (0.50, 0.75),
    (0.75, 1.0),
    (1.0, 1.25),
    (1.25, 1.50),
    (1.50, 1.75),
    (1.75, 2.0),
]

perturbation_mapper = {
    (0.1, 0.25): '0.1-0.25',
    (0.25, 0.50): '0.25-0.50',
    (0.50, 0.75): '0.50-0.75',
    (0.75, 1.0): '0.75-1.0',
    (1.0, 1.25): '1.0-1.25',
    (1.25, 1.50): '1.25-1.50',
    (1.50, 1.75): '1.50-1.75',
    (1.75, 2.0): '1.75-2.0',
}

print("PROTEOFORM GROUPING BENCHMARK ANALYSIS")
print("-" * 52)
print(f"Simulation ID: {simID}")
print(f"Methods: {', '.join(methods)} ({len(methods)} total)")
print(f"Perturbation Ranges: {', '.join([perturbation_mapper[pr] for pr in perturbationRanges])}")
print(f"Output Path: {output_path}")    

# Progress tracking
total_combinations = len(methods) * len(perturbationRanges)
files_found = 0
files_missing = 0
current_combination = 0
print(f"Total combinations to process: {total_combinations}")
print("-" * 52)

display("Calculating Grouping Benchmarks:")

combined_results = []
for cur_method in methods:
    for pertRange in perturbationRanges:
        current_combination += 1
        
        # Load the result data
        result_file = f"{output_path}2_{pertRange[0]}_{pertRange[1]}_{cur_method}_ResultData.feather"
        
        if os.path.exists(result_file):
            res_df = pd.read_feather(result_file)
            files_found += 1

            # Show progress for successful loads only
            if verbose:
                progress = f"[{current_combination:2d}/{total_combinations}]"
                print(f"{progress} ✓ {cur_method:12} | {perturbation_mapper[pertRange]:9}")
        else:
            files_missing += 1
            if verbose:
                progress = f"[{current_combination:2d}/{total_combinations}]"
                print(f"{progress} ✗ {cur_method:12} | {perturbation_mapper[pertRange]:9} (MISSING)")
            continue

        # Handle COPF-specific column renaming
        if cur_method == "COPF":
            res_df = res_df.rename(columns={
                "proteoform_score_pval": "adj_pval",
                'protein_id': "Protein"
            })
            metric_data = utils.grouping_performance_copf(
                data=res_df,
                thresholds=thresholds,
                pvalue_col='adj_pval',
                protein_col='Protein',
                cluster_col='cluster',
                perturbation_col='pertPFG',
            )

        elif cur_method == "ProteoForge":
            metric_data = utils.grouping_performance_proteoforge(
                data=res_df,
                thresholds=thresholds,
                pvalue_col='adj_pval',
                protein_col='Protein',
                cluster_col='ClusterID',
                perturbation_col='pertPFG',
            )
        
        metric_data['Method'] = cur_method
        metric_data['PerturbationRange'] = perturbation_mapper[pertRange]
        combined_results.append(metric_data)

# Combine and process data
groupBenchmarkData = pd.concat(combined_results, ignore_index=True)
groupBenchmarkData['PerturbationRange'] = pd.Categorical(
    groupBenchmarkData['PerturbationRange'], 
    categories=[perturbation_mapper[pr] for pr in perturbationRanges],
    ordered=True
)

print(f"\nDATA PREVIEW:")
display(groupBenchmarkData.head(3))

# Save the processed data
output_file = f"{output_path}4_{simID}_Grouping_PerformanceData.feather"
groupBenchmarkData.to_feather(output_file)
groupBenchmarkData.to_csv(output_file.replace('.feather', '.csv'), index=False)

print(f"\nRESULTS SUMMARY:")
print("=" * 52)
execTime = utils.prettyTimer(utils.getTime() - stTime)
print(f"Total execution time:         {execTime}")
print(f"Files processed successfully: {files_found}")
print(f"Files missing/skipped:        {files_missing}")
print(f"Processing success rate:      {files_found/(files_found+files_missing)*100:.1f}%")
print("-" * 52)
print(f"Final dataset shape:          {groupBenchmarkData.shape}")
print(f"Unique Perturbation Ranges:   {groupBenchmarkData['PerturbationRange'].nunique()}")
print(f"Methods analyzed:             {groupBenchmarkData['Method'].nunique()}")
print(f"Data saved to: {output_file}")
print("=" * 52)
```

```
PROTEOFORM GROUPING BENCHMARK ANALYSIS
----------------------------------------------------
Simulation ID: Sim3
Methods: COPF, ProteoForge (2 total)
Perturbation Ranges: 0.1-0.25, 0.25-0.50, 0.50-0.75, 0.75-1.0, 1.0-1.25, 1.25-1.50, 1.50-1.75, 1.75-2.0
Output Path: ./data/Sim3/
Total combinations to process: 16
----------------------------------------------------
```

```
'Calculating Grouping Benchmarks:'
```

```
[ 1/16] ✓ COPF         | 0.1-0.25 
[ 2/16] ✓ COPF         | 0.25-0.50
[ 3/16] ✓ COPF         | 0.50-0.75
[ 4/16] ✓ COPF         | 0.75-1.0 
[ 5/16] ✓ COPF         | 1.0-1.25 
[ 6/16] ✓ COPF         | 1.25-1.50
[ 7/16] ✓ COPF         | 1.50-1.75
[ 8/16] ✓ COPF         | 1.75-2.0 
[ 9/16] ✓ ProteoForge  | 0.1-0.25 
[10/16] ✓ ProteoForge  | 0.25-0.50
[11/16] ✓ ProteoForge  | 0.50-0.75
[12/16] ✓ ProteoForge  | 0.75-1.0 
[13/16] ✓ ProteoForge  | 1.0-1.25 
[14/16] ✓ ProteoForge  | 1.25-1.50
[15/16] ✓ ProteoForge  | 1.50-1.75
[16/16] ✓ ProteoForge  | 1.75-2.0 

DATA PREVIEW:
```

|  | TP | FP | TN | FN | TPR | FPR | FDR | MCC | Precision | Recall | F1 | threshold | Method | PerturbationRange |
| --- | --- | --- | --- | --- | --- | --- | --- | --- | --- | --- | --- | --- | --- | --- |
| 0 | 0 | 0 | 322 | 178 | 0.0000 | 0.0000 | 0.0000 | 0.0000 | 0.0000 | 0.0000 | 0.0000 | 0.0000 | COPF | 0.1-0.25 |
| 1 | 0 | 0 | 322 | 178 | 0.0000 | 0.0000 | 0.0000 | 0.0000 | 0.0000 | 0.0000 | 0.0000 | 0.0000 | COPF | 0.1-0.25 |
| 2 | 0 | 0 | 322 | 178 | 0.0000 | 0.0000 | 0.0000 | 0.0000 | 0.0000 | 0.0000 | 0.0000 | 0.0000 | COPF | 0.1-0.25 |

```
RESULTS SUMMARY:
====================================================
Total execution time:         00h:00m:15s
Files processed successfully: 16
Files missing/skipped:        0
Processing success rate:      100.0%
----------------------------------------------------
Final dataset shape:          (400, 14)
Unique Perturbation Ranges:   8
Methods analyzed:             2
Data saved to: ./data/Sim3/4_Sim3_Grouping_PerformanceData.feather
====================================================
```

##### MCC Sensitivity Curves: Magnitude Response¶

Line plot showing mean MCC across perturbation magnitude ranges with 95% CI bands. MCC interpretation thresholds shown as reference lines. Black markers indicate MCC at p = 10⁻³.

```
# Line plot showing MCC scores across perturbation ranges 
# line plot marker at the mean MCC and CI as a shaded area
# for each method and data type
fig, ax = plt.subplots(1, 1, figsize=(12, 6))

sns.lineplot(
    ax=ax,
    data=groupBenchmarkData,
    x="PerturbationRange",
    y="MCC",
    hue="Method",
    style="Method",
    palette=method_palette,
    linewidth=3,
    markers=method_markers,
    markersize=12,
    markeredgewidth=0.5,
    markeredgecolor="black", 
    
    # Estimation and confidence interval
    estimator='mean',
    errorbar=('ci', 95),
    err_style="band",
    rasterized=True,
)

ax.set_xlabel("Perturbation Range", fontsize=14, fontweight="bold")
ax.set_ylabel("MCC Score", fontsize=14, fontweight="bold")
ax.set_title("Peptide Identification Benchmark with MCC across Perturbation Ranges", fontsize=16, fontweight="bold")
ax.set_ylim(-0.15, 1.0)
ax.grid("y", linestyle="--", linewidth=0.75, alpha=0.5, color="lightgrey")
# ax.set_xticklabels(ax.get_xticklabels(), rotation=45, ha="right", fontsize=10)
ax.legend(
    loc='upper right', title="Method", fontsize=10, title_fontsize=12, 
    frameon=False, 
)   

# # Draw MCC interpretation thresholds
# for thresh, label in mcc_thresholds.items():
#     ax.axhline(
#         thresh, color=mcc_colors[label], alpha=1,
#         linestyle="dotted", linewidth=1.5, 
#         label=label, zorder=0
#         )
#     # Use mixed transform: relative x, data y
#     ax.text(
#         0.01,
#         thresh,
#         label,
#         color=mcc_colors[label],
#         ha="left",
#         va="center",  # Center on the line
#         fontsize=10,
#         fontweight="bold",
#         transform=ax.get_yaxis_transform(),  # x in axes coords, y in data coords
#         bbox=dict(boxstyle='round,pad=0.3', facecolor='white', edgecolor=mcc_colors[label], alpha=0.8)
#     )

# Add smaller scatter points for MCC values at p-value threshold
pthr_data = groupBenchmarkData[groupBenchmarkData['threshold'] == pthr]
for i, method in enumerate(method_palette.keys()):
    method_data = pthr_data[pthr_data['Method'] == method]
    ax.scatter(
        method_data['PerturbationRange'],
        method_data['MCC'],
        color=method_palette[method],
        s=50,
        edgecolor='black',
        linewidth=1.25,
        marker=method_markers[method],
        zorder=10,
        alpha=0.9
    )

# Add a text to clarify the various annotations of the plot
# Include the p-value threshold in the text
fig.text(
    1,
    1.05,
    f"Black markers represent the MCC values at p-value threshold of {pthr}\nMCC interpretation thresholds are indicated by dotted lines\nShaded areas represent 95% confidence intervals around the mean MCC",
    ha="right",
    va="center",
    fontsize=12,
    fontstyle="italic",
    color="gray",
)
# Finalize the plot
plt.tight_layout()
plots.finalize_plot(
    fig, 
    show=True,
    save=save_to_folder,
    filename=f"{simID}_GroupingBenchmark_MCC_Curve_Perturbation",
    filepath=figure_path,
    formats=figure_formats, 
    transparent=transparent_bg,
    dpi=figure_dpi,
)
```

##### AUROC Heatmap: Discrimination by Magnitude¶

Heatmap showing AUROC scores for each method across perturbation magnitude ranges. Visualizes how grouping discrimination improves with larger effect sizes.

```
# Heatmap for AUROC values across perturbation ranges and methods
fig, ax = plt.subplots(1, 1, figsize=(12, 4))

# Calculate AUC for ROC curves
auc_results = []
for (method, pert_range), group in groupBenchmarkData.groupby(
        ['Method', 'PerturbationRange']
    ):
    # ROC AUC
    group = utils.complete_curve_data(group, 'ROC', 'FPR', 'TPR')
    roc_data = group.sort_values('FPR')
    roc_auc = np.trapezoid(roc_data['TPR'], roc_data['FPR'])
    
    auc_results.append({
        'Method': method,
        'PerturbationRange': pert_range,
        'AUC': roc_auc,
    })
auc_df = pd.DataFrame(auc_results)
# print("\nAUC Data Preview:")
# display(auc_df.head())

# Pivot to create the matrix: Methods as rows, PerturbationRange as columns
pivot_data = auc_df.pivot(index='Method', columns='PerturbationRange', values='AUC')

# Create heatmap
sns.heatmap(
    pivot_data,
    annot=True,
    fmt='.2f',
    cmap='Blues',
    cbar_kws={'label': 'AUROC Score', 'shrink': 0.8},
    square=False,
    linewidths=0.5,
    linecolor='white',
    annot_kws={'size': 10, 'weight': 'bold'},
    ax=ax
)

ax.set_title("Peptide Grouping Benchmark with AUROC across Perturbation Ranges", fontsize=16, fontweight="bold")
ax.set_xlabel("Perturbation Range", fontsize=14, fontweight="bold")
ax.set_ylabel("Method", fontsize=14, fontweight="bold")
# Rotate x-axis labels for better readabilitys
ax.set_xticklabels(ax.get_xticklabels(), rotation=0, ha="center", fontsize=10)
ax.set_yticklabels(ax.get_yticklabels(), rotation=0, ha="right", fontsize=10)

plt.tight_layout()
plots.finalize_plot(
    fig, 
    show=True,
    save=save_to_folder,
    filename=f"{simID}_GroupingBenchmark_AUROC_Heatmap_Perturbation",
    filepath=figure_path,
    formats=figure_formats, 
    transparent=transparent_bg,
    dpi=figure_dpi,
)
```

#### Simulation 4 - Impact of Experimental Complexity on Grouping Performance¶

##### Objective¶

Assess how increasing the number of experimental conditions affects peptide grouping accuracy across different overlap and direction scenarios.

##### Input Data¶

- **Pattern**: `2_{N}Cond_{overlap}_{dir}Dir_{method}_ResultData.feather`
- **Methods**: COPF, ProteoForge

##### Experimental Design¶

| Factor | Levels | Description |
| --- | --- | --- |
| Conditions | 2, 3, 4, 5, 6 | Number of experimental conditions |
| Overlap | Overlap, NonOverlap | Whether perturbed groups share peptides |
| Direction | same, random | Perturbation direction consistency |

##### Key Questions¶

1. Does grouping accuracy scale with experimental complexity?
2. How does peptide overlap between groups affect performance?
3. Does consistent vs. random perturbation direction matter for grouping?

##### Metrics Generated¶

- Four-panel MCC barplot (Overlap × Direction facets)

```
stTime = utils.getTime()

simID = "Sim4"  
# Set up a path for the simulation
output_path, figure_path = setup_simulation_paths( simID )

methods = list(method_palette.keys())

# Systematic parameter combinations
overlap_types = [True, False]    # Overlap vs Non-overlap
direction_types = ["random", "same"]  # Direction of perturbations

# Number of conditions to generate (with shifts)
conditions = {
    2: [0, .5],
    3: [0, .5, 1],
    4: [0, .5, 1, 1.5],
    5: [0, .5, 1, 1.5, 2],
    6: [0, .5, 1, 1.5, 2, 2.5]
}


print("PROTEOFORM GROUPING BENCHMARK ANALYSIS")
print("-" * 52)
print(f"Simulation ID: {simID}")
print(f"Methods: {', '.join(methods)} ({len(methods)} total)")
print(f"Overlap Types: {', '.join(['Overlap' if ot else 'Non-overlap' for ot in overlap_types])}")
print(f"Direction Types: {', '.join(direction_types)}")
print(f"Conditions: {', '.join([str(c) for c in conditions.keys()])} (with shifts)")
print(f"Output Path: {output_path}")

# Progress tracking
total_combinations = len(methods) * len(overlap_types) * len(direction_types) * len(conditions)
files_found = 0
files_missing = 0
current_combination = 0
print(f"Total combinations to process: {total_combinations}")
print("-" * 52)

display("Calculating Grouping Benchmarks:")

combined_results = []
for cur_method in methods:
    for cur_overlap in overlap_types:
        for cur_direction in direction_types:
            for num_conds, cond_shifts in conditions.items():
                current_combination += 1
                
                overlap_str = "Overlap" if cur_overlap else "NonOverlap"
                shifts_str = "random" if cur_direction == "random" else "same"

                result_file = f"{output_path}2_{num_conds}Cond_{overlap_str}_{shifts_str}Dir_{cur_method}_ResultData.feather"

                
                if os.path.exists(result_file):
                    res_df = pd.read_feather(result_file)
                    files_found += 1

                    if verbose:
                        progress = f"[{current_combination:2d}/{total_combinations}]"
                        print(f"{progress} ✓ {cur_method:12} | {num_conds} Conditions | {overlap_str} | {shifts_str}")
                else:
                    files_missing += 1
                    if verbose:
                        progress = f"[{current_combination:2d}/{total_combinations}]"
                        print(f"{progress} ✗ {cur_method:12} | {num_conds} Conditions | {overlap_str} | {shifts_str} (MISSING)")
                    continue

                # Handle COPF-specific column renaming
                if cur_method == "COPF":
                    res_df = res_df.rename(columns={
                        "proteoform_score_pval": "adj_pval",
                        'protein_id': "Protein"
                    })
                    metric_data = utils.grouping_performance_copf(
                        data=res_df,
                        thresholds=thresholds,
                        pvalue_col='adj_pval',
                        protein_col='Protein',
                        cluster_col='cluster',
                        perturbation_col='pertPFG',
                    )

                elif cur_method == "ProteoForge":
                    metric_data = utils.grouping_performance_proteoforge(
                        data=res_df,
                        thresholds=thresholds,
                        pvalue_col='adj_pval',
                        protein_col='Protein',
                        cluster_col='ClusterID',
                        perturbation_col='pertPFG',
                    )
                
                metric_data['Method'] = cur_method
                metric_data['Overlap'] = cur_overlap
                metric_data['Direction'] = cur_direction
                metric_data['N_Conditions'] = num_conds
                metric_data['Shifts'] = ','.join([str(s) for s in cond_shifts])
                combined_results.append(metric_data)

# Combine and process data
groupBenchmarkData = pd.concat(combined_results, ignore_index=True)
#Save the processed data
output_file = f"{output_path}4_{simID}_Grouping_PerformanceData.feather"
groupBenchmarkData.to_feather(output_file)
groupBenchmarkData.to_csv(output_file.replace('.feather', '.csv'), index=False)
```

```
PROTEOFORM GROUPING BENCHMARK ANALYSIS
----------------------------------------------------
Simulation ID: Sim4
Methods: COPF, ProteoForge (2 total)
Overlap Types: Overlap, Non-overlap
Direction Types: random, same
Conditions: 2, 3, 4, 5, 6 (with shifts)
Output Path: ./data/Sim4/
Total combinations to process: 40
----------------------------------------------------
```

```
'Calculating Grouping Benchmarks:'
```

```
[ 1/40] ✓ COPF         | 2 Conditions | Overlap | random
[ 2/40] ✓ COPF         | 3 Conditions | Overlap | random
[ 3/40] ✓ COPF         | 4 Conditions | Overlap | random
[ 4/40] ✓ COPF         | 5 Conditions | Overlap | random
[ 5/40] ✓ COPF         | 6 Conditions | Overlap | random
[ 6/40] ✓ COPF         | 2 Conditions | Overlap | same
[ 7/40] ✓ COPF         | 3 Conditions | Overlap | same
[ 8/40] ✓ COPF         | 4 Conditions | Overlap | same
[ 9/40] ✓ COPF         | 5 Conditions | Overlap | same
[10/40] ✓ COPF         | 6 Conditions | Overlap | same
[11/40] ✓ COPF         | 2 Conditions | NonOverlap | random
[12/40] ✓ COPF         | 3 Conditions | NonOverlap | random
[13/40] ✓ COPF         | 4 Conditions | NonOverlap | random
[14/40] ✓ COPF         | 5 Conditions | NonOverlap | random
[15/40] ✓ COPF         | 6 Conditions | NonOverlap | random
[16/40] ✓ COPF         | 2 Conditions | NonOverlap | same
[17/40] ✓ COPF         | 3 Conditions | NonOverlap | same
[18/40] ✓ COPF         | 4 Conditions | NonOverlap | same
[19/40] ✓ COPF         | 5 Conditions | NonOverlap | same
[20/40] ✓ COPF         | 6 Conditions | NonOverlap | same
[21/40] ✓ ProteoForge  | 2 Conditions | Overlap | random
[22/40] ✓ ProteoForge  | 3 Conditions | Overlap | random
[23/40] ✓ ProteoForge  | 4 Conditions | Overlap | random
[24/40] ✓ ProteoForge  | 5 Conditions | Overlap | random
[25/40] ✓ ProteoForge  | 6 Conditions | Overlap | random
[26/40] ✓ ProteoForge  | 2 Conditions | Overlap | same
[27/40] ✓ ProteoForge  | 3 Conditions | Overlap | same
[28/40] ✓ ProteoForge  | 4 Conditions | Overlap | same
[29/40] ✓ ProteoForge  | 5 Conditions | Overlap | same
[30/40] ✓ ProteoForge  | 6 Conditions | Overlap | same
[31/40] ✓ ProteoForge  | 2 Conditions | NonOverlap | random
[32/40] ✓ ProteoForge  | 3 Conditions | NonOverlap | random
[33/40] ✓ ProteoForge  | 4 Conditions | NonOverlap | random
[34/40] ✓ ProteoForge  | 5 Conditions | NonOverlap | random
[35/40] ✓ ProteoForge  | 6 Conditions | NonOverlap | random
[36/40] ✓ ProteoForge  | 2 Conditions | NonOverlap | same
[37/40] ✓ ProteoForge  | 3 Conditions | NonOverlap | same
[38/40] ✓ ProteoForge  | 4 Conditions | NonOverlap | same
[39/40] ✓ ProteoForge  | 5 Conditions | NonOverlap | same
[40/40] ✓ ProteoForge  | 6 Conditions | NonOverlap | same
```

##### Comprehensive Barplot: Factor Interactions¶

2×2 panel barplot showing MCC across number of conditions, stratified by overlap type (rows) and perturbation direction (columns). Error bars show 95% CI. Scatter points indicate MCC at p = 10⁻³. Values below bars show mean MCC scores.

```
# Comprehensive bar plot with all perturbation ranges and statistical details
fig, axes = plt.subplots(
    2, 2, figsize=(16, 10), 
    sharey=True, sharex=True,
    gridspec_kw={
        "hspace": 0.01,  # Reduced vertical spacing
        "wspace": 0.01  # Minimal horizontal spacing
    }
)

plot_combinations = [
    (True, "same"),
    (True, "random"),
    (False, "same"),
    (False, "random"),
]

# Store handles and labels for shared legend
handles, labels = None, None

for i, (overlap, direction) in enumerate(plot_combinations):
    cur_data = groupBenchmarkData[
        (groupBenchmarkData["Overlap"] == overlap) & 
        (groupBenchmarkData["Direction"] == direction)
    ]
    ax = axes[i//2, i%2]

    # Get unique conditions and methods for proper positioning
    unique_conditions = sorted(cur_data['N_Conditions'].unique())
    methods = list(method_palette.keys())
    n_methods = len(methods)
    # Calculate bar width and positioning (seaborn default spacing)
    bar_width = 0.8 / n_methods  # Total width of 0.8 divided by number of methods

    bar_plot = sns.barplot(
        ax=ax,
        data=cur_data,
        x="N_Conditions",
        y="MCC",
        hue="Method",
        palette=method_palette,
        edgecolor='black',
        errwidth=1.5,
        capsize=0.1,
        dodge=True,
        estimator='mean',
        errorbar=('ci', 95),
        legend=False  # Disable individual legends
    )
    
    # Store legend info from first plot
    if i == 0:
        handles, labels = ax.get_legend_handles_labels()
    
    # Set direction as title for each column
    if i // 2 == 0:  # Top row
        direction_title = f"{direction.capitalize()} Direction"
        ax.set_title(direction_title, fontsize=14, fontweight="bold")
    
    # Add overlap/non-overlap labels on the right side
    if i % 2 == 1:  # Right column
        overlap_label = "Overlap" if overlap else "Non-overlap"
        ax.text(
            1.05, 0.5, 
            overlap_label,
            transform=ax.transAxes,
            rotation=270,
            va="center",
            ha="left",
            fontsize=14,
            fontweight="bold"
        )
    
    # Configure axis labels
    if i // 2 == 1:  # Bottom row - show x-axis labels
        ax.set_xlabel("Number of Conditions", fontsize=12, fontweight="bold")
    else:  # Top row - hide x-axis labels
        ax.set_xlabel("")
        # ax.set_xticklabels([])
    
    if i % 2 == 0:  # Left column - show y-axis labels
        ax.set_ylabel("Mean MCC Score", fontsize=12, fontweight="bold")
    else:  # Right column - hide y-axis labels
        ax.set_ylabel("")
    
    # Styling
    ax.set_ylim(-0.15, 1.0)
    ax.grid("y", linestyle="--", linewidth=0.75, alpha=0.5, color="lightgrey")

    # Write the average MCC score for each method at the bottom of each bar group
    for j, condition in enumerate(unique_conditions):
        for k, method in enumerate(methods):
            avg_score = cur_data[
                (cur_data["N_Conditions"] == condition) &
                (cur_data["Method"] == method)
            ]["MCC"].mean()
            if pd.notna(avg_score):
                ax.text(
                    j + (k - n_methods / 2) * bar_width + bar_width / 2,  # Center above each bar
                    -0.05,  # Slightly below x-axis
                    f"{avg_score:.2f}",
                    ha="center",
                    va="top",
                    fontsize=9,
                    color=method_palette[method],
                    rotation=90,
                    fontweight="bold"
                )
    # # Draw MCC interpretation thresholds
    # for thresh, label in mcc_thresholds.items():
    #     ax.axhline(
    #         thresh, color=mcc_colors[label], alpha=1,
    #         linestyle="dotted", linewidth=1.5, 
    #         label=label, zorder=0
    #         )
    #     # Only draw text for first column to avoid clutter
    #     if i % 2 == 1:
    #         continue
    #     # Use mixed transform: relative x, data y
    #     ax.text(
    #         0.01,
    #         thresh,
    #         label,
    #         color=mcc_colors[label],
    #         ha="left",
    #         va="center",  # Center on the line
    #         fontsize=10,
    #         fontweight="bold",
    #         transform=ax.get_yaxis_transform(),  # x in axes coords, y in data coords
    #         bbox=dict(boxstyle='round,pad=0.3', facecolor='white', edgecolor=mcc_colors[label], alpha=0.8)
    #     )

    # Add scatter points for MCC values at p-value threshold
    pthr_data = cur_data[cur_data['threshold'] == pthr]
    
    for j, method in enumerate(methods):
        method_data = pthr_data[pthr_data['Method'] == method]
        
        x_positions = []
        y_positions = []
        
        for condition in unique_conditions:
            condition_data = method_data[method_data['N_Conditions'] == condition]
            if not condition_data.empty:
                # Calculate x position: condition index + offset for this method
                condition_idx = unique_conditions.index(condition)
                x_pos = condition_idx + (j - n_methods / 2) * bar_width + bar_width / 2
                x_positions.append(x_pos)
                y_positions.append(condition_data['MCC'].values[0])

        ax.scatter(
            x_positions,
            y_positions,
            color=method_palette[method],
            s=100,
            edgecolor='black',
            linewidth=1.25,
            marker=method_markers[method],
            zorder=10,
            alpha=0.9
        )

# Create a legend for methods using marker and color
from matplotlib.lines import Line2D
method_legend_elements = [
    Line2D(
        [0], [0],
        marker=method_markers[m],
        color='w',
        markerfacecolor=method_palette[m],
        markeredgecolor='black',
        markersize=12,
        linewidth=0,
        label=m
    )
    for m in methods
]

fig.legend(
    handles=method_legend_elements,
    loc='lower center',
    ncol=len(methods),
    fontsize=12,
    title="Method",
    title_fontsize=14,
    frameon=False,
    bbox_to_anchor=(0.5, 0.05)
)

# Adjust layout to accommodate legend
plt.tight_layout()
plt.subplots_adjust(bottom=0.15)

plt.suptitle("Peptide Grouping Benchmark with MCC across Conditions", fontsize=16, fontweight="bold")
plt.subplots_adjust(top=0.92)
plots.finalize_plot(
    fig, 
    show=True,
    save=save_to_folder,
    filename=f"{simID}_GroupingBenchmark_MCC_Barplot_Conditions",
    filepath=figure_path,
    formats=figure_formats, 
    transparent=transparent_bg,
    dpi=figure_dpi,
)
```


---

#### Summary¶

##### Output Files Generated¶

| Simulation | File | Description |
| --- | --- | --- |
| Sim1 | `Sim1_GroupingBenchmark_ROC_Curve.pdf` | ROC curves by perturbation pattern |
| Sim1 | `Sim1_GroupingBenchmark_PR_Curve.pdf` | PR curves by perturbation pattern |
| Sim1 | `Sim1_GroupingBenchmark_MCC_Curve.pdf` | MCC vs threshold curves |
| Sim1 | `Sim1_GroupingBenchmark_MCC_Bar.pdf` | MCC barplot comparison |
| Sim2 | `Sim2_GroupingBenchmark_AUROC_Heatmap.pdf` | AUROC across missingness |
| Sim2 | `Sim2_GroupingBenchmark_MCC_mean_Heatmap.pdf` | Mean MCC heatmap |
| Sim2 | `Sim2_GroupingBenchmark_MCC_pThr_Heatmap.pdf` | MCC at p-threshold |
| Sim2 | `Sim2_GroupingBenchmark_MCC_max_Heatmap.pdf` | Max MCC heatmap |
| Sim3 | `Sim3_GroupingBenchmark_MCC_Curve_Perturbation.pdf` | MCC sensitivity curves |
| Sim3 | `Sim3_GroupingBenchmark_AUROC_Heatmap_Perturbation.pdf` | AUROC by magnitude |
| Sim4 | `Sim4_GroupingBenchmark_MCC_Barplot_Conditions.pdf` | MCC across conditions |

##### Key Findings by Simulation¶

1. **Sim1 (Imputation)**: Establishes baseline grouping accuracy; compares COPF vs ProteoForge clustering approaches
2. **Sim2 (Missingness)**: Reveals method robustness to missing data at protein and peptide levels
3. **Sim3 (Magnitude)**: Identifies minimum effect size for reliable peptide grouping
4. **Sim4 (Conditions)**: Tests scalability and robustness to overlap/direction scenarios

```
print("Notebook Execution Time:", utils.prettyTimer(utils.getTime() - startTime))
```

```
Notebook Execution Time: 00h:02m:55s
```
