## Supplementary material for "ProteoForge: An Imputation-Aware Framework for Differential Proteoform Discovery in Bottom-Up Proteomics": Analysis notebooks as html render: Notebook S7.html

### Notebook S7

###### Enes Kemal Ergin

#### 2025-12-08 23:00:28

### Supplementary Notebook 7: Comparison of Linear Models Offered by ProteoForge¶

- **License:** Creative Commons Attribution-NonCommercial 4.0 International License
- **Version:** 0.2
- **Edit Log:**
  - 2025-11-28: Initial version of the notebook
  - 2025-12-08: Revise the whole notebook, ensuring clarity and correctness

---

**Requirements:**

The simulation datasets should be created, which is done by `01-SimulatedDatasets.ipynb`. This processe geenrates `Sim{N}` folders in `./data/` directory within the current folder. Sim1 and Sim2 datasets are required for this notebook to properly run.

---

**Data Information:**

Data used in this notebook comes from simulation's 1 and 2, where impact of imputation, and impact of missingness were examined.

---

**Purpose:**

The purpose of this notebook is to benchmark how the linear models (from `statmodels` package) accessed and offered through ProteoForge perform in comparison to each other, and for the ones capable of custom weighting strategies, how different weighting strategies impact the model performance. The models offered by ProteoForge and benchmarked here are:

- ~~`Ordinary Least Squares (OLS/ols)`~~ (This is omited since WLS without weights is equivalent to OLS)
- `Weighted Least Squares (WLS/wls)`, supports custom weights
- `Generalized Linear Model (GLM/glm)`, supports custom weights
- `Robust Linear Model (RLM/rlm)` with Huber's T norm, doesn't support custom weights, but auto-calculates robust weights internally
- `Median Quantile Regression (Median QR/mqr)`, doesn't support custom weights, but is robust to outliers by design

> The contents of this notebook are presented in the Supplementary Note 1 for the main manuscript.

---

#### 1. Setup¶

This section imports required libraries, configures display settings, and defines paths for data and figures.

> **Note:** The HTML rendering of this notebook hides code cells by default. Click the "Code" buttons on the right to expand them.

##### 1.1 Libraries¶

```
import os
import sys
import warnings
warnings.filterwarnings("ignore")

import numpy as np # Numerical computing
import pandas as pd # Data manipulatio

import seaborn as sns # R-like high-level plots
import matplotlib.pyplot as plt # Python's base plotting 
from matplotlib.lines import Line2D # Custom legend handles

sys.path.append('../')

# Utility imports for this analysis
from src import utils, plots

# Main Computational Components of ProteoForge
from ProteoForge import disluster
from ProteoForge import weight, model
from ProteoForge import proteoform_classifier as pfc

# Initialize the timer
startTime = utils.getTime()
```

##### 1.2 Display Settings¶

The cell below configures pandas, matplotlib, and seaborn display options for improved readability of tables and figures, including color palettes and figure export settings.

```
# Define default colors and styles for plots
def_colors = [
    "#139593", "#fca311", "#e54f2a",
    "#c3c3c3", "#555555",
    "#690000", "#5f4a00", "#004549"
]

## Global Variables
seed = 42 # Seed for reproducibility  
pthr = 10**-3  # p-value threshold for significance
thresholds = list(utils.generate_thresholds(10.0, -15, 1, 0, 1, 0.1)) # Thresholds for the analysis

# Models and their plotting styles (removed 'ols' and updated to distinct, non-default colors)
model_palette = {
    "wls": "#139593",   # SeaGreen
    "rlm": "#690000",   # Orange
    "mqr": "#fca311",   # Sky Blue
    "glm": "#004549",   # Rose/Magenta
}
model_styles = {
    "wls": "--",
    "rlm": "-",
    "mqr": ":",
    "glm": (0, (3, 1, 1, 1)),
}
model_markers = {
    "wls": "s",
    "rlm": "^",
    "mqr": "D",
    "glm": "P",
}


# Matthews Correlation Coefficient (MCC) thresholds and colors
mcc_thresholds = {
    0.0 : 'Random',
    0.3 : 'Fair (0.3)',
    0.5 : 'Moderate (0.5)',
    0.7 : 'Strong (0.7)',
    0.9 : 'Almost Perfect (0.9)',
}
mcc_colors = {
    'Random': '#c3c3c3',             
    'Fair (0.3)': '#e0aaff',         
    'Moderate (0.5)': '#9d4edd',     
    'Strong (0.7)': '#5a189a',       
    'Almost Perfect (0.9)': '#240046' 
}


# Set seaborn style
sns.set_theme(
    style="white",
    context="paper",
    palette=def_colors,
    font_scale=1,
    rc={
        "figure.figsize": (6, 4),
        "font.family": "sans-serif",
        "font.sans-serif": ["Arial", "Ubuntu Mono"],
    }
)

# Figure Saving Settings
figure_formats = ["pdf", "png"]
save_to_folder = True
transparent_bg = True
figure_dpi = 300

## Configure dataframe displaying
pd.set_option('display.float_format', lambda x: '%.4f' % x)
pd.set_option('display.max_rows', 50)
pd.set_option('display.max_columns', 50)
pd.set_option('display.width', 500)  # Set a wider display width

## Printing Settings
verbose = True

plots.color_palette(def_colors, save=False, name="Default Colors")
plots.color_palette(list(model_palette.values()), save=False, name="Model Colors")
plots.color_palette(mcc_colors, save=False, name="MCC Interpretation Colors")
```


##### 1.3 Data and Result Paths¶

Data and figures are organized in separate folders:

- `data_path` — Directory for simulation datasets (`data/`).
- `figure_path` — Directory for generated plots and figures (`figures/07-ModelComparison/`).

```
notebook_name = "07-ModelComparison"
data_path = f"./data/"
figure_path = f"./figures/{notebook_name}/"

# Create figure folder structure, if needed
if save_to_folder:
    for i in figure_formats:
        cur_folder = figure_path + i + "/"
        if not os.path.exists(cur_folder):
            os.makedirs(cur_folder)

# Function to format time difference
def format_time_diff(time_diff):
    if time_diff < 60:
        return f"{time_diff:.2f} seconds"
    elif time_diff < 3600:
        return f"{time_diff / 60:.2f} minutes"
    else:
        return f"{time_diff / 3600:.2f} hours"

def min_max_scale(x): return (x - x.min()) / (x.max() - x.min())

correction = {
    'strategy': 'two-step',  # Options: 'global', 'protein-only', 'two-step'
    'methods': ('bonferroni', 'fdr_bh')  # Options first protein-level, then global
}
n_jobs = 16
```

#### 2. Experiment Design¶

This notebook will benchmark ProteoForge's linear model variants across Simulation 1 (perturbation patterns + imputation) and Simulation 2 (missingness levels). It focuses exclusively on running ProteoForge variants (wls, glm, rlm, mqr) while varying weight strategies and using the same preprocessing/clustering pipeline.

Objectives:

- Compare model types fairly using the same weights or controlled weight variants
- Evaluate model sensitivity to imputation (Sim1 complete vs imputed and perturbation patterns)
- Evaluate performance across missingness levels (Sim2: Pro×Pep combinations)
- Produce ROC, MCC, AUC, FDR at p=0.001, and grouping metrics for each model configuration

---

> **Weighting Strategy**
>
> The default weighting strategy for the models that uses custom weights to be passed, uses '90% Imputation + 10% RevTechVar' as default weighting strategy. And others don't need weights passed (RLM, MQR). This default strategy was arbitrarily chosen to represent an approach that places the most importance into the imputation weights. The imputation weights are calculated based again, on an arbitrary basis, to put more importance into the cases where complete missingness is more valuable than sparse missingness, due to them being more likely to represent truly absent peptides. Where sparse missingness' reasons for being missing are more ambiguous and can't really be determined with certainty.

---

##### 2.1 Hypotheses and Expectations¶

- RLM and MQR models (robust regression / quantile regression) will be less sensitive to imputation artifacts than OLS but may trade-off sensitivity for robustness.
- Weighted methods (WLS and GLM with weights) should be more sensitive in detecting perturbed peptides when weights reflect true imputation/reversal tech variability.
- For high missingness (Sim2), the performance gap between weighted and unweighted methods will grow; weights emphasizing imputation quality are expected to improve detection.
- We expect different weight mixing ratios to show a trade-off between sensitivity and false positives. We'll quantify these effects with AUC and MCC.

##### 2.2 Experiment 1: How much missingness impact individual model performance?¶

This experiment uses the Sim2 data, but specifically data from no missingness (complete), 20%, 40%, 60%, and 80% missigness version of the random number of perturbed peptide data. The missingness then is imputed with low-value imputation and kNN imputation for completel condition missingness, and sparse missing values respectively.

```
simID = "Sim2"
combined_results = []
experiments = ['Pro0_Pep0', 'Pro0.2_Pep0.2', 'Pro0.4_Pep0.4', 'Pro0.6_Pep0.6', 'Pro0.8_Pep0.8']
# experiment_file_name_template = f"2_{}_imputed_InputData.feather"
experiments_mapper = {
    'Pro0_Pep0': '0% Missingness',
    'Pro0.2_Pep0.2': '20% Missingness',
    'Pro0.4_Pep0.4': '40% Missingness',
    'Pro0.6_Pep0.6': '60% Missingness',
    'Pro0.8_Pep0.8': '80% Missingness'
}
model_types = ['wls', 'glm', 'mqr', 'rlm']
```

###### 2.2.1 Execute ProteoForge Analysis with Different Models¶

The below is the logic that opens all the datasets to be analyzed and applies ProteoForge's analysis and benchmarking results with different linear model variants.

> This part can be commented out if the results are already computed and saved in the `data/` folder. Unless any changes required/needed re-calculation when restarting the kernel is unnecessary.

```
for exp in experiments:
    experiment_file_name = f"2_{exp}_imputed_InputData.feather"
    input_data = pd.read_feather(f"./data/{simID}/{experiment_file_name}")
    # Calculate weights
    weights_data = weight.generate_weights_data(
        input_data,
        sample_cols=['Sample'],
        log_intensity_col='log10Intensity',
        adj_intensity_col='adjIntensity',
        control_condition='control',
        condition_col='Condition',
        protein_col='Protein',
        peptide_col='Peptide',
        is_real_col='isReal',
        is_comp_miss_col='isCompMiss',
        sparse_imputed_val=1e-5,
        dense_imputed_val=0.5,
        verbose=False,
    )
    input_data['Weight'] = (
        (weights_data['W_Impute'] * 0.90) + 
        (weights_data['W_RevTechVar'] * 0.10)
    )

    for modelType in model_types:
        test_data = input_data.copy()
        model_to_use = 'quantile' if modelType == 'mqr' else modelType
        
        # Set 'Weight' to 1 for models not 'wls' or 'glm
        if modelType not in ['wls', 'glm']:
            test_data['Weight'] = 1.0
        
        cur_model = model.LinearModel(
            data=test_data,
            protein_col="Protein",
            peptide_col="Peptide",
            cond_col="Condition",
            intensity_col="adjIntensity",
            weight_col='Weight',
        )

        test_data = cur_model.run_analysis(
            model_type=model_to_use,
            correction_strategy=correction['strategy'],
            correction_methods=correction['methods'],
            n_jobs=n_jobs
        )

        clusters = disluster.distance_and_cluster(
            data=test_data,
            protein_col='Protein',
            peptide_col='PeptideID',
            cond_col='Condition',
            quant_col='adjIntensity',
            clustering_params={
                'min_clusters': 1,
                'distance_transform': 'corr',
                'clustering_method': 'hybrid_outlier_cut',
                'linkage_method': 'ward',
                'distance_metric': 'euclidean'
            },
            n_jobs=n_jobs,
            verbose=False
        )

        test_data = test_data.merge(
            clusters[['Protein', 'PeptideID', 'cluster_label']],
            on=['Protein', 'PeptideID'],
            how='left'
        ).rename(columns={'cluster_label': 'ClusterID'})

        test_data['pertPeptide'] = np.where(
            test_data['isCompMiss'] == 1, 1, test_data['pertPeptide']
        ).astype(bool)
        test_data['Reason'] = np.where(
            test_data['isCompMiss'] == 1, "Biological Absence", test_data['Reason']
        )
        test_data.loc[
            (test_data['isCompMiss']==1) & (test_data['pertPeptide']), 
            'pertPFG'
        ] = 1

        # Identification Benchmark
        metric_data = utils.create_metric_data(
            test_data, 
            pvalue_thresholds=thresholds,
            label_col='pertPeptide',
            pvalue_col='adj_pval'
        )
        metric_data['Model'] = modelType
        metric_data['Experiment'] = experiments_mapper[exp]
        metric_data['Benchmark'] = 'PepID'
        combined_results.append(metric_data)

        # Peptide Grouping Benchmark
        tmp = test_data[[
            'Protein', 'Peptide','PeptideID', 
            'pertPeptide', 'pertPFG', 'ClusterID', 'adj_pval'
        ]].drop_duplicates().reset_index(drop=True)
        metric_data = utils.grouping_performance_proteoforge(
            data=tmp,
            thresholds=thresholds,
            pvalue_col='adj_pval',
            protein_col='Protein',
            cluster_col='ClusterID',
            perturbation_col='pertPFG',
        )
        metric_data['Model'] = modelType
        metric_data['Experiment'] = experiments_mapper[exp]
        metric_data['Benchmark'] = 'PepGroup'
        combined_results.append(metric_data)

# Combine all results into a single DataFrame
all_results = pd.concat(combined_results, ignore_index=True)
all_results.to_feather(f"./data/{simID}_ModelBenchmark_Results_staticWeight.feather")
```

###### 2.2.2 ROC Curves and AUC Comparison¶

```
all_results = pd.read_feather(f"./data/{simID}_ModelBenchmark_Results_staticWeight.feather")
id_data = all_results[all_results['Benchmark'] == 'PepID'].copy()
pg_data = all_results[all_results['Benchmark'] == 'PepGroup'].copy()
```

```
# ROC Curves
# Facet -> Rows: Benchmark, Columns: Experiment
# Hue: Model, Style: Model, Marker: Model

# Initialize the figure and axes
fig, axes = plt.subplots(
    nrows=2, ncols=len(experiments), figsize=(20, 8), 
    sharex=True, sharey=True,
    constrained_layout=True
)
for i, benchmark in enumerate(['PepID', 'PepGroup']):
    for j, exp in enumerate(experiments):
        ax = axes[i, j]
        subset = all_results[
            (all_results['Benchmark'] == benchmark) & 
            (all_results['Experiment'] == experiments_mapper[exp])
        ].copy()
        if subset.empty:
            ax.set_visible(False)
            continue

        # Build ROC curves per model and compute AUC robustly using the sorted curve
        roc_frames = []
        aucs = {}
        for modelType in model_types:
            model_subset = subset[subset['Model'] == modelType].copy()
            if model_subset.empty:
                continue

            # Create a complete ROC curve for this model, then sort by FPR for AUC calc
            model_roc = utils.complete_curve_data(model_subset, 'ROC', 'FPR', 'TPR')
            if model_roc is None or model_roc.empty:
                continue
            model_roc = model_roc.sort_values('FPR').reset_index(drop=True)
            model_roc['Model'] = modelType
            roc_frames.append(model_roc)

            # Compute AUC using trapezoidal integration on sorted FPR
            try:
                if len(model_roc['FPR'].dropna()) >= 2:
                    auc_val = float(np.trapezoid(model_roc['TPR'].values, model_roc['FPR'].values))
                else:
                    auc_val = np.nan
            except Exception:
                auc_val = np.nan
            aucs[modelType] = auc_val

        if len(roc_frames) == 0:
            ax.set_visible(False)
            continue

        combined_roc_df = pd.concat(roc_frames, ignore_index=True)

        # Plot each model separately to apply specific line style, marker and color
        for modelType in model_types:
            cur_df = combined_roc_df[combined_roc_df['Model'] == modelType]
            if cur_df.empty:
                continue
            auc = aucs.get(modelType, np.nan)
            ax.plot(
                cur_df['FPR'],
                cur_df['TPR'],
                label=f"{modelType}: {auc:.2f}" if not np.isnan(auc) else modelType,
                color=model_palette.get(modelType, 'black'),
                linestyle=model_styles.get(modelType, '-'),
                marker=model_markers.get(modelType, None),
                markersize=6,
                markeredgewidth=0.6,
                alpha=0.95,
                zorder=4
            )

            # Add a large no-facecolor marker to the pthr value for each method if present
            pthr_mask = np.isclose(cur_df['threshold'].astype(float).fillna(np.nan), pthr)
            if pthr_mask.any():
                pthr_row = cur_df.loc[pthr_mask].iloc[0]
                ax.scatter(
                    pthr_row['FPR'],
                    pthr_row['TPR'],
                    color=model_palette.get(modelType, 'black'),
                    s=110,
                    edgecolor='black',
                    linewidth=1.25,
                    marker='o',
                    facecolors='none',
                    zorder=10
                )

        # Add the diagonal line (random classifier)
        ax.plot([0, 1], [0, 1], color="black", linestyle="--", linewidth=0.8, zorder=1)

        # Add AUC legend-like text (ordered descending AUC)
        sorted_auc_items = sorted(aucs.items(), key=lambda x: (np.nan if x[1] is None else -x[1], x[0]))
        for k, (method, auc) in enumerate(sorted_auc_items):
            color = model_palette.get(method, 'black')
            y_pos = 0.05 + k * 0.075
            ax.text(
                0.95,
                y_pos,
                f"{method}: {auc:.2f}" if not np.isnan(auc) else f"{method}: n/a",
                color=color,
                transform=ax.transAxes,
                ha="right",
                va="bottom",
                fontsize=12,
                fontweight="bold",
            )

        ax.set_xlim(-0.05, 1.05)
        ax.set_ylim(-0.05, 1.05)
        ax.grid(True, linestyle="--", linewidth=0.6, alpha=0.45, color="lightgrey")

        # Titles
        if i == 0:
            ax.set_title(f"{experiments_mapper[exp]}", fontsize=13, fontweight="bold")
            ax.set_xlabel("")
        if i == 1:
            ax.set_xlabel("False Positive Rate (FPR)", fontsize=12)

        # Remove y-axis label and tick labels for inner columns
        if j != 0:
            ax.set_ylabel("")                  # remove y-axis label text
            ax.tick_params(axis="y", labelleft=False)  # hide y tick labels
        else:
            ax.set_ylabel("True Positive Rate (TPR)", fontsize=12)

# Add Benchmark labels to right side of the rows
for i, benchmark in enumerate(['PepID', 'PepGroup']):
    ax = axes[i, -1]
    ax.text(
        1.0,
        0.5,
        benchmark,
        transform=ax.transAxes,
        rotation=270,
        fontsize=13,
        fontweight="bold",
        va="center"
    )

plt.tight_layout()
```


---

###### 2.2.3 MCC Across p-Value Thresholds Comparison¶

```
# MCC over thresholds (faceted per benchmark x experiment)
# Compute best thresholds per Model/Benchmark/Experiment for annotation
best_thresholds = all_results.groupby(["Benchmark", "Experiment", "Model"]).apply(
    lambda x: x.loc[x["MCC"].idxmax(), ["threshold", "MCC"]]
).reset_index()
best_thresholds.columns = ["Benchmark", "Experiment", "Model", "threshold", "MCC"]

# Build the grid: rows = benchmarks, cols = experiments
benchmarks = ["PepID", "PepGroup"]
cols = list(experiments)  # same order as experiments list
fig, axes = plt.subplots(
    nrows=len(benchmarks), ncols=len(cols), figsize=(20, 8), sharey='row', constrained_layout=True,
)

for i, benchmark in enumerate(benchmarks):
    for j, exp in enumerate(cols):
        ax = axes[i, j]
        subset = all_results[
            (all_results['Benchmark'] == benchmark) &
            (all_results['Experiment'] == experiments_mapper[exp])
        ].copy()
        if subset.empty:
            ax.set_visible(False)
            continue

        # Convert threshold to float safely and avoid log10(0)
        subset['threshold'] = subset['threshold'].astype(float)
        subset['-log10(threshold)'] = -np.log10(subset['threshold'])

        # Plot each model separately to keep color/marker/style
        for modelType in model_types:
            cur_df = subset[subset['Model'] == modelType].copy()
            if cur_df.empty:
                continue

            ax.plot(
                cur_df['-log10(threshold)'],
                cur_df['MCC'],
                label=modelType,
                color=model_palette.get(modelType, 'black'),
                linestyle=model_styles.get(modelType, '-'),
                marker=model_markers.get(modelType, None),
                markersize=7.5,
                markeredgewidth=0.6,
                alpha=0.95,
                zorder=4,
            )

            # Add the best MCC point and annotation for this Model/Benchmark/Experiment
            best_data = best_thresholds[
                (best_thresholds['Model'] == modelType) &
                (best_thresholds['Benchmark'] == benchmark) &
                (best_thresholds['Experiment'] == experiments_mapper[exp])
            ]
            if not best_data.empty:
                best_threshold = float(best_data['threshold'].values[0])
                best_mcc = float(best_data['MCC'].values[0])
                ax.scatter(
                    -np.log10(best_threshold),
                    best_mcc,
                    color=model_palette.get(modelType, 'black'),
                    s=125,
                    edgecolor='black',
                    linewidth=1.25,
                    marker='o',
                    facecolors='none',
                )
                ax.text(
                    0.95,
                    0.70 + (list(model_types).index(modelType)) * 0.075,
                    f"{modelType}: {best_mcc:.2f}",
                    color=model_palette.get(modelType, 'black'),
                    ha='right', va='bottom', transform=ax.transAxes,
                    fontsize=11, fontweight='bold'
                )

        ax.set_xlim(-0.5, 15.5)
        ax.set_ylim(-0.05, 1.05)
        ax.set_xlabel("-log10(threshold)", fontsize=11)
        if j == 0:
            ax.set_ylabel("MCC", fontsize=12)
        else:
            ax.set_ylabel("")
            ax.tick_params(axis='y', labelleft=False)

        ax.grid(True, linestyle='--', linewidth=0.75, alpha=0.5, color='lightgrey')

        # Title for top row
        if i == 0:
            ax.set_title(f"{experiments_mapper[exp]}", fontsize=13, fontweight='bold')

# Add Benchmark row labels on the right side
for i, benchmark in enumerate(benchmarks):
    ax = axes[i, -1]
    ax.text(
        1.01, 0.5, benchmark, transform=ax.transAxes, rotation=270,
        fontsize=13, fontweight='bold', va='center'
    )

plt.tight_layout()
```


---

###### 2.2.4 High-Summary Figure with Mean MCC Across Missingness Levels¶

This is the default summary figure that can be used to compare model performances using mean mcc from all thresholds calculated across missingness levels in a compact manner.

```
# --- 1. Data Prep ---
safe_df = all_results.copy()
if 'DataType' not in safe_df.columns:
    safe_df['DataType'] = 'imputed'

mcc_by_exp = (
    safe_df.groupby(['Benchmark', 'Experiment', 'Model', 'DataType'], dropna=False)['MCC']
    .mean()
    .reset_index()
)

# Parse X-axis values
exp_labels = [experiments_mapper[e] for e in experiments]
def parse_pct(label):
    try:
        return int(str(label).split('%')[0])
    except Exception:
        return 0 # Fallback or handle accordingly
exp_x = [parse_pct(l) for l in exp_labels]

data_types = sorted(mcc_by_exp['DataType'].dropna().unique()) if 'DataType' in mcc_by_exp.columns else ['imputed']
benchmarks = ['PepID', 'PepGroup']

# --- 2. Figure Setup ---
# Use gridspec_kw to tighten the gap between rows since they share X
fig, axes = plt.subplots(
    nrows=2, ncols=1,
    figsize=(6, 8),      # Slightly reduced height to be more compact
    sharex=True, sharey=True,
    gridspec_kw={'hspace': 0.1} 
)

if not isinstance(axes, (list, np.ndarray)):
    axes = [axes]

# --- 3. Plotting Loop ---
for ax, benchmark in zip(axes, benchmarks):
    df_bench = mcc_by_exp[mcc_by_exp['Benchmark'] == benchmark]
    
    # Aesthetic: Grid behind plot, remove top/right spines
    ax.grid(True, which='major', axis='both', linestyle='--', linewidth=0.5, alpha=0.4, zorder=0)
    ax.spines['top'].set_visible(False)
    ax.spines['right'].set_visible(False)

    for modelType in model_types:
        df_model = df_bench[df_bench['Model'] == modelType]
        if df_model.empty:
            continue

        for dt in data_types:
            df_line = df_model[df_model['DataType'] == dt]
            
            # Align Y values to explicit X positions
            y_vals = []
            for label in exp_labels:
                row = df_line[df_line['Experiment'] == label]
                if row.empty:
                    y_vals.append(np.nan)
                else:
                    val = row['MCC'].values[0]
                    y_vals.append(float(val) if not np.isnan(val) else np.nan)

            if all(np.isnan(v) for v in y_vals):
                continue

            # Style Logic
            color = model_palette.get(modelType, '#333333')
            linestyle = model_styles.get(modelType, '-')
            marker = model_markers.get(modelType, 'o')
            
            is_imputed = str(dt).lower() in ('imputed', 'imputation', 'i')
            mfc = color if is_imputed else 'white' # 'white' usually looks cleaner than 'none' regarding gridlines
            alpha = 1.0 if is_imputed else 0.9

            ax.plot(
                exp_x, y_vals, 
                label=f"{modelType}", # Label simplified, handled in custom legend
                color=color, 
                linestyle=linestyle,
                marker=marker, 
                markersize=7, 
                markeredgewidth=1.2,
                markeredgecolor=color, 
                markerfacecolor=mfc, 
                alpha=alpha, 
                zorder=3
            )

    # Subplot Specifics
    ax.set_ylabel(benchmark, fontsize=12, fontweight='bold', labelpad=10)
    ax.set_ylim(-0.05, 1.05)
    
    # Add a subtle baseline at 0
    ax.axhline(0, color='black', linewidth=0.8, zorder=1)

# --- 4. Axis Formatting ---
# Only format the bottom axis
axes[-1].set_xticks(exp_x)
axes[-1].set_xticklabels(exp_labels, rotation=0, ha='center') # Rotation 0 is usually better if labels are short numbers
axes[-1].set_xlabel('Missingness (%)', fontsize=11, fontweight='bold')

# Common Y-label (placed manually to be perfectly centered across rows)
# Note: I moved individual titles to Y-labels (above) to save vertical space, 
# but if you prefer the global Y-label, keep this:
fig.text(0.02, 0.5, 'Mean MCC', va='center', rotation='vertical', fontsize=12, fontweight='bold')

# --- 5. Comprehensive Legend ---
# We construct a composite legend: Models on left, Data Types on right (or simply appended)

# A. Model Handles
model_handles = []
for m in model_types:
    if m in mcc_by_exp['Model'].unique():
        c = model_palette.get(m, 'k')
        ls = model_styles.get(m, '-')
        mk = model_markers.get(m, 'o')
        model_handles.append(
            Line2D(
                [0], [0], color=c, linestyle=ls, marker=mk, label=m,
                markerfacecolor=c, markersize=8
            )
        )


# Combine or Separate? 
# For document headers, a single line is best.
all_handles = model_handles + [Line2D([], [], linestyle='')]# + type_handles # Spacer in middle
all_labels = [h.get_label() for h in model_handles]# + [''] + [h.get_label() for h in type_handles]

# Place legend
fig.legend(
    handles=all_handles, 
    labels=all_labels,
    loc='lower center',       # Often better at bottom for documents, or 'upper center'
    bbox_to_anchor=(0.5, 0.90), 
    ncol=len(all_handles),    # Try to fit in one row
    frameon=False,            # Clean look
    fontsize=10,
)

# --- 6. Final Layout ---
fig.suptitle('Performance by Missingness', fontsize=14, fontweight='bold', y=0.98)

# Manual adjust is safer than tight_layout when using fig.text or complex legends
plt.subplots_adjust(
    top=0.90, 
    bottom=0.08, 
    left=0.10, 
    right=0.95, 
    hspace=0.1
)

plt.show()
plots.finalize_plot(
    fig, show=True, save=save_to_folder,
    filename='modelBenchmarks_missingness_MCC_exp1',
    filepath=figure_path,
    formats=figure_formats, 
    transparent=transparent_bg,
    dpi=figure_dpi,
)
```

With the increasing density of the missing values that needs to be imputed the high performing models such as RLM and MQR tend to decrease drastically in their performances. Especially a method like MQR that is robust to outliers by design, tends to suffer greatly from the imputation artifacts that are introduced with high missingness levels starting from early 40% missingness and onwards. The RLM is a method that does still provide similar/comparable performance until 60% missingness, after which it also starts to lose its performance significantly. On the other hand the WLS and GLM methods which are dependent on the weighting strategy provided, tend to perform better than RLM and MQR at high missingness levels, given that the weights provided do reflect the imputation quality well enough. In our default weighting strategy as well as the imputation weighting strategy, we can see that both WLS and GLM outperform RLM and MQR at 80% missingness level, while still providing comparable performance at lower missingness levels.

> However an importat thing to note is that 80% missingness level is an extreme case scenario that is not very likely to be observed in real-life datasets. If the data is that sparse, it is likely needs to be re-generated or looked into for potential issues in the data acquisition process. I even considere 60% missingness to be a very high level that should be considered carefully before proceeding with any analysis.

This can be very useful to get a quick overview of which model performed better across missingness levels in general. Additionally I will likely add this to the supplemantary note 1, as the first figure there.

---

##### 2.3 How the Number of Perturbed Peptides affect model performance?¶

This experiment uses the Sim1 data, where different numbers of peptides were perturbed, and the data is copied to make complete and imputed versions. Where the imputed is generate by 35% of the data is amputated, and then imputed with same strategy as mentioned earlier. We then apply the different linear models offered by ProteoForge to see how the number of perturbed peptides affect the model performances.

```
simID = "Sim1"
combined_results = []
data_types = ['complete', 'imputed']
experiments = ['twoPep', 'randomPep', 'halfPep', 'halfPlusPep']
# experiment_file_name_template = f"2_{experiments}_{data_types}_InputData.feather"
experiments_mapper = {
    'twoPep': '2 Perturbed',
    'randomPep': '2-50% Perturbed',
    'halfPep': '50% Perturbed',
    'halfPlusPep': '60-70% Perturbed'
}
model_types = ['wls', 'glm', 'mqr', 'rlm']
```

###### 2.3.1 Execute ProteoForge Analysis with Different Models¶

The below is the logic that opens all the datasets to be analyzed and applies ProteoForge's analysis and benchmarking results with different linear model variants.

> This part can be commented out if the results are already computed and saved in the `data/` folder. Unless any changes required/needed re-calculation when restarting the kernel is unnecessary.

```
for exp in experiments:
    for data_type in data_types:
        experiment_file_name = f"2_{exp}_{data_type}_InputData.feather"
        input_data = pd.read_feather(f"./data/{simID}/{experiment_file_name}")
        # Calculate weights
        weights_data = weight.generate_weights_data(
            input_data,
            sample_cols=['Sample'],
            log_intensity_col='log10Intensity',
            adj_intensity_col='adjIntensity',
            control_condition='control',
            condition_col='Condition',  
            protein_col='Protein',
            peptide_col='Peptide',
            is_real_col='isReal',
            is_comp_miss_col='isCompMiss',
            sparse_imputed_val=1e-5,
            dense_imputed_val=0.5,
            verbose=False,
        )
        input_data['Weight'] = (
            (weights_data['W_Impute'] * 0.90) + 
            (weights_data['W_RevTechVar'] * 0.10)
        )

        for modelType in model_types:
            test_data = input_data.copy()
            model_to_use = 'quantile' if modelType == 'mqr' else modelType
            
            # Set 'Weight' to 1 for models not 'wls' or 'glm
            if modelType not in ['wls', 'glm']:
                test_data['Weight'] = 1.0
            
            cur_model = model.LinearModel(
                data=test_data,
                protein_col="Protein",
                peptide_col="Peptide",
                cond_col="Condition",
                intensity_col="adjIntensity",
                weight_col='Weight',
            )

            test_data = cur_model.run_analysis(
                model_type=model_to_use,
                correction_strategy=correction['strategy'],
                correction_methods=correction['methods'],
                n_jobs=n_jobs
            )

            clusters = disluster.distance_and_cluster(
                data=test_data,
                protein_col='Protein',
                peptide_col='PeptideID',
                cond_col='Condition',
                quant_col='adjIntensity',
                clustering_params={
                    'min_clusters': 1,
                    'distance_transform': 'corr',
                    'clustering_method': 'hybrid_outlier_cut',
                    'linkage_method': 'ward',
                    'distance_metric': 'euclidean'
                },
                n_jobs=n_jobs,
                verbose=False
            )

            test_data = test_data.merge(
                clusters[['Protein', 'PeptideID', 'cluster_label']],
                on=['Protein', 'PeptideID'],
                how='left'
            ).rename(columns={'cluster_label': 'ClusterID'})

            test_data['pertPeptide'] = np.where(
                test_data['isCompMiss'] == 1, 1, test_data['pertPeptide']
            ).astype(bool)
            test_data['Reason'] = np.where(
                test_data['isCompMiss'] == 1, "Biological Absence", test_data['Reason']
            )
            test_data.loc[
                (test_data['isCompMiss']==1) & (test_data['pertPeptide']), 
                'pertPFG'
            ] = 1

            # Identification Benchmark
            metric_data = utils.create_metric_data(
                test_data, 
                pvalue_thresholds=thresholds,
                label_col='pertPeptide',
                pvalue_col='adj_pval'
            )
            metric_data['Model'] = modelType
            metric_data['Experiment'] = experiments_mapper[exp]
            metric_data['DataType'] = data_type
            metric_data['Benchmark'] = 'PepID'
            combined_results.append(metric_data)

            # Peptide Grouping Benchmark
            tmp = test_data[[
                'Protein', 'Peptide','PeptideID', 
                'pertPeptide', 'pertPFG', 'ClusterID', 'adj_pval'
            ]].drop_duplicates().reset_index(drop=True)
            metric_data = utils.grouping_performance_proteoforge(
                data=tmp,
                thresholds=thresholds,
                pvalue_col='adj_pval',
                protein_col='Protein',
                cluster_col='ClusterID',
                perturbation_col='pertPFG',
            )
            metric_data['Model'] = modelType
            metric_data['Experiment'] = experiments_mapper[exp]
            metric_data['DataType'] = data_type
            metric_data['Benchmark'] = 'PepGroup'
            combined_results.append(metric_data)

# Combine all results into a single DataFrame
all_results = pd.concat(combined_results, ignore_index=True)
all_results.to_feather(f"./data/{simID}_ModelBenchmark_Results_staticWeight.feather")
```

###### 2.3.2 ROC Curves and AUC Comparison¶

**Peptide Identification**s

```
all_results = pd.read_feather(f"./data/{simID}_ModelBenchmark_Results_staticWeight.feather")
id_data = all_results[all_results['Benchmark'] == 'PepID'].copy()
pg_data = all_results[all_results['Benchmark'] == 'PepGroup'].copy()
```

```
# ROC Curves (PepID)
# Rows: Data Type (complete, imputed); Columns: Experiments
fig, axes = plt.subplots(
    nrows=2, ncols=len(experiments), figsize=(20, 8),
    sharex=True, sharey=True, constrained_layout=True
)
for k, data_type in enumerate(['complete', 'imputed']):
    for j, exp in enumerate(experiments):
        ax = axes[k, j]
        subset = all_results[
            (all_results['Benchmark'] == 'PepID') &
            (all_results['DataType'] == data_type) &
            (all_results['Experiment'] == experiments_mapper[exp])
        ].copy()
        if subset.empty:
            ax.set_visible(False)
            continue

        # Build ROC curves per model and compute AUC robustly using the sorted curve
        roc_frames = []
        aucs = {}
        for modelType in model_types:
            model_subset = subset[subset['Model'] == modelType].copy()
            if model_subset.empty:
                continue

            model_roc = utils.complete_curve_data(model_subset, 'ROC', 'FPR', 'TPR')
            if model_roc is None or model_roc.empty:
                continue
            model_roc = model_roc.sort_values('FPR').reset_index(drop=True)
            model_roc['Model'] = modelType
            roc_frames.append(model_roc)

            # Compute AUC using trapezoidal integration on sorted FPR
            try:
                if len(model_roc['FPR'].dropna()) >= 2:
                    auc_val = float(np.trapezoid(model_roc['TPR'].values, model_roc['FPR'].values))
                else:
                    auc_val = np.nan
            except Exception:
                auc_val = np.nan
            aucs[modelType] = auc_val

        if len(roc_frames) == 0:
            ax.set_visible(False)
            continue

        combined_roc_df = pd.concat(roc_frames, ignore_index=True)

        # Plot each model separately to apply specific line style, marker and color
        for modelType in model_types:
            cur_df = combined_roc_df[combined_roc_df['Model'] == modelType]
            if cur_df.empty:
                continue
            auc = aucs.get(modelType, np.nan)
            ax.plot(
                cur_df['FPR'],
                cur_df['TPR'],
                label=f"{modelType}: {auc:.2f}" if not np.isnan(auc) else modelType,
                color=model_palette.get(modelType, 'black'),
                linestyle=model_styles.get(modelType, '-'),
                marker=model_markers.get(modelType, None),
                markersize=6,
                markeredgewidth=0.6,
                alpha=0.95,
                zorder=4,
            )

            # Add a large no-facecolor marker to the pthr value for each method if present
            pthr_mask = np.isclose(cur_df['threshold'].astype(float).fillna(np.nan), pthr)
            if pthr_mask.any():
                pthr_row = cur_df.loc[pthr_mask].iloc[0]
                ax.scatter(
                    pthr_row['FPR'],
                    pthr_row['TPR'],
                    color=model_palette.get(modelType, 'black'),
                    s=110,
                    edgecolor='black',
                    linewidth=1.25,
                    marker='o',
                    facecolors='none',
                    zorder=10,
                )

        # Add the diagonal line (random classifier)
        ax.plot([0, 1], [0, 1], color="black", linestyle="--", linewidth=0.8, zorder=1)

        # Add AUC legend-like text (ordered descending AUC)
        sorted_auc_items = sorted(aucs.items(), key=lambda x: (np.nan if x[1] is None else -x[1], x[0]))
        for kk, (method, auc) in enumerate(sorted_auc_items):
            color = model_palette.get(method, 'black')
            y_pos = 0.05 + kk * 0.075
            ax.text(
                0.95,
                y_pos,
                f"{method}: {auc:.2f}" if not np.isnan(auc) else f"{method}: n/a",
                color=color,
                transform=ax.transAxes,
                ha="right",
                va="bottom",
                fontsize=12,
                fontweight="bold",
            )

        ax.set_xlim(-0.05, 1.05)
        ax.set_ylim(-0.05, 1.05)
        ax.grid(True, linestyle="--", linewidth=0.6, alpha=0.45, color="lightgrey")

        # Titles
        if k == 0:
            ax.set_title(f"{experiments_mapper[exp]}", fontsize=13, fontweight="bold")
            ax.set_xlabel("")
        if k == 1:
            ax.set_xlabel("False Positive Rate (FPR)", fontsize=12)

        # Remove y-axis label and tick labels for inner columns
        if j != 0:
            ax.set_ylabel("")                  # remove y-axis label text
            ax.tick_params(axis="y", labelleft=False)  # hide y tick labels
        else:
            ax.set_ylabel("True Positive Rate (TPR)", fontsize=12)

# Add DataType labels on the right side of the rows
for k, data_type in enumerate(['complete', 'imputed']):
    ax = axes[k, -1]
    ax.text(
        1.0,
        0.5,
        f"PepID - {data_type.capitalize()} Data",
        transform=ax.transAxes,
        rotation=270,
        fontsize=13,
        fontweight="bold",
        va="center",
    )

plt.tight_layout()
```

**Peptide Grouping**

```
# ROC Curves (PepGroup)
# Rows: Data Type (complete, imputed); Columns: Experiments
fig, axes = plt.subplots(
    nrows=2, ncols=len(experiments), figsize=(20, 8),
    sharex=True, sharey=True, constrained_layout=True
)
for k, data_type in enumerate(['complete', 'imputed']):
    for j, exp in enumerate(experiments):
        ax = axes[k, j]
        subset = all_results[
            (all_results['Benchmark'] == 'PepGroup') &
            (all_results['DataType'] == data_type) &
            (all_results['Experiment'] == experiments_mapper[exp])
        ].copy()
        if subset.empty:
            ax.set_visible(False)
            continue

        # Build ROC curves per model and compute AUC robustly using the sorted curve
        roc_frames = []
        aucs = {}
        for modelType in model_types:
            model_subset = subset[subset['Model'] == modelType].copy()
            if model_subset.empty:
                continue

            model_roc = utils.complete_curve_data(model_subset, 'ROC', 'FPR', 'TPR')
            if model_roc is None or model_roc.empty:
                continue
            model_roc = model_roc.sort_values('FPR').reset_index(drop=True)
            model_roc['Model'] = modelType
            roc_frames.append(model_roc)

            # Compute AUC using trapezoidal integration on sorted FPR
            try:
                if len(model_roc['FPR'].dropna()) >= 2:
                    auc_val = float(np.trapezoid(model_roc['TPR'].values, model_roc['FPR'].values))
                else:
                    auc_val = np.nan
            except Exception:
                auc_val = np.nan
            aucs[modelType] = auc_val

        if len(roc_frames) == 0:
            ax.set_visible(False)
            continue

        combined_roc_df = pd.concat(roc_frames, ignore_index=True)

        # Plot each model separately to apply specific line style, marker and color
        for modelType in model_types:
            cur_df = combined_roc_df[combined_roc_df['Model'] == modelType]
            if cur_df.empty:
                continue
            auc = aucs.get(modelType, np.nan)
            ax.plot(
                cur_df['FPR'],
                cur_df['TPR'],
                label=f"{modelType}: {auc:.2f}" if not np.isnan(auc) else modelType,
                color=model_palette.get(modelType, 'black'),
                linestyle=model_styles.get(modelType, '-'),
                marker=model_markers.get(modelType, None),
                markersize=6,
                markeredgewidth=0.6,
                alpha=0.95,
                zorder=4,
            )

            # Add a large no-facecolor marker to the pthr value for each method if present
            pthr_mask = np.isclose(cur_df['threshold'].astype(float).fillna(np.nan), pthr)
            if pthr_mask.any():
                pthr_row = cur_df.loc[pthr_mask].iloc[0]
                ax.scatter(
                    pthr_row['FPR'],
                    pthr_row['TPR'],
                    color=model_palette.get(modelType, 'black'),
                    s=110,
                    edgecolor='black',
                    linewidth=1.25,
                    marker='o',
                    facecolors='none',
                    zorder=10,
                )

        # Add the diagonal line (random classifier)
        ax.plot([0, 1], [0, 1], color="black", linestyle="--", linewidth=0.8, zorder=1)

        # Add AUC legend-like text (ordered descending AUC)
        sorted_auc_items = sorted(aucs.items(), key=lambda x: (np.nan if x[1] is None else -x[1], x[0]))
        for kk, (method, auc) in enumerate(sorted_auc_items):
            color = model_palette.get(method, 'black')
            y_pos = 0.05 + kk * 0.075
            ax.text(
                0.95,
                y_pos,
                f"{method}: {auc:.2f}" if not np.isnan(auc) else f"{method}: n/a",
                color=color,
                transform=ax.transAxes,
                ha="right",
                va="bottom",
                fontsize=12,
                fontweight="bold",
            )

        ax.set_xlim(-0.05, 1.05)
        ax.set_ylim(-0.05, 1.05)
        ax.grid(True, linestyle="--", linewidth=0.6, alpha=0.45, color="lightgrey")

        # Titles
        if k == 0:
            ax.set_title(f"{experiments_mapper[exp]}", fontsize=13, fontweight="bold")
            ax.set_xlabel("")
        if k == 1:
            ax.set_xlabel("False Positive Rate (FPR)", fontsize=12)

        # Remove y-axis label and tick labels for inner columns
        if j != 0:
            ax.set_ylabel("")                  # remove y-axis label text
            ax.tick_params(axis="y", labelleft=False)  # hide y tick labels
        else:
            ax.set_ylabel("True Positive Rate (TPR)", fontsize=12)

# Add DataType labels on the right side of the rows
for k, data_type in enumerate(['complete', 'imputed']):
    ax = axes[k, -1]
    ax.text(
        1.0,
        0.5,
        f"PepGroup - {data_type.capitalize()} Data",
        transform=ax.transAxes,
        rotation=270,
        fontsize=13,
        fontweight="bold",
        va="center",
    )

plt.tight_layout()
```


---

###### 2.3.2 High Level Summary Figure with Mean MCC Across Number of Perturbed Peptides¶

```
# Mean MCC Summary Plot — Summarize mean MCC across Benchmark x Experiment x DataType x Model
# Build MCC summary DataFrame (mean across matched rows in all_results)
mcc_list = []

# Determine data types present (fallback to 'imputed' if missing)
if 'DataType' in all_results.columns:
    data_types = sorted(all_results['DataType'].astype(str).unique())
else:
    data_types = ['imputed']

for benchmark in ['PepID', 'PepGroup']:
    for exp in experiments:
        for dt in data_types:
            for m in model_types:
                subset = all_results[
                    (all_results['Benchmark'] == benchmark) &
                    (all_results['Experiment'] == experiments_mapper[exp]) &
                    (all_results['Model'] == m)
                ].copy()
                if 'DataType' in subset.columns:
                    subset = subset[subset['DataType'] == dt]

                mean_mcc = np.nan
                if 'MCC' in subset.columns and not subset['MCC'].dropna().empty:
                    mean_mcc = float(subset['MCC'].mean())

                mcc_list.append({
                    'Benchmark': benchmark,
                    'Experiment': experiments_mapper[exp],
                    'DataType': dt,
                    'Model': m,
                    'Mean_MCC': mean_mcc
                })

mcc_summary_df = pd.DataFrame(mcc_list)

# Compute delta (IMPUTED - COMPLETE) per Benchmark/Experiment/Model if both data types exist
mcc_pivot = None
if {'complete', 'imputed'}.issubset(set(mcc_summary_df['DataType'])):
    mcc_pivot = mcc_summary_df.pivot_table(
        index=['Benchmark', 'Experiment', 'Model'],
        columns='DataType',
        values='Mean_MCC',
        aggfunc='mean'
    ).reset_index()
    mcc_pivot.columns.name = None
    mcc_pivot['MCC_delta'] = mcc_pivot.get('imputed', np.nan) - mcc_pivot.get('complete', np.nan)
else:
    mcc_pivot = None

# Prepare plotting order
model_order = model_types
data_type_order = ['complete', 'imputed'] if 'complete' in data_types else data_types
experiment_order = [experiments_mapper[e] for e in experiments]

# Plot: Facet rows=Benchmark, cols=Experiment, x=Model, hue=DataType
# Use global styling; do not set theme or save
mg = sns.catplot(
    data=mcc_summary_df,
    kind='bar',
    x='Model',
    y='Mean_MCC',
    hue='DataType',
    col='Experiment',
    row='Benchmark',
    hue_order=data_type_order,
    order=model_order,
    col_order=experiment_order,
    row_order=['PepID', 'PepGroup'],
    height=3.5,
    aspect=1.5,
    sharey=True,
    legend=True,
)

# Styling & annotation: annotate bars and deltas (Imputed - Complete)
for row_idx, row in enumerate(['PepID', 'PepGroup']):
    for col_idx, exp_name in enumerate(experiment_order):
        ax = mg.axes[row_idx, col_idx]
        if ax is None:
            continue

        # annotate each bar height
        for p in ax.patches:
            h = p.get_height()
            if not np.isnan(h):
                ax.annotate(f"{h:.2f}", (p.get_x() + p.get_width() / 2., h), ha='center', va='bottom', fontsize=9)

        # Add delta annotation above the group for each model if computed
        if mcc_pivot is not None:
            for mi, model_name in enumerate(model_order):
                row_mask = (
                    (mcc_pivot['Benchmark'] == row) &
                    (mcc_pivot['Experiment'] == exp_name) &
                    (mcc_pivot['Model'] == model_name)
                )
                if not row_mask.any():
                    continue
                delta_val = mcc_pivot.loc[row_mask, 'MCC_delta'].values[0]
                if np.isnan(delta_val):
                    continue

                sign = '+' if delta_val > 0 else ''
                color = 'green' if delta_val > 0 else 'red' if delta_val < 0 else 'gray'
                delta_label = f"Δ (I - C) {sign}{delta_val:.2f}"

                # Determine center x for the model group bars
                model_patches = [p for p in ax.patches if p.get_x() >= mi - 0.5 and p.get_x() < mi + 0.5]
                if len(model_patches) > 0:
                    x_center = np.mean([p.get_x() + p.get_width() / 2. for p in model_patches])
                else:
                    xticks = ax.get_xticks()
                    x_center = xticks[mi] if mi < len(xticks) else mi

                # compute y position above the higher bar for that model
                model_vals = mcc_summary_df[
                    (mcc_summary_df['Benchmark'] == row) &
                    (mcc_summary_df['Experiment'] == exp_name) &
                    (mcc_summary_df['Model'] == model_name)
                ]
                y_top = 0.0
                if not model_vals.empty:
                    y_top = np.nanmax(model_vals['Mean_MCC'].values)
                y_pos = y_top + 0.04

                ax.text(x_center, y_pos, delta_label, ha='center', va='bottom', fontsize=10, fontweight='bold', color=color)

        ax.set_ylim(-0.05, 1.05)
        ax.set_ylabel('Mean MCC')
        ax.set_xlabel('')
        ax.grid(True, linestyle='--', linewidth=0.6, alpha=0.4, color='lightgrey')

# Tidy up
plt.subplots_adjust(top=0.92)
plt.suptitle('Mean MCC Summary by Model • Benchmark • Experiment • DataType', fontsize=14, fontweight='bold')
plots.finalize_plot(
    plt.gcf(),
    show=True, save=save_to_folder,
    filename='modelBenchmarks_perturbations_MCC_exp2',
    filepath=figure_path,
    formats=figure_formats, 
    transparent=transparent_bg,
    dpi=figure_dpi,
)
```

Looking at the mean MCC across different number of perturbed peptides, couple interesting observations can be made.

- MQR Does work best with all perturbation levels when the data is complete, however its performance drops significantly when imputation is introduced, especially at lower number of perturbations. With imputed versions RLM seems to be more robust across all perturbation levels.
- While at Peptide identification MQR has quite high at complete, peptide grouping, having identifying a lot of individual peptides as significant has down sides, which shows that it is generall the lowest performing model in grouping metrics.
- At grouping metric the RLM also the best performing model with or without imputation, showing its robustness across different scenarios.

> The seemingly contradictory improvement that can be observed across all methods in peptide grouping with imputation, is because by default the amputation strategy some removes about 5% out of 35% of the peptides as completely missing, and they are imputed with low-value imputation, which algorithm can identify meaningful biological signal, to capture this fully, we account for them in the correct labels, which does generally improves the performance metrics across the board for grouping benchmarks.

---

##### 2.4 How Sophistication of Weighting Strategies Can Impact Custom Weighted Models?¶

This section enumerates the weighting strategies used in our benchmarking runs and maps each strategy to the specific capabilities implemented in `ProteoForge/weight.py` (so you can replicate or extend them). The weights returned by these functions are aligned to the original data and are intended to be written into a `Weight` column (used by `wls` and `glm`).

Here we will compare different weighting strategies for WLS and GLM models to see how they affect performance in detecting perturbed peptides. While the other models either doesn't have any weights (OLS), uses median instead of mean (MQR), or auto-calculates weights internally (RLM), for model that required or strengthen by user defined weights proves that selection of weights is necessary to explore. Here in this experiment we will compare different weighting strategies for WLS and GLM:

- no\_weight: set 'Weight' = 1 for all models (baseline)
- default\_mix: calculated weights and (0.9  *W\_Impute + 0.1*  W\_RevTechVar) (current default)
- impute\_only: Weight = W\_Impute only
- revtech\_only: Weight = W\_RevTechVar only
- PLS\_auto: Weight = Auto calculated PLS based weights only

###### 2.4.1 Imputation Weight Strategy¶

This strategy assigns per-row weights based on whether a measurement is real, part of a completely missing peptide for a condition (dense imputation), or a sparsely imputed value. The mapping is:

- is\_real == True → weight = true\_val
- is\_real == False and is\_comp\_miss == True → weight = dense\_imputed\_val
- is\_real == False and is\_comp\_miss == False → weight = sparse\_imputed\_val

**Key behaviors and validations**:

- If neither imputation column is provided the function assumes all values are real and returns an array of ones.
- The function checks that the named columns exist and raises ValueError if they do not.
- All three weight values are cast to floats and must be in [0, 1]; true\_val must be strictly greater than the two imputed values (ensures real measurements are prioritized).
- The returned object is a numpy array of weights aligned with the input dataframe and intended to be written into the dataframe as the `Weight` column for WLS/GLM.

**Rationale and recommended defaults**:

- true\_val (default 1.0): highest confidence for measured values.
- dense\_imputed\_val (default 0.5): moderate weight for values imputed because a peptide was completely missing in a condition (may reflect true biological absence but still useful signal).
- sparse\_imputed\_val (default 1e-5): near-zero weight for isolated imputations (high uncertainty / likely technical artifacts).

**Usage notes**

- Tune dense\_imputed\_val upward if your low-value imputation is known to preserve signal; lower it if dense imputations are mostly noise.
- Keep sparse\_imputed\_val very small to avoid over-counting uncertain imputed entries.
- After generating these imputation weights you can combine them with other components (e.g., reverse-technical-variance, profile-correlation) using linear mixtures (e.g., `0.9*W_Impute + 0.1*W_RevTechVar`) before passing to WLS/GLM.

###### 2.4.2 Customized Function for Generating All Weight Variants¶

The `weight.py` module as of writing comments out majority of the `generate_weights_data` to ensure no additional heavy computations are done when generating weights since they are not used in our analyses as well as benchmarking. Here we will define a custom function that will generate all the weight variants we need for this experiment.

- `W_Impute:` imputation confidence (prioritizes real/dense values vs sparse imputed)
- `W_RepImbalance:` penalty for replicate imbalance across conditions
- `W_NormalizationImpact:` weight for how much normalization affects the peptide/profile
- `W_RevTechVar:` reverse technical-variation score (1 - scaled variance within protein/peptide/condition)
- `W_ProfileCorr:` profile correlation across samples/conditions
- `W_InverseMean(log):` inverse-mean weight on log-intensity (lower mean → higher weight)
- `W_InverseMean(adj):` inverse-mean weight on adjusted-intensity
- `W_InverseVar(log):` inverse-variance weight on log-intensity
- `W_InverseVar(adj):` inverse-variance weight on adjusted-intensity
- `W_InverseStd(log):` inverse-std weight on log-intensity
- `W_InverseStd(adj):` inverse-std weight on adjusted-intensity
- `W_CorrDiscordance(log):` correlation‑discordance on log-intensity (penalizes discordant profiles vs control)
- `W_CorrDiscordance(adj):` correlation‑discordance on adjusted-intensity
- `W_RelativeVariability(log):` relative variability on log-intensity
- `W_RelativeVariability(adj):` relative variability on adjusted-intensity
- `W_SignalToNoise(log):` generalized signal‑to‑noise on log-intensity
- `W_SignalToNoise(adj):` generalized signal‑to‑noise on adjusted-intensity
- `W_ReplicateConcordance(log):` replicate concordance on log-intensity
- `W_ReplicateConcordance(adj):` replicate concordance on adjusted-intensity

###### 2.4.3 PLS Based Auto Calculating the Weights¶

Partial Least Squares (PLS): simple primer

- PLS is a supervised dimensionality‑reduction technique: it finds a small number of components (linear combinations of input features) that maximize covariance with a target variable (here: peptide intensity).
- Compared with PCA (unsupervised), PLS uses the outcome (y) during component extraction, so the components are directly tuned to predict the target.
- In our context, PLS is used to compress many candidate weight components (W\_\*) into a small set of supervised components that best explain the intensity signal; those components are then combined into a single per‑row weight.

What generate\_auto\_pls\_weights does (plain steps)

1. Input
   - X: DataFrame of candidate weight components (columns starting with "W\_").
   - y: Target series (e.g., log10Intensity or adjIntensity).
2. Component scan
   - Fit PLS models with 1..N components (N = min(max\_components, n\_features, n\_samples-1)).
   - Record R² for each number of components.
   - Find an “elbow” in the R² curve to pick an optimal component count (automatically).
3. Feature importance
   - Fit a PLS with the chosen number of components and retrieve the absolute X‑loadings.
   - Weight loadings by incremental R² gains across components to compute an overall importance score per feature.
4. Automatic feature selection
   - Use an elbow on the cumulative importance to choose top‑N features automatically.
5. Refined PLS and final weight
   - Fit a refined PLS on the top features.
   - Transform X into the PLS component space and collapse components (mean across components).
   - Min‑max scale the final score to [0, 1] and return it as the per‑row weight Series.

Practical notes and recommended defaults

- Defaults in weight.py: max\_components=10; the function will cap components by available samples/features.
- Use select\_components(...) first to drop near‑constant or highly correlated components before PLS.
- Typical target: use a vector that reflects expected signal (log10Intensity or adjIntensity).
- If not enough samples/features the function returns a neutral weight (0.5) to avoid misleading extremes.
- Set verbose=True to print diagnostics (R² by component, chosen components, top features, final range).
- Computation: PLS fitting is fast for moderate sizes but can be heavier if many features and many samples — run on a subset if needed.

Caveats and interpretation

- PLS optimizes weights to predict your chosen target (intensity). Good predictive alignment does not guarantee better biological sensitivity to perturbations — validate on held‑out or simulated data.
- Because the method is supervised, avoid leaking labels into X or y (use raw intensity, not downstream p‑values).
- The returned weights emphasize components that cohere with intensity structure; combine with domain priors (e.g., keep W\_Impute high) if desired.

Quick usage recipe (notebook)

- Generate full set of W\_\* with weight.generate\_weights\_data(...)
- Keep informative columns via weight.select\_components(...)
- Call weight.generate\_auto\_pls\_weights(weights\_data, weights\_cols, y\_target=input\_data['log10Intensity'], verbose=True)
- Write the returned Series into input\_data['Weight'] and pass to model.LinearModel for WLS/GLM.

Alternative: generate\_refined\_top5\_mean\_weights(...)

- If you prefer a more conservative approach, there is an alternative helper that selects the top‑5 most important features then averages refined PLS components. It is faster and often robust when you expect only a few informative components.

```
# Initialize the variables
combined_results = []
model_types = ['wls', 'glm']

simID = "Sim2"
combined_results = []
miss_keys = ['Pro0_Pep0', 'Pro0.2_Pep0.2', 'Pro0.4_Pep0.4', 'Pro0.6_Pep0.6', 'Pro0.8_Pep0.8']
# experiment_file_name_template = f"2_{}_imputed_InputData.feather"
miss_mappers = {
    'Pro0_Pep0': '0% Missingness',
    'Pro0.2_Pep0.2': '20% Missingness',
    'Pro0.4_Pep0.4': '40% Missingness',
    'Pro0.6_Pep0.6': '60% Missingness',
    'Pro0.8_Pep0.8': '80% Missingness'
}

weight_keys = ['no_weight', 'default_mix', 'impute_only', 'revtech_only', 'PLS_auto']
weight_mappers = {
    'no_weight': 'No Weights',
    'default_mix': '90% Impute + 10% RevTech',
    'impute_only': '100% Impute',
    'revtech_only': '100% RevTech',
    'PLS_auto': 'Weights Auto-calculated from PLS'
}

def generate_weights_data(
    df: pd.DataFrame,
    sample_cols: list[str] = None,
    log_intensity_col: str = 'log10Intensity',
    adj_intensity_col: str = 'adjIntensity',
    control_condition: str = 'day1',
    condition_col: str = 'day',
    protein_col: str = 'protein_id',
    peptide_col: str = 'peptide_id',
    # Imputation Parameters
    is_real_col: str = None,
    is_comp_miss_col: str = None,
    true_val: float = 1.0,
    sparse_imputed_val: float = 1e-5,
    dense_imputed_val: float = 0.25,
    verbose: bool = False,
) -> pd.DataFrame:
    """
    Generate a DataFrame with all standard weight components calculated for given dataframe.

    Parameters
    ----------
    df : pd.DataFrame
        Input DataFrame containing all required columns.
    sample_cols : list[str], optional
        Sample identifier columns, by default ['filename']
    log_intensity_col : str, optional
        Name of the log intensity column, by default 'log10Intensity'
    adj_intensity_col : str, optional
        Name of the adjusted intensity column, by default 'adjIntensity'
    control_condition : str, optional
        Name of the control condition, by default 'day1'
    condition_col : str, optional
        Name of the condition column, by default 'day'
    protein_col : str, optional
        Name of the protein column, by default 'protein_id'
    peptide_col : str, optional
        Name of the peptide column, by default 'peptide_id'
    verbose : bool, optional
        Whether to print progress and timing, by default True
    is_real_col : str, optional
        Column name indicating if the value is real (not imputed).
    is_comp_miss_col : str, optional
        Column name indicating if the peptide is completely missing for a condition.
    true_val : float, optional
        Weight for real (not imputed) values.
    sparse_imputed_val : float, optional
        Weight for sparsely imputed values.
    dense_imputed_val : float, optional
        Weight for densely imputed values.

    Returns
    -------
    pd.DataFrame
        DataFrame with all weight columns added (prefixed with 'W_').
    """
    group_cols = [protein_col, peptide_col, condition_col]
    if sample_cols is None:
        sample_cols = ['filename']
    intensity_cols = [log_intensity_col, adj_intensity_col]

    weight_names_with_intensity = [
        ('InverseMean', 'mean'),
        ('InverseVar', 'var'),
        ('InverseStd', 'std'),
        ('CorrDiscordance', None),
        ('RelativeVariability', None),
        ('SignalToNoise', None),
        ('ReplicateConcordance', None),
        ('DirectionalAgreement', None),
    ]

    weights_data = df[group_cols + sample_cols + intensity_cols].copy()
    # Add is_real and is_comp_miss columns if provided
    if is_real_col:
        if is_real_col not in df.columns:
            raise ValueError(f"Column '{is_real_col}' not found in the DataFrame.")
        weights_data[is_real_col] = df[is_real_col].astype(bool)
    if is_comp_miss_col:
        if is_comp_miss_col not in df.columns:
            raise ValueError(f"Column '{is_comp_miss_col}' not found in the DataFrame.")
        weights_data[is_comp_miss_col] = df[is_comp_miss_col].astype(bool)

    # Imputation Weights
    if verbose:
        print(" 📏 Calculating Imputation Weights...", end="")
    weights_data['W_Impute'] = weight.calculate_imputation_weights(
        weights_data,
        is_real_col=is_real_col,
        is_comp_miss_col=is_comp_miss_col,
        true_val=true_val,
        sparse_imputed_val=sparse_imputed_val,
        dense_imputed_val=dense_imputed_val,
        verbose=verbose
    )
    # # Replicate Imbalance Weights
    weights_data['W_RepImbalance'] = weight.calculate_replicate_imbalance_weights(
        weights_data,
        condition_col=condition_col,
        protein_col=protein_col,
        peptide_col=peptide_col,
        is_real_col=is_real_col,
        verbose=verbose
    )


    # # Normalization Impact Weights
    weights_data['W_NormalizationImpact'] = weight.calculate_normalization_impact_weights(
        weights_data,
        log_intensity_col=log_intensity_col,
        adj_intensity_col=adj_intensity_col,
        protein_col=protein_col,
        peptide_col=peptide_col,
        verbose=verbose
    )

    # Reverse Technical Variation Weights
    weights_data['W_RevTechVar'] = weights_data.groupby(
        [protein_col, peptide_col, condition_col], observed=True
    )[adj_intensity_col].transform('var').fillna(0.0)
    weights_data['W_RevTechVar'] = 1 - min_max_scale(weights_data['W_RevTechVar'])

    # Profile Correlation Weights
    weights_data['W_ProfileCorr'] = weight.calculate_profile_correlation_weights(
        weights_data,
        log_intensity_col=log_intensity_col,
        adj_intensity_col=adj_intensity_col,
        condition_col=condition_col,
        protein_col=protein_col,
        peptide_col=peptide_col, 
        verbose=verbose
    )

    # Weights with intensity
    for wName, metric in weight_names_with_intensity:
        for icol, suffix in zip([log_intensity_col, adj_intensity_col], ['(log)', '(adj)']):
            colname = f'W_{wName}{suffix}'
            if wName == 'InverseMean':
                if verbose:
                    print(f" 📏 Calculating {wName} Weights ({suffix})...", end="")
                weights_data[colname] = weight.calculate_inverse_metric_weights(
                    weights_data,
                    metric='mean',
                    intensity_col=icol,
                    group_cols=group_cols
                )
            elif wName == 'InverseVar':
                if verbose:
                    print(f" 📏 Calculating {wName} Weights ({suffix})...", end="")
                weights_data[colname] = weight.calculate_inverse_metric_weights(
                    weights_data,
                    metric='var',
                    intensity_col=icol,
                    group_cols=group_cols
                )
            elif wName == 'InverseStd':
                if verbose:
                    print(f" 📏 Calculating {wName} Weights ({suffix})...", end="")
                weights_data[colname] = weight.calculate_inverse_metric_weights(
                    weights_data,
                    metric='std',
                    intensity_col=icol,
                    group_cols=group_cols
                )
            elif wName == 'CorrDiscordance':
                if verbose:
                    print(f" 📏 Calculating {wName} Weights ({suffix})...", end="")
                weights_data[colname] = weight.calculate_correlation_discordance_weights(
                    weights_data,
                    intensity_col=icol,
                    control_condition=control_condition,
                    condition_col=condition_col,
                    protein_col=protein_col,
                    peptide_col=peptide_col, 
                    verbose=verbose
                )
            elif wName == 'RelativeVariability':
                if verbose:
                    print(f" 📏 Calculating {wName} Weights ({suffix})...", end="")
                weights_data[colname] = weight.calculate_relative_variability_weights(
                    weights_data,
                    intensity_col=icol,
                    condition_col=condition_col,
                    protein_col=protein_col,
                    peptide_col=peptide_col, 
                    verbose=verbose
                )
            elif wName == 'SignalToNoise':
                if verbose:
                    print(f" 📏 Calculating {wName} Weights ({suffix})...", end="")
                weights_data[colname] = weight.calculate_generalized_signal_to_noise_weights(
                    weights_data,
                    intensity_col=icol,
                    condition_col=condition_col,
                    protein_col=protein_col,
                    peptide_col=peptide_col, 
                    verbose=verbose
                )
            elif wName == 'ReplicateConcordance':
                if verbose:
                    print(f" 📏 Calculating {wName} Weights ({suffix})...", end="")
                weights_data[colname] = weight.calculate_replicate_concordance_weights(
                    weights_data,
                    intensity_col=icol,
                    condition_col=condition_col,
                    protein_col=protein_col,
                    peptide_col=peptide_col, 
                    verbose=verbose
                )

    # # Normalize all W_ columns
    # weights_cols = [col for col in weights_data.columns if col.startswith('W_')]
    # weights_data[weights_cols] = weights_data[weights_cols].apply(min_max_scale, axis=0)
    if verbose:
        print("All weights calculated and normalized.")
    return weights_data
```

###### 2.4.5 Execute ProteoForge Analysis with Different Models¶

The below is the logic that opens all the datasets to be analyzed and applies ProteoForge's analysis and benchmarking results with different linear model variants.

> This part can be commented out if the results are already computed and saved in the `data/` folder. Unless any changes required/needed re-calculation when restarting the kernel is unnecessary.

```
for exp in weight_keys:
    if exp == 'default_mix':
        continue  # Skip default_mix as it's already included in previous analysis
    for miss_type in miss_keys:
        file_path = f"./data/{simID}/2_{miss_type}_imputed_InputData.feather"
        input_data = pd.read_feather(file_path)
        weights_data = generate_weights_data(
            input_data,
            sample_cols=['Sample'],
            log_intensity_col='log10Intensity',
            adj_intensity_col='adjIntensity',
            control_condition='control',
            condition_col='Condition',  
            protein_col='Protein',
            peptide_col='Peptide',
            is_real_col='isReal',
            is_comp_miss_col='isCompMiss',
            sparse_imputed_val=1e-10,
            dense_imputed_val=0.5,
            verbose=False,
        )
        weights_cols = [col for col in weights_data.columns if col.startswith('W_')]
        weight_cols_to_keep = weight.select_components(
            weights_data[weights_cols], 
            std_threshold=0.05, 
            corr_threshold=0.8,
            verbose=False
        )
        # Calculate weights based on the experiment
        if exp == 'no_weight':
            input_data['Weight'] = 1.0
        elif exp == 'impute_only':
            input_data['Weight'] = weights_data['W_Impute']
        elif exp == 'revtech_only':
            input_data['Weight'] = weights_data['W_InverseVar(log)']
        elif exp == 'PLS_top5':
            auto_weights = weight.generate_refined_top5_mean_weights(
                weights_data=weights_data,
                weights_cols=weight_cols_to_keep,
                y_target=input_data['log10Intensity'],
                verbose=False
            )
            input_data['Weight'] = auto_weights
        elif exp == 'PLS_auto':
            auto_weights = weight.generate_auto_pls_weights(
                weights_data=weights_data,
                weights_cols=weight_cols_to_keep,
                y_target=input_data['log10Intensity'],
                max_components=10,
                verbose=False
            )
            input_data['Weight'] = auto_weights

        for modelType in model_types:
            test_data = input_data.copy()
            model_to_use = 'quantile' if modelType == 'mqr' else modelType
            
            cur_model = model.LinearModel(
                data=test_data,
                protein_col="Protein",
                peptide_col="Peptide",
                cond_col="Condition",
                intensity_col="adjIntensity",
                weight_col='Weight',
            )

            test_data = cur_model.run_analysis(
                model_type=model_to_use,
                correction_strategy=correction['strategy'],
                correction_methods=correction['methods'],
                n_jobs=n_jobs
            )

            clusters = disluster.distance_and_cluster(
                data=test_data,
                protein_col='Protein',
                peptide_col='PeptideID',
                cond_col='Condition',
                quant_col='adjIntensity',
                clustering_params={
                    'min_clusters': 1,
                    'distance_transform': 'corr',
                    'clustering_method': 'hybrid_outlier_cut',
                    'linkage_method': 'ward',
                    'distance_metric': 'euclidean'
                },
                n_jobs=n_jobs,
                verbose=False
            )

            test_data = test_data.merge(
                clusters[['Protein', 'PeptideID', 'cluster_label']],
                on=['Protein', 'PeptideID'],
                how='left'
            ).rename(columns={'cluster_label': 'ClusterID'})

            test_data['pertPeptide'] = np.where(
                test_data['isCompMiss'] == 1, 1, test_data['pertPeptide']
            ).astype(bool)
            test_data['Reason'] = np.where(
                test_data['isCompMiss'] == 1, "Biological Absence", test_data['Reason']
            )
            test_data.loc[
                (test_data['isCompMiss']==1) & (test_data['pertPeptide']), 
                'pertPFG'
            ] = 1

            # Identification Benchmark
            metric_data = utils.create_metric_data(
                test_data, 
                pvalue_thresholds=thresholds,
                label_col='pertPeptide',
                pvalue_col='adj_pval'
            )
            metric_data['Model'] = modelType
            metric_data['Experiment'] = weight_mappers[exp]
            metric_data['MissLevel'] = miss_mappers[miss_type]
            metric_data['Benchmark'] = 'PepID'
            combined_results.append(metric_data)

            # Peptide Grouping Benchmark
            # Prepare data for grouping performance
            tmp = test_data[[
                'Protein', 'Peptide','PeptideID', 
                'pertPeptide', 'pertPFG', 'ClusterID', 'adj_pval'
            ]].drop_duplicates().reset_index(drop=True)
            metric_data = utils.grouping_performance_proteoforge(
                data=tmp,
                thresholds=thresholds,
                pvalue_col='adj_pval',
                protein_col='Protein',
                cluster_col='ClusterID',
                perturbation_col='pertPFG',
            )
            metric_data['Model'] = modelType
            metric_data['Experiment'] = weight_mappers[exp]
            metric_data['MissLevel'] = miss_mappers[miss_type]
            metric_data['Benchmark'] = 'PepGroup'
            combined_results.append(metric_data)

# Combine all results into a single DataFrame
all_results = pd.concat(combined_results, ignore_index=True)
# Adding the Default Mix Model
default_mix_file_path = f"./data/Sim2_ModelBenchmark_Results_staticWeight.feather"
default_mix_data = pd.read_feather(default_mix_file_path)
default_mix_data.rename(columns={'Experiment':'MissLevel'}, inplace=True)
default_mix_data['Experiment'] = '90% Impute + 10% RevTech'
# Adding this data to the combined results
all_results = pd.concat([all_results, default_mix_data], ignore_index=True)
all_results.to_feather(f"./data/Sim1_ModelBenchmark_Results_alternativeWeights.feather")
```

```
all_results = pd.read_feather(f"./data/Sim2_ModelBenchmark_Results_alternativeWeights.feather")
mask = all_results['Model'].isin(['mqr', 'rlm'])
all_results.loc[mask, 'Experiment'] = 'No Weights'
# Remove the DataType Column if present
if 'DataType' in all_results.columns:
    all_results = all_results.drop(columns=['DataType'])
id_data = all_results[all_results['Benchmark'] == 'PepID'].copy()
pg_data = all_results[all_results['Benchmark'] == 'PepGroup'].copy()
df = all_results.copy()
```

###### 2.4.6 Mean MCC Difference from No Weight Baseline against Different Weighting Strategies¶

This setup essentially calculates the mean MCC difference from no weight baseline for each weighting strategy used in WLS and GLM models, to see how much each weighting strategy improves or worsens the model performance in comparison to no weights at all, and the results are shown for various missingness levels, going from 0% to 80% missingness.

```
df['MissLevel_Int'] = df['MissLevel'].str.extract(r'(\d+)').astype(int)

# 2. Aggregation: Get Max MCC per condition (Best Threshold)
agg = df.groupby(['Benchmark', 'MissLevel_Int', 'Model', 'Experiment'])['MCC'].mean().reset_index()

# 3. Calculate Deltas (vs '90% Impute + 10% RevTech' baseline)
# Isolate the baselines (set default-mix as baseline)
baseline_label = 'No Weights'
baselines = agg[agg['Experiment'] == baseline_label][['Benchmark', 'MissLevel_Int', 'Model', 'MCC']]
baselines = baselines.rename(columns={'MCC': 'Baseline_MCC'})

# Merge and Subtract (Delta relative to the selected baseline)
plot_df = agg.merge(baselines.rename(columns={'MCC': 'Baseline_MCC'}), on=['Benchmark', 'MissLevel_Int', 'Model'], how='left')
plot_df['Delta_MCC'] = plot_df['MCC'] - plot_df['Baseline_MCC']

# Filter: Comparison only makes sense for models that HAVE weights (WLS, GLM)
plot_df = plot_df[plot_df['Model'].isin(['wls', 'glm'])]
# NOTE: We keep all experiments in the plot so the 'No Weights' strategy remains visible

# 4. Ordering
plot_df['Benchmark'] = pd.Categorical(plot_df['Benchmark'], categories=['PepID', 'PepGroup'], ordered=True)

from matplotlib.ticker import MaxNLocator

# Configuration
models_order = ['wls', 'glm']                      # rows
benchmarks_order = ['PepID', 'PepGroup']           # cols
exp_order = [
    'No Weights',
    '100% Impute',
    '100% RevTech',
    '90% Impute + 10% RevTech',
    'Weights Auto-calculated from PLS'
]
colors = {
    'No Weights': '#7f7f7f',
    '100% Impute': '#1f77b4',
    '100% RevTech': '#ff7f0e',
    'Weights Auto-calculated from PLS': '#d62728',
    '90% Impute + 10% RevTech': '#2ca02c'
}
markers = {
    'No Weights': 'o',
    '100% Impute': 's',
    '100% RevTech': 'D',
    '90% Impute + 10% RevTech': '^',
    'Weights Auto-calculated from PLS': 'P'
}

# Prepare grid
nrows = len(models_order)
ncols = len(benchmarks_order)
fig, axes = plt.subplots(
    nrows=nrows, ncols=ncols,
    figsize=(14, 6.5),
    sharex=True, sharey=True,
    constrained_layout=True
)

# If axes is 1D for any dimension, make it a 2D array
if nrows == 1 and ncols == 1:
    axes = np.array([[axes]])
elif nrows == 1:
    axes = axes[np.newaxis, :]
elif ncols == 1:
    axes = axes[:, np.newaxis]

# X tick positions / labels
x_positions = sorted(plot_df['MissLevel_Int'].unique())
x_labels = [f"{int(x)}%" for x in x_positions]

for i, model in enumerate(models_order):
    for j, bench in enumerate(benchmarks_order):
        ax = axes[i, j]
        ax.set_facecolor("#fbfbfb")
        ax.grid(axis='both', linestyle='--', linewidth=0.6, alpha=0.45, zorder=0)
        ax.spines['top'].set_visible(False)
        ax.spines['right'].set_visible(False)
        ax.spines['left'].set_visible(True)
        ax.spines['bottom'].set_visible(True)

        # Subset for this facet
        facet = plot_df[(plot_df['Model'] == model) & (plot_df['Benchmark'] == bench)].copy()
        if facet.empty:
            ax.text(0.5, 0.5, "No data", ha='center', va='center', fontsize=10, color='gray')
            ax.set_xticks(x_positions)
            ax.set_xticklabels(x_labels, rotation=0)
            continue

        # Plot each experiment line
        for exp in exp_order:
            sub = facet[facet['Experiment'] == exp].copy()
            if sub.empty:
                continue
            sub = sub.sort_values('MissLevel_Int')
            ax.plot(
                sub['MissLevel_Int'],
                sub['Delta_MCC'],
                label=exp,
                color=colors.get(exp, '#333333'),
                marker=markers.get(exp, 'o'),
                markersize=7,
                linewidth=2.2,
                linestyle='-',
                markeredgewidth=0.9,
                alpha=0.95,
                zorder=5
            )

            # Highlight best improvement point for this (model,bench,exp)
            best_idx = sub['Delta_MCC'].idxmax()
            if not np.isnan(best_idx):
                best_row = sub.loc[best_idx]
                ax.scatter(
                    best_row['MissLevel_Int'],
                    best_row['Delta_MCC'],
                    s=110,
                    facecolors='none',
                    edgecolors=colors.get(exp, '#333333'),
                    linewidths=1.25,
                    zorder=10
                )

        # Baseline reference (Delta = 0)
        ax.axhline(0.0, color='black', linestyle='--', linewidth=1.0, alpha=0.7, zorder=1)
        # Axis formatting
        ax.set_xticks(x_positions)
        ax.set_xticklabels(x_labels, rotation=0)
        ax.xaxis.set_major_locator(MaxNLocator(integer=True))
        ax.set_xlim(min(x_positions) - 2, max(x_positions) + 2)
        ax.set_ylim(plot_df['Delta_MCC'].min() - 0.05, plot_df['Delta_MCC'].max() + 0.05)

        # Titles / labels
        if i == 0:
            ax.set_title(bench, fontsize=12, fontweight='bold')

        # Add small label on right side indicating Benchmark row name (only last column)
        if j == ncols - 1:
            ax.text(1.02, 0.5, model.upper(), transform=ax.transAxes, rotation=270, va='center', fontsize=12, fontweight='bold')

# Global legend (unique handles)
handles = []
labels = []
for exp in exp_order:
    handles.append(plt.Line2D([0], [0], color=colors.get(exp), marker=markers.get(exp), linestyle='-', markersize=7, linewidth=2.2))
    labels.append(exp)

# Add legend below the plots
fig.legend(handles, labels, loc='lower center', bbox_to_anchor=(0.5, -0.20), ncol=3, frameon=False, fontsize=10, title="Weighting Strategy")

# Global labels & title
fig.suptitle("Impact of Weighting Strategies — MCC Improvement vs Unweighted Baseline", fontsize=16, fontweight='bold', y=1.04)
fig.text(0.5, -0.05, "Missingness (%)", ha='center', fontsize=12, fontweight='bold')
fig.text(-0.05, 0.5, "MCC Improvement (Delta vs No Weights)", va='center', rotation='vertical', fontsize=12, fontweight='bold')

plots.finalize_plot(
    plt.gcf(), 
    show=True, save=save_to_folder,
    filename='modelBenchmarks_missingness_MCCDeltas_exp3',
    filepath=figure_path,
    formats=figure_formats, 
    transparent=transparent_bg,
    dpi=figure_dpi,
)
```

This is a very interesting case of how the bencmarking at identification and grouping levels shows drastic differences:

- The identification level benchmark shows using only imputations weights proves the highest then the mixed (90% imputation + 10% revtechvar) weights, especially when missingness increases 40% and onwards.
- Auto-calculated PLS based weights seems to perform on par, however with lower missingness it actually performs worse than no weights at all, which is an interesting observation.
- RevTech is by far the worst across all missingness levels and comparisons, as alone it doesn't provide any meaningful information about the imputation quality, which is the most important factor when dealing with high missingness levels.
- However at grouping level benchmarks, the mixed weights (90% imputation + 10% revtechvar) seems to perform the best across all missingness levels, followed by auto-calculated PLS based weights.
- At the highest missingness Auto-calculated PLS based weights seems to perform quite well at grouping levels, which shows that it is able to capture other meaningful information, that can compensate for the imputation quality, as well as other weight components that are useful for grouping level benchmarks.
- Imputation only drastically reduces performance at grouping levels, showing that while it is important to have good imputation quality weights, other weight components are also necessary to capture the full picture to generaete meaningful and correct dPFs.

> The difference that is observed from identification to grouping benchmarking results is due to grouping uses the peptide identificaiton levels (aka, significantly discordant peptides) to build dPFs from the clusterin methods. The cluster part of the ProteoForge is not the most robust method, which makes the generally grouping more sensitive to small imperfections. They are also in a way the summarized, where only one peptide needs to be identified as significant to make the whole dPF significant, which can be quite sensitive to small changes in the peptide identification levels.

I think I will use this since it gives a good overview of how much each weighting strategy improves or worsens the model performance in comparison to no weights at all.

---

###### 2.4.7 High Level Summary Figure with Mean MCC¶

This last visualization is the high-level summary figure that shows the mean MCC for each weighting strategy used in WLS and GLM models, as well as the RLM and MQR, to see how much each weighting strategy improves or worsens the model performance in comparison to no weights at all, and the results are shown for various missingness levels, going from 0% to 80% missingness.

It is designed to show the mean MCC directly, and uses hue (color) grouping for the 'No Weights', 'Mixed Strategy (90% Imp)'Auto-PLS Weights'. Highlights the top performing at each missingness level and benchmarking type with a larger marker size. Additionally sets RLM's mean MCC as the reference to ease of comparison.

```
df['MissLevel_Int'] = df['MissLevel'].str.extract(r'(\d+)').astype(int)

# Aggregate (Max MCC per condition)
agg = df.groupby(['Benchmark', 'MissLevel_Int', 'Model', 'Experiment'])['MCC'].max().reset_index()

# Filter Data
experiments_show = ['No Weights', '90% Impute + 10% RevTech', 'Weights Auto-calculated from PLS']
models_show = ['mqr', 'rlm', 'wls', 'glm']
plot_df = agg[agg['Experiment'].isin(experiments_show) & agg['Model'].isin(models_show)].copy()

# Enforce Ordering
plot_df['Model'] = pd.Categorical(plot_df['Model'], categories=models_show, ordered=True)
plot_df['Experiment'] = pd.Categorical(plot_df['Experiment'], categories=experiments_show, ordered=True)

# 2. Setup Grid
benchmarks = ['PepID', 'PepGroup']
miss_levels = sorted(plot_df['MissLevel_Int'].unique())
fig, axes = plt.subplots(
    len(benchmarks), len(miss_levels), 
    figsize=(16, 7), 
    sharey=True, sharex=True,
    constrained_layout=True
)

# Colors
colors = {
    'No Weights': '#bababa',                       # Neutral Grey
    '90% Impute + 10% RevTech': '#1f77b4',         # Blue (Explicit Strategy)
    'Weights Auto-calculated from PLS': '#d62728'  # Red (PLS Strategy)
}

# 3. Plotting Loop
for i, bench in enumerate(benchmarks):
    for j, miss in enumerate(miss_levels):
        ax = axes[i, j]
        
        # Subset
        subset = plot_df[(plot_df['Benchmark'] == bench) & (plot_df['MissLevel_Int'] == miss)]
        
        # Draw Bars
        sns.barplot(
            data=subset, 
            x='Model', y='MCC', hue='Experiment',
            palette=colors,
            hue_order=experiments_show,
            order=models_show,
            ax=ax,
            edgecolor='#333333', linewidth=0.5,
            errorbar=None,
            alpha=0.9
        )
        
        # --- Annotations ---
        
        # A. Highlight the Winner (Star)
        if not subset.empty:
            best_idx = subset['MCC'].idxmax()
            best_row = subset.loc[best_idx]
            best_val = best_row['MCC']
            
            # Calculate x position for the star
            n_hues = len(experiments_show)
            hue_width = 0.8 / n_hues 
            hue_idx = experiments_show.index(best_row['Experiment'])
            model_idx = models_show.index(best_row['Model'])
            x_pos = (model_idx - 0.4) + (hue_width * hue_idx) + (hue_width / 2)
            
            ax.text(x_pos, best_val - 0.05, '*', ha='center', va='bottom', 
                    fontsize=25, color='#ffd700', fontweight='bold', zorder=10) # Gold Star

        # B. RLM Baseline Line
        rlm_row = subset[(subset['Model'] == 'rlm') & (subset['Experiment'] == 'No Weights')]
        rlm_val = 0
        if not rlm_row.empty:
            rlm_val = rlm_row['MCC'].values[0]
            ax.axhline(rlm_val, color='#333333', linestyle='--', linewidth=1.5, alpha=0.7, zorder=5)
            if j == 0:
                ax.text(-0.45, rlm_val + 0.02, "RLM Baseline", color='#333333', fontsize=8, fontweight='bold')

        # C. Annotate Delta for WLS (Auto-PLS) vs RLM
        wls_pls = subset[(subset['Model'] == 'wls') & (subset['Experiment'] == 'Weights Auto-calculated from PLS')]
        if not wls_pls.empty and not rlm_row.empty:
            wls_val = wls_pls['MCC'].values[0]
            delta = wls_val - rlm_val
            
            # WLS is index 2. PLS is index 2.
            x_pos_wls = (2 - 0.4) + (hue_width * 2) + (hue_width / 2)
            
            color_txt = '#d62728' if delta > -0.01 else 'gray'
            fw = 'bold' if delta > 0 else 'normal'
            txt = f"{delta:+.2f}"
            
            # Show delta if it's high missingness or significant difference
            if miss >= 40:
                ax.text(x_pos_wls, wls_val + 0.1, txt, ha='center', va='bottom', 
                        fontsize=8, color=color_txt, fontweight=fw)

        # Styling
        ax.set_ylim(0, 1.2)
        ax.grid(axis='y', linestyle='--', alpha=0.3)
        ax.set_xlabel('')
        ax.set_ylabel('')
        ax.get_legend().remove()
        
        if i == 0:
            ax.set_title(f"{miss}% Missing", fontsize=12, fontweight='bold')
        if j == len(miss_levels) - 1:
            ax.text(1.05, 0.5, bench, transform=ax.transAxes, rotation=270, va='center', fontsize=14, fontweight='bold')

# Global Labels & Legend
fig.text(0.5, -0.02, 'Model', ha='center', fontsize=13, fontweight='bold')
fig.text(-0.01, 0.5, 'Mean MCC', va='center', rotation='vertical', fontsize=13, fontweight='bold')

handles = [
    Line2D([0], [0], color=colors['No Weights'], lw=6, label='No Weights (Baseline)'),
    Line2D([0], [0], color=colors['90% Impute + 10% RevTech'], lw=6, label='Mixed Strategy (90% Imp)'),
    Line2D([0], [0], color=colors['Weights Auto-calculated from PLS'], lw=6, label='Auto-PLS Weights'),
    Line2D([0], [0], color='#333333', linestyle='--', lw=2, label='RLM Baseline'),
    Line2D([0], [0], marker='*', color='w', markerfacecolor='#ffd700', markersize=14, label='Best in Group')
]

fig.legend(handles=handles, loc='upper center', bbox_to_anchor=(0.5, 1.08), ncol=5, frameon=False, fontsize=10)
plots.finalize_plot(
    plt.gcf(), 
    show=True, save=save_to_folder,
    filename='modelBenchmarks_missingness_MCC_exp3',
    filepath=figure_path,
    formats=figure_formats, 
    transparent=transparent_bg,
    dpi=figure_dpi,
)
```

This is alternative way to show the RLM and MQRs performance drops with higher missingness. However It also shows how customized weighting strategies passed with WLS and GLM models do shine through, especially at higher missingness levels, where they are able to outperform RLM and MQR, given that the weights provided do reflect the imputation quality well enough.

Auto-PLS weights performs either better or comparable to imputation heavy weights in grouping level benchmarks where as the identification generally favours heavier emphasis on imputation quality weights.

---

#### 3. Summary and conclusions¶

##### 3.1 Main findings¶

- RLM is the most robust model overall.
  - Stable for low → moderate missingness.
  - Performance drops start ≈ 50% missingness and become severe by 60–80%.
- MQR is extremely niche
  - Only excels at identification of single or multiple perturbed peptides with complete data.
  - Suffers greatly from imputation artifacts; avoid if imputation is used.
- Weighted methods (WLS, GLM)
  - Outperform robust methods at high missingness when weights emphasize imputation quality.
  - Weight composition matters: imputation‑heavy mixes (e.g., 90% W\_Impute + 10% W\_RevTechVar) improve PepID at high missingness; mixed/Auto‑PLS weights favor PepGroup.
- Auto‑PLS
  - Competitive for grouping metrics.
  - May underperform vs. no weights at low missingness (overfitting to intensity structure rather than perturbations).

##### 3.2 Actionable recommendations¶

- Low-Mid missingness (< 20%): use RLM as a safe default.
- Moderate missingness (20–50%): prefer RLM or WLS/GLM with moderate weighting (∼50–75% W\_Impute). Validate subgroups.
- High missingness (≥ 60%): prefer WLS/GLM with imputation‑heavy weights (0.8–0.95 W\_Impute) or Auto‑PLS; validate top hits manually and on held‑out simulations.
- Peptide grouping: always include at least one mixed strategy or Auto‑PLS weight — grouping is sensitive to single‑peptide calls.

---

##### 3.3 Reproducible default settings used here¶

- Imputation weights: true\_val = 1.0; dense\_imputed\_val = 0.5 (sometimes 0.25); sparse\_imputed\_val = 1e‑5.
- Default mix: 0.90  *W\_Impute + 0.10*  W\_RevTechVar.
- PLS auto: max\_components = 10 (capped by n\_features and n\_samples‑1); run select\_components(...) before PLS.

---

##### 3.4 Limitations & caveats¶

- Clustering (ProteoForge disluster) strongly affects PepGroup — small differences in peptide calls can amplify grouping metrics.
- PLS is supervised on intensity: do NOT leak labels (p‑values, perturbation flags) into X or y when generating Auto‑PLS weights.
- Weight components depend on the imputation method used in Sim1/Sim2; conclusions are conditional on that choice.

```
print("Notebook Execution Time:", utils.prettyTimer(utils.getTime() - startTime))
```

```
Notebook Execution Time: 00h:12m:55s
```
