## Supplementary material for "ProteoForge: An Imputation-Aware Framework for Differential Proteoform Discovery in Bottom-Up Proteomics": Analysis notebooks as html render: Notebook S8.html

### Notebook S8

###### Enes Kemal Ergin

#### 2025-12-08 23:00:34

### Supplementary Notebook 8: Highly Summarized Plots for Assembling the Figure 2¶

- **License:** Creative Commons Attribution-NonCommercial 4.0 International License
- **Version:** 0.2
- **Edit Log:**
  - 2025-11-28: Initial version of the notebook
  - 2025-12-08: Revise the whole notebook, ensuring clarity and correctness

---

**Requirements:**

The following preprocessing and method runs must be completed (in order) before executing this notebook:

1. `01-SimulatedDatasets.ipynb` - Prepares simulated datasets with various perturbation scenarios.
2. `02-runCOPF.R` - Executes the COPF method on the simulated datasets.
3. `03-runPeCorA.R` - Executes the PeCorA method on the simulated datasets.
4. `04-runProteoForge.R` - Executes the ProteoForge method on the simulated datasets.
5. `05-IdentificationBenchmark.ipynb` - Generates the peptide identification benchmark figures, and combines results from all methods.
6. `06-GroupingBenchmark.ipynb` - Generates the peptide grouping benchmark figures, and combines results from all methods.

---

**Data Information:**

This notebooks uses the final combined benchmarking results table from the real-data benchmarking (../Benchmark/) and simulated benchmarking (./) analyses. From 1 real-data, and 4 simulated setups, 2 benchmarks (peptide identification and peptide grouping) were used.

---

**Purpose:**

This notebook compiles key performance plots from previous analyses into a cohesive figure for publication. It focuses on summarizing the ROC curves and optimal F1/MCC thresholds across different perturbation scenarios and methods (COPF, PeCorA, ProteoForge). The final figure highlights comparative strengths and weaknesses of each approach in detecting proteoform changes under various conditions. This figure is meant to be very high-level and concise to go into the main figures, whereas the detailed plots and analyses are available in the supplementary notebooks as well as placed into supplementary figures.

---

#### Setup¶

This section imports required libraries, configures display settings, and defines paths for data and figures.

> **Note:** The HTML rendering of this notebook hides code cells by default. Click the "Code" buttons on the right to expand them.

##### Libraries¶

```
import os
import sys
import warnings

import numpy as np # Numerical computing
import pandas as pd # Data manipulatio

import seaborn as sns # R-like high-level plots
import matplotlib.pyplot as plt # Python's base plotting 

sys.path.append('../')
from src import utils, plots

warnings.filterwarnings('ignore')

# Initialize the timer
startTime = utils.getTime()
```

##### Display Settings¶

The cell below configures pandas, matplotlib, and seaborn display options for improved readability of tables and figures, including color palettes and figure export settings.

```
# Define default colors and styles for plots
def_colors = [
    "#139593", "#fca311", "#e54f2a",
    "#c3c3c3", "#555555",
    "#690000", "#5f4a00", "#004549"
]

# Set seaborn style
sns.set_theme(
    style="white",
    context="paper",
    palette=def_colors,
    font_scale=1,
    rc={
        "figure.figsize": (6, 4),
        "font.family": "sans-serif",
        "font.sans-serif": ["Arial", "Ubuntu Mono"],
    }
)

# Figure Saving Settings
figure_formats = ["pdf"]
save_to_folder = True
transparent_bg = True
figure_dpi = 300

## Configure dataframe displaying
pd.set_option('display.float_format', lambda x: '%.4f' % x)
pd.set_option('display.max_rows', 50)
pd.set_option('display.max_columns', 50)
pd.set_option('display.width', 500)  # Set a wider display width

## Printing Settings
verbose = True
```

##### Data and Result Paths¶

Data and figures are organized in separate folders:

- `data_path` — Not one directory is needed as data is read from both the real-data benchmarking (`../Benchmark/data/prepared/`) and simulated benchmarking (`./data/prepared/`) analyses. They are explicitly added to each file-reading step.
- `figure_path` — Directory for generated plots and figures (`figures/08-FigureAssembly/`).

```
notebook_name = "08-FigureAssembly"
figure_path = f"./figures/{notebook_name}/"

# Create figure folder structure, if needed
if save_to_folder:
    for i in figure_formats:
        cur_folder = figure_path + i + "/"
        if not os.path.exists(cur_folder):
            os.makedirs(cur_folder)
```

##### Global Variables¶

Defines constants and visual styling for consistent analysis and publication-quality figures:

- **`seed`** — Random seed for reproducibility
- **`pthr`** — P-value threshold (10⁻³) for statistical significance
- **`method_palette`** / **`method_markers`** — Color and marker scheme for methods (COPF, PeCorA, ProteoForge)
- **`mcc_thresholds`** / **`mcc_colors`** — MCC interpretation scale with semantic labels (Random → Almost Perfect)

```
# Global variables
seed = 42 # Seed for reproducibility  
pthr = 10**-3  # p-value threshold for significance
thresholds = list(utils.generate_thresholds(10.0, -15, 1, 0, 1, 0.1)) # Thresholds for the analysis

# Methods and their plotting styles
method_palette = {
    "COPF": "#139593",
    "PeCorA": "#fca311",
    "ProteoForge": "#e54f2a",
}
method_styles = {
    "COPF": "--",
    "PeCorA": "-.",
    "ProteoForge": ":",
}
method_markers = {
    "COPF": "o",
    "PeCorA": "s",
    "ProteoForge": "^",
}


# Matthews Correlation Coefficient (MCC) thresholds and colors
mcc_thresholds = {
    0.0 : 'Random',
    0.3 : 'Fair (0.3)',
    0.5 : 'Moderate (0.5)',
    0.7 : 'Strong (0.7)',
    0.9 : 'Almost Perfect (0.9)',
}
mcc_colors = {
    'Random': '#c3c3c3',             
    'Fair (0.3)': '#e0aaff',         
    'Moderate (0.5)': '#9d4edd',     
    'Strong (0.7)': '#5a189a',       
    'Almost Perfect (0.9)': '#240046' 
}

plots.color_palette(mcc_colors, save=False, name="MCC Interpretation Colors")
plots.color_palette(method_palette, save=False, name="Method Colors")
```


---

#### Figure 2: Method Benchmarking Overview¶

This figure compiles performance comparisons of ProteoForge against established methods (COPF, PeCorA) using both real-world SWATH-MS data and controlled simulations. The benchmarks evaluate two core tasks: **Peptide Identification** (detecting which peptides are perturbed) and **Peptide Grouping** (correctly assigning perturbed peptides to proteoform groups).

##### 2.1 SWATH-MS Benchmark: Real-World Performance¶

**Data Source:** COPF manuscript benchmark dataset (analyzed in `../Benchmark/` notebooks)

**Key Question:** How do methods compare on experimentally validated proteoform perturbations?

```
swath_pepid = pd.read_feather("../Benchmark/data/results/peptide_identification_performance_data.feather")
swath_pepgrp = pd.read_feather("../Benchmark/data/results/peptide_grouping_performance_data.feather")
# Combine as swath with Benchmark column indicating pepID, or pepGrp
swath_pepgrp['Benchmark'] = 'Peptide Grouping'
swath_pepid['Benchmark'] = 'Peptide Identification'
swath_combined = pd.concat(
    [swath_pepid, swath_pepgrp], ignore_index=True
).rename(columns={
    'perturbation': 'Experiment',
    'method': 'Method',
})
# swath_combined['Experiment'].value_counts()
# Replace the Experiment to standardize the naming:
swath_combined['Experiment'] = swath_combined['Experiment'].replace({
    '1 Peptide': '1 Peptide',
    '2 Peptides': '2 Peptides',
    '%50 Peptides': '50% Peptides',
    'Random (1 to %50) Peptides': 'Random Peptides',
    'Random (2 to %50) Peptides': 'Random Peptides',
})
swath_combined['Experiment'].value_counts()
```

```
Experiment
2 Peptides         125
50% Peptides       125
Random Peptides    125
1 Peptide           75
Name: count, dtype: int64
```

###### MCC Performance Across Perturbation Scenarios¶

The barplot compares Matthews Correlation Coefficient (MCC) across four perturbation scenarios:

- **1 Peptide / 2 Peptides**: Single or paired peptide perturbations
- **Random Peptides**: Variable number (2–50%) of peptides perturbed
- **50% Peptides**: Half of all peptides perturbed

**Visual Elements:**

- Bars show mean MCC (± 95% CI) across thresholds
- Scatter points indicate MCC at p-value threshold (10⁻³)
- Horizontal reference lines denote MCC interpretation thresholds

```
from matplotlib.lines import Line2D
from matplotlib.gridspec import GridSpec

def compute_offset(n_methods, idx_method, total_width=0.8):
    # Calculates the horizontal offset for aligning elements with grouped bars.
    if n_methods <= 1:
        return 0.0
    single_width = total_width / n_methods
    center_offset = (n_methods - 1) / 2.0
    return (idx_method - center_offset) * single_width

def plot_benchmark_panel(ax, data, x_order, hue_order, palette, markers, p_threshold):
    # Generates a single panel with a bar plot, score annotations, and scatter points.
    sns.barplot(
        ax=ax, data=data, x='Experiment', y='MCC', hue='Method',
        order=x_order, hue_order=hue_order, palette=palette,
        edgecolor="black", linewidth=1.0, errorbar="ci",
        capsize=0.03, err_kws={'linewidth': 1.0}
    )

    n_methods = len(hue_order)
    pthr_data = data[data['threshold'] == p_threshold]

    for i, experiment in enumerate(x_order):
        for j, method in enumerate(hue_order):
            offset = compute_offset(n_methods, j)
            x_pos = i + offset

            mean_mcc = data[(data["Experiment"] == experiment) & (data["Method"] == method)]["MCC"].mean()
            if not np.isnan(mean_mcc):
                ax.text(
                    x_pos, -0.03, f"{mean_mcc:.2f}", color=palette.get(method, 'k'),
                    ha="center", va="top", fontsize=8, fontweight="bold", rotation=90,
                )

            method_pthr_data = pthr_data[(pthr_data['Method'] == method) & (pthr_data['Experiment'] == experiment)]
            if not method_pthr_data.empty:
                y_pos = method_pthr_data['MCC'].iloc[0]
                ax.scatter(
                    x_pos, y_pos, color=palette.get(method, 'k'), s=60,
                    edgecolor='black', linewidth=1.0, marker=markers.get(method, 'o'),
                    zorder=10, alpha=0.95
                )
    
    ax.grid(axis="y", linestyle="--", linewidth=0.5, alpha=0.7, color="gray")
    ax.set_xlabel('')
    ax.tick_params(axis='x', labelsize=9, rotation=90) # Rotate x-axis labels
    ax.tick_params(axis='y', labelsize=9)
    if ax.get_legend() is not None:
        ax.get_legend().remove()

# Filter and categorize data for plotting
experiments_ordered = ['1 Peptide', '2 Peptides', 'Random Peptides', '50% Peptides']
methods_ordered_pepid = ['ProteoForge', 'PeCorA', 'COPF']
methods_ordered_pepgrp = ['ProteoForge', 'COPF']

subset_pepid = swath_combined[(swath_combined['Benchmark'] == 'Peptide Identification') & (swath_combined['Experiment'].isin(experiments_ordered))].copy()
subset_pepid['Experiment'] = pd.Categorical(subset_pepid['Experiment'], categories=experiments_ordered, ordered=True)
subset_pepid['Method'] = pd.Categorical(subset_pepid['Method'], categories=methods_ordered_pepid, ordered=True)

subset_pepgrp = swath_combined[(swath_combined['Benchmark'] == 'Peptide Grouping') & (swath_combined['Experiment'].isin(experiments_ordered))].copy()
subset_pepgrp['Experiment'] = pd.Categorical(subset_pepgrp['Experiment'], categories=experiments_ordered, ordered=True)
subset_pepgrp['Method'] = pd.Categorical(subset_pepgrp['Method'], categories=methods_ordered_pepgrp, ordered=True)
    
# --- Figure Creation ---
fig = plt.figure(figsize=(12, 5))
# Adjust width ratios: ax1 wider (3 groups), ax2 narrower (2 groups)
gs = GridSpec(2, 20, height_ratios=[1, 0.15], hspace=0.4, wspace=0.025)
ax1 = fig.add_subplot(gs[0, :12])  # ax1: wider (12/20)
ax2 = fig.add_subplot(gs[0, 12:18], sharey=ax1)  # ax2: narrower (6/20)
ax_legend = fig.add_subplot(gs[1, :])
ax_legend.axis('off')

# Plot Peptide Identification panel
plot_benchmark_panel(ax1, subset_pepid, experiments_ordered, methods_ordered_pepid, method_palette, method_markers, pthr)
ax1.set_title("Peptide Identification", fontsize=12, fontweight="bold")
ax1.set_ylabel("MCC Score", fontsize=10)
ax1.set_ylim(-0.1, 1.05)
plt.setp(ax2.get_yticklabels(), visible=False)

# Draw MCC interpretation thresholds
for i, (thresh, label) in enumerate(mcc_thresholds.items()):
    ax1.axhline(
        thresh, color=mcc_colors[label], alpha=1,
        linestyle="dotted", linewidth=1.5, 
        label=label, zorder=0
    )
    ax1.text(
        0.01,
        thresh,
        label,
        color=mcc_colors[label],
        ha="left",
        va="center",
        fontsize=8,
        fontweight="bold",
        transform=ax1.get_yaxis_transform(),
        bbox=dict(boxstyle='round,pad=0.3', facecolor='white', edgecolor=mcc_colors[label], alpha=0.8),
        zorder=0
    )

# Plot Peptide Grouping panel
plot_benchmark_panel(ax2, subset_pepgrp, experiments_ordered, methods_ordered_pepgrp, method_palette, method_markers, pthr)
ax2.set_title("Peptide Grouping", fontsize=12, fontweight="bold")

# Set ylims (-0.15, to 0.75)
ax1.set_ylim(-0.15, 0.75)
ax2.set_ylim(-0.15, 0.75)

# Add threshold lines (without text) to the second plot
for thresh, label in mcc_thresholds.items():
    ax2.axhline(thresh, color=mcc_colors[label], alpha=0.9, linestyle=":", linewidth=1.2, zorder=0)

# --- Unified Legend ---
legend_elements = [
    Line2D([0], [0], marker=method_markers[m], color='w', markerfacecolor=method_palette[m],
           markeredgecolor='black', markersize=9, label=m)
    for m in methods_ordered_pepid
]
ax_legend.legend(
    handles=legend_elements, loc='center', ncol=len(methods_ordered_pepid), 
    fontsize=10, title="Method", title_fontsize=11, frameon=False
)

# --- Final Touches ---
plt.tight_layout(rect=[0, 0.1, 1, 1])
plt.subplots_adjust(bottom=0.2)
plots.finalize_plot(
    fig, show=True, save=save_to_folder,
    filename='benchmark_compact_fig2ab',
    filepath=figure_path,
    formats=figure_formats, 
    transparent=transparent_bg,
    dpi=figure_dpi,
)
```

**Key Observations:**

- **ProteoForge** consistently outperforms **COPF** and **PeCorA** across all scenarios except 50% Peptides where COPF edges out by a large margin.
  - This is due to 50% peptide can sometimes be 3/5 for odd numbers which would put PeCora and ProteoForge at a disadvantage since they rely on discordance.
- Even at 50% peptide perturbation, **ProteoForge** shows competitive performance in peptide grouping, which is notable given its challenges in peptide identification under this scenario.
- **PeCorA** is competitive in simpler scenarios (1 or 2 peptides) but lags in complex perturbations. And can't be in the grouping benchmark since it does not group peptides.

---

##### 2.2 Simulation 1: Impact of Data Imputation¶

**Objective:** Assess whether imputing missing values affects method performance compared to complete (fully quantified) datasets.

**Design:** Random perturbation scenario (2–50% peptides) with matched complete vs. imputed versions.

```
sim1_pepid = pd.read_feather('./data/Sim1/4_Sim1_PeptideIdentification_PerformanceData.feather')
sim1_pepgrp = pd.read_feather('./data/Sim1/4_Sim1_Grouping_PerformanceData.feather')
# Combine as sim1 with Benchmark column indicating pepID, or pepGrp
sim1_pepgrp['Benchmark'] = 'Peptide Grouping'
sim1_pepid['Benchmark'] = 'Peptide Identification'
sim1_combined = pd.concat([sim1_pepid, sim1_pepgrp], ignore_index=True)
sim1_combined['Experiment'].value_counts()
# Subset to show: '2>50% Peptides'
sim1_combined = sim1_combined[sim1_combined['Experiment'] == '2>50% Peptides']
# sim1_combined
```

###### Visualization Option 1: Horizontal Dumbbell Plot¶

Shows performance change between complete (●) and imputed (■) data with connecting lines indicating magnitude and direction of change.

```
fig, axs = plt.subplots(2, 1, figsize=(6, 4), sharex=True)
fig.suptitle('Imputation Impact on Performance', fontsize=12, fontweight='bold')

benchmarks = ['Peptide Identification', 'Peptide Grouping']
methods_order = {
    'Peptide Identification': ['COPF', 'PeCorA', 'ProteoForge'],
    'Peptide Grouping': ['COPF', 'ProteoForge']
}

for idx, benchmark in enumerate(benchmarks):
    ax = axs[idx]
    df_bench = sim1_combined[sim1_combined['Benchmark'] == benchmark]
    
    # Calculate mean MCC for complete and imputed data for each method
    means = df_bench.groupby(['Method', 'DataType'])['MCC'].mean().unstack()
    methods = methods_order[benchmark]
    means = means.reindex(methods)

    y_pos = np.arange(len(methods))
    
    # Draw compact dumbbell elements
    for i, method in enumerate(methods):
        mcc_complete = means.loc[method, 'complete']
        mcc_imputed = means.loc[method, 'imputed']
        color = method_palette.get(method, 'grey')
        
        # Thinner connecting line
        ax.plot([mcc_complete, mcc_imputed], [i, i], color=color, alpha=0.8, 
                linewidth=2, solid_capstyle='round', zorder=1)
        
        # Smaller markers
        ax.scatter(mcc_complete, i, color='black', marker='o', s=40, zorder=2, alpha=0.9)
        ax.scatter(mcc_imputed, i, color=color, marker='s', s=35, zorder=2)
        
        # Add difference annotation
        diff = mcc_imputed - mcc_complete
        mid_x = (mcc_complete + mcc_imputed) / 2
        ax.text(mid_x, i + 0.15, f'{diff:+.3f}', ha='center', va='bottom', 
                fontsize=8, color=color, fontweight='bold')

    # Compact aesthetics
    ax.set_title(benchmark, fontsize=10, fontweight='bold', pad=8)
    ax.grid(axis='x', linestyle=':', alpha=0.5)
    ax.set_yticks(y_pos)
    ax.set_yticklabels(methods, fontsize=9)
    ax.spines['top'].set_visible(False)
    ax.spines['right'].set_visible(False)
    ax.spines['left'].set_visible(False)
    ax.tick_params(axis='y', length=0)
    ax.tick_params(axis='x', labelsize=8)

axs[-1].set_xlabel('MCC Score', fontsize=9)

# Compact legend
legend_elements = [
    Line2D([0], [0], color='gray', marker='o', linestyle='None', markersize=6, 
           markerfacecolor='black', label='Complete'),
    Line2D([0], [0], color='gray', marker='s', linestyle='None', markersize=6, 
           label='Imputed'),
]

fig.legend(handles=legend_elements, loc='lower center', ncol=2, 
           bbox_to_anchor=(0.5, -0.02), fontsize=8, frameon=False)

plt.tight_layout(rect=[0, 0.05, 1, 0.95])
plt.show()
```

**Assessment:** Clear visualization of magnitude changes; distinct but uses significant whitespace.

###### Visualization Option 2: Slope Graph¶

Connects performance from "Complete" to "Imputed" conditions—effective for showing direction when changes are large.

```
fig, axs = plt.subplots(1, 2, figsize=(8, 3), sharey=True)
fig.suptitle('Performance: Complete vs Imputed Data', fontsize=12, fontweight='bold')

benchmarks = ['Peptide Identification', 'Peptide Grouping']
methods_order = {'Peptide Identification': ['COPF', 'PeCorA', 'ProteoForge'],
                 'Peptide Grouping': ['COPF', 'ProteoForge']}

for idx, benchmark in enumerate(benchmarks):
    ax = axs[idx]
    df_bench = sim1_combined[sim1_combined['Benchmark'] == benchmark]
    
    means = df_bench.groupby(['Method', 'DataType'])['MCC'].mean().unstack()
    methods = methods_order[benchmark]
    means = means.reindex(methods)
    
    # Create slope lines
    x_positions = [0, 1]  # Complete at 0, Imputed at 1
    
    for i, method in enumerate(methods):
        mcc_complete = means.loc[method, 'complete']
        mcc_imputed = means.loc[method, 'imputed']
        color = method_palette.get(method, 'grey')
        
        # Draw slope line
        ax.plot(x_positions, [mcc_complete, mcc_imputed], 
                color=color, linewidth=2, alpha=0.8, marker='o', markersize=6,
                label=method if idx == 0 else "")
        
        # Method labels at the midpoint
        mid_y = (mcc_complete + mcc_imputed) / 2
        ax.text(0.5, mid_y, method, ha='center', va='center', 
                fontsize=8, fontweight='bold', color=color,
                bbox=dict(boxstyle='round,pad=0.2', facecolor='white', alpha=0.8))

    ax.set_title(benchmark, fontsize=10, fontweight='bold')
    ax.set_xticks(x_positions)
    ax.set_xticklabels(['Complete', 'Imputed'], fontsize=9)
    ax.set_ylabel('MCC Score' if idx == 0 else '', fontsize=9)
    ax.grid(axis='y', linestyle=':', alpha=0.5)
    ax.spines['top'].set_visible(False)
    ax.spines['right'].set_visible(False)
    ax.tick_params(labelsize=8)
    ax.set_ylim(0, 0.8)  # Set consistent y-axis range

plt.tight_layout(rect=[0, 0, 1, 0.95])
plt.show()
```

**Assessment:** Small performance differences result in nearly horizontal lines—less effective for this dataset.

###### Visualization Option 3: Grouped Bar Chart¶

Traditional side-by-side comparison with hierarchical x-axis grouping methods by benchmark.

```
fig, ax = plt.subplots(figsize=(10, 4))
fig.suptitle('Impact of Data Imputation', fontsize=14, fontweight='bold')

# Prepare data for plotting
plot_data = []
benchmarks = ['Peptide Identification', 'Peptide Grouping']

for benchmark in benchmarks:
    df_bench = sim1_combined[sim1_combined['Benchmark'] == benchmark]
    means = df_bench.groupby(['Method', 'DataType'])['MCC'].mean().unstack()
    
    for method in means.index:
        complete_val = means.loc[method, 'complete']
        imputed_val = means.loc[method, 'imputed']
        diff = imputed_val - complete_val
        
        plot_data.append({
            'Benchmark': benchmark,
            'Method': method,
            'Complete': complete_val,
            'Imputed': imputed_val,
            'Difference': diff
        })

plot_df = pd.DataFrame(plot_data)

# Create hierarchical x positions
n_benchmarks = len(benchmarks)
n_methods_per_bench = {'Peptide Identification': 3, 'Peptide Grouping': 2}  # COPF, PeCorA, ProteoForge vs COPF, ProteoForge
method_order = {'Peptide Identification': ['COPF', 'PeCorA', 'ProteoForge'], 
                'Peptide Grouping': ['COPF', 'ProteoForge']}

x_positions = []
labels = []
benchmark_centers = []

x = 0
for i, benchmark in enumerate(benchmarks):
    methods = method_order[benchmark]
    start_x = x
    
    for j, method in enumerate(methods):
        method_data = plot_df[(plot_df['Benchmark'] == benchmark) & (plot_df['Method'] == method)]
        if not method_data.empty:
            row = method_data.iloc[0]
            
            # Plot bars
            width = 0.35
            bar1 = ax.bar(x - width/2, row['Complete'], width, color='lightgray', 
                         edgecolor='black', alpha=0.8, label='Complete' if i == 0 and j == 0 else "")
            bar2 = ax.bar(x + width/2, row['Imputed'], width,
                         color=method_palette.get(method, 'gray'), edgecolor='black', alpha=0.9,
                         label='Imputed' if i == 0 and j == 0 else "")
            
            # Add change indicators
            diff = row['Difference']
            arrow_color = 'green' if diff > 0 else 'red'
            arrow_symbol = '↑' if diff > 0 else '↓'
            
            ax.text(x, max(row['Complete'], row['Imputed']) + 0.05, 
                   f'{arrow_symbol} {abs(diff):.3f}', 
                   ha='center', va='bottom', fontsize=9, 
                   color=arrow_color, fontweight='bold')
            
            x_positions.append(x)
            labels.append(method)
        
        x += 1
    
    # Calculate benchmark center for labeling
    end_x = x - 1
    benchmark_centers.append((start_x + end_x) / 2)
    x += 0.5  # Space between benchmarks

# Set labels and ticks
ax.set_xticks(x_positions)
ax.set_xticklabels(labels, fontsize=10, rotation=45, ha='right')

# Add benchmark labels as secondary x-axis
ax2 = ax.twiny()
ax2.set_xlim(ax.get_xlim())
ax2.set_xticks(benchmark_centers)
ax2.set_xticklabels(benchmarks, fontsize=11, fontweight='bold')
ax2.tick_params(axis='x', length=0)

# Styling
ax.set_ylabel('MCC Score', fontsize=11)
ax.grid(axis='y', linestyle=':', alpha=0.5)
ax.spines['top'].set_visible(False)
ax.spines['right'].set_visible(False)
ax2.spines['top'].set_visible(False)
ax2.spines['right'].set_visible(False)
ax2.spines['bottom'].set_visible(False)
ax2.spines['left'].set_visible(False)

# Add vertical separator between benchmarks
separator_x = x_positions[2] + 0.75  # Between ProteoForge (PepID) and COPF (PepGrp)
ax.axvline(separator_x, color='gray', linestyle='--', alpha=0.5)

# Legend
handles, labels_leg = ax.get_legend_handles_labels()
ax.legend(handles, labels_leg, loc='upper left', fontsize=10, frameon=False)

plt.tight_layout()
plt.show()
```

**Assessment:** Intuitive layout but horizontally expansive—better suited for supplementary material.

###### Visualization Option 4: Vertical Overlapping Bars *(Selected)*¶

Compact vertical layout overlaying imputed (colored) on complete (gray) bars with annotated differences.

```
fig, axs = plt.subplots(2, 1, figsize=(4, 6), sharex=True)
fig.suptitle('MCC Performance: Complete vs Imputed Data (Random Perturbation (Sim1))', fontsize=12, fontweight='bold')

benchmarks = ['Peptide Identification', 'Peptide Grouping']
methods_order = {'Peptide Identification': ['COPF', 'PeCorA', 'ProteoForge'],
                 'Peptide Grouping': ['COPF', 'ProteoForge']}

for idx, benchmark in enumerate(benchmarks):
    ax = axs[idx]
    df_bench = sim1_combined[sim1_combined['Benchmark'] == benchmark]
    
    means = df_bench.groupby(['Method', 'DataType'])['MCC'].mean().unstack()
    methods = methods_order[benchmark]
    means = means.reindex(methods)
    
    y_pos = np.arange(len(methods))
    colors = [method_palette.get(method, 'gray') for method in methods]
    
    # Plot complete data (baseline) as light bars
    complete_bars = ax.barh(y_pos, means['complete'], height=0.6, 
                           color='lightgray', alpha=0.6, edgecolor='black',
                           label='Complete Data')
    
    # Plot imputed data as colored bars
    imputed_bars = ax.barh(y_pos, means['imputed'], height=0.4, 
                          color=colors, alpha=0.9, edgecolor='black',
                          label='Imputed Data')
    
    # Add MCC values and differences as text
    for i, method in enumerate(methods):
        complete_val = means.loc[method, 'complete']
        imputed_val = means.loc[method, 'imputed']
        diff = imputed_val - complete_val
        
        # Show complete MCC value
        ax.text(complete_val + 0.01, i + 0.15, f'{complete_val:.3f}', 
               va='center', ha='left', fontsize=9, color='black', fontweight='bold')
        
        # Show imputed MCC value  
        ax.text(imputed_val + 0.01, i - 0.15, f'{imputed_val:.3f}', 
               va='center', ha='left', fontsize=9, color=colors[i], fontweight='bold')
        
        # Show difference with arrow
        diff_color = 'green' if diff > 0 else 'red'
        arrow_symbol = '↑' if diff > 0 else '↓'
        
        # Position difference text at the right edge
        max_val = max(complete_val, imputed_val)
        ax.text(max_val + 0.08, i, f'{arrow_symbol}{abs(diff):.3f}', 
               va='center', ha='left', fontsize=8, 
               color=diff_color, fontweight='bold',
               bbox=dict(boxstyle='round,pad=0.2', facecolor='white', 
                        edgecolor=diff_color, alpha=0.8))
    
    # Styling
    # ax.set_title(benchmark, fontsize=11, fontweight='bold', pad=5)
    ax.set_yticks(y_pos)
    ax.set_yticklabels(methods, fontsize=10, fontweight='bold')
    if idx == 1:  # Only show x-axis label on bottom plot
        ax.set_xlabel('MCC Score', fontsize=10)
    ax.grid(axis='x', linestyle=':', alpha=0.4)
    ax.spines['top'].set_visible(False)
    ax.spines['right'].set_visible(False)
    ax.tick_params(labelsize=12)
    
    # Set consistent x-axis limits for all subplots
    ax.set_xlim(0, 0.7)  # Fixed range to accommodate all data

# Unified legend at the bottom
handles, labels_leg = axs[0].get_legend_handles_labels()
fig.legend(handles, labels_leg, loc='lower center', ncol=2, 
           bbox_to_anchor=(0.5, -0.05), fontsize=10, frameon=False)

# Add explanation
fig.text(0.5, -0.12, 'Green ↑: Improvement with imputation | Red ↓: Decline with imputation', 
         ha='center', fontsize=9, style='italic')

plt.tight_layout(rect=[0, 0.12, 1, 0.95])
plt.show()
plots.finalize_plot(
    fig, show=True, save=save_to_folder,
    filename='benchmark_sim1_imputation_effect',
    filepath=figure_path,
    formats=figure_formats, 
    transparent=transparent_bg,
    dpi=figure_dpi,
)
```

**Key Findings — Simulation 1:**

- In peptide identification the imputation has small impact for ProteoForge and COPF but PeCorA declines sharply
- For peptide grouping, ProteoForge actually improves slightly with imputation while COPF decline slightly.
  - This is due to with imputation, especially complete missingness does introduce potential new groups of peptides, proteoforge is capable of grouping them correctly whereas COPF fails to do so.
- Selected visualization (Option 4) provides optimal information density for main figure

---

##### 2.3 Simulation 2: Effect of Missingness Level¶

**Objective:** Quantify how increasing proportions of missing data affect detection performance.

**Design:** Factorial combinations of protein-level (0–80%) × peptide-level (0–80%) missingness.

**Key Questions:**

- At what missingness threshold does performance critically degrade?
- Do methods differ in their tolerance to incomplete data?

```
# Load and explore Sim2 data (Missingness analysis)
sim2_pepid = pd.read_feather('./data/Sim2/4_Sim2_PeptideIdentification_PerformanceData.feather')
sim2_pepgrp = pd.read_feather('./data/Sim2/4_Sim2_Grouping_PerformanceData.feather')

# Combine data
sim2_pepgrp['Benchmark'] = 'Peptide Grouping'
sim2_pepid['Benchmark'] = 'Peptide Identification'
sim2_combined = pd.concat([sim2_pepid, sim2_pepgrp], ignore_index=True)

# Clean up the missingness data
def clean_missingness_values(x):
    if isinstance(x, str):
        return float(x.rstrip('%')) / 100.0
    return float(x)

sim2_combined['ProteinMissingness'] = sim2_combined['ProteinMissingness'].apply(clean_missingness_values)
sim2_combined['PeptideMissingness'] = sim2_combined['PeptideMissingness'].apply(clean_missingness_values)

sim2_summary = sim2_combined.groupby(['Method', 'ProteinMissingness', 'PeptideMissingness', 'Benchmark'])['MCC'].mean().reset_index()
```

###### Visualization Option 1: Degradation Line Plots¶

MCC trajectories as peptide missingness increases (x-axis), stratified by protein missingness levels (line styles: solid=0%, dashed=40%, dotted=80%).

```
def create_degradation_lines(figsize=(6, 5)):
    """Show how performance degrades with increasing missingness - Vertical Layout"""
    
    fig, axs = plt.subplots(2, 1, figsize=figsize, sharex=True)
    fig.suptitle('Performance Degradation with Missingness', fontsize=11, fontweight='bold')
    
    benchmarks = ['Peptide Identification', 'Peptide Grouping']
    method_sets = {
        'Peptide Identification': ['COPF', 'PeCorA', 'ProteoForge'],
        'Peptide Grouping': ['COPF', 'ProteoForge']  # No PeCorA for grouping
    }
    
    for idx, benchmark in enumerate(benchmarks):
        ax = axs[idx]
        methods = method_sets[benchmark]
        
        # Plot lines for each method at different protein missingness levels
        protein_levels = [0.0, 0.4, 0.8]  # Show key levels only
        
        for method in methods:
            color = method_palette.get(method, 'gray')
            
            for i, prot_miss in enumerate(protein_levels):
                method_data = sim2_summary[(sim2_summary['Method'] == method) & 
                                          (sim2_summary['Benchmark'] == benchmark) &
                                          (sim2_summary['ProteinMissingness'] == prot_miss)]
                
                # Sort by peptide missingness
                method_data = method_data.sort_values('PeptideMissingness')
                
                linestyle = ['-', '--', ':'][i]  # Different styles for different protein missingness
                alpha = 1.0 if i == 0 else 0.7
                linewidth = 2.5 if method == 'ProteoForge' else 2
                
                ax.plot(method_data['PeptideMissingness'] * 100, method_data['MCC'], 
                       color=color, linestyle=linestyle, marker='o', markersize=3,
                       alpha=alpha, linewidth=linewidth)
        
        ax.set_title(benchmark, fontsize=10, fontweight='bold', pad=5)
        if idx == 1:  # Only bottom plot gets x-axis label
            ax.set_xlabel('Peptide Missingness (%)', fontsize=9)
        ax.set_ylabel('MCC Score', fontsize=9)
        ax.grid(True, linestyle=':', alpha=0.4)
        ax.spines['top'].set_visible(False)
        ax.spines['right'].set_visible(False)
        ax.tick_params(labelsize=8)
        ax.set_ylim(0, 0.85)
    
    # Create custom legend
    legend_elements = []
    for method in ['COPF', 'PeCorA', 'ProteoForge']:
        color = method_palette.get(method, 'gray')
        legend_elements.append(Line2D([0], [0], color=color, lw=2, label=method))
    
    # Add line style legend
    legend_elements.extend([
        Line2D([0], [0], color='gray', linestyle='-', lw=1, label='Protein 0%'),
        Line2D([0], [0], color='gray', linestyle='--', lw=1, label='Protein 40%'),
        Line2D([0], [0], color='gray', linestyle=':', lw=1, label='Protein 80%')
    ])
    
    fig.legend(handles=legend_elements, loc='lower center', ncol=3, 
              bbox_to_anchor=(0.5, -0.05), fontsize=8, frameon=False)
    
    plt.tight_layout(rect=[0, 0.08, 1, 0.95])
    return fig

create_degradation_lines()
plt.show()
```

**Assessment:** Shows interaction between protein and peptide missingness, easy to spot the most obvious point, however not clean and details such as the protein level missingness is hard to read.

###### Visualization Option 2: Compact Line Plot with Change Annotations *(Selected)*¶

Shows MCC at diagonal missingness levels (Complete → Low → Moderate → High → Extreme) with Δ annotations, calculated from Complete - Others.

```
fig, axs = plt.subplots(2, 1, figsize=(5, 5), sharex=True)
fig.suptitle('Performance Change Across Missingness Levels', fontsize=12, fontweight='bold')

benchmarks = ['Peptide Identification', 'Peptide Grouping']
method_sets = {
    'Peptide Identification': ['COPF', 'PeCorA', 'ProteoForge'],
    'Peptide Grouping': ['COPF', 'ProteoForge']
}

change_levels = [
    ('Complete\n(0%,0%)', 0.0, 0.0),
    ('Low\n(20%,20%)', 0.2, 0.2),
    ('Moderate\n(40%,40%)', 0.4, 0.4),
    ('High\n(60%,60%)', 0.6, 0.6),
    ('Extreme\n(80%,80%)', 0.8, 0.8)
]
change_labels = [x[0] for x in change_levels]

for idx, benchmark in enumerate(benchmarks):
    ax = axs[idx]
    methods = method_sets[benchmark]
    for method in methods:
        color = method_palette.get(method, 'gray')
        marker = method_markers.get(method, 'o')
        mcc_vals = []
        for _, prot_miss, pep_miss in change_levels:
            mcc_val = sim2_summary[(sim2_summary['Method'] == method) &
                                   (sim2_summary['Benchmark'] == benchmark) &
                                   (sim2_summary['ProteinMissingness'] == prot_miss) &
                                   (sim2_summary['PeptideMissingness'] == pep_miss)]['MCC'].values
            mcc_vals.append(mcc_val[0] if len(mcc_val) > 0 else np.nan)
        ax.plot(change_labels, mcc_vals, color=color, marker=marker, markersize=8, linewidth=2.5,
                label=method, alpha=0.9, markeredgecolor='black', markeredgewidth=1.5)
        # Annotate difference from Complete
        for i in range(1, len(mcc_vals)):
            if not np.isnan(mcc_vals[0]) and not np.isnan(mcc_vals[i]):
                diff = mcc_vals[i] - mcc_vals[0]
                arrow = '↑' if diff > 0 else '↓'
                color_diff = 'green' if diff > 0 else 'red'
                ax.text(change_labels[i], mcc_vals[i] + 0.03, f'{arrow}{abs(diff):.2f}',
                        ha='center', va='bottom', fontsize=8, color=color_diff, fontweight='bold')
        # Annotate the complete MCC value
        if not np.isnan(mcc_vals[0]):
            ax.text(change_labels[0], mcc_vals[0] - 0.04, f'{mcc_vals[0]:.2f}',
                    ha='center', va='top', fontsize=8, color=color, fontweight='bold',
                    bbox=dict(boxstyle='round,pad=0.2', facecolor='white', edgecolor=color, alpha=0.7))
    ax.set_title(benchmark, fontsize=10, fontweight='bold', pad=5)
    ax.set_ylabel('MCC Score', fontsize=9)
    ax.grid(axis='y', linestyle=':', alpha=0.4)
    ax.spines['top'].set_visible(False)
    ax.spines['right'].set_visible(False)
    ax.tick_params(labelsize=8)
    ax.set_ylim(-.15, 0.75)

axs[-1].set_xlabel('Missingness Level (Protein%, Peptide%)', fontsize=9)
axs[-1].set_xticks(change_labels)
axs[-1].set_xticklabels(change_labels, fontsize=9)

# Unified legend
handles, labels = axs[0].get_legend_handles_labels()
fig.legend(handles, labels, loc='lower center', ncol=3, bbox_to_anchor=(0.5, -0.08), fontsize=9, frameon=False)

plt.tight_layout(rect=[0, 0.08, 1, 0.95])
plt.show()
plots.finalize_plot(
    fig, show=True, save=save_to_folder,
    filename='benchmark_sim2_missingness_effect',
    filepath=figure_path,
    formats=figure_formats, 
    transparent=transparent_bg,
    dpi=figure_dpi,
)
```

**Assessment:** Clearly shows degradation trend; annotations highlight performance loss magnitude.

**Key Findings — Simulation 2:**

- Peptide Identification:
  - Performance degrades progressively with increasing missingness
  - Methods maintain reasonable accuracy (~70% retention) up to 40% missingness
  - ProteoForge shows minimal effect until 60% High missingness, at 80% drops extremely sharply
    - This is due to RLM's internal robust weighting mechanism consider the imputed values to be more reliable than real values, since they are more prevalent.
    - In WLS with imputation weighting system passed, the decline is less sharp, however the lower missingness gains from RLM is lost.
  - Beyond 60% missingness, detection reliability decreases substantially
- Peptide Grouping:
  - Peptide grouping improves slightly with low, but shows decline at higher missingness for ProteoForge, again at extreme it drops sharply.
  - COPF also shows it shows that it gets down by .85-75 at all levels, meaning the amount of missingness doesn't matter much.

---

##### 2.4 Simulation 3: Detection Sensitivity vs. Perturbation Magnitude¶

**Objective:** Determine the minimum perturbation magnitude required for reliable detection.

**Design:** Perturbation magnitudes ranging from subtle (0.10–0.25 log2) to strong (1.75–2.00 log2) with complete data (no missing values).

**Key Questions:**

- What is the practical detection threshold for each method?
- How does sensitivity scale with perturbation size?

```
# Load and explore Sim3 data (Magnitude of perturbation levels)
sim3_pepid = pd.read_feather('./data/Sim3/4_Sim3_PeptideIdentification_PerformanceData.feather')
sim3_pepid['Benchmark'] = 'Peptide Identification'
sim3_pepgrp = pd.read_feather('./data/Sim3/4_Sim3_Grouping_PerformanceData.feather')
sim3_pepgrp['Benchmark'] = 'Peptide Grouping'
sim3_combined = pd.concat([sim3_pepid, sim3_pepgrp], ignore_index=True)
# sim3_combined

pert_ranges = sorted(sim3_combined['PerturbationRange'].unique())

# Process sim3 data for visualization
def process_sim3_data():
    """Process sim3 data and normalize perturbation ranges"""
    
    # Create a copy to work with
    sim3_processed = sim3_combined.copy()
    
    # Normalize perturbation range formatting and create magnitude column
    def extract_magnitude(pert_range):
        """Extract average magnitude from perturbation range string"""
        if isinstance(pert_range, str) and '-' in pert_range:
            try:
                start, end = pert_range.split('-')
                return (float(start) + float(end)) / 2
            except:
                return None
        return None
    
    sim3_processed['Magnitude'] = sim3_processed['PerturbationRange'].apply(extract_magnitude)
    
    # Normalize perturbation range names (handle 0.1 vs 0.10 inconsistency)
    def normalize_range(pert_range):
        if isinstance(pert_range, str) and '-' in pert_range:
            try:
                start, end = pert_range.split('-')
                start_f = float(start)
                end_f = float(end)
                return f"{start_f:.2f}-{end_f:.2f}"
            except:
                return pert_range
        return pert_range
    
    sim3_processed['PerturbationRange_norm'] = sim3_processed['PerturbationRange'].apply(normalize_range)
    
    # Get best MCC for each combination (since there are multiple thresholds)
    sim3_summary = sim3_processed.groupby(['Method', 'Benchmark', 'PerturbationRange_norm', 'Magnitude'])['MCC'].mean().reset_index()
    
    # Sort by magnitude for proper ordering
    sim3_summary = sim3_summary.sort_values('Magnitude')
    
    print("Processed sim3 summary:")
    print(f"Shape: {sim3_summary.shape}")
    print(f"Magnitude levels: {sorted(sim3_summary['Magnitude'].unique())}")
    print(f"Perturbation ranges: {sorted(sim3_summary['PerturbationRange_norm'].unique())}")
    
    return sim3_summary

sim3_summary = process_sim3_data()
sim3_summary.head()
```

```
Processed sim3 summary:
Shape: (40, 5)
Magnitude levels: [np.float64(0.175), np.float64(0.375), np.float64(0.625), np.float64(0.875), np.float64(1.125), np.float64(1.375), np.float64(1.625), np.float64(1.875)]
Perturbation ranges: ['0.10-0.25', '0.25-0.50', '0.50-0.75', '0.75-1.00', '1.00-1.25', '1.25-1.50', '1.50-1.75', '1.75-2.00']
```

|  | Method | Benchmark | PerturbationRange\_norm | Magnitude | MCC |
| --- | --- | --- | --- | --- | --- |
| 0 | COPF | Peptide Grouping | 0.10-0.25 | 0.1750 | 0.0014 |
| 8 | COPF | Peptide Identification | 0.10-0.25 | 0.1750 | -0.0050 |
| 24 | ProteoForge | Peptide Grouping | 0.10-0.25 | 0.1750 | 0.4368 |
| 16 | PeCorA | Peptide Identification | 0.10-0.25 | 0.1750 | 0.0296 |
| 32 | ProteoForge | Peptide Identification | 0.10-0.25 | 0.1750 | 0.1927 |

###### Visualization Option 1: Detection Curves with Endpoint Annotations *(Selected)*¶

MCC vs. perturbation magnitude with method-specific trajectories and Δ(MCC) summaries.

```
def create_detection_curves_combined(figsize=(3.2, 6)):
    """Stylized detection curves with distribution-of-change summaries to the right."""
    fig, ax = plt.subplots(figsize=figsize)
    fig.suptitle('Perturbation Detection Sensitivity', fontsize=11, fontweight='bold', y=0.98)

    # Ensure sim3_combined has a Magnitude column (average of perturbation range endpoints)
    sim3_full = sim3_combined.copy()
    def _extract_mag(pert):
        if isinstance(pert, str) and ('-' in pert):
            try:
                a, b = pert.split('-')
                return (float(a) + float(b)) / 2.0
            except:
                return np.nan
        return np.nan
    if 'Magnitude' not in sim3_full.columns:
        sim3_full['Magnitude'] = sim3_full['PerturbationRange'].apply(_extract_mag)

    benchmarks = ['Peptide Identification', 'Peptide Grouping']
    method_sets = {
        'Peptide Identification': ['COPF', 'PeCorA', 'ProteoForge'],
        'Peptide Grouping': ['COPF', 'ProteoForge']
    }
    linestyle_map = {
        'Peptide Identification': '-',
        'Peptide Grouping': '--'
    }

    # Plot curves
    plotted_info = []  # collect info for summary plotting (method, benchmark, first_mag, last_mag)
    for benchmark in benchmarks:
        methods = method_sets[benchmark]
        for method in methods:
            method_data = sim3_summary[
                (sim3_summary['Method'] == method) & 
                (sim3_summary['Benchmark'] == benchmark)
            ].sort_values('Magnitude')
            if len(method_data) > 0:
                color = method_palette.get(method, 'black')
                marker = method_markers.get(method, 'o')
                ls = linestyle_map[benchmark]
                ax.plot(
                    method_data['Magnitude'], method_data['MCC'],
                    linestyle=ls, color=color, linewidth=2.5,
                    marker=marker, markersize=7, alpha=0.9,
                    markerfacecolor=color, markeredgecolor='black'
                )

                mag_vals = method_data['Magnitude'].values
                mcc_vals = method_data['MCC'].values

                # Directly label at the end of the curve
                label_text = "ID" if benchmark == "Peptide Identification" else "Group"
                ax.text(
                    mag_vals[-1] + 0.03, mcc_vals[-1],
                    f"{label_text}",
                    ha='left', va='center', fontsize=8, fontweight='bold',
                    color=color, bbox=dict(boxstyle='round,pad=0.2', facecolor='white', alpha=0.8)
                )

                plotted_info.append({
                    'method': method,
                    'benchmark': benchmark,
                    'first_mag': mag_vals[0],
                    'last_mag': mag_vals[-1],
                    'color': color
                })

    ax.set_xlabel('Mean Perturbation Magnitude (log2)', fontsize=9)
    ax.set_ylabel('MCC Score', fontsize=10)
    ax.set_xlim(0.1, 2.0)
    ax.set_ylim(-0.05, 1.0)
    ax.grid(True, alpha=0.3, linestyle='--')
    mag_ticks = [0.2, 0.6, 1.0, 1.4, 1.8]
    ax.set_xticks(mag_ticks)
    ax.set_xticklabels([str(x) for x in mag_ticks], fontsize=8)
    ax.tick_params(axis='y', labelsize=8)

    # --- Summary to the right: vertical lines with annotation spanning MCC change ---
    if len(plotted_info) > 0:
        max_mag = sim3_summary['Magnitude'].max()
        x_base = max_mag + 0.08
        # create small offsets so multiple methods don't overlap
        max_methods = max(len(method_sets[b]) for b in benchmarks)
        offsets = np.linspace(-0.06, 0.06, max_methods)

        for bench_idx, benchmark in enumerate(benchmarks):
            methods = method_sets[benchmark]
            for m_idx, method in enumerate(methods):
                info = next((it for it in plotted_info if it['method'] == method and it['benchmark'] == benchmark), None)
                if info is None:
                    continue

                first_mag = info['first_mag']
                last_mag = info['last_mag']
                color = info['color']

                # Get MCC at first and last magnitude
                mrow_first = sim3_summary[
                    (sim3_summary['Method'] == method) &
                    (sim3_summary['Benchmark'] == benchmark) &
                    (sim3_summary['Magnitude'] == first_mag)
                ]
                mrow_last = sim3_summary[
                    (sim3_summary['Method'] == method) &
                    (sim3_summary['Benchmark'] == benchmark) &
                    (sim3_summary['Magnitude'] == last_mag)
                ]
                if len(mrow_first) > 0 and len(mrow_last) > 0:
                    mcc_first = mrow_first['MCC'].median()
                    mcc_last = mrow_last['MCC'].median()
                    median_diff = mcc_last - mcc_first
                    norm_diff = median_diff / (abs(mcc_first) + 1e-6)

                    x_pos = x_base + offsets[m_idx]
                    # Draw vertical line spanning MCC range
                    ax.plot([x_pos, x_pos], [mcc_first, mcc_last], color=color, linewidth=2.5, alpha=0.8, linestyle=linestyle_map[benchmark])
                    # Annotate at midpoint
                    mid_y = (mcc_first + mcc_last) / 2
                    txt = (f"Δ={median_diff:.3f}")
                    ax.text(x_pos + 0.03, mid_y, txt, fontsize=7, va='center', ha='left',
                            color=color, bbox=dict(boxstyle='round,pad=0.2', facecolor='white', alpha=0.8))

        # expand x-limits to accommodate the summaries
        ax.set_xlim(ax.get_xlim()[0], x_base + 0.18)


    plt.tight_layout()
    return fig

fig = create_detection_curves_combined(figsize=(4, 7))
plt.show()
plots.finalize_plot(
    fig, show=True, save=save_to_folder,
    filename='benchmark_sim3_perturbation_detection',
    filepath=figure_path,
    formats=figure_formats, 
    transparent=transparent_bg,
    dpi=figure_dpi,
)
```

**Assessment:** Comprehensive view of sensitivity curves; endpoint Δ annotations aid interpretation.

###### Visualization Option 2: Detection Efficiency Bars¶

Compact comparison at categorical magnitude levels (Low → Mid-Low → Mid-High → High).

```
# SIM3 Compact Visualization Approach 3: Detection Efficiency Bar Chart (VERTICAL)
def create_detection_efficiency_bars(figsize=(5, 4)):
    """Show detection efficiency at key perturbation levels"""
    
    fig, axs = plt.subplots(2, 1, figsize=figsize, sharex=True)
    fig.suptitle('Detection Efficiency at Key Perturbation Levels', fontsize=11, fontweight='bold')
    
    # Select key perturbation levels for comparison
    key_magnitudes = [0.175, 0.625, 1.125, 1.875]  # Low, Mid-low, Mid-high, High
    mag_labels = ['Low\n(~0.2)', 'Mid-Low\n(~0.6)', 'Mid-High\n(~1.1)', 'High\n(~1.9)']
    
    benchmarks = ['Peptide Identification', 'Peptide Grouping']
    method_sets = {
        'Peptide Identification': ['COPF', 'PeCorA', 'ProteoForge'],
        'Peptide Grouping': ['COPF', 'ProteoForge']
    }
    
    for i, benchmark in enumerate(benchmarks):
        ax = axs[i]
        methods = method_sets[benchmark]
        
        # Prepare data for bar chart
        x_positions = np.arange(len(key_magnitudes))
        width = 0.8 / len(methods)
        
        for m_idx, method in enumerate(methods):
            method_scores = []
            
            for magnitude in key_magnitudes:
                method_data = sim3_summary[
                    (sim3_summary['Method'] == method) & 
                    (sim3_summary['Benchmark'] == benchmark) &
                    (sim3_summary['Magnitude'] == magnitude)
                ]
                
                if len(method_data) > 0:
                    method_scores.append(method_data['MCC'].iloc[0])
                else:
                    method_scores.append(0)
            
            # Plot bars
            color = method_palette.get(method, 'black')
            x_pos = x_positions + (m_idx - len(methods)/2 + 0.5) * width
            
            bars = ax.bar(x_pos, method_scores, width, 
                         color=color, alpha=0.8, label=method,
                         edgecolor='white', linewidth=0.5)
            
            # Add value labels on bars
            for bar, score in zip(bars, method_scores):
                if score > 0.05:  # Only show if meaningful
                    ax.text(bar.get_x() + bar.get_width()/2, bar.get_height() + 0.02,
                           f'{score:.2f}', ha='center', va='bottom', 
                           fontsize=6, fontweight='bold', color=color)
        
        # Formatting
        ax.set_title(benchmark, fontsize=10, fontweight='bold', loc='left')
        ax.set_ylabel('MCC Score', fontsize=9)
        ax.set_ylim(0, 1.1)
        ax.set_xticks(x_positions)
        ax.grid(True, alpha=0.3, axis='y', linestyle='--')
        
        if i == 0:
            ax.legend(bbox_to_anchor=(1.02, 1), loc='upper left', fontsize=8)
        
        if i == 1:  # Bottom plot
            ax.set_xlabel('Perturbation Magnitude Level', fontsize=9)
            ax.set_xticklabels(mag_labels, fontsize=8)
        else:
            ax.set_xticklabels([])
    
    plt.tight_layout()
    return fig

create_detection_efficiency_bars()
plt.show()
```

**Key Findings — Simulation 3:**

- MCC increases monotonically with perturbation magnitude across all methods
- Practical detection threshold: ~0.5 log2 (MCC improves sharply above this level)
- ProteoForge shows strong sensitivity for the Peptide Identification benchmark
- Methods converge at high magnitudes—differences are most pronounced for subtle perturbations

---

##### 2.5 Simulation 4: Complex Multi-Variable Experimental Designs¶

**Objective:** Evaluate method robustness across experimental designs with multiple interacting factors.

**Design Factors:**

- **Number of conditions**: 2, 4, 6, or 8 experimental groups
- **Overlap pattern**: Same peptides perturbed across conditions (True) vs. different peptides (False)
- **Direction**: All perturbations in same direction vs. random up/down changes

**Key Questions:**

- Does adding more experimental conditions improve detection?
- Which perturbation patterns are easier to detect?

```
# Load and explore Sim3 data (Magnitude of perturbation levels)
sim4_pepid = pd.read_feather('./data/Sim4/4_Sim4_PeptideIdentification_PerformanceData.feather')
sim4_pepid['Benchmark'] = 'Peptide Identification'
sim4_pepgrp = pd.read_feather('./data/Sim4/4_Sim4_Grouping_PerformanceData.feather')
sim4_pepgrp['Benchmark'] = 'Peptide Grouping'
sim4_combined = pd.concat([sim4_pepid, sim4_pepgrp], ignore_index=True)

sim4_summary = sim4_combined.groupby([
        'Method', 'Benchmark', 'Overlap', 'Direction', 'N_Conditions'
    ])['MCC'].mean().unstack().reset_index()
# sim4_summary
```

###### Robustness Across Experimental Setups¶

Line plots showing MCC vs. number of conditions, with line styles encoding overlap (solid=True, dashed=False) and markers encoding direction (●=random, ■=same).

```
from matplotlib.lines import Line2D

fig, axes = plt.subplots(2, 1, figsize=(5.0, 5.0), sharex=True)
fig.suptitle('Method Robustness Across Experimental Setups', fontsize=10, fontweight='bold', y=0.98)

# Aggregate
agg = sim4_combined.groupby(['Benchmark','Method','Overlap','Direction','N_Conditions'])['MCC'].mean().reset_index()

# Order methods by overall mean performance so best are drawn last (on top)
method_order = agg.groupby('Method')['MCC'].mean().sort_values(ascending=True).index.tolist()

benchmarks = ['Peptide Identification', 'Peptide Grouping']
direction_marker = {'random': 'o', 'same': 's'}
overlap_ls = {True: '-', False: '--'}

# Plot each benchmark in its own small panel
for i, benchmark in enumerate(benchmarks):
    ax = axes[i]
    sub = agg[agg['Benchmark'] == benchmark].copy()
    # draw in method order so higher performers are on top
    for method in method_order:
        sub_m = sub[sub['Method'] == method]
        if sub_m.empty:
            continue
        # within method, draw combinations of overlap/direction
        combos = sorted(sub_m.groupby(['Overlap','Direction']), key=lambda x: (x[0][0], x[0][1]))
        for (overlap, direction), grp in combos:
            grp = grp.sort_values('N_Conditions')
            color = method_palette.get(method, '#333333')
            ls = overlap_ls.get(overlap, '-')
            mk = direction_marker.get(direction, 'o')
            # Plot line with clearly visible marker edge for small sizes
            ax.plot(
                grp['N_Conditions'], grp['MCC'],
                color=color, linestyle=ls, marker=mk,
                markersize=5, markeredgecolor='black', markeredgewidth=0.7,
                linewidth=1.6, alpha=0.95, zorder=3
            )
            # small endpoint label (method short) to aid reading in compact panel
            last_x = grp['N_Conditions'].iloc[-1]
            last_y = grp['MCC'].iloc[-1]
            ax.text(
                last_x + 0.12, last_y,
                method if (overlap, direction) == (True, 'random') else "",
                fontsize=10, fontweight='bold', color=color, va='center'
            )

    # Styling for compact embedding
    ax.set_title(benchmark, fontsize=9, fontweight='bold', loc='left')
    ax.set_ylabel('MCC', fontsize=8)
    ax.set_ylim(-0.05, 0.95)
    ax.grid(axis='y', linestyle=':', linewidth=0.6, alpha=0.5)
    ax.tick_params(axis='both', which='major', labelsize=7)
    sns.despine(ax=ax)

    # # Panel label (A / B) for later placement in multi-panel figures
    # panel_label = 'A' if i == 0 else 'B'
    # ax.text(0.01, 0.98, panel_label, transform=ax.transAxes,
    #         fontsize=9, fontweight='bold', va='top', ha='left',
    #         bbox=dict(boxstyle='round,pad=0.15', facecolor='white', edgecolor='none', alpha=0.8))

# X-axis formatting (shared)
unique_conds = sorted(agg['N_Conditions'].unique())
axes[-1].set_xticks(unique_conds)
axes[-1].set_xlabel('Number of Conditions', fontsize=8)
axes[-1].tick_params(axis='x', labelsize=7)

# Build compact legends (method color, overlap linestyle, direction marker)
methods_present = [m for m in method_order if m in agg['Method'].unique()]
method_handles = [
    Line2D([0], [0], color=method_palette.get(m, 'gray'), lw=2.5, label=m)
    for m in methods_present
]
overlap_handles = [
    Line2D([0], [0], color='gray', lw=1.8, linestyle='-', label='Overlap: True'),
    Line2D([0], [0], color='gray', lw=1.8, linestyle='--', label='Overlap: False'),
]
direction_handles = [
    Line2D([0], [0], color='black', marker='o', linestyle='None', markersize=5, label='Direction: random'),
    Line2D([0], [0], color='black', marker='s', linestyle='None', markersize=5, label='Direction: same'),
]

# # Place method legend in top-left of first panel and style legend for patterns in second
# leg1 = axes[0].legend(handles=method_handles, title='Method', loc='upper left',
#                       fontsize=7, title_fontsize=8, frameon=False, ncol=1)
# axes[0].add_artist(leg1)
axes[1].legend(handles=overlap_handles + direction_handles, loc='upper left',
               fontsize=7, frameon=False, ncol=1)

plt.tight_layout(rect=[0, 0, 1, 0.96])
plt.show()
plots.finalize_plot(
    fig, show=True, save=save_to_folder,
    filename='benchmark_sim4_experimental_setups',
    filepath=figure_path,
    formats=figure_formats, 
    transparent=transparent_bg,
    dpi=figure_dpi,
)
```

**Key Findings — Simulation 4:**

- Higher number of conditions help with peptide identification for linear model based methods like PeCorA and ProteoForge
- When it comes to peptide identification the pattern accross different setups for PeCorA and ProteoForge is extremely similar just with different MCC levels due to the data being missing/imputed version.
- Allowing overlap seems to improve the performance, likely resulting in less predicted groups of peptides, making it easier to identify them correctly.
- Direction of the perturbations (same vs. random) has less effect than other factors
- ProteoForge consistently outperforms other methods across design configurations

---

#### Summary and Figure Assembly¶

##### Generated Figure Panels¶

| Panel | Source | Description | File |
| --- | --- | --- | --- |
| **2A-B** | SWATH-MS | MCC barplots: Identification & Grouping | `benchmark_compact_fig2ab.pdf` |
| **2C** | Sim1 | Imputation impact (vertical bars) | `benchmark_sim1_imputation_effect.pdf` |
| **2D** | Sim2 | Missingness degradation (line plot) | `benchmark_sim2_missingness_effect.pdf` |
| **2E** | Sim3 | Detection sensitivity curves | `benchmark_sim3_perturbation_detection.pdf` |
| **2F** | Sim4 | Experimental design robustness | `benchmark_sim4_experimental_setups.pdf` |

##### Summary of Benchmark Findings¶

1. **Real-world performance (SWATH-MS):** ProteoForge achieves competitive MCC across perturbation scenarios, with strongest relative performance in identification tasks.
2. **Imputation robustness (Sim1):** ProteoForge has minimal performance impact from imputated and robust to appropriately handled missing values.
3. **Missingness tolerance (Sim2):** ~40%-60% missingness threshold before substantial degradation; all methods show similar sensitivity patterns.
4. **Detection sensitivity (Sim3):** ~0.5 log2 perturbation magnitude marks the practical detection threshold; larger effects are consistently detected.
5. **Design complexity (Sim4):** More conditions improve detection; overlapping perturbation patterns offer detection advantages.

> **Note:** Figures exported as PDF vectors for final assembly in Inkscape. Panel labels and minor styling adjustments to be applied during composition.

```
print("Notebook Execution Time:", utils.prettyTimer(utils.getTime() - startTime))
```

```
Notebook Execution Time: 00h:00m:03s
```
