## Supplementary material for "ProteoForge: An Imputation-Aware Framework for Differential Proteoform Discovery in Bottom-Up Proteomics": Analysis notebooks as html render: Notebook S9.html

### Notebook S9

###### Enes Kemal Ergin

#### 2025-12-08 23:00:27

### Supplementary Notebook 9: Data Preparation for Hypoxia Experiment of H358 Cell Lines¶

- **License:** Creative Commons Attribution-NonCommercial 4.0 International License
- **Version:** 0.2
- **Edit Log:**
  - 2025-11-28: Initial version of the notebook
  - 2025-12-08: Revise the whole notebook, ensuring clarity and correctness

---

**Requirements:**

- Raw DIA-NN output: `report.parquet`, `report.pg_matrix.tsv`
- Reference databases: UniProt FASTA, UniProt annotations, iPTMNet, MEROPS
- Prior notebook: None (this is the first notebook in the NSCLC analysis pipeline)

**Data Information:**

- **Cell Line**: H358 (Non-Small Cell Lung Cancer)
- **Conditions**: True Hypoxia (1% O₂), Normoxia (21% O₂)
- **Timepoints**: 48hr, 72hr
- **Replicates**: 4 biological replicates per condition-timepoint
- **Total Samples**: 16 (before QC filtering)

**Purpose:**  
Prepare peptide-level quantitative proteomics data for downstream proteoform analysis:

1. Load and integrate reference databases (FASTA, UniProt, iPTMNet, MEROPS)
2. Parse experimental metadata and quantitative data from DIA-NN output
3. Map peptides to protein sequences with positional information
4. Apply quality control filters (overlap removal, outlier detection)
5. Normalize and impute missing values
6. Calculate protein coverage metrics and save cleaned data

---

#### 01. Setup¶

This section imports required libraries, configures display settings, and defines paths for data and figures.

> **Note:** The HTML rendering of this notebook hides code cells by default. Click the "Code" buttons on the right to expand them.

##### 01.1 Libraries¶

```
# Libraries Used
import re
import os
import sys

import numpy as np # Numerical computing
import pandas as pd # Data manipulation

import seaborn as sns # R-like high-level plots
import matplotlib.pyplot as plt # Python's base plotting 

# home path
sys.path.append('../')
from src import utils, plots
from ProteoForge import annotate, parse 
from ProteoForge import process, coverage
from ProteoForge import normalize, impute
# Initialize the timer
startTime = utils.getTime()
```

##### 01.2 Configure Notebook Settings¶

Configure visualization styles, color palettes, and display options for consistent output formatting throughout the analysis.

```
## Configure Plotting Settings
def_colors = [
    "#139593", "#fca311", "#e54f2a",
    "#c3c3c3", "#555555",
    "#690000", "#5f4a00", "#004549"
]

condition_colors = {
    'True Hypoxia (1% O2) - 48 Hr': '#e9c46a',
    'True Hypoxia (1% O2) - 72 Hr': '#f4a261',
    'Normoxia (21% O2) - 48 Hr': '#ade8f4',
    'Normoxia (21% O2) - 72 Hr': '#0077b6',
    # Won't be used since the manuscript didn't cover these conditions
    # 'Chemical Hypoxia (CoCl2) - 48 Hr': '#c77dff',
    # 'Chemical Hypoxia (CoCl2) - 72 Hr': '#8338ec',
    # 'Oxidative Stress (H2O2) - 48 Hr': '#ff006e',
    # 'Oxidative Stress (H2O2) - 72 Hr': '#fb5607',
}

cv_group_palettes = {
    "<10%":  "#081c15",
    "<25%":  "#1b4332",
    "<50%": "#2d6a4f",
    "<100%": "#52b788",
    ">100%": "#95d5b2",
}

# Set seaborn style
sns.set_theme(
    style="white",
    context="paper",
    palette=def_colors,
    font_scale=1,
    rc={
        "figure.figsize": (6, 4),
        "font.family": "sans-serif",
        "font.sans-serif": ["Arial", "Ubuntu Mono"],
    }
)

# Figure Saving Settings
figure_formats = ["png", "pdf"]
save_to_folder = True
transparent_bg = True
figure_dpi = 300

## Configure dataframe displaying
pd.set_option('display.float_format', lambda x: '%.4f' % x)
pd.set_option('display.max_rows', 50)
pd.set_option('display.max_columns', 50)
pd.set_option('display.width', 500)  # Set a wider display width


## Printing Settings
verbose = True

plots.color_palette( def_colors, save=False )
plots.color_palette( cv_group_palettes, name="CVGroups", save=False )
plots.color_palette( condition_colors, save=False, name='condition', size=2.5)
```


##### 01.3 Data and Result Paths¶

Defines input/output directory structure:

- **Input**: `./data/input/hypoxia/` — DIA-NN output files
- **References**: `./data/input/` — Reference tables such as FASTA and annotations
- **Output**: `./data/cleaned/hypoxia/` — Processed data tables
- **Figures**: `./figures/hypoxia/01-DataPreparation/` — QC visualizations

```
home_path = './'
data_name = "hypoxia"
notebook_name = "01-DataPreparation"
data_path = f"{home_path}data/"
input_path = f"{data_path}input/{data_name}/"
output_path = f"{data_path}cleaned/{data_name}/"
figure_path = f"{home_path}figures/{data_name}/{notebook_name}/"

# Create the output folder if it does not exist
if not os.path.exists(output_path):
    os.makedirs(output_path)

# Create figure folder structure, if needed
if save_to_folder:
    for i in figure_formats:
        cur_folder = figure_path + i + "/"
        if not os.path.exists(cur_folder):
            os.makedirs(cur_folder)
```

---

#### 02. Load Datasets¶

This section loads all required reference databases and experimental data. Multiple data sources are integrated to enable comprehensive proteoform analysis.

##### 02.1 FASTA Reference Sequences¶

The **UniProt human reference proteome** (`UP000005640`) provides canonical protein sequences for peptide mapping. Specifically the `20230316_HumanCr_20806.fasta` file that is from the Hypoxia project folder is loaded and parsed.

```
fasta_data = parse.fasta_to_df(
    fasta_path = f'./data/input/20230316_HumanCr_20806.fasta',
    gene_only = False,
    column_order=['entry', 'geneName', 'proteinDescription', 'sequenceLength', 'molecularWeight_kDa', 'sequence']
).rename(
    columns={
        'entry': 'Protein',
        'geneName': 'Gene',
        'proteinDescription': 'Description',
        'sequenceLength': 'Length',
        'molecularWeight_kDa': 'Weight(kDa)',
        'sequence': 'Sequence'
    }
)
fasta_data.head()
```

```
Processing FASTA file: ./data/input/20230316_HumanCr_20806.fasta
Auto-selected: multiprocessing with 4 processes
Loading FASTA entries...
Processing 26855 entries in chunks of 2828
Processed 26772 entries, skipped 83
  - invalid_sequence: 78
  - length_filter: 5
Generated DataFrame with 26772 rows and 6 columns
```

|  | Protein | Gene | Description | Length | Weight(kDa) | Sequence |
| --- | --- | --- | --- | --- | --- | --- |
| 0 | A0A024B7W1 | None | Genome polyprotein | 3423 | 378.8692 | MKNPKKKSGGFRIVNMLKRGVARVSPFGGLKRLPAGLLLGHGPIRM... |
| 1 | A0A024R1R8 | TMA7B | Translation machinery-associated protein 7B | 64 | 7.0870 | MSSHEGGKKKALKQPKKQAKEMDEEEKAFKQKQKEEQKKLEVLKAK... |
| 2 | A0A024RBG1 | NUDT4B | Diphosphoinositol polyphosphate phosphohydrola... | 181 | 20.4213 | MMKFKPNQTRTYDREGFKKRAACLCFRSEQEDEVLLVSSSRYPDQW... |
| 3 | A0A075B6H7 | IGKV3-7 | Probable non-functional immunoglobulin kappa v... | 116 | 12.7754 | MEAPAQLLFLLLLWLPDTTREIVMTQSPPTLSLSPGERVTLSCRAS... |
| 4 | A0A075B6H8 | IGKV1D-42 | Probable non-functional immunoglobulin kappa v... | 117 | 13.0056 | MDMRVPAQLLGLLLLWLPGVRFDIQMTQSPSFLSASVGDRVSIICW... |

The FASTA reference contains ~20,000 human protein sequences with UniProt accessions, gene names, descriptions, lengths, molecular weights, and full amino acid sequences. These will be used for peptide-to-protein position mapping, and other relevant analyses.

---

##### 02.2 UniProt Annotations¶

Here we prepared the UniProt annotations file to assist with the mapping of any additional protein information to make sense of the proteoforms or PTMs we observe in the data. The `annotate` module from `proteoforge` is used to load and parse the `uniprotkb_Human_AND_model_organism_9606_2025_07_22.txt` file. Additionaly functions makes some annotations more accessible by additional processing steps.

```
# Parse the UniProt features data from the large text file
print("\n--- 📄 Parsing UniProt Features Data ---")
uniprot_data = annotate.preprocess_uniprot(f"./data/input/uniprotkb_Human_AND_model_organism_9606_2025_07_22.txt")
# Expand the uniprot
uniprot_data.columns = ['Protein', 'feature', 'isoform', 'start', 'end', 'description']
# Set up additional columns for further information keeping
uniprot_data['agent'] = ''
uniprot_data['note'] = ''
uniprot_data['group'] = ''
uniprot_data = uniprot_data[['isoform', 'Protein', 'feature', 'start', 'end', 'group', 'agent', 'note', 'description']]
uniprot_data['start'] = uniprot_data['start'].fillna(0).astype(int)
uniprot_data['end'] = uniprot_data['end'].fillna(uniprot_data['start']).astype(int)
uniprot_data.sort_values(by=['Protein', 'start', 'end'], inplace=True)

# Cleaning up and expanding 
print("\n--- 🧹 Cleaning UniProt Features Data ---")
print("Processing the individual features for better info extraction...")
print(" - SITE to Cleavage and Breakpoints...")
uniprot_data = annotate.process_site_features(uniprot_data)
print(" - LIPID features...")
uniprot_data = annotate.process_lipid_features(uniprot_data)
print(" - CROSSLNK features...")
uniprot_data = annotate.process_crosslink_features(uniprot_data)
print(" - CARBOHYD features...")
uniprot_data = annotate.process_carbohydrate_features(uniprot_data)
print(" - DISULFID features...")
uniprot_data = annotate.process_disulfide_features(uniprot_data)
print(" - VAR_SEQ features...")
uniprot_data = annotate.process_varseq_features(uniprot_data)
print(" - MUTAGEN features...")
uniprot_data = annotate.process_mutagen_features(uniprot_data)
print(" - CONFLICT features...")
uniprot_data = annotate.process_conflict_features(uniprot_data)
print(" - MOD_RES features...")
uniprot_data = annotate.process_modres_features(uniprot_data)
print(" - VARIANT features...")
uniprot_data = annotate.process_variant_features(uniprot_data)

# Clean up the empty group, agent, note, and description columns
uniprot_data['group'] = uniprot_data['group'].fillna('')
uniprot_data['agent'] = uniprot_data['agent'].fillna('')
uniprot_data['note'] = uniprot_data['note'].fillna('')
uniprot_data['description'] = uniprot_data['description'].fillna('')

# Display the first few rows of the parsed UniProt features data
print("\nParsed UniProt Features Data Sample:")
print(uniprot_data.head())
print("\nUniProt Features by Occurrence:")
print(uniprot_data['feature'].value_counts())
```

```
--- 📄 Parsing UniProt Features Data ---

--- 🧹 Cleaning UniProt Features Data ---
Processing the individual features for better info extraction...
 - SITE to Cleavage and Breakpoints...
 - LIPID features...
 - CROSSLNK features...
 - CARBOHYD features...
 - DISULFID features...
 - VAR_SEQ features...
 - MUTAGEN features...
 - CONFLICT features...
 - MOD_RES features...
 - VARIANT features...

Parsed UniProt Features Data Sample:
        isoform     Protein   feature  start  end group agent note                                        description
1158389          A0A023HJ61   NON_TER     16   16                                                                    
813523           A0A023I7H2    SIGNAL      1   21                                                                    
813524           A0A023I7H2     CHAIN     22  603                              NADH-ubiquinone oxidoreductase chain 5
813539           A0A023I7H2    DOMAIN     68  118                   NADH-Ubiquinone oxidoreductase (complex I) cha...
813525           A0A023I7H2  TRANSMEM     84  105                                                             Helical

UniProt Features by Occurrence:
feature
DOMAIN        205497
TRANSMEM      174613
NON_TER       167406
COMPBIAS      163351
REGION        131723
STRAND         93341
HELIX          92360
VARIANT        87787
BINDING        66264
MOD_RES        55836
CHAIN          42318
CONFLICT       35418
MUTAGEN        33909
VAR_SEQ        28741
DISULFID       27471
REPEAT         27354
SIGNAL         24658
TURN           23679
TOPO_DOM       20046
CARBOHYD       18542
COILED         13370
ZN_FING         9644
ACT_SITE        7586
CROSSLNK        7126
MOTIF           4160
SITE            2912
INIT_MET        1984
DNA_BIND        1775
LIPID           1209
CLEAVAGE         842
PROPEP           831
TRANSIT          513
PEPTIDE          425
INTRAMEM         422
BREAKPOINT       181
UNSURE            81
NON_STD           45
GLRA2              3
Name: count, dtype: int64
```

UniProt feature annotations have been parsed and processed, including domains, PTM sites, variants, and cleavage sites. Individual feature types (SITE, LIPID, CROSSLNK, CARBOHYD, DISULFID, VAR\_SEQ, MUTAGEN, CONFLICT, MOD\_RES, VARIANT) have been expanded for detailed annotation.

---

##### 02.3 iPTMNet Database¶

**iPTMNet** is a comprehensive database of post-translational modifications (PTMs) integrating data from multiple sources. This database provides additional PTM context that is not available in UniProt alone, both ptms and their scores are downloaded from iPTMNet, parsed, and merged with the uniprot annotations to expand the PTM information.

```
merged_data = annotate.load_and_process_iptmnet(
    ptm_path="./data/input/iPTMnet/ptm.txt",
    score_path="./data/input/iPTMnet/score.txt",
    verbose=verbose,
)
print(f"Shape of Merged PTM Data: {merged_data.shape}")
print("\nMerged PTM Data Sample:")
print(merged_data.head())
print("\nPTM Types by Occurrence:")
print(merged_data['ptm_type'].value_counts())
```

```
--- Loading and Processing ./data/input/iPTMnet/ptm.txt ---
Initial shape: (1055800, 10)
Filtered to Homo sapiens, new shape: (584690, 10)
Shape after cleaning: (584690, 10)

--- Loading and Processing ./data/input/iPTMnet/score.txt ---
Initial shape: (851361, 5)
Shape after cleaning: (851361, 5)

--- Aggregating Scores ---
Aggregated scores into 851,361 unique events.

--- Merging DataFrames and Finalizing Columns ---
Merged scores: 536,531 / 584,690 rows have a non-zero score.
Created 'start' and 'end' columns from 'site'.

Shape of Merged PTM Data: (584690, 13)

Merged PTM Data Sample:
      ptm_type source substrate_protein substrate_gene              organism   site enzyme_protein enzyme_gene          note pmid  score  start   end
0  ACETYLATION   iedb            A8TX70         COL6A5  Homo sapiens (Human)   K652           <NA>              iedb:1797193           1    652   652
1  ACETYLATION   iedb            B2RTY4          MYO9A  Homo sapiens (Human)  K2393           <NA>              iedb:1770492           1   2393  2393
2  ACETYLATION   iedb            O00159          MYO1C  Homo sapiens (Human)   K396           <NA>              iedb:1837008           1    396   396
3  ACETYLATION   iedb            O14672         ADAM10  Homo sapiens (Human)   K659           <NA>              iedb:1772798           1    659   659
4  ACETYLATION   iedb            O15078         CEP290  Homo sapiens (Human)  K1350           <NA>              iedb:1781939           1   1350  1350

PTM Types by Occurrence:
ptm_type
PHOSPHORYLATION      393181
UBIQUITINATION        98115
ACETYLATION           27754
METHYLATION           22555
O-GLYCOSYLATION       22164
N-GLYCOSYLATION        9875
SUMOYLATION            8417
S-NITROSYLATION        2362
MYRISTOYLATION          138
C-GLYCOSYLATION         115
DIHYDROXYLATION           6
FARNESYLATION             3
CARBOXYLATION             2
S-GLYCOSYLATION           2
N-PHOSPHORYLATION         1
Name: count, dtype: int64
```

```
uniprot_data = annotate.add_iptmnet_to_uniprot(uniprot_data, merged_data)
print("\nUpdated UniProt Data with iPTMNet PTM Types:")
print(uniprot_data['feature'].value_counts())
print()
print("The score distribution (3=UniProt and Maybe iPTMNet, highest the better):")
print(uniprot_data['score'].value_counts())
print()
print(uniprot_data.head())
```

```
--- Integrating iPTMnet and UniProt Data ---
Added default score of 3 to 1,573,423 UniProt annotations.
Standardized 584,690 iPTMnet annotations.
Concatenated data. New total annotations: 2,158,113
Final DataFrame sorted by Protein, start, and end positions.

Updated UniProt Data with iPTMNet PTM Types:
feature
MOD_RES       501697
DOMAIN        205497
TRANSMEM      174613
NON_TER       167406
COMPBIAS      163351
REGION        131723
CROSSLNK      113658
STRAND         93341
HELIX          92360
VARIANT        87787
BINDING        66264
CARBOHYD       50698
CHAIN          42318
CONFLICT       35418
MUTAGEN        33909
VAR_SEQ        28741
DISULFID       27471
REPEAT         27354
SIGNAL         24658
TURN           23679
TOPO_DOM       20046
COILED         13370
ZN_FING         9644
ACT_SITE        7586
MOTIF           4160
SITE            2912
INIT_MET        1984
DNA_BIND        1775
LIPID           1350
CLEAVAGE         842
PROPEP           831
TRANSIT          513
PEPTIDE          425
INTRAMEM         422
BREAKPOINT       181
UNSURE            81
NON_STD           45
GLRA2              3
Name: count, dtype: int64

The score distribution (3=UniProt and Maybe iPTMNet, highest the better):
score
3    1597927
1     365128
2     115775
0      48159
4      31124
Name: count, dtype: int64

      Protein   feature  start  end group agent note                                        description  score isoform
0  A0A023HJ61   NON_TER     16   16                                                                          3        
1  A0A023I7H2    SIGNAL      1   21                                                                          3        
2  A0A023I7H2     CHAIN     22  603                              NADH-ubiquinone oxidoreductase chain 5      3        
3  A0A023I7H2    DOMAIN     68  118                   NADH-Ubiquinone oxidoreductase (complex I) cha...      3        
4  A0A023I7H2  TRANSMEM     84  105                                                             Helical      3
```

---

##### 02.4 MEROPS Protease Database¶

**MEROPS** is the comprehensive database of peptidases (proteases) and their substrates. Loading both family annotations and cleavage site details enables identification of potential proteolytic processing events. This enables a lot more cleavage and proteolytic event annotations for the peptides we observe in the data.

```
print("--- Processing parsed merops table ---")
merops_data = annotate.process_merops_data(f"./data/input/merops/mer.tab", verbose=verbose)
print()
print("MEROPS Families by Occurrence:")
print(merops_data['agent'].value_counts())
print()
print(f"Shape of MEROPS Data: {merops_data.shape}")
print(merops_data.head())
```

```
--- Processing parsed merops table ---
--- Processing MEROPS data from ./data/input/merops/mer.tab ---
Processed 67,781 MEROPS cleavage sites for Homo sapiens.

MEROPS Families by Occurrence:
agent
trypsin 1                                                         20792
glutamyl endopeptidase I                                           4096
cathepsin S                                                        3098
cathepsin L                                                        2883
matrix metallopeptidase-3                                          2424
                                                                  ...  
IgA1-specific serine peptidase type 2 ({Haemophilus} sp.)             1
hemorrhagic metallopeptidase b ({Trimeresurus mucrosquamatus})        1
bothrolysin                                                           1
IgA1-specific metallopeptidase                                        1
membrane Pro-Xaa carboxypeptidase                                     1
Name: count, Length: 586, dtype: int64

Shape of MEROPS Data: (67781, 10)
  Protein   feature  start   end                 group                       agent                                               note                                        description  score  isoform
0  P29401  CLEAVAGE    609   609  Proteolytic Cleavage             DESC1 peptidase        N-Ph cleavage at None609 by DESC1 peptidase  MEROPS Cleavage Site; Protease: DESC1 peptidas...      2      NaN
1  O14950  CLEAVAGE    144   144  Proteolytic Cleavage             DESC1 peptidase        N-Ph cleavage at None144 by DESC1 peptidase  MEROPS Cleavage Site; Protease: DESC1 peptidas...      2      NaN
2  P01011  CLEAVAGE    384   384  Proteolytic Cleavage   matrix metallopeptidase-1   Ph cleavage at A384 by matrix metallopeptidase-1  MEROPS Cleavage Site; Protease: matrix metallo...      2      NaN
3  P01009  CLEAVAGE    374   374  Proteolytic Cleavage  matrix metallopeptidase-11  Ph cleavage at A374 by matrix metallopeptidase-11  MEROPS Cleavage Site; Protease: matrix metallo...      2      NaN
4  Q15201  CLEAVAGE   1218  1218  Proteolytic Cleavage     procollagen C-peptidase    Ph cleavage at A1218 by procollagen C-peptidase  MEROPS Cleavage Site; Protease: procollagen C-...      2      NaN
```

```
uniprot_data = annotate.add_merops_to_uniprot(uniprot_data, merops_data, verbose=verbose)
print("\nUpdated UniProt Data with iPTMNet PTM Types:")
print(uniprot_data['feature'].value_counts())
print()
print("The score distribution (3=UniProt and Maybe iPTMNet, highest the better):")
print(uniprot_data['score'].value_counts())
print()
print(uniprot_data.head())
```

```
--- Adding MEROPS data to UniProt DataFrame ---
Added 67,781 MEROPS annotations. New total: 2,225,894
Final DataFrame sorted by Protein, start, and end positions.

Updated UniProt Data with iPTMNet PTM Types:
feature
MOD_RES       501697
DOMAIN        205497
TRANSMEM      174613
NON_TER       167406
COMPBIAS      163351
REGION        131723
CROSSLNK      113658
STRAND         93341
HELIX          92360
VARIANT        87787
CLEAVAGE       68623
BINDING        66264
CARBOHYD       50698
CHAIN          42318
CONFLICT       35418
MUTAGEN        33909
VAR_SEQ        28741
DISULFID       27471
REPEAT         27354
SIGNAL         24658
TURN           23679
TOPO_DOM       20046
COILED         13370
ZN_FING         9644
ACT_SITE        7586
MOTIF           4160
SITE            2912
INIT_MET        1984
DNA_BIND        1775
LIPID           1350
PROPEP           831
TRANSIT          513
PEPTIDE          425
INTRAMEM         422
BREAKPOINT       181
UNSURE            81
NON_STD           45
GLRA2              3
Name: count, dtype: Int64

The score distribution (3=UniProt and Maybe iPTMNet, highest the better):
score
3    1597927
1     365128
2     183556
0      48159
4      31124
Name: count, dtype: int64

  isoform     Protein   feature  start  end group agent note                                        description  score
0          A0A023HJ61   NON_TER     16   16                                                                          3
1          A0A023I7H2    SIGNAL      1   21                                                                          3
2          A0A023I7H2     CHAIN     22  603                              NADH-ubiquinone oxidoreductase chain 5      3
3          A0A023I7H2    DOMAIN     68  118                   NADH-Ubiquinone oxidoreductase (complex I) cha...      3
4          A0A023I7H2  TRANSMEM     84  105                                                             Helical      3
```

##### 02.5 Sample Metadata¶

The **experimental metadata** file contains sample-to-condition mappings, including treatment groups (hypoxia vs. normoxia), timepoints (48hr, 72hr), and replicate information. This is generated from the `report.pg_matrix.tsv` file by analyzing the column names to extract condition and replicate details and overall standardizing the sample names for consistency.

```
# Load the raw metadata file
print("Loading sample metadata (from pg_matrix column names)...")
tmp = pd.read_csv(f'{input_path}report.pg_matrix.tsv', sep='\t')
meta = tmp.columns[4:].to_list()
# Build a meta dataframe
meta = pd.DataFrame(
    data=meta,
    columns=['filename']
)
# --- Clean and Standardize Metadata ---
print("\n🧹 Cleaning and standardizing metadata columns...")

meta['Run'] = meta['filename'].str.split(r'\\').str[-1].str.replace('.d', '')
# Extract condition identifier
def get_condition(x):
    if '1%' in x:
        return 'True Hypoxia (1% O2)'
    elif 'CoCl' in x:
        return 'Chemical Hypoxia (CoCl2)'
    elif 'H2O2' in x:
        return 'Oxidative Stress (H2O2)'
    else:
        return 'Normoxia (21% O2)'
meta['Condition'] = meta['Run'].map(get_condition)
# Extract the replicate number (special rule for 1% O2)
meta['Replicate'] = meta['Run'].str.split('_').apply(lambda x: x[2] if '1%' in x[1] else x[1][-1])
meta['Duration'] = meta['Run'].str.split('_').str[-5] # last 5th element from the end
meta['Colname'] = meta['Run'].str.split('_').map(
    # Simplify the colname based on the condition
    lambda x: f"{x[1]}_{x[3]}_{x[2]}" if '1%' in x[1] else f"{x[1][:-1]}_{x[2]}_{x[1][-1]}"
)
# --- Finalize DataFrame ---
print("Selecting final columns and renaming for clarity...")
print("\n✅ Metadata processing complete.")
print(f"Final DataFrame shape: {meta.shape}")
print(f"Final columns: {meta.columns.tolist()}")
print()

# Sort the metadata DataFrame for better readability
meta = meta.sort_values(
    ['Condition', 'Duration', 'Replicate'], 
    ascending=[False, True, True]
).reset_index(drop=True)

# Keep only the relevant conditions for the study
relevant_conditions = [
    'True Hypoxia (1% O2)',
    'Normoxia (21% O2)',
]
print(f"Filtering metadata to keep only relevant conditions: {relevant_conditions}")
meta = meta[meta['Condition'].isin(relevant_conditions)].reset_index(drop=True)
print(f"Filtered DataFrame shape: {meta.shape}")
meta
```

```
Loading sample metadata (from pg_matrix column names)...

🧹 Cleaning and standardizing metadata columns...
Selecting final columns and renaming for clarity...

✅ Metadata processing complete.
Final DataFrame shape: (32, 6)
Final columns: ['filename', 'Run', 'Condition', 'Replicate', 'Duration', 'Colname']

Filtering metadata to keep only relevant conditions: ['True Hypoxia (1% O2)', 'Normoxia (21% O2)']
Filtered DataFrame shape: (16, 6)
```

|  | filename | Run | Condition | Replicate | Duration | Colname |
| --- | --- | --- | --- | --- | --- | --- |
| 0 | D:\Data\Tam\RevisionLungDIA\H358\H358\_1%\_1\_48\_... | H358\_1%\_1\_48\_DIA\_BB4\_1\_5432 | True Hypoxia (1% O2) | 1 | 48 | 1%\_48\_1 |
| 1 | D:\Data\Tam\RevisionLungDIA\H358\H358\_1%\_2\_48\_... | H358\_1%\_2\_48\_DIA\_BC4\_1\_5440 | True Hypoxia (1% O2) | 2 | 48 | 1%\_48\_2 |
| 2 | D:\Data\Tam\RevisionLungDIA\H358\H358\_1%\_3\_48\_... | H358\_1%\_3\_48\_DIA\_BD4\_1\_5448 | True Hypoxia (1% O2) | 3 | 48 | 1%\_48\_3 |
| 3 | D:\Data\Tam\RevisionLungDIA\H358\H358\_1%\_4\_48\_... | H358\_1%\_4\_48\_DIA\_BE4\_1\_5456 | True Hypoxia (1% O2) | 4 | 48 | 1%\_48\_4 |
| 4 | D:\Data\Tam\RevisionLungDIA\H358\H358\_1%\_1\_72\_... | H358\_1%\_1\_72\_DIA\_BB8\_1\_5436 | True Hypoxia (1% O2) | 1 | 72 | 1%\_72\_1 |
| 5 | D:\Data\Tam\RevisionLungDIA\H358\H358\_1%\_2\_72\_... | H358\_1%\_2\_72\_DIA\_BC8\_1\_5444 | True Hypoxia (1% O2) | 2 | 72 | 1%\_72\_2 |
| 6 | D:\Data\Tam\RevisionLungDIA\H358\H358\_1%\_3\_72\_... | H358\_1%\_3\_72\_DIA\_BD8\_1\_5452 | True Hypoxia (1% O2) | 3 | 72 | 1%\_72\_3 |
| 7 | D:\Data\Tam\RevisionLungDIA\H358\H358\_1%\_4\_72\_... | H358\_1%\_4\_72\_DIA\_BE8\_1\_5460 | True Hypoxia (1% O2) | 4 | 72 | 1%\_72\_4 |
| 8 | D:\Data\Tam\RevisionLungDIA\H358\H358\_C1\_48\_DI... | H358\_C1\_48\_DIA\_BB1\_1\_5429 | Normoxia (21% O2) | 1 | 48 | C\_48\_1 |
| 9 | D:\Data\Tam\RevisionLungDIA\H358\H358\_C2\_48\_DI... | H358\_C2\_48\_DIA\_BC1\_1\_5437 | Normoxia (21% O2) | 2 | 48 | C\_48\_2 |
| 10 | D:\Data\Tam\RevisionLungDIA\H358\H358\_C3\_48\_DI... | H358\_C3\_48\_DIA\_BD1\_1\_5445 | Normoxia (21% O2) | 3 | 48 | C\_48\_3 |
| 11 | D:\Data\Tam\RevisionLungDIA\H358\H358\_C4\_48\_DI... | H358\_C4\_48\_DIA\_BE1\_1\_5453 | Normoxia (21% O2) | 4 | 48 | C\_48\_4 |
| 12 | D:\Data\Tam\RevisionLungDIA\H358\H358\_C1\_72\_DI... | H358\_C1\_72\_DIA\_BB5\_1\_5433 | Normoxia (21% O2) | 1 | 72 | C\_72\_1 |
| 13 | D:\Data\Tam\RevisionLungDIA\H358\H358\_C2\_72\_DI... | H358\_C2\_72\_DIA\_BC5\_1\_5441 | Normoxia (21% O2) | 2 | 72 | C\_72\_2 |
| 14 | D:\Data\Tam\RevisionLungDIA\H358\H358\_C3\_72\_DI... | H358\_C3\_72\_DIA\_BD5\_1\_5449 | Normoxia (21% O2) | 3 | 72 | C\_72\_3 |
| 15 | D:\Data\Tam\RevisionLungDIA\H358\H358\_C4\_72\_DI... | H358\_C4\_72\_DIA\_BE5\_1\_5457 | Normoxia (21% O2) | 4 | 72 | C\_72\_4 |

The metadata defines the experimental design with 16 samples across two conditions (True Hypoxia and Normoxia) at two timepoints (48hr and 72hr). Each condition-timepoint combination has 4 biological replicates. filename, Run, Condition, Duration, Replicate, and Colname is created to be in the metadata. Colname is shortened and standardized version of the sample names that will be used throughout the analysis.

---

##### 02.6 Quantitative Proteomics Data¶

The **quantitative data** from DIA-NN output contains peptide-level intensity values across all samples. The `report.parquet` file is loaded to obtain peptide sequences, protein accessions, and intensity measurements for each of the 16 samples. This reports precursor-level data that will be aggregated to peptide-level and mapped to proteins.

```
print("Loading precursors long data...")
# Load and preprocess hypoxia experiment data efficiently
data = pd.read_parquet(f'{input_path}report.parquet')
print(f"Data shape (long format): {data.shape}")
print(f"Columns: {data.columns.tolist()}")

print("\n--- 📊 Data Preparation: Pivoting and Cleaning ---")
print("Pivoting the data to wide format (use sum of multiple precursors)...")
print(" - Using normalized precursor intensities for quantification.")
print(" - Doesn't have modified sequences using stripped sequences as peptide identifiers.")
print(" - Protein IDs may contain isoforms or multiple entries, which will be handled later.")
data_wide = (
    data.pivot_table(
        index=['Protein.Ids', 'Stripped.Sequence'],
        columns='Run',
        values='Precursor.Normalised',
        aggfunc='sum'
    )
    .reset_index()
    .rename(columns={
        'Protein.Ids': 'Protein',
        'Stripped.Sequence': 'Peptide'
    })
 )

# Display summary information
print(f"Data shape (wide format): {data_wide.shape}")
print(f"Columns: {data_wide.columns.tolist()}")

print("\n--- 🧬 Protein ID Representation ---")
# Check if the protein contains '-' indicating it has isoforms
has_isoform = data_wide['Protein'].str.contains('-').any()
has_multiples = data_wide['Protein'].str.contains(';').any()
# Friendly print statements to explain the protein ID representation
if has_isoform:
    print("The dataset contains protein IDs with isoforms (e.g., 'P12345-2'). These represent specific variants of a protein.")
else:
    print("The dataset does not contain protein IDs with isoforms.")

if has_multiples:
    print("The dataset contains protein IDs with multiple entries (e.g., 'P12345; P67890'). These represent proteins that may have multiple forms or related proteins.")
    print()
    print("Will explode these entries to create a more detailed DataFrame.")
    data_wide = process.explode_aligned(data_wide, ['Protein'], sep=';', verbose=verbose)
    print()
    print(f"After exploding, the dataset now has {len(data_wide)} rows and {len(data_wide.columns)} columns.")
else:
    print("The dataset does not contain protein IDs with multiple entries.")

print(f"Values below 2 should be set as NaN, they might be artifacts of quantification.")
print(f"Number of missing values in the DataFrame before cleanup: {data_wide.isnull().sum().sum()}")
# Identify numeric columns in the DataFrame
numeric_columns = data_wide.select_dtypes(include=[np.number]).columns
data_wide[numeric_columns] = data_wide[numeric_columns].mask(
    data_wide[numeric_columns] < 2
)
print(f"Number of missing values in the DataFrame after cleanup: {data_wide.isnull().sum().sum()}")

data_wide = data_wide.set_index(['Protein', 'Peptide'])[meta['Run']]
print("Set cleaned column names from metadata")
data_wide.columns = meta['Colname']
data_wide = data_wide.sort_index()
data_wide.head()
```

```
Loading precursors long data...
Data shape (long format): (2988832, 67)
Columns: ['Run.Index', 'Run', 'Channel', 'Precursor.Id', 'Precursor.Channels.Group', 'Modified.Sequence', 'Stripped.Sequence', 'Precursor.Charge', 'Precursor.Lib.Index', 'Proteotypic', 'Precursor.Mz', 'Protein.Ids', 'Protein.Group', 'Protein.Names', 'Genes', 'RT', 'iRT', 'Predicted.RT', 'Predicted.iRT', 'IM', 'iIM', 'Predicted.IM', 'Predicted.iIM', 'Precursor.Quantity', 'Precursor.Normalised', 'Ms1.Area', 'Ms1.Normalised', 'Ms1.Apex.Area', 'Normalisation.Factor', 'Quantity.Quality', 'Empirical.Quality', 'Normalisation.Noise', 'Ms1.Profile.Corr', 'Averagine', 'Evidence', 'Mass.Evidence', 'Ms1.Total.Signal.Before', 'Ms1.Total.Signal.After', 'RT.Start', 'RT.Stop', 'FWHM', 'PG.Normalised', 'PG.MaxLFQ', 'Genes.Normalised', 'Genes.MaxLFQ', 'Genes.MaxLFQ.Unique', 'PG.MaxLFQ.Quality', 'Q.Value', 'PEP', 'Global.Q.Value', 'Lib.Q.Value', 'Peptidoform.Q.Value', 'Global.Peptidoform.Q.Value', 'Lib.Peptidoform.Q.Value', 'PTM.Q.Value', 'PTM.Site.Confidence', 'Site.Occupancy.Probabilities', 'Protein.Sites', 'Lib.PTM.Site.Confidence', 'Translated.Q.Value', 'Channel.Q.Value', 'PG.Q.Value', 'PG.PEP', 'GG.Q.Value', 'Protein.Q.Value', 'Global.PG.Q.Value', 'Lib.PG.Q.Value']

--- 📊 Data Preparation: Pivoting and Cleaning ---
Pivoting the data to wide format (use sum of multiple precursors)...
 - Using normalized precursor intensities for quantification.
 - Doesn't have modified sequences using stripped sequences as peptide identifiers.
 - Protein IDs may contain isoforms or multiple entries, which will be handled later.
Data shape (wide format): (104861, 34)
Columns: ['Protein', 'Peptide', 'H358_1%_1_48_DIA_BB4_1_5432', 'H358_1%_1_72_DIA_BB8_1_5436', 'H358_1%_2_48_DIA_BC4_1_5440', 'H358_1%_2_72_DIA_BC8_1_5444', 'H358_1%_3_48_DIA_BD4_1_5448', 'H358_1%_3_72_DIA_BD8_1_5452', 'H358_1%_4_48_DIA_BE4_1_5456', 'H358_1%_4_72_DIA_BE8_1_5460', 'H358_C1_48_DIA_BB1_1_5429', 'H358_C1_72_DIA_BB5_1_5433', 'H358_C2_48_DIA_BC1_1_5437', 'H358_C2_72_DIA_BC5_1_5441', 'H358_C3_48_DIA_BD1_1_5445', 'H358_C3_72_DIA_BD5_1_5449', 'H358_C4_48_DIA_BE1_1_5453', 'H358_C4_72_DIA_BE5_1_5457', 'H358_CoCl1_48_DIA_BB2_1_5430', 'H358_CoCl1_72_DIA_BB6_1_5434', 'H358_CoCl2_48_DIA_BC2_1_5438', 'H358_CoCl2_72_DIA_BC6_1_5442', 'H358_CoCl3_48_DIA_BD2_1_5446', 'H358_CoCl3_72_DIA_BD6_1_5450', 'H358_CoCl4_48_DIA_BE2_1_5454', 'H358_CoCl4_72_DIA_BE6_1_5458', 'H358_H2O21_48_DIA_BB3_1_5431', 'H358_H2O21_72_DIA_BB7_1_5435', 'H358_H2O22_48_DIA_BC3_1_5439', 'H358_H2O22_72_DIA_BC7_1_5443', 'H358_H2O23_48_DIA_BD3_1_5447', 'H358_H2O23_72_DIA_BD7_1_5451', 'H358_H2O24_48_DIA_BE3_1_5455', 'H358_H2O24_72_DIA_BE7_1_5459']

--- 🧬 Protein ID Representation ---
The dataset does not contain protein IDs with isoforms.
The dataset contains protein IDs with multiple entries (e.g., 'P12345; P67890'). These represent proteins that may have multiple forms or related proteins.

Will explode these entries to create a more detailed DataFrame.
🚀 Starting aligned explosion for columns: ['Protein']
Original DataFrame shape: (104861, 34)
Final DataFrame shape: (112248, 34)
✅ Transformation complete in 1.0261 seconds.

After exploding, the dataset now has 112248 rows and 34 columns.
Values below 2 should be set as NaN, they might be artifacts of quantification.
Number of missing values in the DataFrame before cleanup: 610106
Number of missing values in the DataFrame after cleanup: 610106
Set cleaned column names from metadata
```

|  | Colname | 1%\_48\_1 | 1%\_48\_2 | 1%\_48\_3 | 1%\_48\_4 | 1%\_72\_1 | 1%\_72\_2 | 1%\_72\_3 | 1%\_72\_4 | C\_48\_1 | C\_48\_2 | C\_48\_3 | C\_48\_4 | C\_72\_1 | C\_72\_2 | C\_72\_3 | C\_72\_4 |
| --- | --- | --- | --- | --- | --- | --- | --- | --- | --- | --- | --- | --- | --- | --- | --- | --- | --- |
| Protein | Peptide |  |  |  |  |  |  |  |  |  |  |  |  |  |  |  |  |
| A0A024B7W1 | LAILMGATFAEMNTGGDVAHLALIAAFK | 2394.7788 | 2653.8262 | 4374.3252 | 4618.3652 | NaN | 3560.3354 | 5652.7100 | NaN | 3485.5161 | 2289.0959 | 4028.9673 | 4263.6060 | NaN | NaN | 3707.3254 | NaN |
| A0A024RBG1 | EVYEEAGVK | 56351.1875 | 63097.6523 | 49170.1328 | 55968.3789 | NaN | 41009.9844 | 51193.7461 | 54011.7969 | 62612.5977 | 63970.3516 | 37216.3477 | 44602.7344 | 58097.2539 | 73113.1484 | 65346.1016 | 56678.7344 |
|  | EVYEEAGVKGK | NaN | NaN | 4063.8176 | NaN | NaN | NaN | NaN | NaN | NaN | NaN | NaN | NaN | NaN | NaN | NaN | NaN |
|  | EWFKVEDAIK | 1051.2585 | NaN | 4486.0947 | 5052.0957 | NaN | 7127.1855 | 6379.5830 | 3082.0085 | NaN | 2105.7913 | 4573.0874 | 5685.3628 | 2177.9431 | NaN | 6287.4355 | 6404.9004 |
|  | LLGIFEQNQDR | 16599.4629 | 13334.4561 | 12436.4561 | 16408.7402 | NaN | 16807.1602 | 13422.2295 | 11881.2012 | 14877.3135 | 11963.4307 | 13533.5078 | 18642.8730 | 16161.4492 | 18011.0938 | 16386.9121 | 13474.7529 |

**Findings:** The DIA-NN quantitative data has been loaded and pivoted to wide format with:

- Peptide sequences (stripped) as row identifiers
- Sample intensities as columns (normalized precursor values)
- Multi-protein assignments exploded to individual rows
- Low-value artifacts (<2) set to missing

Now the all data tables is ready for the next steps of sequence mapping and quality control.

---

#### 03. Sequence Mapping¶

This section maps detected peptide sequences to their parent protein sequences to establish positional information and calculate sequence coverage metrics.

```
# Build a dictionary to match condition_colors keys: 'Condition - Duration Hr'
condition_to_samples = {}
for condition in meta['Condition'].unique():
    for duration in sorted(meta.loc[meta['Condition'] == condition, 'Duration'].unique()):
        key = f"{condition} - {duration} Hr"
        samples = meta.loc[(meta['Condition'] == condition) & (meta['Duration'] == duration), 'Colname'].tolist()
        condition_to_samples[key] = samples

# samples_to_condition mapping
samples_to_condition = {sample: condition for condition, samples in condition_to_samples.items() for sample in samples}
# samples_to_colors mapping
samples_to_colors = {sample: condition_colors[condition] for sample, condition in samples_to_condition.items()}
```

##### 03.1 Protein Overlap Analysis¶

First, all identified proteins compared against the fasta reference to ensure they exist in the database and to retrieve their sequences for peptide mapping. While might not be necessary, we also examine the intersection, and differences between identified proteins and reference proteome (from UniProt FASTA).

```
# Extract protein information and reset indices
info_data = data_wide.reset_index()[['Protein', 'Peptide']]
data_wide = data_wide.reset_index(drop=True)
print(f"✓ Info data extracted: {info_data.shape[0]:,} peptides")
print()

# =============================================================================
# PROTEIN REFERENCE COMPARISON
# =============================================================================

print("🧬 PROTEIN OVERLAP ANALYSIS")
print("=" * 50)

# Calculate protein sets and overlap
fasta_set = set(fasta_data['Protein'].tolist())
identified_set = set(info_data['Protein'].tolist())

overlap = len(fasta_set & identified_set)
only_reference = len(fasta_set - identified_set)
only_identified = len(identified_set - fasta_set)
total_reference = len(fasta_set)

# Create summary statistics
print(f"📋 PROTEIN INVENTORY:")
print(f"  • Reference proteins:     {total_reference:,}")
print(f"  • Identified proteins:    {len(identified_set):,}")
print(f"  • Overlap:               {overlap:,} ({overlap/total_reference*100:.1f}% coverage)")
print(f"  • Unmatched:             {only_identified:,}")
print()

# Create clean horizontal bar chart
fig, ax = plt.subplots(1, 1, figsize=(10, 4))

categories = ['Reference Only', 'Overlap\n(Identified)', 'Dataset Only\n(Unmatched)']
values = [only_reference, overlap, only_identified]
colors = ['#c3c3c3', '#139593', '#e54f2a']

bars = ax.barh(categories, values, color=colors, alpha=0.8)

# Add value labels on bars
for i, (bar, value) in enumerate(zip(bars, values)):
    width = bar.get_width()
    percentage = (value / total_reference * 100) if i != 2 else (value / len(identified_set) * 100)
    ax.text(width + max(values)*0.01, bar.get_y() + bar.get_height()/2, 
            f'{value:,}\n({percentage:.1f}%)', 
            ha='left', va='center', fontweight='bold')

ax.set_xlabel('Number of Proteins', fontweight='bold')
ax.set_title('Protein Dataset Coverage Analysis', fontweight='bold', pad=15)
ax.spines['top'].set_visible(False)
ax.spines['right'].set_visible(False)
plt.tight_layout()
plots.finalize_plot( 
    fig, show=True, save=save_to_folder, 
    filename="fasta_overlap",
    # Arguments
    filepath=figure_path,
    formats=figure_formats,
    transparent=transparent_bg,
    dpi=figure_dpi
)

# Handle unmatched proteins
if only_identified > 0:
    unmatched_proteins = identified_set - fasta_set
    print("⚠️  UNMATCHED PROTEINS ANALYSIS")
    print("-" * 35)
    print(f"• {only_identified:,} proteins not in reference (may be isoforms, contaminants, or outdated IDs)")
    
    if only_identified <= 5:
        print("• Complete list:")
        for protein in sorted(unmatched_proteins):
            print(f"  - {protein}")
    else:
        print(f"• Sample (showing 5 of {only_identified:,}):")
        for protein in sorted(list(unmatched_proteins))[:5]:
            print(f"  - {protein}")
    
    # =============================================================================
    # DATA CLEANUP
    # =============================================================================
    
    print("� CLEANING DATA - REMOVING UNMATCHED PROTEINS")
    print("-" * 50)
    print(f"• Removing {only_identified:,} unmatched proteins from dataset")
    print(f"• Before cleanup: {info_data.shape[0]:,} peptides")
    
    # Remove unmatched proteins from info_data
    info_data = info_data[~info_data["Protein"].isin(unmatched_proteins)]
    print(f"• After cleanup:  {info_data.shape[0]:,} peptides")
    
    # Update main data to match filtered info_data
    data_wide = data_wide.loc[info_data.index].reset_index(drop=True)
    info_data = info_data.reset_index(drop=True)
    
    # Verify data integrity
    if info_data.shape[0] == data_wide.shape[0]:
        print("✓ Data shapes match after cleanup")
    else:
        print("❌ Warning: Data shape mismatch after cleanup")
        
else:
    print("✅ Perfect coverage: All identified proteins found in reference")
```

```
✓ Info data extracted: 112,248 peptides

🧬 PROTEIN OVERLAP ANALYSIS
==================================================
📋 PROTEIN INVENTORY:
  • Reference proteins:     26,772
  • Identified proteins:    11,042
  • Overlap:               11,035 (41.2% coverage)
  • Unmatched:             7
```

```
⚠️  UNMATCHED PROTEINS ANALYSIS
-----------------------------------
• 7 proteins not in reference (may be isoforms, contaminants, or outdated IDs)
• Sample (showing 5 of 7):
  - O90368
  - P05878
  - P27283
  - Q306W8
  - Q5XXP4
� CLEANING DATA - REMOVING UNMATCHED PROTEINS
--------------------------------------------------
• Removing 7 unmatched proteins from dataset
• Before cleanup: 112,248 peptides
• After cleanup:  112,240 peptides
✓ Data shapes match after cleanup
```

There are 7 unmatched proteins (not in reference) and have been removed to ensure valid peptide-to-sequence mapping. The bar chart visualizes the comparison between identified and reference proteins.

---

##### 03.2 Position Mapping¶

The **sequence position mapping** locates each peptide within its parent protein sequence. This establishes the start and end positions required for coverage visualization and proteoform analysis.

```
# =============================================================================
# MERGE FASTA DATA WITH INFO DATA
# =============================================================================
print("🔗 MERGING FASTA DATA WITH INFO DATA")
print("=" * 50)

info_data = info_data.merge(
    fasta_data[['Protein', 'Gene', 'Description', 'Length', 'Weight(kDa)', 'Sequence']],
    on='Protein',
    how='left',
    validate='m:1'
)
print(f"✓ Merged info data: {info_data.shape[0]:,} peptides with protein details")
print()

# =============================================================================
# MAP PEPTIDE POSITIONS TO PROTEIN SEQUENCES
# =============================================================================

new_info_data = process.map_peptide_to_protein_positions(
    data=info_data,
    sequence_col='Sequence',
    peptide_col='Peptide',
    protein_col='Protein',
    # modified_col='Modified Peptide',
    # Outputs
    peptide_start_col='peptide_start',
    peptide_end_col='peptide_end',
    unique_id_col='unique_id',
    total_occurrences_col='total_occurrences',
    occurrence_index_col='occurrence_index',
    original_index_col='original_index',
    # Processing Options
    expand_multiple=True,
    add_unique_id=True,
    add_occurrence_info=True,
    position_ordered=True,
    verbose=verbose
)

print()
print(f'Original Info Data shape: {info_data.shape}')
print(f'Original Quant Data shape: {data_wide.shape}')

print()
print("Reindexing the quantitative data to match the new info data...")
info_data, data_wide = process.reindex_quantitative_data(
    info_data=new_info_data,
    quan_data=data_wide,
    original_index_col='original_index',
    remove_unmatched=True,
    position_col='peptide_start',
    verbose=verbose
)

print()

print(f"Final Info Data shape: {info_data.shape}")
print(f"Final Quant Data shape: {data_wide.shape}")
```

```
🔗 MERGING FASTA DATA WITH INFO DATA
==================================================
✓ Merged info data: 112,240 peptides with protein details

============================================================
PEPTIDE-TO-PROTEIN POSITION MAPPING
============================================================
Input dataset: 112,240 rows
Detected columns:
  - Protein sequence: 'Sequence'
  - Peptide sequence: 'Peptide'
  - Protein ID: 'Protein'
Processing options:
  - Expand multiple occurrences: True
  - Add unique IDs: True
  - Add occurrence info: True
  - Position ordered: True
  - Preserve original indices: True

Finding peptide positions...
POSITION FINDING RESULTS:
  - Total peptide entries: 112,240
  - Successfully matched: 112,240 (100.000%)
  - Unmatched peptides: 0 (0.000%)

OCCURRENCE DETAILS:
  - Single occurrences: 111,800
  - Multiple occurrences: 440
  - Total positions found: 113,512
  - Max occurrences per peptide: 22
  - Average occurrences (matched): 1.01

EXPANSION INFO:
  - Expected output rows (with expansion): 113,512
    → 113,512 positioned rows + 0 unmatched rows
  - Row expansion factor: 1.01x

Expanding multiple occurrences...
Adding unique proteoform identifiers...
FINAL PROCESSING SUMMARY:
  - Input rows: 112,240
  - Output rows: 113,512

POSITIONING RESULTS (EXPANDED MODE):
  - Original peptides matched: 112,240 (100.000%)
  - Original peptides unmatched: 0 (0.000%)
  - Total positioned peptides (expanded): 113,512
  - Total unpositioned rows: 0
  - Expected positioned peptides: 113,512
  - Expected unpositioned rows: 0
  - Expected total output: 113,512
  - Row expansion factor: 1.01x

INSIGHTS:
  - Total unique positions mapped: 113,512
  - Average positions per mapped peptide: 1.01
  - Peptides with >5 occurrences: 77 (max: 22)
============================================================

Ordering peptides by positions using columns: ['Protein', 'peptide_start', 'peptide_end']

Original Info Data shape: (112240, 7)
Original Quant Data shape: (112240, 16)

Reindexing the quantitative data to match the new info data...
============================================================
REINDEXING AND FILTERING QUANTITATIVE DATA
============================================================
Info data shape: (113512, 13)
Quantitative data shape: (112240, 16)
Remove unmatched peptides: True
Position column: 'peptide_start'
Filtered info data shape: (113512, 13)
Original index column: 'original_index'
Original indices range: 0 to 112239

REINDEXING RESULTS:
  - Total output rows: 113,512
  - Matched rows (copied quantitative data): 113,512
  - Remaining unmatched rows (NaN values): 0
  - Output shape: (113512, 16)
  - Data fill rate: 83.4% (1,514,557/1,816,192 values)
  - Row expansion factor: 1.01x
  - Max duplications per original row: 22
============================================================


Final Info Data shape: (113512, 12)
Final Quant Data shape: (113512, 16)
```

The `info_data` is expanded with peptide positions (start, end). Peptides with multiple occurrences within a protein are expanded with unique identifiers, while this might inflate the number of peptides in some repeating peptides (eg. chain-like peptides), it is done to ensure we are not randomly mapping it to one place and present them all. With this additional expansion, the quantitative data has been reindexed to match the positional information.

---

#### 04. Data Cleaning¶

This section applies quality control filters and normalization procedures to prepare the data for statistical analysis. The cleaning workflow ensures robust and reproducible downstream results. Additionally, visualizations are generated to assess data quality at each step.

##### 04.1 Overlapping Peptide Grouping¶

There could be many reasons why there are highly overlapping peptides for the same protein identified. While exact reason for them might be hard to pin down, to reduce the complexity and redundancy in the data, we group highly overlapping peptides together. This helps in simplifying the dataset and focusing on unique peptide signals for downstream analysis. The maximum allowed difference for overlap detection is set as 3 amino acids. This ensures if the peptides are overlapping within 3 amino acids, they will be grouped together.

> (Note: This is not if two peptide overlap by 3 amino-acids, this means the difference between their start and end positions is less than or equal to 3 amino-acids)

```
from ProteoForge import peptide_grouper as pg
prv_unq_peps = info_data.shape[0]
prv_unq_prts = info_data['Protein'].nunique()
info_data, data_wide = pg.process_peptide_overlaps(
    info_df=info_data,
    quan_df=data_wide,
    protein_col='Protein',
    startpos_col='peptide_start',
    endpos_col='peptide_end',
    max_diff=3,
    verbose=verbose
)

# Check if the indices match
if info_data.shape[0] == data_wide.shape[0]:
    print("✓ Data shapes match after peptide grouping")
if info_data.index.equals(data_wide.index):
    print("✓ Data indices contents and orders match after peptide grouping")

# Reset indices for consistency
info_data = info_data.reset_index(drop=True)
data_wide = data_wide.reset_index(drop=True)

upd_unq_peps = info_data.shape[0]
upd_unq_prts = info_data['Protein'].nunique()
print()
print(f"Peptide grouping reduced peptides from {prv_unq_peps:,} to {upd_unq_peps:,} ({(1 - upd_unq_peps/prv_unq_peps)*100:.1f}% reduction)")
print(f"Number of proteins remained the same: {prv_unq_prts:,} proteins")
print()
```

```
Processing 113512 peptides across 11035 proteins
Converting object columns to numeric...
✅ Completed in 12.0s: 102179 representative peptides from 102179 groups
✓ Data shapes match after peptide grouping
✓ Data indices contents and orders match after peptide grouping

Peptide grouping reduced peptides from 113,512 to 102,179 (10.0% reduction)
Number of proteins remained the same: 11,035 proteins
```

Overlapping peptides (within 3 positions) have been merged to reduce redundancy. This consolidation reduces the peptide count while maintaining protein coverage, ensuring cleaner downstream proteoform analysis. The both `info_data` and `data_wide` (quant data) have been updated to reflect these groupings.

---

##### 04.2 Protein Coverage Calculation¶

**Protein coverage** quantifies the proportion of each protein sequence covered by detected peptides. The `coverage.calculate_protein_coverage()` function computes coverage percentages and trace patterns for quality assessment. The smart calculations allows some overlap considerations to avoid overestimating coverage due to overlapping peptides.

```
print("📏 CALCULATING PROTEIN COVERAGE & TRACE")
info_data = coverage.calculate_protein_coverage(
    info_data, 
    protein_col='Protein',
    peptide_col='Peptide',
    seq_len_col='Length',
    start_pos_col='peptide_start',
    end_pos_col='peptide_end',
    unique_id_col='unique_id',
    gap=0,
    validate_input=True,
    n_jobs='auto', 
    verbose=verbose
)
```

```
📏 CALCULATING PROTEIN COVERAGE & TRACE
Processing 11035 proteins with 102179 total peptides
Auto-selected: multiprocessing with 8 processes
Starting multiprocessing coverage calculation...
Multiprocessing completed in 3.58s
Applying results to DataFrame...
Building trace mappings (large dataset)...
Finalizing results...
Average protein coverage: 20.6%
Generated DataFrame with 102179 rows and 15 columns
```

Protein sequence coverage has been calculated for all proteins (Avg 20.6%). Coverage percentages (Cov%) indicate the fraction of each protein's amino acid sequence represented by detected peptides. This information is added to `info_data` for downstream filtering and analysis.

---

###### Proteomic Coverage Visualization¶

The data-wide protein coverage plots visualize the relationship between protein length, peptide count, and sequence coverage. This high-level overview helps put in perspective how well proteins are represented in the dataset. This is not crucial but as a nice-to-have visualization for understanding the dataset better.

```
# 1. Aggregate and transform the data
# This step remains the same as your original code
plot_data = info_data.groupby("Protein").agg(
    Length=("Length", "first"),
    PeptideCount=("Peptide", "count"),
    CovPerc=("Cov%", "first")
).reset_index()
plot_data["Length"] = np.log10(plot_data["Length"])
plot_data["PeptideCount"] = np.log10(plot_data["PeptideCount"])


# 2. Set up the figure and subplots
# Create a figure with 1 row and 2 columns of subplots
fig, axes = plt.subplots(1, 2, figsize=(12, 5.5), constrained_layout=True)
fig.suptitle(
    "Proteomic Quality Assessment: Linking Protein Length with Peptide Count and Sequence Coverage",
    fontsize=16,
    fontweight="bold"
)

# --- Subplot A: Protein Length vs. Peptide Count ---
ax1 = axes[0]
sns.scatterplot(
    ax=ax1,
    data=plot_data,
    x="Length",
    y="PeptideCount",
    hue="CovPerc",
    size="CovPerc",
    palette="viridis_r", # Reversed Viridis is great for this
    sizes=(10, 200),
    alpha=0.7,
    edgecolor="black",
    linewidth=0.4,
)
ax1.set_title("Length vs. Peptide Count by Coverage", fontsize=12, style="italic")
ax1.set_xlabel("$Log_{10}$(Protein Length)", fontsize=11)
ax1.set_ylabel("$Log_{10}$(Peptide Count)", fontsize=11)
ax1.grid(True, which="both", linestyle="--", linewidth=0.5, alpha=0.7)
ax1.legend(
    title="Coverage %", loc="upper left", frameon=False, 
    facecolor="#FFFFFFCC", edgecolor="black", 
    fontsize=9, title_fontsize=10
)
ax1.text(-0.1, 1.05, 'A', transform=ax1.transAxes, size=18, weight='bold') # Subplot label


# --- Subplot B: Protein Coverage Distribution ---
ax2 = axes[1]
# Use a complementary color from the start of the viridis palette
hist_color = "#440154"
sns.histplot(
    ax=ax2,
    data=plot_data,
    x="CovPerc",
    bins=15,
    kde=True,
    color=hist_color,
    alpha=0.75,
    edgecolor="white",
    linewidth=0.8,
)
ax2.set_title("Protein Coverage Distribution", fontsize=12, style="italic")
ax2.set_xlabel("Sequence Coverage (%)", fontsize=11)
ax2.set_ylabel("Protein Frequency", fontsize=11)
ax2.grid(True, which="both", linestyle="--", linewidth=0.5, alpha=0.7)
ax2.text(-0.1, 1.05, 'B', transform=ax2.transAxes, size=18, weight='bold') # Subplot label


# 3. Final adjustments
# Despine removes the top and right spines for a cleaner look
sns.despine(fig=fig, top=True, right=True, left=True, bottom=False)

plots.finalize_plot( 
    fig, show=True, save=save_to_folder, 
    filename='proteinLength_peptideCount_coverage',
    filepath=figure_path,
    formats=figure_formats,
    transparent=transparent_bg,
    dpi=figure_dpi
)

# Print the number of proteins with higher than 75% coverage
high_coverage_count = plot_data[plot_data['CovPerc'] > 75].shape[0]
print(f"Number of proteins with coverage > 75%: {high_coverage_count}")
print(f"Number of proteins with coverage > 90%: {plot_data[plot_data['CovPerc'] > 90].shape[0]}")
high_coverage_proteins = plot_data[plot_data['CovPerc'] > 90].sort_values(by='CovPerc', ascending=False)['Protein'].tolist()
print()
for protein in high_coverage_proteins:
    gene = info_data.loc[info_data['Protein'] == protein, 'Gene'].iloc[0]
    description = info_data.loc[info_data['Protein'] == protein, 'Description'].iloc[0]
    prtLength = info_data.loc[info_data['Protein'] == protein, 'Length'].iloc[0]
    print(f"{protein:<10} ({gene:<5}) - {description:<50} - Length: {prtLength:>4} aa")
```

```
Number of proteins with coverage > 75%: 33
Number of proteins with coverage > 90%: 3

P61604     (HSPE1) - 10 kDa heat shock protein, mitochondrial           - Length:  102 aa
P08727     (KRT19) - Keratin, type I cytoskeletal 19                    - Length:  400 aa
O76070     (SNCG ) - Gamma-synuclein                                    - Length:  127 aa
```

This figure, provides a scatter plot and histogram to visualize protein coverage statistics. In scatter plot the log10 transformed protein lengths and peptide counts are plotted, with point size and color indicating sequence coverage percentage. The histogram displays the distribution of sequence coverage across all proteins. The histogram is almost always right-skewed, indicating that many proteins have low coverage while a few have high coverage. Unless the dataset is extremely well-covered, this pattern is expected. The numbers of proteins with more than 75% and 90% coverage is printed above, and 90% > ones are generally very few in typical datasets, and they are individually printed out for reference.

```
# Check if info_data has the unique_id column
info_data = info_data.set_index(['Protein', 'unique_id'], drop=False)
data_wide.index = info_data.index
# Ensure the columns are numeric
data_wide = data_wide.apply(pd.to_numeric, errors='coerce')
```

---

##### 04.3 Filter Proteins by Peptide Count¶

**Minimum peptide requirement**: ProteoForge requires at least 4 peptides per protein for reliable proteoform detection. Proteins with fewer peptides are excluded from downstream analysis. While this might look too stringent, having multiple peptides per proteoform is crucial to have an accurate outlier detection and clustering. Going lower than this threshold can compromise the reliability of proteoform assignments.

```
tmp = info_data.reset_index(drop=True).copy()
prot_size = tmp.groupby('Protein')['Peptide'].nunique().sort_values(ascending=False)
# Print summary statistics
total_proteins = len(prot_size)
num_proteins_ge4 = (prot_size >= 4).sum()
num_proteins_lt4 = (prot_size < 4).sum()

print(f"Total proteins detected: {total_proteins}")
print(f"Proteins with ≥4 peptides: {num_proteins_ge4}")
print(f"Proteins with <4 peptides: {num_proteins_lt4}")

print("\nMinimum of 4 peptides per protein is required for ProteoForge, only proteins with ≥4 peptides will be used for downstream analysis.")
print(f"This would retain {num_proteins_ge4} proteins and exclude {num_proteins_lt4} proteins.")

# Enhanced visualization of protein size distribution
fig, ax = plt.subplots(figsize=(10, 5))
sns.histplot(np.log10(prot_size), bins=30, kde=True, color="#139593", ax=ax, edgecolor='black', alpha=0.8)

# Add vertical line at log10(4)
ax.axvline(np.log10(4), color='red', linestyle='--', linewidth=2, label='Peptide Size = 4')

# Add text box summarizing the filtering effect
ax.text(
    0.75, ax.get_ylim()[1]*0.8, 
    f"Total proteins: {total_proteins}\nRetained (≥4 peptides): {num_proteins_ge4}\nExcluded (<4 peptides): {num_proteins_lt4}", 
    fontsize=11, color='navy', bbox=dict(facecolor='white', alpha=0.7, boxstyle='round,pad=0.5')
)

ax.set_ylabel('Frequency', fontsize=13)
ax.set_xlabel('Log10(Number of Peptides per Protein)', fontsize=13)
ax.set_title('Distribution of Protein Sizes (Peptides per Protein)', fontsize=15, fontweight='bold')

ax.legend(fontsize=12)
ax.grid(which='both', linestyle='--', linewidth=0.5, color='lightgray', alpha=0.75)

plt.tight_layout()
plots.finalize_plot( 
    fig, show=True, save=save_to_folder, 
    filename='proteinSize_distribution',
    filepath=figure_path,
    formats=figure_formats,
    transparent=transparent_bg,
    dpi=figure_dpi
)

print()
print("Updating the main data by keeping only proteins with at least 4 peptides...")
print(f"- Data shape before filtering: {data_wide.shape}")
print(f"- Proteins before filtering: {data_wide.index.get_level_values('Protein').nunique()}")
print(f"- Peptides before filtering: {data_wide.index.get_level_values('unique_id').nunique()}")
data_wide = data_wide.loc[data_wide.index.get_level_values('Protein').isin(prot_size[prot_size >= 4].index)]
print("Filtering complete.")
print(f"- Data shape after filtering: {data_wide.shape}")
print(f"- Proteins after filtering: {data_wide.index.get_level_values('Protein').nunique()}")
print(f"- Peptides after filtering: {data_wide.index.get_level_values('unique_id').nunique()}")
```

```
Total proteins detected: 11035
Proteins with ≥4 peptides: 7161
Proteins with <4 peptides: 3874

Minimum of 4 peptides per protein is required for ProteoForge, only proteins with ≥4 peptides will be used for downstream analysis.
This would retain 7161 proteins and exclude 3874 proteins.
```

```
Updating the main data by keeping only proteins with at least 4 peptides...
- Data shape before filtering: (102179, 16)
- Proteins before filtering: 11035
- Peptides before filtering: 102179
Filtering complete.
- Data shape after filtering: (95369, 16)
- Proteins after filtering: 7161
- Peptides after filtering: 95369
```

The protein size distribution shows the number of peptides quantified per protein. The vertical red dashed line marks the ≥4 peptide threshold required for ProteoForge analysis. Proteins below this threshold lack sufficient number of peptide requirements and have been filtered out, which is approximately 35% of the proteins in this dataset.

> Note: Again this might look too stringent and aggressive filtering, and in some special cases, one might consider lowering this threshold to 3 peptides, but it is not recommended.

---

##### 04.4 Outlier Sample Detection¶

This is a standard way of detecting if any sample behaves abnormally compared to the rest of the dataset, specifically within their biological groups. Such outlier samples can skew the analysis, cause lower confidence by increasing variability, and lead to misleading conclusions. Therefore, identifying and removing these outliers is crucial for maintaining data integrity. There can be many reasons for a sample to behave abnormally, including technical issues during sample preparation, instrument errors during data acquisition, or biological variability. Identifying these outliers helps ensure that the analysis focuses on reliable and representative data.

The method that we have been using is to examine correlation, number of quantified samples, and coefficient of variation within biological groups. Within each and between all these metrics if a sample is behaving outside of allowed IQR ranges or has low correlation with other samples in the same group, it is flagged as an outlier. If it is flagged multiple times (here is 3/7), it is considered an outlier and removed from the dataset.

**Outlier detection** identifies samples with anomalous intensity distributions using pairwise correlation, quantification ratios, and coefficient of variation metrics. Samples flagged as outliers across multiple metrics are removed to ensure data quality.

```
metric_weights = {  # Weights for the metrics used in pairwise combined score calculation
    'correlation': 1.0,
    'quantification': 1.0,
    'cv': 1.0
}
minOutlier_occurrance = 3 # minimum number of outlier occurrences to consider a sample behaves like an outlier

conditions_to_check = list(condition_to_samples.keys())  # List of conditions to check for outlier detection

print("\nCalculating Pairwise Combined Scores...")
print(" - Using metric weights:", metric_weights)
print(" - Using Spearman correlation, quantification as ratio, and CV as 1 - CV")

all_outlier_samples, combined_group_reports = utils.identify_outlier_samples(
    data=data_wide,
    sample_groups=condition_to_samples,
    analysis_func=utils.run_metric_combination_analysis,
    min_outlier_occurrence=minOutlier_occurrance,
    verbose=False
)
print(f"\nComplete list of unique outlier samples to remove (after min_outlier_occurrence >= {minOutlier_occurrance})\n - {all_outlier_samples}")
print()

plot_data = data_wide.reset_index().melt(
    id_vars=['Protein', 'unique_id'],
    var_name='Sample',
    value_name='Intensity'
).dropna().assign(log2_Intensity=lambda x: np.log2(x['Intensity']))
fig, ax = plt.subplots(figsize=(12, 4))
sns.boxplot(
    x='Sample', y='log2_Intensity', hue='Sample', data=plot_data, ax=ax, 
    palette=samples_to_colors,
    fliersize=0.5,
    linewidth=0.5,
    notch=False,
    showmeans=True,
    meanprops={"marker": "o", "markerfacecolor": "white", "markeredgecolor": "black"},
    medianprops={"color": "white", "linewidth": 2.5},
    whiskerprops={"color": "black", "linewidth": 1},
    capprops={"color": "black", "linewidth": 0},
    legend=False
)
y_min = 0
y_max = plot_data['log2_Intensity'].max() + 1
ax.set_ylim(y_min, y_max)
ax.tick_params(axis='x', labelrotation=90)
ax.set_title('Boxplot of Log2 Intensities Across Samples', fontsize=16, fontweight='bold')
ax.set_xlabel('Sample', fontsize=14)
ax.set_ylabel('Log2(Intensity)', fontsize=14)
ax.grid(which='both', linestyle='--', linewidth=0.5, color='lightgray', alpha=0.7)

import matplotlib.patches as mpatches

# Highlight background for outlier samples
for i, label in enumerate(ax.get_xticklabels()):
    sample_label = label.get_text()
    if sample_label in all_outlier_samples:
        # Add a rectangle spanning the y-range at the x position
        ax.add_patch(
            mpatches.Rectangle(
                (i - 0.5, y_min),  # x position, y position
                1,                 # width (one boxplot)
                y_max - y_min,     # height
                color='#ffcccc',   # light red
                zorder=0,          # behind the boxplot
                alpha=0.5
            )
        )
# Add legend for outlier highlight
outlier_patch = mpatches.Patch(color='#ffcccc', label='Identified Outlier Sample', alpha=0.5)
ax.legend(handles=[outlier_patch], loc='upper right', fontsize=12)
plt.tight_layout()
plots.finalize_plot( 
    fig, show=True, save=save_to_folder, 
    filename='boxplot_log2Intensity_with_outliers',
    filepath=figure_path,
    formats=figure_formats,
    transparent=transparent_bg,
    dpi=figure_dpi
)
# Remove the identified outlier samples from the main data
print("\nRemoving identified outlier samples from the main data...")
print(f"- Samples to remove: {all_outlier_samples}")
print(f"- Data shape before removal: {data_wide.shape}")
cleaned_data = data_wide.drop(columns=all_outlier_samples)
print(f"- Data shape after removal: {cleaned_data.shape}")
print("Outlier removal complete.")
```

```
Calculating Pairwise Combined Scores...
 - Using metric weights: {'correlation': 1.0, 'quantification': 1.0, 'cv': 1.0}
 - Using Spearman correlation, quantification as ratio, and CV as 1 - CV

Complete list of unique outlier samples to remove (after min_outlier_occurrence >= 3)
 - ['1%_72_1', 'C_72_4']
```

```
Removing identified outlier samples from the main data...
- Samples to remove: ['1%_72_1', 'C_72_4']
- Data shape before removal: (95369, 16)
- Data shape after removal: (95369, 14)
Outlier removal complete.
```

```
meta = meta[~meta['Colname'].isin(all_outlier_samples)].reset_index(drop=True)
# Build a dictionary to match condition_colors keys: 'Condition - Duration Hr'
condition_to_samples = {}
for condition in meta['Condition'].unique():
    for duration in sorted(meta.loc[meta['Condition'] == condition, 'Duration'].unique()):
        key = f"{condition} - {duration} Hr"
        samples = meta.loc[(meta['Condition'] == condition) & (meta['Duration'] == duration), 'Colname'].tolist()
        condition_to_samples[key] = samples

# samples_to_condition mapping
samples_to_condition = {sample: condition for condition, samples in condition_to_samples.items() for sample in samples}
# samples_to_colors mapping
samples_to_colors = {sample: condition_colors[condition] for sample, condition in samples_to_condition.items()}
```

Outlier detection identified 2 samples with anomalous intensity distributions based on pairwise correlation, quantification ratios, and CV metrics. Samples flagged as outliers (≥3 metric violations) are highlighted in the boxplot with light red background and removed from further analysis. The metadata and sample mappings are updated accordingly.

> Note1: This is one thing that is highly dataset dependent. In some datasets, there might be no outliers at all, while in some datasets there might be multiple outliers. The key is to have a robust method to identify them based on multiple metrics rather than relying on a single criterion.
>
> Note2: I wasn't sure if I should do this step, since I already have low N, but wanted to keep the data consistent with my other analyses.

##### 04.5 Filter by Minimum Quantification¶

Removes peptides with insufficient quantification across conditions. Entries must have at least 2 valid measurements in at least one condition to be retained for analysis. This ensures that only peptides with adequate data for statistical comparisons are included in downstream analyses. This enables we remove peptides that are mostly missing across all samples or conditions, which would not provide reliable information for proteoform detection, and don't want to include them to later impute with low-values which would not really add any value to the analysis.

```
min_limit = 2 # Minimum number of peptides or intensity to consider a valid quantification
metric_type = 'Count'  # 'Count' for number of peptides, 'Intensity' for summed intensity
filter_type = 'or'   # 'or' for any condition, 'and' for all conditions

# --- Highlighted Print Statements ---
print("="*80)
print(f"🔬 CONFIGURATION: Filtering by '{metric_type}' with a threshold of ≥ {min_limit}.")
print(f"Logic: Keep entries if {'ANY' if filter_type == 'or' else 'ALL'} condition(s) meet the threshold.")
print("="*80)

# --- Calculating Statistics ---
print("\n📊 Calculating quantification statistics...")
all_stats = {}
for cond, cols in condition_to_samples.items():
    quantified_count = cleaned_data[cols].notna().sum(axis=1)
    total_cols_in_condition = len(cols)
    quantified_pctl = (quantified_count / total_cols_in_condition) * 100
     # Add the results to our dictionary
    all_stats[f'{cond} Quantified Count'] = quantified_count
    all_stats[f'{cond} Quantified Percentage'] = quantified_pctl.round(2) # Rounding for clarity

stats_df = pd.DataFrame(all_stats)
print(f"Total entries identified: {len(stats_df):,}")

# --- Filtering Logic and Results ---
print("\n" + "-"*80)
print("🔍 APPLYING FILTER...")

if filter_type == 'or':
    # Keep rows where at least one condition meets the limit
    idx_to_keep = (stats_df.loc[:, stats_df.columns.str.contains(metric_type)] >= min_limit).any(axis=1)
elif filter_type == 'and':
    # Keep rows where all conditions meet the limit
    idx_to_keep = (stats_df.loc[:, stats_df.columns.str.contains(metric_type)] >= min_limit).all(axis=1)
else:
    raise ValueError("Invalid filter_type. Use 'or' or 'and'.")

# --- Summarize Results ---
total_count = len(stats_df)
kept_count = idx_to_keep.sum()
percentage_kept = (kept_count / total_count) * 100

print("\n" + "="*80)
print("✅ FILTERING COMPLETE: SUMMARY")
print("="*80)
print(f"{'Initial Entries:':<20} {total_count:>10,}")
print(f"{'Entries Kept:':<20} {kept_count:>10,}")
print(f"{'Entries Discarded:':<20} {total_count - kept_count:>10,}")
print(f"{'Percentage Kept:':<20} {percentage_kept:>9.2f}%")
print("="*80)
```

```
================================================================================
🔬 CONFIGURATION: Filtering by 'Count' with a threshold of ≥ 2.
Logic: Keep entries if ANY condition(s) meet the threshold.
================================================================================

📊 Calculating quantification statistics...
Total entries identified: 95,369

--------------------------------------------------------------------------------
🔍 APPLYING FILTER...

================================================================================
✅ FILTERING COMPLETE: SUMMARY
================================================================================
Initial Entries:         95,369
Entries Kept:            93,309
Entries Discarded:        2,060
Percentage Kept:         97.84%
================================================================================
```

The quantification filter retains peptides with at least 2 valid intensity measurements in at least one experimental condition. This kept close to 98% of the data, ensuring only the sufficient data points for reliable statistical analysis while maintaining coverage across the dataset.

---

##### 04.6 Median Centering Normalization¶

**Median centering** corrects for systematic intensity differences between samples by aligning sample medians while preserving the original intensity magnitude scale.

```
centered_data = normalize.by_median_centering(
    df=cleaned_data,
    rescale_to_original_magnitude=True,
    condition_map=None, # Pass dictionary if want to apply condition-specific centering
    verbose=verbose
)
# plots.normalized_density_comparison(
#     before_df=data_wide_cleaned,
#     after_df=centered_data,
#     condition_map=condition_to_samples,
#     color_palette=condition_colors,
#     log_transform=True,
#     title="\nDensity of Samples",
#     show=True,
#     save=save_to_folder,
#     filename="DensityPlot_Normalization_Comparison",
#     fileformats=figure_formats,
#     filepath=figure_path,
#     transparent=transparent_bg,
#     dpi=figure_dpi,
# )
```

```
Applying Median Centering...
  - Applying global median centering...
    - Rescaling by a global factor of: 16384.00
  - Data after median centering has shape: (95369, 14)
  - Missing values (Before/After): (145644/145644)
```

Median centering is standard, non-invasive normalization technique to adjust for systematic intensity differences between samples. It aligns sample medians while preserving the original intensity magnitude scale, ensuring comparability across samples without distorting relative abundance relationships. One can do within biological groups as well, but here since we have only 2 conditions with low N, we are doing it across all samples.

---

##### 04.7 Missing Value Imputation¶

**Multi-strategy imputation** addresses different missingness patterns:

1. **Amputate**: Set sparse values (1 of 4 replicates) to complete missing
   - *Rationale*: Single missing values in a condition with 4 replicates are likely random dropouts (MCAR).
2. **Fill dense**: Mean-fill single missing values per condition
   - *Rationale*: When only one value is missing in a condition, it is reasonable to impute it with the mean of the other replicates, assuming the missingness is random.
3. **Downshift**: Low-value imputation for completely missing features (MNAR)
   - *Rationale*: Features missing in all replicates of a condition are likely below detection limits (MNAR) and are imputed with low values to reflect this.
4. **k-NN**: Neighbor-based imputation for remaining missing values
   - *Rationale*: For remaining missing values that do not fit the above patterns, k-NN imputation leverages similarity between peptides to estimate missing intensities based on neighboring peptide profiles.

###### Examining the Downshifted Low-Value Imputation Profiles¶

Before determining some of the parameters for downshifted low-value imputation, it is important to understand the low-percentile intensity distributions of the dataset. The two version of figures below examine the 0.1 percentile and 0.05 percentile intensity distributions across all samples combined. Then the downshift magnitude 1.5, and 5 are examined with respectively. This allows me to visualize how much overlap between the downshifted low values and the actual low-intensity values in the dataset. Based on these visualizations, I can make an informed decision on the downshift magnitude to use for imputation.

```
print("\nChecking the downshifted low-value imputation parameter's impact...")
print(" The magnitude is taking log2 into account if data is already log2, is_log2 needs to be set True.")
shift_mag = 1.5
low_percentile=0.10
print(f" - Using the default values shift magnitude: {shift_mag}, low percentile: {low_percentile}")

plots.downshift_effect(
    df=centered_data,
    cond_dict=condition_to_samples,
    shift_magnitude=shift_mag,
    low_percentile=low_percentile,
    is_log2=False,
    show=True,
    save=save_to_folder,
    filename="Data_DownshiftedImputation_Effect_default",
    fileformats=figure_formats,
    filepath=figure_path,
    transparent=transparent_bg,
    dpi=figure_dpi,
)
shift_mag = 5
low_percentile = 0.05
print(f"\n - Adjusting the shift magnitude to {shift_mag} and low percentile to {low_percentile} for downshifted imputation.")
plots.downshift_effect(
    df=centered_data,
    cond_dict=condition_to_samples,
    shift_magnitude=shift_mag,
    low_percentile=low_percentile,
    is_log2=False,
    show=True,
    save=save_to_folder,
    filename="Data_DownshiftedImputation_Effect_adjusted",
    fileformats=figure_formats,
    filepath=figure_path,
    transparent=transparent_bg,
    dpi=figure_dpi,
)
print(" - Accepting the adjusted parameters for downshifted imputation.")
```

```
Checking the downshifted low-value imputation parameter's impact...
 The magnitude is taking log2 into account if data is already log2, is_log2 needs to be set True.
 - Using the default values shift magnitude: 1.5, low percentile: 0.1
```

```
 - Adjusting the shift magnitude to 5 and low percentile to 0.05 for downshifted imputation.
```

```
 - Accepting the adjusted parameters for downshifted imputation.
```

The downshift parameter visualization compares default (shift=1.5, percentile=0.10) and adjusted (shift=5, percentile=0.05) settings. The adjusted parameters provide greater separation between imputed low-values and the real data distribution, reducing potential false positive abundance changes from imputed values. The parameters chosen here (shift=5, percentile=0.05) provide a clearer separation between imputed low-values and the actual data distribution. This reduces the risk of imputed values overlapping with real low-intensity measurements, which could lead to false positives in downstream analyses.

###### Applying Imputation Strategies¶

The multi-strategy imputation is provide in the `impute` module and requires a list of dictionaries defining the strategies to apply in order. Here the strategies defined above are implemented sequentially to address different missingness patterns in the data.

```
imputation_pipeline_scheme = [
    { # First is the remove sparsely quantified features to set complete missing for low-value imputation 
        'method': 'amputate',
        'params': { 'min_quantified': 2 } # below 2 meaning if 1 quantified out of 4 replicates, then set to all missing
    },
    # I generally don't use this step, but it can be useful for some datasets
    { # Second step is generally to fill the sparse missingness with features' mean or median 
        'method': 'fill_dense', 
        'params': {'max_missing': 1, 'strategy': 'mean'}
    },
    # This is the best for complete or mostly missing features
    { # Adds downshifted low-values to completely missing (1.0) or mostly missing (< 1.0) features
        'method': 'downshift',
        'params': {'missingness_threshold': 1.0, 'shift_magnitude': 5, 'low_percentile': 0.05}
    },
    { # Final step is to use k-NN imputation to fill the remaining missing values
        'method': 'knn',
        'params': {'n_neighbors': 5}
    }
]
```

```
imputation_pipeline_scheme = [
    { # First is the remove sparsely quantified features to set complete missing for low-value imputation 
        'method': 'amputate',
        'params': { 'min_quantified': 2 } # below 2 meaning if 1 quantified out of 4 replicates, then set to all missing
    },
    # I generally don't use this step, but it can be useful for some datasets
    { # Second step is generally to fill the sparse missingness with features' mean or median 
        'method': 'fill_dense', 
        'params': {'max_missing': 1, 'strategy': 'mean'}
    },
    # This is the best for complete or mostly missing features
    { # Adds downshifted low-values to completely missing (1.0) or mostly missing (< 1.0) features
        'method': 'downshift',
        'params': {'missingness_threshold': 1.0, 'shift_magnitude': 5, 'low_percentile': 0.05}
    },
    { # Final step is to use k-NN imputation to fill the remaining missing values
        'method': 'knn',
        'params': {'n_neighbors': 5}
    }
]

imputed_data = impute.run_imputation_pipeline(
    data=centered_data,
    cond_dict=condition_to_samples,
    scheme=imputation_pipeline_scheme,
    is_log2=False,  
    return_log2=False,
    verbose=verbose
)

# print("\nVisualizating the distribution of imputed and real values:")
# # if number of columns higher than 20, then use 
# if len(imputed_data.columns) > 20:
#     print(" - Visualize per condition, as there are too many samples to visualize per sample")
#     plots.imputation_distribution_per_condition(
#         original_data=centered_data,
#         imputed_data=imputed_data,
#         title="\nDistribution of Imputed and Real Values per Condition",
#         is_log2=False,
#         cond_dict=condition_to_samples,
#         show=True,
#         save=save_to_folder,
#         filename="Imputation_Distribution_PerCondition",
#         fileformats=figure_formats,
#         filepath=figure_path,
#         transparent=transparent_bg,
#         dpi=figure_dpi,
#     )
# else:
#     print(" - Visualize per sample, as there are not too many samples to visualize per sample")
#     plots.imputation_distribution_per_sample(
#         original_data=centered_data,
#         imputed_data=imputed_data,
#         title="\nDistribution of Imputed and Real Values per Sample",
#         is_log2=False,
#         show=True,
#         save=save_to_folder,
#         filename="Imputation_Distribution_PerSample",
#         fileformats=figure_formats,
#         filepath=figure_path,
#         transparent=transparent_bg,
#         dpi=figure_dpi,
#     )
```

```
====== Starting Imputation Pipeline ======
Initial missing values: 145644

--- Running Step: Sparse Feature Amputation ---
  Parameters: min_quantified=2
  Condition 'True Hypoxia (1% O2) - 48 Hr': Amputating 3017 features with < 2 quantified values.
  Condition 'True Hypoxia (1% O2) - 72 Hr': Amputating 5478 features with < 2 quantified values.
  Condition 'Normoxia (21% O2) - 48 Hr': Amputating 2870 features with < 2 quantified values.
  Condition 'Normoxia (21% O2) - 72 Hr': Amputating 6090 features with < 2 quantified values.
  Summary: Created 17455 new missing values by amputation.
Total missing values after step 1 ('amputate'): 163099

--- Running Step: Dense Feature Filling ---
  Parameters: max_missing=1, strategy='mean'
  Condition 'True Hypoxia (1% O2) - 48 Hr': Filling 9961 features with 1 to 1 missing values.
  Condition 'True Hypoxia (1% O2) - 72 Hr': Filling 9163 features with 1 to 1 missing values.
  Condition 'Normoxia (21% O2) - 48 Hr': Filling 9457 features with 1 to 1 missing values.
  Condition 'Normoxia (21% O2) - 72 Hr': Filling 10031 features with 1 to 1 missing values.
  Summary: Filled 38612 missing values.
Total missing values after step 2 ('fill_dense'): 124487

--- Running Step: Downshifted Imputation ---
  Parameters: missingness_threshold=1.0, shift_magnitude=5, low_percentile=0.05
  Condition 'True Hypoxia (1% O2) - 48 Hr': Imputing 5109 features with >= 4 missing values.
  Condition 'True Hypoxia (1% O2) - 72 Hr': Imputing 9166 features with >= 3 missing values.
  Condition 'Normoxia (21% O2) - 48 Hr': Imputing 4877 features with >= 4 missing values.
  Condition 'Normoxia (21% O2) - 72 Hr': Imputing 9349 features with >= 3 missing values.
  Summary: Imputed 95489 missing values.
Total missing values after step 3 ('downshift'): 28998

--- Running Step: k-NN Imputation ---
  Parameters: n_neighbors=5
  Condition 'True Hypoxia (1% O2) - 48 Hr': Applying k-NN to 14412 remaining missing values.
  Condition 'Normoxia (21% O2) - 48 Hr': Applying k-NN to 14586 remaining missing values.
  Summary: Imputed 28998 missing values.
Total missing values after step 4 ('knn'): 0

====== Imputation Pipeline Finished ======
```

The multi-strategy imputation pipeline addresses different missingness patterns sequentially: (1) amputate sparse values, (2) mean-fill single missing values per condition, (3) downshift completely missing values, and (4) k-NN imputation for remaining gaps. This approach treats Missing Completely At Random (MCAR), Missing At Random (MAR), and Missing Not At Random (MNAR) patterns appropriately.

You can see the imputation results summary above, which shows how many values were imputed by each strategy. The majority of imputations were done using the downshift method for completely missing features, followed by k-NN for remaining gaps. This indicates that many peptides had missing values likely due to being below detection limits in certain conditions.

---

##### 04.8 Coefficient of Variation Analysis¶

**Coefficient of Variation (CV)** measures within-condition reproducibility. Lower CV values indicate more consistent measurements across biological replicates, with CV <25% generally considered acceptable for quantitative proteomics.

```
print("\nCalculating Coefficient of Variation (CV) from proteins and group by condition...")

cv_data = pd.DataFrame(index = imputed_data.index)
# Calculate the coefficient of variation per biological sample
for k, v in condition_to_samples.items():
    cv_data[k] = utils.cv_numpy(
        imputed_data[v].values,
        axis=1,
        ignore_nan=True,
        format="percent"
    )

plot_data = utils.create_cv_group_plot_data(
    cv_data=cv_data,
    cv_group_palettes=cv_group_palettes
).dropna(subset=['CV'])

plots.cv_comparison(
    plot_data=plot_data,
    cv_group_palettes=cv_group_palettes,
    condition_colors=condition_colors,
    main_title="\nCoefficient of Variation (CV) Comparison",
    show=True,
    save=save_to_folder,
    filename="CV_Comparison",
    fileformats=figure_formats,
    filepath=figure_path,
    transparent=transparent_bg,
    dpi=figure_dpi,
)
```

```
Calculating Coefficient of Variation (CV) from proteins and group by condition...
```

The CV comparison visualizes measurement reproducibility across experimental conditions. While the table overshadows the boxplots, we can see the mean and median CV values are generally below 25%, indicating good reproducibility. Some outliers with higher CVs exist, which is common in proteomics data. Additionally the proportion of proteins with lower than 10% CV is generally 15-25%, which is a good sign of high-quality data.

---

#### 05. Complete Final Checks and Save the Data¶

##### 05.1 Final Checks on the Data¶

Before we finish and finalize the cleaned data tables to be saved for downstream analysis, we perform some final checks to ensure data integrity and consistency. This includes verifying that all filtering and normalization steps have been correctly applied, checking for any remaining missing values, and confirming that the metadata accurately reflects the cleaned dataset. Once these checks are complete, the cleaned data tables will be saved to the specified output directory for use in subsequent proteoform analysis steps.

```
print("Making sure the info_data is aligned with the imputed data (latest version)...")
info_data = info_data.loc[imputed_data.index].reset_index(drop=True)
print()

print("Establish protein data, will be making use in later steps.")
protein_data = info_data.groupby('Protein').agg(
    Gene=('Gene', 'first'),
    PeptideCount=('Peptide', 'count'),
    Coverage=('Cov%', 'mean'),
    Length=('Length', 'first'),
    Weight_kDa=('Weight(kDa)', 'first'),
    Description=('Description', 'first'),
    Sequence=('Sequence', 'first')
).reset_index()
print(f"Protein data shape: {protein_data.shape}")
protein_data.head()
```

```
Making sure the info_data is aligned with the imputed data (latest version)...

Establish protein data, will be making use in later steps.
Protein data shape: (7161, 8)
```

|  | Protein | Gene | PeptideCount | Coverage | Length | Weight\_kDa | Description | Sequence |
| --- | --- | --- | --- | --- | --- | --- | --- | --- |
| 0 | A0A024RBG1 | NUDT4B | 5 | 30.3867 | 181 | 20.4213 | Diphosphoinositol polyphosphate phosphohydrola... | MMKFKPNQTRTYDREGFKKRAACLCFRSEQEDEVLLVSSSRYPDQW... |
| 1 | A0A0B4J2D5 | GATD3B | 7 | 35.4478 | 268 | 28.1247 | Putative glutamine amidotransferase-like class... | MAAVRALVASRLAAASAFTSLSPGGRTPSQRAALHLSVPRPAARVA... |
| 2 | A0A1B0GTU1 | ZC3H11B | 5 | 8.1988 | 805 | 88.8824 | Zinc finger CCCH domain-containing protein 11B | MPNQGEDCYFFFYSTCTKGDSCPFRHCEAALGNETVCTLWQEGRCF... |
| 3 | A0A1W2PQL4 | ZNF722 | 5 | 11.7188 | 384 | 44.5719 | Zinc finger protein 722 | MAERPGSPGSREMRLLTFRDIAIEFSLEEWQCLDCAQQNLYRDVML... |
| 4 | A0A2R8Y4L2 | HNRNPA1L3 | 12 | 44.7273 | 275 | 29.1558 | Heterogeneous nuclear ribonucleoprotein A1-like 3 | MRDPNTKRSRGFGFVTYATVEEVDAAMNARPHKVDGRVVEPKRAVS... |

The alignmend of information is the most crucial. Then a protein data is generate (above), where some info like PeptideCount and Coverage is added to the fasta like table to ease of access.

---

##### 05.2 Subset of UniProt Annotations¶

The UniProt annotations is a large table, and we don't need it all only the 7161 proteins that we have in our final dataset. Therefore, we subset the UniProt annotations to only include entries corresponding to the proteins present in our cleaned data. This reduces file size and improves efficiency for downstream analyses that utilize these annotations.

```
print("Filtering UniProt data to match the identified proteins...")
uniprot_data = uniprot_data[uniprot_data['Protein'].isin(protein_data['Protein'].tolist())]
print(f"Filtered UniProt data shape: {uniprot_data.shape}")
print("Renaming UniProt columns for clarity...")
uniprot_data = uniprot_data.reset_index(drop=True)
print("UniProt data columns:", uniprot_data.columns.tolist())
uniprot_data.head()
```

```
Filtering UniProt data to match the identified proteins...
Filtered UniProt data shape: (774552, 10)
Renaming UniProt columns for clarity...
UniProt data columns: ['isoform', 'Protein', 'feature', 'start', 'end', 'group', 'agent', 'note', 'description', 'score']
```

|  | isoform | Protein | feature | start | end | group | agent | note | description | score |
| --- | --- | --- | --- | --- | --- | --- | --- | --- | --- | --- |
| 0 |  | A0A024RBG1 | CHAIN | 1 | 181 |  |  |  | Diphosphoinositol polyphosphate phosphohydrola... | 3 |
| 1 |  | A0A024RBG1 | CROSSLNK | 5 | 5 | Ubiquitination/SUMOylation |  | Ubiquitination/SUMOylation at K5 | Source: psp; Type: UBIQUITINATION | 1 |
| 2 |  | A0A024RBG1 | BINDING | 10 | 10 |  |  |  |  | 3 |
| 3 |  | A0A024RBG1 | BINDING | 18 | 20 |  |  |  |  | 3 |
| 4 |  | A0A024RBG1 | DOMAIN | 18 | 145 |  |  |  | Nudix hydrolase | 3 |

##### 05.3 Save All the Cleaned and Processed Tables¶

Finally, all cleaned and processed data tables are saved to the designated output directory. They are mainly saved in `feather` format for efficient i/o, which is well-suited for large tabular datasets commonly encountered in proteomics analyses. They can't be opened and viewed directly like tsv or csv files, but they are much faster to read and write, which is crucial when dealing with large datasets.

```
print("\n📁 Saving cleaned table...")
cleaned_data.to_feather(f'{output_path}cleaned_data.feather')
print(f"Cleaned data saved to: {output_path}cleaned_data.feather")

print("\n📁 Saving normalized data...")
centered_data.to_feather(f'{output_path}centered_data.feather')
print(f"Centered data saved to: {output_path}centered_data.feather")

print("\n📁 Saving imputed data...")
imputed_data.to_feather(f'{output_path}imputed_data.feather')
print(f"Imputed data saved to: {output_path}imputed_data.feather")

print("\n📁 Saving metadata...")
meta.to_csv(f'{output_path}metadata.csv', index=False)
print(f"Metadata saved to: {output_path}metadata.csv")

print("\n📁 Saving protein data...")
protein_data.to_feather(f'{output_path}protein_data.feather')
print(f"Protein data saved to: {output_path}protein_data.feather")

print("\n📁 Saving UniProt data...")
uniprot_data.to_feather(f'{output_path}uniprot_data.feather')
print(f"UniProt data saved to: {output_path}uniprot_data.feather")

print("\n📁 Saving info data...")
info_data.to_feather(f'{output_path}info_data.feather')
print(f"Info data saved to: {output_path}info_data.feather")
```

```
📁 Saving cleaned table...
Cleaned data saved to: ./data/cleaned/hypoxia/cleaned_data.feather

📁 Saving normalized data...
Centered data saved to: ./data/cleaned/hypoxia/centered_data.feather

📁 Saving imputed data...
Imputed data saved to: ./data/cleaned/hypoxia/imputed_data.feather

📁 Saving metadata...
Metadata saved to: ./data/cleaned/hypoxia/metadata.csv

📁 Saving protein data...
Protein data saved to: ./data/cleaned/hypoxia/protein_data.feather

📁 Saving UniProt data...
UniProt data saved to: ./data/cleaned/hypoxia/uniprot_data.feather

📁 Saving info data...
Info data saved to: ./data/cleaned/hypoxia/info_data.feather
```

---

#### Summary¶

This notebook prepared peptide-level proteomics data from H358 NSCLC cells under hypoxia treatment for proteoform analysis with ProteoForge.

##### Output Files¶

| File | Description |
| --- | --- |
| `cleaned_data.feather` | Data after outlier removal and filtering |
| `centered_data.feather` | Median-centered normalized intensities |
| `imputed_data.feather` | Complete dataset after multi-strategy imputation |
| `metadata.csv` | Sample metadata with condition and duration |
| `protein_data.feather` | Protein-level summary (gene, coverage, length) |
| `uniprot_data.feather` | Filtered UniProt annotations |
| `info_data.feather` | Peptide-to-protein mapping with sequences |

##### Key Processing Steps¶

1. **Data Loading**: Imported DIA-NN output, UniProt sequences, and annotations
2. **Sequence Mapping**: Mapped peptide positions and calculated sequence coverage
3. **Quality Filtering**: Removed proteins with <4 peptides, identified outlier samples
4. **Normalization**: Applied median centering with original magnitude preservation
5. **Imputation**: Multi-strategy approach (amputate → fill dense → downshift → k-NN)
6. **Quality Assessment**: Evaluated CV distribution across experimental conditions

The prepared data is now ready for ProteoForge proteoform detection analysis.

```
print("Notebook Execution Time:", utils.prettyTimer(utils.getTime() - startTime))
```

```
Notebook Execution Time: 00h:02m:38s
```
