## Supplementary material for "ProteoForge: An Imputation-Aware Framework for Differential Proteoform Discovery in Bottom-Up Proteomics": Analysis notebooks as html render: Notebook S10.html

### Notebook S10

###### Enes Kemal Ergin

#### 2025-12-08 23:00:23

### Supplementary Notebook 10: Applying ProteoForge to 48hr Hypoxia vs Normoxia Comparison¶

- **License:** Creative Commons Attribution-NonCommercial 4.0 International License
- **Version:** 0.2
- **Edit Log:**
  - 2025-11-28: Initial version of the notebook
  - 2025-12-08: Revise the whole notebook, ensuring clarity and correctness

---

**Requirements:**

This notebook requires the `01-DataPreparation.ipynb` to be run first to generate the cleaned data files used here. Ensure the `./data/cleaned/` directory contains the necessary input files.

---

**Data Information:**

To apply proteoforge to single comparison at a specific timepoint, we focus on the 48-hour hypoxia vs normoxia comparison. This is to get the differential proteoforms (dPFs) specific to hypoxia response at this timepoint.

- **Dataset:** NSCLC cell lines under hypoxia vs normoxia at 48 hours
- **Conditions:** Hypoxia (1% O2) vs Normoxia (21% O2)
- **Timepoint:** 48 hours
- **Biological Replicates:** 4 per condition

---

**Purpose:**

This notebook applies the ProteoForge pipeline to identify differential proteoforms (dPFs) between hypoxia and normoxia conditions at the 48-hour timepoint. The pipeline includes data preparation and normalization, weight calculation (not used for completeness), linear modeling, hierarchical clustering, and proteoform classification.

> **Note:** Focusing on a single timepoint allows us to capture proteoform changes specifically associated with hypoxia response, minimizing confounding effects from temporal dynamics. Additionally results from this will be applied to downstream statistical and functional analyses in `04-DownstreamComparison.ipynb`, when calculating dPF level protein quants.

---

#### 01. Setup¶

This section imports required libraries, configures display settings, and defines paths for data and figures.

> **Note:** The HTML rendering of this notebook hides code cells by default. Click the "Code" buttons on the right to expand them.

##### 01.1 Libraries¶

```
# Libraries Used

import os
import sys

import numpy as np # Numerical computing
import pandas as pd # Data manipulation

import seaborn as sns # R-like high-level plots
import matplotlib.pyplot as plt # Python's base plotting 

# home path
sys.path.append('../')
from src import utils, plots

# Functionality to Process the Data (Weight, Normalize, Model, Correlate, Cluster)
from ProteoForge import normalize, impute
# Functionality to Predict Discordant Peptides (id)
from ProteoForge import correct, weight, model
# Functionality to Cluster Peptides for Grouping (group)
from ProteoForge import disluster
# Functionality to Generate dPFs (differential proteoforms)
from ProteoForge import proteoform_classifier as pfc
# Initialize the timer
startTime = utils.getTime()
```

##### 01.2 Configure Notebook Settings¶

Configure visualization styles, color palettes, and display options for consistent output formatting throughout the analysis.

```
## Configure Plotting Settings
def_colors = [
    "#139593", "#fca311", "#e54f2a",
    "#c3c3c3", "#555555",
    "#690000", "#5f4a00", "#004549"
]

condition_colors = {
    'True Hypoxia (1% O2) - 48 Hr': '#e9c46a',
    'True Hypoxia (1% O2) - 72 Hr': '#f4a261',
    'Normoxia (21% O2) - 48 Hr': '#ade8f4',
    'Normoxia (21% O2) - 72 Hr': '#0077b6',
    # Won't be used since the manuscript didn't cover these conditions
    # 'Chemical Hypoxia (CoCl2) - 48 Hr': '#c77dff',
    # 'Chemical Hypoxia (CoCl2) - 72 Hr': '#8338ec',
    # 'Oxidative Stress (H2O2) - 48 Hr': '#ff006e',
    # 'Oxidative Stress (H2O2) - 72 Hr': '#fb5607',
}

# Set seaborn style
sns.set_theme(
    style="white",
    context="paper",
    palette=def_colors,
    font_scale=1,
    rc={
        "figure.figsize": (6, 4),
        "font.family": "sans-serif",
        "font.sans-serif": ["Arial", "Ubuntu Mono"],
    }
)

# Figure Saving Settings
figure_formats = ["png", "pdf"]
save_to_folder = True
transparent_bg = True
figure_dpi = 300

## Configure dataframe displaying
pd.set_option('display.float_format', lambda x: '%.4f' % x)
pd.set_option('display.max_rows', 50)
pd.set_option('display.max_columns', 50)
pd.set_option('display.width', 500)  # Set a wider display width


## Printing Settings
verbose = True

plots.color_palette( def_colors, save=False )
plots.color_palette( condition_colors, save=False, name='condition', size=2)
```


##### 01.3 Data and Result Paths¶

- `input_path` — Cleaned data from `01-DataPreparation.ipynb`
- `output_path` — ProteoForge analysis results (`./data/results/hypoxia`)
- `figure_path` — Generated figures for 48hr comparison
- `notebook_name` — Current notebook name for logging (`02-ProteoForge`)

```
home_path = './'
data_name = "hypoxia"
notebook_name = "02-ProteoForge"
data_path = f"{home_path}data/"
input_path = f"{data_path}cleaned/{data_name}/"
output_path = f"{data_path}results/{data_name}/"
figure_path = f"{home_path}figures/{data_name}/{notebook_name}/"

# Create the output folder if it does not exist
if not os.path.exists(output_path):
    os.makedirs(output_path)

# Create figure folder structure, if needed
if save_to_folder:
    for i in figure_formats:
        cur_folder = figure_path + i + "/"
        if not os.path.exists(cur_folder):
            os.makedirs(cur_folder)
```

##### 01.4 Load Input Data¶

Load preprocessed datasets from the data preparation notebook including UniProt annotations, protein-level summaries, peptide mappings, cleaned and imputed intensity data, and sample metadata.

**Uniprot Data**: Contains expanded annotations for proteins in the data.

```
uniprot_data = pd.read_feather(f"{input_path}uniprot_data.feather")
print(uniprot_data.shape)
uniprot_data.head()
```

```
(774552, 10)
```

|  | isoform | Protein | feature | start | end | group | agent | note | description | score |
| --- | --- | --- | --- | --- | --- | --- | --- | --- | --- | --- |
| 0 |  | A0A024RBG1 | CHAIN | 1 | 181 |  |  |  | Diphosphoinositol polyphosphate phosphohydrola... | 3 |
| 1 |  | A0A024RBG1 | CROSSLNK | 5 | 5 | Ubiquitination/SUMOylation |  | Ubiquitination/SUMOylation at K5 | Source: psp; Type: UBIQUITINATION | 1 |
| 2 |  | A0A024RBG1 | BINDING | 10 | 10 |  |  |  |  | 3 |
| 3 |  | A0A024RBG1 | BINDING | 18 | 20 |  |  |  |  | 3 |
| 4 |  | A0A024RBG1 | DOMAIN | 18 | 145 |  |  |  | Nudix hydrolase | 3 |

**Protein Data**: Summarized protein level information of the proteins that will be used in the analysis.

```
protein_data = pd.read_feather(f"{input_path}protein_data.feather")
print(protein_data.shape)
protein_data.head()
```

```
(7161, 8)
```

|  | Protein | Gene | PeptideCount | Coverage | Length | Weight\_kDa | Description | Sequence |
| --- | --- | --- | --- | --- | --- | --- | --- | --- |
| 0 | A0A024RBG1 | NUDT4B | 5 | 30.3867 | 181 | 20.4213 | Diphosphoinositol polyphosphate phosphohydrola... | MMKFKPNQTRTYDREGFKKRAACLCFRSEQEDEVLLVSSSRYPDQW... |
| 1 | A0A0B4J2D5 | GATD3B | 7 | 35.4478 | 268 | 28.1247 | Putative glutamine amidotransferase-like class... | MAAVRALVASRLAAASAFTSLSPGGRTPSQRAALHLSVPRPAARVA... |
| 2 | A0A1B0GTU1 | ZC3H11B | 5 | 8.1988 | 805 | 88.8824 | Zinc finger CCCH domain-containing protein 11B | MPNQGEDCYFFFYSTCTKGDSCPFRHCEAALGNETVCTLWQEGRCF... |
| 3 | A0A1W2PQL4 | ZNF722 | 5 | 11.7188 | 384 | 44.5719 | Zinc finger protein 722 | MAERPGSPGSREMRLLTFRDIAIEFSLEEWQCLDCAQQNLYRDVML... |
| 4 | A0A2R8Y4L2 | HNRNPA1L3 | 12 | 44.7273 | 275 | 29.1558 | Heterogeneous nuclear ribonucleoprotein A1-like 3 | MRDPNTKRSRGFGFVTYATVEEVDAAMNARPHKVDGRVVEPKRAVS... |

**Info Data**: The expanded information about the peptides and proteins in the dataset, containing index of peptide within the protein (ordered from start and end positions), and all other information that will be used in the analysis.

```
info_data = pd.read_feather(f"{input_path}info_data.feather")
print(info_data.shape)
info_data.head()
```

```
(95369, 15)
```

|  | Protein | index | Peptide | Gene | Description | Length | Weight(kDa) | Sequence | peptide\_start | peptide\_end | occurrence\_index | total\_occurrences | unique\_id | Cov% | trace |
| --- | --- | --- | --- | --- | --- | --- | --- | --- | --- | --- | --- | --- | --- | --- | --- |
| 0 | A0A024RBG1 | 1 | TYDREGFK | NUDT4B | Diphosphoinositol polyphosphate phosphohydrola... | 181 | 20.4213 | MMKFKPNQTRTYDREGFKKRAACLCFRSEQEDEVLLVSSSRYPDQW... | 11.0000 | 18.0000 | 1.0000 | 1 | A0A024RBG1-TYDREGFK-11.0 | 30.3867 | 0 |
| 1 | A0A024RBG1 | 2 | SEQEDEVLLVSSSR | NUDT4B | Diphosphoinositol polyphosphate phosphohydrola... | 181 | 20.4213 | MMKFKPNQTRTYDREGFKKRAACLCFRSEQEDEVLLVSSSRYPDQW... | 28.0000 | 41.0000 | 1.0000 | 1 | A0A024RBG1-SEQEDEVLLVSSSR-28.0 | 30.3867 | 0 |
| 2 | A0A024RBG1 | 3 | EVYEEAGVKGK | NUDT4B | Diphosphoinositol polyphosphate phosphohydrola... | 181 | 20.4213 | MMKFKPNQTRTYDREGFKKRAACLCFRSEQEDEVLLVSSSRYPDQW... | 66.0000 | 76.0000 | 1.0000 | 1 | A0A024RBG1-EVYEEAGVKGK-66.0 | 30.3867 | 0 |
| 3 | A0A024RBG1 | 4 | LLGIFEQNQDRK | NUDT4B | Diphosphoinositol polyphosphate phosphohydrola... | 181 | 20.4213 | MMKFKPNQTRTYDREGFKKRAACLCFRSEQEDEVLLVSSSRYPDQW... | 80.0000 | 91.0000 | 1.0000 | 1 | A0A024RBG1-LLGIFEQNQDRK-80.0 | 30.3867 | 0 |
| 4 | A0A024RBG1 | 5 | EWFKVEDAIK | NUDT4B | Diphosphoinositol polyphosphate phosphohydrola... | 181 | 20.4213 | MMKFKPNQTRTYDREGFKKRAACLCFRSEQEDEVLLVSSSRYPDQW... | 119.0000 | 128.0000 | 1.0000 | 1 | A0A024RBG1-EWFKVEDAIK-119.0 | 30.3867 | 0 |

**Quantitative Data**: The cleaned and imputed versions of the peptide level data in wide-format. The wide format entails that each row is a peptide and each column is a sample, with intensity values as the entries.

```
cleaned_data = pd.read_feather(f"{input_path}cleaned_data.feather")
print(cleaned_data.shape)
cleaned_data.head()

imputed_data = pd.read_feather(f"{input_path}imputed_data.feather")
print(imputed_data.shape)
imputed_data.head()
```

```
(95369, 14)
(95369, 14)
```

|  | Colname | 1%\_48\_1 | 1%\_48\_2 | 1%\_48\_3 | 1%\_48\_4 | 1%\_72\_2 | 1%\_72\_3 | 1%\_72\_4 | C\_48\_1 | C\_48\_2 | C\_48\_3 | C\_48\_4 | C\_72\_1 | C\_72\_2 | C\_72\_3 |
| --- | --- | --- | --- | --- | --- | --- | --- | --- | --- | --- | --- | --- | --- | --- | --- |
| Protein | unique\_id |  |  |  |  |  |  |  |  |  |  |  |  |  |  |
| A0A024RBG1 | A0A024RBG1-TYDREGFK-11.0 | 11907.9257 | 13382.8577 | 12383.7035 | 10530.5089 | 8083.9621 | 6812.1680 | 9593.1931 | 14181.9304 | 12918.3702 | 11056.6977 | 13999.6338 | 8248.7570 | 11734.2555 | 13588.2897 |
|  | A0A024RBG1-SEQEDEVLLVSSSR-28.0 | 85176.7848 | 96480.5830 | 83977.0036 | 99640.0878 | 86286.4272 | 81336.9615 | 75096.5430 | 105689.9042 | 90460.1718 | 98407.4817 | 113661.0614 | 88837.1056 | 118683.7065 | 104112.7786 |
|  | A0A024RBG1-EVYEEAGVKGK-66.0 | 43679.2578 | 52024.5608 | 42553.7306 | 44072.4676 | 31219.7739 | 38914.0183 | 42793.7742 | 50800.5659 | 52223.7104 | 29521.7964 | 34854.6125 | 45086.6695 | 60278.9107 | 53039.1121 |
|  | A0A024RBG1-LLGIFEQNQDRK-80.0 | 15667.9096 | 10994.3745 | 12312.3053 | 15382.4000 | 16444.1212 | 15101.0793 | 12342.9230 | 12070.6690 | 11760.2174 | 15145.2258 | 19111.7337 | 12542.1749 | 14849.4373 | 15791.4696 |
|  | A0A024RBG1-EWFKVEDAIK-119.0 | 814.8576 | 2265.3315 | 3586.0586 | 3978.2879 | 5425.7305 | 4849.3269 | 2441.8884 | 3025.9374 | 1719.1125 | 3627.5928 | 4442.8020 | 1690.2038 | 2936.9368 | 5103.2884 |

**Metadata**: Sample metadata including condition labels and other experimental variables, which will be used for mapping, modelling, and other analyses.

```
meta = pd.read_csv(f"{input_path}metadata.csv")
meta
```

|  | filename | Run | Condition | Replicate | Duration | Colname |
| --- | --- | --- | --- | --- | --- | --- |
| 0 | D:\Data\Tam\RevisionLungDIA\H358\H358\_1%\_1\_48\_... | H358\_1%\_1\_48\_DIA\_BB4\_1\_5432 | True Hypoxia (1% O2) | 1 | 48 | 1%\_48\_1 |
| 1 | D:\Data\Tam\RevisionLungDIA\H358\H358\_1%\_2\_48\_... | H358\_1%\_2\_48\_DIA\_BC4\_1\_5440 | True Hypoxia (1% O2) | 2 | 48 | 1%\_48\_2 |
| 2 | D:\Data\Tam\RevisionLungDIA\H358\H358\_1%\_3\_48\_... | H358\_1%\_3\_48\_DIA\_BD4\_1\_5448 | True Hypoxia (1% O2) | 3 | 48 | 1%\_48\_3 |
| 3 | D:\Data\Tam\RevisionLungDIA\H358\H358\_1%\_4\_48\_... | H358\_1%\_4\_48\_DIA\_BE4\_1\_5456 | True Hypoxia (1% O2) | 4 | 48 | 1%\_48\_4 |
| 4 | D:\Data\Tam\RevisionLungDIA\H358\H358\_1%\_2\_72\_... | H358\_1%\_2\_72\_DIA\_BC8\_1\_5444 | True Hypoxia (1% O2) | 2 | 72 | 1%\_72\_2 |
| 5 | D:\Data\Tam\RevisionLungDIA\H358\H358\_1%\_3\_72\_... | H358\_1%\_3\_72\_DIA\_BD8\_1\_5452 | True Hypoxia (1% O2) | 3 | 72 | 1%\_72\_3 |
| 6 | D:\Data\Tam\RevisionLungDIA\H358\H358\_1%\_4\_72\_... | H358\_1%\_4\_72\_DIA\_BE8\_1\_5460 | True Hypoxia (1% O2) | 4 | 72 | 1%\_72\_4 |
| 7 | D:\Data\Tam\RevisionLungDIA\H358\H358\_C1\_48\_DI... | H358\_C1\_48\_DIA\_BB1\_1\_5429 | Normoxia (21% O2) | 1 | 48 | C\_48\_1 |
| 8 | D:\Data\Tam\RevisionLungDIA\H358\H358\_C2\_48\_DI... | H358\_C2\_48\_DIA\_BC1\_1\_5437 | Normoxia (21% O2) | 2 | 48 | C\_48\_2 |
| 9 | D:\Data\Tam\RevisionLungDIA\H358\H358\_C3\_48\_DI... | H358\_C3\_48\_DIA\_BD1\_1\_5445 | Normoxia (21% O2) | 3 | 48 | C\_48\_3 |
| 10 | D:\Data\Tam\RevisionLungDIA\H358\H358\_C4\_48\_DI... | H358\_C4\_48\_DIA\_BE1\_1\_5453 | Normoxia (21% O2) | 4 | 48 | C\_48\_4 |
| 11 | D:\Data\Tam\RevisionLungDIA\H358\H358\_C1\_72\_DI... | H358\_C1\_72\_DIA\_BB5\_1\_5433 | Normoxia (21% O2) | 1 | 72 | C\_72\_1 |
| 12 | D:\Data\Tam\RevisionLungDIA\H358\H358\_C2\_72\_DI... | H358\_C2\_72\_DIA\_BC5\_1\_5441 | Normoxia (21% O2) | 2 | 72 | C\_72\_2 |
| 13 | D:\Data\Tam\RevisionLungDIA\H358\H358\_C3\_72\_DI... | H358\_C3\_72\_DIA\_BD5\_1\_5449 | Normoxia (21% O2) | 3 | 72 | C\_72\_3 |

```
# Build a dictionary to match condition_colors keys: 'Condition - Duration Hr'
condition_to_samples = {}
for condition in meta['Condition'].unique():
    for duration in sorted(meta.loc[meta['Condition'] == condition, 'Duration'].unique()):
        key = f"{condition} - {duration} Hr"
        samples = meta.loc[(meta['Condition'] == condition) & (meta['Duration'] == duration), 'Colname'].tolist()
        condition_to_samples[key] = samples

# samples_to_condition mapping
samples_to_condition = {sample: condition for condition, samples in condition_to_samples.items() for sample in samples}
# samples_to_colors mapping
samples_to_colors = {sample: condition_colors[condition] for sample, condition in samples_to_condition.items()}
```

---

#### 02. ProteoForge Analysis Pipeline¶

This section shows how the ProteoForge pipeline is applied to the 48-hour hypoxia vs normoxia comparison, including data preparation, weight calculation, linear modeling, hierarchical clustering, and proteoform classification.

##### 02.1 Prepare Analysis Data¶

Prepare the test dataset by merging peptide information with quantification data, subsetting to 48-hour timepoint conditions (Hypoxia vs Normoxia), normalizing intensities against the control condition, and tracking imputation status for downstream weighting.

```
test_data = info_data.sort_values(
    by=['Protein', 'peptide_start', 'peptide_end']
)[['unique_id', 'Protein', 'Peptide', 'peptide_start', 'peptide_end']]
# peptide_end and start should be int
test_data['peptide_start'] = test_data['peptide_start'].astype(int)
test_data['peptide_end'] = test_data['peptide_end'].astype(int)
# Generate PeptideID from cum
test_data['PeptideID'] = test_data.groupby('Protein').cumcount() + 1

print()
print("📊 MELTING CLEANED AND IMPUTED DATA FOR ANALYSIS")
# Get the imputed and cleaned (with missing values) data into long format
melt_imputed = imputed_data.reset_index().drop(
    columns='Protein'
).melt(
    id_vars='unique_id',
    var_name='Sample',
    value_name='Intensity'
).set_index(['unique_id', 'Sample'])
melt_cleaned = cleaned_data.reset_index().drop(
    columns='Protein'
).melt(
    id_vars='unique_id',
    var_name='Sample',
    value_name='Intensity(Raw)'
).set_index(['unique_id', 'Sample'])

combined_data = pd.concat(
    [melt_imputed, melt_cleaned],
    axis=1
).reset_index()

print()
print("📊 ADDITIONAL INFO ADDED")
combined_data['log10Intensity'] = np.log10(combined_data['Intensity'])
combined_data['Condition'] = combined_data['Sample'].map(samples_to_condition)
combined_data['isReal'] = combined_data['Intensity(Raw)'].notna()

compMiss_table = pd.DataFrame(index=cleaned_data.index)
for cond, col in condition_to_samples.items():
    compMiss_table[cond] = cleaned_data[col].isna().all(axis=1)

compMiss_table = compMiss_table.reset_index().drop(columns='Protein').melt(
    id_vars='unique_id',
    var_name='Condition',
    value_name='isCompleteMiss'
)
combined_data = combined_data.merge(
    compMiss_table,
    on=['unique_id', 'Condition'],
    how='left'
)
combined_data = combined_data[[
    'unique_id', 'Condition', 'Sample', 
    'isCompleteMiss', 'isReal',
    'Intensity(Raw)', 'Intensity', 'log10Intensity', 
]]

print()
print("📊 COMBINING INFO AND QUANT DATA")

# Merge the test_data with combined_data
test_data = test_data.merge(
    combined_data,
    on='unique_id',
    how='left'
)

# conditions to subset for 
conditions = [
    # # Removed the chemical and oxidative versions since they were not used in the manuscript
    # 'Oxidative Stress (H2O2) - 48 Hr',
    # 'Oxidative Stress (H2O2) - 72 Hr',
    # 'Chemical Hypoxia (CoCl2) - 48 Hr',
    # 'Chemical Hypoxia (CoCl2) - 72 Hr',
    'True Hypoxia (1% O2) - 48 Hr',
    # 'True Hypoxia (1% O2) - 72 Hr',
    'Normoxia (21% O2) - 48 Hr',
    # 'Normoxia (21% O2) - 72 Hr',
]

# Update the condition_to_samples to only include the conditions of interest
condition_to_samples = {
    cond: condition_to_samples[cond] for cond in conditions if cond in condition_to_samples
}

test_data = test_data[
    test_data['Condition'].isin(condition_to_samples.keys())
].reset_index(drop=True)

print()
print("📏 NORMALIZING INTENSITIES AGAINST CONTROL CONDITION")
test_data = normalize.against_condition(
    long_data=test_data,
    cond_run_dict=condition_to_samples,
    run_col='Sample',
    index_cols=['Protein', 'PeptideID'],
    norm_against='Normoxia (21% O2) - 48 Hr',
    intensity_col='Intensity',
    is_log2=False,
    norm_intensity_col='AdjIntensity'
)
print()
# print("🛡️ CALCULATING ROBUST INTENSITIES FOR CLUSTERING")
# test_data = normalize.build_robust_intensity(
#     test_data,
#     log_col='log10Intensity',
#     adj_col='AdjIntensity',
#     protein_col='Protein',
#     cond_col='Condition',
#     out_col='RobIntensity',
#     verbose=verbose

# )
print(f"Test Data shape after normalization: {test_data.shape}")
test_data.head()
```

```
📊 MELTING CLEANED AND IMPUTED DATA FOR ANALYSIS

📊 ADDITIONAL INFO ADDED

📊 COMBINING INFO AND QUANT DATA

📏 NORMALIZING INTENSITIES AGAINST CONTROL CONDITION

Test Data shape after normalization: (762952, 14)
```

|  | unique\_id | Protein | Peptide | peptide\_start | peptide\_end | PeptideID | Condition | Sample | isCompleteMiss | isReal | Intensity(Raw) | Intensity | log10Intensity | AdjIntensity |
| --- | --- | --- | --- | --- | --- | --- | --- | --- | --- | --- | --- | --- | --- | --- |
| 0 | A0A024RBG1-TYDREGFK-11.0 | A0A024RBG1 | TYDREGFK | 11 | 18 | 1 | True Hypoxia (1% O2) - 48 Hr | 1%\_48\_1 | False | True | 15362.5723 | 11907.9257 | 4.0758 | -0.0330 |
| 1 | A0A024RBG1-TYDREGFK-11.0 | A0A024RBG1 | TYDREGFK | 11 | 18 | 1 | True Hypoxia (1% O2) - 48 Hr | 1%\_48\_2 | False | True | 16231.3125 | 13382.8577 | 4.1265 | 0.0213 |
| 2 | A0A024RBG1-TYDREGFK-11.0 | A0A024RBG1 | TYDREGFK | 11 | 18 | 1 | True Hypoxia (1% O2) - 48 Hr | 1%\_48\_3 | False | True | 15491.7900 | 12383.7035 | 4.0929 | -0.0303 |
| 3 | A0A024RBG1-TYDREGFK-11.0 | A0A024RBG1 | TYDREGFK | 11 | 18 | 1 | True Hypoxia (1% O2) - 48 Hr | 1%\_48\_4 | False | True | 13372.8730 | 10530.5089 | 4.0224 | -0.0881 |
| 4 | A0A024RBG1-TYDREGFK-11.0 | A0A024RBG1 | TYDREGFK | 11 | 18 | 1 | Normoxia (21% O2) - 48 Hr | C\_48\_1 | False | True | 17479.4805 | 14181.9304 | 4.1517 | 0.0386 |

Analysis data prepared with peptide positions, intensities, and condition assignments. Data is reshaped into a long-format. Intensities normalized against Normoxia (21% O₂) - 48 Hr as the control condition. Imputation status tracked via `isReal` and `isCompleteMiss` flags, which will be used in weight calculation.

---

##### 02.2 Generate Sample Weights¶

Weight calculation accounts for data quality differences. The imputation weight (90%) penalizes imputed values based on missingness pattern, while the technical variance weight (10%) down-weights high-variance measurements. This ensures statistical modeling prioritizes reliable measurements, but mainly focuses on imputation status.

> **Note:** The 90-10 split is chosen to heavily prioritize imputation status, as it is the main source of uncertainty in the data. The more sophisticated modelling of weights can be explored withing the `weight` module however I was not able to see any meaningful difference in the results when changing these values.
>
> **Note:** The sparse imputed value is set to a very low intensity (10^-5) to reflect the uncertainty of sparse missingness, while the dense imputed value is set to a moderate intensity (0.5) to reflect the higher confidence in dense missingness imputation.
>
> **IMPORTANT NOTE:** The weights calculated here are not used in the subsequent linear modeling, only when WLS (weighted least squares) or GLM (Generalized Linear modeling) is selected they are relevant. The weighting can be very sophisticate and `weight` module does provide a quite a bit of variation, however when using RLM or Quantile Regressions, custom weights can't be passed. The internal weights are calculated especially in the case of RLM. Here I am showing them for the sake of completeness for the pipeline demonstration.

```
weights_data = weight.generate_weights_data(
    test_data,
    sample_cols=['Sample'],
    log_intensity_col='log10Intensity',
    adj_intensity_col='AdjIntensity',
    control_condition='Normoxia (21% O2) - 48 Hr',
    condition_col='Condition',
    protein_col='Protein',
    peptide_col='Peptide',
    is_real_col='isReal',
    is_comp_miss_col='isCompleteMiss',
    sparse_imputed_val=10**-5,
    dense_imputed_val=0.5,
    verbose=verbose,
)
test_data['Weight'] = (
    (weights_data['W_Impute'] * 0.90) + 
    (weights_data['W_RevTechVar'] * 0.10)
)

print(f"Test Data shape after weight generation: {test_data.shape}")
test_data.head()
```

```
 📏 Calculating Imputation Weights... (done in 2.84 ms)
 📏 Calculating Reverse Technical Variation Weights... (done in 201.89 ms)
All weights calculated and normalized.
Test Data shape after weight generation: (762952, 15)
```

|  | unique\_id | Protein | Peptide | peptide\_start | peptide\_end | PeptideID | Condition | Sample | isCompleteMiss | isReal | Intensity(Raw) | Intensity | log10Intensity | AdjIntensity | Weight |
| --- | --- | --- | --- | --- | --- | --- | --- | --- | --- | --- | --- | --- | --- | --- | --- |
| 0 | A0A024RBG1-TYDREGFK-11.0 | A0A024RBG1 | TYDREGFK | 11 | 18 | 1 | True Hypoxia (1% O2) - 48 Hr | 1%\_48\_1 | False | True | 15362.5723 | 11907.9257 | 4.0758 | -0.0330 | 0.9999 |
| 1 | A0A024RBG1-TYDREGFK-11.0 | A0A024RBG1 | TYDREGFK | 11 | 18 | 1 | True Hypoxia (1% O2) - 48 Hr | 1%\_48\_2 | False | True | 16231.3125 | 13382.8577 | 4.1265 | 0.0213 | 0.9999 |
| 2 | A0A024RBG1-TYDREGFK-11.0 | A0A024RBG1 | TYDREGFK | 11 | 18 | 1 | True Hypoxia (1% O2) - 48 Hr | 1%\_48\_3 | False | True | 15491.7900 | 12383.7035 | 4.0929 | -0.0303 | 0.9999 |
| 3 | A0A024RBG1-TYDREGFK-11.0 | A0A024RBG1 | TYDREGFK | 11 | 18 | 1 | True Hypoxia (1% O2) - 48 Hr | 1%\_48\_4 | False | True | 13372.8730 | 10530.5089 | 4.0224 | -0.0881 | 0.9999 |
| 4 | A0A024RBG1-TYDREGFK-11.0 | A0A024RBG1 | TYDREGFK | 11 | 18 | 1 | Normoxia (21% O2) - 48 Hr | C\_48\_1 | False | True | 17479.4805 | 14181.9304 | 4.1517 | 0.0386 | 0.9999 |

##### 02.3 Linear Modeling¶

ProteoForge support multiple linear modelling and regression based methods to identify significantly discordant peptides within their protein context. Here we use the RLM (Robust Linear Modeling) method as the default, which is robust to outliers and can handle heteroscedasticity in the data, additionally if the data is not overly noisy, or contains many missing values that are imputed (> 50%), RLM can provide better specificity and sensitivity compared to WLS.

**Methods Supported:**

- RLM (Robust Linear Modeling) - No custom weights necessary and unnecessary to calculate and pass weights.
- WLS (Weighted Least Squares) - Custom weights can be calculated and passed to the model.
- GLM (Generalized Linear Modeling) - Custom weights can be calculated and passed to the model.
- Quantile Regression - No custom weights necessary and unnecessary to calculate and pass weights. (mqr)
- OLS (Ordinary Least Squares) - No weights can be passed, this is the simplest model.

RLM (Robust Linear Modeling) used to identify significantly discordant peptides. Each peptide is tested independently within its protein context with auto-calculated robust weights calculated within RLM. Wald-test for interaction term is used to extract the p-value for interaction.Then two-step correction is applied: Bonferroni within protein, then FDR (Benjamini-Hochberg) across all tests.

> Note: The two-level correction might be too conservative but it is important to introduce due to amount of tests being performed, within the protein and then across all proteins.

```
n_jobs = 16
model_to_use = 'rlm' # Options: 'ols', 'wls', 'rlm', 'robust', 'quantile', glm
correction = {
    'strategy': 'two-step',
    'methods': ['bonferroni', 'fdr_bh']
}
# Initialize the linear model with parameters
cur_model = model.LinearModel(
    data=test_data,
    protein_col="Protein",
    peptide_col="Peptide",
    cond_col="Condition",
    intensity_col="AdjIntensity",
    weight_col="Weight",
)

test_data = cur_model.run_analysis(
    model_type=model_to_use,
    correction_strategy= correction['strategy'],
    correction_methods=correction['methods'],
    n_jobs=n_jobs
)

if correction['strategy'] == 'two-step' and isinstance(correction['methods'], tuple):
    methods_str = " and ".join(correction['methods'])
    print(f"  Ran weighted least squares (WLS) model with two-step correction")
    print(f"  Used {n_jobs} parallel jobs")
    print(f"  Applied {methods_str} corrections")
else:
    method_str = correction['methods'] if isinstance(correction['methods'], str) else ", ".join(correction['methods'])
    print(f"  Ran weighted least squares (WLS) model with {correction['strategy']} correction")
    print(f"  Used {n_jobs} parallel jobs")
    print(f"  Applied {method_str} correction")
print()
```

```
  Ran weighted least squares (WLS) model with two-step correction
  Used 16 parallel jobs
  Applied bonferroni, fdr_bh correction
```

##### 02.4 Visualize Statistical Results¶

Compare p-value distributions across correction stages: raw p-values, within-protein Bonferroni correction, and FDR-adjusted p-values. The combined analysis may reveal different significance patterns compared to individual timepoint analyses.

```
pThr = 0.001
palette = {
    'pval': '#4c72b0',
    'prt_pval': '#dd8452',
    'adj_pval': '#55a868'
}
# --- 3. Plotting ---
# Define the columns to plot and their corresponding titles
plot_configs = {
    'pval': 'Raw p-value',
    'prt_pval': 'p-value Within Protein (bonferroni)',
    'adj_pval': 'Adjusted p-value (fdr_bh)'
}

# Calculate log10 p-values for each column, replacing zeros with minimum nonzero value for stability
for col in plot_configs.keys():
    min_nonzero = test_data[col][test_data[col] > 0].min() if (test_data[col] > 0).any() else 1e-300
    test_data[f'log10_{col}'] = np.log10(test_data[col].replace(0, min_nonzero))

fig, axes = plt.subplots(1, 3, figsize=(20, 7), sharex=True, sharey=True)

for ax, (col, title) in zip(axes, plot_configs.items()):
    significant_mask = test_data[col] < pThr
    significant_hits = significant_mask.sum()
    total_hits = test_data[col].count()
    percent_hits = 100 * significant_hits / total_hits if total_hits > 0 else 0
    # Proteins
    total_proteins = test_data['Protein'].nunique()
    significant_proteins = test_data.loc[significant_mask, 'Protein'].nunique()
    percent_proteins = 100 * significant_proteins / total_proteins if total_proteins > 0 else 0
    # Peptides
    total_peptides = test_data['Peptide'].nunique()
    significant_peptides = test_data.loc[significant_mask, 'Peptide'].nunique()
    percent_peptides = 100 * significant_peptides / total_peptides if total_peptides > 0 else 0

    # Drop NaN values for histogram
    log10_vals = test_data[f'log10_{col}'].dropna()
    if len(log10_vals) == 0:
        ax.text(0.5, 0.5, 'No data available', transform=ax.transAxes, ha='center', va='center', fontsize=14)
        continue

    counts, bins = np.histogram(log10_vals, bins=50)
    counts_log10 = np.log10(np.where(counts == 0, 1, counts))

    ax.bar(bins[:-1], counts_log10, width=np.diff(bins), align='edge', color=palette[col], edgecolor='white', linewidth=0.5, alpha=0.8)

    # Add vertical line for threshold
    x_thr = np.log10(pThr)
    ax.axvline(x=x_thr, color='crimson', linestyle='--', linewidth=2)

    # Add vertical annotation next to the line
    ylim = ax.get_ylim()
    ax.text(
        x_thr, ylim[1]*0.95, 
        f'log10={x_thr:.0f}', 
        color='crimson', rotation=90, 
        va='top', ha='right', 
        fontsize=12, fontweight='bold', 
        bbox=dict(boxstyle='round,pad=0.2', fc='white', ec='crimson', alpha=0.7)
    )

    annotation = (
        f'Hits: {significant_hits:,} / {total_hits:,} ({percent_hits:.2f}%)\n'
        f'Proteins: {significant_proteins:,} / {total_proteins:,} ({percent_proteins:.2f}%)\n'
        f'Peptides: {significant_peptides:,} / {total_peptides:,} ({percent_peptides:.2f}%)'
    )
    ax.text(
        0.05, 0.90,
        annotation,
        transform=ax.transAxes,
        fontsize=12, fontweight='bold',
        ha='left', va='top',
        bbox=dict(boxstyle='round,pad=0.3', fc='white', ec='black', alpha=0.75)
    )

    ax.set_title(title + ' (log10)', fontsize=16, fontweight='bold')
    ax.set_xlabel('log10(p-value)', fontsize=12)
    ax.set_ylabel('log10(Frequency)', fontsize=14)
    ax.grid(axis='y', linestyle='--', alpha=0.6)
    ax.tick_params(axis='x', rotation=45)

axes[1].set_ylabel('')
axes[2].set_ylabel('')

fig.suptitle(
    'Comparison of log10 p-value Distributions',
    fontsize=22,
    fontweight='bold',
    y=1.02
    )

sns.despine()
plt.tight_layout(rect=[0, 0, 1, 0.96])
plots.finalize_plot( 
    fig, show=True, save=save_to_folder, 
    filename='pvalue_distributions_48hr',
    filepath=figure_path,
    formats=figure_formats,
    transparent=transparent_bg,
    dpi=figure_dpi
)

test_data['isSignificant'] = test_data['adj_pval'] < pThr
test_data.head()
```

|  | unique\_id | Protein | Peptide | peptide\_start | peptide\_end | PeptideID | Condition | Sample | isCompleteMiss | isReal | Intensity(Raw) | Intensity | log10Intensity | AdjIntensity | Weight | pval | prt\_pval | adj\_pval | log10\_pval | log10\_prt\_pval | log10\_adj\_pval | isSignificant |
| --- | --- | --- | --- | --- | --- | --- | --- | --- | --- | --- | --- | --- | --- | --- | --- | --- | --- | --- | --- | --- | --- | --- |
| 0 | A0A024RBG1-TYDREGFK-11.0 | A0A024RBG1 | TYDREGFK | 11 | 18 | 1 | True Hypoxia (1% O2) - 48 Hr | 1%\_48\_1 | False | True | 15362.5723 | 11907.9257 | 4.0758 | -0.0330 | 0.9999 | 0.8481 | 1.0000 | 1.0000 | -0.0716 | 0.0000 | 0.0000 | False |
| 1 | A0A024RBG1-TYDREGFK-11.0 | A0A024RBG1 | TYDREGFK | 11 | 18 | 1 | True Hypoxia (1% O2) - 48 Hr | 1%\_48\_2 | False | True | 16231.3125 | 13382.8577 | 4.1265 | 0.0213 | 0.9999 | 0.8481 | 1.0000 | 1.0000 | -0.0716 | 0.0000 | 0.0000 | False |
| 2 | A0A024RBG1-TYDREGFK-11.0 | A0A024RBG1 | TYDREGFK | 11 | 18 | 1 | True Hypoxia (1% O2) - 48 Hr | 1%\_48\_3 | False | True | 15491.7900 | 12383.7035 | 4.0929 | -0.0303 | 0.9999 | 0.8481 | 1.0000 | 1.0000 | -0.0716 | 0.0000 | 0.0000 | False |
| 3 | A0A024RBG1-TYDREGFK-11.0 | A0A024RBG1 | TYDREGFK | 11 | 18 | 1 | True Hypoxia (1% O2) - 48 Hr | 1%\_48\_4 | False | True | 13372.8730 | 10530.5089 | 4.0224 | -0.0881 | 0.9999 | 0.8481 | 1.0000 | 1.0000 | -0.0716 | 0.0000 | 0.0000 | False |
| 4 | A0A024RBG1-TYDREGFK-11.0 | A0A024RBG1 | TYDREGFK | 11 | 18 | 1 | Normoxia (21% O2) - 48 Hr | C\_48\_1 | False | True | 17479.4805 | 14181.9304 | 4.1517 | 0.0386 | 0.9999 | 0.8481 | 1.0000 | 1.0000 | -0.0716 | 0.0000 | 0.0000 | False |

##### 02.5 Peptide Clustering¶

Hierarchical clustering groups peptides with similar abundance patterns using Euclidean distance on adjusted intensities with Ward's linkage method. Cluster determination uses hybrid outlier cut for automatic cluster number selection. Peptides within the same cluster are assumed to derive from the same proteoform.

```
clusters = disluster.distance_and_cluster(
    data=test_data,
    protein_col='Protein',
    peptide_col='PeptideID',
    cond_col='Condition',
    quant_col='AdjIntensity',
    pdist_metric='euclidean',
    clustering_params={
        'min_clusters': 1,
        'distance_transform': 'corr',
        'clustering_method': 'hybrid_outlier_cut',
        'linkage_method': 'ward',
        'distance_metric': 'euclidean'
    },
    n_jobs=n_jobs,
    verbose=False
)

test_data = test_data.merge(
    clusters[['Protein', 'PeptideID', 'cluster_label']],
    on=['Protein', 'PeptideID'],
    how='left'
).rename(columns={'cluster_label': 'ClusterID'})

# Count proteins and peptides processed
proteins_processed = test_data['Protein'].nunique()
peptides_processed = test_data['Peptide'].nunique()

print(f"  Performed hierarchical clustering with average linkage and correlation distance")
print(f"  Auto-determined optimal number of clusters (1-25 range)")
print(f"  Calculated clusters with {n_jobs} parallel jobs")
print(f"  Processed {proteins_processed} proteins with {peptides_processed} peptides")
```

```
  Performed hierarchical clustering with average linkage and correlation distance
  Auto-determined optimal number of clusters (1-25 range)
  Calculated clusters with 16 parallel jobs
  Processed 7161 proteins with 89651 peptides
```

##### 02.6 Identify Differential Proteoforms¶

Differential proteoform (dPF) classification identifies proteins with at least one cluster containing significant peptides AND at least one cluster containing non-significant peptides. This pattern indicates that different regions of the protein respond differently to hypoxia, suggesting proteoform-level regulation.

```
summary_data = test_data[[
    'Protein', 'PeptideID', 'Peptide',
    'peptide_start', 'peptide_end',
    'pval', 'adj_pval',
    'isSignificant', 'ClusterID'
]].drop_duplicates().copy()

summary_data = pfc.process_peptide_data(
    summary_data, 
    protein_col='Protein',
    cluster_col='ClusterID',
    significance_col='isSignificant',
    dpf_col='dPF', 
    validate=False,
    verbose=verbose
)
print(f"  Processed {summary_data['dPF'].sum()} differential proteoforms (dPFs)")

summary_data = protein_data.merge(
    summary_data,
    on='Protein',
    how='left'
)

# Add the dPF to the test_data
test_data = test_data.merge(
    summary_data[['Protein', 'PeptideID', 'dPF']],
    on=['Protein', 'PeptideID'],
    how='left'
)
```

```
Processing 95,369 peptides from 7,161 proteins...
Auto-selected algorithm: ultra_fast
Ultra-fast algorithm: Processing 95369 peptides...
  ✓ Ultra-fast processing complete: maximum performance achieved

Processing completed in 0.590 seconds
Performance: 161,600 peptides/second

Classification Summary:
  Single PTMs (dPF=-1): 3,038
  Non-differential (dPF=0): 80,219
  Differential proteoforms (dPF>0): 12,112 peptides
  Maximum proteoform group: dPF=3
  Processed 14268 differential proteoforms (dPFs)
```

#### 03. Save Results¶

Export the complete analysis results for downstream interpretation and visualization. Output includes peptide-level data with statistics and clusters (`test_data_48hr.feather`), protein-level summary with dPF classifications (`summary_data_48hr.feather`), and UniProt annotations (`uniprot_data_48hr.feather`).

```
test_data.to_feather(f"{output_path}test_data_48hr.feather")
summary_data.to_feather(f"{output_path}summary_data_48hr.feather")
uniprot_data.to_feather(f"{output_path}uniprot_data_48hr.feather")
```

---

#### Summary¶

This notebook applied the ProteoForge pipeline to compare True Hypoxia (1% O₂) vs Normoxia (21% O₂) at 48 hours in H358 NSCLC cells. The analysis prepared peptide data with condition-normalized intensities, generated quality-aware sample weights, performed weighted linear modeling with two-step multiple testing correction, clustered peptides by abundance patterns, and identified proteins with differential proteoform regulation.

```
print("Notebook Execution Time:", utils.prettyTimer(utils.getTime() - startTime))
```

```
Notebook Execution Time: 00h:01m:47s
```
