## Supplementary material for "ProteoForge: An Imputation-Aware Framework for Differential Proteoform Discovery in Bottom-Up Proteomics": Analysis notebooks as html render: Notebook S11.html

### Notebook S11

###### Enes Kemal Ergin

#### 2025-12-08 23:00:19

### Supplementary Notebook 11: Applying ProteoForge to 72hr Hypoxia vs Normoxia Comparison¶

- **License:** Creative Commons Attribution-NonCommercial 4.0 International License
- **Version:** 0.2
- **Edit Log:**
  - 2025-11-28: Initial version of the notebook
  - 2025-12-08: Revise the whole notebook, ensuring clarity and correctness

---

**Requirements:**

This notebook requires the `01-DataPreparation.ipynb` to be run first to generate the cleaned data files used here. Ensure the `./data/cleaned/` directory contains the necessary input files.

---

**Data Information:**

To apply proteoforge to single comparison at a specific timepoint, we focus on the 72-hour hypoxia vs normoxia comparison. This is to get the differential proteoforms (dPFs) specific to hypoxia response at this timepoint.

- **Dataset:** NSCLC cell lines under hypoxia vs normoxia at 72 hours
- **Conditions:** Hypoxia (1% O2) vs Normoxia (21% O2)
- **Timepoint:** 72 hours
- **Biological Replicates:** 4 per condition

---

**Purpose:**

This notebook applies the ProteoForge pipeline to identify differential proteoforms (dPFs) between hypoxia and normoxia conditions at the 48-hour timepoint. The pipeline includes data preparation and normalization, weight calculation (not used for completeness), linear modeling, hierarchical clustering, and proteoform classification.

> **Note:** Focusing on a single timepoint allows us to capture proteoform changes specifically associated with hypoxia response, minimizing confounding effects from temporal dynamics. Additionally results from this will be applied to downstream statistical and functional analyses in `04-DownstreamComparison.ipynb`, when calculating dPF level protein quants.

---

#### 01. Setup¶

This section imports required libraries, configures display settings, and defines paths for data and figures.

> **Note:** The HTML rendering of this notebook hides code cells by default. Click the "Code" buttons on the right to expand them.

##### 01.1 Libraries¶

```
# Libraries Used

import os
import sys

import numpy as np # Numerical computing
import pandas as pd # Data manipulation

import seaborn as sns # R-like high-level plots
import matplotlib.pyplot as plt # Python's base plotting 

# home path
sys.path.append('../')
from src import utils, plots

# Functionality to Process the Data (Weight, Normalize, Model, Correlate, Cluster)
from ProteoForge import normalize, impute
# Functionality to Predict Discordant Peptides (id)
from ProteoForge import correct, weight, model
# Functionality to Cluster Peptides for Grouping (group)
from ProteoForge import corluster, disluster
# Functionality to Generate dPFs (differential proteoforms)
from ProteoForge import proteoform_classifier as pfc
# Initialize the timer
startTime = utils.getTime()
```

---

#### 02. ProteoForge Analysis Pipeline¶

This section shows how the ProteoForge pipeline is applied to the 72-hour hypoxia vs normoxia comparison, including data preparation, weight calculation, weighted linear modeling, hierarchical clustering, and proteoform classification.

##### 02.1 Prepare Analysis Data¶

Prepare the test dataset by merging peptide information with quantification data, subsetting to 72-hour timepoint conditions (Hypoxia vs Normoxia), normalizing intensities against the control condition, and tracking imputation status for downstream weighting.

```
test_data = info_data.sort_values(
    by=['Protein', 'peptide_start', 'peptide_end']
)[['unique_id', 'Protein', 'Peptide', 'peptide_start', 'peptide_end']]
# peptide_end and start should be int
test_data['peptide_start'] = test_data['peptide_start'].astype(int)
test_data['peptide_end'] = test_data['peptide_end'].astype(int)
# Generate PeptideID from cum
test_data['PeptideID'] = test_data.groupby('Protein').cumcount() + 1

print()
print("📊 MELTING CLEANED AND IMPUTED DATA FOR ANALYSIS")
# Get the imputed and cleaned (with missing values) data into long format
melt_imputed = imputed_data.reset_index().drop(
    columns='Protein'
).melt(
    id_vars='unique_id',
    var_name='Sample',
    value_name='Intensity'
).set_index(['unique_id', 'Sample'])
melt_cleaned = cleaned_data.reset_index().drop(
    columns='Protein'
).melt(
    id_vars='unique_id',
    var_name='Sample',
    value_name='Intensity(Raw)'
).set_index(['unique_id', 'Sample'])

combined_data = pd.concat(
    [melt_imputed, melt_cleaned],
    axis=1
).reset_index()

print()
print("📊 ADDITIONAL INFO ADDED")
combined_data['log10Intensity'] = np.log10(combined_data['Intensity'])
combined_data['Condition'] = combined_data['Sample'].map(samples_to_condition)
combined_data['isReal'] = combined_data['Intensity(Raw)'].notna()

compMiss_table = pd.DataFrame(index=cleaned_data.index)
for cond, col in condition_to_samples.items():
    compMiss_table[cond] = cleaned_data[col].isna().all(axis=1)

compMiss_table = compMiss_table.reset_index().drop(columns='Protein').melt(
    id_vars='unique_id',
    var_name='Condition',
    value_name='isCompleteMiss'
)
combined_data = combined_data.merge(
    compMiss_table,
    on=['unique_id', 'Condition'],
    how='left'
)
combined_data = combined_data[[
    'unique_id', 'Condition', 'Sample', 
    'isCompleteMiss', 'isReal',
    'Intensity(Raw)', 'Intensity', 'log10Intensity', 
]]

print()
print("📊 COMBINING INFO AND QUANT DATA")

# Merge the test_data with combined_data
test_data = test_data.merge(
    combined_data,
    on='unique_id',
    how='left'
)

# conditions to subset for 
conditions = [
    # # Removed the chemical and oxidative versions since they were not used in the manuscript
    # 'Oxidative Stress (H2O2) - 48 Hr',
    # 'Oxidative Stress (H2O2) - 72 Hr',
    # 'Chemical Hypoxia (CoCl2) - 48 Hr',
    # 'Chemical Hypoxia (CoCl2) - 72 Hr',
    # 'True Hypoxia (1% O2) - 48 Hr',
    'True Hypoxia (1% O2) - 72 Hr',
    # 'Normoxia (21% O2) - 48 Hr', # Keeping it simple with longest control
    'Normoxia (21% O2) - 72 Hr',
]

# Update the condition_to_samples to only include the conditions of interest
condition_to_samples = {
    cond: condition_to_samples[cond] for cond in conditions if cond in condition_to_samples
}

test_data = test_data[
    test_data['Condition'].isin(condition_to_samples.keys())
].reset_index(drop=True)

print()
print("📏 NORMALIZING INTENSITIES AGAINST CONTROL CONDITION")
test_data = normalize.against_condition(
    long_data=test_data,
    cond_run_dict=condition_to_samples,
    run_col='Sample',
    index_cols=['Protein', 'PeptideID'],
    norm_against='Normoxia (21% O2) - 72 Hr',
    intensity_col='Intensity',
    is_log2=False,
    norm_intensity_col='AdjIntensity'
)
print()
# print("🛡️ CALCULATING ROBUST INTENSITIES FOR CLUSTERING")
# test_data = normalize.build_robust_intensity(
#     test_data,
#     log_col='log10Intensity',
#     adj_col='AdjIntensity',
#     protein_col='Protein',
#     cond_col='Condition',
#     out_col='RobIntensity',
#     verbose=verbose

# )
print(f"Test Data shape after normalization: {test_data.shape}")
test_data.head()
```

```
📊 MELTING CLEANED AND IMPUTED DATA FOR ANALYSIS

📊 ADDITIONAL INFO ADDED

📊 COMBINING INFO AND QUANT DATA

📏 NORMALIZING INTENSITIES AGAINST CONTROL CONDITION

Test Data shape after normalization: (572214, 14)
```

|  | unique\_id | Protein | Peptide | peptide\_start | peptide\_end | PeptideID | Condition | Sample | isCompleteMiss | isReal | Intensity(Raw) | Intensity | log10Intensity | AdjIntensity |
| --- | --- | --- | --- | --- | --- | --- | --- | --- | --- | --- | --- | --- | --- | --- |
| 0 | A0A024RBG1-TYDREGFK-11.0 | A0A024RBG1 | TYDREGFK | 11 | 18 | 1 | True Hypoxia (1% O2) - 72 Hr | 1%\_72\_2 | False | False | NaN | 8083.9621 | 3.9076 | -0.1312 |
| 1 | A0A024RBG1-TYDREGFK-11.0 | A0A024RBG1 | TYDREGFK | 11 | 18 | 1 | True Hypoxia (1% O2) - 72 Hr | 1%\_72\_3 | False | True | 8961.8193 | 6812.1680 | 3.8333 | -0.2018 |
| 2 | A0A024RBG1-TYDREGFK-11.0 | A0A024RBG1 | TYDREGFK | 11 | 18 | 1 | True Hypoxia (1% O2) - 72 Hr | 1%\_72\_4 | False | True | 12107.9668 | 9593.1931 | 3.9820 | -0.0825 |
| 3 | A0A024RBG1-TYDREGFK-11.0 | A0A024RBG1 | TYDREGFK | 11 | 18 | 1 | Normoxia (21% O2) - 72 Hr | C\_72\_1 | False | True | 10629.0869 | 8248.7570 | 3.9164 | -0.1126 |
| 4 | A0A024RBG1-TYDREGFK-11.0 | A0A024RBG1 | TYDREGFK | 11 | 18 | 1 | Normoxia (21% O2) - 72 Hr | C\_72\_2 | False | True | 14232.6455 | 11734.2555 | 4.0695 | 0.0287 |

```
 📏 Calculating Imputation Weights... (done in 2.04 ms)
 📏 Calculating Reverse Technical Variation Weights... (done in 166.24 ms)
All weights calculated and normalized.
Test Data shape after weight generation: (572214, 15)
```

|  | unique\_id | Protein | Peptide | peptide\_start | peptide\_end | PeptideID | Condition | Sample | isCompleteMiss | isReal | Intensity(Raw) | Intensity | log10Intensity | AdjIntensity | Weight |
| --- | --- | --- | --- | --- | --- | --- | --- | --- | --- | --- | --- | --- | --- | --- | --- |
| 0 | A0A024RBG1-TYDREGFK-11.0 | A0A024RBG1 | TYDREGFK | 11 | 18 | 1 | True Hypoxia (1% O2) - 72 Hr | 1%\_72\_2 | False | False | NaN | 8083.9621 | 3.9076 | -0.1312 | 0.0999 |
| 1 | A0A024RBG1-TYDREGFK-11.0 | A0A024RBG1 | TYDREGFK | 11 | 18 | 1 | True Hypoxia (1% O2) - 72 Hr | 1%\_72\_3 | False | True | 8961.8193 | 6812.1680 | 3.8333 | -0.2018 | 0.9999 |
| 2 | A0A024RBG1-TYDREGFK-11.0 | A0A024RBG1 | TYDREGFK | 11 | 18 | 1 | True Hypoxia (1% O2) - 72 Hr | 1%\_72\_4 | False | True | 12107.9668 | 9593.1931 | 3.9820 | -0.0825 | 0.9999 |
| 3 | A0A024RBG1-TYDREGFK-11.0 | A0A024RBG1 | TYDREGFK | 11 | 18 | 1 | Normoxia (21% O2) - 72 Hr | C\_72\_1 | False | True | 10629.0869 | 8248.7570 | 3.9164 | -0.1126 | 0.9997 |
| 4 | A0A024RBG1-TYDREGFK-11.0 | A0A024RBG1 | TYDREGFK | 11 | 18 | 1 | Normoxia (21% O2) - 72 Hr | C\_72\_2 | False | True | 14232.6455 | 11734.2555 | 4.0695 | 0.0287 | 0.9997 |

```
pThr = 0.001
palette = {
    'pval': '#4c72b0',
    'prt_pval': '#dd8452',
    'adj_pval': '#55a868'
}
# --- 3. Plotting ---
# Define the columns to plot and their corresponding titles
plot_configs = {
    'pval': 'Raw p-value',
    'prt_pval': 'p-value Within Protein (bonferroni)',
    'adj_pval': 'Adjusted p-value (fdr_bh)'
}

# Calculate log10 p-values for each column, replacing zeros with minimum nonzero value for stability
for col in plot_configs.keys():
    min_nonzero = test_data[col][test_data[col] > 0].min() if (test_data[col] > 0).any() else 1e-300
    test_data[f'log10_{col}'] = np.log10(test_data[col].replace(0, min_nonzero))

fig, axes = plt.subplots(1, 3, figsize=(20, 7), sharex=True, sharey=True)

for ax, (col, title) in zip(axes, plot_configs.items()):
    significant_mask = test_data[col] < pThr
    significant_hits = significant_mask.sum()
    total_hits = test_data[col].count()
    percent_hits = 100 * significant_hits / total_hits if total_hits > 0 else 0
    # Proteins
    total_proteins = test_data['Protein'].nunique()
    significant_proteins = test_data.loc[significant_mask, 'Protein'].nunique()
    percent_proteins = 100 * significant_proteins / total_proteins if total_proteins > 0 else 0
    # Peptides
    total_peptides = test_data['Peptide'].nunique()
    significant_peptides = test_data.loc[significant_mask, 'Peptide'].nunique()
    percent_peptides = 100 * significant_peptides / total_peptides if total_peptides > 0 else 0

    # Drop NaN values for histogram
    log10_vals = test_data[f'log10_{col}'].dropna()
    if len(log10_vals) == 0:
        ax.text(0.5, 0.5, 'No data available', transform=ax.transAxes, ha='center', va='center', fontsize=14)
        continue

    counts, bins = np.histogram(log10_vals, bins=50)
    counts_log10 = np.log10(np.where(counts == 0, 1, counts))

    ax.bar(bins[:-1], counts_log10, width=np.diff(bins), align='edge', color=palette[col], edgecolor='white', linewidth=0.5, alpha=0.8)

    # Add vertical line for threshold
    x_thr = np.log10(pThr)
    ax.axvline(x=x_thr, color='crimson', linestyle='--', linewidth=2)

    # Add vertical annotation next to the line
    ylim = ax.get_ylim()
    ax.text(
        x_thr, ylim[1]*0.95, 
        f'log10={x_thr:.0f}', 
        color='crimson', rotation=90, 
        va='top', ha='right', 
        fontsize=12, fontweight='bold', 
        bbox=dict(boxstyle='round,pad=0.2', fc='white', ec='crimson', alpha=0.7)
    )

    annotation = (
        f'Hits: {significant_hits:,} / {total_hits:,} ({percent_hits:.2f}%)\n'
        f'Proteins: {significant_proteins:,} / {total_proteins:,} ({percent_proteins:.2f}%)\n'
        f'Peptides: {significant_peptides:,} / {total_peptides:,} ({percent_peptides:.2f}%)'
    )
    ax.text(
        0.05, 0.90,
        annotation,
        transform=ax.transAxes,
        fontsize=12, fontweight='bold',
        ha='left', va='top',
        bbox=dict(boxstyle='round,pad=0.3', fc='white', ec='black', alpha=0.75)
    )

    ax.set_title(title + ' (log10)', fontsize=16, fontweight='bold')
    ax.set_xlabel('log10(p-value)', fontsize=12)
    ax.set_ylabel('log10(Frequency)', fontsize=14)
    ax.grid(axis='y', linestyle='--', alpha=0.6)
    ax.tick_params(axis='x', rotation=45)

axes[1].set_ylabel('')
axes[2].set_ylabel('')

fig.suptitle(
    'Comparison of log10 p-value Distributions',
    fontsize=22,
    fontweight='bold',
    y=1.02
    )

sns.despine()
plt.tight_layout(rect=[0, 0, 1, 0.96])
plots.finalize_plot( 
    fig, show=True, save=save_to_folder, 
    filename='pvalue_distributions_72hr',
    filepath=figure_path,
    formats=figure_formats,
    transparent=transparent_bg,
    dpi=figure_dpi
)

test_data['isSignificant'] = test_data['adj_pval'] < pThr
test_data.head()
```

|  | unique\_id | Protein | Peptide | peptide\_start | peptide\_end | PeptideID | Condition | Sample | isCompleteMiss | isReal | Intensity(Raw) | Intensity | log10Intensity | AdjIntensity | Weight | pval | prt\_pval | adj\_pval | log10\_pval | log10\_prt\_pval | log10\_adj\_pval | isSignificant |
| --- | --- | --- | --- | --- | --- | --- | --- | --- | --- | --- | --- | --- | --- | --- | --- | --- | --- | --- | --- | --- | --- | --- |
| 0 | A0A024RBG1-TYDREGFK-11.0 | A0A024RBG1 | TYDREGFK | 11 | 18 | 1 | True Hypoxia (1% O2) - 72 Hr | 1%\_72\_2 | False | False | NaN | 8083.9621 | 3.9076 | -0.1312 | 0.0999 | 0.3949 | 1.0000 | 1.0000 | -0.4035 | 0.0000 | 0.0000 | False |
| 1 | A0A024RBG1-TYDREGFK-11.0 | A0A024RBG1 | TYDREGFK | 11 | 18 | 1 | True Hypoxia (1% O2) - 72 Hr | 1%\_72\_3 | False | True | 8961.8193 | 6812.1680 | 3.8333 | -0.2018 | 0.9999 | 0.3949 | 1.0000 | 1.0000 | -0.4035 | 0.0000 | 0.0000 | False |
| 2 | A0A024RBG1-TYDREGFK-11.0 | A0A024RBG1 | TYDREGFK | 11 | 18 | 1 | True Hypoxia (1% O2) - 72 Hr | 1%\_72\_4 | False | True | 12107.9668 | 9593.1931 | 3.9820 | -0.0825 | 0.9999 | 0.3949 | 1.0000 | 1.0000 | -0.4035 | 0.0000 | 0.0000 | False |
| 3 | A0A024RBG1-TYDREGFK-11.0 | A0A024RBG1 | TYDREGFK | 11 | 18 | 1 | Normoxia (21% O2) - 72 Hr | C\_72\_1 | False | True | 10629.0869 | 8248.7570 | 3.9164 | -0.1126 | 0.9997 | 0.3949 | 1.0000 | 1.0000 | -0.4035 | 0.0000 | 0.0000 | False |
| 4 | A0A024RBG1-TYDREGFK-11.0 | A0A024RBG1 | TYDREGFK | 11 | 18 | 1 | Normoxia (21% O2) - 72 Hr | C\_72\_2 | False | True | 14232.6455 | 11734.2555 | 4.0695 | 0.0287 | 0.9997 | 0.3949 | 1.0000 | 1.0000 | -0.4035 | 0.0000 | 0.0000 | False |

```
Processing 95,369 peptides from 7,161 proteins...
Auto-selected algorithm: ultra_fast
Ultra-fast algorithm: Processing 95369 peptides...
  ✓ Ultra-fast processing complete: maximum performance achieved

Processing completed in 0.579 seconds
Performance: 164,574 peptides/second

Classification Summary:
  Single PTMs (dPF=-1): 4,376
  Non-differential (dPF=0): 63,545
  Differential proteoforms (dPF>0): 27,448 peptides
  Maximum proteoform group: dPF=3
  Processed 40459 differential proteoforms (dPFs)
```

#### 03. Save Results¶

Export the complete analysis results for downstream interpretation and visualization. Output includes peptide-level data with statistics and clusters (`test_data_72hr.feather`), protein-level summary with dPF classifications (`summary_data_72hr.feather`), and UniProt annotations (`uniprot_data_72hr.feather`).

```
test_data.to_feather(f"{output_path}test_data_72hr.feather")
summary_data.to_feather(f"{output_path}summary_data_72hr.feather")
uniprot_data.to_feather(f"{output_path}uniprot_data_72hr.feather")
```

---

#### Summary¶

This notebook applied the ProteoForge pipeline to compare True Hypoxia (1% O₂) vs Normoxia (21% O₂) at 72 hours in H358 NSCLC cells. The analysis prepared peptide data with condition-normalized intensities, generated quality-aware sample weights, performed weighted linear modeling with two-step multiple testing correction, clustered peptides by abundance patterns, and identified proteins with differential proteoform regulation.

```
print("Notebook Execution Time:", utils.prettyTimer(utils.getTime() - startTime))
```

```
Notebook Execution Time: 00h:01m:50s
```
