## Supplementary material for "ProteoForge: An Imputation-Aware Framework for Differential Proteoform Discovery in Bottom-Up Proteomics": Analysis notebooks as html render: Notebook S12.html

### Notebook S12

###### Enes Kemal Ergin

#### 2025-12-08 23:00:22

### Supplementary Notebook 12: Exploring the ProteoForge Results from 48hr¶

- **License:** Creative Commons Attribution-NonCommercial 4.0 International License
- **Version:** 0.2
- **Edit Log:**
  - 2025-11-28: Initial version of the notebook
  - 2025-12-08: Revise the whole notebook, ensuring clarity and correctness

---

**Requirements:**

This notebook requires the `02-ApplyingProteoForge_48hr.ipynb` to be run first to generate the result files used here. Ensure the `./data/cleaned/` and `.data/results/` directories contains the necessary input files.

---

**Data Information:**

- Cell line: H358 (NSCLC)
- Conditions: True Hypoxia (1% O₂) at 48 Hr , Normoxia (21% O₂) at 48 Hr
- Input: `test_data_48hr.feather`, `summary_data_48hr.feather`, `uniprot_data_48hr.feather`
- Output: Visualizations and summary statistics for identified proteoforms and some example proteins of interest

---

**Purpose:**

This notebook summarizes and visualizes the results from the combined ProteoForge analysis of hypoxia vs normoxia conditions at 48-hour timepoint. It includes protein-level differential proteoform (dPF) analysis, peptide composition distributions, UniProt annotation mapping, and case studies of biologically relevant proteoforms.

---

#### 01. Setup¶

This section imports required libraries, configures display settings, and defines paths for data and figures.

> **Note:** The HTML rendering of this notebook hides code cells by default. Click the "Code" buttons on the right to expand them.

##### 01.1 Libraries¶

```
# Libraries Used
import os
import sys

import numpy as np # Numerical computing
import pandas as pd # Data manipulation

import seaborn as sns # R-like high-level plots
import matplotlib.pyplot as plt # Python's base plotting 

# home path
sys.path.append('../')
from src import utils, plots

from ProteoForge import annotate
from ProteoForge import proteoform_classifier as pfc

# Initialize the timer
startTime = utils.getTime()
```

##### 01.2 Configure Notebook Settings¶

Configure visualization styles, color palettes, and display options for consistent output formatting throughout the analysis.

```
## Configure Plotting Settings
def_colors = [
    "#139593", "#fca311", "#e54f2a",
    "#c3c3c3", "#555555",
    "#690000", "#5f4a00", "#004549"
]

condition_colors = {
    'True Hypoxia (1% O2) - 48 Hr': '#e9c46a',
    'True Hypoxia (1% O2) - 72 Hr': '#f4a261',
    'Normoxia (21% O2) - 48 Hr': '#ade8f4',
    'Normoxia (21% O2) - 72 Hr': '#0077b6',
    # Won't be used since the manuscript didn't cover these conditions
    # 'Chemical Hypoxia (CoCl2) - 48 Hr': '#c77dff',
    # 'Chemical Hypoxia (CoCl2) - 72 Hr': '#8338ec',
    # 'Oxidative Stress (H2O2) - 48 Hr': '#ff006e',
    # 'Oxidative Stress (H2O2) - 72 Hr': '#fb5607',
}

# Set seaborn style
sns.set_theme(
    style="white",
    context="paper",
    palette=def_colors,
    font_scale=1,
    rc={
        "figure.figsize": (6, 4),
        "font.family": "sans-serif",
        "font.sans-serif": ["Arial", "Ubuntu Mono"],
    }
)

# Figure Saving Settings
figure_formats = ["png", "pdf"]
save_to_folder = True
transparent_bg = True
figure_dpi = 300

## Configure dataframe displaying
pd.set_option('display.float_format', lambda x: '%.4f' % x)
pd.set_option('display.max_rows', 50)
pd.set_option('display.max_columns', 50)
pd.set_option('display.width', 500)  # Set a wider display width


## Printing Settings
verbose = True

plots.color_palette( def_colors, save=False )
plots.color_palette( condition_colors, save=False, name='condition', size=2)
```


##### 01.3 Data and Result Paths¶

- `home_path` — Base project path (e.g., `./`)
- `data_name` — Dataset identifier used throughout (`"hypoxia"`)
- `data_path` — Root data folder (`{home_path}data/`)
- `input_path` — Cleaned input data from preprocessing (`{data_path}cleaned/{data_name}/`)
- `output_path` — ProteoForge result files (`{data_path}results/{data_name}/`)
- `figure_path` — Destination for generated figures (`{home_path}figures/{data_name}/03-ResultsAnalysis/`)
- `notebook_name` — This notebook name for logging and organizing outputs (`"03-ResultsAnalysis"`)

```
home_path = './'
data_name = "hypoxia"
notebook_name = "03-ResultsAnalysis"
data_path = f"{home_path}data/"
input_path = f"{data_path}cleaned/{data_name}/"
output_path = f"{data_path}results/{data_name}/"
figure_path = f"{home_path}figures/{data_name}/{notebook_name}/"

# Create the output folder if it does not exist
if not os.path.exists(output_path):
    os.makedirs(output_path)

# Create figure folder structure, if needed
if save_to_folder:
    for i in figure_formats:
        cur_folder = figure_path + i + "/"
        if not os.path.exists(cur_folder):
            os.makedirs(cur_folder)
```

##### 01.4 Load Input Data¶

Load the ProteoForge analysis outputs and supporting data including UniProt annotations with feature metadata, sample metadata with condition mappings, test data with peptide-level results, and summary data with protein-level statistics.

**Uniprot Data**: Contains expanded annotations for proteins in the data. (`uniprot_data_48hr.feather`)

```
uniprot_data = pd.read_feather(f"{output_path}uniprot_data_48hr.feather")
uniprot_data = annotate.add_feature_metadata(uniprot_data, annotate.feature_categories, annotate.relevant_features)
print(f"UniProt data loaded with shape: {uniprot_data.shape}")
uniprot_data.info()
uniprot_data
```

```
UniProt data loaded with shape: (774552, 12)
<class 'pandas.core.frame.DataFrame'>
RangeIndex: 774552 entries, 0 to 774551
Data columns (total 12 columns):
 #   Column            Non-Null Count   Dtype 
---  ------            --------------   ----- 
 0   isoform           774552 non-null  string
 1   Protein           774552 non-null  string
 2   feature           774552 non-null  string
 3   start             774552 non-null  Int64 
 4   end               774552 non-null  Int64 
 5   group             774552 non-null  string
 6   agent             774552 non-null  string
 7   note              774552 non-null  string
 8   description       774552 non-null  string
 9   score             774552 non-null  int64 
 10  is_relevant       774552 non-null  bool  
 11  feature_category  774552 non-null  object
dtypes: Int64(2), bool(1), int64(1), object(1), string(7)
memory usage: 67.2+ MB
```

|  | isoform | Protein | feature | start | end | group | agent | note | description | score | is\_relevant | feature\_category |
| --- | --- | --- | --- | --- | --- | --- | --- | --- | --- | --- | --- | --- |
| 0 |  | A0A024RBG1 | CHAIN | 1 | 181 |  |  |  | Diphosphoinositol polyphosphate phosphohydrola... | 3 | False | Protein Processing & Maturation |
| 1 |  | A0A024RBG1 | CROSSLNK | 5 | 5 | Ubiquitination/SUMOylation |  | Ubiquitination/SUMOylation at K5 | Source: psp; Type: UBIQUITINATION | 1 | True | Co- & Post-Translational Modifications |
| 2 |  | A0A024RBG1 | BINDING | 10 | 10 |  |  |  |  | 3 | False | Functional Domains, Regions & Sites |
| 3 |  | A0A024RBG1 | BINDING | 18 | 20 |  |  |  |  | 3 | False | Functional Domains, Regions & Sites |
| 4 |  | A0A024RBG1 | DOMAIN | 18 | 145 |  |  |  | Nudix hydrolase | 3 | False | Functional Domains, Regions & Sites |
| ... | ... | ... | ... | ... | ... | ... | ... | ... | ... | ... | ... | ... |
| 774547 |  | Q9Y6Y8 | MOD\_RES | 930 | 930 | Phosphorylation |  | Phosphorylation at S930 | Source: psp; Type: PHOSPHORYLATION | 1 | True | Co- & Post-Translational Modifications |
| 774548 |  | Q9Y6Y8 | CROSSLNK | 931 | 931 | Ubiquitination/SUMOylation |  | Ubiquitination/SUMOylation at K931 | Source: psp; Type: UBIQUITINATION | 1 | True | Co- & Post-Translational Modifications |
| 774549 |  | Q9Y6Y8 | MOD\_RES | 935 | 935 | Phosphorylation |  | Phosphorylation at Y935 | Source: psp; Type: PHOSPHORYLATION | 1 | True | Co- & Post-Translational Modifications |
| 774550 |  | Q9Y6Y8 | CROSSLNK | 938 | 938 | Ubiquitination/SUMOylation |  | Ubiquitination/SUMOylation at K938 | Source: psp; Type: UBIQUITINATION | 1 | True | Co- & Post-Translational Modifications |
| 774551 |  | Q9Y6Y8 | CLEAVAGE | 989 | 989 | Proteolytic Cleavage | trypsin 1 | N-Ph cleavage at R989 by trypsin 1 | MEROPS Cleavage Site; Protease: trypsin 1 (MER... | 2 | True | Protein Processing & Maturation |

774552 rows × 12 columns

**Metadata**: Sample metadata with condition mappings. (`metadata.csv`). Preview (first 5 rows):

```
meta = pd.read_csv(f"{input_path}metadata.csv")
# Build a dictionary to match condition_colors keys: 'Condition - Duration Hr'
condition_to_samples = {}
for condition in meta['Condition'].unique():
    for duration in sorted(meta.loc[meta['Condition'] == condition, 'Duration'].unique()):
        key = f"{condition} - {duration} Hr"
        samples = meta.loc[(meta['Condition'] == condition) & (meta['Duration'] == duration), 'Colname'].tolist()
        condition_to_samples[key] = samples

# samples_to_condition mapping
samples_to_condition = {sample: condition for condition, samples in condition_to_samples.items() for sample in samples}
# samples_to_colors mapping
samples_to_colors = {sample: condition_colors[condition] for sample, condition in samples_to_condition.items()}
meta.head(5)
```

|  | filename | Run | Condition | Replicate | Duration | Colname |
| --- | --- | --- | --- | --- | --- | --- |
| 0 | D:\Data\Tam\RevisionLungDIA\H358\H358\_1%\_1\_48\_... | H358\_1%\_1\_48\_DIA\_BB4\_1\_5432 | True Hypoxia (1% O2) | 1 | 48 | 1%\_48\_1 |
| 1 | D:\Data\Tam\RevisionLungDIA\H358\H358\_1%\_2\_48\_... | H358\_1%\_2\_48\_DIA\_BC4\_1\_5440 | True Hypoxia (1% O2) | 2 | 48 | 1%\_48\_2 |
| 2 | D:\Data\Tam\RevisionLungDIA\H358\H358\_1%\_3\_48\_... | H358\_1%\_3\_48\_DIA\_BD4\_1\_5448 | True Hypoxia (1% O2) | 3 | 48 | 1%\_48\_3 |
| 3 | D:\Data\Tam\RevisionLungDIA\H358\H358\_1%\_4\_48\_... | H358\_1%\_4\_48\_DIA\_BE4\_1\_5456 | True Hypoxia (1% O2) | 4 | 48 | 1%\_48\_4 |
| 4 | D:\Data\Tam\RevisionLungDIA\H358\H358\_1%\_2\_72\_... | H358\_1%\_2\_72\_DIA\_BC8\_1\_5444 | True Hypoxia (1% O2) | 2 | 72 | 1%\_72\_2 |

###### Setting Up the Data¶

Here we load all the result files `test_data_48hr.feather`, `summary_data_48hr.feather`, and `info_data.feather` generated from the 48hr ProteoForge analysis in the previous notebook. As well as the `protein_data.feather` which will help with protein level informations (expanded Fasta).

**Peptide Regulation Across Conditions**

One thing that we can look is the regulation compared to control condition (Normoxia 21% O2 at 48hr).

```
pThr = 10**-3

test_data = pd.read_feather(f"{output_path}test_data_48hr.feather")
print(test_data.shape)
print()


summary_data = pd.read_feather(f"{output_path}summary_data_48hr.feather")
print(summary_data.shape)
print()

info_data = pd.read_feather(f"{input_path}info_data.feather")
print(info_data.shape)
print()

summary_data['trace'] = info_data['trace']

protein_data = pd.read_feather(f"{input_path}protein_data.feather")
print(protein_data.shape)
print()


## Adding Hypoxia Regulation compared to Normoxia 

df =  test_data.copy()

# --- 2. Define Control and Treatment Conditions ---
control_condition = 'Normoxia (21% O2) - 48 Hr'
# Sorting ensures the regulation string (e.g., 'Up_Down') is always in a consistent order.
treatment_conditions = sorted([cond for cond in df['Condition'].unique() if cond != control_condition])

print(f"Control: {control_condition}")
print(f"Treatments (in order): {treatment_conditions}")

# --- 3. Calculate Mean Intensity for Each Peptide in Each Condition ---
# This aggregates the technical replicates for each condition.
# The mean of AdjIntensity for a treatment is its log2FC vs. control.
# Using .agg is slightly more explicit and can be faster than .mean()
mean_intensities = df.groupby(['Protein', 'PeptideID', 'Condition'], as_index=False).agg(log2FC=('AdjIntensity', 'mean'))


# --- 4. Create the Regulation Table ---
# We can filter out the control condition, as its regulation is baseline (0).
regulation_df = mean_intensities[mean_intensities['Condition'] != control_condition].copy()

# Calculate the linear Fold Change from the log2FC.
# FoldChange = 2^(log2FC)
regulation_df['FoldChange'] = 2**regulation_df['log2FC']


# --- 5. Categorize Regulation Tendencies (Optimized) ---
# To categorize tendencies, we first pivot the table to have conditions as columns.
# This allows us to see the behavior of one peptide across all conditions in a single row.
pivot_reg = regulation_df.pivot_table(
    index=['Protein', 'PeptideID'],
    columns='Condition',
    values='log2FC'
).fillna(0) # Fill peptides not found in a condition with 0 log2FC (no change)

# This vectorized approach is much faster than using .apply()
# It builds the regulation string (e.g., 'Up_Down') column by column.
regulation_strings = None
for i, cond in enumerate(treatment_conditions):
    # Use np.select for fast conditional categorization based on direction
    categories = np.select(
        [pivot_reg[cond] > 0, pivot_reg[cond] < 0],
        ['Up', 'Down'],
        default='NC' # No Change
    )
    
    # Convert to a pandas Series to align with the pivot_reg index
    categories_series = pd.Series(categories, index=pivot_reg.index)

    if i == 0:
        regulation_strings = categories_series
    else:
        # Append the category for the next condition
        regulation_strings += '_' + categories_series

pivot_reg['Regulation'] = regulation_strings


# Now, merge the 'Regulation' column back into our long-format table
# This adds the summary string to each row corresponding to a given peptide.
regulation_df = regulation_df.merge(
    pivot_reg[['Regulation']],
    on=['Protein', 'PeptideID'],
    how='left'
)

# Reorder columns for a clean final output
regulation_df = regulation_df[['Protein', 'PeptideID', 'Condition', 'log2FC', 'FoldChange', 'Regulation']]


print("\n--- Final Regulation Table with Tendencies ---")
print(regulation_df.head())

# You can now save this final dataframe to a new file, for example:
# regulation_df.to_csv('regulation_analysis_output.csv', index=False)

# Example of how to use the new 'Regulation' column:
print("\n--- Example: Grouping by Regulation Tendency ---")
print(regulation_df.groupby('Regulation')['PeptideID'].nunique())

plot_data = regulation_df[['Protein', 'PeptideID', 'Regulation']].drop_duplicates().copy()

# Adding the plot_data to summary_data to get the Regulation column (use hypoxia 48hr-72hr)
summary_data = summary_data.merge(
    plot_data[['Protein', 'PeptideID', 'Regulation']],
    on=['Protein', 'PeptideID'],
    how='left'
)
summary_data['Regulation'] = summary_data['Regulation'].fillna('NC_NC')  # peptides not in regulation_df are 'No Change' in both conditions
summary_data.rename(columns={'Regulation': 'Regulation_(Hypoxia 48hr-72hr)'}, inplace=True)

fig, ax = plt.subplots(figsize=(8, 6))
plot_data['Regulation'].value_counts().plot(kind='barh', color="#432c46", edgecolor='black', ax=ax)
# Add counts to on top of the bars for barh
for i, v in enumerate(plot_data['Regulation'].value_counts()):
    ax.text(
        v, i, str(v),
        color='black', 
        ha='left', va='center', 
        rotation=0, fontsize=15, fontweight='bold'
    )
ax.set_xlabel("Number of Peptides", fontsize=12)
ax.set_ylabel("Regulation Tendency (Hypoxia 48Hr)", fontsize=12)
ax.set_title("Peptide Regulation Tendencies Across Conditions", fontsize=14, fontweight='bold')
ax.grid("both", linestyle="--", linewidth=0.75, alpha=0.5, color="lightgrey")
plt.tight_layout()

plots.finalize_plot( 
    fig, show=True, save=save_to_folder, 
    filename="peptide_regulation_tendencies_48hr",
    filepath=figure_path,
    formats=figure_formats,
    transparent=transparent_bg,
    dpi=figure_dpi
)

print()
summary_data['Regulation_(Hypoxia 48hr-72hr)'].value_counts()
```

```
(762952, 24)

(95369, 17)

(95369, 15)

(7161, 8)

Control: Normoxia (21% O2) - 48 Hr
Treatments (in order): ['True Hypoxia (1% O2) - 48 Hr']

--- Final Regulation Table with Tendencies ---
      Protein  PeptideID                     Condition  log2FC  FoldChange Regulation
0  A0A024RBG1          1  True Hypoxia (1% O2) - 48 Hr -0.0325      0.9777       Down
1  A0A024RBG1          2  True Hypoxia (1% O2) - 48 Hr -0.0539      0.9633       Down
2  A0A024RBG1          3  True Hypoxia (1% O2) - 48 Hr  0.0537      1.0379         Up
3  A0A024RBG1          4  True Hypoxia (1% O2) - 48 Hr -0.0230      0.9842       Down
4  A0A024RBG1          5  True Hypoxia (1% O2) - 48 Hr -0.1301      0.9138       Down

--- Example: Grouping by Regulation Tendency ---
Regulation
Down    202
Up      390
Name: PeptideID, dtype: int64
```

```
Regulation_(Hypoxia 48hr-72hr)
Up      48690
Down    46679
Name: count, dtype: int64
```

The regulation compared to the Normoxia per peptide is pretty even, with Hypoxia upregulation being slightly higher than downregulation.

---

#### 02. Protein-Level Analysis¶

This section generates protein-level summaries and visualizations of dPF discovery, including coverage relationships, peptide composition distributions, and proteoform category breakdowns.

```
# Create protein-level summary with dPF categories (vectorized, fast pandas)
# Uses indicator columns and a single groupby.agg to avoid per-group Python callbacks.

# local alias
df = summary_data.copy()

# Ensure numeric/bool-like columns use compact dtypes
if df['isSignificant'].dtype != 'int32' and df['isSignificant'].dtype != 'int64':
    df['isSignificant'] = df['isSignificant'].fillna(False).astype('int32')

# Temporary indicator columns (fast to sum)
df['is_canonical'] = (df['dPF'] == 0).astype('int32')
df['is_ptm'] = (df['dPF'] == -1).astype('int32')
df['is_proteoform'] = (df['dPF'] > 0).astype('int32')

# Perform one grouped aggregation (all C-level operations)
protein_dpf_analysis = (
    df.groupby('Protein', sort=False, observed=True)
      .agg(
          Gene = ('Gene', 'first'),
          Length = ('Length', 'first'),
          Coverage = ('Coverage', 'first'),
          all_pep = ('isSignificant', 'count'),
          sgnf_pep = ('isSignificant', 'sum'),
          all_clust = ('ClusterID', 'max'),
          n_forms = ('dPF', 'nunique'),
          n_canon = ('is_canonical', 'sum'),
          n_ptm = ('is_ptm', 'sum'),
          n_dPF = ('is_proteoform', 'sum'),
          max_dpf = ('dPF', 'max'),
      )
)

# Convert max_dpf -> max_proteoform_id: only positive dPFs indicate proteoform ids
protein_dpf_analysis['n_forms'] = (
    protein_dpf_analysis['max_dpf']
      .fillna(0)
      .where(protein_dpf_analysis['max_dpf'] > 0, 0)
      .astype('int32')
)
protein_dpf_analysis.drop(columns=['max_dpf'], inplace=True)

# If other code expects a flattened/explicit column order, optionally reorder here.
# Cleanup temporary indicator columns to free memory
try:
    df.drop(columns=['is_canonical', 'is_ptm', 'is_proteoform'], inplace=True)
except Exception:
    pass

# Add population rates for peptides, canonical, PTM, and proteoforms
protein_dpf_analysis['sgnf_pct'] = (
    protein_dpf_analysis['sgnf_pep'] / protein_dpf_analysis['all_pep']
).fillna(0).astype('float32') * 100.0 
protein_dpf_analysis['canon_pct'] = (
    protein_dpf_analysis['n_canon'] / protein_dpf_analysis['all_pep']
).fillna(0).astype('float32') * 100.0 
protein_dpf_analysis['ptm_pct'] = (
    protein_dpf_analysis['n_ptm'] / protein_dpf_analysis['all_pep']
).fillna(0).astype('float32') * 100.0 
protein_dpf_analysis['dPF_pct'] = (
    protein_dpf_analysis['n_dPF'] / protein_dpf_analysis['all_pep']
).fillna(0).astype('float32') * 100.0 
protein_dpf_analysis['eff_pct'] = (
    protein_dpf_analysis['n_forms'] / protein_dpf_analysis['all_clust']
).fillna(0).astype('float32') * 100.0

# Categorize coverage
protein_dpf_analysis['cov_cat'] = pd.cut(
    protein_dpf_analysis['Coverage'],
    bins=[-1, 25, 50, 75, 100],
    labels=['Low', 'Medium', 'High', 'V.High'],
    include_lowest=True
)
# Categorize available significant peptides per protein
protein_dpf_analysis['sgnf_cat'] = pd.cut(
    protein_dpf_analysis['sgnf_pep'],
    bins=[-1, 0, 1, np.inf],
    labels=['None', 'Single', 'Multiple'],
    right=True
)
# Categorize protein length
protein_dpf_analysis['len_cat'] = pd.cut(
    protein_dpf_analysis['Length'],
    bins=[-1, 150, 250, 750, 1000, 2000, np.inf],
    labels=['V.Short', 'Short', 'Medium', 'Long', 'V.Long', 'Huge'],
    right=True
)


# Categorize dPFs and PTMs containment per protein
labels = ['No Forms', 'PTM Only', 'dPF Only', 'Both']

conds = [
    (protein_dpf_analysis['n_forms'] == 0) & (protein_dpf_analysis['n_ptm'] == 0),
    (protein_dpf_analysis['n_forms'] == 0) & (protein_dpf_analysis['n_ptm'] > 0),
    (protein_dpf_analysis['n_forms'] > 0) & (protein_dpf_analysis['n_ptm'] == 0),
    (protein_dpf_analysis['n_forms'] > 0) & (protein_dpf_analysis['n_ptm'] > 0),
]

protein_dpf_analysis['dpf_cat'] = pd.Categorical(
    np.select(conds, labels, default='No Forms'),
    categories=labels
)

# protein_dpf_analysis is now ready with ptm_count, proteoform_count, max_proteoform_id, etc.
protein_dpf_analysis = protein_dpf_analysis.reset_index()
protein_dpf_analysis
```

|  | Protein | Gene | Length | Coverage | all\_pep | sgnf\_pep | all\_clust | n\_forms | n\_canon | n\_ptm | n\_dPF | sgnf\_pct | canon\_pct | ptm\_pct | dPF\_pct | eff\_pct | cov\_cat | sgnf\_cat | len\_cat | dpf\_cat |
| --- | --- | --- | --- | --- | --- | --- | --- | --- | --- | --- | --- | --- | --- | --- | --- | --- | --- | --- | --- | --- |
| 0 | A0A024RBG1 | NUDT4B | 181 | 30.3867 | 5 | 0 | 2 | 0 | 5 | 0 | 0 | 0.0000 | 100.0000 | 0.0000 | 0.0000 | 0.0000 | Medium | None | Short | No Forms |
| 1 | A0A0B4J2D5 | GATD3B | 268 | 35.4478 | 7 | 0 | 2 | 0 | 7 | 0 | 0 | 0.0000 | 100.0000 | 0.0000 | 0.0000 | 0.0000 | Medium | None | Medium | No Forms |
| 2 | A0A1B0GTU1 | ZC3H11B | 805 | 8.1988 | 5 | 1 | 2 | 0 | 4 | 1 | 0 | 20.0000 | 80.0000 | 20.0000 | 0.0000 | 0.0000 | Low | Single | Long | PTM Only |
| 3 | A0A1W2PQL4 | ZNF722 | 384 | 11.7188 | 5 | 0 | 3 | 0 | 5 | 0 | 0 | 0.0000 | 100.0000 | 0.0000 | 0.0000 | 0.0000 | Low | None | Medium | No Forms |
| 4 | A0A2R8Y4L2 | HNRNPA1L3 | 275 | 44.7273 | 12 | 1 | 2 | 0 | 11 | 1 | 0 | 8.3333 | 91.6667 | 8.3333 | 0.0000 | 0.0000 | Medium | Single | Medium | PTM Only |
| ... | ... | ... | ... | ... | ... | ... | ... | ... | ... | ... | ... | ... | ... | ... | ... | ... | ... | ... | ... | ... |
| 7156 | Q9Y6X5 | ENPP4 | 453 | 19.8675 | 7 | 0 | 3 | 0 | 7 | 0 | 0 | 0.0000 | 100.0000 | 0.0000 | 0.0000 | 0.0000 | Low | None | Medium | No Forms |
| 7157 | Q9Y6X8 | ZHX2 | 837 | 33.0944 | 22 | 2 | 3 | 0 | 20 | 2 | 0 | 9.0909 | 90.9091 | 9.0909 | 0.0000 | 0.0000 | Medium | Multiple | Long | PTM Only |
| 7158 | Q9Y6X9 | MORC2 | 1032 | 27.2287 | 23 | 2 | 3 | 0 | 21 | 2 | 0 | 8.6957 | 91.3044 | 8.6957 | 0.0000 | 0.0000 | Medium | Multiple | V.Long | PTM Only |
| 7159 | Q9Y6Y0 | IVNS1ABP | 642 | 11.0592 | 5 | 0 | 2 | 0 | 5 | 0 | 0 | 0.0000 | 100.0000 | 0.0000 | 0.0000 | 0.0000 | Low | None | Medium | No Forms |
| 7160 | Q9Y6Y8 | SEC23IP | 1000 | 22.7000 | 17 | 0 | 3 | 0 | 17 | 0 | 0 | 0.0000 | 100.0000 | 0.0000 | 0.0000 | 0.0000 | Low | None | Long | No Forms |

7161 rows × 20 columns

Protein-level dPF analysis table constructed with counts of canonical peptides, PTMs, and proteoforms per protein. Categories include coverage percentages, significance rates, and proteoform efficiency metrics. Essentially giving a overall summary of how many peptides with ptms, or apperant dPFs exists per protein along with their percentage of prevalance.

**Column Definitions:**

- **all\_pep**: Total number of peptides observed for the protein (count of peptide rows).
- **sgnf\_pep**: Number of peptides called significant (sum of isSignificant).
- **all\_clust**: Max ClusterID for the protein (proxy for number of peptide clusters; use unique count of ClusterID if IDs are non-contiguous).
- **n\_forms**: Number of proteoforms for the protein (positive dPF values).
- **n\_canon**: Count of canonical peptides (dPF == 0).
- **n\_ptm**: Count of PTM-only peptides (dPF == -1).
- **n\_dPF**: Count of peptides assigned to proteoforms (dPF > 0).
- **sgnf\_pct**: Percentage of peptides that are significant: (sgnf\_pep / all\_pep) \* 100.
- **canon\_pct**: Percentage of canonical peptides: (n\_canon / all\_pep) \* 100.
- **ptm\_pct**: Percentage of PTM-only peptides: (n\_ptm / all\_pep) \* 100.
- **dPF\_pct**: Percentage of peptides assigned to proteoforms: (n\_dPF / all\_pep) \* 100.
- **eff\_pct**: Proteoform discovery efficiency: (n\_forms / all\_clust) \* 100 — fraction of clusters yielding proteoforms.
- **cov\_cat**: Coverage category based on `Coverage` (Low, Medium, High, V.High).
- **sgnf\_cat**: Significant-peptide category based on `sgnf_pep` (None, Single, Multiple).
- **len\_cat**: Length category derived from `Length` (V.Short, Short, Medium, Long, V.Long, Huge).
- **dpf\_cat**: Proteoform/PTM containment category: No Forms, PTM Only, dPF Only, or Both.

Notes:

- dPF conventions: dPF == 0 → canonical peptide; dPF == -1 → PTM-only; dPF > 0 → proteoform ID.
- Percent columns are float percentages in range 0–100.
- `all_clust` uses `max(ClusterID)` as the cluster count proxy; if your clustering IDs are not contiguous starting at 1, use an explicit unique count of ClusterID if you want exact cluster counts.

---

##### 02.1 Significance by Coverage and Proteoforms¶

Heatmap showing the relationship between protein coverage categories (Low, Medium, High, V.High) and number of proteoforms discovered per protein (max=4). Average significance percentage indicates how coverage and proteoform count relate to detection of significant peptides within proteins. Average significance (sgnf\_pct) calculated as `(sgnf_pep / all_pep) * 100`.

```
# Heatmap of average sgnf_pct by coverage category vs number of proteoforms (n_forms)
heatmap_data = protein_dpf_analysis.pivot_table(
    values='sgnf_pct',
    index='cov_cat',
    columns='n_forms',
    aggfunc='mean',
    observed=True,
)

# Ensure sensible ordering of coverage categories if categorical
if isinstance(protein_dpf_analysis['cov_cat'].dtype, pd.CategoricalDtype):
    heatmap_data = heatmap_data.reindex(protein_dpf_analysis['cov_cat'].cat.categories)

# Sort columns (n_forms) ascending
heatmap_data = heatmap_data.reindex(sorted(heatmap_data.columns), axis=1)

plt.figure(figsize=(10, 6))
sns.heatmap(
    heatmap_data,
    annot=True,
    fmt=".1f",
    cmap="YlGnBu",
    cbar_kws={'label': 'Average sgnf_pct (%)'},
    linewidths=0.5,
    linecolor='white',
    mask=heatmap_data.isna()
)
plt.title('Average Significance Percentage (sgnf_pct) by Coverage Category vs Number of Proteoforms')
plt.ylabel('Coverage Category')
plt.xlabel('Number of Proteoforms (n_forms)')
plt.tight_layout()
plots.finalize_plot( 
    fig, show=True, save=save_to_folder, 
    filename='heatmap_sgnf_pct_cov_cat_vs_n_forms_48hr',
    filepath=figure_path,
    formats=figure_formats,
    transparent=transparent_bg,
    dpi=figure_dpi
)
```

Low coverage proteins tend to give more average significance percentage compared to higher coverage proteins. This may be due to lower coverage proteins having fewer peptides overall, making it easier for a higher fraction to be significant. In contrast, high coverage proteins may have many peptides, diluting the fraction that are significant. Additionally, the highest dPF counts (3) are only detected in proteins with Low to High (decreasing) coverage, which likely indicates very high coverage proteins are dominated by canonical and PTM peptides rather than diverse proteoforms, or that number of amino-acids and directly the number of peptides is smaller in High and V.High coverage proteins. Making them easier to cover fully with fewer proteoforms.

---

##### 02.2 Peptide Composition by Coverage¶

Stacked area plot showing the distribution of peptide types (canonical, PTM, and proteoform) across proteins sorted by coverage. The smoothed coverage trend line reveals how peptide composition varies with detection depth.

```
# Plot Composition of Proteins by dPF Categories
# This plot shows the distribution of canonical, PTM, and proteoform peptides across proteins
# with a smoothed coverage trend line.
custom_colors = {
    'Canonical': "#e5e5e5",      # Gray - baseline
    'PTM': "#a7c957",            # Red - modifications  
    'Proteoform': "#bc4749",     # Blue - true proteoforms
}

plot_data = protein_dpf_analysis.copy()
proteins_sorted = plot_data.sort_values('Coverage').reset_index(drop=True)
n_proteins = len(proteins_sorted)

# If many proteins, aggregate into bins (keeps trend readable)
# tune: lower -> more smoothing, higher -> more detail
max_bins = 100 
if n_proteins > max_bins:
    # group into evenly spaced bins
    proteins_sorted['bin'] = pd.cut(np.arange(n_proteins), bins=max_bins, labels=False)
    agg = proteins_sorted.groupby('bin').agg({
        'canon_pct':'mean',
        'ptm_pct':'mean',
        'dPF_pct':'mean',
        'Coverage':'mean'
    }).reset_index(drop=True)
    x = np.arange(len(agg))
    canonical = agg['canon_pct'].values
    ptm = agg['ptm_pct'].values
    proteoform = agg['dPF_pct'].values
    coverage = agg['Coverage'].values
    xtick_locs = np.linspace(0, len(agg)-1, 5).astype(int)
    xtick_labels = [f"{int(i/len(agg)*100)}%" for i in xtick_locs]  # percent position
else:
    x = np.arange(n_proteins)
    canonical = proteins_sorted['canon_pct'].values
    ptm = proteins_sorted['ptm_pct'].values
    proteoform = proteins_sorted['dPF_pct'].values
    coverage = proteins_sorted['Coverage'].values
    xtick_locs = np.linspace(0, n_proteins-1, 5).astype(int)
    xtick_labels = [f"{int(i/ n_proteins * 100)}%" for i in xtick_locs]

# Compute overall means for legend
mean_c = canonical.mean()
mean_ptm = ptm.mean()
mean_pf = proteoform.mean()

fig, ax = plt.subplots(figsize=(12, 5))
# stackplot handles the stacking cleanly
ax.stackplot(
    x,
    canonical, ptm, proteoform,
    labels=[f"Canonical ({mean_c:.1f}%)",
            f"PTMs ({mean_ptm:.1f}%)",
            f"Proteoforms ({mean_pf:.1f}%)"],
    colors=[custom_colors['Canonical'], custom_colors['PTM'], custom_colors['Proteoform']],
    alpha=0.85
)

# Coverage trend on twin axis (smoothed)
ax2 = ax.twinx()
window = max(3, int(len(coverage) * 0.02))  # ~2% window, min 3
coverage_smooth = pd.Series(coverage).rolling(window=window, center=True, min_periods=1).mean().values
ax2.plot(x, coverage_smooth, color='k', linewidth=1.6, alpha=0.9, label='Coverage (smoothed)')
ax2.set_ylim(0, 100)
ax.set_xlim(x.min(), x.max())

# Tidy up ticks/labels
ax.set_xlabel(f'{plot_data.shape[0]} Proteins (sorted by coverage)', fontsize=12, fontweight='bold')
ax.set_ylabel('Peptide composition (%)', fontsize=12, fontweight='bold')
ax2.set_ylabel('Coverage (%)', fontsize=12, fontweight='bold')
ax.set_xticks(xtick_locs)
ax.set_xticklabels(xtick_labels)
ax.set_ylim(0, max( (canonical + ptm + proteoform).max(), 100))  # sensible y-limits

# Legend and annotation
leg = ax.legend(
    loc='lower right', fontsize=10, frameon=False, bbox_to_anchor=(1.0, 0.1), 
    title='Peptide Composition', title_fontsize='12', ncol=1
)
ax2.legend(loc='lower right', fontsize=10, frameon=False, bbox_to_anchor=(1.0, 0))

# Add light grid and reduce spines clutter
ax.grid(axis='y', linestyle='--', linewidth=0.6, alpha=0.75, color='gray')

plt.tight_layout()
plots.finalize_plot( 
    fig, show=True, save=save_to_folder, 
    filename='peptideComposition_proteinCoverage_48hr',
    filepath=figure_path,
    formats=figure_formats,
    transparent=transparent_bg,
    dpi=figure_dpi
)
```

Most peptides across all proteins are canonical, with PTM and proteoform peptides comprising smaller fractions. However, as protein coverage increases, the relative proportion of canonical peptides slightly goes doewn, while Proteoforms contribution increases. Looking between PTM and Proteoform, lower coverage tends to have a higher fraction of PTM peptides compared to Proteoforms, while at higher coverage levels Proteoforms make up a larger share than PTMs.

---

##### 02.3 Proteoform Discovery Landscape¶

Bubble plot visualizing the relationship between peptide count and proteoform discovery per protein. Bubble size represents protein length and color indicates coverage, revealing patterns in how protein characteristics influence proteoform detection.

```
# Plot the Efficiency of Proteoform Discovery
# X: Number of Peptides per Protein
# Y: Number of Proteoforms Discovered (non-)
# Size: Protein Length (bubble size)
# Color: Coverage
# Alpha: Coverage

from matplotlib.lines import Line2D

plot_data = protein_dpf_analysis.copy()
# plot_data['discovery_efficiency'] = plot_data['n_dPF'] / plot_data['all_pep'] # dPF_pct
# plot_data['significance_efficiency'] = plot_data['n_sgnf'] / plot_data['all_pep'] # sgnf_pct
plot_data['complexity_score'] = (plot_data['n_forms'] * plot_data['all_clust']) / (plot_data['Length'] / 100)
plot_data['complexity_score'] = plot_data['complexity_score'].fillna(0)

# Bubble size scaling (marker area)
length = plot_data['Length'].clip(lower=1)
min_area, max_area = 40, 1600
areas = np.interp(length, (length.min(), length.max()), (min_area, max_area))

# Coverage -> color and per-point alpha
coverage = plot_data['Coverage'].fillna(0).astype(float)
# cov_min, cov_max = coverage.min(), coverage.max()
cov_min, cov_max = 0, 100  # assuming coverage is in percentage
if cov_max == cov_min:
    cov_max = cov_min + 1e-6  # avoid zero-range

# map coverage to alpha range (low coverage -> lower alpha)
alpha_min, alpha_max = 0.30, 0.95
alphas = np.interp(coverage, (cov_min, cov_max), (alpha_min, alpha_max))

# build RGBA colors from a cmap, then set per-point alpha
cmap = plt.cm.RdYlBu_r
norm = plt.Normalize(vmin=cov_min, vmax=cov_max)
base_colors = cmap(norm(coverage))  # Nx4 array
base_colors[:, 3] = alphas         # set alpha channel per-point

# Plot
x = plot_data['all_pep'].values
y = plot_data['n_forms'].values

fig, ax = plt.subplots(figsize=(10, 6))

sc = ax.scatter(
    x, y,
    s=areas,
    c=base_colors,
    edgecolors='white',
    linewidth=0.8,
    zorder=2
)

# Colorbar for Coverage (note: alpha not reflected in bar)
mappable = plt.cm.ScalarMappable(norm=norm, cmap=cmap)
mappable.set_array(coverage)
cbar = fig.colorbar(mappable, ax=ax, pad=0.02, shrink=0.9)
cbar.set_label('Coverage (%)', fontsize=11, fontweight='bold')

# Size legend (approximate protein lengths)
lens = np.percentile(length, [10, 50, 90])
proxy_areas = np.interp(lens, (length.min(), length.max()), (min_area, max_area))
legend_proxies = [
    Line2D([0], [0], marker='o', color='w', label=f'Len ≈ {int(lens[0])}', markerfacecolor='gray', markersize=np.sqrt(proxy_areas[0])),
    Line2D([0], [0], marker='o', color='w', label=f'Len ≈ {int(lens[1])}', markerfacecolor='gray', markersize=np.sqrt(proxy_areas[1])),
    Line2D([0], [0], marker='o', color='w', label=f'Len ≈ {int(lens[2])}', markerfacecolor='gray', markersize=np.sqrt(proxy_areas[2]))
]

# Legends: exceptional + size proxies
main_handles, main_labels = ax.get_legend_handles_labels()
# ax.legend(loc='upper left', fontsize=10)
ax2 = ax.legend(handles=legend_proxies, title='Protein length (approx)', loc='upper right', fontsize=9, title_fontsize=10, frameon=False)
ax.add_artist(ax2)

# Labels and title
ax.set_xlabel('Number of Peptides per Protein', fontsize=12, fontweight='bold')
ax.set_ylabel('Number of Proteoforms Discovered', fontsize=12, fontweight='bold')
ax.set_title(
    'Proteoform Discovery Landscape\n(Size ~ Protein length, Color = Coverage; alpha ~ Coverage)',
    fontsize=14, fontweight='bold'
)
ax.grid(True, which="both", linestyle="--", linewidth=0.5, alpha=0.7)

plt.tight_layout()
plots.finalize_plot( 
    fig, show=True, save=save_to_folder, 
    filename='proteoform_discovery_efficiency_48hr',
    filepath=figure_path,
    formats=figure_formats,
    transparent=transparent_bg,
    dpi=figure_dpi
)
```

Similar to the previous visualization (heatmap), this shows the number of proteoforms discovered tends to increase with peptide count, but with diminishing returns at higher counts. Larger proteins (bigger bubbles) and those with higher coverage (warmer colors) generally yield more proteoforms, indicating that both protein size and detection depth facilitate proteoform discovery.

---

##### 02.4 Discovery Efficiency by Cluster Count¶

Combined visualization showing proteoform category composition (stacked bars) and efficiency metrics (line plots) across cluster counts. This reveals how clustering behavior relates to proteoform discovery rates.

```
# Revised layout: stacked bar (top) + percentage lines + efficiency (bottom) with shared x-axis
import matplotlib.patches as mpatches

plot_data = protein_dpf_analysis.copy()

# 1. Prepare data (reuse same ordering)
composition_data = pd.crosstab(plot_data['all_clust'], plot_data['dpf_cat'])
category_order = ['dPF Only', 'PTM Only', 'No Forms', 'Both']
composition_data = composition_data.reindex(columns=category_order, fill_value=0).sort_index()
totals = composition_data.sum(axis=1).astype(int)
x_coords = np.arange(len(composition_data))

# colors in same order as columns
col_color_map = {"No Forms":"#e5e5e5","PTM Only":"#a7c957","dPF Only":"#bc4749","Both":"#967aa1"}
bar_colors = [col_color_map[c] for c in composition_data.columns]

# 2. Create stacked bar (top) and line/CI plot (bottom) sharing X
fig, (ax_bar, ax_line) = plt.subplots(2, 1, figsize=(12, 8), sharex=True, gridspec_kw={'height_ratios':[1, 1]})

# Top: stacked bars
composition_data.plot(
    kind='bar',
    stacked=True,
    ax=ax_bar,
    color=bar_colors,
    width=0.8,
    legend=False
)
ax_bar.set_ylabel('Number of Proteins', fontsize=12)
ax_bar.set_title('Composition of Proteoform Discoveries by Cluster Count', fontsize=16, pad=12)
ax_bar.grid(True, which="both", linestyle="--", linewidth=0.5, alpha=0.7)
ax_bar.set_ylim(0, int(totals.max() * 1.12))

# totals above bars
y_offset = max(2, totals.max() * 0.02)
for i, total in enumerate(totals.values):
    if total > 0:
        ax_bar.text(i, total + y_offset, f"{total:,}", ha='center', va='bottom', fontsize=10, fontweight='bold')

# tidy x tick labels on the bottom axis
ax_line.set_xlabel('Number of Clusters (all_clust)', fontsize=12)
ax_line.set_xticks(x_coords)
ax_line.set_xticklabels([str(x) for x in composition_data.index], rotation=0)

# 3. Bottom: percentage lines per category with 95% CI (binomial) + efficiency mean with CI (t-approx)
# Percentage per category (p) and 95% CI using binomial approximation: CI = 1.96 * sqrt(p*(1-p)/N)
for col, color in zip(composition_data.columns, bar_colors):
    counts = composition_data[col].values.astype(float)
    N = totals.values.astype(float)
    # avoid division by zero
    p = np.divide(counts, N, out=np.zeros_like(counts, dtype=float), where=N!=0)
    sem_binom = np.sqrt(np.divide(p * (1 - p), np.where(N == 0, 1, N)))
    ci95 = sem_binom * 1.96
    ax_line.errorbar(
        x_coords, p * 100,
        yerr=ci95 * 100,
        marker='o', linewidth=4, color=color, capsize=8, label=f"{col} (%)", 
        linestyle='-', markersize=8, alpha=0.8, elinewidth=2,
        markeredgecolor='black', capthick=2,
    )

# Efficiency stats by cluster: mean and 95% CI from group std / sqrt(n)
eff_stats = (
    plot_data.reset_index()
             .groupby('all_clust')['eff_pct']
             .agg(['mean', 'std', 'count'])
             .reindex(composition_data.index)
)
eff_mean = eff_stats['mean'].fillna(0)
eff_std = eff_stats['std'].fillna(0)
eff_count = eff_stats['count'].fillna(0).replace(0, np.nan)  # avoid div-by-zero
eff_sem = eff_std / np.sqrt(eff_count)
eff_ci95 = (eff_sem * 1.96).fillna(0)

# Plot efficiency mean and CI as emphasized black line + shaded band
ax_line.plot(x_coords, eff_mean.values, marker='s', color='k', linewidth=2.5, markersize=7, label='Avg Efficiency (eff_pct)')
ax_line.fill_between(x_coords, (eff_mean - eff_ci95).values, (eff_mean + eff_ci95).values, color='k', alpha=0.12)

# 4. Styling & legends
ax_line.set_ylabel('Percentage / Efficiency (%)', fontsize=12)
ax_line.set_ylim(0, 100)
ax_line.grid(True, which="both", linestyle="--", linewidth=0.5, alpha=0.7)

# Bar legend (from patches) and line legend
bar_handles = [mpatches.Patch(color=col_color_map[c], label=c) for c in composition_data.columns]
leg1 = ax_bar.legend(handles=bar_handles, title='Proteoform Category', bbox_to_anchor=(1.02, 1), loc='upper left', frameon=False)

leg2 = ax_line.legend(loc='upper center', bbox_to_anchor=(0.5, -0.18), ncol=5, frameon=False, title='Percent lines / Efficiency')

# keep bar legend visible
ax_bar.add_artist(leg1)

plt.tight_layout()
plots.finalize_plot( 
    fig, show=True, save=save_to_folder, 
    filename='proteoform_discovery_efficiency_by_cluster_48hr',
    filepath=figure_path,
    formats=figure_formats,
    transparent=transparent_bg,
    dpi=figure_dpi
)
```

This plot directly looks at the number of clusters not just dPF or PTMs. The number of clusters doesn't really affect much the proteoform discovery efficiency or the proteoform categories. With higher number of clusters percentage of dPFs decreases as well as PTM percentage, however at 9 clusters (which is only one protein) PTM is the only category observed.

> Note: This plot is not the most informative, however it does give some insight into how clustering relates to proteoform discovery. More clusters do not necessarily lead to higher proteoform detection, indicating that cluster quality and peptide characteristics may play larger roles.
>
> Note: Another thing that I can take from this is the clustering is the main bottleneck for proteoform discovery rather than the actual proteoform detection itself. As the number of clusters increase the proteoform discovery efficiency and percentages decrease which indicates that either the clustering is not optimal or the peptides themselves are not informative enough to distinguish proteoforms.

---

##### 02.5 Significant Peptide Distribution¶

Donut chart showing the distribution of proteins by their number of statistically significant peptides from the linear model: none, single, or multiple significant peptides per protein.

```
# Plot donut chart for proteins with no signf, single signf, multiple signf
plot_data = protein_dpf_analysis[[ 'Gene', 'sgnf_pep' ]].copy()

# Categorize proteins
labels = ['None', 'Single', 'Multiple']
plot_data['sgnf_cat'] = pd.cut(
    plot_data['sgnf_pep'],
    bins=[-1, 0, 1, np.inf],
    labels=labels,
    right=True
)

# Ordered counts and percentages
counts = plot_data['sgnf_cat'].value_counts().reindex(labels, fill_value=0)
total = counts.sum()
percent = counts / total

# Save the counts table
counts_table = pd.DataFrame({
    'Count': counts,
    'Percentage': percent
})
counts_table.to_csv(f"{output_path}protein_significantPeptides_counts_48hr.csv", index_label='Significant Peptides Category')
print("\nProtein Significant Peptides Counts:")
print(counts_table)

# slightly separate very small slices
explode = [0.02 if p < 0.05 else 0.0 for p in percent]

# Use a list of color strings (in the same order as `labels`) instead of a dict.
# colors_list = ['#c3c3c3', '#fca311', '#e54f2a'] # Autumn palette
colors_list = ["#e5e5e5", "#669bbc", "#003049"] # Blue-Gray palette

fig, ax = plt.subplots(figsize=(8, 6))
wedges, texts = ax.pie(
    counts.values,
    labels=None,             
    startangle=140,
    colors=colors_list,
    wedgeprops=dict(width=0.40, edgecolor='w'),
    explode=explode,
    counterclock=False
)

# Add percent labels placed on wedges
bbox_props = dict(boxstyle="round,pad=0.3", fc="white", ec="none", alpha=0.8)
for w, p in zip(wedges, percent):
    angle = (w.theta2 + w.theta1) / 2.0
    x, y = w.r * 0.75 * np.cos(np.deg2rad(angle)), w.r * 0.75 * np.sin(np.deg2rad(angle))
    ax.text(x, y, f"{p:.1%}", ha='center', va='center', fontsize=15, bbox=bbox_props, fontweight='bold')

# Center annotation: total proteins
ax.text(0, 0, f"Total\n{total}", ha='center', va='center', fontsize=15, fontweight='bold')

# Legend with counts and percentages
legend_labels = [f"{lab}: {cnt:,} ({pct:.1%})" for lab, cnt, pct in zip(labels, counts, percent)]
ax.legend(
    wedges, legend_labels,
    title="Number of Significant Peptides",
    loc="upper center",
    bbox_to_anchor=(0.5, -0.12),
    ncol=len(labels),
    fontsize=11,
    title_fontsize=12,
    frameon=False, 
    handletextpad=0.5,
    columnspacing=1.5,
)

ax.axis('equal')
plt.title('Distribution of Proteins by Number of \nSignificant Peptides from Linear Model', fontsize=16, fontweight='bold')
plt.tight_layout()
plots.finalize_plot( 
    fig, show=True, save=save_to_folder, 
    filename='significantPeptides_perProtein_Donut_48hr',
    filepath=figure_path,
    formats=figure_formats,
    transparent=transparent_bg,
    dpi=figure_dpi
)
```

```
Protein Significant Peptides Counts:
          Count  Percentage
sgnf_cat                   
None       3920      0.5474
Single     2185      0.3051
Multiple   1056      0.1475
```

This figure simply calculates the proportion of significant peptides detected per protein with splitting single or multiple significant peptides into their own categories. Shows ~55% of proteins without any significant peptides, which is an early indicator that comparing hypoxia vs normoxia at 48hr doesn't have a strong effect on most proteins at the peptide level. However, a substantial fraction (~45%) of proteins have at least one significant peptides, suggesting that a subset of proteins are responsive to the conditions tested.

---

##### 02.6 Proteoform Category Distribution¶

Donut chart showing the overall distribution of proteins across proteoform categories: no forms detected, PTM-only, dPF-only, or both PTMs and dPFs identified.

```
# Plot donut chart for proteins with no additional forms, only dPFs, only PTMs, with Both
plot_data = protein_dpf_analysis[[ 'Gene', 'n_forms', 'n_canon', 'n_ptm', 'n_dPF' ]].copy()

without_forms = set(plot_data[(plot_data['n_forms'] == 0)].index)
with_PTM_only = set(plot_data[(plot_data['n_forms'] == 0) & (plot_data['n_ptm'] > 0)].index)
with_dPF_only = set(plot_data[(plot_data['n_forms'] > 0) & (plot_data['n_ptm'] == 0)].index)
with_both = set(plot_data[(plot_data['n_forms'] > 0) & (plot_data['n_ptm'] > 0)].index)
without_forms = without_forms.difference(with_PTM_only)
print(f"Proteins with no additional forms: {len(without_forms):,}")
print(f"Proteins with only PTMs: {len(with_PTM_only):,}")
print(f"Proteins with only dPFs: {len(with_dPF_only):,}")
print(f"Proteins with both PTMs and dPFs: {len(with_both):,}")

# Categorize proteins
labels = ['No Forms', 'PTM Only', 'dPF Only', 'Both']
counts = [
    len(without_forms),
    len(with_PTM_only),
    len(with_dPF_only),
    len(with_both)
]
# Calculate percentages
percent = np.array(counts) / sum(counts)

# Save the counts table
counts_table = pd.DataFrame({
    'Count': counts,
    'Percentage': percent
}, index=labels)
counts_table.to_csv(f"{output_path}protein_formCategories_counts_48hr.csv", index_label='Form Categories')
print("\nProtein Form Categories Counts:")
print(counts_table)

# slightly separate very small slices
explode = [0.02 if p < 0.05 else 0.0 for p in percent]
# Use a list of color strings (in the same order as `labels`) instead of a dict.
colors_list = ["#e5e5e5", "#a7c957", "#bc4749", "#967aa1"]

fig, ax = plt.subplots(figsize=(8, 6))
wedges, texts = ax.pie(
    counts,
    labels=None,             
    startangle=140,
    colors=colors_list,
    wedgeprops=dict(width=0.40, edgecolor='w'),
    explode=explode,
    counterclock=False
)
# Add percent labels placed on wedges
bbox_props = dict(boxstyle="round,pad=0.3", fc="white", ec="none", alpha=0.8)
for w, p in zip(wedges, percent):
    angle = (w.theta2 + w.theta1) / 2.0
    x, y = w.r * 0.75 * np.cos(np.deg2rad(angle)), w.r * 0.75 * np.sin(np.deg2rad(angle))
    ax.text(x, y, f"{p:.1%}", ha='center', va='center', fontsize=15, bbox=bbox_props, fontweight='bold')

# Center annotation: total proteins
ax.text(0, 0, f"Total\n{sum(counts)}", ha='center', va='center', fontsize=15, fontweight='bold')

# Legend with counts and percentages
legend_labels = [f"{lab}: {cnt:,} ({pct:.1%})" for lab, cnt, pct in zip(labels, counts, percent)]
ax.legend(
    wedges, legend_labels,
    title="Number of Proteins",
    loc="upper center",
    bbox_to_anchor=(0.5, -0.12),
    ncol=len(labels),
    fontsize=11,
    title_fontsize=12,
    frameon=False, 
    handletextpad=0.5,
    columnspacing=1.5,
)
ax.axis('equal')
plt.title('Distribution of Proteins by Additional Forms and PTMs', fontsize=16, fontweight='bold')
plt.tight_layout()
plots.finalize_plot( 
    fig, show=True, save=save_to_folder, 
    filename='formCategories_perProtein_Donut_48hr',
    filepath=figure_path,
    formats=figure_formats,
    transparent=transparent_bg,
    dpi=figure_dpi
)
```

```
Proteins with no additional forms: 3,920
Proteins with only PTMs: 2,385
Proteins with only dPFs: 522
Proteins with both PTMs and dPFs: 334

Protein Form Categories Counts:
          Count  Percentage
No Forms   3920      0.5474
PTM Only   2385      0.3331
dPF Only    522      0.0729
Both        334      0.0466
```

Moving on to the categorization of the signficant peptides, there can be two things they can be. A signficant peptide can either doesn't cluster with any other peptides becomes a PTM, or it can cluster with other peptides (either significant or not) and become part of a dPF. This chart shows the overall distribution of proteins across these categories. Added "Both" which includes proteins that have both PTM-only peptides and peptides assigned to dPFs. Not surprising that most significant peptides are PTM-only. Majority are PTM-only, whereas the only ~5% of proteins with both PTM and dPF peptides.

---

#### 03. Peptide-UniProt Annotation Mapping¶

Now that we have an idea about the discovery of peptides as PTMs and/or dPFs, we can move on to mapping these peptides to UniProt annotations to understand their functional implications. This is the most crucial step to connect the peptide-level findings to biological context.

##### 03.1 Build Enhanced Annotation Map¶

Create a comprehensive mapping between identified peptides and UniProt annotations, including miscleavage detection. This connects peptide-level ProteoForge results with functional annotation data for biological interpretation.

```
def create_enhanced_peptide_uniprot_map_with_miscleavage(
    summary_data: pd.DataFrame,
    uniprot_data: pd.DataFrame,
    protein_dpf_analysis: pd.DataFrame,
    overlap_threshold: float = 0.5,
    verbose: bool = False
) -> pd.DataFrame:
    """
    Create an enhanced peptide-UniProt annotation mapping with miscleavage detection.

    Parameters
    ----------
    summary_data : pd.DataFrame
        Peptide summary data with columns: Protein, PeptideID, peptide_start, peptide_end, dPF.
    uniprot_data : pd.DataFrame
        UniProt annotation data with columns: Protein, feature_category, feature, start, end, group, agent, note, score, is_relevant.
    protein_dpf_analysis : pd.DataFrame
        Protein dPF analysis with at least columns: Protein, dpf_cat.
    overlap_threshold : float, optional
        Fractional overlap threshold for miscleavage detection (default is 0.5).
    verbose : bool, optional
        If True, print detailed progress and statistics.

    Returns
    -------
    pd.DataFrame
        DataFrame with columns:
        Protein, PeptideID, pepStart, pepEnd, startpos, endpos, feature_category, feature,
        group, agent, note, score, dPF.
    """
    import time
    t0 = time.time()
    if verbose:
        print("=== Enhanced Peptide-UniProt Mapping with Miscleavage Detection ===")

    # Filter proteins of interest
    proteins = protein_dpf_analysis.loc[
        protein_dpf_analysis['dpf_cat'] != 'No Forms', 'Protein'
    ].unique()
    if verbose:
        print(f"  Proteins to check: {len(proteins):,}")

    # Prepare peptides
    peptides = (
        summary_data[summary_data['Protein'].isin(proteins)]
        .rename(columns={'peptide_start': 'pepStart', 'peptide_end': 'pepEnd'})
        [['Protein', 'PeptideID', 'pepStart', 'pepEnd', 'dPF']]
        .reset_index(drop=True)
    )
    if verbose:
        print(f"  Peptides to process: {len(peptides):,}")

    # Prepare relevant UniProt annotations
    annotations = (
        uniprot_data[
            (uniprot_data['is_relevant']) &
            (uniprot_data['Protein'].isin(proteins))
        ]
        .rename(columns={'start': 'ann_start', 'end': 'ann_end'})
        [['Protein', 'feature_category', 'feature', 'ann_start', 'ann_end', 'group', 'agent', 'note', 'score']]
        .reset_index(drop=True)
    )
    if verbose:
        print(f"  Relevant UniProt annotations: {len(annotations):,}")

    # Find peptide-annotation overlaps
    merged = peptides.merge(annotations, on='Protein', how='left')
    overlap_mask = (
        ((merged['ann_start'] >= merged['pepStart']) & (merged['ann_start'] <= merged['pepEnd'])) |
        ((merged['ann_end'] >= merged['pepStart']) & (merged['ann_end'] <= merged['pepEnd'])) |
        ((merged['ann_start'] <= merged['pepStart']) & (merged['ann_end'] >= merged['pepEnd']))
    )
    ann_overlap = merged[overlap_mask].copy()

    # Build annotation result
    result_ann = pd.DataFrame()
    if not ann_overlap.empty:
        result_ann = pd.DataFrame({
            'Protein': ann_overlap['Protein'],
            'PeptideID': ann_overlap['PeptideID'],
            'pepStart': ann_overlap['pepStart'],
            'pepEnd': ann_overlap['pepEnd'],
            'startpos': ann_overlap['ann_start'],
            'endpos': ann_overlap['ann_end'],
            'feature_category': ann_overlap['feature_category'],
            'feature': ann_overlap['feature'],
            'group': ann_overlap['group'],
            'agent': ann_overlap['agent'],
            'note': ann_overlap['note'],
            'score': ann_overlap['score'],
            'dPF': ann_overlap['dPF']
        })

    # Miscleavage/peptide-peptide overlap detection
    if verbose:
        print("  Detecting miscleavage/peptide-peptide overlaps...")

    pep2pep = peptides.merge(peptides, on='Protein', suffixes=('', '_other'))
    pep2pep = pep2pep[pep2pep['PeptideID'] != pep2pep['PeptideID_other']]
    pep2pep['pep_length'] = pep2pep['pepEnd'] - pep2pep['pepStart'] + 1
    pep2pep['overlap_start'] = np.maximum(pep2pep['pepStart'], pep2pep['pepStart_other'])
    pep2pep['overlap_end'] = np.minimum(pep2pep['pepEnd'], pep2pep['pepEnd_other'])
    actual_overlap = pep2pep[pep2pep['overlap_start'] <= pep2pep['overlap_end']].copy()
    actual_overlap['overlap_length'] = actual_overlap['overlap_end'] - actual_overlap['overlap_start'] + 1
    actual_overlap['overlap_fraction'] = actual_overlap['overlap_length'] / actual_overlap['pep_length']
    sig_overlap = actual_overlap[actual_overlap['overlap_fraction'] > overlap_threshold].copy()
    sig_overlap['overlap_type'] = np.where(sig_overlap['overlap_fraction'] > 0.8, 'Miscleavage', 'Partial_Overlap')

    result_mis = pd.DataFrame()
    if not sig_overlap.empty:
        result_mis = pd.DataFrame({
            'Protein': sig_overlap['Protein'],
            'PeptideID': sig_overlap['PeptideID'],
            'pepStart': sig_overlap['pepStart'],
            'pepEnd': sig_overlap['pepEnd'],
            'startpos': sig_overlap['overlap_start'],
            'endpos': sig_overlap['overlap_end'],
            'feature_category': 'Cleavage Analysis',
            'feature': 'MISCLEAVAGE',
            'group': sig_overlap['overlap_type'],
            'agent': 'ProteoForge_Analysis',
            'note': 'Overlap with PeptideID ' + sig_overlap['PeptideID_other'].astype(str) +
                    ' (fraction: ' + sig_overlap['overlap_fraction'].round(3).astype(str) + ')',
            'score': sig_overlap['overlap_fraction'],
            'dPF': sig_overlap['dPF']
        })

    # Peptides with no annotation or miscleavage
    all_pep_ids = set(zip(peptides['Protein'], peptides['PeptideID']))
    annotated_ids = set()
    if not result_ann.empty:
        annotated_ids.update(zip(result_ann['Protein'], result_ann['PeptideID']))
    if not result_mis.empty:
        annotated_ids.update(zip(result_mis['Protein'], result_mis['PeptideID']))
    no_annot_ids = all_pep_ids - annotated_ids

    result_none = pd.DataFrame()
    if no_annot_ids:
        no_annot_df = peptides[
            peptides.set_index(['Protein', 'PeptideID']).index.isin(no_annot_ids)
        ].copy()
        result_none = pd.DataFrame({
            'Protein': no_annot_df['Protein'],
            'PeptideID': no_annot_df['PeptideID'],
            'pepStart': no_annot_df['pepStart'],
            'pepEnd': no_annot_df['pepEnd'],
            'startpos': -1,
            'endpos': -1,
            'feature_category': '',
            'feature': 'No Annotation',
            'group': '',
            'agent': '',
            'note': '',
            'score': 0.0,
            'dPF': no_annot_df['dPF']
        })

    # Combine all results
    result = pd.concat([df for df in [result_ann, result_mis, result_none] if not df.empty], ignore_index=True)

    # Standardize dtypes
    dtype_map = {
        'Protein': 'string',
        'PeptideID': 'int32',
        'pepStart': 'int32',
        'pepEnd': 'int32',
        'startpos': 'int32',
        'endpos': 'int32',
        'feature_category': 'string',
        'feature': 'string',
        'group': 'string',
        'agent': 'string',
        'note': 'string',
        'score': 'float32',
        'dPF': 'int8'
    }
    for col, typ in dtype_map.items():
        if col in result.columns:
            result[col] = result[col].astype(typ)
    string_cols = ['feature_category', 'group', 'agent', 'note']
    result[string_cols] = result[string_cols].fillna('')

    # Verbose summary
    if verbose:
        t1 = time.time()
        miscleavage_count = (result['feature'] == 'MISCLEAVAGE').sum()
        annotation_count = (result['feature'] != 'MISCLEAVAGE').sum()
        no_annotation_count = (result['feature'] == 'No Annotation').sum()
        print(f"\n=== Mapping Summary ===")
        print(f"  Total entries: {len(result):,}")
        print(f"    - UniProt annotations: {annotation_count - no_annotation_count:,}")
        print(f"    - Miscleavage features: {miscleavage_count:,}")
        print(f"    - No annotation: {no_annotation_count:,}")
        print(f"  Processing time: {t1-t0:.2f} seconds")

    return result[[
        'Protein', 'PeptideID', 'pepStart', 'pepEnd', 
        'feature_category', 'feature', 'startpos', 'endpos', 
        'group', 'agent', 'note', 'score', 'dPF'
    ]].sort_values(['Protein', 'PeptideID', 'feature', 'startpos']).reset_index(drop=True)

peptide_uniprot_map = create_enhanced_peptide_uniprot_map_with_miscleavage(
    summary_data, uniprot_data, protein_dpf_analysis, 
    overlap_threshold=0.5, verbose=True
)
print()
print("Sample of Peptide-UniProt Mapping with Miscleavage Detection:")
peptide_uniprot_map.head(10)
```

```
=== Enhanced Peptide-UniProt Mapping with Miscleavage Detection ===
  Proteins to check: 3,241
  Peptides to process: 56,822
  Relevant UniProt annotations: 315,401
  Detecting miscleavage/peptide-peptide overlaps...

=== Mapping Summary ===
  Total entries: 180,294
    - UniProt annotations: 160,410
    - Miscleavage features: 7,123
    - No annotation: 12,761
  Processing time: 1.78 seconds

Sample of Peptide-UniProt Mapping with Miscleavage Detection:
```

|  | Protein | PeptideID | pepStart | pepEnd | feature\_category | feature | startpos | endpos | group | agent | note | score | dPF |
| --- | --- | --- | --- | --- | --- | --- | --- | --- | --- | --- | --- | --- | --- |
| 0 | A0A1B0GTU1 | 1 | 89 | 99 |  | No Annotation | -1 | -1 |  |  |  | 0.0000 | 0 |
| 1 | A0A1B0GTU1 | 2 | 115 | 124 |  | No Annotation | -1 | -1 |  |  |  | 0.0000 | 0 |
| 2 | A0A1B0GTU1 | 3 | 179 | 193 |  | No Annotation | -1 | -1 |  |  |  | 0.0000 | -1 |
| 3 | A0A1B0GTU1 | 4 | 714 | 724 |  | No Annotation | -1 | -1 |  |  |  | 0.0000 | 0 |
| 4 | A0A1B0GTU1 | 5 | 758 | 776 | Co- & Post-Translational Modifications | CROSSLNK | 765 | 765 | Ubiquitination/SUMOylation |  | Ubiquitination/SUMOylation at K765 | 1.0000 | 0 |
| 5 | A0A2R8Y4L2 | 1 | 11 | 33 |  | No Annotation | -1 | -1 |  |  |  | 0.0000 | 0 |
| 6 | A0A2R8Y4L2 | 2 | 62 | 77 | Cleavage Analysis | MISCLEAVAGE | 69 | 77 | Partial\_Overlap | ProteoForge\_Analysis | Overlap with PeptideID 3 (fraction: 0.562) | 0.5625 | 0 |
| 7 | A0A2R8Y4L2 | 2 | 62 | 77 | Cleavage Analysis | MISCLEAVAGE | 69 | 77 | Partial\_Overlap | ProteoForge\_Analysis | Overlap with PeptideID 4 (fraction: 0.562) | 0.5625 | 0 |
| 8 | A0A2R8Y4L2 | 3 | 69 | 77 | Cleavage Analysis | MISCLEAVAGE | 69 | 77 | Miscleavage | ProteoForge\_Analysis | Overlap with PeptideID 2 (fraction: 1.0) | 1.0000 | 0 |
| 9 | A0A2R8Y4L2 | 3 | 69 | 77 | Cleavage Analysis | MISCLEAVAGE | 69 | 77 | Miscleavage | ProteoForge\_Analysis | Overlap with PeptideID 4 (fraction: 1.0) | 1.0000 | 0 |

Mapped about 180K annotations-peptides to generate a full Peptide-UniProt annotation map. As it is scene if a peptide doesn't have any annotation it is still kept in the map with empty annotations for `feature_category`, `group`, `agent`, etc. Additionally information such as score, dPF, and position are also kept for further analysis.

> This step is critical and I wanted to provide as much information from as many angles/sources into a single mapping table to facilitate downstream analyses linking peptide-level findings to functional annotations. However that you will see in upcoming parts, some are too complicated to automatically calculate so that I had to count manually for some of them.

---

#### 04. PTM and Proteoform Characterization¶

##### 04.1 PTM Annotation Analysis¶

Analyze peptides classified as PTM-only (dPF = -1) by examining their overlap with UniProt annotations. This reveals which PTM types are most commonly associated with single-peptide discordance patterns.

```
# Filter for only PTM peptides (dPF == -1)
final_res = peptide_uniprot_map[
    (peptide_uniprot_map['dPF'] == -1)
]

# Number of Proteins with No Annotation
no_annot_proteins = final_res[final_res['feature'] == 'No Annotation']['Protein'].unique()
with_annotation_proteins = final_res[final_res['feature'] != 'No Annotation']['Protein'].unique()
print(f"Number of Proteins with No Annotation: {len(no_annot_proteins)}")
print(f"Number of Proteins with Annotation: {len(with_annotation_proteins)}")

# Create a donut chart
data = [len(no_annot_proteins), len(with_annotation_proteins)]
# Labels
labels = ["No Annotation", "With Annotation"]
# Colors
colors = ["#e5e5e5", "#a7c957"]

# Save the counts table
counts_table = pd.DataFrame({
    'Count': data,
    'Percentage': np.array(data) / sum(data)
}, index=labels)
counts_table.to_csv(f"{output_path}proteins_withPTM_peptides_UniprotAnnotation_counts_48hr.csv", index_label='UniProt Annotation Availability')
# print("\nProteins with PTM Peptides UniProt Annotation Availability Counts:")
# print(counts_table)

# Create donut chart
fig, ax = plt.subplots(figsize=(8, 6))
wedges, texts = ax.pie(
    data,
    labels=None,             
    startangle=140,
    colors=colors,
    wedgeprops=dict(width=0.60, edgecolor='w'),
    explode=[0.02, 0.0],  # separate the first slice slightly
    counterclock=False
)
# Add percent labels placed on wedges
bbox_props = dict(boxstyle="round,pad=0.3", fc="white", ec="none", alpha=0.8)
for w, p in zip(wedges, np.array(data) / sum(data)):
    angle = (w.theta2 + w.theta1) / 2.0
    x, y = w.r * 0.75 * np.cos(np.deg2rad(angle)), w.r * 0.75 * np.sin(np.deg2rad(angle))
    ax.text(x, y, f"{p:.1%}", ha='center', va='center', fontsize=15, bbox=bbox_props, fontweight='bold')
# Center annotation: total proteins
ax.text(0, 0, f"Total\n{sum(data)}", ha='center', va='center', fontsize=15, fontweight='bold')
# Legend with counts and percentages
legend_labels = [f"{lab}: {cnt:,} ({pct:.1%})" for lab, cnt, pct in zip(labels, data, np.array(data) / sum(data))]
ax.legend(
    wedges, legend_labels,
    title="Number of Proteins",
    loc="upper center",
    bbox_to_anchor=(0.5, -0.12),
    ncol=len(labels),
    fontsize=11,
    title_fontsize=12,
    frameon=False, 
    handletextpad=0.5,
    columnspacing=1.5,
)
ax.axis('equal')
plt.title('Distribution of Proteins with PTM Peptides\nAnnotation Availability', fontsize=16, fontweight='bold')
plt.tight_layout()
plots.finalize_plot( 
    fig, show=True, save=save_to_folder, 
    filename='proteins_withPTM_peptides_UniprotAnnotation_Donut_48hr',
    filepath=figure_path,
    formats=figure_formats,
    transparent=transparent_bg,
    dpi=figure_dpi
)

print()

# Create a visualization to show the Number of Proteins with UniProt Annotations
fig, ax = plt.subplots(figsize=(5, 5))
sub_data = final_res[final_res['feature'] != 'No Annotation']

# Get counts of each feature type
feature_counts = sub_data['feature'].value_counts()

# Save the counts table
counts_table = pd.DataFrame({
    'Count': feature_counts
})
counts_table.to_csv(f"{output_path}proteins_withPTM_peptides_UniprotAnnotation_featureCounts_48hr.csv", index_label='Uniprot Feature Type')
# print("Proteins with PTM Peptides UniProt Annotation Feature Counts:")
# print(counts_table)

# Create a bar plot
feature_counts.plot(kind='barh', color="#a7c957", ax=ax, edgecolor='black')
# Add counts to on top of the bars
for i, v in enumerate(sub_data['feature'].value_counts()):
    ax.text(
        # v, i, str(v),
        # Adjust the position of the text
        v + 2.5, i-0.1, str(v),
        color='black', 
        ha='left', va='center', 
        rotation=0, fontsize=12, fontweight='bold'
    )
ax.set_xlabel("Number of Annotations", fontsize=12)
ax.set_ylabel("Uniprot Feature Type", fontsize=12)
ax.set_title("Annotations Matched to Single PTMs", fontsize=14, fontweight='bold')
ax.grid("both", linestyle="--", linewidth=0.75, alpha=0.5, color="lightgrey")
plt.tight_layout()
plt.show()
plots.finalize_plot( 
    fig, show=True, save=save_to_folder, 
    filename='proteins_withPTM_peptides_UniprotAnnotation_Bar_48hr',
    filepath=figure_path,
    formats=figure_formats,
    transparent=transparent_bg,
    dpi=figure_dpi
)
print()
print("Modified Residues (MOD_RES) Annotations:")
print((sub_data.loc[sub_data['feature']=='MOD_RES', 'group'].str.split(';').str[0]).value_counts())
print()
print("Cleavage Annotations:")
print((sub_data.loc[sub_data['feature']=='CLEAVAGE', 'group'].str.split(';').str[0]).value_counts())
print()
```

```
Number of Proteins with No Annotation: 705
Number of Proteins with Annotation: 2097

Proteins with PTM Peptides UniProt Annotation Availability Counts:
                 Count  Percentage
No Annotation      705      0.2516
With Annotation   2097      0.7484
```

```
Proteins with PTM Peptides UniProt Annotation Feature Counts:
             Count
feature           
MOD_RES       3663
CROSSLNK      1136
VAR_SEQ        954
CLEAVAGE       859
VARIANT        649
MISCLEAVAGE    450
CARBOHYD       188
MUTAGEN        179
CONFLICT       168
DISULFID        42
INIT_MET         1
```

```
Modified Residues (MOD_RES) Annotations:
group
Phosphorylation            3056
Acetylation                 360
Methylation                 188
Acylation                    33
Hydroxylation                13
Other Modification            6
Nitration/Nitrosylation       4
ADP-ribosylation              2
Citrullination                1
Name: count, dtype: int64

Cleavage Annotations:
group
Proteolytic Cleavage    859
Name: count, dtype: int64
```

There are 2802 peptides classified as PTM-only (dPF = -1) across all proteins (proteins with at least one peptide with ptm = 2097). Out of these, 2097 peptides (approximately 75%) have at least one annotation found within the peptide. This means quite a significant portion of PTM-only peptides can be linked to known features, however there are still ~25% of PTM-only peptides (proteins with at least one peptide with ptm but those don't match to any annotation = 705) that do not overlap with any known annotations, indicating potential novel modifications or limitations in current annotation databases, or simply peptides behaving like technical outliers without biological basis.

> Note: There can be multiple peptides as PTMs in a single proteins, thats why the 2802 total equal to the number of proteins with PTM-only and Both number of 2719.

Of those 2097 PTM-only peptides with annotations quite a large number of annotations is observed the numbers can be see in the second figure above. With the largest category being the MOD\_RES which makes sense as these are post-translational modifications. Other large categories like CROSSLNK, VAR\_SEQ, CLEAVAGE also make sense in the context of PTMs.

> Note: Another thing is that a peptide can host many annotations, which is why we see very large number of annotations compared to peptides.

Looking closely at the MOD\_RES annotations which are directly related to PTMs we see a wide variety of modification types associated with these PTM-only peptides. The most common modifications include Phosphorylation (Phosphoserine, Phosphothreonine, and Phosphotyrosine), which are well-known regulatory PTMs involved in signaling pathways. They are overwhelmingly represented here, indicating that many of the PTM-only peptides may be phosphorylated forms of the canonical sequences.

Additionally I have checked for what are the cleavage annotations, and only proteolytic cleavages are observed (Signal peptide cleavage, Transit peptide cleavage, and Propeptide cleavage). Which indicates that some of these PTM-only peptides may arise from specific proteolytic processing events rather than chemical modifications.

---

##### 04.2 Modification Type Distribution¶

Breakdown of specific modification types mapped to identified PTMs. The MOD\_RES annotations reveal the chemical nature of post-translational modifications associated with discordant peptide behavior.

```
# Data preparation
sub_data = final_res[final_res['feature'] != 'No Annotation']
mod_res_data = sub_data[sub_data['feature'] == 'MOD_RES']['group'].str.split(';').str[0].value_counts()

# Calculate percentages
mod_total = mod_res_data.sum()
mod_percentages = (mod_res_data / mod_total * 100)

# Save the counts table
counts_table = pd.DataFrame({
    'Count': mod_res_data,
    'Percentage': mod_percentages
})
counts_table.to_csv(f"{output_path}mod_res_modificationTypes_counts_48hr.csv", index_label='Modification Type')
print("\nMOD_RES Modification Types Counts:")
print(counts_table)

# Create figure
fig, ax = plt.subplots(figsize=(12, 4))

# Create stacked horizontal bar
left = 0
colors_mod = plt.cm.Set3(np.linspace(0, 1, len(mod_res_data)))

for i, (mod_type, count) in enumerate(mod_res_data.items()):
    width = mod_percentages.iloc[i]
    bar = ax.barh(0, width, left=left, color=colors_mod[i], 
                  edgecolor='white', linewidth=2, height=0.6)
    
    # Add percentage label if there's enough space (>3% threshold)
    if width > 3:
        ax.text(left + width/2, 0, f'{width:.1f}%\n({count:,})', 
               ha='center', va='center', fontsize=11, fontweight='bold',
               bbox=dict(boxstyle="round,pad=0.2", facecolor='white', alpha=0.8))
    
    left += width

# Styling
ax.set_xlim(0, 100)
ax.set_ylim(-0.5, 0.5)
ax.set_title(
    f'Distribution of Modification Types mapped to identified PTMs (n={mod_total:,})', 
    fontsize=16, fontweight='bold', pad=20
)
ax.set_yticks([])
ax.set_xticks([])

# Remove spines for cleaner look
ax.spines['top'].set_visible(False)
ax.spines['right'].set_visible(False)
ax.spines['left'].set_visible(False)
ax.spines['bottom'].set_visible(False)

# Add grid for easier reading
ax.grid(axis='x', linestyle='--', alpha=0.3)

# Create legend with counts
legend_elements = [
    plt.Rectangle((0,0),1,1, facecolor=colors_mod[i], label=f'{mod_type}: {count:,} ({mod_percentages.iloc[i]:.1f}%)')
    for i, (mod_type, count) in enumerate(mod_res_data.items())
]

ax.legend(
    handles=legend_elements, 
    loc='center', bbox_to_anchor=(0.5, -0.25), ncol=4, 
    fontsize=10, frameon=False, columnspacing=1.5
)

plt.tight_layout()

# Save the figure
plots.finalize_plot( 
    fig, show=True, save=save_to_folder, 
    filename='mod_res_modifications_stacked_bar_48hr',
    filepath=figure_path,
    formats=figure_formats,
    transparent=transparent_bg,
    dpi=figure_dpi
)

print(f"MOD_RES Analysis:")
print(f"Total modifications: {mod_total:,}")
for mod_type, count in mod_res_data.items():
    print(f"• {mod_type}: {count:,} ({count/mod_total*100:.1f}%)")
```

```
MOD_RES Modification Types Counts:
                         Count  Percentage
group                                     
Phosphorylation           3056     83.4289
Acetylation                360      9.8280
Methylation                188      5.1324
Acylation                   33      0.9009
Hydroxylation               13      0.3549
Other Modification           6      0.1638
Nitration/Nitrosylation      4      0.1092
ADP-ribosylation             2      0.0546
Citrullination               1      0.0273
```

```
MOD_RES Analysis:
Total modifications: 3,663
• Phosphorylation: 3,056 (83.4%)
• Acetylation: 360 (9.8%)
• Methylation: 188 (5.1%)
• Acylation: 33 (0.9%)
• Hydroxylation: 13 (0.4%)
• Other Modification: 6 (0.2%)
• Nitration/Nitrosylation: 4 (0.1%)
• ADP-ribosylation: 2 (0.1%)
• Citrullination: 1 (0.0%)
```

Above is the detailed distribution of specific MOD\_RES modification types observed among the PTM-only peptides. Phosphorylation events (Phosphoserine, Phosphothreonine, Phosphotyrosine) dominate the landscape, highlighting their prevalence in regulatory processes. Other modifications such as Acetylation, Methylation are with small but notable counts, indicating a diversity of PTM types contributing to peptide discordance.

---

##### 04.3 Differential Proteoform (dPF) Analysis¶

Analyze peptides classified as differential proteoforms (dPF > 0), examining annotation coverage and categorizing reasons for proteoform identification based on manual review results.

```
# Smartly build dPF color palette for all data
dPF_colors = {
    # PTM-Only
    -1: "#a7c957",  # PTM Only
    # Canonical
    0: "#c3c3c3",  # Canonical
    # Proteoforms
    # Calculate reds (#bc4749) for dPF > 0
    **({} if (len(summary_data) == 0 or int(summary_data['dPF'].max()) <= 0) 
        else {i: sns.color_palette("Reds_r", n_colors=int(summary_data['dPF'].max())).as_hex()[i-1] for i in range(1, int(summary_data['dPF'].max()) + 1)})
}
print(f"Proteoform Group Colors: {dPF_colors}")
# sns.palplot(list(dPF_colors.values()));
print()

protein_with_dpf = protein_dpf_analysis.loc[
    (protein_dpf_analysis['n_forms'] > 0),
    'Protein'
].copy()

dPF_uniprot_maps = peptide_uniprot_map[
    (peptide_uniprot_map['dPF'] > 0)
].copy()
print(f"Number of Proteins with dPFs: {len(protein_with_dpf):,}")
print(f"Number of generated mappings: {len(dPF_uniprot_maps):,}")
print(f"Number of dPF Annotations found: {len(dPF_uniprot_maps[dPF_uniprot_maps['feature'] != 'No Annotation']):,}")
# dPF_uniprot_maps.head(20)

# protein_dpf_analysis.sort_values('dPF_pct', ascending=False).head(30)
```

```
Proteoform Group Colors: {-1: '#a7c957', 0: '#c3c3c3', 1: '#cb181d', 2: '#fb6b4b', 3: '#fcbca2'}

Number of Proteins with dPFs: 856
Number of generated mappings: 43,640
Number of dPF Annotations found: 41,458
```

```
# # Manual Check Results for dPFs
# # Reasons Categorized
# high_level_dict = {
#     'No Annotation': 192,
#     'Mixed/Unclear Reasons': 857,
#     'Clear Reasons': 405,
# }
# # Categories of Clear Reasons
# clear_reasons_dict = {
#     'Missed Cleavages': 193,
#     'Isoforms': 51,
#     'Multiple PTMs': 37,
#     'N-terminal': 34,
#     'C-terminal': 31,
#     'Variants': 20,
#     'Mutagen': 20,
#     'Cleavage': 19,
# }

# # Number of dPFs 
# dPF_counts = protein_dpf_analysis['n_forms'].value_counts().sort_index()
# print("dPF Counts per Protein:")
# for n_forms, count in dPF_counts.items():
#     print(f"  {n_forms}: {count:,}")

# print("Number of Unique dPFs > 0:", protein_dpf_analysis[protein_dpf_analysis['n_forms'] > 0]['n_forms'].sum())
# print()

# print("Manual Check Results for dPFs:")
# for reason, count in high_level_dict.items():
#     print(f"  {reason}: {count:,}")

# # Donut chart for dPF annotation reasons
# labels = list(high_level_dict.keys())
# counts = list(high_level_dict.values())
# colors = ["#e5e5e5",  '#fdcab5', "#bc4749"]
# fig, ax = plt.subplots(figsize=(8, 6))
# wedges, texts = ax.pie(
#     counts,
#     labels=None,             
#     startangle=140,
#     colors=colors,
#     wedgeprops=dict(width=0.60, edgecolor='w'),
#     explode=[0.02 if c < 0.05 * sum(counts) else 0.0 for c in counts],
#     counterclock=False
# )
# # Add percent labels placed on wedges
# bbox_props = dict(boxstyle="round,pad=0.3", fc="white", ec="none", alpha=0.8)
# for w, p in zip(wedges, np.array(counts) / sum(counts)):
#     angle = (w.theta2 + w.theta1) / 2.0
#     x, y = w.r * 0.75 * np.cos(np.deg2rad(angle)), w.r * 0.75 * np.sin(np.deg2rad(angle))
#     ax.text(x, y, f"{p:.1%}", ha='center', va='center', fontsize=15, bbox=bbox_props, fontweight='bold')
# # Center annotation: total proteins
# ax.text(0, 0, f"Total\n{sum(counts)}", ha='center', va='center', fontsize=15, fontweight='bold')
# # Legend with counts and percentages
# legend_labels = [f"{lab}: {cnt:,} ({pct:.1%})" for lab, cnt, pct in zip(labels, counts, np.array(counts) / sum(counts))]
# ax.legend(
#     wedges, legend_labels,
#     title="dPF Annotation Reasons",
#     loc="upper center",
#     bbox_to_anchor=(0.5, -0.12),
#     ncol=len(labels),
#     fontsize=11,
#     title_fontsize=12,
#     frameon=False, 
#     handletextpad=0.5,  
#     columnspacing=1.5,
# )
# ax.axis('equal')
# plt.title('Distribution of dPF Annotation Reasons', fontsize=16, fontweight='bold')
# plt.tight_layout()
# plots.finalize_plot( 
#     fig, show=True, save=save_to_folder, 
#     filename='dPF_annotation_reasons_Donut_48hr',
#     filepath=figure_path,
#     formats=figure_formats,
#     transparent=transparent_bg,
#     dpi=figure_dpi
# )

# # Create a horizontal bar chart for Clear Reasons
# clear_reasons = list(clear_reasons_dict.keys())
# clear_counts = list(clear_reasons_dict.values())
# clear_color = "#bc4749"
# fig, ax = plt.subplots(figsize=(5, 5))
# bars = ax.barh(clear_reasons, clear_counts, color=clear_color, edgecolor='black')
# # Add counts to on top of the bars
# for i, v in enumerate(clear_counts):
#     ax.text(
#         # v, i, str(v),
#         # Adjust the position of the text
#         v + 2.5, i-0.1, str(v),
#         color='black', 
#         ha='left', va='center', 
#         rotation=0, fontsize=12, fontweight='bold'
#     )
# ax.set_xlabel("Number of dPFs", fontsize=12)
# ax.set_ylabel("Clear Reason Categories", fontsize=12)
# ax.set_title("Breakdown of Clear Reasons for dPF Annotations", fontsize=14, fontweight='bold')
# ax.grid("both", linestyle="--", linewidth=0.75, alpha=0.5, color="lightgrey")
# plt.tight_layout()
# plots.finalize_plot( 
#     fig, show=True, save=save_to_folder, 
#     filename='dPF_annotation_clear_reasons_Bar_48hr',
#     filepath=figure_path,
#     formats=figure_formats,
#     transparent=transparent_bg,
#     dpi=figure_dpi
# )
```

Due to the complexity of dPF peptides and the variety of potential reasons for their discordant behavior, a manual review was conducted to categorize the underlying causes. I was able to categorize them into three groups: No Annotation Found, Clear Reasons (annotations that clearly explain the dPF behavior), and Mixed/Unclear Reasons (annotations present but not obviously explaining the discordance or many different annotations making it hard to pinpoint a single cause).

While the automated annotation mapping provided a wealth of information, the diversity of annotations and their interpretations necessitated manual curation to accurately classify the dPF peptides. This manual review helped to clarify the biological relevance of the observed proteoform discordance.

About 28% (405) do have clear reasons for their discordant behavior based on the annotations present. These include well-characterized PTMs, alternative splicing events, or known functional domains that could explain the differential regulation observed. Howver majority 59% don't have well separated reasons, there are annotations present but they either don't clearly explain the discordance or there are many different annotations making it hard to pinpoint a single cause.

Upon examining the annotations associated with dPF peptides, several common themes emerged among those with clear reasons for their discordant behavior. Many dPF peptides were linked: Missed cleavages which can be due to incomplete enzymatic digestion during sample preparation, leading to unexpected peptide forms; Alternative splicing events (mostly isoforms) resulting in proteoforms with distinct sequences and functional properties; Post-translational modifications (PTMs) such as phosphorylation, acetylation, or ubiquitination that can alter peptide behavior and detection; Functional domains or motifs known to influence protein activity, localization, or interactions, which may contribute to differential regulation. There are also cleavage, neo-N/C terminal events.

> **Note:** The manual categorization process is inherently subjective and may vary between reviewers. Future work could focus on developing more objective criteria or automated methods for classifying dPF peptides based on annotation data.
>
> **Future Directions:** With a web-based interface and interactive analysis and visualization tools, this process can be streamlined and left to the biologist, or experts to examine based on their domain knowledge rather than relying on a single reviewer. Additionally having access to more centralized or comprehensive annotations or even proteoforms themselves could make this step more automated and less subjective.

---

##### 04.4 Consecutive Peptide Validation¶

Identify proteins where all positive dPF peptides are consecutive along the protein sequence. Consecutive dPF peptides suggest true proteoform regions rather than scattered technical artifacts.

```
tmp = summary_data.set_index('Protein').loc[protein_with_dpf].copy()

candidate_proteins = []

# Build a vectorized table for positive dPF groups across all proteins
df_pos = tmp.reset_index()[['Protein', 'dPF', 'PeptideID']].copy()
# Keep only positive dPFs
df_pos = df_pos[df_pos['dPF'] > 0].copy()

# Ensure PeptideID numeric and drop invalid rows
df_pos['PeptideID'] = pd.to_numeric(df_pos['PeptideID'], errors='coerce')
df_pos = df_pos.dropna(subset=['PeptideID'])
if df_pos.empty:
    if verbose:
        print("No positive dPFs found in the selected proteins.")
else:
    # Aggregate per (Protein, dPF)
    agg = (
        df_pos
        .groupby(['Protein', 'dPF'], observed=True)
        .PeptideID
        .agg(min_id='min', max_id='max', n_unique=lambda x: x.nunique())
        .reset_index()
    )
    # Compute consecutiveness: n_unique == (max_id - min_id + 1)
    agg['consecutive'] = agg['n_unique'] == (agg['max_id'] - agg['min_id'] + 1)

    # For each protein, require ALL positive-dPF groups to be consecutive
    per_protein_ok = agg.groupby('Protein', observed=True)['consecutive'].all()

    # Candidate proteins are those with True
    candidate_proteins = per_protein_ok[per_protein_ok].index.tolist()

    # # Optional verbose reporting (summarize kept / rejected)
    # if verbose:
    #     kept = per_protein_ok[per_protein_ok].index.tolist()
    #     rejected = per_protein_ok[~per_protein_ok].index.tolist()
    #     print(f"KEPT count: {len(kept):,}; REJECTED count: {len(rejected):,}")

    #     # Print examples (up to 10) with details to aid inspection
    #     for p in kept[:10]:
    #         rows = agg[agg['Protein'] == p].set_index('dPF')[['min_id','max_id','n_unique','consecutive']]
    #         print(f"KEPT: {p} -> all dPFs consecutive: {rows.to_dict(orient='index')}")

    #     for p in rejected[:10]:
    #         rows = agg[agg['Protein'] == p].set_index('dPF')[['min_id','max_id','n_unique','consecutive']]
    #         failed = rows[~rows['consecutive']]
    #         print(f"REJECT: {p} -> non-consecutive dPFs: {failed.index.tolist()}; details:")
    #         print(failed)

print(f"Number of Candidate Proteins with consistent (non-gap) dPFs: {len(candidate_proteins):,}")
print(f"Candidate Proteins: {candidate_proteins}")
print()
# utils.view_table(
#     protein_dpf_analysis[
#         (protein_dpf_analysis['Protein'].isin(candidate_proteins)) & 
#         (protein_dpf_analysis['dPF_pct'] <= 50)
#     ].sort_values('dPF_pct', ascending=False),
#     page_size=10,
#     page_number=5
# )
```

```
Number of Candidate Proteins with consistent (non-gap) dPFs: 98
Candidate Proteins: ['O14503', 'O15091', 'O43149', 'O60493', 'O94813', 'O95249', 'O95678', 'P04406', 'P05089', 'P07951', 'P08729', 'P0DTE7', 'P0DTE8', 'P0DUB6', 'P16401', 'P18850', 'P19012', 'P19793', 'P20042', 'P20645', 'P23921', 'P26447', 'P27816', 'P28340', 'P28370', 'P28702', 'P35527', 'P38646', 'P43007', 'P51178', 'P52306', 'P52735', 'P52824', 'P54764', 'P61201', 'Q00577', 'Q02218', 'Q03468', 'Q12901', 'Q12913', 'Q13283', 'Q13330', 'Q13950', 'Q14147', 'Q14588', 'Q14CX7', 'Q15003', 'Q3SY52', 'Q4KMQ2', 'Q4KWH8', 'Q5SSJ5', 'Q5SYE7', 'Q69YH5', 'Q6NUQ4', 'Q6NXT6', 'Q6P4Q7', 'Q6P9B6', 'Q6PIJ6', 'Q6UW63', 'Q6UWP8', 'Q6UXD5', 'Q7L1W4', 'Q7Z5K2', 'Q7Z736', 'Q86SQ0', 'Q86YC2', 'Q8IU60', 'Q8NBU5', 'Q8ND83', 'Q8NEN9', 'Q8WWM7', 'Q8WXG6', 'Q92600', 'Q96AY2', 'Q96GN5', 'Q96HS1', 'Q96RL7', 'Q96SB8', 'Q99650', 'Q99933', 'Q99986', 'Q9BTC8', 'Q9BZL6', 'Q9C000', 'Q9H000', 'Q9HC35', 'Q9NQ11', 'Q9NVA2', 'Q9NWU2', 'Q9NZI7', 'Q9NZL3', 'Q9UKE5', 'Q9Y2L5', 'Q9Y314', 'Q9Y5E3', 'Q9Y5I2', 'Q9Y6E2', 'Q9Y6I4']
```

This is one category that looked promising and interesting to look at. The idea here is that if all the dPF peptides for a given protein are consecutive along the protein sequence, it strongly suggests that these peptides represent a true proteoform region rather than scattered technical artifacts. Consecutive peptides indicate a coherent segment of the protein that may be differentially regulated or modified, supporting the biological relevance of the observed proteoform discordance.

> Note: I have used these to start the manual checkups, as well as the interesting and biologically relevant proteins that I know of from the hypoxia experiment.

---

#### 05. Case Studies and Indivudal Examination of Proteins¶

##### 05.1 Relevant Genes and Proteins from Hypoxia Experiment¶

Based on the hypoxia experiment detailed in the paper, the following genes and proteins are of particular interest. The experiment exposed H358 lung cancer cells to hypoxic conditions (1% oxygen) and analyzed changes in their proteome. Key enzymes related to glycolysis and the glyoxalase system showed altered abundance, with an inverse correlation between hypoxic markers (LDHA, HK2, GAPDH upregulation) and GLO2 downregulation.

**Key Proteins of Interest:**

- **LDHA** (P00338): Lactate production, upregulated in hypoxia
- **HK2** (P52790): Glycolysis entry, upregulated in hypoxia
- **GAPDH** (P04406): Glycolysis, upregulated in hypoxia
- **GLO1** (Q04760): Methylglyoxal metabolism, unaffected
- **HAGH/GLO2** (Q16775): Methylglyoxal metabolism, downregulated in hypoxia

```
glycolytic_up_by_HIF_1 = [
    'P00338', #LDHA                 # PTM
    'P52789', #HK2                  # PTM
    'P04406', #GAPDH                # dPF * Picked
    'Q04760', #GLO1                 # dPF * Picked 
    'Q16775', #HAGH (GLO2)          # PTM
]

is_demo = False # False if want to save figures

pass_dict = {
    'data': {
        'uniprot': uniprot_data,
        'detailed': test_data,
        'summary': summary_data,
    },
    'styles': {
        'dpf_colors': dPF_colors,
    },
    'config': {
        'is_demo': is_demo,
        'figure_path': figure_path,
        'figure_formats': figure_formats,
        'transparent_bg': transparent_bg,
        'figure_dpi': figure_dpi,
        'str_add': '48hr',
    }
}
```

###### 05.1.1 LDHA (P00338)¶

The LDHA in the original hypoxia experiment showed significant upregulation under hypoxic conditions, consistent with its role in anaerobic metabolism.

**ProteoForge Protein Summary Visualization**

Generate detailed peptide-level visualization showing intensity patterns, significance, and cluster assignments for selected proteins of interest. This enables in-depth examination of proteoform behavior under hypoxic conditions.

This figure summarizes how the ProteoForge sees the protein, and at raw, adjusted intentsities peptides across conditions act, as well as the p-value of the interaction model per peptide as well as the clustering assignments.

```
current_protein = 'P00338'
cur_gene = protein_dpf_analysis.loc[
    protein_dpf_analysis['Protein'] == current_protein, 'Gene'
].values[0]
print(f"Current Protein: {current_protein} ({cur_gene})")
plots.detailed_peptide_with_clusters(
    data=test_data,
    cur_protein=current_protein,
    pThr=pThr,
    pvalue_col='adj_pval',
    cluster_col='ClusterID',
    rawIntensity_col='log10Intensity',
    adjIntensity_col='AdjIntensity',
    condition_palette=condition_colors,
    
    save = not is_demo,
    show=True,
    filename=f'detailed_peptide_look_{cur_gene}_48hr',
    filepath=figure_path,
    fileformats=figure_formats,
    transparent=transparent_bg,
    dpi=figure_dpi
)
```

```
Current Protein: P00338 (LDHA)
```

From the adjusted intensity there is a pattern where hypoxia is slightly higher than normoxia, but the difference is not very pronounced. And only one PTM peptide is observed, where the difference is very pronounced between hypoxia and normoxia. For that peptide the intensity is not high in raw intensity, however the normoxia is very low, this could peptide that is imputed with low-values due to it being completely missing in normoxia samples.

---

**Detailed Protein Annotation Visualization**

Create comprehensive protein-level visualization integrating UniProt annotations with peptide coverage. Multiple annotation layers show sequence features, modifications, and functional domains alongside identified peptides colored by dPF category.

```
plots.annotations_on_protein(
    target_protein=current_protein,
    control_package=pass_dict
)
```

**Various Data Views around the Peptides of Interest(s)**

LDAH's 16th peptide is marked as potential PTM, let's see what are the annotations mapped to that region of the protein.

```
peptide_uniprot_map[
    (peptide_uniprot_map['Protein']==current_protein) & # Current Protein
    (peptide_uniprot_map['PeptideID'].isin([16])) # Peptide to inspect
]
```

|  | Protein | PeptideID | pepStart | pepEnd | feature\_category | feature | startpos | endpos | group | agent | note | score | dPF |
| --- | --- | --- | --- | --- | --- | --- | --- | --- | --- | --- | --- | --- | --- |
| 20628 | P00338 | 16 | 229 | 243 | Protein Processing & Maturation | CLEAVAGE | 231 | 231 | Proteolytic Cleavage | peptidyl-Lys metallopeptidase | N-Ph cleavage at H231 by peptidyl-Lys metallop... | 2.0000 | -1 |
| 20629 | P00338 | 16 | 229 | 243 | Protein Processing & Maturation | CLEAVAGE | 232 | 232 | Proteolytic Cleavage | trypsin 1 | N-Ph cleavage at K232 by trypsin 1 | 2.0000 | -1 |
| 20630 | P00338 | 16 | 229 | 243 | Protein Processing & Maturation | CLEAVAGE | 235 | 235 | Proteolytic Cleavage | cathepsin K | Ph cleavage at V235 by cathepsin K | 2.0000 | -1 |
| 20631 | P00338 | 16 | 229 | 243 | Protein Processing & Maturation | CLEAVAGE | 236 | 236 | Proteolytic Cleavage | cathepsin K | Ph cleavage at E236 by cathepsin K | 2.0000 | -1 |
| 20632 | P00338 | 16 | 229 | 243 | Protein Processing & Maturation | CLEAVAGE | 236 | 236 | Proteolytic Cleavage | cathepsin S | N-Ph cleavage at E236 by cathepsin S | 2.0000 | -1 |
| 20633 | P00338 | 16 | 229 | 243 | Protein Processing & Maturation | CLEAVAGE | 242 | 242 | Proteolytic Cleavage | peptidyl-Lys metallopeptidase | N-Ph cleavage at I242 by peptidyl-Lys metallop... | 2.0000 | -1 |
| 20634 | P00338 | 16 | 229 | 243 | Co- & Post-Translational Modifications | CROSSLNK | 232 | 232 | Ubiquitination/SUMOylation |  | Ubiquitination/SUMOylation at K232 | 1.0000 | -1 |
| 20635 | P00338 | 16 | 229 | 243 | Co- & Post-Translational Modifications | CROSSLNK | 243 | 243 | Ubiquitination/SUMOylation |  | Ubiquitination/SUMOylation at K243 | 1.0000 | -1 |
| 20636 | P00338 | 16 | 229 | 243 | Cleavage Analysis | MISCLEAVAGE | 233 | 243 | Partial\_Overlap | ProteoForge\_Analysis | Overlap with PeptideID 17 (fraction: 0.733) | 0.7333 | -1 |
| 20637 | P00338 | 16 | 229 | 243 | Co- & Post-Translational Modifications | MOD\_RES | 232 | 232 | Acetylation |  | N6-acetyllysine | 3.0000 | -1 |
| 20638 | P00338 | 16 | 229 | 243 | Co- & Post-Translational Modifications | MOD\_RES | 232 | 232 | Acetylation |  | Acetylation at K232 | 1.0000 | -1 |
| 20639 | P00338 | 16 | 229 | 243 | Co- & Post-Translational Modifications | MOD\_RES | 237 | 237 | Phosphorylation |  | Phosphorylation at S237 | 1.0000 | -1 |
| 20640 | P00338 | 16 | 229 | 243 | Co- & Post-Translational Modifications | MOD\_RES | 239 | 239 | Phosphorylation |  | Phosphotyrosine | 3.0000 | -1 |
| 20641 | P00338 | 16 | 229 | 243 | Co- & Post-Translational Modifications | MOD\_RES | 239 | 239 | Phosphorylation |  | Phosphorylation at Y239 | 2.0000 | -1 |
| 20642 | P00338 | 16 | 229 | 243 | Co- & Post-Translational Modifications | MOD\_RES | 239 | 239 | Phosphorylation |  | Phosphorylation at Y239 | 2.0000 | -1 |
| 20643 | P00338 | 16 | 229 | 243 | Co- & Post-Translational Modifications | MOD\_RES | 243 | 243 | Acetylation |  | N6-acetyllysine | 3.0000 | -1 |
| 20644 | P00338 | 16 | 229 | 243 | Co- & Post-Translational Modifications | MOD\_RES | 243 | 243 | Acetylation |  | Acetylation at K243 | 1.0000 | -1 |
| 20645 | P00338 | 16 | 229 | 243 | Sequence Heterogeneity & Isoforms | VAR\_SEQ | 230 | 274 | Modified | 2 | Modified (in isoform 2) | 3.0000 | -1 |
| 20646 | P00338 | 16 | 229 | 243 | Sequence Heterogeneity & Isoforms | VAR\_SEQ | 237 | 241 | Modified | 5 | Modified (in isoform 5) | 3.0000 | -1 |
| 20647 | P00338 | 16 | 229 | 243 | Sequence Heterogeneity & Isoforms | VAR\_SEQ | 242 | 332 | Missing | 5 | Missing (in isoform 5) | 3.0000 | -1 |

#### 05.1.2 HK2 (P52789)¶

```
current_protein = 'P52789'
cur_gene = protein_dpf_analysis.loc[
    protein_dpf_analysis['Protein'] == current_protein, 'Gene'
].values[0]
print(f"Current Protein: {current_protein} ({cur_gene})")
plots.detailed_peptide_with_clusters(
    data=test_data,
    cur_protein=current_protein,
    pThr=pThr,
    pvalue_col='adj_pval',
    cluster_col='ClusterID',
    rawIntensity_col='log10Intensity',
    adjIntensity_col='AdjIntensity',
    condition_palette=condition_colors,
    
    save = not is_demo,
    show=True,
    filename=f'detailed_peptide_look_{cur_gene}_48hr',
    filepath=figure_path,
    fileformats=figure_formats,
    transparent=transparent_bg,
    dpi=figure_dpi
)
```

```
Current Protein: P52789 (HK2)
```

```
plots.annotations_on_protein(
    target_protein=current_protein,
    control_package=pass_dict
)
```

```
peptide_uniprot_map[
    (peptide_uniprot_map['Protein']==current_protein) & # Current Protein
    (peptide_uniprot_map['PeptideID'].isin([1])) # Peptide to inspect
]
```

|  | Protein | PeptideID | pepStart | pepEnd | feature\_category | feature | startpos | endpos | group | agent | note | score | dPF |
| --- | --- | --- | --- | --- | --- | --- | --- | --- | --- | --- | --- | --- | --- |
| 69426 | P52789 | 1 | 22 | 30 | Protein Processing & Maturation | CLEAVAGE | 30 | 30 | Proteolytic Cleavage | trypsin 1 | N-Ph cleavage at R30 by trypsin 1 | 2.0000 | -1 |

###### 05.1.2 GAPDH (P04406)¶

```
current_protein = 'P04406'
cur_gene = protein_dpf_analysis.loc[
    protein_dpf_analysis['Protein'] == current_protein, 'Gene'
].values[0]
print(f"Current Protein: {current_protein} ({cur_gene})")
plots.detailed_peptide_with_clusters(
    data=test_data,
    cur_protein=current_protein,
    pThr=pThr,
    pvalue_col='adj_pval',
    cluster_col='ClusterID',
    rawIntensity_col='log10Intensity',
    adjIntensity_col='AdjIntensity',
    condition_palette=condition_colors,
    
    save = not is_demo,
    show=True,
    filename=f'detailed_peptide_look_{cur_gene}_48hr',
    filepath=figure_path,
    fileformats=figure_formats,
    transparent=transparent_bg,
    dpi=figure_dpi
)
```

```
Current Protein: P04406 (GAPDH)
```

```
plots.annotations_on_protein(
    target_protein=current_protein,
    control_package=pass_dict
)
```

```
peptide_uniprot_map[
    (peptide_uniprot_map['Protein']==current_protein) & # Current Protein
    (peptide_uniprot_map['PeptideID'].isin([17, 18])) # Peptide to inspect
]
```

|  | Protein | PeptideID | pepStart | pepEnd | feature\_category | feature | startpos | endpos | group | agent | note | score | dPF |
| --- | --- | --- | --- | --- | --- | --- | --- | --- | --- | --- | --- | --- | --- |
| 24791 | P04406 | 17 | 235 | 248 | Co- & Post-Translational Modifications | CARBOHYD | 239 | 239 | N-linked |  | N-linked at N239 | 1.0000 | 1 |
| 24792 | P04406 | 17 | 235 | 248 | Protein Processing & Maturation | CLEAVAGE | 235 | 235 | Proteolytic Cleavage | trypsin 1 | N-Ph cleavage at V235 by trypsin 1 | 2.0000 | 1 |
| 24793 | P04406 | 17 | 235 | 248 | Protein Processing & Maturation | CLEAVAGE | 238 | 238 | Proteolytic Cleavage | matrix metallopeptidase-2 | N-Ph cleavage at A238 by matrix metallopeptida... | 2.0000 | 1 |
| 24794 | P04406 | 17 | 235 | 248 | Protein Processing & Maturation | CLEAVAGE | 238 | 238 | Proteolytic Cleavage | meprin beta subunit | N-Ph cleavage at A238 by meprin beta subunit | 2.0000 | 1 |
| 24795 | P04406 | 17 | 235 | 248 | Protein Processing & Maturation | CLEAVAGE | 238 | 238 | Proteolytic Cleavage | meprin alpha subunit | N-Ph cleavage at A238 by meprin alpha subunit | 2.0000 | 1 |
| 24796 | P04406 | 17 | 235 | 248 | Protein Processing & Maturation | CLEAVAGE | 238 | 238 | Proteolytic Cleavage | matrix metallopeptidase-3 | Ph cleavage at A238 by matrix metallopeptidase-3 | 2.0000 | 1 |
| 24797 | P04406 | 17 | 235 | 248 | Protein Processing & Maturation | CLEAVAGE | 239 | 239 | Proteolytic Cleavage | meprin alpha subunit | N-Ph cleavage at N239 by meprin alpha subunit | 2.0000 | 1 |
| 24798 | P04406 | 17 | 235 | 248 | Protein Processing & Maturation | CLEAVAGE | 239 | 239 | Proteolytic Cleavage | trypsin 1 | N-Ph cleavage at N239 by trypsin 1 | 2.0000 | 1 |
| 24799 | P04406 | 17 | 235 | 248 | Protein Processing & Maturation | CLEAVAGE | 239 | 239 | Proteolytic Cleavage | matrix metallopeptidase-3 | Ph cleavage at N239 by matrix metallopeptidase-3 | 2.0000 | 1 |
| 24800 | P04406 | 17 | 235 | 248 | Protein Processing & Maturation | CLEAVAGE | 241 | 241 | Proteolytic Cleavage | cathepsin S | N-Ph cleavage at S241 by cathepsin S | 2.0000 | 1 |
| 24801 | P04406 | 17 | 235 | 248 | Protein Processing & Maturation | CLEAVAGE | 244 | 244 | Proteolytic Cleavage | matrix metallopeptidase-3 | Ph cleavage at D244 by matrix metallopeptidase-3 | 2.0000 | 1 |
| 24802 | P04406 | 17 | 235 | 248 | Protein Processing & Maturation | CLEAVAGE | 246 | 246 | Proteolytic Cleavage | matrix metallopeptidase-3 | Ph cleavage at T246 by matrix metallopeptidase-3 | 2.0000 | 1 |
| 24803 | P04406 | 17 | 235 | 248 | Protein Processing & Maturation | CLEAVAGE | 248 | 248 | Proteolytic Cleavage | trypsin 1 | N-Ph cleavage at R248 by trypsin 1 | 2.0000 | 1 |
| 24804 | P04406 | 17 | 235 | 248 | Protein Processing & Maturation | CLEAVAGE | 248 | 248 | Proteolytic Cleavage | cathepsin S | N-Ph cleavage at R248 by cathepsin S | 2.0000 | 1 |
| 24805 | P04406 | 17 | 235 | 248 | Co- & Post-Translational Modifications | MOD\_RES | 237 | 237 | Phosphorylation |  | Phosphothreonine | 3.0000 | 1 |
| 24806 | P04406 | 17 | 235 | 248 | Co- & Post-Translational Modifications | MOD\_RES | 237 | 237 | Phosphorylation |  | Phosphorylation at T237 | 4.0000 | 1 |
| 24807 | P04406 | 17 | 235 | 248 | Co- & Post-Translational Modifications | MOD\_RES | 237 | 237 | Phosphorylation | P31751 | Phosphorylation at T237 | 2.0000 | 1 |
| 24808 | P04406 | 17 | 235 | 248 | Co- & Post-Translational Modifications | MOD\_RES | 237 | 237 | Phosphorylation |  | Phosphorylation at T237 | 4.0000 | 1 |
| 24809 | P04406 | 17 | 235 | 248 | Co- & Post-Translational Modifications | MOD\_RES | 237 | 237 | Phosphorylation |  | Phosphorylation at T237 | 4.0000 | 1 |
| 24810 | P04406 | 17 | 235 | 248 | Co- & Post-Translational Modifications | MOD\_RES | 241 | 241 | Phosphorylation |  | Phosphoserine | 3.0000 | 1 |
| 24811 | P04406 | 17 | 235 | 248 | Co- & Post-Translational Modifications | MOD\_RES | 241 | 241 | Phosphorylation |  | Phosphorylation at S241 | 2.0000 | 1 |
| 24812 | P04406 | 17 | 235 | 248 | Co- & Post-Translational Modifications | MOD\_RES | 241 | 241 | Phosphorylation |  | Phosphorylation at S241 | 2.0000 | 1 |
| 24813 | P04406 | 17 | 235 | 248 | Co- & Post-Translational Modifications | MOD\_RES | 246 | 246 | Phosphorylation |  | Phosphorylation at T246 | 3.0000 | 1 |
| 24814 | P04406 | 17 | 235 | 248 | Co- & Post-Translational Modifications | MOD\_RES | 246 | 246 | Phosphorylation | Q05655 | Phosphorylation at T246 | 2.0000 | 1 |
| 24815 | P04406 | 17 | 235 | 248 | Co- & Post-Translational Modifications | MOD\_RES | 246 | 246 | Phosphorylation |  | Phosphorylation at T246 | 3.0000 | 1 |
| 24816 | P04406 | 17 | 235 | 248 | Co- & Post-Translational Modifications | MOD\_RES | 247 | 247 | Acylation |  | S-(2-succinyl)cysteine | 3.0000 | 1 |
| 24817 | P04406 | 17 | 235 | 248 | Co- & Post-Translational Modifications | MOD\_RES | 247 | 247 | Nitration/Nitrosylation |  | S-nitrosocysteine | 3.0000 | 1 |
| 24818 | P04406 | 17 | 235 | 248 | Co- & Post-Translational Modifications | MOD\_RES | 247 | 247 | Methylation |  | Methylation at C247 | 1.0000 | 1 |
| 24819 | P04406 | 17 | 235 | 248 | Co- & Post-Translational Modifications | MOD\_RES | 247 | 247 | Nitration/Nitrosylation |  | Nitration/Nitrosylation at C247 | 0.0000 | 1 |
| 24820 | P04406 | 17 | 235 | 248 | Co- & Post-Translational Modifications | MOD\_RES | 247 | 247 | Nitration/Nitrosylation |  | Nitration/Nitrosylation at C247 | 0.0000 | 1 |
| 24821 | P04406 | 17 | 235 | 248 | Co- & Post-Translational Modifications | MOD\_RES | 248 | 248 | Methylation |  | Methylation at R248 | 1.0000 | 1 |
| 24822 | P04406 | 17 | 235 | 248 | Sequence Heterogeneity & Isoforms | MUTAGEN | 241 | 241 | No Significant Effect | S->A | Does not affect glycosylation by C.rodentium p... | 3.0000 | 1 |
| 24823 | P04406 | 17 | 235 | 248 | Sequence Heterogeneity & Isoforms | MUTAGEN | 245 | 245 | Complex/Specific Effect | L->M | Inhibits S-nitrosylation of Cys-247; when asso... | 3.0000 | 1 |
| 24824 | P04406 | 17 | 235 | 248 | Sequence Heterogeneity & Isoforms | MUTAGEN | 246 | 246 | No Significant Effect | T->A | Does not affect glycosylation by C.rodentium p... | 3.0000 | 1 |
| 24825 | P04406 | 17 | 235 | 248 | Sequence Heterogeneity & Isoforms | MUTAGEN | 247 | 247 | Reduced Function | C->S | Markedly reduced glycolytic activity; when ass... | 3.0000 | 1 |
| 24826 | P04406 | 18 | 261 | 271 | Protein Processing & Maturation | CLEAVAGE | 263 | 263 | Proteolytic Cleavage | trypsin 1 | N-Ph cleavage at K263 by trypsin 1 | 2.0000 | 1 |
| 24827 | P04406 | 18 | 261 | 271 | Protein Processing & Maturation | CLEAVAGE | 271 | 271 | Proteolytic Cleavage | trypsin 1 | N-Ph cleavage at K271 by trypsin 1 | 2.0000 | 1 |
| 24828 | P04406 | 18 | 261 | 271 | Co- & Post-Translational Modifications | CROSSLNK | 263 | 263 | Ubiquitination/SUMOylation |  | Ubiquitination/SUMOylation at K263 | 1.0000 | 1 |
| 24829 | P04406 | 18 | 261 | 271 | Co- & Post-Translational Modifications | CROSSLNK | 263 | 263 | Ubiquitination/SUMOylation |  | Ubiquitination/SUMOylation at K263 | 1.0000 | 1 |
| 24830 | P04406 | 18 | 261 | 271 | Co- & Post-Translational Modifications | CROSSLNK | 271 | 271 | Ubiquitination/SUMOylation |  | Ubiquitination/SUMOylation at K271 | 1.0000 | 1 |
| 24831 | P04406 | 18 | 261 | 271 | Co- & Post-Translational Modifications | MOD\_RES | 263 | 263 | Methylation |  | N6,N6-dimethyllysine | 3.0000 | 1 |
| 24832 | P04406 | 18 | 261 | 271 | Co- & Post-Translational Modifications | MOD\_RES | 263 | 263 | Acetylation |  | Acetylation at K263 | 1.0000 | 1 |
| 24833 | P04406 | 18 | 261 | 271 | Co- & Post-Translational Modifications | MOD\_RES | 263 | 263 | Methylation |  | Methylation at K263 | 3.0000 | 1 |
| 24834 | P04406 | 18 | 261 | 271 | Co- & Post-Translational Modifications | MOD\_RES | 263 | 263 | Methylation |  | Methylation at K263 | 3.0000 | 1 |
| 24835 | P04406 | 18 | 261 | 271 | Co- & Post-Translational Modifications | MOD\_RES | 266 | 266 | Phosphorylation |  | Phosphorylation at S266 | 1.0000 | 1 |

###### 05.1.2 GLO1 (Q04760)¶

```
current_protein = 'Q04760'
cur_gene = protein_dpf_analysis.loc[
    protein_dpf_analysis['Protein'] == current_protein, 'Gene'
].values[0]
print(f"Current Protein: {current_protein} ({cur_gene})")
plots.detailed_peptide_with_clusters(
    data=test_data,
    cur_protein=current_protein,
    pThr=pThr,
    pvalue_col='adj_pval',
    cluster_col='ClusterID',
    rawIntensity_col='log10Intensity',
    adjIntensity_col='AdjIntensity',
    condition_palette=condition_colors,
    
    save = not is_demo,
    show=True,
    filename=f'detailed_peptide_look_{cur_gene}_48hr',
    filepath=figure_path,
    fileformats=figure_formats,
    transparent=transparent_bg,
    dpi=figure_dpi
)
```

```
Current Protein: Q04760 (GLO1)
```

```
plots.annotations_on_protein(
    target_protein=current_protein,
    control_package=pass_dict
)
```

```
peptide_uniprot_map[
    (peptide_uniprot_map['Protein']==current_protein) & # Current Protein
    (peptide_uniprot_map['PeptideID'].isin([1])) # Peptide to inspect
]
```

|  | Protein | PeptideID | pepStart | pepEnd | feature\_category | feature | startpos | endpos | group | agent | note | score | dPF |
| --- | --- | --- | --- | --- | --- | --- | --- | --- | --- | --- | --- | --- | --- |
| 82870 | Q04760 | 1 | 29 | 38 | Protein Processing & Maturation | CLEAVAGE | 38 | 38 | Proteolytic Cleavage | trypsin 1 | N-Ph cleavage at R38 by trypsin 1 | 2.0000 | -1 |
| 82871 | Q04760 | 1 | 29 | 38 | Co- & Post-Translational Modifications | MOD\_RES | 35 | 35 | Phosphorylation |  | Phosphorylation at T35 | 1.0000 | -1 |
| 82872 | Q04760 | 1 | 29 | 38 | Co- & Post-Translational Modifications | MOD\_RES | 38 | 38 | Methylation |  | Methylation at R38 | 1.0000 | -1 |
| 82873 | Q04760 | 1 | 29 | 38 | Sequence Heterogeneity & Isoforms | MUTAGEN | 34 | 34 | Reduced Function | Q->E | Reduces enzyme activity by 99%. | 3.0000 | -1 |

###### 05.1.2 HAGH/GLO2 (Q16775)¶

```
current_protein = 'Q16775'
cur_gene = protein_dpf_analysis.loc[
    protein_dpf_analysis['Protein'] == current_protein, 'Gene'
].values[0]
print(f"Current Protein: {current_protein} ({cur_gene})")
plots.detailed_peptide_with_clusters(
    data=test_data,
    cur_protein=current_protein,
    pThr=pThr,
    pvalue_col='adj_pval',
    cluster_col='ClusterID',
    rawIntensity_col='log10Intensity',
    adjIntensity_col='AdjIntensity',
    condition_palette=condition_colors,
    
    save = not is_demo,
    show=True,
    filename=f'detailed_peptide_look_{cur_gene}_48hr',
    filepath=figure_path,
    fileformats=figure_formats,
    transparent=transparent_bg,
    dpi=figure_dpi
)
```

```
Current Protein: Q16775 (HAGH)
```

```
plots.annotations_on_protein(
    target_protein=current_protein,
    control_package=pass_dict
)
```

```
peptide_uniprot_map[
    (peptide_uniprot_map['Protein']==current_protein) & # Current Protein
    (peptide_uniprot_map['PeptideID'].isin([4])) # Peptide to inspect
]
```

|  | Protein | PeptideID | pepStart | pepEnd | feature\_category | feature | startpos | endpos | group | agent | note | score | dPF |
| --- | --- | --- | --- | --- | --- | --- | --- | --- | --- | --- | --- | --- | --- |
| 106933 | Q16775 | 4 | 233 | 242 | Sequence Heterogeneity & Isoforms | VAR\_SEQ | 146 | 236 | Modified | 3 | Modified (in isoform 3) | 3.0000 | -1 |
| 106934 | Q16775 | 4 | 233 | 242 | Sequence Heterogeneity & Isoforms | VAR\_SEQ | 237 | 308 | Missing | 3 | Missing (in isoform 3) | 3.0000 | -1 |

##### 05.2 Other Notable/Interesting Proteins¶

###### 05.2.1 SARG (Q9BW04)¶

```
current_protein = 'Q9BW04'
cur_gene = protein_dpf_analysis.loc[
    protein_dpf_analysis['Protein'] == current_protein, 'Gene'
].values[0]
print(f"Current Protein: {current_protein} ({cur_gene})")
plots.detailed_peptide_with_clusters(
    data=test_data,
    cur_protein=current_protein,
    pThr=pThr,
    pvalue_col='adj_pval',
    cluster_col='ClusterID',
    rawIntensity_col='log10Intensity',
    adjIntensity_col='AdjIntensity',
    condition_palette=condition_colors,
    
    save = not is_demo,
    show=True,
    filename=f'detailed_peptide_look_{cur_gene}_48hr',
    filepath=figure_path,
    fileformats=figure_formats,
    transparent=transparent_bg,
    dpi=figure_dpi
)
```

```
Current Protein: Q9BW04 (SARG)
```

```
plots.annotations_on_protein(
    target_protein=current_protein,
    control_package=pass_dict
)
```

```
peptide_uniprot_map[
    (peptide_uniprot_map['Protein']==current_protein) & # Current Protein
    (peptide_uniprot_map['PeptideID'].isin([11, 12, 15, 16])) # Peptide to inspect
]
```

|  | Protein | PeptideID | pepStart | pepEnd | feature\_category | feature | startpos | endpos | group | agent | note | score | dPF |
| --- | --- | --- | --- | --- | --- | --- | --- | --- | --- | --- | --- | --- | --- |
| 151365 | Q9BW04 | 11 | 300 | 308 | Co- & Post-Translational Modifications | MOD\_RES | 308 | 308 | Methylation |  | Methylation at K308 | 1.0000 | 1 |
| 151366 | Q9BW04 | 12 | 354 | 369 |  | No Annotation | -1 | -1 |  |  |  | 0.0000 | 1 |
| 151369 | Q9BW04 | 15 | 429 | 439 | Co- & Post-Translational Modifications | MOD\_RES | 429 | 429 | Phosphorylation |  | Phosphorylation at S429 | 1.0000 | 1 |
| 151370 | Q9BW04 | 15 | 429 | 439 | Sequence Heterogeneity & Isoforms | VARIANT | 434 | 434 | Polymorphism | rs35267170 | N -> S | 3.0000 | 1 |
| 151371 | Q9BW04 | 16 | 451 | 475 | Co- & Post-Translational Modifications | MOD\_RES | 453 | 453 | Phosphorylation |  | Phosphorylation at S453 | 1.0000 | 1 |
| 151372 | Q9BW04 | 16 | 451 | 475 | Co- & Post-Translational Modifications | MOD\_RES | 456 | 456 | Phosphorylation |  | Phosphorylation at T456 | 1.0000 | 1 |
| 151373 | Q9BW04 | 16 | 451 | 475 | Co- & Post-Translational Modifications | MOD\_RES | 462 | 462 | Phosphorylation |  | Phosphorylation at S462 | 1.0000 | 1 |
| 151374 | Q9BW04 | 16 | 451 | 475 | Co- & Post-Translational Modifications | MOD\_RES | 471 | 471 | Phosphorylation |  | Phosphorylation at T471 | 1.0000 | 1 |

###### 05.2.2 FPGT (O14772)¶

```
current_protein = 'O14772'
cur_gene = protein_dpf_analysis.loc[
    protein_dpf_analysis['Protein'] == current_protein, 'Gene'
].values[0]
print(f"Current Protein: {current_protein} ({cur_gene})")
plots.detailed_peptide_with_clusters(
    data=test_data,
    cur_protein=current_protein,
    pThr=pThr,
    pvalue_col='adj_pval',
    cluster_col='ClusterID',
    rawIntensity_col='log10Intensity',
    adjIntensity_col='AdjIntensity',
    condition_palette=condition_colors,
    
    save = not is_demo,
    show=True,
    filename=f'detailed_peptide_look_{cur_gene}_48hr',
    filepath=figure_path,
    fileformats=figure_formats,
    transparent=transparent_bg,
    dpi=figure_dpi
)
```

```
Current Protein: O14772 (FPGT)
```

```
plots.annotations_on_protein(
    target_protein=current_protein,
    control_package=pass_dict
)
```

```
# peptide_uniprot_map[
#     (peptide_uniprot_map['Protein']==current_protein) & # Current Protein
#     (peptide_uniprot_map['PeptideID'].isin([11, 12, 15, 16])) # Peptide to inspect
# ]
```

```
print("Notebook Execution Time:", utils.prettyTimer(utils.getTime() - startTime))
```

```
Notebook Execution Time: 00h:00m:48s
```
