## Supplementary material for "ProteoForge: An Imputation-Aware Framework for Differential Proteoform Discovery in Bottom-Up Proteomics": Analysis notebooks as html render: Notebook S13.html

### Notebook S13

###### Enes Kemal Ergin

#### 2025-12-08 23:00:17

### Supplementary Notebook 13: Exploring the ProteoForge Results from 72hr¶

- **License:** Creative Commons Attribution-NonCommercial 4.0 International License
- **Version:** 0.2
- **Edit Log:**
  - 2025-11-28: Initial version of the notebook
  - 2025-12-08: Revise the whole notebook, ensuring clarity and correctness

---

**Requirements:**

This notebook requires the `02-ApplyingProteoForge_72hr.ipynb` to be run first to generate the result files used here. Ensure the `./data/cleaned/` and `.data/results/` directories contains the necessary input files.

---

**Data Information:**

- Cell line: H358 (NSCLC)
- Conditions: True Hypoxia (1% O₂) at 72 Hr, Normoxia (21% O₂) at 72 Hr
- Input: `test_data_72hr.feather`, `summary_data_72hr.feather`, `uniprot_data_72hr.feather`
- Output: Visualizations and summary statistics for identified proteoforms and some example proteins of interest

---

**Purpose:**

This notebook summarizes and visualizes the results from the ProteoForge analysis of hypoxia vs normoxia conditions at 72-hour timepoint. It includes protein-level differential proteoform (dPF) analysis, peptide composition distributions, UniProt annotation mapping, and case studies of biologically relevant proteoforms.

###### Setting Up the Data¶

Here we load all the result files `test_data_72hr.feather`, `summary_data_72hr.feather`, and `info_data.feather` generated from the ProteoForge analysis in the previous notebook for 72hr. As well as the `protein_data.feather` which will help with protein level informations (expanded Fasta).

**Peptide Regulation Across Conditions**

One thing that we can look is the regulation compared to control condition (Normoxia 21% O2 at 72hr).

```
pThr = 10**-3

test_data = pd.read_feather(f"{output_path}test_data_72hr.feather")
print(test_data.shape)
print()


summary_data = pd.read_feather(f"{output_path}summary_data_72hr.feather")
print(summary_data.shape)
print()

info_data = pd.read_feather(f"{input_path}info_data.feather")
print(info_data.shape)
print()

summary_data['trace'] = info_data['trace']

protein_data = pd.read_feather(f"{input_path}protein_data.feather")
print(protein_data.shape)
print()


## Adding Hypoxia Regulation compared to Normoxia 

df =  test_data.copy()

# --- 2. Define Control and Treatment Conditions ---
control_condition = 'Normoxia (21% O2) - 72 Hr'
# Sorting ensures the regulation string (e.g., 'Up_Down') is always in a consistent order.
treatment_conditions = sorted([cond for cond in df['Condition'].unique() if cond != control_condition])

print(f"Control: {control_condition}")
print(f"Treatments (in order): {treatment_conditions}")

# --- 3. Calculate Mean Intensity for Each Peptide in Each Condition ---
# This aggregates the technical replicates for each condition.
# The mean of AdjIntensity for a treatment is its log2FC vs. control.
# Using .agg is slightly more explicit and can be faster than .mean()
mean_intensities = df.groupby(['Protein', 'PeptideID', 'Condition'], as_index=False).agg(log2FC=('AdjIntensity', 'mean'))


# --- 4. Create the Regulation Table ---
# We can filter out the control condition, as its regulation is baseline (0).
regulation_df = mean_intensities[mean_intensities['Condition'] != control_condition].copy()

# Calculate the linear Fold Change from the log2FC.
# FoldChange = 2^(log2FC)
regulation_df['FoldChange'] = 2**regulation_df['log2FC']


# --- 5. Categorize Regulation Tendencies (Optimized) ---
# To categorize tendencies, we first pivot the table to have conditions as columns.
# This allows us to see the behavior of one peptide across all conditions in a single row.
pivot_reg = regulation_df.pivot_table(
    index=['Protein', 'PeptideID'],
    columns='Condition',
    values='log2FC'
).fillna(0) # Fill peptides not found in a condition with 0 log2FC (no change)

# This vectorized approach is much faster than using .apply()
# It builds the regulation string (e.g., 'Up_Down') column by column.
regulation_strings = None
for i, cond in enumerate(treatment_conditions):
    # Use np.select for fast conditional categorization based on direction
    categories = np.select(
        [pivot_reg[cond] > 0, pivot_reg[cond] < 0],
        ['Up', 'Down'],
        default='NC' # No Change
    )
    
    # Convert to a pandas Series to align with the pivot_reg index
    categories_series = pd.Series(categories, index=pivot_reg.index)

    if i == 0:
        regulation_strings = categories_series
    else:
        # Append the category for the next condition
        regulation_strings += '_' + categories_series

pivot_reg['Regulation'] = regulation_strings


# Now, merge the 'Regulation' column back into our long-format table
# This adds the summary string to each row corresponding to a given peptide.
regulation_df = regulation_df.merge(
    pivot_reg[['Regulation']],
    on=['Protein', 'PeptideID'],
    how='left'
)

# Reorder columns for a clean final output
regulation_df = regulation_df[['Protein', 'PeptideID', 'Condition', 'log2FC', 'FoldChange', 'Regulation']]


print("\n--- Final Regulation Table with Tendencies ---")
print(regulation_df.head())

# You can now save this final dataframe to a new file, for example:
# regulation_df.to_csv('regulation_analysis_output.csv', index=False)

# Example of how to use the new 'Regulation' column:
print("\n--- Example: Grouping by Regulation Tendency ---")
print(regulation_df.groupby('Regulation')['PeptideID'].nunique())

plot_data = regulation_df[['Protein', 'PeptideID', 'Regulation']].drop_duplicates().copy()

# Adding the plot_data to summary_data to get the Regulation column (use hypoxia 48hr-72hr)
summary_data = summary_data.merge(
    plot_data[['Protein', 'PeptideID', 'Regulation']],
    on=['Protein', 'PeptideID'],
    how='left'
)
summary_data['Regulation'] = summary_data['Regulation'].fillna('NC_NC')  # peptides not in regulation_df are 'No Change' in both conditions
summary_data.rename(columns={'Regulation': 'Regulation_(Hypoxia 72hr)'}, inplace=True)

fig, ax = plt.subplots(figsize=(8, 6))
plot_data['Regulation'].value_counts().plot(kind='barh', color="#432c46", edgecolor='black', ax=ax)
# Add counts to on top of the bars for barh
for i, v in enumerate(plot_data['Regulation'].value_counts()):
    ax.text(
        v, i, str(v),
        color='black', 
        ha='left', va='center', 
        rotation=0, fontsize=15, fontweight='bold'
    )
ax.set_xlabel("Number of Peptides", fontsize=12)
ax.set_ylabel("Regulation Tendency (Hypoxia 48Hr-72Hr)", fontsize=12)
ax.set_title("Peptide Regulation Tendencies Across Conditions", fontsize=14, fontweight='bold')
ax.grid("both", linestyle="--", linewidth=0.75, alpha=0.5, color="lightgrey")
plt.tight_layout()

plots.finalize_plot( 
    fig, show=True, save=save_to_folder, 
    filename="peptide_regulation_tendencies_72hr",
    filepath=figure_path,
    formats=figure_formats,
    transparent=transparent_bg,
    dpi=figure_dpi
)

print()
summary_data['Regulation_(Hypoxia 72hr)'].value_counts()
```

```
(572214, 24)

(95369, 17)

(95369, 15)

(7161, 8)

Control: Normoxia (21% O2) - 72 Hr
Treatments (in order): ['True Hypoxia (1% O2) - 72 Hr']

--- Final Regulation Table with Tendencies ---
      Protein  PeptideID                     Condition  log2FC  FoldChange Regulation
0  A0A024RBG1          1  True Hypoxia (1% O2) - 72 Hr -0.1385      0.9085       Down
1  A0A024RBG1          2  True Hypoxia (1% O2) - 72 Hr -0.1047      0.9300       Down
2  A0A024RBG1          3  True Hypoxia (1% O2) - 72 Hr -0.1472      0.9030       Down
3  A0A024RBG1          4  True Hypoxia (1% O2) - 72 Hr -0.0052      0.9964       Down
4  A0A024RBG1          5  True Hypoxia (1% O2) - 72 Hr  0.1125      1.0811         Up

--- Example: Grouping by Regulation Tendency ---
Regulation
Down    221
Up      387
Name: PeptideID, dtype: int64
```

```
Regulation_(Hypoxia 72hr)
Down    53825
Up      41544
Name: count, dtype: int64
```

Unlike the previous notebook (48hr only summary), here 72hr the peptide regulation is to be downregulation in hypoxia conditions is more prevalent compared to normoxia. This is inline with the biological expectation that hypoxia generally leads to a reduction in overall protein synthesis and metabolic activity as cells adapt to low oxygen conditions.

|  | Protein | Gene | Length | Coverage | all\_pep | sgnf\_pep | all\_clust | n\_forms | n\_canon | n\_ptm | n\_dPF | sgnf\_pct | canon\_pct | ptm\_pct | dPF\_pct | eff\_pct | cov\_cat | sgnf\_cat | len\_cat | dpf\_cat |
| --- | --- | --- | --- | --- | --- | --- | --- | --- | --- | --- | --- | --- | --- | --- | --- | --- | --- | --- | --- | --- |
| 0 | A0A024RBG1 | NUDT4B | 181 | 30.3867 | 5 | 0 | 3 | 0 | 5 | 0 | 0 | 0.0000 | 100.0000 | 0.0000 | 0.0000 | 0.0000 | Medium | None | Short | No Forms |
| 1 | A0A0B4J2D5 | GATD3B | 268 | 35.4478 | 7 | 0 | 4 | 0 | 7 | 0 | 0 | 0.0000 | 100.0000 | 0.0000 | 0.0000 | 0.0000 | Medium | None | Medium | No Forms |
| 2 | A0A1B0GTU1 | ZC3H11B | 805 | 8.1988 | 5 | 2 | 3 | 0 | 3 | 2 | 0 | 40.0000 | 60.0000 | 40.0000 | 0.0000 | 0.0000 | Low | Multiple | Long | PTM Only |
| 3 | A0A1W2PQL4 | ZNF722 | 384 | 11.7188 | 5 | 1 | 2 | 0 | 4 | 1 | 0 | 20.0000 | 80.0000 | 20.0000 | 0.0000 | 0.0000 | Low | Single | Medium | PTM Only |
| 4 | A0A2R8Y4L2 | HNRNPA1L3 | 275 | 44.7273 | 12 | 2 | 2 | 1 | 0 | 1 | 11 | 16.6667 | 0.0000 | 8.3333 | 91.6667 | 50.0000 | Medium | Multiple | Medium | Both |
| ... | ... | ... | ... | ... | ... | ... | ... | ... | ... | ... | ... | ... | ... | ... | ... | ... | ... | ... | ... | ... |
| 7156 | Q9Y6X5 | ENPP4 | 453 | 19.8675 | 7 | 0 | 2 | 0 | 7 | 0 | 0 | 0.0000 | 100.0000 | 0.0000 | 0.0000 | 0.0000 | Low | None | Medium | No Forms |
| 7157 | Q9Y6X8 | ZHX2 | 837 | 33.0944 | 22 | 2 | 2 | 1 | 20 | 0 | 2 | 9.0909 | 90.9091 | 0.0000 | 9.0909 | 50.0000 | Medium | Multiple | Long | dPF Only |
| 7158 | Q9Y6X9 | MORC2 | 1032 | 27.2287 | 23 | 4 | 3 | 2 | 19 | 0 | 4 | 17.3913 | 82.6087 | 0.0000 | 17.3913 | 66.6667 | Medium | Multiple | V.Long | dPF Only |
| 7159 | Q9Y6Y0 | IVNS1ABP | 642 | 11.0592 | 5 | 0 | 3 | 0 | 5 | 0 | 0 | 0.0000 | 100.0000 | 0.0000 | 0.0000 | 0.0000 | Low | None | Medium | No Forms |
| 7160 | Q9Y6Y8 | SEC23IP | 1000 | 22.7000 | 17 | 2 | 2 | 1 | 15 | 0 | 2 | 11.7647 | 88.2353 | 0.0000 | 11.7647 | 50.0000 | Low | Multiple | Long | dPF Only |

This is shows a clear difference from the 48hr only summary, where we saw the increase of PTM/ProteoForms with increasing coverage, which is the same, but the overall peptide composition is a lot higher, indicating that at 72hr there are more peptide level variations, leading ProteoForge to identify more peptides as PTM or proteoform associated.

```
# Plot donut chart for proteins with no signf, single signf, multiple signf
plot_data = protein_dpf_analysis[[ 'Gene', 'sgnf_pep' ]].copy()

# Categorize proteins
labels = ['None', 'Single', 'Multiple']
plot_data['sgnf_cat'] = pd.cut(
    plot_data['sgnf_pep'],
    bins=[-1, 0, 1, np.inf],
    labels=labels,
    right=True
)

# Ordered counts and percentages
counts = plot_data['sgnf_cat'].value_counts().reindex(labels, fill_value=0)
total = counts.sum()
percent = counts / total

# Save the counts table
counts_table = pd.DataFrame({
    'Count': counts,
    'Percentage': percent
})
counts_table.to_csv(f"{output_path}protein_significantPeptides_counts_72hr.csv", index_label='Significant Peptides Category')
print("\nProtein Significant Peptides Counts:")
print(counts_table)

# slightly separate very small slices
explode = [0.02 if p < 0.05 else 0.0 for p in percent]

# Use a list of color strings (in the same order as `labels`) instead of a dict.
# colors_list = ['#c3c3c3', '#fca311', '#e54f2a'] # Autumn palette
colors_list = ["#e5e5e5", "#669bbc", "#003049"] # Blue-Gray palette

fig, ax = plt.subplots(figsize=(8, 6))
wedges, texts = ax.pie(
    counts.values,
    labels=None,             
    startangle=140,
    colors=colors_list,
    wedgeprops=dict(width=0.40, edgecolor='w'),
    explode=explode,
    counterclock=False
)

# Add percent labels placed on wedges
bbox_props = dict(boxstyle="round,pad=0.3", fc="white", ec="none", alpha=0.8)
for w, p in zip(wedges, percent):
    angle = (w.theta2 + w.theta1) / 2.0
    x, y = w.r * 0.75 * np.cos(np.deg2rad(angle)), w.r * 0.75 * np.sin(np.deg2rad(angle))
    ax.text(x, y, f"{p:.1%}", ha='center', va='center', fontsize=15, bbox=bbox_props, fontweight='bold')

# Center annotation: total proteins
ax.text(0, 0, f"Total\n{total}", ha='center', va='center', fontsize=15, fontweight='bold')

# Legend with counts and percentages
legend_labels = [f"{lab}: {cnt:,} ({pct:.1%})" for lab, cnt, pct in zip(labels, counts, percent)]
ax.legend(
    wedges, legend_labels,
    title="Number of Significant Peptides",
    loc="upper center",
    bbox_to_anchor=(0.5, -0.12),
    ncol=len(labels),
    fontsize=11,
    title_fontsize=12,
    frameon=False, 
    handletextpad=0.5,
    columnspacing=1.5,
)

ax.axis('equal')
plt.title('Distribution of Proteins by Number of \nSignificant Peptides from Linear Model', fontsize=16, fontweight='bold')
plt.tight_layout()
plots.finalize_plot( 
    fig, show=True, save=save_to_folder, 
    filename='significantPeptides_perProtein_Donut_72hr',
    filepath=figure_path,
    formats=figure_formats,
    transparent=transparent_bg,
    dpi=figure_dpi
)
```

```
Protein Significant Peptides Counts:
          Count  Percentage
sgnf_cat                   
None       2109      0.2945
Single     2477      0.3459
Multiple   2575      0.3596
```

This figure simply calculates the proportion of significant peptides detected per protein with splitting single or multiple significant peptides into their own categories. Shows ~29-30% of proteins without any significant peptides. And most have at least one significant peptide detected.

---

##### 02.6 Proteoform Category Distribution¶

Donut chart showing the overall distribution of proteins across proteoform categories: no forms detected, PTM-only, dPF-only, or both PTMs and dPFs identified.

```
# Plot donut chart for proteins with no additional forms, only dPFs, only PTMs, with Both
plot_data = protein_dpf_analysis[[ 'Gene', 'n_forms', 'n_canon', 'n_ptm', 'n_dPF' ]].copy()

without_forms = set(plot_data[(plot_data['n_forms'] == 0)].index)
with_PTM_only = set(plot_data[(plot_data['n_forms'] == 0) & (plot_data['n_ptm'] > 0)].index)
with_dPF_only = set(plot_data[(plot_data['n_forms'] > 0) & (plot_data['n_ptm'] == 0)].index)
with_both = set(plot_data[(plot_data['n_forms'] > 0) & (plot_data['n_ptm'] > 0)].index)
without_forms = without_forms.difference(with_PTM_only)
print(f"Proteins with no additional forms: {len(without_forms):,}")
print(f"Proteins with only PTMs: {len(with_PTM_only):,}")
print(f"Proteins with only dPFs: {len(with_dPF_only):,}")
print(f"Proteins with both PTMs and dPFs: {len(with_both):,}")

# Categorize proteins
labels = ['No Forms', 'PTM Only', 'dPF Only', 'Both']
counts = [
    len(without_forms),
    len(with_PTM_only),
    len(with_dPF_only),
    len(with_both)
]
# Calculate percentages
percent = np.array(counts) / sum(counts)

# Save the counts table
counts_table = pd.DataFrame({
    'Count': counts,
    'Percentage': percent
}, index=labels)
counts_table.to_csv(f"{output_path}protein_formCategories_counts_72hr.csv", index_label='Form Categories')
print("\nProtein Form Categories Counts:")
print(counts_table)

# slightly separate very small slices
explode = [0.02 if p < 0.05 else 0.0 for p in percent]
# Use a list of color strings (in the same order as `labels`) instead of a dict.
colors_list = ["#e5e5e5", "#a7c957", "#bc4749", "#967aa1"]

fig, ax = plt.subplots(figsize=(8, 6))
wedges, texts = ax.pie(
    counts,
    labels=None,             
    startangle=140,
    colors=colors_list,
    wedgeprops=dict(width=0.40, edgecolor='w'),
    explode=explode,
    counterclock=False
)
# Add percent labels placed on wedges
bbox_props = dict(boxstyle="round,pad=0.3", fc="white", ec="none", alpha=0.8)
for w, p in zip(wedges, percent):
    angle = (w.theta2 + w.theta1) / 2.0
    x, y = w.r * 0.75 * np.cos(np.deg2rad(angle)), w.r * 0.75 * np.sin(np.deg2rad(angle))
    ax.text(x, y, f"{p:.1%}", ha='center', va='center', fontsize=15, bbox=bbox_props, fontweight='bold')

# Center annotation: total proteins
ax.text(0, 0, f"Total\n{sum(counts)}", ha='center', va='center', fontsize=15, fontweight='bold')

# Legend with counts and percentages
legend_labels = [f"{lab}: {cnt:,} ({pct:.1%})" for lab, cnt, pct in zip(labels, counts, percent)]
ax.legend(
    wedges, legend_labels,
    title="Number of Proteins",
    loc="upper center",
    bbox_to_anchor=(0.5, -0.12),
    ncol=len(labels),
    fontsize=11,
    title_fontsize=12,
    frameon=False, 
    handletextpad=0.5,
    columnspacing=1.5,
)
ax.axis('equal')
plt.title('Distribution of Proteins by Additional Forms and PTMs', fontsize=16, fontweight='bold')
plt.tight_layout()
plots.finalize_plot( 
    fig, show=True, save=save_to_folder, 
    filename='formCategories_perProtein_Donut_72hr',
    filepath=figure_path,
    formats=figure_formats,
    transparent=transparent_bg,
    dpi=figure_dpi
)
```

```
Proteins with no additional forms: 2,109
Proteins with only PTMs: 2,818
Proteins with only dPFs: 1,255
Proteins with both PTMs and dPFs: 979

Protein Form Categories Counts:
          Count  Percentage
No Forms   2109      0.2945
PTM Only   2818      0.3935
dPF Only   1255      0.1753
Both        979      0.1367
```

```
=== Enhanced Peptide-UniProt Mapping with Miscleavage Detection ===
  Proteins to check: 5,052
  Peptides to process: 78,956
  Relevant UniProt annotations: 447,422
  Detecting miscleavage/peptide-peptide overlaps...

=== Mapping Summary ===
  Total entries: 248,015
    - UniProt annotations: 220,435
    - Miscleavage features: 9,675
    - No annotation: 17,905
  Processing time: 2.23 seconds

Sample of Peptide-UniProt Mapping with Miscleavage Detection:
```

|  | Protein | PeptideID | pepStart | pepEnd | feature\_category | feature | startpos | endpos | group | agent | note | score | dPF |
| --- | --- | --- | --- | --- | --- | --- | --- | --- | --- | --- | --- | --- | --- |
| 0 | A0A1B0GTU1 | 1 | 89 | 99 |  | No Annotation | -1 | -1 |  |  |  | 0.0000 | 0 |
| 1 | A0A1B0GTU1 | 2 | 115 | 124 |  | No Annotation | -1 | -1 |  |  |  | 0.0000 | 0 |
| 2 | A0A1B0GTU1 | 3 | 179 | 193 |  | No Annotation | -1 | -1 |  |  |  | 0.0000 | -1 |
| 3 | A0A1B0GTU1 | 4 | 714 | 724 |  | No Annotation | -1 | -1 |  |  |  | 0.0000 | -1 |
| 4 | A0A1B0GTU1 | 5 | 758 | 776 | Co- & Post-Translational Modifications | CROSSLNK | 765 | 765 | Ubiquitination/SUMOylation |  | Ubiquitination/SUMOylation at K765 | 1.0000 | 0 |
| 5 | A0A1W2PQL4 | 1 | 43 | 50 |  | No Annotation | -1 | -1 |  |  |  | 0.0000 | 0 |
| 6 | A0A1W2PQL4 | 2 | 234 | 242 |  | No Annotation | -1 | -1 |  |  |  | 0.0000 | 0 |
| 7 | A0A1W2PQL4 | 3 | 290 | 298 |  | No Annotation | -1 | -1 |  |  |  | 0.0000 | -1 |
| 8 | A0A1W2PQL4 | 4 | 346 | 354 |  | No Annotation | -1 | -1 |  |  |  | 0.0000 | 0 |
| 9 | A0A1W2PQL4 | 5 | 373 | 382 |  | No Annotation | -1 | -1 |  |  |  | 0.0000 | 0 |

Mapped about 250K annotations-peptides to generate a full Peptide-UniProt annotation map. As it is scene if a peptide doesn't have any annotation it is still kept in the map with empty annotations for `feature_category`, `group`, `agent`, etc. Additionally information such as score, dPF, and position are also kept for further analysis.

> This step is critical and I wanted to provide as much information from as many angles/sources into a single mapping table to facilitate downstream analyses linking peptide-level findings to functional annotations. However that you will see in upcoming parts, some are too complicated to automatically calculate so that I had to count manually for some of them.

---

#### 04. PTM and Proteoform Characterization¶

##### 04.1 PTM Annotation Analysis¶

Analyze peptides classified as PTM-only (dPF = -1) by examining their overlap with UniProt annotations. This reveals which PTM types are most commonly associated with single-peptide discordance patterns.

```
# Filter for only PTM peptides (dPF == -1)
final_res = peptide_uniprot_map[
    (peptide_uniprot_map['dPF'] == -1)
]

# Number of Proteins with No Annotation
no_annot_proteins = final_res[final_res['feature'] == 'No Annotation']['Protein'].unique()
with_annotation_proteins = final_res[final_res['feature'] != 'No Annotation']['Protein'].unique()
print(f"Number of Proteins with No Annotation: {len(no_annot_proteins)}")
print(f"Number of Proteins with Annotation: {len(with_annotation_proteins)}")

# Create a donut chart
data = [len(no_annot_proteins), len(with_annotation_proteins)]
# Labels
labels = ["No Annotation", "With Annotation"]
# Colors
colors = ["#e5e5e5", "#a7c957"]

# Save the counts table
counts_table = pd.DataFrame({
    'Count': data,
    'Percentage': np.array(data) / sum(data)
}, index=labels)
counts_table.to_csv(f"{output_path}proteins_withPTM_peptides_UniprotAnnotation_counts_72hr.csv", index_label='UniProt Annotation Availability')
# print("\nProteins with PTM Peptides UniProt Annotation Availability Counts:")
# print(counts_table)

fig, ax = plt.subplots(figsize=(8, 6))
wedges, texts = ax.pie(
    data,
    labels=None,             
    startangle=140,
    colors=colors,
    wedgeprops=dict(width=0.60, edgecolor='w'),
    explode=[0.02, 0.0],  # separate the first slice slightly
    counterclock=False
)
# Add percent labels placed on wedges
bbox_props = dict(boxstyle="round,pad=0.3", fc="white", ec="none", alpha=0.8)
for w, p in zip(wedges, np.array(data) / sum(data)):
    angle = (w.theta2 + w.theta1) / 2.0
    x, y = w.r * 0.75 * np.cos(np.deg2rad(angle)), w.r * 0.75 * np.sin(np.deg2rad(angle))
    ax.text(x, y, f"{p:.1%}", ha='center', va='center', fontsize=15, bbox=bbox_props, fontweight='bold')
# Center annotation: total proteins
ax.text(0, 0, f"Total\n{sum(data)}", ha='center', va='center', fontsize=15, fontweight='bold')
# Legend with counts and percentages
legend_labels = [f"{lab}: {cnt:,} ({pct:.1%})" for lab, cnt, pct in zip(labels, data, np.array(data) / sum(data))]
ax.legend(
    wedges, legend_labels,
    title="Number of Proteins",
    loc="upper center",
    bbox_to_anchor=(0.5, -0.12),
    ncol=len(labels),
    fontsize=11,
    title_fontsize=12,
    frameon=False, 
    handletextpad=0.5,
    columnspacing=1.5,
)
ax.axis('equal')
plt.title('Distribution of Proteins with PTM Peptides\nAnnotation Availability', fontsize=16, fontweight='bold')
plt.tight_layout()
plots.finalize_plot( 
    fig, show=True, save=save_to_folder, 
    filename='proteins_withPTM_peptides_UniprotAnnotation_Donut_72hr',
    filepath=figure_path,
    formats=figure_formats,
    transparent=transparent_bg,
    dpi=figure_dpi
)

print()

# Create a visualization to show the Number of Proteins with UniProt Annotations
fig, ax = plt.subplots(figsize=(5, 5))
sub_data = final_res[final_res['feature'] != 'No Annotation']

# Get counts of each feature type
feature_counts = sub_data['feature'].value_counts()

# Save the counts table
counts_table = pd.DataFrame({
    'Count': feature_counts
})
counts_table.to_csv(f"{output_path}proteins_withPTM_peptides_UniprotAnnotation_featureCounts_72hr.csv", index_label='Uniprot Feature Type')
# print("Proteins with PTM Peptides UniProt Annotation Feature Counts:")
# print(counts_table)

# Create a bar plot
feature_counts.plot(kind='barh', color="#a7c957", ax=ax, edgecolor='black')
# Add counts to on top of the bars
for i, v in enumerate(sub_data['feature'].value_counts()):
    ax.text(
        # v, i, str(v),
        # Adjust the position of the text
        v + 2.5, i-0.1, str(v),
        color='black', 
        ha='left', va='center', 
        rotation=0, fontsize=12, fontweight='bold'
    )
ax.set_xlabel("Number of Annotations", fontsize=12)
ax.set_ylabel("Uniprot Feature Type", fontsize=12)
ax.set_title("Annotations Matched to Single PTMs", fontsize=14, fontweight='bold')
ax.grid("both", linestyle="--", linewidth=0.75, alpha=0.5, color="lightgrey")
plt.tight_layout()
plt.show()
plots.finalize_plot( 
    fig, show=True, save=save_to_folder, 
    filename='proteins_withPTM_peptides_UniprotAnnotation_Bar_72hr',
    filepath=figure_path,
    formats=figure_formats,
    transparent=transparent_bg,
    dpi=figure_dpi
)
print()
print("Modified Residues (MOD_RES) Annotations:")
print((sub_data.loc[sub_data['feature']=='MOD_RES', 'group'].str.split(';').str[0]).value_counts())
print()
print("Cleavage Annotations:")
print((sub_data.loc[sub_data['feature']=='CLEAVAGE', 'group'].str.split(';').str[0]).value_counts())
print()
```

```
Number of Proteins with No Annotation: 1053
Number of Proteins with Annotation: 2892
```


```
Modified Residues (MOD_RES) Annotations:
group
Phosphorylation            3966
Acetylation                 467
Methylation                 271
Acylation                    37
Hydroxylation                15
ADP-ribosylation              3
Nitration/Nitrosylation       3
Citrullination                2
Name: count, dtype: int64

Cleavage Annotations:
group
Proteolytic Cleavage    1283
Name: count, dtype: int64
```

There are 4484 peptides classified as PTM-only (dPF = -1) across all proteins. Out of these, 3245 peptides (approximately 72%) have at least one annotation found within the peptide. This means quite a significant portion of PTM-only peptides can be linked to known features, however there are still ~28% of PTM-only peptides that do not overlap with any known annotations, indicating potential novel modifications or limitations in current annotation databases, or simply peptides behaving like technical outliers without biological basis.

> Note: There can be multiple peptides as PTMs in a single proteins, thats why the 4484 total doesn't correspond to the number of proteins with PTM-only and Both number of 4216.

Of those 3245 PTM-only peptides with annotations quite a large number of annotations is observed the numbers can be see in the second figure above. With the largest category being the MOD\_RES which makes sense as these are post-translational modifications. Other large categories like CROSSLNK, VAR\_SEQ, CLEAVAGE also make sense in the context of PTMs.

> Note: Another thing is that a peptide can host many annotations, which is why we see very large number of annotations compared to peptides.

Looking closely at the MOD\_RES annotations which are directly related to PTMs we see a wide variety of modification types associated with these PTM-only peptides. The most common modifications include Phosphorylation (Phosphoserine, Phosphothreonine, and Phosphotyrosine), which are well-known regulatory PTMs involved in signaling pathways. They are overwhelmingly represented here, indicating that many of the PTM-only peptides may be phosphorylated forms of the canonical sequences.

Additionally I have checked for what are the cleavage annotations, and only proteolytic cleavages are observed (Signal peptide cleavage, Transit peptide cleavage, and Propeptide cleavage). Which indicates that some of these PTM-only peptides may arise from specific proteolytic processing events rather than chemical modifications.

---

##### 04.2 Modification Type Distribution¶

Breakdown of specific modification types mapped to identified PTMs. The MOD\_RES annotations reveal the chemical nature of post-translational modifications associated with discordant peptide behavior.

```
# Data preparation
sub_data = final_res[final_res['feature'] != 'No Annotation']
mod_res_data = sub_data[sub_data['feature'] == 'MOD_RES']['group'].str.split(';').str[0].value_counts()

# Calculate percentages
mod_total = mod_res_data.sum()
mod_percentages = (mod_res_data / mod_total * 100)

# Save the counts table
counts_table = pd.DataFrame({
    'Count': mod_res_data,
    'Percentage': mod_percentages
})
counts_table.to_csv(f"{output_path}mod_res_modificationTypes_counts_72hr.csv", index_label='Modification Type')

# Create figure
fig, ax = plt.subplots(figsize=(12, 4))

# Create stacked horizontal bar
left = 0
colors_mod = plt.cm.Set3(np.linspace(0, 1, len(mod_res_data)))

for i, (mod_type, count) in enumerate(mod_res_data.items()):
    width = mod_percentages.iloc[i]
    bar = ax.barh(0, width, left=left, color=colors_mod[i], 
                  edgecolor='white', linewidth=2, height=0.6)
    
    # Add percentage label if there's enough space (>3% threshold)
    if width > 3:
        ax.text(left + width/2, 0, f'{width:.1f}%\n({count:,})', 
               ha='center', va='center', fontsize=11, fontweight='bold',
               bbox=dict(boxstyle="round,pad=0.2", facecolor='white', alpha=0.8))
    
    left += width

# Styling
ax.set_xlim(0, 100)
ax.set_ylim(-0.5, 0.5)
ax.set_title(
    f'Distribution of Modification Types mapped to identified PTMs (n={mod_total:,})', 
    fontsize=16, fontweight='bold', pad=20
)
ax.set_yticks([])
ax.set_xticks([])

# Remove spines for cleaner look
ax.spines['top'].set_visible(False)
ax.spines['right'].set_visible(False)
ax.spines['left'].set_visible(False)
ax.spines['bottom'].set_visible(False)

# Add grid for easier reading
ax.grid(axis='x', linestyle='--', alpha=0.3)

# Create legend with counts
legend_elements = [
    plt.Rectangle((0,0),1,1, facecolor=colors_mod[i], label=f'{mod_type}: {count:,} ({mod_percentages.iloc[i]:.1f}%)')
    for i, (mod_type, count) in enumerate(mod_res_data.items())
]

ax.legend(
    handles=legend_elements, 
    loc='center', bbox_to_anchor=(0.5, -0.25), ncol=4, 
    fontsize=10, frameon=False, columnspacing=1.5
)

plt.tight_layout()

# Save the figure
plots.finalize_plot( 
    fig, show=True, save=save_to_folder, 
    filename='mod_res_modifications_stacked_bar_72hr',
    filepath=figure_path,
    formats=figure_formats,
    transparent=transparent_bg,
    dpi=figure_dpi
)

print(f"MOD_RES Analysis:")
print(f"Total modifications: {mod_total:,}")
for mod_type, count in mod_res_data.items():
    print(f"• {mod_type}: {count:,} ({count/mod_total*100:.1f}%)")
```

```
MOD_RES Analysis:
Total modifications: 4,764
• Phosphorylation: 3,966 (83.2%)
• Acetylation: 467 (9.8%)
• Methylation: 271 (5.7%)
• Acylation: 37 (0.8%)
• Hydroxylation: 15 (0.3%)
• ADP-ribosylation: 3 (0.1%)
• Nitration/Nitrosylation: 3 (0.1%)
• Citrullination: 2 (0.0%)
```

Above is the detailed distribution of specific MOD\_RES modification types observed among the PTM-only peptides. Phosphorylation events (Phosphoserine, Phosphothreonine, Phosphotyrosine) dominate the landscape, highlighting their prevalence in regulatory processes. Other modifications such as Acetylation, Methylation are with small but notable counts, indicating a diversity of PTM types contributing to peptide discordance.

---

##### 04.3 Differential Proteoform (dPF) Analysis¶

Analyze peptides classified as differential proteoforms (dPF > 0), examining annotation coverage and categorizing reasons for proteoform identification based on manual review results.

```
# Smartly build dPF color palette for all data
dPF_colors = {
    # PTM-Only
    -1: "#a7c957",  # PTM Only
    # Canonical
    0: "#c3c3c3",  # Canonical
    # Proteoforms
    # Calculate reds (#bc4749) for dPF > 0
    **({} if (len(summary_data) == 0 or int(summary_data['dPF'].max()) <= 0) 
        else {i: sns.color_palette("Reds_r", n_colors=int(summary_data['dPF'].max())).as_hex()[i-1] for i in range(1, int(summary_data['dPF'].max()) + 1)})
}
print(f"Proteoform Group Colors: {dPF_colors}")
# sns.palplot(list(dPF_colors.values()));
print()

protein_with_dpf = protein_dpf_analysis.loc[
    (protein_dpf_analysis['n_forms'] > 0),
    'Protein'
].copy()

dPF_uniprot_maps = peptide_uniprot_map[
    (peptide_uniprot_map['dPF'] > 0)
].copy()
print(f"Number of Proteins with dPFs: {len(protein_with_dpf):,}")
print(f"Number of generated mappings: {len(dPF_uniprot_maps):,}")
print(f"Number of dPF Annotations found: {len(dPF_uniprot_maps[dPF_uniprot_maps['feature'] != 'No Annotation']):,}")
# dPF_uniprot_maps.head(20)

# protein_dpf_analysis.sort_values('dPF_pct', ascending=False).head(30)
```

```
Proteoform Group Colors: {-1: '#a7c957', 0: '#c3c3c3', 1: '#cb181d', 2: '#fb6b4b', 3: '#fcbca2'}

Number of Proteins with dPFs: 2,234
Number of generated mappings: 94,781
Number of dPF Annotations found: 89,445
```

```
# # Manual Check Results for dPFs
# # Reasons Categorized
# high_level_dict = {
#     'No Annotation': 192,
#     'Mixed/Unclear Reasons': 857,
#     'Clear Reasons': 405,
# }
# # Categories of Clear Reasons
# clear_reasons_dict = {
#     'Missed Cleavages': 193,
#     'Isoforms': 51,
#     'Multiple PTMs': 37,
#     'N-terminal': 34,
#     'C-terminal': 31,
#     'Variants': 20,
#     'Mutagen': 20,
#     'Cleavage': 19,
# }

# # Number of dPFs 
# dPF_counts = protein_dpf_analysis['n_forms'].value_counts().sort_index()
# print("dPF Counts per Protein:")
# for n_forms, count in dPF_counts.items():
#     print(f"  {n_forms}: {count:,}")

# print("Number of Unique dPFs > 0:", protein_dpf_analysis[protein_dpf_analysis['n_forms'] > 0]['n_forms'].sum())
# print()

# print("Manual Check Results for dPFs:")
# for reason, count in high_level_dict.items():
#     print(f"  {reason}: {count:,}")

# # Donut chart for dPF annotation reasons
# labels = list(high_level_dict.keys())
# counts = list(high_level_dict.values())
# colors = ["#e5e5e5",  '#fdcab5', "#bc4749"]
# fig, ax = plt.subplots(figsize=(8, 6))
# wedges, texts = ax.pie(
#     counts,
#     labels=None,             
#     startangle=140,
#     colors=colors,
#     wedgeprops=dict(width=0.60, edgecolor='w'),
#     explode=[0.02 if c < 0.05 * sum(counts) else 0.0 for c in counts],
#     counterclock=False
# )
# # Add percent labels placed on wedges
# bbox_props = dict(boxstyle="round,pad=0.3", fc="white", ec="none", alpha=0.8)
# for w, p in zip(wedges, np.array(counts) / sum(counts)):
#     angle = (w.theta2 + w.theta1) / 2.0
#     x, y = w.r * 0.75 * np.cos(np.deg2rad(angle)), w.r * 0.75 * np.sin(np.deg2rad(angle))
#     ax.text(x, y, f"{p:.1%}", ha='center', va='center', fontsize=15, bbox=bbox_props, fontweight='bold')
# # Center annotation: total proteins
# ax.text(0, 0, f"Total\n{sum(counts)}", ha='center', va='center', fontsize=15, fontweight='bold')
# # Legend with counts and percentages
# legend_labels = [f"{lab}: {cnt:,} ({pct:.1%})" for lab, cnt, pct in zip(labels, counts, np.array(counts) / sum(counts))]
# ax.legend(
#     wedges, legend_labels,
#     title="dPF Annotation Reasons",
#     loc="upper center",
#     bbox_to_anchor=(0.5, -0.12),
#     ncol=len(labels),
#     fontsize=11,
#     title_fontsize=12,
#     frameon=False, 
#     handletextpad=0.5,  
#     columnspacing=1.5,
# )
# ax.axis('equal')
# plt.title('Distribution of dPF Annotation Reasons', fontsize=16, fontweight='bold')
# plt.tight_layout()
# plots.finalize_plot( 
#     fig, show=True, save=save_to_folder, 
#     filename='dPF_annotation_reasons_Donut_72hr',
#     filepath=figure_path,
#     formats=figure_formats,
#     transparent=transparent_bg,
#     dpi=figure_dpi
# )

# # Create a horizontal bar chart for Clear Reasons
# clear_reasons = list(clear_reasons_dict.keys())
# clear_counts = list(clear_reasons_dict.values())
# clear_color = "#bc4749"
# fig, ax = plt.subplots(figsize=(5, 5))
# bars = ax.barh(clear_reasons, clear_counts, color=clear_color, edgecolor='black')
# # Add counts to on top of the bars
# for i, v in enumerate(clear_counts):
#     ax.text(
#         # v, i, str(v),
#         # Adjust the position of the text
#         v + 2.5, i-0.1, str(v),
#         color='black', 
#         ha='left', va='center', 
#         rotation=0, fontsize=12, fontweight='bold'
#     )
# ax.set_xlabel("Number of dPFs", fontsize=12)
# ax.set_ylabel("Clear Reason Categories", fontsize=12)
# ax.set_title("Breakdown of Clear Reasons for dPF Annotations", fontsize=14, fontweight='bold')
# ax.grid("both", linestyle="--", linewidth=0.75, alpha=0.5, color="lightgrey")
# plt.tight_layout()
# plots.finalize_plot( 
#     fig, show=True, save=save_to_folder, 
#     filename='dPF_annotation_clear_reasons_Bar_72hr',
#     filepath=figure_path,
#     formats=figure_formats,
#     transparent=transparent_bg,
#     dpi=figure_dpi
# )
```

```
Number of Candidate Proteins with consistent (non-gap) dPFs: 216
Candidate Proteins: ['A4D1U4', 'A6NJ78', 'B2RTY4', 'O00231', 'O00425', 'O00459', 'O14772', 'O14939', 'O15075', 'O43272', 'O43524', 'O43660', 'O43663', 'O60294', 'O60343', 'O60502', 'O60506', 'O60890', 'O94776', 'O94813', 'O95363', 'O95782', 'O95817', 'O96005', 'P02768', 'P04183', 'P0DPH7', 'P0DPH8', 'P12081', 'P21675', 'P23469', 'P25685', 'P27540', 'P28074', 'P28288', 'P31146', 'P31751', 'P31942', 'P33240', 'P35609', 'P35813', 'P40937', 'P42680', 'P42765', 'P48200', 'P48556', 'P49590', 'P49674', 'P49755', 'P50479', 'P51116', 'P51178', 'P51398', 'P51589', 'P52907', 'P53804', 'P54136', 'P55060', 'P60983', 'P61158', 'P61964', 'P62191', 'P62491', 'P62847', 'P68363', 'Q00653', 'Q02241', 'Q03701', 'Q07890', 'Q07960', 'Q08345', 'Q08945', 'Q0JRZ9', 'Q12769', 'Q12851', 'Q12905', 'Q13084', 'Q13131', 'Q13228', 'Q13438', 'Q13618', 'Q14114', 'Q15003', 'Q15125', 'Q15382', 'Q15631', 'Q15738', 'Q15833', 'Q15907', 'Q16401', 'Q16566', 'Q16587', 'Q16665', 'Q16775', 'Q2M2I8', 'Q3SY52', 'Q4G0N4', 'Q53GQ0', 'Q5BJF2', 'Q5EBL4', 'Q5H8A4', 'Q5IFJ8', 'Q5JVF3', 'Q5QJE6', 'Q5SY16', 'Q5W111', 'Q641Q2', 'Q68DA7', 'Q6NZY4', 'Q6P444', 'Q6Q0C0', 'Q6VY07', 'Q71U36', 'Q7Z5Q1', 'Q7Z7A1', 'Q86TJ2', 'Q86UX6', 'Q86VW0', 'Q86XK2', 'Q86XP1', 'Q8IUS5', 'Q8IVL5', 'Q8IXQ4', 'Q8IY33', 'Q8N163', 'Q8N392', 'Q8N556', 'Q8N5C6', 'Q8N5C8', 'Q8N6H7', 'Q8ND82', 'Q8NFT2', 'Q8NG68', 'Q8NI77', 'Q8TAA9', 'Q8TBZ6', 'Q8TCN5', 'Q8TDB6', 'Q8TEP8', 'Q8TEU7', 'Q8WUF5', 'Q92733', 'Q92766', 'Q92802', 'Q92922', 'Q93034', 'Q96B23', 'Q96BT7', 'Q96HV5', 'Q96IV0', 'Q96JC4', 'Q96PD2', 'Q96QU8', 'Q96S21', 'Q96S66', 'Q96T37', 'Q99459', 'Q99460', 'Q99567', 'Q99623', 'Q99986', 'Q9BPX6', 'Q9BQE3', 'Q9BQG2', 'Q9BRX2', 'Q9BW04', 'Q9BWF3', 'Q9BZV1', 'Q9C0C7', 'Q9GZZ9', 'Q9H223', 'Q9H2U1', 'Q9H3M7', 'Q9H5H4', 'Q9H7E2', 'Q9H992', 'Q9H9B1', 'Q9H9T3', 'Q9HAV7', 'Q9NP73', 'Q9NQ84', 'Q9NU19', 'Q9NVP1', 'Q9NW82', 'Q9NX08', 'Q9NXZ2', 'Q9NYK5', 'Q9NZJ0', 'Q9NZN5', 'Q9P209', 'Q9P2D1', 'Q9UGN5', 'Q9UHV9', 'Q9UIC8', 'Q9UJW7', 'Q9UKE5', 'Q9UKI8', 'Q9UKM7', 'Q9ULJ3', 'Q9ULV4', 'Q9UNM6', 'Q9UPM8', 'Q9UQN3', 'Q9Y2E5', 'Q9Y2I1', 'Q9Y2R4', 'Q9Y2S7', 'Q9Y314', 'Q9Y3D9', 'Q9Y448', 'Q9Y4E1', 'Q9Y4X0', 'Q9Y639', 'Q9Y692', 'Q9Y6E0', 'Q9Y6Y8']
```

This is one category that looked promising and interesting to look at. The idea here is that if all the dPF peptides for a given protein are consecutive along the protein sequence, it strongly suggests that these peptides represent a true proteoform region rather than scattered technical artifacts. Consecutive peptides indicate a coherent segment of the protein that may be differentially regulated or modified, supporting the biological relevance of the observed proteoform discordance.

> Note: I have used these to start the manual checkups, as well as the interesting and biologically relevant proteins that I know of from the hypoxia experiment.

---

#### 05. Case Studies and Indivudal Examination of Proteins¶

##### 05.1 Relevant Genes and Proteins from Hypoxia Experiment¶

Based on the hypoxia experiment detailed in the paper, the following genes and proteins are of particular interest. The experiment exposed H358 lung cancer cells to hypoxic conditions (1% oxygen) and analyzed changes in their proteome. Key enzymes related to glycolysis and the glyoxalase system showed altered abundance, with an inverse correlation between hypoxic markers (LDHA, HK2, GAPDH upregulation) and GLO2 downregulation.

**Key Proteins of Interest:**

- **LDHA** (P00338): Lactate production, upregulated in hypoxia
- **HK2** (P52790): Glycolysis entry, upregulated in hypoxia
- **GAPDH** (P04406): Glycolysis, upregulated in hypoxia
- **GLO1** (Q04760): Methylglyoxal metabolism, unaffected
- **HAGH/GLO2** (Q16775): Methylglyoxal metabolism, downregulated in hypoxia

```
glycolytic_up_by_HIF_1 = [
    'P00338', #LDHA                 
    'P52789', #HK2                  
    'P04406', #GAPDH                
    'Q04760', #GLO1                 
    'Q16775', #HAGH (GLO2)         
]

is_demo = False # False if want to save figures

pass_dict = {
    'data': {
        'uniprot': uniprot_data,
        'detailed': test_data,
        'summary': summary_data,
    },
    'styles': {
        'dpf_colors': dPF_colors,
    },
    'config': {
        'is_demo': is_demo,
        'figure_path': figure_path,
        'figure_formats': figure_formats,
        'transparent_bg': transparent_bg,
        'figure_dpi': figure_dpi,
        'str_add': '72hr',
    }
}
```

###### 05.1.1 LDHA (P00338)¶

The LDHA in the original hypoxia experiment showed significant upregulation under hypoxic conditions, consistent with its role in anaerobic metabolism.

**ProteoForge Protein Summary Visualization**

Generate detailed peptide-level visualization showing intensity patterns, significance, and cluster assignments for selected proteins of interest. This enables in-depth examination of proteoform behavior under hypoxic conditions.

This figure summarizes how the ProteoForge sees the protein, and at raw, adjusted intentsities peptides across conditions act, as well as the p-value of the interaction model per peptide as well as the clustering assignments.

```
current_protein = 'P00338'
cur_gene = protein_dpf_analysis.loc[
    protein_dpf_analysis['Protein'] == current_protein, 'Gene'
].values[0]
print(f"Current Protein: {current_protein} ({cur_gene})")
plots.detailed_peptide_with_clusters(
    data=test_data,
    cur_protein=current_protein,
    pThr=pThr,
    pvalue_col='adj_pval',
    cluster_col='ClusterID',
    rawIntensity_col='log10Intensity',
    adjIntensity_col='AdjIntensity',
    condition_palette=condition_colors,
    
    save = not is_demo,
    show=True,
    filename=f'detailed_peptide_look_{cur_gene}_72hr',
    filepath=figure_path,
    fileformats=figure_formats,
    transparent=transparent_bg,
    dpi=figure_dpi
)
```

```
Current Protein: P00338 (LDHA)
```

From the adjusted intensity there is a pattern where hypoxia is slightly higher than normoxia, but the difference is not very pronounced. And only one PTM peptide is observed, where the difference is very pronounced between hypoxia and normoxia. For that peptide the intensity is not high in raw intensity, however the normoxia is very low, this could peptide that is imputed with low-values due to it being completely missing in normoxia samples.

---

**Detailed Protein Annotation Visualization**

Create comprehensive protein-level visualization integrating UniProt annotations with peptide coverage. Multiple annotation layers show sequence features, modifications, and functional domains alongside identified peptides colored by dPF category.

```
plots.annotations_on_protein(
    target_protein=current_protein,
    control_package=pass_dict
)
```

**Various Data Views around the Peptides of Interest(s)**

LDAH's 16th peptide is marked as potential PTM, let's see what are the annotations mapped to that region of the protein.

```
peptide_uniprot_map[
    (peptide_uniprot_map['Protein']==current_protein) & # Current Protein
    (peptide_uniprot_map['PeptideID'].isin([11, 13, 12, 16])) # Peptide to inspect
]
```

|  | Protein | PeptideID | pepStart | pepEnd | feature\_category | feature | startpos | endpos | group | agent | note | score | dPF |
| --- | --- | --- | --- | --- | --- | --- | --- | --- | --- | --- | --- | --- | --- |
| 27448 | P00338 | 11 | 133 | 149 | Co- & Post-Translational Modifications | CARBOHYD | 137 | 137 | O-linked |  | O-linked at S137 | 1.0000 | 1 |
| 27449 | P00338 | 11 | 133 | 149 | Cleavage Analysis | MISCLEAVAGE | 133 | 149 | Miscleavage | ProteoForge\_Analysis | Overlap with PeptideID 12 (fraction: 1.0) | 1.0000 | 1 |
| 27450 | P00338 | 11 | 133 | 149 | Co- & Post-Translational Modifications | MOD\_RES | 145 | 145 | Phosphorylation |  | Phosphorylation at Y145 | 1.0000 | 1 |
| 27451 | P00338 | 11 | 133 | 149 | Sequence Heterogeneity & Isoforms | VAR\_SEQ | 82 | 139 | Missing | 4 | Missing (in isoform 4) | 3.0000 | 1 |
| 27452 | P00338 | 12 | 133 | 155 | Co- & Post-Translational Modifications | CARBOHYD | 137 | 137 | O-linked |  | O-linked at S137 | 1.0000 | 2 |
| 27453 | P00338 | 12 | 133 | 155 | Co- & Post-Translational Modifications | CROSSLNK | 155 | 155 | Ubiquitination/SUMOylation |  | Ubiquitination/SUMOylation at K155 | 1.0000 | 2 |
| 27454 | P00338 | 12 | 133 | 155 | Cleavage Analysis | MISCLEAVAGE | 133 | 149 | Partial\_Overlap | ProteoForge\_Analysis | Overlap with PeptideID 11 (fraction: 0.739) | 0.7391 | 2 |
| 27455 | P00338 | 12 | 133 | 155 | Co- & Post-Translational Modifications | MOD\_RES | 145 | 145 | Phosphorylation |  | Phosphorylation at Y145 | 1.0000 | 2 |
| 27456 | P00338 | 12 | 133 | 155 | Sequence Heterogeneity & Isoforms | VAR\_SEQ | 82 | 139 | Missing | 4 | Missing (in isoform 4) | 3.0000 | 2 |
| 27457 | P00338 | 13 | 170 | 177 | Protein Processing & Maturation | CLEAVAGE | 177 | 177 | Proteolytic Cleavage | trypsin 1 | N-Ph cleavage at R177 by trypsin 1 | 2.0000 | 1 |
| 27458 | P00338 | 13 | 170 | 177 | Co- & Post-Translational Modifications | MOD\_RES | 172 | 172 | Phosphorylation |  | Phosphorylation at Y172 | 1.0000 | 1 |
| 27479 | P00338 | 16 | 229 | 243 | Protein Processing & Maturation | CLEAVAGE | 231 | 231 | Proteolytic Cleavage | peptidyl-Lys metallopeptidase | N-Ph cleavage at H231 by peptidyl-Lys metallop... | 2.0000 | 2 |
| 27480 | P00338 | 16 | 229 | 243 | Protein Processing & Maturation | CLEAVAGE | 232 | 232 | Proteolytic Cleavage | trypsin 1 | N-Ph cleavage at K232 by trypsin 1 | 2.0000 | 2 |
| 27481 | P00338 | 16 | 229 | 243 | Protein Processing & Maturation | CLEAVAGE | 235 | 235 | Proteolytic Cleavage | cathepsin K | Ph cleavage at V235 by cathepsin K | 2.0000 | 2 |
| 27482 | P00338 | 16 | 229 | 243 | Protein Processing & Maturation | CLEAVAGE | 236 | 236 | Proteolytic Cleavage | cathepsin K | Ph cleavage at E236 by cathepsin K | 2.0000 | 2 |
| 27483 | P00338 | 16 | 229 | 243 | Protein Processing & Maturation | CLEAVAGE | 236 | 236 | Proteolytic Cleavage | cathepsin S | N-Ph cleavage at E236 by cathepsin S | 2.0000 | 2 |
| 27484 | P00338 | 16 | 229 | 243 | Protein Processing & Maturation | CLEAVAGE | 242 | 242 | Proteolytic Cleavage | peptidyl-Lys metallopeptidase | N-Ph cleavage at I242 by peptidyl-Lys metallop... | 2.0000 | 2 |
| 27485 | P00338 | 16 | 229 | 243 | Co- & Post-Translational Modifications | CROSSLNK | 232 | 232 | Ubiquitination/SUMOylation |  | Ubiquitination/SUMOylation at K232 | 1.0000 | 2 |
| 27486 | P00338 | 16 | 229 | 243 | Co- & Post-Translational Modifications | CROSSLNK | 243 | 243 | Ubiquitination/SUMOylation |  | Ubiquitination/SUMOylation at K243 | 1.0000 | 2 |
| 27487 | P00338 | 16 | 229 | 243 | Cleavage Analysis | MISCLEAVAGE | 233 | 243 | Partial\_Overlap | ProteoForge\_Analysis | Overlap with PeptideID 17 (fraction: 0.733) | 0.7333 | 2 |
| 27488 | P00338 | 16 | 229 | 243 | Co- & Post-Translational Modifications | MOD\_RES | 232 | 232 | Acetylation |  | N6-acetyllysine | 3.0000 | 2 |
| 27489 | P00338 | 16 | 229 | 243 | Co- & Post-Translational Modifications | MOD\_RES | 232 | 232 | Acetylation |  | Acetylation at K232 | 1.0000 | 2 |
| 27490 | P00338 | 16 | 229 | 243 | Co- & Post-Translational Modifications | MOD\_RES | 237 | 237 | Phosphorylation |  | Phosphorylation at S237 | 1.0000 | 2 |
| 27491 | P00338 | 16 | 229 | 243 | Co- & Post-Translational Modifications | MOD\_RES | 239 | 239 | Phosphorylation |  | Phosphotyrosine | 3.0000 | 2 |
| 27492 | P00338 | 16 | 229 | 243 | Co- & Post-Translational Modifications | MOD\_RES | 239 | 239 | Phosphorylation |  | Phosphorylation at Y239 | 2.0000 | 2 |
| 27493 | P00338 | 16 | 229 | 243 | Co- & Post-Translational Modifications | MOD\_RES | 239 | 239 | Phosphorylation |  | Phosphorylation at Y239 | 2.0000 | 2 |
| 27494 | P00338 | 16 | 229 | 243 | Co- & Post-Translational Modifications | MOD\_RES | 243 | 243 | Acetylation |  | N6-acetyllysine | 3.0000 | 2 |
| 27495 | P00338 | 16 | 229 | 243 | Co- & Post-Translational Modifications | MOD\_RES | 243 | 243 | Acetylation |  | Acetylation at K243 | 1.0000 | 2 |
| 27496 | P00338 | 16 | 229 | 243 | Sequence Heterogeneity & Isoforms | VAR\_SEQ | 230 | 274 | Modified | 2 | Modified (in isoform 2) | 3.0000 | 2 |
| 27497 | P00338 | 16 | 229 | 243 | Sequence Heterogeneity & Isoforms | VAR\_SEQ | 237 | 241 | Modified | 5 | Modified (in isoform 5) | 3.0000 | 2 |
| 27498 | P00338 | 16 | 229 | 243 | Sequence Heterogeneity & Isoforms | VAR\_SEQ | 242 | 332 | Missing | 5 | Missing (in isoform 5) | 3.0000 | 2 |

#### 05.1.2 HK2 (P52789)¶

```
current_protein = 'P52789'
cur_gene = protein_dpf_analysis.loc[
    protein_dpf_analysis['Protein'] == current_protein, 'Gene'
].values[0]
print(f"Current Protein: {current_protein} ({cur_gene})")
plots.detailed_peptide_with_clusters(
    data=test_data,
    cur_protein=current_protein,
    pThr=pThr,
    pvalue_col='adj_pval',
    cluster_col='ClusterID',
    rawIntensity_col='log10Intensity',
    adjIntensity_col='AdjIntensity',
    condition_palette=condition_colors,
    
    save = not is_demo,
    show=True,
    filename=f'detailed_peptide_look_{cur_gene}_72hr',
    filepath=figure_path,
    fileformats=figure_formats,
    transparent=transparent_bg,
    dpi=figure_dpi
)
```

```
Current Protein: P52789 (HK2)
```

```
plots.annotations_on_protein(
    target_protein=current_protein,
    control_package=pass_dict
)
```

```
peptide_uniprot_map[
    (peptide_uniprot_map['Protein']==current_protein) & # Current Protein
    (peptide_uniprot_map['PeptideID'].isin([13])) # Peptide to inspect
]
```

|  | Protein | PeptideID | pepStart | pepEnd | feature\_category | feature | startpos | endpos | group | agent | note | score | dPF |
| --- | --- | --- | --- | --- | --- | --- | --- | --- | --- | --- | --- | --- | --- |
| 92474 | P52789 | 13 | 338 | 346 | Co- & Post-Translational Modifications | CROSSLNK | 346 | 346 | Ubiquitination/SUMOylation |  | Ubiquitination/SUMOylation at K346 | 1.0000 | -1 |
| 92475 | P52789 | 13 | 338 | 346 | Cleavage Analysis | MISCLEAVAGE | 338 | 346 | Miscleavage | ProteoForge\_Analysis | Overlap with PeptideID 14 (fraction: 1.0) | 1.0000 | -1 |
| 92476 | P52789 | 13 | 338 | 346 | Co- & Post-Translational Modifications | MOD\_RES | 340 | 340 | Phosphorylation |  | Phosphorylation at S340 | 1.0000 | -1 |

###### 05.1.2 GAPDH (P04406)¶

```
current_protein = 'P04406'
cur_gene = protein_dpf_analysis.loc[
    protein_dpf_analysis['Protein'] == current_protein, 'Gene'
].values[0]
print(f"Current Protein: {current_protein} ({cur_gene})")
plots.detailed_peptide_with_clusters(
    data=test_data,
    cur_protein=current_protein,
    pThr=pThr,
    pvalue_col='adj_pval',
    cluster_col='ClusterID',
    rawIntensity_col='log10Intensity',
    adjIntensity_col='AdjIntensity',
    condition_palette=condition_colors,
    
    save = not is_demo,
    show=True,
    filename=f'detailed_peptide_look_{cur_gene}_72hr',
    filepath=figure_path,
    fileformats=figure_formats,
    transparent=transparent_bg,
    dpi=figure_dpi
)
```

```
Current Protein: P04406 (GAPDH)
```

```
plots.annotations_on_protein(
    target_protein=current_protein,
    control_package=pass_dict
)
```

```
peptide_uniprot_map[
    (peptide_uniprot_map['Protein']==current_protein) & # Current Protein
    (peptide_uniprot_map['PeptideID'].isin([15, 17, 18, 20])) # Peptide to inspect
]
```

|  | Protein | PeptideID | pepStart | pepEnd | feature\_category | feature | startpos | endpos | group | agent | note | score | dPF |
| --- | --- | --- | --- | --- | --- | --- | --- | --- | --- | --- | --- | --- | --- |
| 32380 | P04406 | 15 | 216 | 227 | Protein Processing & Maturation | CLEAVAGE | 218 | 218 | Proteolytic Cleavage | cathepsin L | N-Ph cleavage at G218 by cathepsin L | 2.0000 | 1 |
| 32381 | P04406 | 15 | 216 | 227 | Protein Processing & Maturation | CLEAVAGE | 218 | 218 | Proteolytic Cleavage | cathepsin S | N-Ph cleavage at G218 by cathepsin S | 2.0000 | 1 |
| 32382 | P04406 | 15 | 216 | 227 | Protein Processing & Maturation | CLEAVAGE | 218 | 218 | Proteolytic Cleavage | cathepsin V | N-Ph cleavage at G218 by cathepsin V | 2.0000 | 1 |
| 32383 | P04406 | 15 | 216 | 227 | Protein Processing & Maturation | CLEAVAGE | 218 | 218 | Proteolytic Cleavage | matrix metallopeptidase-3 | Ph cleavage at G218 by matrix metallopeptidase-3 | 2.0000 | 1 |
| 32384 | P04406 | 15 | 216 | 227 | Protein Processing & Maturation | CLEAVAGE | 219 | 219 | Proteolytic Cleavage | trypsin 1 | N-Ph cleavage at K219 by trypsin 1 | 2.0000 | 1 |
| ... | ... | ... | ... | ... | ... | ... | ... | ... | ... | ... | ... | ... | ... |
| 32542 | P04406 | 20 | 310 | 334 | Co- & Post-Translational Modifications | MOD\_RES | 334 | 334 | Methylation |  | N6,N6-dimethyllysine | 3.0000 | 1 |
| 32543 | P04406 | 20 | 310 | 334 | Co- & Post-Translational Modifications | MOD\_RES | 334 | 334 | Acetylation |  | Acetylation at K334 | 1.0000 | 1 |
| 32544 | P04406 | 20 | 310 | 334 | Co- & Post-Translational Modifications | MOD\_RES | 334 | 334 | Methylation |  | Methylation at K334 | 3.0000 | 1 |
| 32545 | P04406 | 20 | 310 | 334 | Co- & Post-Translational Modifications | MOD\_RES | 334 | 334 | Methylation |  | Methylation at K334 | 3.0000 | 1 |
| 32546 | P04406 | 20 | 310 | 334 | Sequence Heterogeneity & Isoforms | MUTAGEN | 320 | 320 | No Significant Effect | Y->F | No effect on free radical-induced aggregation. | 3.0000 | 1 |

111 rows × 13 columns

###### 05.1.2 GLO1 (Q04760)¶

```
current_protein = 'Q04760'
cur_gene = protein_dpf_analysis.loc[
    protein_dpf_analysis['Protein'] == current_protein, 'Gene'
].values[0]
print(f"Current Protein: {current_protein} ({cur_gene})")
plots.detailed_peptide_with_clusters(
    data=test_data,
    cur_protein=current_protein,
    pThr=pThr,
    pvalue_col='adj_pval',
    cluster_col='ClusterID',
    rawIntensity_col='log10Intensity',
    adjIntensity_col='AdjIntensity',
    condition_palette=condition_colors,
    
    save = not is_demo,
    show=True,
    filename=f'detailed_peptide_look_{cur_gene}_72hr',
    filepath=figure_path,
    fileformats=figure_formats,
    transparent=transparent_bg,
    dpi=figure_dpi
)
```

```
Current Protein: Q04760 (GLO1)
```

```
plots.annotations_on_protein(
    target_protein=current_protein,
    control_package=pass_dict
)
```

```
peptide_uniprot_map[
    (peptide_uniprot_map['Protein']==current_protein) & # Current Protein
    (peptide_uniprot_map['PeptideID'].isin([1, 2, 8])) # Peptide to inspect
]
```

|  | Protein | PeptideID | pepStart | pepEnd | feature\_category | feature | startpos | endpos | group | agent | note | score | dPF |
| --- | --- | --- | --- | --- | --- | --- | --- | --- | --- | --- | --- | --- | --- |
| 112187 | Q04760 | 1 | 29 | 38 | Protein Processing & Maturation | CLEAVAGE | 38 | 38 | Proteolytic Cleavage | trypsin 1 | N-Ph cleavage at R38 by trypsin 1 | 2.0000 | 1 |
| 112188 | Q04760 | 1 | 29 | 38 | Co- & Post-Translational Modifications | MOD\_RES | 35 | 35 | Phosphorylation |  | Phosphorylation at T35 | 1.0000 | 1 |
| 112189 | Q04760 | 1 | 29 | 38 | Co- & Post-Translational Modifications | MOD\_RES | 38 | 38 | Methylation |  | Methylation at R38 | 1.0000 | 1 |
| 112190 | Q04760 | 1 | 29 | 38 | Sequence Heterogeneity & Isoforms | MUTAGEN | 34 | 34 | Reduced Function | Q->E | Reduces enzyme activity by 99%. | 3.0000 | 1 |
| 112191 | Q04760 | 2 | 44 | 51 | Protein Processing & Maturation | CLEAVAGE | 51 | 51 | Proteolytic Cleavage | trypsin 1 | N-Ph cleavage at R51 by trypsin 1 | 2.0000 | 1 |
| 112192 | Q04760 | 2 | 44 | 51 | Co- & Post-Translational Modifications | CROSSLNK | 44 | 44 | Ubiquitination/SUMOylation |  | Ubiquitination/SUMOylation at K44 | 1.0000 | 1 |
| 112193 | Q04760 | 2 | 44 | 51 | Co- & Post-Translational Modifications | MOD\_RES | 45 | 45 | Phosphorylation |  | Phosphorylation at S45 | 1.0000 | 1 |
| 112194 | Q04760 | 2 | 44 | 51 | Sequence Heterogeneity & Isoforms | MUTAGEN | 45 | 45 | No Significant Effect | S->A | No effect on phosphorylation. | 3.0000 | 1 |
| 112236 | Q04760 | 8 | 141 | 148 | Protein Processing & Maturation | CLEAVAGE | 144 | 144 | Proteolytic Cleavage | glutamyl endopeptidase I | N-Ph cleavage at E144 by glutamyl endopeptidase I | 2.0000 | 1 |
| 112237 | Q04760 | 8 | 141 | 148 | Co- & Post-Translational Modifications | CROSSLNK | 148 | 148 | Ubiquitination/SUMOylation |  | Ubiquitination/SUMOylation at K148 | 1.0000 | 1 |
| 112238 | Q04760 | 8 | 141 | 148 | Co- & Post-Translational Modifications | MOD\_RES | 148 | 148 | Acetylation |  | N6-acetyllysine; alternate | 3.0000 | 1 |
| 112239 | Q04760 | 8 | 141 | 148 | Co- & Post-Translational Modifications | MOD\_RES | 148 | 148 | Acylation |  | N6-succinyllysine; alternate | 3.0000 | 1 |
| 112240 | Q04760 | 8 | 141 | 148 | Co- & Post-Translational Modifications | MOD\_RES | 148 | 148 | Acetylation |  | Acetylation at K148 | 2.0000 | 1 |
| 112241 | Q04760 | 8 | 141 | 148 | Co- & Post-Translational Modifications | MOD\_RES | 148 | 148 | Acetylation |  | Acetylation at K148 | 2.0000 | 1 |

###### 05.1.2 HAGH/GLO2 (Q16775)¶

```
current_protein = 'Q16775'
cur_gene = protein_dpf_analysis.loc[
    protein_dpf_analysis['Protein'] == current_protein, 'Gene'
].values[0]
print(f"Current Protein: {current_protein} ({cur_gene})")
plots.detailed_peptide_with_clusters(
    data=test_data,
    cur_protein=current_protein,
    pThr=pThr,
    pvalue_col='adj_pval',
    cluster_col='ClusterID',
    rawIntensity_col='log10Intensity',
    adjIntensity_col='AdjIntensity',
    condition_palette=condition_colors,
    
    save = not is_demo,
    show=True,
    filename=f'detailed_peptide_look_{cur_gene}_72hr',
    filepath=figure_path,
    fileformats=figure_formats,
    transparent=transparent_bg,
    dpi=figure_dpi
)
```

```
Current Protein: Q16775 (HAGH)
```

```
plots.annotations_on_protein(
    target_protein=current_protein,
    control_package=pass_dict
)
```

```
peptide_uniprot_map[
    (peptide_uniprot_map['Protein']==current_protein) & # Current Protein
    (peptide_uniprot_map['PeptideID'].isin([4, 5])) # Peptide to inspect
]
```

|  | Protein | PeptideID | pepStart | pepEnd | feature\_category | feature | startpos | endpos | group | agent | note | score | dPF |
| --- | --- | --- | --- | --- | --- | --- | --- | --- | --- | --- | --- | --- | --- |
| 143380 | Q16775 | 4 | 233 | 242 | Sequence Heterogeneity & Isoforms | VAR\_SEQ | 146 | 236 | Modified | 3 | Modified (in isoform 3) | 3.0000 | 1 |
| 143381 | Q16775 | 4 | 233 | 242 | Sequence Heterogeneity & Isoforms | VAR\_SEQ | 237 | 308 | Missing | 3 | Missing (in isoform 3) | 3.0000 | 1 |
| 143382 | Q16775 | 5 | 252 | 274 | Sequence Heterogeneity & Isoforms | VAR\_SEQ | 237 | 308 | Missing | 3 | Missing (in isoform 3) | 3.0000 | 1 |

##### 05.2 Other Notable/Interesting Proteins¶

###### 05.2.1 SARG (Q9BW04)¶

```
current_protein = 'Q9BW04'
cur_gene = protein_dpf_analysis.loc[
    protein_dpf_analysis['Protein'] == current_protein, 'Gene'
].values[0]
print(f"Current Protein: {current_protein} ({cur_gene})")
plots.detailed_peptide_with_clusters(
    data=test_data,
    cur_protein=current_protein,
    pThr=pThr,
    pvalue_col='adj_pval',
    cluster_col='ClusterID',
    rawIntensity_col='log10Intensity',
    adjIntensity_col='AdjIntensity',
    condition_palette=condition_colors,
    
    save = not is_demo,
    show=True,
    filename=f'detailed_peptide_look_{cur_gene}_72hr',
    filepath=figure_path,
    fileformats=figure_formats,
    transparent=transparent_bg,
    dpi=figure_dpi
)
```

```
Current Protein: Q9BW04 (SARG)
```

```
plots.annotations_on_protein(
    target_protein=current_protein,
    control_package=pass_dict
)
```

```
peptide_uniprot_map[
    (peptide_uniprot_map['Protein']==current_protein) & # Current Protein
    (peptide_uniprot_map['PeptideID'].isin([11, 12])) # Peptide to inspect
]
```

|  | Protein | PeptideID | pepStart | pepEnd | feature\_category | feature | startpos | endpos | group | agent | note | score | dPF |
| --- | --- | --- | --- | --- | --- | --- | --- | --- | --- | --- | --- | --- | --- |
| 207481 | Q9BW04 | 11 | 300 | 308 | Co- & Post-Translational Modifications | MOD\_RES | 308 | 308 | Methylation |  | Methylation at K308 | 1.0000 | 1 |
| 207482 | Q9BW04 | 12 | 354 | 369 |  | No Annotation | -1 | -1 |  |  |  | 0.0000 | 1 |

###### 05.2.2 FPGT (O14772)¶

```
current_protein = 'O14772'
cur_gene = protein_dpf_analysis.loc[
    protein_dpf_analysis['Protein'] == current_protein, 'Gene'
].values[0]
print(f"Current Protein: {current_protein} ({cur_gene})")
plots.detailed_peptide_with_clusters(
    data=test_data,
    cur_protein=current_protein,
    pThr=pThr,
    pvalue_col='adj_pval',
    cluster_col='ClusterID',
    rawIntensity_col='log10Intensity',
    adjIntensity_col='AdjIntensity',
    condition_palette=condition_colors,
    
    save = not is_demo,
    show=True,
    filename=f'detailed_peptide_look_{cur_gene}_72hr',
    filepath=figure_path,
    fileformats=figure_formats,
    transparent=transparent_bg,
    dpi=figure_dpi
)
```

```
Current Protein: O14772 (FPGT)
```

```
plots.annotations_on_protein(
    target_protein=current_protein,
    control_package=pass_dict
)
```

```
peptide_uniprot_map[
    (peptide_uniprot_map['Protein']==current_protein) & # Current Protein
    (peptide_uniprot_map['PeptideID'].isin([1,2])) # Peptide to inspect
]
```

|  | Protein | PeptideID | pepStart | pepEnd | feature\_category | feature | startpos | endpos | group | agent | note | score | dPF |
| --- | --- | --- | --- | --- | --- | --- | --- | --- | --- | --- | --- | --- | --- |
| 6169 | O14772 | 1 | 62 | 73 | Co- & Post-Translational Modifications | CROSSLNK | 73 | 73 | Ubiquitination/SUMOylation |  | Ubiquitination/SUMOylation at K73 | 1.0000 | 1 |
| 6170 | O14772 | 2 | 78 | 95 |  | No Annotation | -1 | -1 |  |  |  | 0.0000 | 1 |

```
peptide_uniprot_map[
    (peptide_uniprot_map['Protein']==current_protein) & # Current Protein
    (peptide_uniprot_map['feature']=='VAR_SEQ') # Where the Variant Sequence annotations are
]
```

|  | Protein | PeptideID | pepStart | pepEnd | feature\_category | feature | startpos | endpos | group | agent | note | score | dPF |
| --- | --- | --- | --- | --- | --- | --- | --- | --- | --- | --- | --- | --- | --- |
| 6171 | O14772 | 3 | 296 | 303 | Sequence Heterogeneity & Isoforms | VAR\_SEQ | 128 | 607 | Modified | 6 | Modified (in isoform 6) | 3.0000 | 0 |
| 6172 | O14772 | 3 | 296 | 303 | Sequence Heterogeneity & Isoforms | VAR\_SEQ | 150 | 403 | Missing | 5 | Missing (in isoform 5) | 3.0000 | 0 |
| 6174 | O14772 | 4 | 332 | 347 | Sequence Heterogeneity & Isoforms | VAR\_SEQ | 128 | 607 | Modified | 6 | Modified (in isoform 6) | 3.0000 | 0 |
| 6175 | O14772 | 4 | 332 | 347 | Sequence Heterogeneity & Isoforms | VAR\_SEQ | 150 | 403 | Missing | 5 | Missing (in isoform 5) | 3.0000 | 0 |
| 6176 | O14772 | 5 | 356 | 368 | Sequence Heterogeneity & Isoforms | VAR\_SEQ | 128 | 607 | Modified | 6 | Modified (in isoform 6) | 3.0000 | 0 |
| 6177 | O14772 | 5 | 356 | 368 | Sequence Heterogeneity & Isoforms | VAR\_SEQ | 150 | 403 | Missing | 5 | Missing (in isoform 5) | 3.0000 | 0 |

---

```
other_interesting_proteins = [
    ### Add other proteins of interest here ###
    'Q08345', # DDR1 -> 72hr  last 3 peptide dPF lost normoxia (missing in isoform 3)
    'Q5W111', # SPRYD7 -> 72hr  first 2 peptide no normoxia (perhaps an isoform)
    

    
]

is_demo = True # False if want to save figures

pass_dict = {
    'data': {
        'uniprot': uniprot_data,
        'detailed': test_data,
        'summary': summary_data,
    },
    'styles': {
        'dpf_colors': dPF_colors,
    },
    'config': {
        'is_demo': is_demo,
        'figure_path': figure_path,
        'figure_formats': figure_formats,
        'transparent_bg': transparent_bg,
        'figure_dpi': figure_dpi,
        'str_add': '72hr',
    }
}
```

```
utils.view_table(
    protein_dpf_analysis[
        (protein_dpf_analysis['Protein'].isin(candidate_proteins)) & 
        (protein_dpf_analysis['dPF_pct'] <= 50)
    ].sort_values('dPF_pct', ascending=False),
    page_size=10,
    page_number=11
)
```

```
Total pages: 16, Current page: 11
```

|  | Protein | Gene | Length | Coverage | all\_pep | sgnf\_pep | all\_clust | n\_forms | n\_canon | n\_ptm | n\_dPF | sgnf\_pct | canon\_pct | ptm\_pct | dPF\_pct | eff\_pct | cov\_cat | sgnf\_cat | len\_cat | dpf\_cat |
| --- | --- | --- | --- | --- | --- | --- | --- | --- | --- | --- | --- | --- | --- | --- | --- | --- | --- | --- | --- | --- |
| 2646 | Q0JRZ9 | FCHO2 | 810 | 21.4815 | 12 | 3 | 3 | 1 | 9 | 1 | 2 | 25.0000 | 75.0000 | 8.3333 | 16.6667 | 33.3333 | Low | Multiple | Long | Both |
| 491 | O60502 | OGA | 916 | 17.4672 | 12 | 2 | 2 | 1 | 10 | 0 | 2 | 16.6667 | 83.3333 | 0.0000 | 16.6667 | 50.0000 | Low | Multiple | Long | dPF Only |
| 1476 | P23469 | PTPRE | 700 | 21.7143 | 13 | 2 | 2 | 1 | 11 | 0 | 2 | 15.3846 | 84.6154 | 0.0000 | 15.3846 | 50.0000 | Low | Multiple | Medium | dPF Only |
| 5424 | Q99986 | VRK1 | 396 | 38.1313 | 13 | 1 | 2 | 1 | 11 | 0 | 2 | 7.6923 | 84.6154 | 0.0000 | 15.3846 | 50.0000 | Medium | Single | Medium | dPF Only |
| 6913 | Q9Y2R4 | DDX52 | 599 | 34.0568 | 13 | 2 | 2 | 1 | 11 | 0 | 2 | 15.3846 | 84.6154 | 0.0000 | 15.3846 | 50.0000 | Medium | Multiple | Medium | dPF Only |
| 6883 | Q9Y2E5 | MAN2B2 | 1009 | 18.7314 | 13 | 2 | 2 | 1 | 11 | 0 | 2 | 15.3846 | 84.6154 | 0.0000 | 15.3846 | 50.0000 | Low | Multiple | V.Long | dPF Only |
| 4836 | Q92766 | RREB1 | 1687 | 9.9585 | 13 | 2 | 2 | 1 | 11 | 0 | 2 | 15.3846 | 84.6154 | 0.0000 | 15.3846 | 50.0000 | Low | Multiple | V.Long | dPF Only |
| 4360 | Q8N556 | AFAP1 | 730 | 20.5479 | 13 | 2 | 2 | 1 | 11 | 0 | 2 | 15.3846 | 84.6154 | 0.0000 | 15.3846 | 50.0000 | Low | Multiple | Medium | dPF Only |
| 5828 | Q9H223 | EHD4 | 541 | 42.5139 | 20 | 4 | 3 | 1 | 16 | 1 | 3 | 20.0000 | 80.0000 | 5.0000 | 15.0000 | 33.3333 | Medium | Multiple | Medium | Both |
| 1816 | P42680 | TEC | 631 | 26.1490 | 14 | 2 | 2 | 1 | 12 | 0 | 2 | 14.2857 | 85.7143 | 0.0000 | 14.2857 | 50.0000 | Medium | Multiple | Medium | dPF Only |

```
current_protein = 'Q9H223'
cur_gene = protein_dpf_analysis.loc[
    protein_dpf_analysis['Protein'] == current_protein, 'Gene'
].values[0]
print(f"Current Protein: {current_protein} ({cur_gene})")
plots.detailed_peptide_with_clusters(
    data=test_data,
    cur_protein=current_protein,
    pThr=pThr,
    pvalue_col='adj_pval',
    cluster_col='ClusterID',
    rawIntensity_col='log10Intensity',
    adjIntensity_col='AdjIntensity',
    condition_palette=condition_colors,
    
    save = not is_demo,
    show=True,
    filename=f'detailed_peptide_look_{cur_gene}_72hr',
    filepath=figure_path,
    fileformats=figure_formats,
    transparent=transparent_bg,
    dpi=figure_dpi
)
```

```
Current Protein: Q9H223 (EHD4)
```

```
plots.annotations_on_protein(
    target_protein=current_protein,
    control_package=pass_dict
)
```

```
peptide_uniprot_map[
    (peptide_uniprot_map['Protein']==current_protein) & # Current Protein
    (peptide_uniprot_map['PeptideID'].isin([5, 11, 12, 13])) # Peptide to inspect
]
```

|  | Protein | PeptideID | pepStart | pepEnd | feature\_category | feature | startpos | endpos | group | agent | note | score | dPF |
| --- | --- | --- | --- | --- | --- | --- | --- | --- | --- | --- | --- | --- | --- |
| 212668 | Q9H223 | 5 | 188 | 196 | Cleavage Analysis | MISCLEAVAGE | 188 | 196 | Miscleavage | ProteoForge\_Analysis | Overlap with PeptideID 6 (fraction: 1.0) | 1.0000 | -1 |
| 212677 | Q9H223 | 11 | 273 | 283 |  | No Annotation | -1 | -1 |  |  |  | 0.0000 | 1 |
| 212678 | Q9H223 | 12 | 309 | 319 | Co- & Post-Translational Modifications | CROSSLNK | 319 | 319 | Ubiquitination/SUMOylation |  | Ubiquitination/SUMOylation at K319 | 1.0000 | 1 |
| 212679 | Q9H223 | 12 | 309 | 319 | Co- & Post-Translational Modifications | MOD\_RES | 312 | 312 | Phosphorylation |  | Phosphorylation at Y312 | 1.0000 | 1 |
| 212680 | Q9H223 | 12 | 309 | 319 | Co- & Post-Translational Modifications | MOD\_RES | 316 | 316 | Phosphorylation |  | Phosphorylation at Y316 | 1.0000 | 1 |
| 212681 | Q9H223 | 12 | 309 | 319 | Co- & Post-Translational Modifications | MOD\_RES | 319 | 319 | Methylation |  | Methylation at K319 | 1.0000 | 1 |
| 212682 | Q9H223 | 13 | 333 | 347 | Cleavage Analysis | MISCLEAVAGE | 338 | 347 | Partial\_Overlap | ProteoForge\_Analysis | Overlap with PeptideID 14 (fraction: 0.667) | 0.6667 | 1 |

```
peptide_uniprot_map[
    (peptide_uniprot_map['Protein']==current_protein) & # Current Protein
    # (peptide_uniprot_map['feature']=='VAR_SEQ') # Where the Variant Sequence annotations are
    (peptide_uniprot_map['feature'].isin(['VAR_SEQ','CLEAVAGE'])) # Where the Variant Sequence annotations are
]
```

|  | Protein | PeptideID | pepStart | pepEnd | feature\_category | feature | startpos | endpos | group | agent | note | score | dPF |
| --- | --- | --- | --- | --- | --- | --- | --- | --- | --- | --- | --- | --- | --- |
| 212675 | Q9H223 | 10 | 246 | 254 | Protein Processing & Maturation | CLEAVAGE | 254 | 254 | Proteolytic Cleavage | trypsin 1 | N-Ph cleavage at R254 by trypsin 1 | 2.0000 | 0 |
| 212684 | Q9H223 | 15 | 348 | 360 | Protein Processing & Maturation | CLEAVAGE | 360 | 360 | Proteolytic Cleavage | trypsin 1 | N-Ph cleavage at K360 by trypsin 1 | 2.0000 | 0 |

```
print("Notebook Execution Time:", utils.prettyTimer(utils.getTime() - startTime))
```

```
Notebook Execution Time: 00h:15m:17s
```
