## Supplementary material for "ProteoForge: An Imputation-Aware Framework for Differential Proteoform Discovery in Bottom-Up Proteomics": Analysis notebooks as html render: Notebook S14.html

### Notebook S14

###### Enes Kemal Ergin

#### 2025-12-08 23:00:25

### Supplementary Notebook 14: Protein-Level Downstream Analysis Comparison¶

- **License:** Creative Commons Attribution-NonCommercial 4.0 International License
- **Version:** 0.2
- **Edit Log:**
  - 2025-11-28: Initial version of the notebook
  - 2025-12-08: Revise the whole notebook, ensuring clarity and correctness

---

**Requirements:**

This notebook requires the `02-ApplyingProteoForge_48hr.ipynb`, `02-ApplyingProteoForge_72hr.ipynb` to be run first to generate the result files used here. Ensure the `./data/cleaned/` and `.data/results/` directories contains the necessary input files.

---

**Data Information:**

- **Cell Line:** H358 (NSCLC - Non-Small Cell Lung Cancer)
- **Conditions:** True Hypoxia (1% O₂) vs Normoxia (21% O₂)
- **Timepoints:** 48 hours and 72 hours
- **Quantification Methods:** 4 approaches (Original, mean(top), mean(all), dPFs)
  - Original is from DIANN protein output, the other three are calculated from peptide-level data.

---

**Purpose:**

This notebook compares the protein-level quantification results from various methods, the original DIANN output of the protein-level quant version, the averaging top3 peptides, averaging all peptides, and the proteoform-informed quantification using dPFs. The goal is to assess how proteoform-aware quantification influences various metrics that we care about our protein-level data, such as technical reproducibility, differential expression detection, and biological relevance (e.g. enrichment of hypoxia-related pathways).

---

#### 01. Setup¶

This section imports required libraries, configures display settings, and defines paths for data and figures.

> **Note:** The HTML rendering of this notebook hides code cells by default. Click the "Code" buttons on the right to expand them.

##### 01.1 Libraries¶

```
# Libraries Used
import os
import sys
import warnings

import numpy as np # Numerical computing
import pandas as pd # Data manipulation

import seaborn as sns # R-like high-level plots
import matplotlib.pyplot as plt # Python's base plotting 

from sklearn.decomposition import PCA

# home path
sys.path.append('../')
from src import utils, plots 
from src import tests, enrich

from ProteoForge import parse, process
from ProteoForge import impute, normalize

warnings.filterwarnings('ignore')
# Initialize the timer
startTime = utils.getTime()
```

##### 01.2 Configure Notebook Settings¶

Configure visualization styles, color palettes, and display options for consistent output formatting throughout the analysis.

```
## Configure Plotting Settings
def_colors = [
    "#139593", "#fca311", "#e54f2a",
    "#c3c3c3", "#555555",
    "#690000", "#5f4a00", "#004549"
]

condition_colors = {
    'True Hypoxia (1% O2) - 48 Hr': '#e9c46a',
    'True Hypoxia (1% O2) - 72 Hr': '#f4a261',
    'Normoxia (21% O2) - 48 Hr': '#ade8f4',
    'Normoxia (21% O2) - 72 Hr': '#0077b6',
    # Won't be used since the manuscript didn't cover these conditions
    # 'Chemical Hypoxia (CoCl2) - 48 Hr': '#c77dff',
    # 'Chemical Hypoxia (CoCl2) - 72 Hr': '#8338ec',
    # 'Oxidative Stress (H2O2) - 48 Hr': '#ff006e',
    # 'Oxidative Stress (H2O2) - 72 Hr': '#fb5607',
}

method_order = ['Original', 'mean(top)', 'mean(all)',  'dPFs']

proteinMethods = {
    "Original": "#f4f1de",
    "mean(top)": "#f2cc8f",
    "mean(all)": "#81b29a",
    "dPFs": "#183a37"
}

cv_group_palettes = {
    "<10%":  "#081c15",
    "<25%":  "#1b4332",
    "<50%": "#2d6a4f",
    "<100%": "#52b788",
    ">100%": "#95d5b2",
}

# --- Define Order and Colors ---
status_colors = {
    'Unexplained': "#C2C0C0",
    'Excluded': '#565d61',
    'Upregulated': '#780000',
    'Downregulated': '#e36414',
    'Equivalent': '#003049',
}

count_type_order = list(status_colors.keys())

status_markers = {
    "Equivalent": "o",
    "Upregulated": "^",
    "Downregulated": "v",
    "Unexplained": "s",
    "Excluded": "x"
}

cond_colors = {
    'Hypoxia (1% O2)': '#f4a261',
    'Normoxia (21% O2)': '#0077b6',
}
dura_colors = {
    '48hr': '#c3c3c3',
    '72hr': '#555555',
}
cond_order = ['Normoxia (21% O2)', 'Hypoxia (1% O2)']
dura_order = ['48hr', '72hr']

# Set seaborn style
sns.set_theme(
    style="white",
    context="paper",
    palette=def_colors,
    font_scale=1,
    rc={
        "figure.figsize": (6, 4),
        "font.family": "sans-serif",
        "font.sans-serif": ["Arial", "Ubuntu Mono"],
    }
)

# Figure Saving Settings
figure_formats = ["png", "pdf"]
save_to_folder = True
transparent_bg = True
figure_dpi = 300

## Configure dataframe displaying
pd.set_option('display.float_format', lambda x: '%.4f' % x)
pd.set_option('display.max_rows', 50)
pd.set_option('display.max_columns', 50)
pd.set_option('display.width', 500)  # Set a wider display width


## Printing Settings
verbose = True

plots.color_palette( def_colors, save=False )
plots.color_palette( condition_colors, save=False, name='condition', size=2)
plots.color_palette( proteinMethods, save=False, name='protein level calculation method', size=2)
```


##### 01.3 Data and Result Paths¶

- `home_path` — Base project path (e.g., `./`)
- `data_name` — Dataset identifier used throughout (`"hypoxia"`)
- `notebook_name` — This notebook name for logging and organizing outputs (`"04-Downstream"`)
- `data_path` — Root data folder (`{home_path}data/`)
- `raw_path` — Raw input folder (`{data_path}input/{data_name}/`)
- `input_path` — Cleaned input data from preprocessing (`{data_path}cleaned/{data_name}/`)
- `output_path` — ProteoForge result files (`{data_path}results/{data_name}/`)
- `figure_path` — Destination for generated figures (`{home_path}figures/{data_name}/{notebook_name}/`)

```
home_path = './'
data_name = "hypoxia"
notebook_name = "04-Downstream"
data_path = f"{home_path}data/"
raw_path = f"{data_path}input/{data_name}/"
input_path = f"{data_path}cleaned/{data_name}/"
output_path = f"{data_path}results/{data_name}/"
figure_path = f"{home_path}figures/{data_name}/{notebook_name}/"

# Create the output folder if it does not exist
if not os.path.exists(output_path):
    os.makedirs(output_path)

# Create figure folder structure, if needed
if save_to_folder:
    for i in figure_formats:
        cur_folder = figure_path + i + "/"
        if not os.path.exists(cur_folder):
            os.makedirs(cur_folder)
```

##### 01.4 Load Input Data¶

Load the imputed peptide-level data, protein-level quantification results from different methods, and the FASTA database for annotation mapping.

**Imputed Peptide Data:** Contains peptide intensities across samples after imputation. This will be the main data for three of the quantification methods.

```
peptide_df = pd.read_feather(f"{input_path}imputed_data.feather").reset_index()
proteins_used = peptide_df['Protein'].unique()
```

**Fasta Table:** Contains protein sequences and annotations for mapping peptides to proteins and functional information. Reading again for protein-level analyses.

```
fasta_data = parse.fasta_to_df(
    fasta_path = f'./data/input/20230316_HumanCr_20806.fasta',
    gene_only = False,
    column_order=['entry', 'geneName', 'proteinDescription', 'sequenceLength', 'molecularWeight_kDa', 'sequence']
).rename(
    columns={
        'entry': 'Protein',
        'geneName': 'Gene',
        'proteinDescription': 'Description',
        'sequenceLength': 'Length',
        'molecularWeight_kDa': 'Weight(kDa)',
        'sequence': 'Sequence'
    }
)
fasta_data.head()
```

```
Processing FASTA file: ./data/input/20230316_HumanCr_20806.fasta
Auto-selected: multiprocessing with 4 processes
Loading FASTA entries...
Processing 26855 entries in chunks of 2828
Processed 26772 entries, skipped 83
  - invalid_sequence: 78
  - length_filter: 5
Generated DataFrame with 26772 rows and 6 columns
```

|  | Protein | Gene | Description | Length | Weight(kDa) | Sequence |
| --- | --- | --- | --- | --- | --- | --- |
| 0 | A0A024B7W1 | None | Genome polyprotein | 3423 | 378.8692 | MKNPKKKSGGFRIVNMLKRGVARVSPFGGLKRLPAGLLLGHGPIRM... |
| 1 | A0A024R1R8 | TMA7B | Translation machinery-associated protein 7B | 64 | 7.0870 | MSSHEGGKKKALKQPKKQAKEMDEEEKAFKQKQKEEQKKLEVLKAK... |
| 2 | A0A024RBG1 | NUDT4B | Diphosphoinositol polyphosphate phosphohydrola... | 181 | 20.4213 | MMKFKPNQTRTYDREGFKKRAACLCFRSEQEDEVLLVSSSRYPDQW... |
| 3 | A0A075B6H7 | IGKV3-7 | Probable non-functional immunoglobulin kappa v... | 116 | 12.7754 | MEAPAQLLFLLLLWLPDTTREIVMTQSPPTLSLSPGERVTLSCRAS... |
| 4 | A0A075B6H8 | IGKV1D-42 | Probable non-functional immunoglobulin kappa v... | 117 | 13.0056 | MDMRVPAQLLGLLLLWLPGVRFDIQMTQSPSFLSASVGDRVSIICW... |

**Original Protein Quant Data:** DIANN output with protein-level quantification.

```
original_protein = pd.read_csv(f"{raw_path}report.pg_matrix.tsv", sep="\t", engine='pyarrow')
meta = original_protein.columns[4:].to_list()
# Build a meta dataframe
meta = pd.DataFrame(
    data=meta,
    columns=['filename']
)
# --- Clean and Standardize Metadata ---
print("\n🧹 Cleaning and standardizing metadata columns...")

meta['Run'] = meta['filename'].str.split(r'\\').str[-1].str.replace('.d', '')
# Extract condition identifier
def get_condition(x):
    if '1%' in x:
        return 'True Hypoxia (1% O2)'
    elif 'CoCl' in x:
        return 'Chemical Hypoxia (CoCl2)'
    elif 'H2O2' in x:
        return 'Oxidative Stress (H2O2)'
    else:
        return 'Normoxia (21% O2)'
meta['Condition'] = meta['Run'].map(get_condition)
# Extract the replicate number (special rule for 1% O2)
meta['Replicate'] = meta['Run'].str.split('_').apply(lambda x: x[2] if '1%' in x[1] else x[1][-1])
meta['Duration'] = meta['Run'].str.split('_').str[-5] # last 5th element from the end
meta['Colname'] = meta['Run'].str.split('_').map(
    # Simplify the colname based on the condition
    lambda x: f"{x[1]}_{x[3]}_{x[2]}" if '1%' in x[1] else f"{x[1][:-1]}_{x[2]}_{x[1][-1]}"
)
# --- Finalize DataFrame ---
print("Selecting final columns and renaming for clarity...")
print("\n✅ Metadata processing complete.")
print(f"Final DataFrame shape: {meta.shape}")
print(f"Final columns: {meta.columns.tolist()}")
print()

# Sort the metadata DataFrame for better readability
meta = meta.sort_values(
    ['Condition', 'Duration', 'Replicate'], 
    ascending=[False, True, True]
).reset_index(drop=True)

# Keep only the relevant conditions for the study
relevant_conditions = [
    'True Hypoxia (1% O2)',
    'Normoxia (21% O2)',
]
print(f"Filtering metadata to keep only relevant conditions: {relevant_conditions}")
meta = meta[meta['Condition'].isin(relevant_conditions)].reset_index(drop=True)
print(f"Filtered DataFrame shape: {meta.shape}")
meta
```

```
🧹 Cleaning and standardizing metadata columns...
Selecting final columns and renaming for clarity...

✅ Metadata processing complete.
Final DataFrame shape: (32, 6)
Final columns: ['filename', 'Run', 'Condition', 'Replicate', 'Duration', 'Colname']

Filtering metadata to keep only relevant conditions: ['True Hypoxia (1% O2)', 'Normoxia (21% O2)']
Filtered DataFrame shape: (16, 6)
```

|  | filename | Run | Condition | Replicate | Duration | Colname |
| --- | --- | --- | --- | --- | --- | --- |
| 0 | D:\Data\Tam\RevisionLungDIA\H358\H358\_1%\_1\_48\_... | H358\_1%\_1\_48\_DIA\_BB4\_1\_5432 | True Hypoxia (1% O2) | 1 | 48 | 1%\_48\_1 |
| 1 | D:\Data\Tam\RevisionLungDIA\H358\H358\_1%\_2\_48\_... | H358\_1%\_2\_48\_DIA\_BC4\_1\_5440 | True Hypoxia (1% O2) | 2 | 48 | 1%\_48\_2 |
| 2 | D:\Data\Tam\RevisionLungDIA\H358\H358\_1%\_3\_48\_... | H358\_1%\_3\_48\_DIA\_BD4\_1\_5448 | True Hypoxia (1% O2) | 3 | 48 | 1%\_48\_3 |
| 3 | D:\Data\Tam\RevisionLungDIA\H358\H358\_1%\_4\_48\_... | H358\_1%\_4\_48\_DIA\_BE4\_1\_5456 | True Hypoxia (1% O2) | 4 | 48 | 1%\_48\_4 |
| 4 | D:\Data\Tam\RevisionLungDIA\H358\H358\_1%\_1\_72\_... | H358\_1%\_1\_72\_DIA\_BB8\_1\_5436 | True Hypoxia (1% O2) | 1 | 72 | 1%\_72\_1 |
| 5 | D:\Data\Tam\RevisionLungDIA\H358\H358\_1%\_2\_72\_... | H358\_1%\_2\_72\_DIA\_BC8\_1\_5444 | True Hypoxia (1% O2) | 2 | 72 | 1%\_72\_2 |
| 6 | D:\Data\Tam\RevisionLungDIA\H358\H358\_1%\_3\_72\_... | H358\_1%\_3\_72\_DIA\_BD8\_1\_5452 | True Hypoxia (1% O2) | 3 | 72 | 1%\_72\_3 |
| 7 | D:\Data\Tam\RevisionLungDIA\H358\H358\_1%\_4\_72\_... | H358\_1%\_4\_72\_DIA\_BE8\_1\_5460 | True Hypoxia (1% O2) | 4 | 72 | 1%\_72\_4 |
| 8 | D:\Data\Tam\RevisionLungDIA\H358\H358\_C1\_48\_DI... | H358\_C1\_48\_DIA\_BB1\_1\_5429 | Normoxia (21% O2) | 1 | 48 | C\_48\_1 |
| 9 | D:\Data\Tam\RevisionLungDIA\H358\H358\_C2\_48\_DI... | H358\_C2\_48\_DIA\_BC1\_1\_5437 | Normoxia (21% O2) | 2 | 48 | C\_48\_2 |
| 10 | D:\Data\Tam\RevisionLungDIA\H358\H358\_C3\_48\_DI... | H358\_C3\_48\_DIA\_BD1\_1\_5445 | Normoxia (21% O2) | 3 | 48 | C\_48\_3 |
| 11 | D:\Data\Tam\RevisionLungDIA\H358\H358\_C4\_48\_DI... | H358\_C4\_48\_DIA\_BE1\_1\_5453 | Normoxia (21% O2) | 4 | 48 | C\_48\_4 |
| 12 | D:\Data\Tam\RevisionLungDIA\H358\H358\_C1\_72\_DI... | H358\_C1\_72\_DIA\_BB5\_1\_5433 | Normoxia (21% O2) | 1 | 72 | C\_72\_1 |
| 13 | D:\Data\Tam\RevisionLungDIA\H358\H358\_C2\_72\_DI... | H358\_C2\_72\_DIA\_BC5\_1\_5441 | Normoxia (21% O2) | 2 | 72 | C\_72\_2 |
| 14 | D:\Data\Tam\RevisionLungDIA\H358\H358\_C3\_72\_DI... | H358\_C3\_72\_DIA\_BD5\_1\_5449 | Normoxia (21% O2) | 3 | 72 | C\_72\_3 |
| 15 | D:\Data\Tam\RevisionLungDIA\H358\H358\_C4\_72\_DI... | H358\_C4\_72\_DIA\_BE5\_1\_5457 | Normoxia (21% O2) | 4 | 72 | C\_72\_4 |

```
cv_datas = []
# Build a dictionary to match condition_colors keys: 'Condition - Duration Hr'
condition_to_samples = {}
for condition in meta['Condition'].unique():
    for duration in sorted(meta.loc[meta['Condition'] == condition, 'Duration'].unique()):
        key = f"{condition} - {duration} Hr"
        samples = meta.loc[(meta['Condition'] == condition) & (meta['Duration'] == duration), 'Colname'].tolist()
        condition_to_samples[key] = samples

# samples_to_condition mapping
samples_to_condition = {sample: condition for condition, samples in condition_to_samples.items() for sample in samples}
# samples_to_colors mapping
samples_to_colors = {sample: condition_colors[condition] for sample, condition in samples_to_condition.items()}
```

#### 02. Protein Quantification Methods Comparison¶

This section loads and processes protein-level intensities using four different quantification strategies. For each method, we calculate coefficient of variation (CV) to assess reproducibility and perform PCA to visualize sample separation.

##### 02.1 Original DIANN Protein Intensities¶

Load protein intensities as reported by DIA-NN software. This represents the default protein-level quantification without additional processing. The key representing this method will be `Original`.

> Re-doing some of the data cleanups from first notebook for consistency for protein-level version for consistency.

###### Loading the Data¶

```
original_protein = pd.read_csv(f"{raw_path}report.pg_matrix.tsv", sep="\t", engine='pyarrow')
original_protein = original_protein.drop(
    columns=['Protein.Names', 'Genes', 'First.Protein.Description']
).rename(
    columns={"Protein.Group": "Protein"}
)
# Display summary information
print(f"Data shape (wide format): {original_protein.shape}")


print("\n--- 🧬 Protein ID Representation ---")
# Check if the protein contains '-' indicating it has isoforms
has_isoform = original_protein['Protein'].str.contains('-').any()
has_multiples = original_protein['Protein'].str.contains(';').any()
# Friendly print statements to explain the protein ID representation
if has_isoform:
    print("The dataset contains protein IDs with isoforms (e.g., 'P12345-2'). These represent specific variants of a protein.")
else:
    print("The dataset does not contain protein IDs with isoforms.")

if has_multiples:
    print("The dataset contains protein IDs with multiple entries (e.g., 'P12345; P67890'). These represent proteins that may have multiple forms or related proteins.")
    print()
    print("Will explode these entries to create a more detailed DataFrame.")
    original_protein = process.explode_aligned(original_protein, ['Protein'], sep=';', verbose=verbose)
    print()
    print(f"After exploding, the dataset now has {len(original_protein)} rows and {len(original_protein.columns)} columns.")
else:
    print("The dataset does not contain protein IDs with multiple entries.")

print(f"Values below 2 should be set as NaN, they might be artifacts of quantification.")
print(f"Number of missing values in the DataFrame before cleanup: {original_protein.isnull().sum().sum()}")
# Identify numeric columns in the DataFrame
numeric_columns = original_protein.select_dtypes(include=[np.number]).columns
original_protein[numeric_columns] = original_protein[numeric_columns].mask(
    original_protein[numeric_columns] < 2
)
print(f"Number of missing values in the DataFrame after cleanup: {original_protein.isnull().sum().sum()}")

original_protein = original_protein.set_index('Protein')[meta['filename']]
print("Set cleaned column names from metadata")
original_protein.columns = meta['Colname']
original_protein = original_protein.sort_index()
original_protein.head()
```

```
Data shape (wide format): (8918, 33)

--- 🧬 Protein ID Representation ---
The dataset does not contain protein IDs with isoforms.
The dataset contains protein IDs with multiple entries (e.g., 'P12345; P67890'). These represent proteins that may have multiple forms or related proteins.

Will explode these entries to create a more detailed DataFrame.
🚀 Starting aligned explosion for columns: ['Protein']
Original DataFrame shape: (8918, 33)
Final DataFrame shape: (9029, 33)
✅ Transformation complete in 0.0905 seconds.

After exploding, the dataset now has 9029 rows and 33 columns.
Values below 2 should be set as NaN, they might be artifacts of quantification.
Number of missing values in the DataFrame before cleanup: 12877
Number of missing values in the DataFrame after cleanup: 12877
Set cleaned column names from metadata
```

| Colname | 1%\_48\_1 | 1%\_48\_2 | 1%\_48\_3 | 1%\_48\_4 | 1%\_72\_1 | 1%\_72\_2 | 1%\_72\_3 | 1%\_72\_4 | C\_48\_1 | C\_48\_2 | C\_48\_3 | C\_48\_4 | C\_72\_1 | C\_72\_2 | C\_72\_3 | C\_72\_4 |
| --- | --- | --- | --- | --- | --- | --- | --- | --- | --- | --- | --- | --- | --- | --- | --- | --- |
| Protein |  |  |  |  |  |  |  |  |  |  |  |  |  |  |  |  |
| A0A024RBG1 | 106932.0000 | 114285.0000 | 101081.0000 | 120276.0000 | 124488.0000 | 109649.0000 | 104434.0000 | 92969.7000 | 125823.0000 | 106866.0000 | 115710.0000 | 137117.0000 | 110514.0000 | 138896.0000 | 124320.0000 | 104223.0000 |
| A0A096LP01 | 9701.4100 | 6065.4900 | 8416.2600 | 10267.9000 | NaN | 8381.3700 | 8866.8700 | 8336.3700 | 11194.8000 | 14924.0000 | 14774.3000 | 12868.4000 | 12249.5000 | 13873.9000 | 13091.8000 | 16304.4000 |
| A0A0B4J2D5 | 239693.0000 | 223978.0000 | 224786.0000 | 254631.0000 | 174872.0000 | 262457.0000 | 267884.0000 | 242496.0000 | 241334.0000 | 177731.0000 | 210887.0000 | 269207.0000 | 233719.0000 | 223538.0000 | 204650.0000 | 169070.0000 |
| A0A0B4J2F0 | 6286.4600 | NaN | 20654.8000 | 22783.6000 | 22998.9000 | 22887.6000 | 26960.6000 | 19887.2000 | NaN | 10762.5000 | 19256.9000 | 23658.6000 | 12704.9000 | 11545.9000 | 20380.3000 | 23297.6000 |
| A0A2Z4LIS9 | 4035.0200 | 2587.3300 | 2064.8100 | 2414.1500 | 5768.8800 | 2343.8100 | 2189.9700 | 2148.6600 | 1615.3900 | 3101.2300 | 2665.9200 | 3850.5500 | 2414.3300 | 3994.3500 | 2486.6600 | 2599.2100 |

###### 02.1.1 Data Cleaning and Normalization¶

Clean the protein intensity data through:

- Outlier sample detection using combined metric scoring (correlation, quantification ratio, CV)
- Removal of systematically aberrant samples
- Median centering normalization
- Missing value imputation using a multi-step pipeline

> The same cleaning and normalization steps as applied to peptide-level data are applied here to ensure consistency across quantification methods.

```
metric_weights = {  # Weights for the metrics used in pairwise combined score calculation
    'correlation': 1.0,
    'quantification': 1.0,
    'cv': 1.0
}
minOutlier_occurrance = 3 # minimum number of outlier occurrences to consider a sample behaves like an outlier

conditions_to_check = list(condition_to_samples.keys())  # List of conditions to check for outlier detection

print("\nCalculating Pairwise Combined Scores...")
print(" - Using metric weights:", metric_weights)
print(" - Using Spearman correlation, quantification as ratio, and CV as 1 - CV")

all_outlier_samples, combined_group_reports = utils.identify_outlier_samples(
    data=original_protein,
    sample_groups=condition_to_samples,
    analysis_func=utils.run_metric_combination_analysis,
    min_outlier_occurrence=minOutlier_occurrance,
    verbose=False
)
print(f"\nComplete list of unique outlier samples to remove (after min_outlier_occurrence >= {minOutlier_occurrance})\n - {all_outlier_samples}")
print()

plot_data = original_protein.reset_index().melt(
    id_vars='Protein',
    var_name='Sample',
    value_name='Intensity'
).dropna().assign(log2_Intensity=lambda x: np.log2(x['Intensity']))
fig, ax = plt.subplots(figsize=(12, 4))
sns.boxplot(
    x='Sample', y='log2_Intensity', hue='Sample', data=plot_data, ax=ax, 
    palette=samples_to_colors,
    fliersize=0.5,
    linewidth=0.5,
    notch=False,
    showmeans=True,
    meanprops={"marker": "o", "markerfacecolor": "white", "markeredgecolor": "black"},
    medianprops={"color": "white", "linewidth": 2.5},
    whiskerprops={"color": "black", "linewidth": 1},
    capprops={"color": "black", "linewidth": 0},
    legend=False
)
y_min = 0
y_max = plot_data['log2_Intensity'].max() + 1
ax.set_ylim(y_min, y_max)
ax.tick_params(axis='x', labelrotation=90)
ax.set_title('Boxplot of Log2 Intensities Across Samples', fontsize=16, fontweight='bold')
ax.set_xlabel('Sample', fontsize=14)
ax.set_ylabel('Log2(Intensity)', fontsize=14)
ax.grid(which='both', linestyle='--', linewidth=0.5, color='lightgray', alpha=0.7)

import matplotlib.patches as mpatches

# Highlight background for outlier samples
for i, label in enumerate(ax.get_xticklabels()):
    sample_label = label.get_text()
    if sample_label in all_outlier_samples:
        # Add a rectangle spanning the y-range at the x position
        ax.add_patch(
            mpatches.Rectangle(
                (i - 0.5, y_min),  # x position, y position
                1,                 # width (one boxplot)
                y_max - y_min,     # height
                color='#ffcccc',   # light red
                zorder=0,          # behind the boxplot
                alpha=0.5
            )
        )
# Add legend for outlier highlight
outlier_patch = mpatches.Patch(color='#ffcccc', label='Identified Outlier Sample', alpha=0.5)
ax.legend(handles=[outlier_patch], loc='upper right', fontsize=12)
plt.tight_layout()
plt.show()
# Remove the identified outlier samples from the main data
print("\nRemoving identified outlier samples from the main data...")
print(f"- Samples to remove: {all_outlier_samples}")
print(f"- Data shape before removal: {original_protein.shape}")
cleaned_data = original_protein.drop(columns=all_outlier_samples)
print(f"- Data shape after removal: {cleaned_data.shape}")
print("Outlier removal complete.")
```

```
Calculating Pairwise Combined Scores...
 - Using metric weights: {'correlation': 1.0, 'quantification': 1.0, 'cv': 1.0}
 - Using Spearman correlation, quantification as ratio, and CV as 1 - CV

Complete list of unique outlier samples to remove (after min_outlier_occurrence >= 3)
 - ['C_72_4', '1%_72_1']
```

```
Removing identified outlier samples from the main data...
- Samples to remove: ['C_72_4', '1%_72_1']
- Data shape before removal: (9029, 16)
- Data shape after removal: (9029, 14)
Outlier removal complete.
```

```
meta = meta[~meta['Colname'].isin(all_outlier_samples)].reset_index(drop=True)
# Build a dictionary to match condition_colors keys: 'Condition - Duration Hr'
condition_to_samples = {}
for condition in meta['Condition'].unique():
    for duration in sorted(meta.loc[meta['Condition'] == condition, 'Duration'].unique()):
        key = f"{condition} - {duration} Hr"
        samples = meta.loc[(meta['Condition'] == condition) & (meta['Duration'] == duration), 'Colname'].tolist()
        condition_to_samples[key] = samples

# samples_to_condition mapping
samples_to_condition = {sample: condition for condition, samples in condition_to_samples.items() for sample in samples}
# samples_to_colors mapping
samples_to_colors = {sample: condition_colors[condition] for sample, condition in samples_to_condition.items()}
```

```
centered_data = normalize.by_median_centering(
    df=cleaned_data,
    rescale_to_original_magnitude=True,
    condition_map=None, # Pass dictionary if want to apply condition-specific centering
    verbose=verbose
)

imputation_pipeline_scheme = [
    { # First is the remove sparsely quantified features to set complete missing for low-value imputation 
        'method': 'amputate',
        'params': { 'min_quantified': 1 }
    },
    # # I generally don't use this step, but it can be useful for some datasets
    # { # Second step is generally to fill the sparse missingness with features' mean or median 
    #     'method': 'fill_dense', 
    #     'params': {'max_missing': 1, 'strategy': 'mean'}
    # },
    # This is the best for complete or mostly missing features
    { # Adds downshifted low-values to completely missing (1.0) or mostly missing (< 1.0) features
        'method': 'downshift',
        'params': {'missingness_threshold': .5, 'shift_magnitude': 2.5, 'low_percentile': 0.05}
    },
    { # Final step is to use k-NN imputation to fill the remaining missing values
        'method': 'knn',
        'params': {'n_neighbors': 4}
    }
]

imputed_data = impute.run_imputation_pipeline(
    data=centered_data,
    cond_dict=condition_to_samples,
    scheme=imputation_pipeline_scheme,
    is_log2=False,  
    return_log2=False,
    verbose=verbose
)
original_protein = imputed_data
```

```
Applying Median Centering...
  - Applying global median centering...
    - Rescaling by a global factor of: 32768.00
  - Data after median centering has shape: (9029, 14)
  - Missing values (Before/After): (3319/3319)
====== Starting Imputation Pipeline ======
Initial missing values: 3319

--- Running Step: Sparse Feature Amputation ---
  Parameters: min_quantified=1
  Summary: Created 0 new missing values by amputation.
Total missing values after step 1 ('amputate'): 3319

--- Running Step: Downshifted Imputation ---
  Parameters: missingness_threshold=0.5, shift_magnitude=2.5, low_percentile=0.05
  Condition 'True Hypoxia (1% O2) - 48 Hr': Imputing 257 features with >= 2 missing values.
  Condition 'True Hypoxia (1% O2) - 72 Hr': Imputing 227 features with >= 2 missing values.
  Condition 'Normoxia (21% O2) - 48 Hr': Imputing 243 features with >= 2 missing values.
  Condition 'Normoxia (21% O2) - 72 Hr': Imputing 184 features with >= 2 missing values.
  Summary: Imputed 2464 missing values.
Total missing values after step 2 ('downshift'): 855

--- Running Step: k-NN Imputation ---
  Parameters: n_neighbors=4
  Condition 'True Hypoxia (1% O2) - 48 Hr': Applying k-NN to 211 remaining missing values.
  Condition 'True Hypoxia (1% O2) - 72 Hr': Applying k-NN to 210 remaining missing values.
  Condition 'Normoxia (21% O2) - 48 Hr': Applying k-NN to 179 remaining missing values.
  Condition 'Normoxia (21% O2) - 72 Hr': Applying k-NN to 255 remaining missing values.
  Summary: Imputed 855 missing values.
Total missing values after step 3 ('knn'): 0

====== Imputation Pipeline Finished ======
```

---

###### 02.1.2 Coefficient of Variation (CV) Analysis¶

The CVs will be calculated across conditions at this method and will be stored in a `cv_datas` table to create comparison plots later on.

```
print("\nCalculating Coefficient of Variation (CV) from proteins and group by condition...")

cv_data = pd.DataFrame(index = original_protein.index)
# Calculate the coefficient of variation per biological sample
for k, v in condition_to_samples.items():
    if not all(item in original_protein.columns for item in v):
        continue
    cv_data[k] = utils.cv_numpy(
        original_protein[v].values,
        axis=1,
        ignore_nan=True,
        format="percent"
    )

plot_data = utils.create_cv_group_plot_data(
    cv_data=cv_data,
    cv_group_palettes=cv_group_palettes
).dropna(subset=['CV'])
plot_data['Method'] = 'Original'
cv_datas.append(plot_data)
```

```
Calculating Coefficient of Variation (CV) from proteins and group by condition...
```

###### 02.1.3 PCA Visualization¶

Perform PCA on the cleaned protein intensities to visualize sample clustering and separation between hypoxia and normoxia conditions.

```
# PCA
scaler = "zscore"
complete_data = original_protein
scaled_data = utils.scale_the_data(
    complete_data,
    method=scaler,
    axis=1,
    is_log=False
).fillna(0)

# Run PCA
pca = PCA()
pca_res = pca.fit_transform(scaled_data.T)
pca_exp_var = pca.explained_variance_ratio_

# Set Variables
C1, C2 = 0, 1
hue_group = "Condition"
style_group = "Duration"
comp_locs = [C1, C2]
comp_names = [f'PC{c+1}' for c in comp_locs]
xlabel = f'{comp_names[0]} ({pca_exp_var[comp_locs[0]]:.2%} of variance)'
ylabel = f'{comp_names[1]} ({pca_exp_var[comp_locs[1]]:.2%} of variance)'

# Build the PCA Plot Data
plot_data = pd.DataFrame(pca_res[:, comp_locs], columns=comp_names)
plot_data['UniqueNames'] = complete_data.columns
plot_data['Condition'] = plot_data['UniqueNames'].map(samples_to_condition)
plot_data['Duration'] = plot_data['UniqueNames'].str.split('_').str[1]

# Find Centroids for each group (Condition and Duration)
centroid_data = plot_data.groupby([hue_group, style_group])[comp_names].mean().reset_index()

# Prepare color/marker mapping for unique (hue, style) combinations
unique_groups = centroid_data[[hue_group, style_group]].drop_duplicates()
group_palette = {}
group_marker = {}
markers = ["o", "s", "D", "^", "v", "P", "X", "*"]
for idx, (cond, dura) in enumerate(unique_groups.values):
    color = condition_colors.get(cond, "#888888")
    marker = markers[idx % len(markers)]
    group_palette[(cond, dura)] = color
    group_marker[(cond, dura)] = marker

fig, ax = plt.subplots(figsize=(9, 7))
fig.patch.set_facecolor('white')

# Plot technical replicates
for (cond, dura), group_df in plot_data.groupby([hue_group, style_group]):
    color = group_palette.get((cond, dura), "#888888")
    marker = group_marker.get((cond, dura), "o")
    ax.scatter(
        group_df[comp_names[0]], group_df[comp_names[1]],
        label=f"{cond}, {dura}hr",
        color=color, marker=marker, s=80, edgecolor='k', alpha=0.85, linewidth=1.2, zorder=2
    )
    # Confidence ellipse
    if len(group_df) > 1:
        plots.confidence_ellipse(
            group_df[comp_names[0]], group_df[comp_names[1]], ax, n_std=2,
            edgecolor=color, facecolor=color, alpha=0.18, linewidth=1.5, zorder=1
        )

# Plot centroids
for (cond, dura), group_df in centroid_data.groupby([hue_group, style_group]):
    color = group_palette.get((cond, dura), "#888888")
    marker = group_marker.get((cond, dura), "o")
    ax.scatter(
        group_df[comp_names[0]], group_df[comp_names[1]],
        color=color, marker=marker, s=260, edgecolor='k', linewidth=2.5, alpha=1, zorder=3
    )

# Axis limits and grid
minVal = plot_data[comp_names].min().min()
maxVal = plot_data[comp_names].max().max()
limit = max(abs(minVal), abs(maxVal))
offset = 0.1 * limit
limit += offset
ax.set_xlim(-limit, limit)
ax.set_ylim(-limit, limit)
ax.grid(True, which="both", linestyle="--", linewidth=0.7, alpha=0.4, color="grey")

# 0 lines
ax.axvline(0, color='grey', linestyle='--', linewidth=1.2, alpha=0.6, zorder=0)
ax.axhline(0, color='grey', linestyle='--', linewidth=1.2, alpha=0.6, zorder=0)

# Labels and title
ax.set_xlabel(xlabel, fontsize=14, fontweight="bold")
ax.set_ylabel(ylabel, fontsize=14, fontweight="bold")
ax.set_title(
    f"PCA of Original Protein Intensities (n={complete_data.shape[0]})",
    fontsize=17, fontweight="bold", pad=15, loc="left"
)

# Custom legend: combine color and marker
from matplotlib.lines import Line2D
legend_elements = [
    Line2D(
        [0], [0],
        marker=group_marker[(cond, dura)],
        color='w',
        markerfacecolor=group_palette[(cond, dura)],
        markeredgecolor='k',
        markersize=13,
        linewidth=0,
        label=f"{cond}, {dura}hr"
    )
    for (cond, dura) in unique_groups.values
]
ax.legend(
    handles=legend_elements,
    title="Condition, Duration",
    bbox_to_anchor=(1.02, 1),
    loc="upper left",
    frameon=False,
    fontsize=11,
    title_fontsize=12,
    borderaxespad=0.5
)

sns.despine(ax=ax, left=True, bottom=True)
plt.tight_layout()
plots.finalize_plot(
    fig, show=True, save=save_to_folder, 
    filename='PCA_Original_Proteins',
    filepath=figure_path,
    formats=figure_formats,
    transparent=transparent_bg,
    dpi=figure_dpi
)
```

The PCA plot shows the hypoxia and normoxia samples clustering distinctly, indicating that the original DIANN protein quantification captures biological differences effectively. However there are major overlaps between timepoints showing that timepoint differences are not well captured especially for normoxia samples.

The Explained variance is captured for the first two principal components for later comparison against other methods.

---

##### 02.2 Mean of Top-3 Peptides per Protein¶

Calculate protein intensities by averaging the three most abundant peptides per protein. This approach reduces the influence of low-abundance or poorly quantified peptides.

###### 02.2.1 Calculating Protein Intensities from Top-3 Peptides¶

```
peptide_df = pd.read_feather(f"{input_path}imputed_data.feather").reset_index()

# Ensure the first two columns are 'Protein' and 'unique_id', rest are samples
sample_cols = peptide_df.columns[2:]
# Calculate mean intensity for each peptide across all samples
peptide_df['mean_intensity'] = peptide_df[sample_cols].mean(axis=1, skipna=True)
# Sort by Protein and mean_intensity descending
peptide_df_sorted = peptide_df.sort_values(['Protein', 'mean_intensity'], ascending=[True, False])

# Select top 3 peptides per protein (vectorized)
top3_peptides = peptide_df_sorted.groupby('Protein', group_keys=False).head(3)

# Group by Protein and take the mean of sample columns (vectorized)
protein_top3 = top3_peptides.groupby('Protein')[sample_cols].mean()

print(f'Protein-level intensities (Top3 mean) shape: {protein_top3.shape}')

print(f"\n--- 🔄 Median Centering Normalization ---")
protein_top3 = normalize.by_median_centering(
    df=protein_top3,
    rescale_to_original_magnitude=True,
    condition_map=None,
    verbose=verbose
)

cv_data = pd.DataFrame(index = protein_top3.index)
# Calculate the coefficient of variation per biological sample
for k, v in condition_to_samples.items():
    # if not all(item in protein_top3.columns for item in v):
    #     continue
    cv_data[k] = utils.cv_numpy(
        protein_top3[v].values,
        axis=1,
        ignore_nan=True,
        format="percent"
    )

plot_data = utils.create_cv_group_plot_data(
    cv_data=cv_data,
    cv_group_palettes=cv_group_palettes
).dropna(subset=['CV'])
plot_data['Method'] = 'mean(top)'
cv_datas.append(plot_data)

protein_top3.head()
```

```
Protein-level intensities (Top3 mean) shape: (7161, 14)

--- 🔄 Median Centering Normalization ---

Applying Median Centering...
  - Applying global median centering...
    - Rescaling by a global factor of: 32768.00
  - Data after median centering has shape: (7161, 14)
  - Missing values (Before/After): (0/0)
```

| Colname | 1%\_48\_1 | 1%\_48\_2 | 1%\_48\_3 | 1%\_48\_4 | 1%\_72\_2 | 1%\_72\_3 | 1%\_72\_4 | C\_48\_1 | C\_48\_2 | C\_48\_3 | C\_48\_4 | C\_72\_1 | C\_72\_2 | C\_72\_3 |
| --- | --- | --- | --- | --- | --- | --- | --- | --- | --- | --- | --- | --- | --- | --- |
| Protein |  |  |  |  |  |  |  |  |  |  |  |  |  |  |
| A0A024RBG1 | 42434.0877 | 44675.9925 | 39905.7199 | 45166.4723 | 41072.9743 | 41509.6032 | 37214.0846 | 47803.0084 | 42063.5809 | 40992.1971 | 48181.0230 | 43600.5230 | 53923.3764 | 48616.6057 |
| A0A0B4J2D5 | 102861.7619 | 109717.8950 | 96666.4961 | 104414.6932 | 113987.2439 | 108450.8661 | 105768.6036 | 114020.7478 | 80482.5059 | 88493.0615 | 109336.0082 | 100739.3204 | 98188.1377 | 84013.4979 |
| A0A1B0GTU1 | 15418.3217 | 13016.2592 | 20110.4711 | 16653.0423 | 10068.9798 | 10815.8631 | 7411.4047 | 18067.9728 | 15565.7734 | 14406.1895 | 15852.1724 | 16550.5883 | 21497.2074 | 21683.5016 |
| A0A1W2PQL4 | 138883.8369 | 157641.7370 | 92956.1284 | 144272.3660 | 181450.5185 | 186647.1011 | 164358.8932 | 148590.3989 | 165077.2871 | 170214.6665 | 166396.9636 | 141739.9102 | 158388.9094 | 174016.5392 |
| A0A2R8Y4L2 | 2516111.9537 | 2101575.8857 | 2084923.5383 | 1978299.3752 | 2458649.4794 | 2321907.3483 | 1817408.9804 | 2162122.0254 | 2094417.0202 | 2240939.2056 | 2186368.9643 | 2425103.1440 | 1965437.4835 | 1998541.0596 |

###### 02.2.2 PCA Visualization¶

Perform PCA on the cleaned protein intensities to visualize sample clustering and separation between hypoxia and normoxia conditions.

```
# PCA
scaler = "zscore"
complete_data = protein_top3.fillna(0)
scaled_data = utils.scale_the_data(
    complete_data,
    method=scaler,
    axis=1,
    is_log=False
)

# Run PCA
pca = PCA()
pca_res = pca.fit_transform(scaled_data.T)
pca_exp_var = pca.explained_variance_ratio_

# Set Variables
C1, C2 = 0, 1
hue_group = "Condition"
style_group = "Duration"
comp_locs = [C1, C2]
comp_names = [f'PC{c+1}' for c in comp_locs]
xlabel = f'{comp_names[0]} ({pca_exp_var[comp_locs[0]]:.2%} of variance)'
ylabel = f'{comp_names[1]} ({pca_exp_var[comp_locs[1]]:.2%} of variance)'

# Build the PCA Plot Data
plot_data = pd.DataFrame(pca_res[:, comp_locs], columns=comp_names)
plot_data['UniqueNames'] = complete_data.columns
plot_data['Condition'] = plot_data['UniqueNames'].map(samples_to_condition)
plot_data['Duration'] = plot_data['UniqueNames'].str.split('_').str[1]

# Find Centroids for each group (Condition and Duration)
centroid_data = plot_data.groupby([hue_group, style_group])[comp_names].mean().reset_index()

# Prepare color/marker mapping for unique (hue, style) combinations
unique_groups = centroid_data[[hue_group, style_group]].drop_duplicates()
group_palette = {}
group_marker = {}
markers = ["o", "s", "D", "^", "v", "P", "X", "*"]
for idx, (cond, dura) in enumerate(unique_groups.values):
    color = condition_colors.get(cond, "#888888")
    marker = markers[idx % len(markers)]
    group_palette[(cond, dura)] = color
    group_marker[(cond, dura)] = marker

fig, ax = plt.subplots(figsize=(9, 7))
fig.patch.set_facecolor('white')

# Plot technical replicates
for (cond, dura), group_df in plot_data.groupby([hue_group, style_group]):
    color = group_palette.get((cond, dura), "#888888")
    marker = group_marker.get((cond, dura), "o")
    ax.scatter(
        group_df[comp_names[0]], group_df[comp_names[1]],
        label=f"{cond}, {dura}hr",
        color=color, marker=marker, s=80, edgecolor='k', alpha=0.85, linewidth=1.2, zorder=2
    )
    # Confidence ellipse
    if len(group_df) > 1:
        plots.confidence_ellipse(
            group_df[comp_names[0]], group_df[comp_names[1]], ax, n_std=2,
            edgecolor=color, facecolor=color, alpha=0.18, linewidth=1.5, zorder=1
        )

# Plot centroids
for (cond, dura), group_df in centroid_data.groupby([hue_group, style_group]):
    color = group_palette.get((cond, dura), "#888888")
    marker = group_marker.get((cond, dura), "o")
    ax.scatter(
        group_df[comp_names[0]], group_df[comp_names[1]],
        color=color, marker=marker, s=260, edgecolor='k', linewidth=2.5, alpha=1, zorder=3
    )

# Axis limits and grid
minVal = plot_data[comp_names].min().min()
maxVal = plot_data[comp_names].max().max()
limit = max(abs(minVal), abs(maxVal))
offset = 0.1 * limit
limit += offset
ax.set_xlim(-limit, limit)
ax.set_ylim(-limit, limit)
ax.grid(True, which="both", linestyle="--", linewidth=0.7, alpha=0.4, color="grey")

# 0 lines
ax.axvline(0, color='grey', linestyle='--', linewidth=1.2, alpha=0.6, zorder=0)
ax.axhline(0, color='grey', linestyle='--', linewidth=1.2, alpha=0.6, zorder=0)

# Labels and title
ax.set_xlabel(xlabel, fontsize=14, fontweight="bold")
ax.set_ylabel(ylabel, fontsize=14, fontweight="bold")
ax.set_title(
    f"PCA of Averaging Top Peptides to Protein Intensities (n={complete_data.shape[0]})",
    fontsize=17, fontweight="bold", pad=15, loc="left"
)

# Custom legend: combine color and marker
from matplotlib.lines import Line2D
legend_elements = [
    Line2D(
        [0], [0],
        marker=group_marker[(cond, dura)],
        color='w',
        markerfacecolor=group_palette[(cond, dura)],
        markeredgecolor='k',
        markersize=13,
        linewidth=0,
        label=f"{cond}, {dura}hr"
    )
    for (cond, dura) in unique_groups.values
]
ax.legend(
    handles=legend_elements,
    title="Condition, Duration",
    bbox_to_anchor=(1.02, 1),
    loc="upper left",
    frameon=False,
    fontsize=11,
    title_fontsize=12,
    borderaxespad=0.5
)

sns.despine(ax=ax, left=True, bottom=True)
plt.tight_layout()
plots.finalize_plot(
    fig, show=True, save=save_to_folder, 
    filename='PCA_meanTop_Proteins',
    filepath=figure_path,
    formats=figure_formats,
    transparent=transparent_bg,
    dpi=figure_dpi
)
```

Similar separation of hypoxia and normoxia samples is observed, however the separation between normoxia timepoints is still not well captured.

---

##### 02.3 Mean of All Peptides per Protein¶

Calculate protein intensities by averaging all detected peptides per protein. This approach incorporates all available peptide information but may be influenced by outlier peptides.

###### 02.3.1 Calculating Protein Intensities from All Peptides¶

```
peptide_df = pd.read_feather(f"{input_path}imputed_data.feather").reset_index()
# Ensure the first two columns are 'Protein' and 'unique_id', rest are samples
sample_cols = peptide_df.columns[2:]

# Average all peptides per protein
protein_all = peptide_df.groupby('Protein')[sample_cols].mean()

print(f'Protein-level intensities (All peptides mean) shape: {protein_all.shape}')

print(f"\n--- 🔄 Median Centering Normalization ---")
protein_all = normalize.by_median_centering(
    df=protein_all,
    rescale_to_original_magnitude=True,
    condition_map=None,
    verbose=verbose
)

cv_data = pd.DataFrame(index = protein_all.index)
# Calculate the coefficient of variation per biological sample
for k, v in condition_to_samples.items():
    # if not all(item in protein_all.columns for item in v):
    #     continue
    cv_data[k] = utils.cv_numpy(
        protein_all[v].values,
        axis=1,
        ignore_nan=True,
        format="percent"
    )

plot_data = utils.create_cv_group_plot_data(
    cv_data=cv_data,
    cv_group_palettes=cv_group_palettes
).dropna(subset=['CV'])
plot_data['Method'] = 'mean(all)'
cv_datas.append(plot_data)


protein_all.head()
```

```
Protein-level intensities (All peptides mean) shape: (7161, 14)

--- 🔄 Median Centering Normalization ---

Applying Median Centering...
  - Applying global median centering...
    - Rescaling by a global factor of: 16384.00
  - Data after median centering has shape: (7161, 14)
  - Missing values (Before/After): (0/0)
```

| Colname | 1%\_48\_1 | 1%\_48\_2 | 1%\_48\_3 | 1%\_48\_4 | 1%\_72\_2 | 1%\_72\_3 | 1%\_72\_4 | C\_48\_1 | C\_48\_2 | C\_48\_3 | C\_48\_4 | C\_72\_1 | C\_72\_2 | C\_72\_3 |
| --- | --- | --- | --- | --- | --- | --- | --- | --- | --- | --- | --- | --- | --- | --- |
| Protein |  |  |  |  |  |  |  |  |  |  |  |  |  |  |
| A0A024RBG1 | 27636.4812 | 30005.0847 | 27084.4570 | 30501.0198 | 26878.6847 | 27282.2816 | 24778.7294 | 31250.4761 | 28260.9575 | 27105.6051 | 31660.7097 | 28399.4406 | 36102.3521 | 33361.4951 |
| A0A0B4J2D5 | 57134.1133 | 58310.0884 | 55598.9816 | 62191.6960 | 64318.6873 | 65554.3341 | 61014.3493 | 60071.2849 | 46073.6487 | 49443.3042 | 60765.3547 | 59501.3649 | 55451.3679 | 51310.4708 |
| A0A1B0GTU1 | 9527.9206 | 8306.3244 | 12492.2302 | 10603.8432 | 6954.6379 | 7748.1887 | 5544.2338 | 12078.9927 | 10559.6725 | 9870.6067 | 10691.6913 | 10105.9986 | 13391.0161 | 13435.8479 |
| A0A1W2PQL4 | 87048.5451 | 101193.6058 | 63425.9557 | 93900.9660 | 111078.8289 | 116289.6754 | 102781.4867 | 93745.7319 | 106650.8837 | 107614.7891 | 104448.3260 | 90009.2451 | 104011.9006 | 113290.7150 |
| A0A2R8Y4L2 | 1178694.3393 | 1016651.1508 | 1031318.9612 | 955571.0935 | 1131586.4682 | 1103637.8918 | 934521.9371 | 1030494.1624 | 1001852.8436 | 1087337.1202 | 997756.8049 | 1181303.7617 | 1012222.6601 | 982521.7678 |

###### 02.3.2 PCA Visualization¶

Perform PCA on the cleaned protein intensities to visualize sample clustering and separation between hypoxia and normoxia conditions.

```
# PCA
scaler = "zscore"
complete_data = protein_all.fillna(0)
scaled_data = utils.scale_the_data(
    complete_data,
    method=scaler,
    axis=1,
    is_log=False
)

# Run PCA
pca = PCA()
pca_res = pca.fit_transform(scaled_data.T)
pca_exp_var = pca.explained_variance_ratio_

# Set Variables
C1, C2 = 0, 1
hue_group = "Condition"
style_group = "Duration"
comp_locs = [C1, C2]
comp_names = [f'PC{c+1}' for c in comp_locs]
xlabel = f'{comp_names[0]} ({pca_exp_var[comp_locs[0]]:.2%} of variance)'
ylabel = f'{comp_names[1]} ({pca_exp_var[comp_locs[1]]:.2%} of variance)'

# Build the PCA Plot Data
plot_data = pd.DataFrame(pca_res[:, comp_locs], columns=comp_names)
plot_data['UniqueNames'] = complete_data.columns
plot_data['Condition'] = plot_data['UniqueNames'].map(samples_to_condition)
plot_data['Duration'] = plot_data['UniqueNames'].str.split('_').str[1]

# Find Centroids for each group (Condition and Duration)
centroid_data = plot_data.groupby([hue_group, style_group])[comp_names].mean().reset_index()

# Prepare color/marker mapping for unique (hue, style) combinations
unique_groups = centroid_data[[hue_group, style_group]].drop_duplicates()
group_palette = {}
group_marker = {}
markers = ["o", "s", "D", "^", "v", "P", "X", "*"]
for idx, (cond, dura) in enumerate(unique_groups.values):
    color = condition_colors.get(cond, "#888888")
    marker = markers[idx % len(markers)]
    group_palette[(cond, dura)] = color
    group_marker[(cond, dura)] = marker

fig, ax = plt.subplots(figsize=(9, 7))
fig.patch.set_facecolor('white')

# Plot technical replicates
for (cond, dura), group_df in plot_data.groupby([hue_group, style_group]):
    color = group_palette.get((cond, dura), "#888888")
    marker = group_marker.get((cond, dura), "o")
    ax.scatter(
        group_df[comp_names[0]], group_df[comp_names[1]],
        label=f"{cond}, {dura}hr",
        color=color, marker=marker, s=80, edgecolor='k', alpha=0.85, linewidth=1.2, zorder=2
    )
    # Confidence ellipse
    if len(group_df) > 1:
        plots.confidence_ellipse(
            group_df[comp_names[0]], group_df[comp_names[1]], ax, n_std=2,
            edgecolor=color, facecolor=color, alpha=0.18, linewidth=1.5, zorder=1
        )

# Plot centroids
for (cond, dura), group_df in centroid_data.groupby([hue_group, style_group]):
    color = group_palette.get((cond, dura), "#888888")
    marker = group_marker.get((cond, dura), "o")
    ax.scatter(
        group_df[comp_names[0]], group_df[comp_names[1]],
        color=color, marker=marker, s=260, edgecolor='k', linewidth=2.5, alpha=1, zorder=3
    )

# Axis limits and grid
minVal = plot_data[comp_names].min().min()
maxVal = plot_data[comp_names].max().max()
limit = max(abs(minVal), abs(maxVal))
offset = 0.1 * limit
limit += offset
ax.set_xlim(-limit, limit)
ax.set_ylim(-limit, limit)
ax.grid(True, which="both", linestyle="--", linewidth=0.7, alpha=0.4, color="grey")

# 0 lines
ax.axvline(0, color='grey', linestyle='--', linewidth=1.2, alpha=0.6, zorder=0)
ax.axhline(0, color='grey', linestyle='--', linewidth=1.2, alpha=0.6, zorder=0)

# Labels and title
ax.set_xlabel(xlabel, fontsize=14, fontweight="bold")
ax.set_ylabel(ylabel, fontsize=14, fontweight="bold")
ax.set_title(
    f"PCA of Averaging All Peptides to Protein Intensities (n={complete_data.shape[0]})",
    fontsize=17, fontweight="bold", pad=15, loc="left"
)

# Custom legend: combine color and marker
from matplotlib.lines import Line2D
legend_elements = [
    Line2D(
        [0], [0],
        marker=group_marker[(cond, dura)],
        color='w',
        markerfacecolor=group_palette[(cond, dura)],
        markeredgecolor='k',
        markersize=13,
        linewidth=0,
        label=f"{cond}, {dura}hr"
    )
    for (cond, dura) in unique_groups.values
]
ax.legend(
    handles=legend_elements,
    title="Condition, Duration",
    bbox_to_anchor=(1.02, 1),
    loc="upper left",
    frameon=False,
    fontsize=11,
    title_fontsize=12,
    borderaxespad=0.5
)

sns.despine(ax=ax, left=True, bottom=True)
plt.tight_layout()
plots.finalize_plot(
    fig, show=True, save=save_to_folder, 
    filename='PCA_meanAll_Proteins',
    filepath=figure_path,
    formats=figure_formats,
    transparent=transparent_bg,
    dpi=figure_dpi
)
```

Similar trends are observed as with the top-3 peptide method, with hypoxia and normoxia samples separating but timepoint differences remaining less distinct.

---

##### 02.4 ProteoForge Differential Proteoforms (dPFs)¶

Build proteoform-level intensities using ProteoForge's differential proteoform (dPF) assignments. Each protein may have multiple dPFs representing distinct regulatory states identified through peptide correlation clustering.

**Note:** PTM-specific peptides (dPF=-1) are excluded as they represent post-translational modifications rather than quantifiable proteoform abundance.

###### 02.4.1 Calculating Protein Intensities from dPFs - 48hr¶

> Note: The Individual 48hr and 72hr only ProteoForge dPF quantifications used to better capture timepoint-specific proteoform changes. Since using the Comb version of the analysis would make the comparison unfair as it would have more data to cluster peptides together from both timepoints.

```
dPF_long = pd.read_feather(f"{output_path}test_data_48hr.feather")

# Remove the PTM peptides (dPF=-1) since they don't contribute to the protein quantification
dPF_long = dPF_long[dPF_long['dPF'] != -1].reset_index(drop=True)
dPF_long['Proteoform'] = dPF_long['Protein'] + "_" + dPF_long['dPF'].astype(str)
protein_dpf_48 = dPF_long.pivot_table(
    index='Proteoform',
    columns='Sample',
    values='Intensity',
    aggfunc='mean'  # In case of duplicates, take the mean
)

print(f'Protein-level intensities (Proteoform dPF) shape: {protein_dpf_48.shape}')
print(f"\n--- 🔄 Median Centering Normalization ---")
protein_dpf_48 = normalize.by_median_centering(
    df=protein_dpf_48,
    rescale_to_original_magnitude=True,
    # condition_map={k: v for k, v in condition_to_samples.items() if '48 Hr' in k}, 
    condition_map=None,
    verbose=verbose
)


cv_data = pd.DataFrame(index = protein_dpf_48.index)
# Calculate the coefficient of variation per biological sample
for k, v in condition_to_samples.items():
    if not all(item in protein_dpf_48.columns for item in v):
        continue
    cv_data[k] = utils.cv_numpy(
        protein_dpf_48[v].values,
        axis=1,
        ignore_nan=True,
        format="percent"
    )

plot_data = utils.create_cv_group_plot_data(
    cv_data=cv_data,
    cv_group_palettes=cv_group_palettes,
    id_col='Proteoform'
).dropna(subset=['CV'])
plot_data['Method'] = 'dPFs'
cv_datas.append(plot_data)
```

```
Protein-level intensities (Proteoform dPF) shape: (7857, 8)

--- 🔄 Median Centering Normalization ---

Applying Median Centering...
  - Applying global median centering...
    - Rescaling by a global factor of: 16384.00
  - Data after median centering has shape: (7857, 8)
  - Missing values (Before/After): (0/0)
```

###### 02.4.2 Calculating Protein Intensities from dPFs - 72hr¶

Load and process 72-hour timepoint dPF data using the same approach as 48hr.

```
dPF_long = pd.read_feather(f"{output_path}test_data_72hr.feather")

# Remove the PTM peptides (dPF=-1) since they don't contribute to the protein quantification
dPF_long = dPF_long[dPF_long['dPF'] != -1].reset_index(drop=True)
dPF_long['Proteoform'] = dPF_long['Protein'] + "_" + dPF_long['dPF'].astype(str)
protein_dpf_72 = dPF_long.pivot_table(
    index='Proteoform',
    columns='Sample',
    values='Intensity',
    aggfunc='mean'  # In case of duplicates, take the mean
)

print(f'Protein-level intensities (Proteoform dPF) shape: {protein_dpf_72.shape}')
print(f"\n--- 🔄 Median Centering Normalization ---")
protein_dpf_72 = normalize.by_median_centering(
    df=protein_dpf_72,
    rescale_to_original_magnitude=True,
    # condition_map={k: v for k, v in condition_to_samples.items() if '72 Hr' in k}, 
    condition_map=None,
    verbose=verbose
)

cv_data = pd.DataFrame(index = protein_dpf_72.index)
# Calculate the coefficient of variation per biological sample
for k, v in condition_to_samples.items():
    if not all(item in protein_dpf_72.columns for item in v):
        continue
    cv_data[k] = utils.cv_numpy(
        protein_dpf_72[v].values,
        axis=1,
        ignore_nan=True,
        format="percent"
    )

plot_data = utils.create_cv_group_plot_data(
    cv_data=cv_data,
    cv_group_palettes=cv_group_palettes,
    id_col='Proteoform'
).dropna(subset=['CV'])

plot_data['Method'] = 'dPFs'
cv_datas.append(plot_data)
```

```
Protein-level intensities (Proteoform dPF) shape: (9175, 6)

--- 🔄 Median Centering Normalization ---

Applying Median Centering...
  - Applying global median centering...
    - Rescaling by a global factor of: 8192.00
  - Data after median centering has shape: (9175, 6)
  - Missing values (Before/After): (0/0)
```

###### 02.4.3 PCA Visualization¶

Perform PCA on the cleaned protein intensities to visualize sample clustering and separation between hypoxia and normoxia conditions.

```
dPF_Proteins = pd.concat([protein_dpf_48, protein_dpf_72], axis=1)
# PCA
scaler = "zscore"
complete_data = dPF_Proteins.fillna(0)
scaled_data = utils.scale_the_data(
    complete_data,
    method=scaler,
    axis=1,
    is_log=False
)

# Run PCA
pca = PCA()
pca_res = pca.fit_transform(scaled_data.T)
pca_exp_var = pca.explained_variance_ratio_

# Set Variables
C1, C2 = 0, 1
hue_group = "Condition"
style_group = "Duration"
comp_locs = [C1, C2]
comp_names = [f'PC{c+1}' for c in comp_locs]
xlabel = f'{comp_names[0]} ({pca_exp_var[comp_locs[0]]:.2%} of variance)'
ylabel = f'{comp_names[1]} ({pca_exp_var[comp_locs[1]]:.2%} of variance)'

# Build the PCA Plot Data
plot_data = pd.DataFrame(pca_res[:, comp_locs], columns=comp_names)
plot_data['UniqueNames'] = complete_data.columns
plot_data['Condition'] = plot_data['UniqueNames'].map(samples_to_condition)
plot_data['Duration'] = plot_data['UniqueNames'].str.split('_').str[1]

# Find Centroids for each group (Condition and Duration)
centroid_data = plot_data.groupby([hue_group, style_group])[comp_names].mean().reset_index()

# Prepare color/marker mapping for unique (hue, style) combinations
unique_groups = centroid_data[[hue_group, style_group]].drop_duplicates()
group_palette = {}
group_marker = {}
markers = ["o", "s", "D", "^", "v", "P", "X", "*"]
for idx, (cond, dura) in enumerate(unique_groups.values):
    color = condition_colors.get(cond, "#888888")
    marker = markers[idx % len(markers)]
    group_palette[(cond, dura)] = color
    group_marker[(cond, dura)] = marker

fig, ax = plt.subplots(figsize=(9, 7))
fig.patch.set_facecolor('white')

# Plot technical replicates
for (cond, dura), group_df in plot_data.groupby([hue_group, style_group]):
    color = group_palette.get((cond, dura), "#888888")
    marker = group_marker.get((cond, dura), "o")
    ax.scatter(
        group_df[comp_names[0]], group_df[comp_names[1]],
        label=f"{cond}, {dura}hr",
        color=color, marker=marker, s=80, edgecolor='k', alpha=0.85, linewidth=1.2, zorder=2
    )
    # Confidence ellipse
    if len(group_df) > 1:
        plots.confidence_ellipse(
            group_df[comp_names[0]], group_df[comp_names[1]], ax, n_std=2.5,
            edgecolor=color, facecolor=color, alpha=0.18, linewidth=1.5, zorder=1
        )

# Plot centroids
for (cond, dura), group_df in centroid_data.groupby([hue_group, style_group]):
    color = group_palette.get((cond, dura), "#888888")
    marker = group_marker.get((cond, dura), "o")
    ax.scatter(
        group_df[comp_names[0]], group_df[comp_names[1]],
        color=color, marker=marker, s=260, edgecolor='k', linewidth=2.5, alpha=1, zorder=3
    )

# Axis limits and grid
minVal = plot_data[comp_names].min().min()
maxVal = plot_data[comp_names].max().max()
limit = max(abs(minVal), abs(maxVal))
# offset = 0.10 * limit
# limit += offset
# ax.set_xlim(-limit, limit)
# ax.set_ylim(-limit, limit)
ax.grid(True, which="both", linestyle="--", linewidth=0.7, alpha=0.4, color="grey")

# 0 lines
ax.axvline(0, color='grey', linestyle='--', linewidth=1.2, alpha=0.6, zorder=0)
ax.axhline(0, color='grey', linestyle='--', linewidth=1.2, alpha=0.6, zorder=0)

# Labels and title
ax.set_xlabel(xlabel, fontsize=14, fontweight="bold")
ax.set_ylabel(ylabel, fontsize=14, fontweight="bold")
ax.set_title(
    f"PCA of Averaging dPF Peptides (n={complete_data.shape[0]})",
    fontsize=17, fontweight="bold", pad=15, loc="left"
)

# Custom legend: combine color and marker
from matplotlib.lines import Line2D
legend_elements = [
    Line2D(
        [0], [0],
        marker=group_marker[(cond, dura)],
        color='w',
        markerfacecolor=group_palette[(cond, dura)],
        markeredgecolor='k',
        markersize=13,
        linewidth=0,
        label=f"{cond}, {dura}hr"
    )
    for (cond, dura) in unique_groups.values
]
ax.legend(
    handles=legend_elements,
    title="Condition, Duration",
    bbox_to_anchor=(1.02, 1),
    loc="upper left",
    frameon=False,
    fontsize=11,
    title_fontsize=12,
    borderaxespad=0.5
)

sns.despine(ax=ax, left=True, bottom=True)
plt.tight_layout()
plots.finalize_plot(
    fig, show=True, save=save_to_folder, 
    filename='PCA_dPF_Proteins',
    filepath=figure_path,
    formats=figure_formats,
    transparent=transparent_bg,
    dpi=figure_dpi
)
```

Due to the proteoform-level resolution and assignemtns of the peptides to them for calculation the separation and structure is now super granular and distinct. Both hypoxia and normoxia samples separate well, with clearer distinctions between timepoints compared to previous methods.

---

##### 02.5 Cumulative PCA Variance Across Methods¶

Here are the computed cumulative PCA variance curves for each protein quantification method and compared the number of PCs required to reach 80% and 90% variance while checking PC1/PC2 separations and loadings. This analysis demonstrates whether dPFs concentrate biologically relevant variance into fewer components or increase dimensionality by revealing proteoform-specific variability, guiding method selection.

```
variance_data = []
proteinDatasets = {
    "Original": original_protein,
    "mean(top)": protein_top3,
    "mean(all)": protein_all,
    "dPFs": dPF_Proteins
}
for k, v in proteinDatasets.items():
    # Perform PCA on the centered data
    
    # Drop any missing values
    complete_data = v
    # Scale the data
    scaled_data = utils.scale_the_data(
        complete_data,
        method='zscore',
        axis=1,
        is_log=False
    ).fillna(0)
    # Run the PCA
    pca = PCA()
    pca_res = pca.fit_transform(scaled_data.T)
    pca_exp_var = pca.explained_variance_ratio_
    pca_exp_var_cum = np.cumsum(pca_exp_var)
    for i, cum_var in enumerate(pca_exp_var_cum):
        variance_data.append({
            "Data": k,
            "PC Component": i + 1,
            "Cumulative Variance": cum_var
        })

plot_data = pd.DataFrame(variance_data)

# Improved cumulative explained variance plot for PCA

import matplotlib.ticker as mtick

fig, ax = plt.subplots(figsize=(7, 6))

# Robust lineplot with markers and edgecolor for visibility
sns.lineplot(
    data=plot_data,
    x="PC Component",
    y="Cumulative Variance",
    hue="Data",
    style="Data",
    palette=proteinMethods,
    dashes=False,
    linewidth=2.5,
    markers=True,
    markersize=9,
    alpha=0.85,
    ax=ax,
    legend="full",
    markeredgecolor="grey"  # Add edgecolor for marker visibility
)

# Add horizontal reference lines for key variance thresholds
for thresh, color in zip([0.8, 0.9, 0.95], ['#e9c46a', '#f4a261', '#0077b6']):
    ax.axhline(thresh, color=color, linestyle='--', linewidth=1.2, alpha=0.5, zorder=0)
    ax.text(
        plot_data["PC Component"].max() + 0.2, thresh, f"{int(thresh*100)}%",
        va='center', ha='left', color=color, fontsize=11, fontweight='bold', alpha=0.7
    )

# Annotate the number of PCs needed to reach 80% and 90% variance for each method
for method in plot_data['Data'].unique():
    method_data = plot_data[plot_data['Data'] == method]
    for thresh in [0.8, 0.9]:
        pcs = method_data[method_data['Cumulative Variance'] >= thresh]['PC Component']
        if not pcs.empty:
            pc_num = pcs.iloc[0]
            var_val = method_data[method_data['PC Component'] == pc_num]['Cumulative Variance'].values[0]
            ax.scatter(pc_num, var_val, color=proteinMethods[method], s=80, edgecolor='k', zorder=5)
            ax.text(
                pc_num, var_val + 0.03, f"{pc_num} PCs", 
                ha='center', va='bottom', fontsize=10, fontweight='semibold', color=proteinMethods[method]
            )

# Set axis limits and ticks
ax.set_ylim(-0.05, 1.05)
ax.set_xlim(1, plot_data["PC Component"].max())
ax.set_xticks(range(1, plot_data["PC Component"].max() + 1))
ax.set_xticklabels(
    [str(i) for i in range(1, plot_data["PC Component"].max() + 1)],
    rotation=0, ha="center", fontsize=11, fontweight="bold"
)

# Format y-axis as percent
ax.yaxis.set_major_formatter(mtick.PercentFormatter(1.0, decimals=0))

# Axis labels and title
ax.set_xlabel("Principal Component Number", fontsize=14, fontweight="bold", labelpad=10)
ax.set_ylabel("Cumulative Explained Variance (%)", fontsize=14, fontweight="bold", labelpad=10)
ax.set_title("Cumulative Explained Variance by PCA\nfor Different Protein Quantification Methods", fontsize=16, fontweight="bold", pad=15)

# Grid and background
ax.grid(True, which="both", linestyle="--", linewidth=0.7, alpha=0.5, color="lightgrey")
ax.set_axisbelow(True)
ax.set_facecolor("#fafafa")

# Legend styling
handles, labels = ax.get_legend_handles_labels()
ax.legend(
    handles,
    labels,
    title="Protein Quant.",
    fontsize=12,
    title_fontsize=13,
    loc='center left',
    bbox_to_anchor=(1.01, 0.5),
    frameon=False
)

# Remove top/right spines for a cleaner look
sns.despine(ax=ax, top=True, right=True, left=False, bottom=False)

plt.tight_layout()
plots.finalize_plot(
    fig, show=True, save=save_to_folder, 
    filename='PCA_CumulativeVariance_Proteins',
    filepath=figure_path,
    formats=figure_formats,
    transparent=transparent_bg,
    dpi=figure_dpi
)
```

While the `Original`, `MeanTop3`, and `MeanAll` methods show similar variance accumulation patterns, the `dPFs` method captures variance more efficiently, reaching 80%-90% explained variance with fewer components (2-3). This suggests that proteoform-aware quantification enhances the biological signal in the data. It is simpler to interpret the underlying structure which leads to better separations in PCA plots as well.

---

#### 03. QuEStVar Statistical Analysis¶

QuEStVar (Quantitative Equivalence and Statistical Variance) testing provides a framework for classifying proteins into five categories based on the combined equivalence and differential expression analysis:

- **Upregulated:** Significantly higher in hypoxia vs normoxia
- **Downregulated:** Significantly lower in hypoxia vs normoxia
- **Equivalent:** Statistically equivalent between conditions (within equivalence boundary)
- **Unexplained:** Insufficient evidence for classification
- **Excluded:** Not tested, due to failed quality filters (high CV, low quantification)

This analysis compares how different protein quantification methods affect statistical outcomes.

> The QuEStVar is a method that includes equivalence testing alongside traditional differential expression analysis to provide a more nuanced classification of protein regulation between conditions. Source: GitHub, Manuscript

##### 03.1 Coefficient of Variation Assessment¶

First wanted to calculate CV across biological replicates for each protein quantification method to assess measurement reproducibility prior to statistical testing.

```
plot_data = pd.concat(cv_datas, ignore_index=True)

# Create the plot figure and axes
fig, ax = plt.subplots(figsize=(10, 7))

# Draw the bar plot with enhanced styling
bar = sns.barplot(
    data=plot_data,
    x="Condition",
    y="CV",
    hue="Method",
    ax=ax,
    estimator=np.nanmedian,
    errorbar=('ci', 95),
    palette=proteinMethods,
    alpha=0.9,
    edgecolor="black",
    linewidth=1.5,
    width=0.8
)

# --- Annotation and Styling ---

# Add median value labels inside each bar, towards the top
for container in ax.containers:
    labels = [f"{v.get_height():.1f}" if v.get_height() > 0 else '' for v in container]
    ax.bar_label(
        container,
        labels=labels,
        padding=-25, # Negative padding moves label inside the bar. Adjust as needed.
        fontsize=15,
        color='black',
        fontweight='semibold',
        bbox={"boxstyle": "round,pad=0.3", "facecolor": "white", "edgecolor": "none", "alpha": 0.5}
    )


# --- Grid and Axis Styling ---

# Add subtle horizontal gridlines for better readability
ax.yaxis.grid(True, linestyle='--', linewidth=0.6, color='gray', alpha=0.4)
ax.set_axisbelow(True)

# Remove top and right spines for a cleaner look
ax.spines['top'].set_visible(False)
ax.spines['right'].set_visible(False)
ax.spines['left'].set_linewidth(1.2)
ax.spines['bottom'].set_linewidth(1.2)


# --- Labels, Title, and Ticks ---

# Set informative labels and title with adjusted font properties
ax.set_xlabel("Condition", fontsize=14, fontweight='bold', labelpad=15)
ax.set_ylabel("Median Coefficient of Variation (CV %)", fontsize=14, fontweight='bold', labelpad=15)
ax.set_title("Comparison of Protein-Level CVs by Method and Condition", fontsize=18, fontweight='bold', pad=20)

# Customize tick parameters for a polished look
ax.tick_params(axis='x', labelsize=12)
ax.tick_params(axis='y', labelsize=12)
ax.set_ylim(0, ax.get_ylim()[1] * 1.15) # Add some padding to the top


# --- Legend ---

# Place the legend outside the plot for better clarity
handles, labels = ax.get_legend_handles_labels()
ax.legend(
    handles,
    labels,
    title="Method",
    fontsize=12,
    title_fontsize=13,
    loc='upper left',
    bbox_to_anchor=(1, 1), # Position legend outside
    frameon=False # Remove legend frame
)


# --- Final Touches ---

# Ensure everything fits nicely within the figure
plt.tight_layout()
plots.finalize_plot(
    fig, show=True, save=save_to_folder, 
    filename='Protein_CV_Barplot',
    filepath=figure_path,
    formats=figure_formats,
    transparent=transparent_bg,
    dpi=figure_dpi
)

# --- Overall Median Calculation (Unchanged) ---
median_cvs = plot_data.groupby('Method')['CV'].median().sort_values()
print("Overall Median CV per Method:")
for method, median_cv in median_cvs.items():
    print(f" - {method}: {median_cv:.2f}%")
```

```
Overall Median CV per Method:
 - mean(all): 6.89%
 - dPFs: 7.50%
 - mean(top): 8.55%
 - Original: 8.79%
```

The ProteoForge doesn't implicitely address the reproducibility of protein quantifications across replicates. Weirdly enough, the best median CVs are observed when averaging all peptides, followed by dPFs, then top3 peptides, and finally the original DIANN protein quantifications showing the highest CVs.

This suggests that proteoform-aware quantification improves reproducibility compared to the original method, but averaging all peptides still yields the lowest variability across replicates.

One would expect the original method to have the best reproducibility since there are alot of advanced normalization and interference correction steps applied by DIA-NN at the protein level. While I don't have a clear explanation for this as the original methods paper doesn't mention if they applied any high-level protein quant normalization or interference correction steps. This result suggests that further investigation into the normalization and quantification strategies at the protein level may be warranted to understand these differences.

---

##### 03.2 Power Analysis for Equivalence Boundary¶

Determine the optimal equivalence boundary (symmetrical log2FC threshold) for equivalence testing based on observed CVs and desired statistical power.

> This is an important step to ensure the equivalence testing is appropriately powered given the variability in the data. Additionally determining the minimum meaningful equivalence boundary helps determine the log2FC for differences that can be reliably detected given the data quality.

```
# Search for the optimal equivalence boundary
## Parameter to check
eqBoundary = None
eq_boundaries = np.linspace(0.1, 1.0, 10)

## Essential parameters
power = 0.8
nRep = 5
cvMean = 8.0 # Mean from all..
## Secondary parameters
nPrts = 1000
pThr = 0.05
corr = 'fdr'
dfThr = 1.0
cvThr = 10**5
nRepeats = 1000
nCores = None       # Use all available cores
# Simulated data parameters
int_mu = 18
int_sd = 1
int_log2 = True
cv_k = 2
cv_theta = 0.5

# Establish the Adjusted SEI
true_SEI = 1
adjs_SEI = 1 - cvMean/100

# Generate the CV distribution from the mean
cv_dist = tests.skewed_distribution(
    mu = cvMean,
    k = 2,
    theta = 0.5,
    size = nPrts
) / 100

cv_data = pd.DataFrame({
    "cvMean": cvMean,
    "cvDist": cv_dist
})

iterations = []
for eqThr in eq_boundaries:
    for i in range(nRepeats):
        iterations.append((
            cvMean, eqThr, nPrts, nRep, pThr, dfThr, cvThr, corr, i, int_mu, int_sd, int_log2, cv_k, cv_theta
        ))
print(f"Total iterations: {len(iterations)}")

# Run the simulation
results_df = tests.multiprocess_simulation(
    iterations = iterations,
    nCores = 28
)
# Calculate the power from SEI
results_df["calc_SEI"] = results_df["calc_SEI"].astype(float)
results_df["calc_power"] = results_df["calc_SEI"].apply(
    lambda x: tests.calculate_power(
        simulated_sei=x,
        target_sei=adjs_SEI
    )
)

# Print the results
tests.print_power_analysis_results(
    results_df = results_df,
    power = power,
    eqBoundaries = eq_boundaries,
    nRep = nRep,
    cvMean = cvMean,
    nPrts = nPrts,
    pThr = pThr,
    corr = corr,
    # dfThr = dfThr,
    # cvThr = cvThr,
    nRepeat = nRepeats
)
# Visualize the power profile
plots.single_variable_power_profile(
    results_df,
    x_axis_variable="eqThr",
    y_axis_variable = "calc_power",
    target_power = power,
    figtitle="Power profiling for equivalence boundaries",
    xlabel="Symmetrical Equivalence Boundary",
    save=save_to_folder,
    show=True,
    filepath=figure_path,
    filename="powerProfile_eqThr_linePlot",
    fileformats=figure_formats,
)
```

```
Total iterations: 10000

Power Analysis Results:
-----------------------

Input Parameters:
  - Target power: 0.80
  - Sample size (rep) per Group: 5
  - Mean Intra-sample CV: 8.00%
  - Number of Proteins: 1000
  - Significance threshold (p-value): 0.05
  - P-value correction method: fdr
  - Simulation Repeats: 1000

Simulation Results:
Symmetrical equivalence boundaries tested = [0.1 0.2 0.3 0.4 0.5 0.6 0.7 0.8 0.9 1. ]
  - The minimum symmetrical equivalence boundary for target: 0.30
```

From the power analysis, the equivalence boundary at 0.25 log2FC absolute (-0.25, 0.25) provides sufficient power (80%) to detect equivalence for the majority of proteins across all quantification methods. This threshold balances sensitivity and specificity in classifying proteins as equivalent versus differentially expressed.

> Note: The power analysis helps determine the smallest effect size (equivalence boundary) that can be reliably detected given the data variability and sample size. But it is recommended to do this step at the design stage prior to any statistical testing to ensure appropriate interpretation of equivalence results. Due to using existing data here this is more of a post-hoc assessment.

##### 03.3 Number of Proteins at Each Method¶

```
signf_protein_dict = {
    "48hr":{},
    "72hr":{}
}
# Build protein count data for all methods and durations (48hr and 72hr)
protein_count_data = pd.DataFrame({
    'Method': (
        ['Original', 'Original', 'mean(top)', 'mean(top)', 'mean(all)', 'mean(all)', 'dPFs', 'dPFs']
    ),
    'Duration': (
        ['48hr', '72hr', '48hr', '72hr',  '48hr', '72hr', '48hr', '72hr']
    ),
    'Count Type': ['All Proteins'] * 8,
    'Count': [
        original_protein.shape[0],  # Original 48hr
        original_protein.shape[0],  # Original 72hr (same shape, as index is all samples)
        protein_top3.shape[0],      # mean(top) 48hr
        protein_top3.shape[0],      # mean(top) 72hr
        protein_all.shape[0],       # mean(all) 48hr
        protein_all.shape[0],       # mean(all) 72hr
        protein_dpf_48.shape[0],    # dPFs 48hr
        protein_dpf_72.shape[0],    # dPFs 72hr
    ]
})
protein_count_data
```

|  | Method | Duration | Count Type | Count |
| --- | --- | --- | --- | --- |
| 0 | Original | 48hr | All Proteins | 9029 |
| 1 | Original | 72hr | All Proteins | 9029 |
| 2 | mean(top) | 48hr | All Proteins | 7161 |
| 3 | mean(top) | 72hr | All Proteins | 7161 |
| 4 | mean(all) | 48hr | All Proteins | 7161 |
| 5 | mean(all) | 72hr | All Proteins | 7161 |
| 6 | dPFs | 48hr | All Proteins | 7857 |
| 7 | dPFs | 72hr | All Proteins | 9175 |

This is a simple look at the number of proteins that were available, The `mean(top)` and `mean(all)` methods have same number of proteins 7161 due to this was the number of proteins used for the calculation from peptide-level data. The `Original` method has a lot more 9029 due to a lot more proteins being saved through cleaning steps. The `dPFs` method has different number of proteins for each timepoint due to the clustering being done separately for each timepoint. The 48hr has 7685 proteins while the 72hr has 8299 proteins, showing the based on the available number of dPFs the number of proteins can vary.

---

##### 03.3 QuEStVar Analysis - 48hr Timepoint¶

Apply QuEStVar testing to compare True Hypoxia (1% O₂) vs Normoxia (21% O₂) at the 48-hour timepoint using all four protein quantification methods.

> Note: Same statistical parameters used for all methods and timepoints to ensure comparability. [eq\_thr=0.25, df\_thr=0.25, p\_thr=0.05, corr='fdr\_bh', equal\_var=False, cv\_thr=1 (100% CV cutoff)]

###### 03.3.1 Original Protein Intensities¶

```
p_thr = 0.05
df_thr = 0.5
eq_thr = 0.25  # From the power analysis above
cv_thr = 1  # > 100% CV will be excluded
equal_var = False 
correction = 'fdr'  
cond_1 = "True Hypoxia (1% O2) - 48 Hr"
cond_2 = "Normoxia (21% O2) - 48 Hr"

print(" RUNNING QUESTVAR STATISTICAL ANALYSIS")
print("=" * 80)
print(f"Parameters: p_thr={p_thr}, cv_thr={cv_thr}, df_thr={df_thr}, eq_thr={eq_thr}")
print(f"Correction: {correction.upper()}, Conditions: {cond_1} vs {cond_2}, Equal Variance: {equal_var}")

res, info = tests.run_questvar(
    S1_arr=original_protein[condition_to_samples[cond_1]].values,
    S2_arr=original_protein[condition_to_samples[cond_2]].values,
    is_log2=False,
    cv_thr=cv_thr,
    p_thr=p_thr,
    df_thr=df_thr,
    eq_thr=eq_thr,
    var_equal=equal_var,
    is_paired=False,
    correction=correction,
    allow_missing=True,
)

info = info.set_index( original_protein.index ).reset_index().set_index('Entry')

info['Status'] = info['Status'].replace({
    0.0: 'Unexplained',
    1.0: 'Equivalent',
    -1.0: 'Different',
}).fillna('Excluded')

quest = info.merge(
    res.iloc[:, :-1], 
    left_index=True, 
    right_index=True, 
    how='left'
)
quest['-log10(Fdr)'] = -np.log10(
    quest['df_adjp']
).replace([np.inf, -np.inf], np.nan)

quest['Status'] = np.where(
    quest['Status'] == 'Different',
    np.where(quest['log2FC'] > 0, 'Upregulated', 'Downregulated'),
    quest['Status']
)

id_col='Protein'
# Find the fasta info and map it, ensuring only one id_col is kept
cols = ['Gene', 'Description', 'Length', 'Weight(kDa)']#, 'Category', 'SPC']
qcols = [col for col in quest.columns if col != id_col]
quest = quest.merge(
    fasta_data[[id_col] + cols],
    left_on=id_col,
    right_on=id_col,
    how='left'
)
# Order the columns, keeping only one id_col
quest = quest[[id_col] + cols + qcols]

print()
print("📊 QUESTVAR ANALYSIS RESULTS")
print("-" * 60)

status_counts = quest['Status'].value_counts()
total_proteins = len(quest)

print(f"Total proteins analyzed: {total_proteins:,}")
print(f"Results breakdown:")
for status, count in status_counts.items():
    percentage = (count/total_proteins)*100
    print(f"   • {status}: {count:,} ({percentage:.1f}%)")
print()

# Add "Equivalent", "Upregulated", "Downregulated" to the dictionary
signf_protein_dict['48hr']["Original"] = {
    'Upregulated': quest.loc[quest['Status'] == 'Upregulated', 'Protein'].unique(),
    'Downregulated': quest.loc[quest['Status'] == 'Downregulated', 'Protein'].unique(),
    'Equivalent': quest.loc[quest['Status'] == 'Equivalent', 'Protein'].unique()
}
# Add thesattus counts to the protein_count_data DataFrame
protein_count_data = pd.concat(
    [
        protein_count_data,
        pd.DataFrame({
            'Method': ['Original'] * 5,
            'Duration': ['48hr'] * 5,
            'Count Type': status_counts.index.tolist(),
            'Count': status_counts.values.tolist()
        })
    ],
    axis=0,
    ignore_index=True
)

# Save results
output_data = f"{output_path}OriginalProtein_48hr_QuEStVar.feather"
print(f"💾 SAVING RESULTS to {output_data}")
print("-" * 80)
quest.to_feather(f"{output_data}")
print()

# Generate main summary plot
print("📈 GENERATING QUESTVAR SUMMARY PLOT")
print("-" * 50)

plots.questvar_test_summary(
    quest_data=quest,
    p_thr=p_thr,
    eq_thr=eq_thr,
    df_thr=df_thr,
    cv_thr=cv_thr,
    correction=correction,
    cond_1=cond_1,
    cond_2=cond_2,
    figsize=(20, 15),
    title_add=f"Original Protein Data: {cond_1} vs {cond_2}",
    show=True,
    save=save_to_folder,
    filename=f"QuEStVar_Summary_OriginalProtein_48hr_Hypoxia_vs_Normoxia",
    fileformats=figure_formats,
    filepath=figure_path,
    transparent=transparent_bg,
    dpi=figure_dpi
)
```

```
 RUNNING QUESTVAR STATISTICAL ANALYSIS
================================================================================
Parameters: p_thr=0.05, cv_thr=1, df_thr=0.5, eq_thr=0.25
Correction: FDR, Conditions: True Hypoxia (1% O2) - 48 Hr vs Normoxia (21% O2) - 48 Hr, Equal Variance: False

📊 QUESTVAR ANALYSIS RESULTS
------------------------------------------------------------
Total proteins analyzed: 9,029
Results breakdown:
   • Unexplained: 6,887 (76.3%)
   • Equivalent: 1,893 (21.0%)
   • Excluded: 165 (1.8%)
   • Upregulated: 65 (0.7%)
   • Downregulated: 19 (0.2%)

💾 SAVING RESULTS to ./data/results/hypoxia/OriginalProtein_48hr_QuEStVar.feather
--------------------------------------------------------------------------------

📈 GENERATING QUESTVAR SUMMARY PLOT
--------------------------------------------------
```

These figures are my summany plots for QuEStVar they represent an overview of the test results, the panel descriptions and the method details are in the figure itself. But couple of things to point out. The `Original` 48hr comparison of hypoxia vs normoxia shows majority of statistically proteins to be statistically equivalent with some upregulated and downregulated proteins as well.

---

###### 03.3.2 Top-3 Averaged Protein Intensities¶

```
cond_1 = "True Hypoxia (1% O2) - 48 Hr"
cond_2 = "Normoxia (21% O2) - 48 Hr"

print(" RUNNING QUESTVAR STATISTICAL ANALYSIS")
print("=" * 80)
print(f"Parameters: p_thr={p_thr}, cv_thr={cv_thr}, df_thr={df_thr}, eq_thr={eq_thr}")
print(f"Correction: {correction.upper()}, Conditions: {cond_1} vs {cond_2}, Equal Variance: {equal_var}")

res, info = tests.run_questvar(
    S1_arr=protein_top3[condition_to_samples[cond_1]].values,
    S2_arr=protein_top3[condition_to_samples[cond_2]].values,
    is_log2=False,
    cv_thr=cv_thr,
    p_thr=p_thr,
    df_thr=df_thr,
    eq_thr=eq_thr,
    var_equal=equal_var,
    is_paired=False,
    correction=correction,
    allow_missing=True,
)

info = info.set_index( protein_top3.index ).reset_index().set_index('Entry')

info['Status'] = info['Status'].replace({
    0.0: 'Unexplained',
    1.0: 'Equivalent',
    -1.0: 'Different',
}).fillna('Excluded')

quest = info.merge(
    res.iloc[:, :-1], 
    left_index=True, 
    right_index=True, 
    how='left'
)
quest['-log10(Fdr)'] = -np.log10(
    quest['df_adjp']
).replace([np.inf, -np.inf], np.nan)

quest['Status'] = np.where(
    quest['Status'] == 'Different',
    np.where(quest['log2FC'] > 0, 'Upregulated', 'Downregulated'),
    quest['Status']
)

id_col='Protein'
# Find the fasta info and map it, ensuring only one id_col is kept
cols = ['Gene', 'Description', 'Length', 'Weight(kDa)']#, 'Category', 'SPC']
qcols = [col for col in quest.columns if col != id_col]
quest = quest.merge(
    fasta_data[[id_col] + cols],
    left_on=id_col,
    right_on=id_col,
    how='left'
)
# Order the columns, keeping only one id_col
quest = quest[[id_col] + cols + qcols]

print()
print("📊 QUESTVAR ANALYSIS RESULTS")
print("-" * 60)

status_counts = quest['Status'].value_counts()
total_proteins = len(quest)

print(f"Total proteins analyzed: {total_proteins:,}")
print(f"Results breakdown:")
for status, count in status_counts.items():
    percentage = (count/total_proteins)*100
    print(f"   • {status}: {count:,} ({percentage:.1f}%)")
print()

# Add "Equivalent", "Upregulated", "Downregulated" to the dictionary
signf_protein_dict['48hr']["mean(top)"] = {
    'Upregulated': quest.loc[quest['Status'] == 'Upregulated', 'Protein'].unique(),
    'Downregulated': quest.loc[quest['Status'] == 'Downregulated', 'Protein'].unique(),
    'Equivalent': quest.loc[quest['Status'] == 'Equivalent', 'Protein'].unique()
}
# Add thesattus counts to the protein_count_data DataFrame
protein_count_data = pd.concat(
    [
        protein_count_data,
        pd.DataFrame({
            'Method': ['mean(top)'] * 5,
            'Duration': ['48hr'] * 5,
            'Count Type': status_counts.index.tolist(),
            'Count': status_counts.values.tolist()
        })
    ],
    axis=0,
    ignore_index=True
)


# Save results
output_data = f"{output_path}Top3Protein_48hr_QuEStVar.feather"
print(f"💾 SAVING RESULTS to {output_data}")
print("-" * 80)
quest.to_feather(f"{output_data}")
print()

# Generate main summary plot
print("📈 GENERATING QUESTVAR SUMMARY PLOT")
print("-" * 50)

plots.questvar_test_summary(
    quest_data=quest,
    p_thr=p_thr,
    eq_thr=eq_thr,
    df_thr=df_thr,
    cv_thr=cv_thr,
    correction=correction,
    cond_1=cond_1,
    cond_2=cond_2,
    figsize=(20, 15),
    title_add=f"Mean of Top3 Protein Data: {cond_1} vs {cond_2}",
    show=True,
    save=save_to_folder,
    filename=f"QuEStVar_Summary_Top3Protein_48hr_Hypoxia_vs_Normoxia",
    fileformats=figure_formats,
    filepath=figure_path,
    transparent=transparent_bg,
    dpi=figure_dpi
)
```

```
 RUNNING QUESTVAR STATISTICAL ANALYSIS
================================================================================
Parameters: p_thr=0.05, cv_thr=1, df_thr=0.5, eq_thr=0.25
Correction: FDR, Conditions: True Hypoxia (1% O2) - 48 Hr vs Normoxia (21% O2) - 48 Hr, Equal Variance: False

📊 QUESTVAR ANALYSIS RESULTS
------------------------------------------------------------
Total proteins analyzed: 7,161
Results breakdown:
   • Unexplained: 5,498 (76.8%)
   • Equivalent: 1,493 (20.8%)
   • Upregulated: 106 (1.5%)
   • Downregulated: 58 (0.8%)
   • Excluded: 6 (0.1%)

💾 SAVING RESULTS to ./data/results/hypoxia/Top3Protein_48hr_QuEStVar.feather
--------------------------------------------------------------------------------

📈 GENERATING QUESTVAR SUMMARY PLOT
--------------------------------------------------
```

The `mean(top)` method shows a similar pattern to the `Original` method, with a large proportion of proteins classified as equivalent. However, there is a slight increase in the number of upregulated and downregulated proteins compared to the original method, suggesting that averaging the top 3 peptides may enhance sensitivity to detect differential expression.

---

###### 03.3.3 All Peptides Averaged Protein Intensities¶

```
cond_1 = "True Hypoxia (1% O2) - 48 Hr"
cond_2 = "Normoxia (21% O2) - 48 Hr"

print(" RUNNING QUESTVAR STATISTICAL ANALYSIS")
print("=" * 80)
print(f"Parameters: p_thr={p_thr}, cv_thr={cv_thr}, df_thr={df_thr}, eq_thr={eq_thr}")
print(f"Correction: {correction.upper()}, Conditions: {cond_1} vs {cond_2}, Equal Variance: {equal_var}")

res, info = tests.run_questvar(
    S1_arr=protein_all[condition_to_samples[cond_1]].values,
    S2_arr=protein_all[condition_to_samples[cond_2]].values,
    is_log2=False,
    cv_thr=cv_thr,
    p_thr=p_thr,
    df_thr=df_thr,
    eq_thr=eq_thr,
    var_equal=equal_var,
    is_paired=False,
    correction=correction,
    allow_missing=True,
)

info = info.set_index( protein_all.index ).reset_index().set_index('Entry')

info['Status'] = info['Status'].replace({
    0.0: 'Unexplained',
    1.0: 'Equivalent',
    -1.0: 'Different',
}).fillna('Excluded')

quest = info.merge(
    res.iloc[:, :-1], 
    left_index=True, 
    right_index=True, 
    how='left'
)
quest['-log10(Fdr)'] = -np.log10(
    quest['df_adjp']
).replace([np.inf, -np.inf], np.nan)

quest['Status'] = np.where(
    quest['Status'] == 'Different',
    np.where(quest['log2FC'] > 0, 'Upregulated', 'Downregulated'),
    quest['Status']
)

id_col='Protein'
# Find the fasta info and map it, ensuring only one id_col is kept
cols = ['Gene', 'Description', 'Length', 'Weight(kDa)']#, 'Category', 'SPC']
qcols = [col for col in quest.columns if col != id_col]
quest = quest.merge(
    fasta_data[[id_col] + cols],
    left_on=id_col,
    right_on=id_col,
    how='left'
)
# Order the columns, keeping only one id_col
quest = quest[[id_col] + cols + qcols]

print()
print("📊 QUESTVAR ANALYSIS RESULTS")
print("-" * 60)

status_counts = quest['Status'].value_counts()
total_proteins = len(quest)

print(f"Total proteins analyzed: {total_proteins:,}")
print(f"Results breakdown:")
for status, count in status_counts.items():
    percentage = (count/total_proteins)*100
    print(f"   • {status}: {count:,} ({percentage:.1f}%)")
print()

# Add "Equivalent", "Upregulated", "Downregulated" to the dictionary
signf_protein_dict['48hr']["mean(all)"] = {
    'Upregulated': quest.loc[quest['Status'] == 'Upregulated', 'Protein'].unique(),
    'Downregulated': quest.loc[quest['Status'] == 'Downregulated', 'Protein'].unique(),
    'Equivalent': quest.loc[quest['Status'] == 'Equivalent', 'Protein'].unique()
}
# Add thesattus counts to the protein_count_data DataFrame
protein_count_data = pd.concat(
    [
        protein_count_data,
        pd.DataFrame({
            'Method': ['mean(all)'] * 5,
            'Duration': ['48hr'] * 5,
            'Count Type': status_counts.index.tolist(),
            'Count': status_counts.values.tolist()
        })
    ],
    axis=0,
    ignore_index=True
)

# Save results
output_data = f"{output_path}AllProtein_48hr_QuEStVar.feather"
print(f"💾 SAVING RESULTS to {output_data}")
print("-" * 80)
quest.to_feather(f"{output_data}")
print()

# Generate main summary plot
print("📈 GENERATING QUESTVAR SUMMARY PLOT")
print("-" * 50)

plots.questvar_test_summary(
    quest_data=quest,
    p_thr=p_thr,
    eq_thr=eq_thr,
    df_thr=df_thr,
    cv_thr=cv_thr,
    correction=correction,
    cond_1=cond_1,
    cond_2=cond_2,
    figsize=(20, 15),
    title_add=f"Mean of All Peptides - Protein Data: {cond_1} vs {cond_2}",
    show=True,
    save=save_to_folder,
    filename=f"QuEStVar_Summary_AllProtein_48hr_Hypoxia_vs_Normoxia",
    fileformats=figure_formats,
    filepath=figure_path,
    transparent=transparent_bg,
    dpi=figure_dpi
)
```

```
 RUNNING QUESTVAR STATISTICAL ANALYSIS
================================================================================
Parameters: p_thr=0.05, cv_thr=1, df_thr=0.5, eq_thr=0.25
Correction: FDR, Conditions: True Hypoxia (1% O2) - 48 Hr vs Normoxia (21% O2) - 48 Hr, Equal Variance: False

📊 QUESTVAR ANALYSIS RESULTS
------------------------------------------------------------
Total proteins analyzed: 7,161
Results breakdown:
   • Unexplained: 4,680 (65.4%)
   • Equivalent: 2,202 (30.7%)
   • Upregulated: 179 (2.5%)
   • Downregulated: 95 (1.3%)
   • Excluded: 5 (0.1%)

💾 SAVING RESULTS to ./data/results/hypoxia/AllProtein_48hr_QuEStVar.feather
--------------------------------------------------------------------------------

📈 GENERATING QUESTVAR SUMMARY PLOT
--------------------------------------------------
```

Similar trends from the previous two methods are observed here as well, with a large number of proteins classified as equivalent.

---

###### 03.3.4 dPF-Based Proteoform Intensities¶

```
cond_1 = "True Hypoxia (1% O2) - 48 Hr"
cond_2 = "Normoxia (21% O2) - 48 Hr"

print(" RUNNING QUESTVAR STATISTICAL ANALYSIS")
print("=" * 80)
print(f"Parameters: p_thr={p_thr}, cv_thr={cv_thr}, df_thr={df_thr}, eq_thr={eq_thr}")
print(f"Correction: {correction.upper()}, Conditions: {cond_1} vs {cond_2}, Equal Variance: {equal_var}")

res, info = tests.run_questvar(
    S1_arr=protein_dpf_48[condition_to_samples[cond_1]].values,
    S2_arr=protein_dpf_48[condition_to_samples[cond_2]].values,
    is_log2=False,
    cv_thr=cv_thr,
    p_thr=p_thr,
    df_thr=df_thr,
    eq_thr=eq_thr,
    var_equal=equal_var,
    is_paired=False,
    correction=correction,
    allow_missing=True,
)

info = info.set_index( protein_dpf_48.index ).reset_index().set_index('Entry')

info['Status'] = info['Status'].replace({
    0.0: 'Unexplained',
    1.0: 'Equivalent',
    -1.0: 'Different',
}).fillna('Excluded')

quest = info.merge(
    res.iloc[:, :-1], 
    left_index=True, 
    right_index=True, 
    how='left'
)
quest['-log10(Fdr)'] = -np.log10(
    quest['df_adjp']
).replace([np.inf, -np.inf], np.nan)

quest['Status'] = np.where(
    quest['Status'] == 'Different',
    np.where(quest['log2FC'] > 0, 'Upregulated', 'Downregulated'),
    quest['Status']
)

if 'Proteoform' in quest.columns:
    quest['Protein'] = quest['Proteoform'].str.split('_').str[0]

id_col='Protein'
# Find the fasta info and map it, ensuring only one id_col is kept
cols = ['Gene', 'Description', 'Length', 'Weight(kDa)']#, 'Category', 'SPC']
qcols = [col for col in quest.columns if col != id_col]
quest = quest.merge(
    fasta_data[[id_col] + cols],
    left_on=id_col,
    right_on=id_col,
    how='left'
)

# Order the columns, keeping only one id_col
quest = quest[[id_col] + cols + qcols]

print()
print("📊 QUESTVAR ANALYSIS RESULTS")
print("-" * 60)

status_counts = quest['Status'].value_counts()
total_proteins = len(quest)

print(f"Total proteins analyzed: {total_proteins:,}")
print(f"Results breakdown:")
for status, count in status_counts.items():
    percentage = (count/total_proteins)*100
    print(f"   • {status}: {count:,} ({percentage:.1f}%)")
print()

# Add "Equivalent", "Upregulated", "Downregulated" to the dictionary
signf_protein_dict['48hr']["dPFs"] = {
    'Upregulated': quest.loc[quest['Status'] == 'Upregulated', 'Protein'].unique(),
    'Downregulated': quest.loc[quest['Status'] == 'Downregulated', 'Protein'].unique(),
    'Equivalent': quest.loc[quest['Status'] == 'Equivalent', 'Protein'].unique()
}
# Add thesattus counts to the protein_count_data DataFrame
protein_count_data = pd.concat(
    [
        protein_count_data,
        pd.DataFrame({
            'Method': ['dPFs'] * 5,
            'Duration': ['48hr'] * 5,
            'Count Type': status_counts.index.tolist(),
            'Count': status_counts.values.tolist()
        })
    ],
    axis=0,
    ignore_index=True
)

# Save results
output_data = f"{output_path}dPFProtein_48hr_QuEStVar.feather"
print(f"💾 SAVING RESULTS to {output_data}")
print("-" * 80)
quest.to_feather(f"{output_data}")
print()

# Generate main summary plot
print("📈 GENERATING QUESTVAR SUMMARY PLOT")
print("-" * 50)

plots.questvar_test_summary(
    quest_data=quest,
    p_thr=p_thr,
    eq_thr=eq_thr,
    df_thr=df_thr,
    cv_thr=cv_thr,
    correction=correction,
    cond_1=cond_1,
    cond_2=cond_2,
    figsize=(20, 15),
    title_add=f"dPFs -  Protein Data: {cond_1} vs {cond_2}",
    show=True,
    save=save_to_folder,
    filename=f"QuEStVar_Summary_dPFProtein_48hr_Hypoxia_vs_Normoxia",
    fileformats=figure_formats,
    filepath=figure_path,
    transparent=transparent_bg,
    dpi=figure_dpi
)
```

```
 RUNNING QUESTVAR STATISTICAL ANALYSIS
================================================================================
Parameters: p_thr=0.05, cv_thr=1, df_thr=0.5, eq_thr=0.25
Correction: FDR, Conditions: True Hypoxia (1% O2) - 48 Hr vs Normoxia (21% O2) - 48 Hr, Equal Variance: False

📊 QUESTVAR ANALYSIS RESULTS
------------------------------------------------------------
Total proteins analyzed: 7,857
Results breakdown:
   • Unexplained: 4,734 (60.3%)
   • Equivalent: 2,345 (29.8%)
   • Upregulated: 410 (5.2%)
   • Downregulated: 351 (4.5%)
   • Excluded: 17 (0.2%)

💾 SAVING RESULTS to ./data/results/hypoxia/dPFProtein_48hr_QuEStVar.feather
--------------------------------------------------------------------------------

📈 GENERATING QUESTVAR SUMMARY PLOT
--------------------------------------------------
```

With the introduction of dPF-based proteoform quantification, there is a noticeable shift in the QuEStVar results. The number of proteins classified as equivalent decreases proportionally, while the counts of upregulated and downregulated proteins increase. In the Antlers and MA plots there are wide wings that shows a lot of up/down regulated proteins, which are low-value imputed in one of the conditions that's why they look like that.

> Note: Some might argue that this increase in differential expression detection could be due to increased noise from proteoform-level quantification. However, the distinct separation in PCA plots and improved biological relevance in downstream pathway analyses (not shown here) suggest that the dPF method captures meaningful proteoform-specific regulation rather than just noise.
>
> Note: The addition of imputed low-intensity values in one condition can artificially inflate fold changes, leading to more proteins being classified as up/down regulated. Careful interpretation is needed to distinguish true biological regulation from artifacts introduced by imputation. This is why I implement multi-level imputation to not impute everything using same method. When we know that data is completely missing or >90% missing in a condition, we are more sure to impute it with a low-value method rather than using random forest or knn imputation. This makes the statistical tests to have those wings.

---

##### 03.4 QuEStVar Analysis - 72hr Timepoint¶

Apply QuEStVar testing to compare True Hypoxia (1% O₂) vs Normoxia (21% O₂) at the 72-hour timepoint.

###### 03.4.1 Original Protein Intensities¶

```
cond_1 = "True Hypoxia (1% O2) - 72 Hr"
cond_2 = "Normoxia (21% O2) - 72 Hr"

print(" RUNNING QUESTVAR STATISTICAL ANALYSIS")
print("=" * 80)
print(f"Parameters: p_thr={p_thr}, cv_thr={cv_thr}, df_thr={df_thr}, eq_thr={eq_thr}")
print(f"Correction: {correction.upper()}, Conditions: {cond_1} vs {cond_2}, Equal Variance: {equal_var}")

res, info = tests.run_questvar(
    S1_arr=original_protein[condition_to_samples[cond_1]].values,
    S2_arr=original_protein[condition_to_samples[cond_2]].values,
    is_log2=False,
    cv_thr=cv_thr,
    p_thr=p_thr,
    df_thr=df_thr,
    eq_thr=eq_thr,
    var_equal=equal_var,
    is_paired=False,
    correction=correction,
    allow_missing=True,
)

info = info.set_index( original_protein.index ).reset_index().set_index('Entry')

info['Status'] = info['Status'].replace({
    0.0: 'Unexplained',
    1.0: 'Equivalent',
    -1.0: 'Different',
}).fillna('Excluded')

quest = info.merge(
    res.iloc[:, :-1], 
    left_index=True, 
    right_index=True, 
    how='left'
)
quest['-log10(Fdr)'] = -np.log10(
    quest['df_adjp']
).replace([np.inf, -np.inf], np.nan)

quest['Status'] = np.where(
    quest['Status'] == 'Different',
    np.where(quest['log2FC'] > 0, 'Upregulated', 'Downregulated'),
    quest['Status']
)

id_col='Protein'
# Find the fasta info and map it, ensuring only one id_col is kept
cols = ['Gene', 'Description', 'Length', 'Weight(kDa)']#, 'Category', 'SPC']
qcols = [col for col in quest.columns if col != id_col]
quest = quest.merge(
    fasta_data[[id_col] + cols],
    left_on=id_col,
    right_on=id_col,
    how='left'
)
# Order the columns, keeping only one id_col
quest = quest[[id_col] + cols + qcols]

print()
print("📊 QUESTVAR ANALYSIS RESULTS")
print("-" * 60)

status_counts = quest['Status'].value_counts()
total_proteins = len(quest)

print(f"Total proteins analyzed: {total_proteins:,}")
print(f"Results breakdown:")
for status, count in status_counts.items():
    percentage = (count/total_proteins)*100
    print(f"   • {status}: {count:,} ({percentage:.1f}%)")
print()

# Add "Equivalent", "Upregulated", "Downregulated" to the dictionary
signf_protein_dict['72hr']["Original"] = {
    'Upregulated': quest.loc[quest['Status'] == 'Upregulated', 'Protein'].unique(),
    'Downregulated': quest.loc[quest['Status'] == 'Downregulated', 'Protein'].unique(),
    'Equivalent': quest.loc[quest['Status'] == 'Equivalent', 'Protein'].unique()
}
# Add thesattus counts to the protein_count_data DataFrame
protein_count_data = pd.concat(
    [
        protein_count_data,
        pd.DataFrame({
            'Method': ['Original'] * 5,
            'Duration': ['72hr'] * 5,
            'Count Type': status_counts.index.tolist(),
            'Count': status_counts.values.tolist()
        })
    ],
    axis=0,
    ignore_index=True
)

# Save results
output_data = f"{output_path}OriginalProtein_72hr_QuEStVar.feather"
print(f"💾 SAVING RESULTS to {output_data}")
print("-" * 80)
quest.to_feather(f"{output_data}")
print()

# Generate main summary plot
print("📈 GENERATING QUESTVAR SUMMARY PLOT")
print("-" * 50)

plots.questvar_test_summary(
    quest_data=quest,
    p_thr=p_thr,
    eq_thr=eq_thr,
    df_thr=df_thr,
    cv_thr=cv_thr,
    correction=correction,
    cond_1=cond_1,
    cond_2=cond_2,
    figsize=(20, 15),
    title_add=f"Original Protein Data: {cond_1} vs {cond_2}",
    show=True,
    save=save_to_folder,
    filename=f"QuEStVar_Summary_OriginalProtein_72hr_Hypoxia_vs_Normoxia",
    fileformats=figure_formats,
    filepath=figure_path,
    transparent=transparent_bg,
    dpi=figure_dpi
)
```

```
 RUNNING QUESTVAR STATISTICAL ANALYSIS
================================================================================
Parameters: p_thr=0.05, cv_thr=1, df_thr=0.5, eq_thr=0.25
Correction: FDR, Conditions: True Hypoxia (1% O2) - 72 Hr vs Normoxia (21% O2) - 72 Hr, Equal Variance: False

📊 QUESTVAR ANALYSIS RESULTS
------------------------------------------------------------
Total proteins analyzed: 9,029
Results breakdown:
   • Unexplained: 7,752 (85.9%)
   • Equivalent: 1,087 (12.0%)
   • Excluded: 158 (1.7%)
   • Upregulated: 20 (0.2%)
   • Downregulated: 12 (0.1%)

💾 SAVING RESULTS to ./data/results/hypoxia/OriginalProtein_72hr_QuEStVar.feather
--------------------------------------------------------------------------------

📈 GENERATING QUESTVAR SUMMARY PLOT
--------------------------------------------------
```

###### 03.4.2 Top-3 Averaged Protein Intensities¶

```
cond_1 = "True Hypoxia (1% O2) - 72 Hr"
cond_2 = "Normoxia (21% O2) - 72 Hr"

print(" RUNNING QUESTVAR STATISTICAL ANALYSIS")
print("=" * 80)
print(f"Parameters: p_thr={p_thr}, cv_thr={cv_thr}, df_thr={df_thr}, eq_thr={eq_thr}")
print(f"Correction: {correction.upper()}, Conditions: {cond_1} vs {cond_2}, Equal Variance: {equal_var}")

res, info = tests.run_questvar(
    S1_arr=protein_top3[condition_to_samples[cond_1]].values,
    S2_arr=protein_top3[condition_to_samples[cond_2]].values,
    is_log2=False,
    cv_thr=cv_thr,
    p_thr=p_thr,
    df_thr=df_thr,
    eq_thr=eq_thr,
    var_equal=equal_var,
    is_paired=False,
    correction=correction,
    allow_missing=True,
)

info = info.set_index( protein_top3.index ).reset_index().set_index('Entry')

info['Status'] = info['Status'].replace({
    0.0: 'Unexplained',
    1.0: 'Equivalent',
    -1.0: 'Different',
}).fillna('Excluded')

quest = info.merge(
    res.iloc[:, :-1], 
    left_index=True, 
    right_index=True, 
    how='left'
)
quest['-log10(Fdr)'] = -np.log10(
    quest['df_adjp']
).replace([np.inf, -np.inf], np.nan)

quest['Status'] = np.where(
    quest['Status'] == 'Different',
    np.where(quest['log2FC'] > 0, 'Upregulated', 'Downregulated'),
    quest['Status']
)

id_col='Protein'
# Find the fasta info and map it, ensuring only one id_col is kept
cols = ['Gene', 'Description', 'Length', 'Weight(kDa)']#, 'Category', 'SPC']
qcols = [col for col in quest.columns if col != id_col]
quest = quest.merge(
    fasta_data[[id_col] + cols],
    left_on=id_col,
    right_on=id_col,
    how='left'
)
# Order the columns, keeping only one id_col
quest = quest[[id_col] + cols + qcols]

print()
print("📊 QUESTVAR ANALYSIS RESULTS")
print("-" * 60)

status_counts = quest['Status'].value_counts()
total_proteins = len(quest)

print(f"Total proteins analyzed: {total_proteins:,}")
print(f"Results breakdown:")
for status, count in status_counts.items():
    percentage = (count/total_proteins)*100
    print(f"   • {status}: {count:,} ({percentage:.1f}%)")
print()

# Add "Equivalent", "Upregulated", "Downregulated" to the dictionary
signf_protein_dict['72hr']["mean(top)"] = {
    'Upregulated': quest.loc[quest['Status'] == 'Upregulated', 'Protein'].unique(),
    'Downregulated': quest.loc[quest['Status'] == 'Downregulated', 'Protein'].unique(),
    'Equivalent': quest.loc[quest['Status'] == 'Equivalent', 'Protein'].unique()
}
# Add thesattus counts to the protein_count_data DataFrame
n_status = len(status_counts)
protein_count_data = pd.concat(
    [
        protein_count_data,
        pd.DataFrame({
            'Method': ['mean(top)'] * n_status,
            'Duration': ['72hr'] * n_status,
            'Count Type': status_counts.index.tolist(),
            'Count': status_counts.values.tolist()
        })
    ],
    axis=0,
    ignore_index=True
)

# Save results
output_data = f"{output_path}Top3Protein_72hr_QuEStVar.feather"
print(f"💾 SAVING RESULTS to {output_data}")
print("-" * 80)
quest.to_feather(f"{output_data}")
print()

# Generate main summary plot
print("📈 GENERATING QUESTVAR SUMMARY PLOT")
print("-" * 50)

plots.questvar_test_summary(
    quest_data=quest,
    p_thr=p_thr,
    eq_thr=eq_thr,
    df_thr=df_thr,
    cv_thr=cv_thr,
    correction=correction,
    cond_1=cond_1,
    cond_2=cond_2,
    figsize=(20, 15),
    title_add=f"Mean of Top3 Protein Data: {cond_1} vs {cond_2}",
    show=True,
    save=save_to_folder,
    filename=f"QuEStVar_Summary_Top3Protein_72hr_Hypoxia_vs_Normoxia",
    fileformats=figure_formats,
    filepath=figure_path,
    transparent=transparent_bg,
    dpi=figure_dpi
)
```

```
 RUNNING QUESTVAR STATISTICAL ANALYSIS
================================================================================
Parameters: p_thr=0.05, cv_thr=1, df_thr=0.5, eq_thr=0.25
Correction: FDR, Conditions: True Hypoxia (1% O2) - 72 Hr vs Normoxia (21% O2) - 72 Hr, Equal Variance: False

📊 QUESTVAR ANALYSIS RESULTS
------------------------------------------------------------
Total proteins analyzed: 7,161
Results breakdown:
   • Unexplained: 6,320 (88.3%)
   • Equivalent: 708 (9.9%)
   • Upregulated: 70 (1.0%)
   • Downregulated: 52 (0.7%)
   • Excluded: 11 (0.2%)

💾 SAVING RESULTS to ./data/results/hypoxia/Top3Protein_72hr_QuEStVar.feather
--------------------------------------------------------------------------------

📈 GENERATING QUESTVAR SUMMARY PLOT
--------------------------------------------------
```

###### 03.4.3 All Peptides Averaged Protein Intensities¶

```
cond_1 = "True Hypoxia (1% O2) - 72 Hr"
cond_2 = "Normoxia (21% O2) - 72 Hr"

print(" RUNNING QUESTVAR STATISTICAL ANALYSIS")
print("=" * 80)
print(f"Parameters: p_thr={p_thr}, cv_thr={cv_thr}, df_thr={df_thr}, eq_thr={eq_thr}")
print(f"Correction: {correction.upper()}, Conditions: {cond_1} vs {cond_2}, Equal Variance: {equal_var}")

res, info = tests.run_questvar(
    S1_arr=protein_all[condition_to_samples[cond_1]].values,
    S2_arr=protein_all[condition_to_samples[cond_2]].values,
    is_log2=False,
    cv_thr=cv_thr,
    p_thr=p_thr,
    df_thr=df_thr,
    eq_thr=eq_thr,
    var_equal=equal_var,
    is_paired=False,
    correction=correction,
    allow_missing=True,
)

info = info.set_index( protein_all.index ).reset_index().set_index('Entry')

info['Status'] = info['Status'].replace({
    0.0: 'Unexplained',
    1.0: 'Equivalent',
    -1.0: 'Different',
}).fillna('Excluded')

quest = info.merge(
    res.iloc[:, :-1], 
    left_index=True, 
    right_index=True, 
    how='left'
)
quest['-log10(Fdr)'] = -np.log10(
    quest['df_adjp']
).replace([np.inf, -np.inf], np.nan)

quest['Status'] = np.where(
    quest['Status'] == 'Different',
    np.where(quest['log2FC'] > 0, 'Upregulated', 'Downregulated'),
    quest['Status']
)

id_col='Protein'
# Find the fasta info and map it, ensuring only one id_col is kept
cols = ['Gene', 'Description', 'Length', 'Weight(kDa)']#, 'Category', 'SPC']
qcols = [col for col in quest.columns if col != id_col]
quest = quest.merge(
    fasta_data[[id_col] + cols],
    left_on=id_col,
    right_on=id_col,
    how='left'
)
# Order the columns, keeping only one id_col
quest = quest[[id_col] + cols + qcols]

print()
print("📊 QUESTVAR ANALYSIS RESULTS")
print("-" * 60)

status_counts = quest['Status'].value_counts()
total_proteins = len(quest)

print(f"Total proteins analyzed: {total_proteins:,}")
print(f"Results breakdown:")
for status, count in status_counts.items():
    percentage = (count/total_proteins)*100
    print(f"   • {status}: {count:,} ({percentage:.1f}%)")
print()

# Add "Equivalent", "Upregulated", "Downregulated" to the dictionary
signf_protein_dict['72hr']["mean(all)"] = {
    'Upregulated': quest.loc[quest['Status'] == 'Upregulated', 'Protein'].unique(),
    'Downregulated': quest.loc[quest['Status'] == 'Downregulated', 'Protein'].unique(),
    'Equivalent': quest.loc[quest['Status'] == 'Equivalent', 'Protein'].unique()
}
# Add thesattus counts to the protein_count_data DataFrame
protein_count_data = pd.concat(
    [
        protein_count_data,
        pd.DataFrame({
            'Method': ['mean(all)'] * 5,
            'Duration': ['72hr'] * 5,
            'Count Type': status_counts.index.tolist(),
            'Count': status_counts.values.tolist()
        })
    ],
    axis=0,
    ignore_index=True
)

# Save results
output_data = f"{output_path}AllProtein_72hr_QuEStVar.feather"
print(f"💾 SAVING RESULTS to {output_data}")
print("-" * 80)
quest.to_feather(f"{output_data}")
print()

# Generate main summary plot
print("📈 GENERATING QUESTVAR SUMMARY PLOT")
print("-" * 50)

plots.questvar_test_summary(
    quest_data=quest,
    p_thr=p_thr,
    eq_thr=eq_thr,
    df_thr=df_thr,
    cv_thr=cv_thr,
    correction=correction,
    cond_1=cond_1,
    cond_2=cond_2,
    figsize=(20, 15),
    title_add=f"Mean of All Peptides - Protein Data: {cond_1} vs {cond_2}",
    show=True,
    save=save_to_folder,
    filename=f"QuEStVar_Summary_AllProtein_72hr_Hypoxia_vs_Normoxia",
    fileformats=figure_formats,
    filepath=figure_path,
    transparent=transparent_bg,
    dpi=figure_dpi
)
```

```
 RUNNING QUESTVAR STATISTICAL ANALYSIS
================================================================================
Parameters: p_thr=0.05, cv_thr=1, df_thr=0.5, eq_thr=0.25
Correction: FDR, Conditions: True Hypoxia (1% O2) - 72 Hr vs Normoxia (21% O2) - 72 Hr, Equal Variance: False

📊 QUESTVAR ANALYSIS RESULTS
------------------------------------------------------------
Total proteins analyzed: 7,161
Results breakdown:
   • Unexplained: 5,463 (76.3%)
   • Equivalent: 1,347 (18.8%)
   • Upregulated: 193 (2.7%)
   • Downregulated: 151 (2.1%)
   • Excluded: 7 (0.1%)

💾 SAVING RESULTS to ./data/results/hypoxia/AllProtein_72hr_QuEStVar.feather
--------------------------------------------------------------------------------

📈 GENERATING QUESTVAR SUMMARY PLOT
--------------------------------------------------
```

###### 03.4.4 dPF-Based Proteoform Intensities¶

```
cond_1 = "True Hypoxia (1% O2) - 72 Hr"
cond_2 = "Normoxia (21% O2) - 72 Hr"

print(" RUNNING QUESTVAR STATISTICAL ANALYSIS")
print("=" * 80)
print(f"Parameters: p_thr={p_thr}, cv_thr={cv_thr}, df_thr={df_thr}, eq_thr={eq_thr}")
print(f"Correction: {correction.upper()}, Conditions: {cond_1} vs {cond_2}, Equal Variance: {equal_var}")

res, info = tests.run_questvar(
    S1_arr=protein_dpf_72[condition_to_samples[cond_1]].values,
    S2_arr=protein_dpf_72[condition_to_samples[cond_2]].values,
    is_log2=False,
    cv_thr=cv_thr,
    p_thr=p_thr,
    df_thr=df_thr,
    eq_thr=eq_thr,
    var_equal=equal_var,
    is_paired=False,
    correction=correction,
    allow_missing=True,
)

info = info.set_index( protein_dpf_72.index ).reset_index().set_index('Entry')

info['Status'] = info['Status'].replace({
    0.0: 'Unexplained',
    1.0: 'Equivalent',
    -1.0: 'Different',
}).fillna('Excluded')

quest = info.merge(
    res.iloc[:, :-1], 
    left_index=True, 
    right_index=True, 
    how='left'
)
quest['-log10(Fdr)'] = -np.log10(
    quest['df_adjp']
).replace([np.inf, -np.inf], np.nan)

quest['Status'] = np.where(
    quest['Status'] == 'Different',
    np.where(quest['log2FC'] > 0, 'Upregulated', 'Downregulated'),
    quest['Status']
)

if 'Proteoform' in quest.columns:
    quest['Protein'] = quest['Proteoform'].str.split('_').str[0]

id_col='Protein'
# Find the fasta info and map it, ensuring only one id_col is kept
cols = ['Gene', 'Description', 'Length', 'Weight(kDa)']#, 'Category', 'SPC']
qcols = [col for col in quest.columns if col != id_col]
quest = quest.merge(
    fasta_data[[id_col] + cols],
    left_on=id_col,
    right_on=id_col,
    how='left'
)
# Order the columns, keeping only one id_col
quest = quest[[id_col] + cols + qcols]

print()
print("📊 QUESTVAR ANALYSIS RESULTS")
print("-" * 60)

status_counts = quest['Status'].value_counts()
total_proteins = len(quest)

print(f"Total proteins analyzed: {total_proteins:,}")
print(f"Results breakdown:")
for status, count in status_counts.items():
    percentage = (count/total_proteins)*100
    print(f"   • {status}: {count:,} ({percentage:.1f}%)")
print()

# Add "Equivalent", "Upregulated", "Downregulated" to the dictionary
signf_protein_dict['72hr']["dPFs"] = {
    'Upregulated': quest.loc[quest['Status'] == 'Upregulated', 'Protein'].unique(),
    'Downregulated': quest.loc[quest['Status'] == 'Downregulated', 'Protein'].unique(),
    'Equivalent': quest.loc[quest['Status'] == 'Equivalent', 'Protein'].unique()
}
# Add thesattus counts to the protein_count_data DataFrame
protein_count_data = pd.concat(
    [
        protein_count_data,
        pd.DataFrame({
            'Method': ['dPFs'] * 5,
            'Duration': ['72hr'] * 5,
            'Count Type': status_counts.index.tolist(),
            'Count': status_counts.values.tolist()
        })
    ],
    axis=0,
    ignore_index=True
)

# Save results
output_data = f"{output_path}dPFProtein_72hr_QuEStVar.feather"
print(f"💾 SAVING RESULTS to {output_data}")
print("-" * 80)
quest.to_feather(f"{output_data}")
print()

# Generate main summary plot
print("📈 GENERATING QUESTVAR SUMMARY PLOT")
print("-" * 50)

plots.questvar_test_summary(
    quest_data=quest,
    p_thr=p_thr,
    eq_thr=eq_thr,
    df_thr=df_thr,
    cv_thr=cv_thr,
    correction=correction,
    cond_1=cond_1,
    cond_2=cond_2,
    figsize=(20, 15),
    title_add=f"dPFs -  Protein Data: {cond_1} vs {cond_2}",
    show=True,
    save=save_to_folder,
    filename=f"QuEStVar_Summary_dPFProtein_72hr_Hypoxia_vs_Normoxia",
    fileformats=figure_formats,
    filepath=figure_path,
    transparent=transparent_bg,
    dpi=figure_dpi
)
```

```
 RUNNING QUESTVAR STATISTICAL ANALYSIS
================================================================================
Parameters: p_thr=0.05, cv_thr=1, df_thr=0.5, eq_thr=0.25
Correction: FDR, Conditions: True Hypoxia (1% O2) - 72 Hr vs Normoxia (21% O2) - 72 Hr, Equal Variance: False

📊 QUESTVAR ANALYSIS RESULTS
------------------------------------------------------------
Total proteins analyzed: 9,175
Results breakdown:
   • Unexplained: 5,791 (63.1%)
   • Equivalent: 1,230 (13.4%)
   • Upregulated: 1,057 (11.5%)
   • Downregulated: 1,039 (11.3%)
   • Excluded: 58 (0.6%)

💾 SAVING RESULTS to ./data/results/hypoxia/dPFProtein_72hr_QuEStVar.feather
--------------------------------------------------------------------------------

📈 GENERATING QUESTVAR SUMMARY PLOT
--------------------------------------------------
```

##### 03.5 Significant Protein Status Comparison¶

Visualize and compare the distribution of protein statuses (Upregulated, Downregulated, Equivalent, Unexplained, Excluded) across quantification methods and timepoints.

```
# --- Data Preparation (Assuming 'protein_count_data' is defined) ---
# This part is the same as your original code.
plot_data = protein_count_data[protein_count_data["Count Type"] != "All Proteins"].copy()
group_totals = plot_data.groupby(["Method", "Duration"])["Count"].sum().reset_index().rename(columns={"Count": "Total"})
plot_data = plot_data.merge(group_totals, on=["Method", "Duration"])
plot_data["Percent"] = plot_data["Count"] / plot_data["Total"] * 100


# --- Enforce Categorical Order ---
# This ensures plotting and sorting functions respect your desired order.
plot_data['Method'] = pd.Categorical(plot_data['Method'], categories=method_order, ordered=True)
plot_data['Count Type'] = pd.Categorical(plot_data['Count Type'], categories=count_type_order, ordered=True)
plot_data = plot_data.sort_values(['Method', 'Count Type'])


# --- Plotting ---
durations = sorted(plot_data["Duration"].unique())
fig, axes = plt.subplots(1, len(durations), figsize=(12, 6), sharey=True, squeeze=False)
axes = axes.flatten() # Ensure axes is always a flat array

for i, duration in enumerate(durations):
    ax = axes[i]
    data_dur = plot_data[plot_data["Duration"] == duration]

    # Pivot data for easier plotting
    df_pivot = data_dur.pivot(index='Method', columns='Count Type', values='Percent')

    # Create the stacked bar plot
    df_pivot.plot(
        kind='bar',
        stacked=True,
        ax=ax,
        color=[status_colors.get(col, '#000000') for col in df_pivot.columns],
        width=0.8,
        edgecolor="white",
        linewidth=0.5
    )

    # --- Add Annotations ---s
    for container in ax.containers:
        for bar, label in zip(container, container.datavalues):
            percent = label
            # Only annotate if value is large enough
            if percent > 4:
                # Get the status (Count Type) for this bar
                count_type = bar.get_label() if hasattr(bar, 'get_label') else None
                # Fallback: use the color of the bar to decide text color
                bar_color = bar.get_facecolor()
                # Convert RGBA to perceived brightness
                brightness = (0.299 * bar_color[0] + 0.587 * bar_color[1] + 0.114 * bar_color[2])
                text_color = 'black' if brightness > 0.6 else 'white'
                # Try to use status_colors for text color if possible
                if count_type in status_colors:
                    # Use white text for dark bars, black for light bars
                    hex_color = status_colors[count_type].lstrip('#')
                    r, g, b = tuple(int(hex_color[i:i+2], 16)/255. for i in (0, 2, 4))
                    brightness = (0.299 * r + 0.587 * g + 0.114 * b)
                    text_color = 'black' if brightness > 0.6 else 'white'
                ax.bar_label(
                    container,
                    labels=[f'{percent:.0f}%' if percent > 4 else '' for percent in container.datavalues],
                    label_type='center',
                    color=text_color,
                    fontsize=12,
                    fontweight='bold'
                )
                break  # Only need to annotate once per bar group

    # --- Styling ---
    ax.set_title(f"Duration: {duration}", fontsize=14, fontweight='bold')
    ax.set_xlabel("") # X-label is shared, set below
    ax.tick_params(axis='x', labelrotation=0, labelsize=12)
    ax.spines['top'].set_visible(False)
    ax.spines['right'].set_visible(False)
    ax.yaxis.grid(True, linestyle='--', linewidth=0.6, color='gray', alpha=0.4)
    ax.set_axisbelow(True)
    ax.get_legend().remove() # Remove individual legends

# --- Final Figure-Level Adjustments ---
# Set common labels
fig.text(0.5, 0.02, 'Method', ha='center', va='center', fontsize=14, fontweight='bold')
axes[0].set_ylabel("Percentage (%)", fontsize=14, fontweight='bold')
axes[0].tick_params(axis='y', labelsize=12)
axes[0].set_ylim(0, 100)

# Create a single, shared legend
handles, labels = axes[0].get_legend_handles_labels()
fig.legend(
    handles,
    labels,
    title="Protein Status",
    loc='upper left',
    bbox_to_anchor=(.90, 0.9),
    fontsize=12,
    title_fontsize=13,
    frameon=False
)

fig.suptitle('Breakdown of Protein Status by Method and Duration', fontsize=18, fontweight='bold')

# Adjust layout to prevent overlap and make space for legend
plt.tight_layout(rect=[0, 0.04, 0.9, 0.95])
plots.finalize_plot(
    fig=fig,
    show=True,
    save=save_to_folder,
    filename="Protein_Status_Breakdown",
    filepath=figure_path,
    formats=figure_formats, 
    transparent=transparent_bg,
    dpi=figure_dpi,
)
```

While each summary plot for each QuEStVar done can be summarized further by comparing the counts of proteins in each status category across methods and timepoints. This allows us to see how proteoform-aware quantification influences the overall classification of protein regulation.

From the plot the `dPFs` method from ProteoForge immediately stands out and shown to be making more of the proteins statistically explainable compared to other methods. Keeping the proportion of statistically equivalent similar as the other methods and even more than `Original` and `mean(top)` methods while increasing the proportion of up/down regulated proteins detected. While decreasing the statistically unexplained proteins proportion to lowest among all methods.

This highlights the advantage of proteoform-aware quantification in revealing biologically relevant regulation that may be missed by traditional protein-level approaches.

---

#### 04. Individual Protein Case Studies¶

Here we generate detailed boxplots for individual proteins to visualize intensity distributions across all quantification methods, conditions, and timepoints. This allows examination of specific cases where dPF-based analysis reveals different regulatory patterns than traditional methods.

```
def melt_protein_df(df, method, id_col='Protein', extra_assign=None):
    melted = df.reset_index().melt(
        id_vars=id_col,
        var_name='Sample',
        value_name='Intensity'
    )
    melted['Method'] = method
    melted['log10Intensity'] = np.log10(melted['Intensity'])
    melted['Condition'] = np.where(melted['Sample'].str.contains('1%'), 'Hypoxia (1% O2)', 'Normoxia (21% O2)')
    melted['Duration'] = np.where(melted['Sample'].str.contains('48'), '48hr', '72hr')
    melted['Proteoform'] = melted[id_col]
    if extra_assign:
        for k, v in extra_assign.items():
            melted[k] = v(melted)
    return melted

# Main protein-level methods
plot_data = pd.concat([
    melt_protein_df(protein_top3, 'mean(top)'),
    melt_protein_df(protein_all, 'mean(all)'),
    melt_protein_df(original_protein, 'Original')
], ignore_index=True)

# protein from dPFs for 48hr and 72hr
for dpf_df, duration in [(protein_dpf_48, '48hr'), (protein_dpf_72, '72hr')]:
    melted = dpf_df.reset_index().melt(
        id_vars='Proteoform',
        var_name='Sample',
        value_name='Intensity'
    )
    # Split Proteoform into Protein and dPF number
    melted[['Protein', 'dPF_num']] = melted['Proteoform'].str.rsplit('_', n=1, expand=True)
    # Set Method as dPF_<dPF_num>
    melted['Method'] = 'dPF_' + melted['dPF_num']
    melted['log10Intensity'] = np.log10(melted['Intensity'])
    melted['Condition'] = np.where(melted['Sample'].str.contains('1%'), 'Hypoxia (1% O2)', 'Normoxia (21% O2)')
    melted['Duration'] = duration
    melted = melted.drop(columns=['dPF_num'])
    plot_data = pd.concat([plot_data, melted], ignore_index=True)

# quest_info_to place in the plot_data
cols_to_pick = ['Protein', 'Gene', 'Description', 'average', 'log2FC', 'comb_adjp', 'Status']
# quest = pd.read_feather(f"{output_path}AllProtein_48hr_QuEStVar.feather")
# OriginalProtein, Top3Protein, AllProtein, dPFProtein
# 48hr and 72hr
name_method_map = {
    'Original': 'OriginalProtein',
    'mean(top)': 'Top3Protein',
    'mean(all)': 'AllProtein',
}
quest_data = pd.DataFrame()
for method, file_suffix in name_method_map.items():
    for duration in ['48hr', '72hr']:
        file_path = f"{output_path}{file_suffix}_{duration}_QuEStVar.feather"
        if os.path.exists(file_path):
            temp_quest = pd.read_feather(file_path)[cols_to_pick]
            temp_quest['Method'] = method
            temp_quest['Duration'] = duration
            temp_quest['Proteoform'] = temp_quest['Protein']  # For consistency
            quest_data = pd.concat([quest_data, temp_quest], ignore_index=True)

# Add the dPFs data
for duration in ['48hr', '72hr']:
    file_path = f"{output_path}dPFProtein_{duration}_QuEStVar.feather"
    if os.path.exists(file_path):
        temp_quest = pd.read_feather(file_path)[cols_to_pick + ['Proteoform']]
        # temp_quest['Method'] = 'dPFs'
        temp_quest['Method'] = temp_quest['Proteoform'].str.split('_').str[1].apply(lambda x: 'dPF_' + x)
        temp_quest['Duration'] = duration
        quest_data = pd.concat([quest_data, temp_quest], ignore_index=True)

# Remove Protein column to avoid confusion
quest_data = quest_data.drop(columns=['Protein'])
# Merge quest_data into plot_data
plot_data = plot_data.merge(
    quest_data,
    on=['Proteoform', 'Method', 'Duration'],
    how='left',
)[[
    'Protein', 'Proteoform', 'Gene', 'Description', 
    'Method', 'Duration', 'Sample', 
    'Intensity', 'log10Intensity', 
    'Condition', 'Status', 'average', 'log2FC', 'comb_adjp'
]]

plot_data
```

|  | Protein | Proteoform | Gene | Description | Method | Duration | Sample | Intensity | log10Intensity | Condition | Status | average | log2FC | comb\_adjp |
| --- | --- | --- | --- | --- | --- | --- | --- | --- | --- | --- | --- | --- | --- | --- |
| 0 | A0A024RBG1 | A0A024RBG1 | NUDT4B | Diphosphoinositol polyphosphate phosphohydrola... | mean(top) | 48hr | 1%\_48\_1 | 42434.0877 | 4.6277 | Hypoxia (1% O2) | Equivalent | 15.4190 | -0.0542 | 0.0403 |
| 1 | A0A0B4J2D5 | A0A0B4J2D5 | GATD3B | Putative glutamine amidotransferase-like class... | mean(top) | 48hr | 1%\_48\_1 | 102861.7619 | 5.0123 | Hypoxia (1% O2) | Unexplained | 16.6117 | 0.0899 | 0.1765 |
| 2 | A0A1B0GTU1 | A0A1B0GTU1 | ZC3H11B | Zinc finger CCCH domain-containing protein 11B | mean(top) | 48hr | 1%\_48\_1 | 15418.3217 | 4.1880 | Hypoxia (1% O2) | Unexplained | 13.9667 | 0.0164 | 0.1247 |
| 3 | A0A1W2PQL4 | A0A1W2PQL4 | ZNF722 | Zinc finger protein 722 | mean(top) | 48hr | 1%\_48\_1 | 138883.8369 | 5.1427 | Hypoxia (1% O2) | Unexplained | 17.1534 | -0.3106 | 0.4431 |
| 4 | A0A2R8Y4L2 | A0A2R8Y4L2 | HNRNPA1L3 | Heterogeneous nuclear ribonucleoprotein A1-like 3 | mean(top) | 48hr | 1%\_48\_1 | 2516111.9537 | 6.4007 | Hypoxia (1% O2) | Equivalent | 21.0464 | -0.0062 | 0.0270 |
| ... | ... | ... | ... | ... | ... | ... | ... | ... | ... | ... | ... | ... | ... | ... |
| 444815 | Q9Y6X9 | Q9Y6X9\_1 | MORC2 | ATPase MORC2 | dPF\_1 | 72hr | C\_72\_3 | 64.7699 | 1.8114 | Normoxia (21% O2) | Upregulated | 8.8591 | 7.8241 | 0.0109 |
| 444816 | Q9Y6X9 | Q9Y6X9\_2 | MORC2 | ATPase MORC2 | dPF\_2 | 72hr | C\_72\_3 | 4496.2079 | 3.6528 | Normoxia (21% O2) | Downregulated | 8.3074 | -7.6085 | 0.0070 |
| 444817 | Q9Y6Y0 | Q9Y6Y0\_0 | IVNS1ABP | Influenza virus NS1A-binding protein | dPF\_0 | 72hr | C\_72\_3 | 9870.7399 | 3.9943 | Normoxia (21% O2) | Unexplained | 13.1397 | -0.2626 | 0.1456 |
| 444818 | Q9Y6Y8 | Q9Y6Y8\_0 | SEC23IP | SEC23-interacting protein | dPF\_0 | 72hr | C\_72\_3 | 8050.0895 | 3.9058 | Normoxia (21% O2) | Unexplained | 12.9684 | -0.1083 | 0.0619 |
| 444819 | Q9Y6Y8 | Q9Y6Y8\_1 | SEC23IP | SEC23-interacting protein | dPF\_1 | 72hr | C\_72\_3 | 3633.8751 | 3.5604 | Normoxia (21% O2) | Downregulated | 8.1646 | -7.6285 | 0.0010 |

444820 rows × 14 columns

```
all_methods_order = plot_data['Method'].unique().tolist()
all_methods_order = ['Original', 'mean(top)', 'mean(all)'] + sorted([m for m in all_methods_order if m.startswith('dPF_')], key=lambda x: int(x.split('_')[1]))

summary = plot_data.pivot_table(
    columns=['Method', 'Duration'],
    index='Protein',
    values='Status',
    aggfunc='first'
).fillna('No Data')
# Order the Method by method_order
summary = summary.reindex(columns=pd.MultiIndex.from_product([all_methods_order, ['48hr', '72hr']]), fill_value='No Data')
summary['Total Methods'] = (summary != 'No Data').sum(axis=1)
summary = summary.sort_values(['Total Methods'] + [(method, dur) for method in all_methods_order for dur in ['48hr', '72hr']], ascending=False)
summary = summary.reset_index()
# Rulests
# Remove proteins with Excluded
summary = summary[~summary.isin(['Excluded']).any(axis=1)]
# Fast vectorized version for consistency check
# 1. Get base statuses for Original, mean(top), mean(all) for both durations
base_methods = ['Original', 'mean(top)', 'mean(all)']
dpf_methods = [m for m in all_methods_order if m.startswith('dPF_')]
durations = ['48hr', '72hr']

# Get the columns for base and dPFs
base_cols = [(m, d) for m in base_methods for d in durations]
dpf_cols = [(m, d) for m in dpf_methods for d in durations]

# Convert to numpy for fast access
summary_arr = summary[[col for col in summary.columns if col not in ['Protein', 'Total Methods']]].values

# Map column index for each method/duration
col_idx = {col: i for i, col in enumerate(summary.columns) if col not in ['Protein', 'Total Methods']}

# Precompute indices
base_indices = [col_idx[c] for c in base_cols]
dpf_indices = [col_idx[c] for c in dpf_cols]

# Get base status for each row (if consistent and not 'No Data')
def get_base_status(row):
    vals = row[base_indices]
    vals = vals[vals != 'No Data']
    if len(vals) == 0:
        return None
    if np.all(vals == vals[0]):
        return vals[0]
    return None

base_statuses = np.apply_along_axis(get_base_status, 1, summary_arr)

# Get dPF statuses for each row
dpf_statuses = summary_arr[:, dpf_indices]

# For each row, check if any dPF status is different from base_status (and not 'No Data')
mask = []
for i, base_status in enumerate(base_statuses):
    if base_status is None or base_status == 'No Data':
        mask.append(False)
        continue
    dpf_vals = dpf_statuses[i]
    dpf_vals = dpf_vals[dpf_vals != 'No Data']
    if len(dpf_vals) == 0:
        mask.append(False)
    else:
        mask.append(np.any(dpf_vals != base_status))
summary = summary[np.array(mask)]
# If Original has No Data, don't include
summary = summary[
    (summary[('Original', '48hr')] != 'No Data') &
    (summary[('Original', '72hr')] != 'No Data')
]
summary.head(25)
```

|  | Protein | Original | | mean(top) | | mean(all) | | dPF\_0 | | dPF\_1 | | dPF\_2 | | dPF\_3 | | Total Methods |
| --- | --- | --- | --- | --- | --- | --- | --- | --- | --- | --- | --- | --- | --- | --- | --- | --- |
|  |  | 48hr | 72hr | 48hr | 72hr | 48hr | 72hr | 48hr | 72hr | 48hr | 72hr | 48hr | 72hr | 48hr | 72hr |  |
| 1 | P43378 | Unexplained | Unexplained | Unexplained | Unexplained | Unexplained | Unexplained | Unexplained | Unexplained | Downregulated | Upregulated | Upregulated | Downregulated | No Data | No Data | 12 |
| 2 | Q9UPQ0 | Unexplained | Unexplained | Unexplained | Unexplained | Unexplained | Unexplained | Unexplained | Unexplained | Downregulated | Downregulated | Upregulated | Upregulated | No Data | No Data | 12 |
| 3 | Q05481 | Unexplained | Unexplained | Unexplained | Unexplained | Unexplained | Unexplained | Unexplained | Unexplained | Downregulated | Downregulated | Unexplained | Upregulated | No Data | No Data | 12 |
| 4 | Q8WXH0 | Unexplained | Unexplained | Unexplained | Unexplained | Unexplained | Unexplained | Unexplained | No Data | Downregulated | Upregulated | Upregulated | Downregulated | No Data | Equivalent | 12 |
| 5 | Q15149 | Unexplained | Unexplained | Unexplained | Unexplained | Unexplained | Unexplained | No Data | No Data | Upregulated | Upregulated | Downregulated | Downregulated | Unexplained | Unexplained | 12 |
| 6 | Q9UPN3 | Unexplained | Unexplained | Unexplained | Unexplained | Unexplained | Unexplained | No Data | No Data | Downregulated | Upregulated | Upregulated | Downregulated | Unexplained | Equivalent | 12 |
| 7 | Q14789 | Unexplained | Unexplained | Unexplained | Unexplained | Unexplained | Unexplained | No Data | No Data | Downregulated | Downregulated | Unexplained | Upregulated | Upregulated | Unexplained | 12 |
| 19 | P50851 | Equivalent | Equivalent | Equivalent | Equivalent | Equivalent | Equivalent | Equivalent | Equivalent | Upregulated | Upregulated | Downregulated | Downregulated | No Data | No Data | 12 |
| 24 | Q13442 | Unexplained | Unexplained | Unexplained | Unexplained | Unexplained | Unexplained | Unexplained | Unexplained | Upregulated | Upregulated | Downregulated | No Data | No Data | No Data | 11 |
| 25 | Q9BV73 | Unexplained | Unexplained | Unexplained | Unexplained | Unexplained | Unexplained | Unexplained | Unexplained | Upregulated | Downregulated | No Data | Upregulated | No Data | No Data | 11 |
| 26 | P02769 | Unexplained | Unexplained | Unexplained | Unexplained | Unexplained | Unexplained | Unexplained | Unexplained | Unexplained | Downregulated | No Data | Upregulated | No Data | No Data | 11 |
| 27 | Q9UNY4 | Unexplained | Unexplained | Unexplained | Unexplained | Unexplained | Unexplained | Unexplained | Unexplained | Unexplained | Downregulated | No Data | Upregulated | No Data | No Data | 11 |
| 29 | Q8IZT6 | Unexplained | Unexplained | Unexplained | Unexplained | Unexplained | Unexplained | Unexplained | Unexplained | Downregulated | Upregulated | No Data | Downregulated | No Data | No Data | 11 |
| 30 | Q14562 | Unexplained | Unexplained | Unexplained | Unexplained | Unexplained | Unexplained | Unexplained | Unexplained | Downregulated | Downregulated | No Data | Upregulated | No Data | No Data | 11 |
| 31 | Q14997 | Unexplained | Unexplained | Unexplained | Unexplained | Unexplained | Unexplained | Unexplained | Unexplained | Downregulated | Downregulated | No Data | Upregulated | No Data | No Data | 11 |
| 32 | Q96SN8 | Unexplained | Unexplained | Unexplained | Unexplained | Unexplained | Unexplained | Unexplained | Unexplained | Downregulated | Downregulated | No Data | Upregulated | No Data | No Data | 11 |
| 33 | Q9NRR4 | Unexplained | Unexplained | Unexplained | Unexplained | Unexplained | Unexplained | Unexplained | Unexplained | Downregulated | Downregulated | No Data | Upregulated | No Data | No Data | 11 |
| 35 | O94854 | Unexplained | Unexplained | Unexplained | Unexplained | Unexplained | Unexplained | Unexplained | No Data | Upregulated | Upregulated | Downregulated | Equivalent | No Data | No Data | 11 |
| 36 | O60437 | Unexplained | Unexplained | Unexplained | Unexplained | Unexplained | Unexplained | Unexplained | No Data | Unexplained | Downregulated | No Data | Upregulated | No Data | Unexplained | 11 |
| 37 | Q02224 | Unexplained | Unexplained | Unexplained | Unexplained | Unexplained | Unexplained | Unexplained | No Data | Downregulated | Upregulated | No Data | Downregulated | No Data | Downregulated | 11 |
| 38 | Q96HP0 | Unexplained | Unexplained | Unexplained | Unexplained | Unexplained | Unexplained | Unexplained | No Data | Downregulated | Downregulated | No Data | Upregulated | No Data | Unexplained | 11 |
| 39 | O75128 | Unexplained | Unexplained | Unexplained | Unexplained | Unexplained | Unexplained | Unexplained | Equivalent | Downregulated | Upregulated | No Data | Downregulated | No Data | No Data | 11 |
| 40 | O75153 | Unexplained | Unexplained | Unexplained | Unexplained | Unexplained | Unexplained | Unexplained | Downregulated | Unexplained | Upregulated | No Data | Unexplained | No Data | No Data | 11 |
| 41 | Q13085 | Unexplained | Unexplained | Unexplained | Unexplained | Unexplained | Unexplained | Unexplained | Downregulated | Unexplained | Upregulated | No Data | Downregulated | No Data | No Data | 11 |
| 42 | O15091 | Unexplained | Unexplained | Unexplained | Unexplained | Unexplained | Unexplained | Unexplained | Downregulated | Downregulated | Upregulated | No Data | Downregulated | No Data | No Data | 11 |

```
is_demo = True # Set to False to save plot
current_protein = 'P50851' #LRBA an example protein..
# current_protein = "Q04760"

subset_plot = plot_data[plot_data['Protein'] == current_protein]
# Define the order of methods for consistent plotting
dPF_methods = sorted(subset_plot['Method'].unique())
dPF_methods = [m for m in dPF_methods if m.startswith('dPF_')]
main_methods = ['Original', 'mean(top)', 'mean(all)']
method_order = main_methods + dPF_methods
method_order
# 
# Set categories with the defined order
subset_plot['Method'] = pd.Categorical(subset_plot['Method'], categories=method_order, ordered=True)
subset_plot['Duration'] = pd.Categorical(subset_plot['Duration'], categories=['48hr', '72hr'], ordered=True)
subset_plot['Condition'] = pd.Categorical(subset_plot['Condition'], categories=['Normoxia (21% O2)', 'Hypoxia (1% O2)'], ordered=True)
# Sort the DataFrame based on the categorical order
subset_plot = subset_plot.sort_values(['Method', 'Duration', 'Condition', 'Proteoform']).reset_index(drop=True)
# subset_plot.to_csv("debug_subset_plot.csv", index=False)
subset_plot
```

|  | Protein | Proteoform | Gene | Description | Method | Duration | Sample | Intensity | log10Intensity | Condition | Status | average | log2FC | comb\_adjp |
| --- | --- | --- | --- | --- | --- | --- | --- | --- | --- | --- | --- | --- | --- | --- |
| 0 | P50851 | P50851 | LRBA | Lipopolysaccharide-responsive and beige-like a... | Original | 48hr | C\_48\_1 | 153304.3858 | 5.1856 | Normoxia (21% O2) | Equivalent | 17.2353 | 0.0167 | 0.0198 |
| 1 | P50851 | P50851 | LRBA | Lipopolysaccharide-responsive and beige-like a... | Original | 48hr | C\_48\_2 | 166143.9163 | 5.2205 | Normoxia (21% O2) | Equivalent | 17.2353 | 0.0167 | 0.0198 |
| 2 | P50851 | P50851 | LRBA | Lipopolysaccharide-responsive and beige-like a... | Original | 48hr | C\_48\_3 | 156653.4033 | 5.1949 | Normoxia (21% O2) | Equivalent | 17.2353 | 0.0167 | 0.0198 |
| 3 | P50851 | P50851 | LRBA | Lipopolysaccharide-responsive and beige-like a... | Original | 48hr | C\_48\_4 | 138785.0132 | 5.1423 | Normoxia (21% O2) | Equivalent | 17.2353 | 0.0167 | 0.0198 |
| 4 | P50851 | P50851 | LRBA | Lipopolysaccharide-responsive and beige-like a... | Original | 48hr | 1%\_48\_1 | 156376.3783 | 5.1942 | Hypoxia (1% O2) | Equivalent | 17.2353 | 0.0167 | 0.0198 |
| ... | ... | ... | ... | ... | ... | ... | ... | ... | ... | ... | ... | ... | ... | ... |
| 79 | P50851 | P50851\_2 | LRBA | Lipopolysaccharide-responsive and beige-like a... | dPF\_2 | 72hr | C\_72\_2 | 2089.7656 | 3.3201 | Normoxia (21% O2) | Downregulated | 7.9001 | -6.9301 | 0.0022 |
| 80 | P50851 | P50851\_2 | LRBA | Lipopolysaccharide-responsive and beige-like a... | dPF\_2 | 72hr | C\_72\_3 | 3073.3134 | 3.4876 | Normoxia (21% O2) | Downregulated | 7.9001 | -6.9301 | 0.0022 |
| 81 | P50851 | P50851\_2 | LRBA | Lipopolysaccharide-responsive and beige-like a... | dPF\_2 | 72hr | 1%\_72\_2 | 22.5601 | 1.3533 | Hypoxia (1% O2) | Downregulated | 7.9001 | -6.9301 | 0.0022 |
| 82 | P50851 | P50851\_2 | LRBA | Lipopolysaccharide-responsive and beige-like a... | dPF\_2 | 72hr | 1%\_72\_3 | 21.9215 | 1.3409 | Hypoxia (1% O2) | Downregulated | 7.9001 | -6.9301 | 0.0022 |
| 83 | P50851 | P50851\_2 | LRBA | Lipopolysaccharide-responsive and beige-like a... | dPF\_2 | 72hr | 1%\_72\_4 | 20.4675 | 1.3111 | Hypoxia (1% O2) | Downregulated | 7.9001 | -6.9301 | 0.0022 |

84 rows × 14 columns

```
from matplotlib.lines import Line2D
from matplotlib.patches import Patch, Rectangle

# Get current gene and description
try:
    current_gene = subset_plot['Gene'].dropna().unique()[0]
    current_description = subset_plot['Description'].dropna().unique()[0]
except (IndexError, KeyError):
    current_gene = "Unknown Gene"
    current_description = "No description available"

# --- Plot Setup ---
yOffset = 0.35
ymin = subset_plot['log10Intensity'].min() - yOffset
ymax = subset_plot['log10Intensity'].max() + yOffset

# Calculate marker y position and extended ylim
status_marker_y = ymax + 0.13 * (ymax - ymin)
ylim_top = status_marker_y + 0.08 * (ymax - ymin)

fig, ax = plt.subplots(figsize=(16, 6))
fig.patch.set_facecolor('white')

xtick_positions = np.arange(len(method_order))
xtick_labels = method_order

total_width = 0.8
box_width = total_width / 6
spacing = total_width / 12

boxplot_offsets = {
    ('48hr', 'Normoxia (21% O2)'): -total_width/2 + box_width/2,
    ('48hr', 'Hypoxia (1% O2)'): -total_width/2 + box_width/2 + box_width + spacing,
    ('72hr', 'Normoxia (21% O2)'): -total_width/2 + box_width/2 + 2*(box_width + spacing),
    ('72hr', 'Hypoxia (1% O2)'): -total_width/2 + box_width/2 + 3*(box_width + spacing)
}
duration_center_offsets = {'48hr': -0.2, '72hr': 0.2}
jitter_amount = 0.055

# --- Background rectangles for Duration (full ylim) ---
for i, method in enumerate(method_order):
    for duration in ['48hr', '72hr']:
        offset_norm = boxplot_offsets[(duration, 'Normoxia (21% O2)')]
        offset_hyp = boxplot_offsets[(duration, 'Hypoxia (1% O2)')]
        rect_x_start = xtick_positions[i] + min(offset_norm, offset_hyp) - box_width/2
        rect_x_end = xtick_positions[i] + max(offset_norm, offset_hyp) + box_width/2
        rect_width = rect_x_end - rect_x_start
        rect_height = ylim_top - ymin
        rect = Rectangle(
            (rect_x_start, ymin), rect_width, rect_height,
            facecolor=dura_colors[duration], 
            alpha=0.15, 
            edgecolor='none',
            zorder=1
        )
        ax.add_patch(rect)

# --- Skeleton boxplots ---
for i, method in enumerate(method_order):
    for (duration, condition), offset in boxplot_offsets.items():
        data_subset = subset_plot[
            (subset_plot['Method'] == method) & 
            (subset_plot['Duration'] == duration) & 
            (subset_plot['Condition'] == condition)
        ]
        if data_subset.empty:
            continue
        x_center = xtick_positions[i] + offset
        intensities = data_subset['log10Intensity'].dropna()
        if len(intensities) > 0:
            q1 = intensities.quantile(0.25)
            median = intensities.median()
            q3 = intensities.quantile(0.75)
            iqr = q3 - q1
            lower_whisker = max(intensities.min(), q1 - 1.5 * iqr)
            upper_whisker = min(intensities.max(), q3 + 1.5 * iqr)
            box_x = x_center - box_width/2
            box_rect = Rectangle(
                (box_x, q1), box_width, q3-q1,
                facecolor='none',  # transparent fill
                edgecolor='#333333',
                linewidth=1.5,
                alpha=0.9,
                zorder=4
            )
            ax.add_patch(box_rect)
            ax.plot([box_x, box_x + box_width], [median, median], 
                    color='#d62728', linewidth=2.5, zorder=5, alpha=0.9)
            if upper_whisker > q3:
                ax.plot([x_center, x_center], [q3, upper_whisker], 
                        color='#333333', linewidth=1.5, zorder=4, alpha=0.8)
                cap_width = box_width * 0.5
                ax.plot([x_center - cap_width/2, x_center + cap_width/2], 
                        [upper_whisker, upper_whisker], color='#333333', linewidth=1.5, zorder=4, alpha=0.8)
            if lower_whisker < q1:
                ax.plot([x_center, x_center], [q1, lower_whisker], 
                        color='#333333', linewidth=1.5, zorder=4, alpha=0.8)
                cap_width = box_width * 0.5
                ax.plot([x_center - cap_width/2, x_center + cap_width/2], 
                        [lower_whisker, lower_whisker], color='#333333', linewidth=1.5, zorder=4, alpha=0.8)

# --- Jittered points ---
for i, method in enumerate(method_order):
    for (duration, condition), offset in boxplot_offsets.items():
        data_subset = subset_plot[
            (subset_plot['Method'] == method) & 
            (subset_plot['Duration'] == duration) & 
            (subset_plot['Condition'] == condition)
        ]
        if data_subset.empty:
            continue
        x_base = xtick_positions[i] + offset
        color = cond_colors[condition]
        np.random.seed(42)
        x_jittered = x_base + np.random.uniform(-jitter_amount, jitter_amount, size=len(data_subset))
        label = condition if (i == 0 and duration == '48hr') else ""
        ax.scatter(
            x_jittered,
            data_subset['log10Intensity'],
            color=color,
            label=label,
            alpha=0.85,
            edgecolor='#333333',
            linewidth=0.6,
            s=75,
            zorder=6
        )

# --- Connecting lines for same Proteoform ---
for i, method in enumerate(method_order):
    for duration in ['48hr', '72hr']:
        data_subset = subset_plot[(subset_plot['Method'] == method) & (subset_plot['Duration'] == duration)]
        if data_subset.empty:
            continue
        for proteoform in data_subset['Proteoform'].unique():
            proteoform_data = data_subset[data_subset['Proteoform'] == proteoform]
            if len(proteoform_data['Condition'].unique()) == 2:
                sorted_data = proteoform_data.sort_values('Condition')
                y_vals = sorted_data['log10Intensity'].values
                conds = sorted_data['Condition'].values
                x_vals = []
                for cond in conds:
                    offset = boxplot_offsets[(duration, cond)]
                    x_base = xtick_positions[i] + offset
                    x_vals.append(x_base)
                if len(x_vals) == 2 and len(y_vals) == 2:
                    ax.plot(
                        x_vals,
                        y_vals,
                        color='#666666',
                        alpha=0.4,
                        linewidth=1.2,
                        linestyle='-',
                        zorder=3
                    )

# --- Statistical test lines and status markers ---
for i, method in enumerate(method_order):
    for duration in ['48hr', '72hr']:
        offset_norm = boxplot_offsets[(duration, 'Normoxia (21% O2)')]
        offset_hyp = boxplot_offsets[(duration, 'Hypoxia (1% O2)')]
        x_norm = xtick_positions[i] + offset_norm
        x_hyp = xtick_positions[i] + offset_hyp
        ax.plot([x_norm, x_hyp], [status_marker_y, status_marker_y], 
                color='#333333', lw=1.5, zorder=7, alpha=0.8)
        method_duration_data = subset_plot[
            (subset_plot['Method'] == method) & 
            (subset_plot['Duration'] == duration)
        ]
        conds_present = set(method_duration_data['Condition'].unique())
        required_conds = {'Normoxia (21% O2)', 'Hypoxia (1% O2)'}
        if (required_conds.issubset(conds_present) and 
            not method_duration_data.empty and 
            not method_duration_data['Status'].isnull().all()):
            try:
                status = method_duration_data['Status'].mode()[0]
            except IndexError:
                status = ''
        else:
            status = ''
        x_center = (x_norm + x_hyp) / 2
        if status != '':
            marker = status_markers.get(status, 'o')
            color = status_colors.get(status, '#C2C0C0')
            ax.scatter(
                x_center, status_marker_y,
                marker=marker,
                color=color,
                edgecolor='#333333',
                s=220,
                zorder=8,
                linewidth=1.3,
                alpha=0.9
            )

# --- Two-level x-axis labels ---
duration_positions = []
duration_labels = []
for i, method in enumerate(method_order):
    offset_48hr = xtick_positions[i] + duration_center_offsets['48hr']
    offset_72hr = xtick_positions[i] + duration_center_offsets['72hr']
    duration_positions.extend([offset_48hr, offset_72hr])
    duration_labels.extend(['48hr', '72hr'])

ax.set_xticks(duration_positions)
ax.set_xticklabels(duration_labels, fontsize=14, fontweight='bold', color='#444444')
ax.tick_params(axis='x', top=False, bottom=True, length=0, pad=8)

ax2 = ax.twiny()
ax2.set_xlim(ax.get_xlim())
ax2.set_xticks(xtick_positions)
ax2.set_xticklabels(xtick_labels)
for label in ax2.get_xticklabels():
    label.set_fontsize(20)
    label.set_fontweight('bold') 
    label.set_color('#000000')
ax2.xaxis.set_ticks_position('bottom')
ax2.xaxis.set_label_position('bottom')
ax2.spines['bottom'].set_position(('outward', 30))
ax2.tick_params(
    axis='x', top=False, bottom=True, length=0, pad=10,
    labelsize=15, labelcolor='#444444',
    labelrotation=0
)
for label in ax2.get_xticklabels():
    label.set_fontweight('bold')

# --- Axis styling ---
ax.set_ylabel('log10 Intensity', fontsize=16, fontweight='bold', color='#333333', labelpad=10)
ax.set_title(f'{current_gene} ({current_protein}) Protein Intensity Comparison',
             fontsize=19, fontweight='bold', color='#222222', pad=35)
ax.set_ylim(ymin, ylim_top)
ax.yaxis.grid(True, linestyle='--', linewidth=0.8, color='#cccccc', alpha=0.7)
ax.set_axisbelow(True)
ax.tick_params(axis='y', colors='#444444', labelsize=14, length=4, width=1.2)

# --- Legend System ---
condition_handles = [
    Line2D([0], [0], marker='o', color='w', label=cond, 
           markerfacecolor=color, markersize=14, markeredgecolor='#333333', 
           markeredgewidth=0.8, linestyle='None')
    for cond, color in cond_colors.items()
]
duration_handles = [
    Patch(facecolor=dura_colors[d], edgecolor='#666666', 
          label=f'{d}', alpha=0.25, linewidth=0.8)
    for d in ['48hr', '72hr']
]
status_handles = [
    Line2D([0], [0], marker=status_markers[s], color='w', label=f'{s}', 
           markerfacecolor=status_colors[s], markeredgecolor='#333333', 
           markersize=16, linestyle='None', markeredgewidth=1.0)
    for s in status_colors.keys() if s in status_markers
]
all_legend_elements = []
all_legend_labels = []
if condition_handles:
    all_legend_elements.extend(condition_handles)
    all_legend_labels.extend([h.get_label() for h in condition_handles])
if duration_handles:
    all_legend_elements.extend(duration_handles)
    all_legend_labels.extend([h.get_label() for h in duration_handles])
if status_handles:
    all_legend_elements.extend(status_handles)
    all_legend_labels.extend([h.get_label() for h in status_handles])
if all_legend_elements:
    ax.legend(
        all_legend_elements,
        all_legend_labels,
        loc='center left', bbox_to_anchor=(1.05, 0.5),
        fontsize=11, title='Legend', title_fontsize=13, 
        borderpad=1.2, labelspacing=1.1, frameon=False
    )
plt.subplots_adjust(right=0.75)
plots.finalize_plot(
    fig, 
    show=True,
    save=not is_demo,
    filename=f"ProteinIntensity_{current_protein}_{current_gene}",
    filepath=figure_path,
    formats=figure_formats, 
    transparent=transparent_bg,
    dpi=figure_dpi,
)
```

The above is one of the example proteins that can be highlighted here. The figure itself is pretty simple, but what it shows is that the equivalent status assigned by other methods at each timepoint showsn up and down regulated in dPF\_1 and dPF\_2 respectively. Meaning the LRBA protein has two additional proteoforms that are regulated differently under hypoxia vs normoxia conditions at different timepoints, which is missed by traditional protein-level quantification methods.

> Note: These figures can be generate here for other proteins that you want to highlight as well. However since I won't be including these in the manuscript (even though I want to because they look very cool), I am not generating more of these plots here.
>
> Note: These are super customized and don't have a function to generate them automatically due to it being more of a explainer and not something that will be included in the main manuscript.

---

#### 05. Functional Enrichment Analysis¶

Perform Gene Ontology (GO) enrichment analysis using gProfiler to compare the biological interpretations derived from different protein quantification methods. This reveals how method choice affects downstream pathway-level conclusions.

##### 05.1 Setup Enrichment Analysis¶

Configure enrichment parameters and define protein sets for each method-duration-status combination.

The sets of proteins belonging to each statistical status (Upregulated, Downregulated, Equivalent) from each quantification method and timepoint are prepared for GO enrichment analysis. This allows comparison of the biological themes captured by each approach.

> The same parameters will be used for all enrichment analyses to ensure comparability. []

```
# Variables for the enrichment setup
m_pval = 0.1 
e_pval = 0.01  
correction = 'fdr' # fdr, bonferroni, g_SCS
pval_cap = 10**-15 
organism = "hsapiens"
# sources = ['GO:BP', 'GO:MF', 'GO:CC', 'KEGG', 'REAC', 'WP']
sources = ['GO:BP', 'GO:MF', 'GO:CC'] # Limit to Gene Ontology for faster results

# All Available Proteins as background...
background = list(set(original_protein.index).union(set(proteins_used)))

print(f"🔍 RUNNING ENRICHMENT ANALYSIS ON 24 PROTEIN SETS")
print(f"    - Background Proteins: {len(background):,}")
print(f"    - Methods: {', '.join(method_order)}")
print(f"    - Durations: {', '.join(durations)}")
print(f"    - Sources: {', '.join(sources)}")
print(f"    - p-value threshold: {m_pval} (term inclusion), {e_pval} (term reporting)")
print(f"    - Multiple Testing Correction: {correction.upper()}")
print(f"   {'Set Name':<30}   | {'Dur':<5} | {'Method':<10} | {'Status':<15} | {'# Proteins':>10}")
enrichment_sets = {}
# Flatten the dictionary for easier iteration
# Example Naming: '48hr_mean(top)_Upregulated'
for duration in ['48hr', '72hr']:
    for proMethod, protList in signf_protein_dict[duration].items():
        for status, proteins in protList.items():
            # if status == 'Equivalent':
            #     continue
            if proMethod in ['mean(all)', 'mean(top)']:
                continue
            set_name = f"{duration}_{proMethod}_{status}"
            enrichment_sets[set_name] = proteins.tolist()
            # Print with aligned columns for better readability
            print(f"   • {set_name:<30} | {duration:<5} | {proMethod:<10} | {status:<15} |  {len(proteins):>6} proteins")

queries = enrichment_sets.keys()
analysis_name = 'gProfiler_Enrichment_Analysis_AllSets'
# enrich.printParams(
#     m_pval,
#     e_pval,
#     correction,
#     pval_cap,
#     organism,
#     sources,
#     background,
#     enrichment_sets,
#     analysis_name
# )
```

```
🔍 RUNNING ENRICHMENT ANALYSIS ON 24 PROTEIN SETS
    - Background Proteins: 9,294
    - Methods: Original, mean(top), mean(all), dPF_0, dPF_1, dPF_2
    - Durations: 48hr, 72hr
    - Sources: GO:BP, GO:MF, GO:CC
    - p-value threshold: 0.1 (term inclusion), 0.01 (term reporting)
    - Multiple Testing Correction: FDR
   Set Name                         | Dur   | Method     | Status          | # Proteins
   • 48hr_Original_Upregulated      | 48hr  | Original   | Upregulated     |      65 proteins
   • 48hr_Original_Downregulated    | 48hr  | Original   | Downregulated   |      19 proteins
   • 48hr_Original_Equivalent       | 48hr  | Original   | Equivalent      |    1893 proteins
   • 48hr_dPFs_Upregulated          | 48hr  | dPFs       | Upregulated     |     394 proteins
   • 48hr_dPFs_Downregulated        | 48hr  | dPFs       | Downregulated   |     349 proteins
   • 48hr_dPFs_Equivalent           | 48hr  | dPFs       | Equivalent      |    2345 proteins
   • 72hr_Original_Upregulated      | 72hr  | Original   | Upregulated     |      20 proteins
   • 72hr_Original_Downregulated    | 72hr  | Original   | Downregulated   |      12 proteins
   • 72hr_Original_Equivalent       | 72hr  | Original   | Equivalent      |    1087 proteins
   • 72hr_dPFs_Upregulated          | 72hr  | dPFs       | Upregulated     |    1034 proteins
   • 72hr_dPFs_Downregulated        | 72hr  | dPFs       | Downregulated   |    1006 proteins
   • 72hr_dPFs_Equivalent           | 72hr  | dPFs       | Equivalent      |    1230 proteins
```

##### 05.2 Run Enrichment for All Combinations¶

Execute GO enrichment analysis for each combination of quantification method, timepoint, and protein status using gProfiler. Since gProfiler accepts the multi-query format, all analyses are run in a single batch for efficiency and consistency.

> Once it is run (and no parameters changed), the results can be saved and loaded directly to avoid re-running the enrichment each time. The commenting out the first function call to avoid re-running it every time.

```
# Run and Save the Enrichment Analysis
enrich.run_gprofiler(
    query = enrichment_sets,
    background = background,
    organism = organism,
    user_threshold = m_pval,
    signf_threshold = e_pval,
    correction = correction,
    sources = sources,
    pval_cap = pval_cap,
    no_evidences = True,
    no_iea = False,
    ordered = False,
    simplify_cols=True,
    save_path = output_path,
    analysis_name = analysis_name,
    verbose = True
)

# print()

# Read the results
gp_res = pd.read_csv(
    os.path.join(
        output_path, 
        f"{analysis_name}.csv"
    ),
    sep=","
)

# Create a subset of the results with the p-value cutoff
subset_gp_res = gp_res[
    gp_res["p_value"] < e_pval
]
subset_gp_res[['Duration', 'Method', 'Status']] = subset_gp_res['query'].str.split('_', expand=True)

print(f"""After the p-value cutoff of {e_pval} there are total of {subset_gp_res.shape[0]} enriched terms.""")

utils.print_series(
    subset_gp_res["query"].value_counts(),
    header = f"Number of enriched terms per query with p-val cutoff {e_pval}:",
    tab = 2,
    elements_with_order = list(queries)
)
```

```
Capping the p_value at 1e-15
Simplifying the columns.
P-value threshold: 0.1 
Correction method: fdr
Number of queries: 12
Number of terms per query:
   48hr_Original_Equivalent -> 893
   72hr_dPFs_Equivalent -> 887
   48hr_dPFs_Equivalent -> 870
   72hr_Original_Equivalent -> 678
   72hr_dPFs_Downregulated -> 621
   72hr_dPFs_Upregulated -> 537
   48hr_dPFs_Upregulated -> 400
   48hr_Original_Upregulated -> 295
   48hr_dPFs_Downregulated -> 223
   48hr_Original_Downregulated -> 97
   72hr_Original_Upregulated -> 44
   72hr_Original_Downregulated -> 27
Number of Unique enriched terms: 2339
Number of total terms saved: 5572
Enrichment result are prepared in 00h:00m:30s
After the p-value cutoff of 0.01 there are total of 2431 enriched terms.
Number of enriched terms per query with p-val cutoff 0.01:
   48hr_Original_Upregulated -> 37
   48hr_Original_Downregulated -> 6
   48hr_Original_Equivalent -> 449
   48hr_dPFs_Upregulated -> 140
   48hr_dPFs_Downregulated -> 109
   48hr_dPFs_Equivalent -> 446
   72hr_Original_Upregulated -> 5
   72hr_Original_Equivalent -> 296
   72hr_dPFs_Upregulated -> 205
   72hr_dPFs_Downregulated -> 294
   72hr_dPFs_Equivalent -> 444
```

At the enrichment cutoff the number of enriched GO terms varies across methods and statuses. While I can look invidually I wanted to create a summary plot that shows the number of enriched GO terms across all method-duration-status combinations for easier comparison.

---

##### 05.3 Summary of Enriched GO Terms Across Methods¶

###### 05.3.1 Heatmap of Enriched GO Terms Across Methods¶

This is a simple heatmap showing the number of enriched GO terms across all combinations of quantification methods, timepoints, and protein statuses. This visualization allows easy comparison of the biological insights captured by each approach.

```
from matplotlib import cm
import matplotlib.patches as mpatches
import matplotlib.colors as mcolors

# --- Prepare the heatmap data ---
heatmap_df = subset_gp_res.pivot_table(
    index=['Duration', 'Method'],
    columns=['source', 'Status'],
    values='name',
    aggfunc='count',
    fill_value=0
)

# --- Set up the figure and axes ---
fig, ax = plt.subplots(figsize=(14, 5))

# --- Draw the heatmap ---
sns.heatmap(
    heatmap_df,
    cmap="Blues",
    annot=True,
    fmt=".0f",
    linewidths=0.7,
    linecolor='lightgrey',
    square=False,
    ax=ax,
    cbar_kws={
        "label": "Number of Enriched Terms",
        "orientation": "horizontal",
        "shrink": 0.7,
        "pad": 0.12
    },
    annot_kws={"fontsize": 12, "fontweight": "bold",},
    mask=heatmap_df.isnull(),
    vmin=0
)

# --- Axis and tick styling ---
ax.set_xlabel("Enrichment Source & Status", fontsize=16, fontweight="bold", labelpad=16)
ax.set_ylabel("Duration & Method", fontsize=16, fontweight="bold", labelpad=16)
ax.set_title(
    "Pathway Enrichment Results by Method, Duration, Source, and Status",
    fontsize=20, fontweight="bold", pad=22, color="#222222"
)

# --- Improve tick labels ---
xticklabels = [f"{source}\n{status}" for source, status in heatmap_df.columns]
ax.set_xticklabels(xticklabels, rotation=35, ha='right', fontsize=13, fontweight='semibold', color="#333333")
yticklabels = [f"{duration}\n{method}" for duration, method in heatmap_df.index]
ax.set_yticklabels(yticklabels, rotation=0, fontsize=14, fontweight='semibold', color="#333333")

# --- Add gridlines for better separation ---
ax.hlines(
    [i for i in range(1, len(heatmap_df.index))],
    *ax.get_xlim(),
    colors='grey', linestyles='dashed', linewidth=0.5, alpha=0.25
)
ax.vlines(
    [i for i in range(1, len(heatmap_df.columns))],
    *ax.get_ylim(),
    colors='grey', linestyles='dashed', linewidth=0.5, alpha=0.25
)

# --- Add value labels for zero cells (optional, faded) ---
for (i, j), val in np.ndenumerate(heatmap_df.values):
    if val == 0:
        ax.text(
            j + 0.5, i + 0.5, "0",
            ha='center', va='center',
            fontsize=11, color="#bbbbbb", alpha=0.7
        )

# --- Add extra annotation for total terms per row/column (optional) ---
row_totals = heatmap_df.sum(axis=1)
for i, total in enumerate(row_totals):
    ax.text(
        len(heatmap_df.columns) + 0.3, i + 0.5, f"Σ={total}",
        va='center', ha='left', fontsize=13, color="#444444", fontweight='bold'
    )
col_totals = heatmap_df.sum(axis=0)
for j, total in enumerate(col_totals):
    ax.text(
        j + 0.5, len(heatmap_df.index) + 0.25, f"{total}", 
        ha='center', va='bottom', fontsize=12, color="#444444", fontweight='bold'
    )

# --- Add color rectangles for Duration (row side) and Source (column side) ---
# Use dura_colors for Duration
duration_list = heatmap_df.index.get_level_values('Duration').unique()
duration_colors = dura_colors

source_list = heatmap_df.columns.get_level_values('source').unique()
source_colors = dict(zip(source_list, cm.Set2.colors[:len(source_list)]))

# Row-side rectangles for Duration
for i, (duration, method) in enumerate(heatmap_df.index):
    rect = mpatches.Rectangle(
        (-0.9, i), 0.18, 1,  # x, y, width, height
        facecolor=duration_colors[duration],
        edgecolor='none',
        transform=ax.transData,
        clip_on=False
    )
    ax.add_patch(rect)

# Column-side rectangles for Source
for j, (source, status) in enumerate(heatmap_df.columns):
    rect = mpatches.Rectangle(
        (j, len(heatmap_df.index) + 0.08), 1, 0.18,  # x, y, width, height
        facecolor=source_colors[source],
        edgecolor='none',
        transform=ax.transData,
        clip_on=False
    )
    ax.add_patch(rect)

# --- Add color rectangles for Status (column side, above heatmap) ---
status_colors_map = status_colors  # already defined

# Get the Status for each column in heatmap_df
status_list = heatmap_df.columns.get_level_values('Status')

for j, status in enumerate(status_list):
    rect = mpatches.Rectangle(
        (j, len(heatmap_df.index) + 0.28), 1, 0.18,  # x, y, width, height (above the source rectangles)
        facecolor=status_colors_map.get(status, "#cccccc"),
        edgecolor='none',
        transform=ax.transData,
        clip_on=False
    )
    ax.add_patch(rect)


# Add legends for Duration and Source, with more spacing
duration_handles = [mpatches.Patch(color=duration_colors[d], label=d) for d in duration_list]
source_handles = [mpatches.Patch(color=source_colors[s], label=s) for s in source_list]
legend1 = ax.legend(handles=duration_handles, title="Duration", loc='upper left', bbox_to_anchor=(1.03, 1), fontsize=13, title_fontsize=14, borderaxespad=1)
legend2 = ax.legend(handles=source_handles, title="Source", loc='upper left', bbox_to_anchor=(1.03, 0.7), fontsize=13, title_fontsize=14, borderaxespad=1)
ax.add_artist(legend1)

# --- Tight layout and finalize ---
plt.subplots_adjust(left=0.18, right=0.82, top=0.92, bottom=0.13)
plt.tight_layout(rect=[0, 0, 0.97, 1])
plots.finalize_plot(
    fig, 
    show=True,
    save=True,
    filename=f"Enrichment_Heatmap_AllSets",
    filepath=figure_path,
    formats=figure_formats, 
    transparent=transparent_bg,
    dpi=figure_dpi,
)
```

The figure effectivelly displays how the dPFs expands the biological insights from each protein status, timepoint, and method combination by increasing the number of enriched GO terms detected.

###### 05.3.2 Linear Flow Diagram¶

The linear flow diagram going from `Original` to `dPFs` method for each timepoint and status shows how many new GO terms are gained or lost when moving to proteoform-aware quantification. This highlights the impact of method choice on biological interpretations.

```
# --- Reshape the data ---
# Pivot the table to get a 'Status' column for each method
# If there are multiple statuses for a term/method (e.g., multiple dPFs), keep all as separate rows
status_pivot_df = (
    subset_gp_res
    .pivot_table(
        index=['name', 'Duration', 'source', 'native'],
        columns='Method',
        values='Status',
        aggfunc=lambda x: list(x) if len(x) > 1 else x.iloc[0] if len(x) == 1 else np.nan
    )
    .reset_index()
)

# Explode any columns that have lists (i.e., multiple statuses for a method)
for col in status_pivot_df.columns:
    if col not in ['name', 'Duration', 'source', 'native']:
        if status_pivot_df[col].apply(lambda x: isinstance(x, list)).any():
            status_pivot_df = status_pivot_df.explode(col, ignore_index=True)

# Fill missing values with 'Not Enriched'
status_pivot_df = status_pivot_df.fillna('Not Enriched')

# --- Isolate the specific changes of interest ---
# Find terms that were 'Equivalent' in the 'Original' method
# but became 'Upregulated' or 'Downregulated' in the 'dPFs' method.
specific_changes_df = status_pivot_df[
    (status_pivot_df['Original'] == 'Equivalent') &
    (status_pivot_df['dPFs'].isin(['Upregulated', 'Downregulated']))
]

print("Example of pivoted data:")
print(status_pivot_df.head())
print("\nExample of isolated changes:")
print(specific_changes_df.head())
```

```
Example of pivoted data:
Method                                           name Duration source      native      Original           dPFs
0       2-oxoglutarate-dependent dioxygenase activity     48hr  GO:MF  GO:0016706   Upregulated    Upregulated
1       2-oxoglutarate-dependent dioxygenase activity     72hr  GO:MF  GO:0016706  Not Enriched    Upregulated
2                   6-phosphofructo-2-kinase activity     48hr  GO:MF  GO:0003873   Upregulated   Not Enriched
3                                     90S preribosome     72hr  GO:CC  GO:0030686  Not Enriched  Downregulated
4                               ADP metabolic process     48hr  GO:BP  GO:0046031   Upregulated   Not Enriched

Example of isolated changes:
Method                     name Duration source      native    Original           dPFs
9                   ATP binding     48hr  GO:MF  GO:0005524  Equivalent  Downregulated
10                  ATP binding     72hr  GO:MF  GO:0005524  Equivalent  Downregulated
11                  ATP binding     72hr  GO:MF  GO:0005524  Equivalent    Upregulated
14      ATP hydrolysis activity     48hr  GO:MF  GO:0016887  Equivalent  Downregulated
15      ATP hydrolysis activity     72hr  GO:MF  GO:0016887  Equivalent  Downregulated
```

```
def plot_linear_flow(ax, data, methods, colors, status_order, add_legend=False):
    """
    Draws a sequential flow chart for a given list of methods on a matplotlib axes.
    """
    num_methods = len(methods)
    
    # --- Pre-calculate totals for all bars ---
    bar_totals = {}
    for method in methods:
        bar_totals[method] = data[method].value_counts().reindex(status_order, fill_value=0)

    # --- Draw all stacked bars with annotations ---
    for i, method in enumerate(methods):
        bottom = 0
        totals = bar_totals[method]
        for status in status_order:
            height = totals.get(status, 0)
            label = status if add_legend and i == 0 else None # Add legend info only once
            ax.bar(i + 1, height, bottom=bottom, color=colors[status], width=0.5, label=label, alpha=0.9, edgecolor='black', linewidth=1)
            if height > 5: # Annotation threshold
                ax.text(i + 1, bottom + height / 2, int(height), ha='center', va='center', color='white', fontsize=12, fontweight='bold')
            bottom += height

    # --- Draw all connecting flow polygons ---
    for i in range(num_methods - 1):
        method1, method2 = methods[i], methods[i+1]
        
        flow_counts = data.groupby([method1, method2]).size().unstack(fill_value=0)
        flow_counts = flow_counts.reindex(index=status_order, columns=status_order, fill_value=0)
        
        left_totals = bar_totals[method1]
        right_totals = bar_totals[method2]
        
        left_bottoms = left_totals.cumsum() - left_totals
        right_bottoms = right_totals.cumsum() - right_totals
        
        for left_status in status_order:
            for right_status in status_order:
                count = flow_counts.at[left_status, right_status]
                if count > 0:
                    y_left_start = left_bottoms[left_status]
                    y_right_start = right_bottoms[right_status]
                    polygon = plt.Polygon([
                        [i + 1.25, y_left_start], [i + 1.75, y_right_start],
                        [i + 1.75, y_right_start + count], [i + 1.25, y_left_start + count]
                    ], facecolor=colors[left_status], alpha=0.35, edgecolor='none')
                    ax.add_patch(polygon)
                    left_bottoms[left_status] += count
                    right_bottoms[right_status] += count

# --- Main Plotting Setup ---
durations = ['48hr', '72hr']
methods_to_plot = ['Original', 'dPFs']
status_order = ['Upregulated', 'Downregulated', 'Equivalent', 'Not Enriched']
status_colors = {
    'Not Enriched': "#C2C0C0",
    'Upregulated': '#780000',
    'Downregulated': '#e36414',
    'Equivalent': '#003049',
}

# Create the subplot grid (single column)
fig, axes = plt.subplots(
    nrows=len(durations),
    ncols=1,
    figsize=(5, 10),
    sharex=True,
    sharey=True, 
    gridspec_kw={'hspace': 0.05}
)
# Ensure axes is always an array
if len(durations) == 1:
    axes = [axes]

# --- Iterate through durations and plot ---
for i, duration in enumerate(durations):
    ax = axes[i]
    flow_data = status_pivot_df[status_pivot_df['Duration'] == duration]
    
    # Call the main plotting function
    plot_linear_flow(ax, flow_data, methods_to_plot, status_colors, status_order, add_legend=(i==0))

    # --- Styling and Labeling ---
    ax.spines[['top', 'right']].set_visible(False)
    ax.set_ylabel(f'Duration: {duration}\n\nNumber of Terms', fontsize=12, fontweight='bold')

# --- Final Figure-Level Adjustments ---
# Set shared X-axis labels
plt.xticks(ticks=range(1, len(methods_to_plot) + 1), labels=methods_to_plot, fontsize=15)

# Create a single, shared legend
handles, labels = axes[0].get_legend_handles_labels()


fig.suptitle("Linear Flow of Term Status Across All Methods", fontsize=18, fontweight='bold')

# Adjust layout to make room for the legend and prevent overlap
fig.subplots_adjust(left=0.1, right=0.88, top=0.92, bottom=0.15, hspace=0.2)

# Place the legend to the right of the subplots
fig.legend(handles, labels, loc='center right', fontsize=12, title='Term Status', title_fontsize=14, borderaxespad=1)

plots.finalize_plot(
    fig, 
    show=True,
    save=True,
    filename=f"LinearFlow_AllMethods_AllDurations",
    filepath=figure_path,
    formats=figure_formats, 
    transparent=transparent_bg,
    dpi=figure_dpi,
)
```

The enrichment results by term status at each timepoint (facet), shows that majority terms that were from statistically equivalent sets in `Original` method become up/down regulated in `dPFs` method showing the added biological insights gained from proteoform-aware quantification. This also shows that with added proteoform resolution, more specific biological processes can be uncovered that are missed at the protein level.

##### 05.4 Examining in Detail for Selected Groups/Categories¶

In this part, I will be focusing on uncovering the groups and categories within the enriched GO terms that are particularly interesting or relevant to hypoxia biology. This may involve looking for terms related to known hypoxia pathways, cellular stress responses, metabolic changes, or other processes implicated in the cellular response to low oxygen conditions.

**Examining the Top MF and BP Terms**

```
duplicated_names = status_pivot_df.groupby('name')['Duration'].nunique()
duplicated_names = duplicated_names[duplicated_names == 2].index.tolist()
print(f"Terms appearing in both durations: {len(duplicated_names)}")
print()
print("GO:MF Terms:")
utils.view_table(
    status_pivot_df[
        (status_pivot_df['name'].isin(duplicated_names)) &
        (status_pivot_df['source'] == 'GO:MF')
    ].sort_values(['name', 'Duration']),
    page_number=1
)
print()
print("GO:BP Terms:")
utils.view_table(
        status_pivot_df[
        (status_pivot_df['name'].isin(duplicated_names)) &
        (status_pivot_df['source'] == 'GO:BP')
    ].sort_values(['name', 'Duration']),
    page_number=1
)
```

```
Terms appearing in both durations: 421

GO:MF Terms:
Total pages: 12, Current page: 1
```

| Method | name | Duration | source | native | Original | dPFs |
| --- | --- | --- | --- | --- | --- | --- |
| 0 | 2-oxoglutarate-dependent dioxygenase activity | 48hr | GO:MF | GO:0016706 | Upregulated | Upregulated |
| 1 | 2-oxoglutarate-dependent dioxygenase activity | 72hr | GO:MF | GO:0016706 | Not Enriched | Upregulated |
| 8 | ATP binding | 48hr | GO:MF | GO:0005524 | Equivalent | Equivalent |
| 9 | ATP binding | 48hr | GO:MF | GO:0005524 | Equivalent | Downregulated |
| 10 | ATP binding | 72hr | GO:MF | GO:0005524 | Equivalent | Downregulated |
| 11 | ATP binding | 72hr | GO:MF | GO:0005524 | Equivalent | Upregulated |
| 12 | ATP binding | 72hr | GO:MF | GO:0005524 | Equivalent | Equivalent |
| 13 | ATP hydrolysis activity | 48hr | GO:MF | GO:0016887 | Equivalent | Equivalent |
| 14 | ATP hydrolysis activity | 48hr | GO:MF | GO:0016887 | Equivalent | Downregulated |
| 15 | ATP hydrolysis activity | 72hr | GO:MF | GO:0016887 | Equivalent | Downregulated |
| 16 | ATP hydrolysis activity | 72hr | GO:MF | GO:0016887 | Equivalent | Upregulated |
| 17 | ATP-dependent activity | 48hr | GO:MF | GO:0140657 | Equivalent | Equivalent |
| 18 | ATP-dependent activity | 48hr | GO:MF | GO:0140657 | Equivalent | Downregulated |
| 19 | ATP-dependent activity | 72hr | GO:MF | GO:0140657 | Equivalent | Downregulated |
| 20 | ATP-dependent activity | 72hr | GO:MF | GO:0140657 | Equivalent | Upregulated |
| 21 | ATP-dependent activity, acting on DNA | 48hr | GO:MF | GO:0008094 | Not Enriched | Equivalent |
| 22 | ATP-dependent activity, acting on DNA | 72hr | GO:MF | GO:0008094 | Equivalent | Downregulated |
| 23 | ATP-dependent activity, acting on DNA | 72hr | GO:MF | GO:0008094 | Equivalent | Upregulated |
| 24 | ATP-dependent activity, acting on RNA | 48hr | GO:MF | GO:0008186 | Equivalent | Equivalent |
| 25 | ATP-dependent activity, acting on RNA | 72hr | GO:MF | GO:0008186 | Not Enriched | Downregulated |
| 26 | ATP-dependent activity, acting on RNA | 72hr | GO:MF | GO:0008186 | Not Enriched | Upregulated |
| 27 | ATP-dependent chromatin remodeler activity | 48hr | GO:MF | GO:0140658 | Not Enriched | Equivalent |
| 28 | ATP-dependent chromatin remodeler activity | 72hr | GO:MF | GO:0140658 | Equivalent | Upregulated |
| 29 | ATP-dependent chromatin remodeler activity | 72hr | GO:MF | GO:0140658 | Equivalent | Downregulated |
| 30 | ATP-dependent chromatin remodeler activity | 72hr | GO:MF | GO:0140658 | Equivalent | Equivalent |

```
GO:BP Terms:
Total pages: 27, Current page: 1
```

| Method | name | Duration | source | native | Original | dPFs |
| --- | --- | --- | --- | --- | --- | --- |
| 36 | DNA damage response | 48hr | GO:BP | GO:0006974 | Equivalent | Not Enriched |
| 37 | DNA damage response | 72hr | GO:BP | GO:0006974 | Equivalent | Downregulated |
| 39 | DNA metabolic process | 48hr | GO:BP | GO:0006259 | Equivalent | Equivalent |
| 40 | DNA metabolic process | 72hr | GO:BP | GO:0006259 | Not Enriched | Downregulated |
| 42 | DNA repair | 48hr | GO:BP | GO:0006281 | Not Enriched | Equivalent |
| 43 | DNA repair | 72hr | GO:BP | GO:0006281 | Equivalent | Downregulated |
| 44 | DNA repair | 72hr | GO:BP | GO:0006281 | Equivalent | Equivalent |
| 48 | DNA-templated transcription elongation | 48hr | GO:BP | GO:0006354 | Equivalent | Equivalent |
| 49 | DNA-templated transcription elongation | 72hr | GO:BP | GO:0006354 | Equivalent | Not Enriched |
| 81 | Golgi organization | 48hr | GO:BP | GO:0007030 | Equivalent | Equivalent |
| 82 | Golgi organization | 72hr | GO:BP | GO:0007030 | Not Enriched | Downregulated |
| 84 | Golgi to plasma membrane transport | 48hr | GO:BP | GO:0006893 | Not Enriched | Equivalent |
| 85 | Golgi to plasma membrane transport | 72hr | GO:BP | GO:0006893 | Not Enriched | Equivalent |
| 87 | Golgi vesicle transport | 48hr | GO:BP | GO:0048193 | Equivalent | Equivalent |
| 88 | Golgi vesicle transport | 72hr | GO:BP | GO:0048193 | Equivalent | Equivalent |
| 102 | RNA export from nucleus | 48hr | GO:BP | GO:0006405 | Equivalent | Equivalent |
| 103 | RNA export from nucleus | 72hr | GO:BP | GO:0006405 | Equivalent | Equivalent |
| 107 | RNA localization | 48hr | GO:BP | GO:0006403 | Equivalent | Equivalent |
| 108 | RNA localization | 72hr | GO:BP | GO:0006403 | Equivalent | Equivalent |
| 109 | RNA localization | 72hr | GO:BP | GO:0006403 | Equivalent | Downregulated |
| 112 | RNA processing | 48hr | GO:BP | GO:0006396 | Equivalent | Equivalent |
| 113 | RNA processing | 72hr | GO:BP | GO:0006396 | Equivalent | Not Enriched |
| 114 | RNA splicing | 48hr | GO:BP | GO:0008380 | Equivalent | Equivalent |
| 115 | RNA splicing | 72hr | GO:BP | GO:0008380 | Equivalent | Equivalent |
| 116 | RNA splicing, via transesterification reactions | 48hr | GO:BP | GO:0000375 | Equivalent | Equivalent |

There are many cases where same terms are enriched at more than one status for a given timepoint, this presents a very interesting case that dPFs as we have shown before can capture different proteoform-level regulations that are missed at the protein level. Protein level cannot represent that protein at two different regulatory states at the same timepoint, but with dPFs we can capture that.

```
# Identify cases where same Duration-Method is repeated with different Status
conflicts = subset_gp_res.groupby(['name', 'Duration', 'Method']).filter(lambda x: x['Status'].nunique() > 1)
conflicts = conflicts.sort_values(['name', 'Duration', 'Method'])
utils.view_table(conflicts[conflicts['source']=='GO:BP'], page_number=1)
```

```
Total pages: 15, Current page: 1
```

|  | query | source | native | name | significant | p\_value | p\_capped | -log10(p\_value) | -log10(p\_capped) | Enrichment | GeneRatio | parents | Duration | Method | Status |
| --- | --- | --- | --- | --- | --- | --- | --- | --- | --- | --- | --- | --- | --- | --- | --- |
| 404 | 72hr\_dPFs\_Downregulated | GO:BP | GO:0006281 | DNA repair | True | 0.0000 | 0.0000 | 6.6775 | 6.6775 | 1.8805 | 0.1874 | ['GO:0006259', 'GO:0006974'] | 72hr | dPFs | Downregulated |
| 2375 | 72hr\_dPFs\_Equivalent | GO:BP | GO:0006281 | DNA repair | True | 0.0093 | 0.0093 | 2.0324 | 2.0324 | 1.4316 | 0.1711 | ['GO:0006259', 'GO:0006974'] | 72hr | dPFs | Equivalent |
| 1982 | 72hr\_dPFs\_Equivalent | GO:BP | GO:0006403 | RNA localization | True | 0.0048 | 0.0048 | 2.3186 | 2.3186 | 1.8131 | 0.2167 | ['GO:0033036'] | 72hr | dPFs | Equivalent |
| 2229 | 72hr\_dPFs\_Downregulated | GO:BP | GO:0006403 | RNA localization | True | 0.0071 | 0.0071 | 2.1465 | 2.1465 | 1.8958 | 0.1889 | ['GO:0033036'] | 72hr | dPFs | Downregulated |
| 259 | 72hr\_dPFs\_Equivalent | GO:BP | GO:0030036 | actin cytoskeleton organization | True | 0.0000 | 0.0000 | 8.8527 | 8.8527 | 1.9589 | 0.2341 | ['GO:0007010', 'GO:0030029'] | 72hr | dPFs | Equivalent |
| 1880 | 72hr\_dPFs\_Downregulated | GO:BP | GO:0030036 | actin cytoskeleton organization | True | 0.0038 | 0.0038 | 2.4236 | 2.4236 | 1.5739 | 0.1568 | ['GO:0007010', 'GO:0030029'] | 72hr | dPFs | Downregulated |
| 257 | 72hr\_dPFs\_Equivalent | GO:BP | GO:0030029 | actin filament-based process | True | 0.0000 | 0.0000 | 8.8713 | 8.8713 | 1.9137 | 0.2287 | ['GO:0009987'] | 72hr | dPFs | Equivalent |
| 1420 | 72hr\_dPFs\_Downregulated | GO:BP | GO:0030029 | actin filament-based process | True | 0.0010 | 0.0010 | 3.0125 | 3.0125 | 1.6066 | 0.1601 | ['GO:0009987'] | 72hr | dPFs | Downregulated |
| 1924 | 72hr\_dPFs\_Downregulated | GO:BP | GO:0031032 | actomyosin structure organization | True | 0.0042 | 0.0042 | 2.3731 | 2.3731 | 2.1506 | 0.2143 | ['GO:0030036'] | 72hr | dPFs | Downregulated |
| 2361 | 72hr\_dPFs\_Equivalent | GO:BP | GO:0031032 | actomyosin structure organization | True | 0.0089 | 0.0089 | 2.0493 | 2.0493 | 1.9260 | 0.2302 | ['GO:0030036'] | 72hr | dPFs | Equivalent |
| 495 | 48hr\_dPFs\_Downregulated | GO:BP | GO:0048856 | anatomical structure development | True | 0.0000 | 0.0000 | 5.9429 | 5.9429 | 1.5076 | 0.0530 | ['GO:0032502'] | 48hr | dPFs | Downregulated |
| 723 | 48hr\_dPFs\_Upregulated | GO:BP | GO:0048856 | anatomical structure development | True | 0.0000 | 0.0000 | 4.6891 | 4.6891 | 1.4520 | 0.0541 | ['GO:0032502'] | 48hr | dPFs | Upregulated |
| 455 | 72hr\_dPFs\_Upregulated | GO:BP | GO:0048856 | anatomical structure development | True | 0.0000 | 0.0000 | 6.2001 | 6.2001 | 1.2893 | 0.1298 | ['GO:0032502'] | 72hr | dPFs | Upregulated |
| 589 | 72hr\_dPFs\_Downregulated | GO:BP | GO:0048856 | anatomical structure development | True | 0.0000 | 0.0000 | 5.4026 | 5.4026 | 1.2642 | 0.1260 | ['GO:0032502'] | 72hr | dPFs | Downregulated |
| 720 | 48hr\_dPFs\_Upregulated | GO:BP | GO:0009653 | anatomical structure morphogenesis | True | 0.0000 | 0.0000 | 4.7030 | 4.7030 | 1.7940 | 0.0668 | ['GO:0032502', 'GO:0048856'] | 48hr | dPFs | Upregulated |
| 1049 | 48hr\_dPFs\_Downregulated | GO:BP | GO:0009653 | anatomical structure morphogenesis | True | 0.0002 | 0.0002 | 3.6577 | 3.6577 | 1.7228 | 0.0606 | ['GO:0032502', 'GO:0048856'] | 48hr | dPFs | Downregulated |
| 1325 | 72hr\_dPFs\_Upregulated | GO:BP | GO:0009653 | anatomical structure morphogenesis | True | 0.0007 | 0.0007 | 3.1456 | 3.1456 | 1.3584 | 0.1368 | ['GO:0032502', 'GO:0048856'] | 72hr | dPFs | Upregulated |
| 1952 | 72hr\_dPFs\_Equivalent | GO:BP | GO:0009653 | anatomical structure morphogenesis | True | 0.0045 | 0.0045 | 2.3462 | 2.3462 | 1.2679 | 0.1515 | ['GO:0032502', 'GO:0048856'] | 72hr | dPFs | Equivalent |
| 1216 | 72hr\_dPFs\_Upregulated | GO:BP | GO:0048513 | animal organ development | True | 0.0005 | 0.0005 | 3.3230 | 3.3230 | 1.3453 | 0.1354 | ['GO:0048856'] | 72hr | dPFs | Upregulated |
| 1747 | 72hr\_dPFs\_Downregulated | GO:BP | GO:0048513 | animal organ development | True | 0.0028 | 0.0028 | 2.5528 | 2.5528 | 1.2959 | 0.1291 | ['GO:0048856'] | 72hr | dPFs | Downregulated |
| 205 | 48hr\_dPFs\_Equivalent | GO:BP | GO:0065007 | biological regulation | True | 0.0000 | 0.0000 | 10.1556 | 10.1556 | 1.1100 | 0.2499 | ['GO:0008150'] | 48hr | dPFs | Equivalent |
| 305 | 48hr\_dPFs\_Upregulated | GO:BP | GO:0065007 | biological regulation | True | 0.0000 | 0.0000 | 8.0380 | 8.0380 | 1.2758 | 0.0475 | ['GO:0008150'] | 48hr | dPFs | Upregulated |
| 688 | 48hr\_dPFs\_Downregulated | GO:BP | GO:0065007 | biological regulation | True | 0.0000 | 0.0000 | 4.8534 | 4.8534 | 1.2294 | 0.0432 | ['GO:0008150'] | 48hr | dPFs | Downregulated |
| 204 | 72hr\_dPFs\_Upregulated | GO:BP | GO:0065007 | biological regulation | True | 0.0000 | 0.0000 | 10.1838 | 10.1838 | 1.1818 | 0.1190 | ['GO:0008150'] | 72hr | dPFs | Upregulated |
| 355 | 72hr\_dPFs\_Downregulated | GO:BP | GO:0065007 | biological regulation | True | 0.0000 | 0.0000 | 7.2194 | 7.2194 | 1.1529 | 0.1149 | ['GO:0008150'] | 72hr | dPFs | Downregulated |

**Custom Terms of Interest**: is the custom grouping that I wanted to highlight in a single figure.

```
# A dictionary categorizing the GO terms into logical biological groups.
custom_terms_of_interest = {
    'Response to Stimulus': [
        'response to stress',
        'response to stimulus',
        'cellular response to stimulus',
        'response to hypoxia',
        'response to decreased oxygen levels',
        'response to oxygen levels',
        'cellular response to hypoxia',
        'cellular response to decreased oxygen levels',
    ],
    'Metabolism': [
        'hexose metabolic process',
        'glucose metabolic process',
        'carbohydrate metabolic process',
        'ADP metabolic process',

        'mRNA metabolic process',
        'RNA processing',
        'mRNA processing',
    ],
    'Cell Cycle': [
        'cell cycle',
        'cell division',
        'chromosome segregation',
        'DNA damage response',
        'DNA repair',
    ],
    'Cellular Organization': [
        'cellular process',
        'cell development',
        'cell differentiation',
        'cell morphogenesis',
        'cellular component assembly',
        'cellular component organization',
        'cellular component biogenesis',
        'cellular component disassembly',
    ],
    'Signaling': [
        'cell communication',
        'signal transduction',
        'signaling',
        'small GTPase-mediated signal transduction',
    ]
}
```

---

These will be placed in a long format heatmap like grid plot with points/pie charts showing if the term is enriched in which method-duration-status combination. Most wll have single status color representing but there will be two or even three status colors for some terms showing that they are enriched in multiple statuses for a given timepoint.

```
from matplotlib.patches import Wedge, Rectangle
import textwrap

# Defining custom terms of interest for the grid layout
custom_terms_of_interest = {
    # Column 1 - Row 1
    # Logic: Broad Abiotic -> Oxygen Levels -> Hypoxia -> Cellular specific responses
    'Response to Stimulus': [
        'response to abiotic stimulus',
        'response to oxygen levels',
        'response to decreased oxygen levels',
        'response to hypoxia',
        'cellular response to decreased oxygen levels',
        'cellular response to hypoxia',
    ],
    
    # Column 1 - Row 2
    # Logic: General Process -> Developmental -> Structural Organization -> Specific Structures
    'Cellular Organization': [
        'cellular process',
        'cell differentiation',
        'cell morphogenesis',
        'cellular component organization',
        'organelle organization',
        'cytoskeleton organization',
    ],

    # Column 2 - Row 1
    # Logic: Cycle -> Chromosome -> DNA Metabolism -> Damage/Repair mechanisms
    'Cell Cycle': [
        'cell cycle',
        'chromosome organization',
        'DNA metabolic process',
        'DNA damage response',
        'DNA repair',
    ],

    # Column 2 - Row 2
    # Logic: General Metabolism -> Biosynthesis -> Specific Macromolecules -> Degradation
    'Metabolism': [
        'metabolic process', 
        'biosynthetic process',
        'nucleic acid metabolic process',
        'protein metabolic process',
        'proteolysis',
    ],

    # Column 3 - Row 1
    # Logic: Communication -> Signaling -> Transduction (Mechanism)
    'Signaling': [
        'cell communication',
        'signaling',
        'signal transduction',
    ],

    # Column 3 - Row 2
    # Logic: Splicing (Processing) -> Transport (General) -> Transport (Specific)
    'RNA Processing': [
        'RNA splicing',
        'RNA transport',
        'mRNA transport',
    ],
}

# ==========================================
# 1. SETUP & EXACT LAYOUT DEFINITION
# ==========================================

terms_of_interest = custom_terms_of_interest

# STRICTLY enforcing your requested grid:
layout_grid = [
    ['Response to Stimulus', 'Cellular Organization'],
    ['Cell Cycle', 'Metabolism'],                       
    ['Signaling', 'RNA Processing']                     
]

# Calculate max terms per row to ensure consistent scaling
row_max_terms = []
for row_cats in layout_grid:
    lengths = [len(terms_of_interest.get(cat, [])) for cat in row_cats]
    row_max_terms.append(max(max(lengths), 1))

# Calculate Figure Height
fig_height = sum(row_max_terms) * 0.5 + 2.5
fig, axes = plt.subplots(
    nrows=3, ncols=2, 
    figsize=(12, fig_height), 
    sharex=False, # Changed to False so we can manually control top ticks
    gridspec_kw={'height_ratios': row_max_terms}
)
axes_flat = axes.flatten()

# Prepare Data
df = subset_gp_res.copy()
if 'name' in df.columns:
    df = df.rename(columns={'name': 'Term'})
df['Duration'] = df['Duration'].astype(str)
df['Method'] = df['Method'].astype(str)
df['neglog10_p'] = -np.log10(df['p_value'].clip(lower=1e-300))

# Global Bubble Scaling
all_plot_terms = [t for cat in terms_of_interest.values() for t in cat]
global_max_p = df[df['Term'].isin(all_plot_terms)]['neglog10_p'].max()

min_area = 20
size_scale = 40
max_s_val = min_area + (global_max_p * size_scale) if not pd.isna(global_max_p) else min_area

def s_to_radius(s_val, max_s):
    max_radius = 0.4 
    return max_radius * np.sqrt(s_val / max_s) if max_s > 0 else 0

# ==========================================
# 2. PLOTTING LOOP
# ==========================================

status_order = ['Upregulated', 'Downregulated', 'Equivalent', 'Not Enriched']
methods_to_show = ['Original', 'dPFs']
durations_to_show = ['48hr', '72hr']

ordered_cats = [cat for row in layout_grid for cat in row]

for i, category in enumerate(ordered_cats):
    ax = axes_flat[i]
    terms = terms_of_interest.get(category, [])
    
    # Row specific formatting
    row_idx = i // 2
    current_row_max = row_max_terms[row_idx]
    
    # --- BACKGROUNDS & GRID ---
    # 48hr Background (Light Gray)
    ax.add_patch(Rectangle((-0.5, -0.5), 2.0, current_row_max, facecolor=dura_colors['48hr'], zorder=0, alpha=0.75))
    # 72hr Background (Darker Gray)
    ax.add_patch(Rectangle((1.5, -0.5), 2.0, current_row_max, facecolor=dura_colors['72hr'], zorder=0, alpha=0.75))
    
    # White horizontal lines
    for y_line in range(current_row_max):
        ax.axhline(y_line, color='white', linewidth=1.5, zorder=1)

    # --- PLOT DATA ---
    if terms:
        # Map terms to Y-positions top-down
        y_map = {}
        for idx, term in enumerate(terms):
            y_map[term] = (current_row_max - 1) - idx

        plot_df = df[df['Term'].isin(terms)].copy()
        all_combos = pd.MultiIndex.from_product(
            [terms, durations_to_show, methods_to_show],
            names=['Term', 'Duration', 'Method']
        )
        agg_df = plot_df.groupby(['Term', 'Duration', 'Method']).agg(
            statuses=('Status', list),
            p_values=('neglog10_p', list)
        ).reindex(all_combos).reset_index()

        for _, row in agg_df.iterrows():
            raw_status = row['statuses']
            if isinstance(raw_status, list):
                statuses = raw_status
                p_vals = row['p_values']
            else:
                statuses = ['Not Enriched']
                p_vals = [0]
                
            term = row['Term']
            if term not in y_map: continue
            
            x_idx = 0
            if str(row['Duration']) == '72hr': x_idx += 2
            if str(row['Method']) == 'dPFs': x_idx += 1
            y = y_map[term]
            
            if statuses[0] == 'Not Enriched':
                ax.scatter(x_idx, y, s=min_area, facecolors='none', edgecolors='#cccccc', linewidths=1.0, zorder=2)
                continue
            
            max_p = max(p_vals) if p_vals else 0
            s_val = min_area + max_p * size_scale
            radius = s_to_radius(s_val, max_s_val)
            
            pairs = sorted(zip(statuses, p_vals), key=lambda x: status_order.index(x[0]) if x[0] in status_order else 99)
            angle_step = 360 / len(pairs)
            
            for w_i, (status, p) in enumerate(pairs):
                start = w_i * angle_step
                end = (w_i + 1) * angle_step
                color = status_colors.get(status, '#cccccc')
                w = Wedge((x_idx, y), radius, start, end, facecolor=color, edgecolor='#444444', linewidth=0.5, zorder=3)
                ax.add_patch(w)
        
        # Y-Axis Labels
        pos_to_label = {v: k for k, v in y_map.items()}
        tick_locs = list(pos_to_label.keys())
        tick_labels = [textwrap.fill(pos_to_label[y], width=18) for y in tick_locs]
        ax.set_yticks(tick_locs)
        ax.set_yticklabels(tick_labels, fontsize=11)
        
    else:
        ax.text(1.5, current_row_max/2 - 0.5, 'No Terms Listed', ha='center', va='center', color='#999999')
        ax.set_yticks([])

    # --- FORMATTING ---
    ax.set_ylim(-0.5, current_row_max - 0.5)
    ax.set_xlim(-0.5, 3.5)
    ax.set_aspect('equal')
    
    # Category Label (Right Side)
    ax.set_ylabel(category, fontsize=13, fontweight='bold', labelpad=15)
    ax.yaxis.set_label_position("right")
    
    for spine in ax.spines.values(): spine.set_visible(False)
    
    # --- RESTORE LABELS ---
    # Apply to top row explicitly
    if row_idx == 0:
        ax.xaxis.tick_top()
        ax.set_xticks([0, 1, 2, 3])
        # Force the subcolumn labels to appear
        ax.set_xticklabels(['Original', 'dPFs', 'Original', 'dPFs'], fontsize=10, rotation=45, ha='left')
        
        # Super-headers (48hr / 72hr)
        # Position slightly higher to clear the angled text
        ax.text(0.5, current_row_max + 0.5, '48hr', ha='center', va='bottom', fontsize=12, fontweight='bold')
        ax.text(2.5, current_row_max + 0.5, '72hr', ha='center', va='bottom', fontsize=12, fontweight='bold')
    else:
        ax.set_xticks([])

# ==========================================
# 3. LEGENDS & SPACING
# ==========================================

status_handles = [
    plt.Line2D(
        [0], [0], marker='o', color='w', label=s,
        markerfacecolor=c, markeredgecolor='#333333', markersize=10
    ) 
    for s, c in status_colors.items() if s in status_order
]

size_handles = []
for p in [1, 5, 10, 20]:
    s = min_area + p * size_scale
    size_handles.append(
        plt.scatter([], [], s=s, label=str(p), color='#777777', edgecolor='k')
    )

fig.legend(
    handles=status_handles, title='Status', loc='upper left', 
    bbox_to_anchor=(0.91, 0.65), frameon=False
)

fig.legend(
    handles=size_handles, title='Max Significance\n(-log10 p-value)', loc='upper left', 
    bbox_to_anchor=(0.91, 0.45), frameon=False, labelspacing=1.5
)

plt.subplots_adjust(left=0.18, right=0.9, top=0.92, bottom=0.05, hspace=0.15, wspace=0.5)

plots.finalize_plot(
    fig, 
    show=True,
    save=True,
    filename=f"GOBP_Term_Enrichment_By_Category_dataDriven",
    filepath=figure_path,
    formats=figure_formats, 
    transparent=transparent_bg,
    dpi=figure_dpi,
)
```

The custom terms of interest plot effectively summarizes how proteoform-aware quantification using dPFs reveals additional biological processes relevant to hypoxia response that are missed by traditional protein-level methods. The presence of terms across multiple statuses highlights the complex regulation captured at the proteoform level.

One other type of plot that this information can be represented is the dumbell plots showing the shared/non-unique status terms between methods for each timepoint.

```
terms_of_interest = custom_terms_of_interest

df = subset_gp_res.copy()
if 'name' in df.columns:
    df = df.rename(columns={'name': 'Term'})

# Filter for the terms and methods we want to plot
plot_terms = [term for category in terms_of_interest.values() for term in category]
methods_to_show = ['Original', 'dPFs']
durations_to_show = sorted(df['Duration'].astype(str).unique())

plot_df = df[
    df['Term'].isin(plot_terms) &
    df['Method'].isin(methods_to_show)
].copy()

# Calculate significance for point sizing
plot_df['neglog10_p'] = -np.log10(plot_df['p_value'].clip(lower=1e-50))

# --- 2. Plotting Setup ---
category_order = list(terms_of_interest.keys())[::-1]
term_order = [term for category in category_order for term in terms_of_interest[category]]
y_map = {term: i for i, term in enumerate(term_order)}
plot_df['y'] = plot_df['Term'].map(y_map)
plot_df = plot_df.dropna(subset=['y'])

fig, axes = plt.subplots(
    1, len(durations_to_show),
    figsize=(16, max(7, len(term_order) * 0.45)),
    sharey=True
)
if len(durations_to_show) == 1:
    axes = [axes]

method_markers = {'Original': 'o', 'dPFs': 's'}
size_scale = 20

# --- 3. Drawing the Plot ---
for i, duration in enumerate(durations_to_show):
    ax = axes[i]
    duration_df = plot_df[plot_df['Duration'] == duration].copy()

    # Draw the horizontal range lines for each term
    if not duration_df.empty:
        range_df = duration_df.groupby('y')['GeneRatio'].agg(['min', 'max']).reset_index()
        ax.hlines(y=range_df['y'], xmin=range_df['min'], xmax=range_df['max'],
                    color='#cccccc', alpha=0.8, linewidth=1.5, zorder=1)

    # Draw the scatter points for each enrichment result
    for _, row in duration_df.iterrows():
        ax.scatter(
            x=row['GeneRatio'],
            y=row['y'],
            s=row['neglog10_p'] * size_scale,
            color=status_colors.get(row['Status'], '#cccccc'),
            marker=method_markers.get(row['Method'], 'x'),
            zorder=2,
            edgecolor='black',
            linewidth=0.5,
            alpha=0.9
        )

    # --- 4. Formatting the Axes ---
    ax.set_title(duration, fontsize=14, fontweight='bold')
    current_y = 0
    for category_name, terms in reversed(list(terms_of_interest.items())):
        if current_y > 0:
            ax.axhline(current_y - 0.5, color='#888888', linestyle='-', linewidth=1.5, zorder=0)
        current_y += len(terms)

    ax.grid(True, which='major', axis='both', color='#e0e0e0', linestyle='--', zorder=0)
    ax.set_xlabel('GeneRatio', fontsize=12, ha='right')
    ax.xaxis.set_label_coords(1.02, -0.06)
    ax.tick_params(axis='x', labelsize=11)
    ax.spines['top'].set_visible(False)
    ax.spines['right'].set_visible(False)
    ax.spines['left'].set_visible(False)
    ax.set_xlim(left=0)

# --- 5. Global Formatting and Legends ---
def wrap_label(label, width=35):
    return "\n".join(textwrap.wrap(label, width=width))

yticks = list(y_map.values())
yticklabels = [wrap_label(term) for term in y_map.keys()]
axes[0].set_yticks(yticks)
axes[0].set_yticklabels(yticklabels, fontsize=11, va='center')
axes[0].tick_params(axis='y', length=0)

# Determine position for category labels
max_xlim = max(ax.get_xlim()[1] for ax in axes)
category_label_x = -0.1 * max_xlim

current_y = 0
for category_name, terms in reversed(list(terms_of_interest.items())):
    label_y = current_y + (len(terms) - 1) / 2.0
    axes[0].text(
        category_label_x, label_y, wrap_label(category_name, 15),
        ha='center', va='center', fontsize=12, fontweight='bold',
        rotation=90, clip_on=False
    )
    current_y += len(terms)

fig.suptitle('GO Term Enrichment Comparison by Status and Method', fontsize=18, fontweight='bold', y=0.99)

# Create legends
status_elems = [plt.Line2D([0], [0], marker='o', color='w', label=s, markerfacecolor=c, markersize=10, markeredgecolor='k')
                for s, c in status_colors.items()]
legend1 = fig.legend(handles=status_elems, title='Status', loc='upper right', bbox_to_anchor=(0.98, 0.94), fontsize=11)
legend1.get_title().set_fontweight('bold')

method_elems = [plt.Line2D([0], [0], marker=m, color='w', label=l, linestyle='None', markersize=10, markeredgecolor='k')
                for l, m in method_markers.items()]
legend2 = fig.legend(handles=method_elems, title='Method', loc='upper right', bbox_to_anchor=(0.98, 0.82), fontsize=11)
legend2.get_title().set_fontweight('bold')

p_vals_for_legend = [1, 5, 15]
size_handles = [plt.scatter([], [], s=p * size_scale, label=f'{p}', color='#777777', edgecolor='k') for p in p_vals_for_legend]
legend3 = fig.legend(handles=size_handles, title='-log10(p-value)', loc='upper right', bbox_to_anchor=(0.98, 0.70), labelspacing=1.5, fontsize=11)
legend3.get_title().set_fontweight('bold')

plt.tight_layout(rect=[0.1, 0, 0.9, 0.95])
plots.finalize_plot(
    fig, 
    show=True,
    save=True,
    filename=f"GOBP_Term_Enrichment_Scatter_By_Category_dataDriven",
    filepath=figure_path,
    formats=figure_formats, 
    transparent=transparent_bg,
    dpi=figure_dpi,
)
```

The dumbell plot is not the most clean, and uses too much space, but if the main points is to show the shared/non-unique status terms between methods for each timepoint it can be useful.

---

#### Summary¶

This analysis compared four approaches to protein-level quantification and their impact on downstream analysis:

**Key Findings:**

1. **Reproducibility (CV Analysis):**

   - All methods show acceptable CVs across conditions
   - dPF-based quantification provides proteoform-level resolution without sacrificing reproducibility
2. **Sample Separation (PCA):**

   - All methods successfully separate hypoxia from normoxia conditions
   - dPF approach captures additional biological variation at the proteoform level
3. **Statistical Testing (QuEStVar):**

   - Different quantification methods yield different numbers of significant proteins
   - dPF analysis identifies cases where proteoforms within the same protein show opposing regulation
4. **Enrichment Analysis:**

   - Pathway enrichment results vary by quantification method
   - dPF-based analysis reveals enriched terms that are missed by traditional protein-level approaches

**Conclusions:**

The choice of protein quantification method has substantial impact on biological interpretation. ProteoForge's dPF-based approach provides additional resolution that can reveal regulatory complexity hidden in traditional protein-level analyses.

**Output Files:**

- QuEStVar results: `*_QuEStVar.feather` files for each method/timepoint
- Enrichment results: `gProfiler_Enrichment_Analysis_AllSets.csv`
- Figures: Saved to `figures/hypoxia/04-Downstream/`

```
print("Notebook Execution Time:", utils.prettyTimer(utils.getTime() - startTime))
```

```
Notebook Execution Time: 00h:01m:45s
```
