## Supplementary material for "ProteoForge: An Imputation-Aware Framework for Differential Proteoform Discovery in Bottom-Up Proteomics": Analysis notebooks as html render: Notebook S15.html

### Notebook S15

###### Enes Kemal Ergin

#### 2025-12-08 23:00:20

### Supplementary Notebook 15: Figure Assembly from both 48hr and 72hr Summaries¶

- **License:** Creative Commons Attribution-NonCommercial 4.0 International License
- **Version:** 0.2
- **Edit Log:**
  - 2025-11-28: Initial version of the notebook
  - 2025-12-08: Revise the whole notebook, ensuring clarity and correctness

---

**Requirements:**

The following preprocessing and method runs must be completed (in order) before executing this notebook:

1. `01-DataPreparation.ipynb` - Prepares the hypoxia data for analysis.
2. `02-ApplyingProteoForge_48hr.ipynb` - Performs ProteoForge analysis on 48-hour hypoxia data.
3. `02-ApplyingProteoForge_72hr.ipynb` - Performs ProteoForge analysis on 72-hour hypoxia data.
4. `03-SummarizeProteoForge_48hr.ipynb` - Summarizes the results from the 48-hour ProteoForge analysis.
5. `03-SummarizeProteoForge_72hr.ipynb` - Summarizes the results from the 72-hour ProteoForge analysis.

---

**Data Information:**

This notebook is an extra figure assembly that is required for combining the summary information from 48hr and 72hr ProteoForge analyses on hypoxia data.

---

**Purpose:**

To provide plots for the figure 3's summary panel, integrating results from both 48-hour and 72-hour analyses.

---

#### 01. Setup¶

This section imports required libraries, configures display settings, and defines paths for data and figures.

> **Note:** The HTML rendering of this notebook hides code cells by default. Click the "Code" buttons on the right to expand them.

##### 01.1 Libraries¶

```
# Libraries Used
import os
import sys
import warnings

import numpy as np # Numerical computing
import pandas as pd # Data manipulation

import seaborn as sns # R-like high-level plots
import matplotlib.pyplot as plt # Python's base plotting 

# home path
sys.path.append('../')
from src import utils, plots 
from src import tests, enrich

from ProteoForge import parse, process
from ProteoForge import impute, normalize

warnings.filterwarnings('ignore')
# Initialize the timer
startTime = utils.getTime()
```

##### 01.2 Configure Notebook Settings¶

Configure visualization styles, color palettes, and display options for consistent output formatting throughout the analysis.

```
## Configure Plotting Settings
def_colors = [
    "#139593", "#fca311", "#e54f2a",
    "#c3c3c3", "#555555",
    "#690000", "#5f4a00", "#004549"
]

condition_colors = {
    'True Hypoxia (1% O2) - 48 Hr': '#e9c46a',
    'True Hypoxia (1% O2) - 72 Hr': '#f4a261',
    'Normoxia (21% O2) - 48 Hr': '#ade8f4',
    'Normoxia (21% O2) - 72 Hr': '#0077b6',
}

cond_colors = {
    'Hypoxia (1% O2)': '#f4a261',
    'Normoxia (21% O2)': '#0077b6',
}
dura_colors = {
    '48hr': '#c3c3c3',
    '72hr': '#555555',
}
cond_order = ['Normoxia (21% O2)', 'Hypoxia (1% O2)']
dura_order = ['48hr', '72hr']

# Set seaborn style
sns.set_theme(
    style="white",
    context="paper",
    palette=def_colors,
    font_scale=1,
    rc={
        "figure.figsize": (6, 4),
        "font.family": "sans-serif",
        "font.sans-serif": ["Arial", "Ubuntu Mono"],
    }
)

# Figure Saving Settings
figure_formats = ["png", "pdf"]
save_to_folder = True
transparent_bg = True
figure_dpi = 300

## Configure dataframe displaying
pd.set_option('display.float_format', lambda x: '%.4f' % x)
pd.set_option('display.max_rows', 50)
pd.set_option('display.max_columns', 50)
pd.set_option('display.width', 500)  # Set a wider display width


## Printing Settings
verbose = True
```

##### 01.3 Data and Result Paths¶

- `home_path` — Base project path (e.g., `./`)
- `data_name` — Dataset identifier used throughout (`"hypoxia"`)
- `notebook_name` — This notebook name for logging and organizing outputs (`"04-Downstream"`)
- `data_path` — Root data folder (`{home_path}data/`)
- `raw_path` — Raw input folder (`{data_path}input/{data_name}/`)
- `input_path` — Cleaned input data from preprocessing (`{data_path}cleaned/{data_name}/`)
- `output_path` — ProteoForge result files (`{data_path}results/{data_name}/`)
- `figure_path` — Destination for generated figures (`{home_path}figures/{data_name}/{notebook_name}/`)

```
home_path = './'
data_name = "hypoxia"
notebook_name = "04-Downstream"
data_path = f"{home_path}data/"
raw_path = f"{data_path}input/{data_name}/"
input_path = f"{data_path}results/{data_name}/"
output_path = f"{data_path}results/{data_name}/"
figure_path = f"{home_path}figures/{data_name}/{notebook_name}/"

# Create the output folder if it does not exist
if not os.path.exists(output_path):
    os.makedirs(output_path)

# Create figure folder structure, if needed
if save_to_folder:
    for i in figure_formats:
        cur_folder = figure_path + i + "/"
        if not os.path.exists(cur_folder):
            os.makedirs(cur_folder)
```

#### 02. Individual Panels¶

##### 02.1 Protein Counts by Significant Peptides¶

This summarize the number of proteins identified, if no peptides are significant, one peptide is significant, or multiple peptides are significant per protein for both 48hr and 72hr analyses.

```
# Mapping for better labels: Single: Single Significant Peptide, Multiple: Multiple Significant Peptides, Nan: No Significant Peptides
label_map = {
    'Single': 'Single Significant Peptide',
    'Multiple': 'Multiple Significant Peptides',
    np.nan: 'No Significant Peptides'
}
color_map = {
    'Single Significant Peptide': '#669bbc',
    'Multiple Significant Peptides': '#003049',
    'No Significant Peptides': '#e5e5e5'
}

# Read 48hr and 72hr files, map labels, and pivot so durations become columns
df48 = pd.read_csv(f"{input_path}protein_significantPeptides_counts_48hr.csv")
df48['Duration'] = '48hr'
df72 = pd.read_csv(f"{input_path}protein_significantPeptides_counts_72hr.csv")
df72['Duration'] = '72hr'

# Concatenate and normalize labels
plot_data = pd.concat([df48, df72], ignore_index=True)
plot_data['Significant Peptides Category'] = (
    plot_data['Significant Peptides Category']
    .where(plot_data['Significant Peptides Category'].notna(), 'No Significant Peptides')
)
# Map short names to long names, but keep existing long names intact
plot_data['Significant Peptides Category'] = plot_data['Significant Peptides Category'].map(label_map).fillna(plot_data['Significant Peptides Category'])

# Ensure category order
plot_data['Significant Peptides Category'] = plot_data['Significant Peptides Category'].astype('category')
plot_data['Significant Peptides Category'] = plot_data['Significant Peptides Category'].cat.set_categories(
    ['No Significant Peptides', 'Single Significant Peptide', 'Multiple Significant Peptides'],
    ordered=True
)

# Create a wide-format table where durations are columns (counts and percentages)
plot_counts = plot_data.pivot_table(index='Significant Peptides Category', columns='Duration', values='Count', aggfunc='sum').reindex(columns=dura_order)
plot_percent = plot_data.pivot_table(index='Significant Peptides Category', columns='Duration', values='Percentage', aggfunc='sum').reindex(columns=dura_order)

# Reset index for a tidy wide-format ready for plotting (columns: Significant Peptides Category, 48hr, 72hr)
plot_data_wide_counts = plot_counts.reset_index().rename_axis(None, axis=1)
print(plot_data_wide_counts)
plot_data_wide_percent = plot_percent.reset_index().rename_axis(None, axis=1)
print(plot_data_wide_percent)

# Stacked bar plot: counts per Duration, stacked by Significant Peptides Category

# Data to plot: use plot_counts (index=categories, columns=durations) and plot_percent for annotations
durations = list(plot_counts.columns)
categories = list(plot_counts.index)
colors = [color_map.get(cat, "#999999") for cat in categories]
# Prepare stacked bars
fig, ax = plt.subplots(figsize=(6, 4))
bottom = np.zeros(len(durations))

for idx, (cat, clr) in enumerate(zip(categories, colors)):
    heights = plot_counts.loc[cat, durations].values
    bars = ax.bar(durations, heights, bottom=bottom, color=clr, label=cat, edgecolor="none", width=0.6)
    
    # Annotate each segment with Count\n(Percentage%)
    for i, bar in enumerate(bars):
        h = heights[i]
        if h <= 0:
            continue
        pct = plot_percent.loc[cat, durations[i]] * 100
        txt = f"{int(h):,}\n({pct:.1f}%)"
        ax.text(
            bar.get_x() + bar.get_width() / 2,
            bottom[i] + h / 2,
            txt,
            ha="center",
            va="center",
            color='k',
            fontsize=12,
            weight="bold"
        )
    bottom += heights  # update bottom for next stacked segment

# Polish plot
ax.set_title("Protein Counts by Significant Peptides Category", fontsize=15, fontweight='bold')
ax.set_ylabel("Protein Count", fontsize=13)
ax.set_xlabel("")  # X-axis is Duration, so label not strictly necessary
ax.set_ylim(0, bottom.max() * 1.05)  # give 10% headroom
ax.legend(title="Significant Peptides", bbox_to_anchor=(1.02, 1), loc="upper left")
ax.spines["top"].set_visible(False)
ax.spines["right"].set_visible(False)
ax.yaxis.grid(True, linestyle="--", linewidth=0.5, alpha=0.6)
# Resize the ticklabels and make them italic
ax.tick_params(axis='both', which='major', labelsize=12)
# Make them italic
for label in ax.get_xticklabels() + ax.get_yticklabels():
    label.set_fontstyle('italic')

plt.tight_layout()
plots.finalize_plot(
    fig, show=True, save=save_to_folder,
    filename='significantPeptides_counts_stackedBar',
    filepath=figure_path,
    formats=figure_formats, 
    transparent=transparent_bg,
    dpi=figure_dpi,
)
```

```
   Significant Peptides Category  48hr  72hr
0        No Significant Peptides  3920  2109
1     Single Significant Peptide  2185  2477
2  Multiple Significant Peptides  1056  2575
   Significant Peptides Category   48hr   72hr
0        No Significant Peptides 0.5474 0.2945
1     Single Significant Peptide 0.3051 0.3459
2  Multiple Significant Peptides 0.1475 0.3596
```

##### 02.2 Protein Counts by Proteoform Categories¶

A protein can have no significant peptides, or significant peptides in singleton clusters are PTM, or dPF if in clusters with multiple peptides. Or a protein can host both PTM and dPF peptides. This panel summarizes the counts of proteins in these categories for both 48hr and 72hr analyses.

```
# Mapping for better labels: Single: Single Significant Peptide, Multiple: Multiple Significant Peptides, Nan: No Significant Peptides
label_map = {
    'No Forms': 'No Significant Peptides',
    'PTM Only': 'Single Peptide PTM',
    'dPF Only': 'With dPF',
    'Both': 'With Single PTM & dPF',

}

color_map = {
    'No Significant Peptides': "#e5e5e5",
    'Single Peptide PTM': "#a7c957",
    'With dPF': "#bc4749",
    'With Single PTM & dPF': "#967aa1",
}

# Read 48hr and 72hr files, map labels, and pivot so durations become columns
df48 = pd.read_csv(f"{input_path}protein_formCategories_counts_48hr.csv")
df48['Duration'] = '48hr'
df72 = pd.read_csv(f"{input_path}protein_formCategories_counts_72hr.csv")
df72['Duration'] = '72hr'
plot_data = pd.concat([df48, df72], ignore_index=True)
plot_data['Form Categories'] = (
    plot_data['Form Categories']
    .where(plot_data['Form Categories'].notna(), 'No Significant Peptides')
)
# Map short names to long names, but keep existing long names intact
plot_data['Form Categories'] = plot_data['Form Categories'].map(label_map).fillna(plot_data['Form Categories'])
# Ensure category order
plot_data['Form Categories'] = plot_data['Form Categories'].astype('category')
plot_data['Form Categories'] = plot_data['Form Categories'].cat.set_categories(
    ['No Significant Peptides', 'Single Peptide PTM', 'With dPF', 'With Single PTM & dPF'],
    ordered=True
)
# Create a wide-format table where durations are columns (counts and percentages)
plot_counts = plot_data.pivot_table(index='Form Categories', columns='Duration', values='Count', aggfunc='sum').reindex(columns=dura_order)
plot_percent = plot_data.pivot_table(index='Form Categories', columns='Duration', values='Percentage', aggfunc='sum').reindex(columns=dura_order)

# Reset index for a tidy wide-format ready for plotting (columns: Form Categories, 48hr, 72hr)
plot_data_wide_counts = plot_counts.reset_index().rename_axis(None, axis=1)
print(plot_data_wide_counts)
print()
plot_data_wide_percent = plot_percent.reset_index().rename_axis(None, axis=1)
print(plot_data_wide_percent)

# Stacked bar plot: counts per Duration, stacked by Form Categories

# Data to plot: use plot_counts (index=categories, columns=durations) and plot_percent for annotations
durations = list(plot_counts.columns)
categories = list(plot_counts.index)
colors = [color_map.get(cat, "#999999") for cat in categories]
# Prepare stacked bars
fig, ax = plt.subplots(figsize=(6, 4))
bottom = np.zeros(len(durations))

for idx, (cat, clr) in enumerate(zip(categories, colors)):
    heights = plot_counts.loc[cat, durations].values
    bars = ax.bar(durations, heights, bottom=bottom, color=clr, label=cat, edgecolor="none", width=0.6)
    
    # Annotate each segment with Count\n(Percentage%)
    for i, bar in enumerate(bars):
        h = heights[i]
        if h <= 0:
            continue
        pct = plot_percent.loc[cat, durations[i]] * 100
        txt = f"{int(h):,}\n({pct:.1f}%)"
        ax.text(
            bar.get_x() + bar.get_width() / 2,
            bottom[i] + h / 2,
            txt,
            ha="center",
            va="center",
            color='k',
            fontsize=12,
            weight="bold"
        )
    bottom += heights  # update bottom for next stacked segment

# Polish plot
ax.set_title("Protein Counts by Proteoform Categories", fontsize=15, fontweight='bold')
ax.set_ylabel("Protein Count", fontsize=13)
ax.set_xlabel("")  # X-axis is Duration, so label not strictly necessary
ax.set_ylim(0, bottom.max() * 1.05)  # give 10% headroom
ax.legend(title="Significant Peptides", bbox_to_anchor=(1.02, 1), loc="upper left")
ax.spines["top"].set_visible(False)
ax.spines["right"].set_visible(False)
ax.yaxis.grid(True, linestyle="--", linewidth=0.5, alpha=0.6)
# Resize the ticklabels and make them italic
ax.tick_params(axis='both', which='major', labelsize=12)
# Make them italic
for label in ax.get_xticklabels() + ax.get_yticklabels():
    label.set_fontstyle('italic')
plt.tight_layout()
plots.finalize_plot(
    fig, show=True, save=save_to_folder,
    filename='proteoformCategories_counts_stackedBar',
    filepath=figure_path,
    formats=figure_formats, 
    transparent=transparent_bg,
    dpi=figure_dpi,
)
```

```
           Form Categories  48hr  72hr
0  No Significant Peptides  3920  2109
1       Single Peptide PTM  2385  2818
2                 With dPF   522  1255
3    With Single PTM & dPF   334   979

           Form Categories   48hr   72hr
0  No Significant Peptides 0.5474 0.2945
1       Single Peptide PTM 0.3331 0.3935
2                 With dPF 0.0729 0.1753
3    With Single PTM & dPF 0.0466 0.1367
```

##### 02.3 Matched Annotations to PTMs Detailed Distribution of PTM Annotations¶

```
from matplotlib.patches import Patch

# This will be y-axis in the 

# Read 48hr and 72hr files, map labels, and pivot so durations become columns
df48 = pd.read_csv(f"{input_path}proteins_withPTM_peptides_UniprotAnnotation_featureCounts_48hr.csv")
df48['Duration'] = '48hr'
df72 = pd.read_csv(f"{input_path}proteins_withPTM_peptides_UniprotAnnotation_featureCounts_72hr.csv")
df72['Duration'] = '72hr'
plot_data = pd.concat([df48, df72], ignore_index=True)
plot_data = plot_data.pivot_table(index='Uniprot Feature Type', columns='Duration', values='Count', aggfunc='sum')
plot_data['Total'] = plot_data.sum(axis=1)
order_features = plot_data.sort_values(by='Total', ascending=False).index.tolist()

# two-panel mirrored horizontal bar plot: counts per Uniprot Feature Type, split by Duration (48hr left, 72hr right)
durations = ['48hr', '72hr']

# Get counts in the order desired
counts_48 = plot_data.loc[order_features, '48hr'].values
counts_72 = plot_data.loc[order_features, '72hr'].values
y_pos = np.arange(len(order_features))

# Symmetric x-limits
max_count = max(counts_72.max(), counts_48.max())

# Create two side-by-side axes sharing y-axis for the mirrored look
fig, (axL, axR) = plt.subplots(ncols=2, sharey=True, figsize=(10, 5), gridspec_kw={'width_ratios': [1, 1]})

# Left panel: 48hr (negative values to show mirror to the left)
barsL = axL.barh(y_pos, -counts_48, color="#a7c957", edgecolor="none", height=0.6)
axL.set_xlim(-max_count * 1.1, 0)

# Right panel: 72hr
barsR = axR.barh(y_pos, counts_72, color="#a7c957", edgecolor="none", height=0.6)
axR.set_xlim(0, max_count * 1.1)

# Ensure both axes share the same y-range
axL.set_ylim(-0.5, len(order_features) - 0.5)
axR.set_ylim(-0.5, len(order_features) - 0.5)

# Make x-ticks readable: left ticks as positive numbers for readability (abs)
axL.set_xticks(np.linspace(-max_count, 0, 5))
axR.set_xticks(np.linspace(0, max_count, 5))
axL.set_xticklabels([f"{int(abs(x)):,}" for x in axL.get_xticks()])
axR.set_xticklabels([f"{int(x):,}" for x in axR.get_xticks()])

# Hide y ticklabels on both axes (we will place central labels)
axL.set_yticks(y_pos)
axL.set_yticklabels([])
axR.set_yticks(y_pos)
axR.set_yticklabels([])

axL.axvline(0, color='gray', linewidth=1.5, linestyle='-', zorder=2) # zorder to be on top of grid
axR.axvline(0, color='gray', linewidth=1.5, linestyle='-', zorder=2)

# Place central y-axis labels between the two panels
bbox = axL.get_position()
ymin, ymax = axL.get_ylim()
for i, lab in enumerate(order_features):
    y_norm = (y_pos[i] - ymin) / (ymax - ymin)
    y_fig = bbox.y0 + y_norm * (bbox.y1 - bbox.y0)
    fig.text(0.5, y_fig, lab, ha='center', va='center', fontsize=11)

# Annotate counts near the center for both sides
label_offset = max_count * 0.02  # relative offset to place labels close to the center
for bar in barsL:
    w = bar.get_width()  # negative value
    axL.text(w + label_offset, bar.get_y() + bar.get_height() / 2, f"{int(abs(w)):,}",
             ha="right", va="center", color='k', fontsize=10, fontweight="bold")
for bar in barsR:
    w = bar.get_width()  # positive value
    axR.text(w - label_offset, bar.get_y() + bar.get_height() / 2, f"{int(w):,}",
             ha="left", va="center", color='k', fontsize=10, fontweight="bold")

# Titles and axes labels
fig.suptitle("Matched Annotations to PTMs - Feature Counts by Duration", fontsize=15, fontweight='bold')
axL.set_xlabel("Protein Count (48hr)", fontsize=12)
axR.set_xlabel("Protein Count (72hr)", fontsize=12)
# Vertical divider line for clarity (between the two subplots)
divider_x = 0.5
fig.axes[0].spines['right'].set_visible(False)
fig.axes[1].spines['left'].set_visible(False)

# Remove top/right spines from both axes and add gridlines
for a in [axL, axR]:
    a.spines["top"].set_visible(False)
    a.spines["right"].set_visible(False)
    a.spines["bottom"].set_visible(True)
    a.spines["left"].set_visible(False)
    a.yaxis.grid(False)
    a.xaxis.grid(True, linestyle='--', linewidth=0.5, alpha=0.6)
    a.tick_params(axis='both', which='major', labelsize=10)
    # italic for x ticklabels
    for lbl in a.get_xticklabels():
        lbl.set_fontstyle('italic')

# plt.tight_layout()
plots.finalize_plot(
    fig, show=True, save=save_to_folder,
    filename='PTM_featureCounts_48v72_mirrored',
    filepath=figure_path,
    formats=figure_formats,
    transparent=transparent_bg,
    dpi=figure_dpi,
)
```
